# Synthesis and evaluation of chemical linchpins for highly selective CK2α targeting

**DOI:** 10.1101/2024.05.16.594086

**Authors:** Francesco A. Greco, Andreas Krämer, Laurenz Wahl, Lewis Elson, Theresa A. L. Ehret, Joshua Gerninghaus, Janina Möckel, Susanne Müller, Thomas Hanke, Stefan Knapp

## Abstract

Casein kinase-2 (CK2) are serine/threonine kinases with dual co-factor (ATP and GTP) specificity, that are involved in the regulation of a wide variety of cellular functions. Small molecules targeting CK2 have been described in the literature targeting different binding pockets of the kinase with a focus on type I inhibitors such as the recently published chemical probe SGC-CK2-1. In this study, we investigated whether known allosteric inhibitors binding to a pocket adjacent to helix αD could be combined with ATP mimetic moieties defining a novel class of ATP competitive compounds with a unique binding mode. Linking both binding sites requires a chemical linking moiety that would introduce a 90-degree angle between the ATP mimetic ring system and the αD targeting moiety, which was realized using a sulfonamide. The synthesized inhibitors were highly selective for CK2 with binding constants in the nM range and low micromolar activity. While these inhibitors need to be further improved, the present work provides a structure-based design strategy for highly selective CK2 inhibitors.

## Introduction

Protein phosphorylation is a crucial regulatory mechanism, playing a key role in signal transduction.^1^ This post-translational modification triggers a change in the activation state of various proteins, leading to a plethora of different effects ranging from regulation of the cell cycle^2^, apoptosis^3^, cell growth^4^ as well as regulatory events in mRNA splicing.^5^ The enzymes that catalyse this reaction, protein kinases (PK), are a large family of regulatory proteins that have been associated with many different diseases including oncological malignancies, inflammation and neurologic disorders.^6–8^ Casein Kinase 2 (CK2) is a PK with probably one of the most extensive interactomes, affecting hundreds of cellular proteins.^9,10^ This suggests that CK2 plays an important role in various cellular processes and that an aberrant function of this key regulator can ultimately lead to the development of various disease phenotypes. CK2 associated diseases range from cancer^11,12^ and neurodegenerative diseases^13–15^ to viral^16–19^ and parasitic infections^20,21^, making CK2 an attractive potential target for the development of small molecule based therapies (**Figure 1**).^22–25^

**Figure 1.**
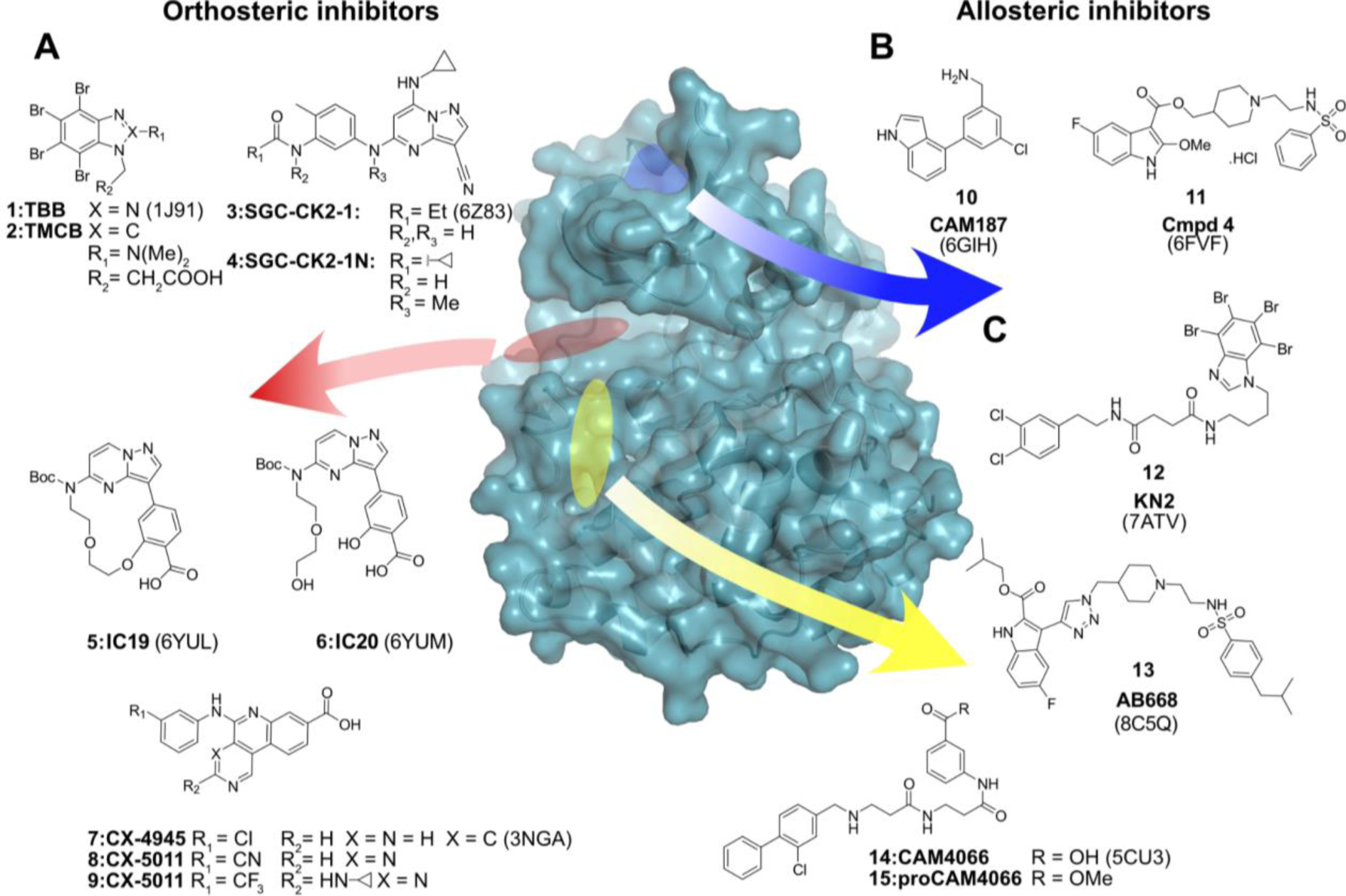
Overview of small molecules targeting CK2α sorted by binding pockets: (A) Orthosteric inhibitors targeting the ATP binding site of CK2α sharing classical hinge binding motifs like pyrazolo[1.5-a]pyrimidines and 2,6-naphtyridines. The PDB code of the corresponding crystal structure is shown in brackets. (B) Allosteric binders targeting the interface between CK2α/CK2β. (C) Ligands targeting the αD out conformation of CK2.

The most frequently used inhibitor in the literature is CX-4945 (Silmitasertib).^26^ CX-4945 is a benzonaphtyridine analogue with the classic features of a type I kinase inhibitor that reached phase-II clinical trials on cholangiocarcinoma.^27^ Apart from its potential as an anti-cancer drug candidate, CX-4945 is not a useful tool compound due to its significant off-target profile that includes potent inhibition of CDK1, CLK1–CLK3, DAPK3, DYRK1A/B, DYRK3, FLT3, HIPK3, PIM1, and TBK1 (*IC*_50_s < 100 nM).^28–30^ Indeed, the recent development of highly selective inhibitors of CK2, so called chemical probes, such as SGC-CK2-1 revealed that targeting of CK2 across different cancer cell lines generally does not lead to antiproliferative effects that have been prominently associated with CX-4945, emphasizing the importance of selective chemical probes linking phenotypic responses to targets.^31^ SGC-CK2-1 is a pyrazolopyrimidine-based type I kinase inhibitor and, like CX-4945, shares critical interactions in the hinge region and the back pocket of CK2. In an SAR study of the pyrazolopyrimidine scaffold leading to the discovery of SGC-CK2-1, it has been shown that even minor structural modifications can lead to a significant loss of selectivity and/or potency. This can be an issue if physicochemical-, or pharmacokinetic properties such as metabolic stability are altered without seriously affecting the overall pharmacodynamic features of the compound. Another important driver for chemical modifications at the compound level is the occurrence of mutations in the catalytic domain of kinases.^32,33^ These point mutations can occur throughout the catalytic domain, typically at the gatekeeper position, located deep in the ATP-binding pocket, which negatively affects the affinity of ATP-competitive compounds.^34^

The divergent phenotypes observed in studies comparing the promiscuous inhibitor CX-4945 with selective compounds such as SGC-CK2-1 challenge the role of CK2 in many disease processes discovered with CX-4945, including its role in regulation cell proliferation and cancer.^35^ However, inhibitors with different binding modes, such as allosteric and orthosteric inhibitors, have been associated with different phenotypes. Such observations may be based on the regulation of CK2 by protein interactions and its quaternary structure consisting of two catalytic (α and α’) and two regulatory (β) subunits.^36^ The discovery of allosteric inhibitors targeting a unique pocket adjacent to αD and the ATP binding site provides an opportunity for the design of selective CK2 inhibitors.^37,38^ However, while the αD pocket provides an interesting anchor point, targeting the pocket alone is unlikely to lead to potent inhibitors. Indeed, the combination of fragments binding to the αD pocket with moieties targeting the CK2 ATP site back pocket led to potent compounds, exemplified by the bivalent inhibitor CAM4066 (K_D_: 320 nM), which was highly selective and, in contrast to SGC-CK2-1, showed anti-proliferative effects at 10-20 µM concentration.^23^ More recently, the inhibitor AB668 demonstrated that bivalent compounds targeting the ATP site back pocket and the αD pocket simultaneously can have excellent selectivity and potency and induce apoptotic cell death in several cancer cell lines.^39^

Although AB668 targets the ATP site, it does not form the canonical hinge hydrogen bonds required for high potency of ATP-competitive orthosteric inhibitors. In this study, we aim to investigate if αD pocket targeting moieties could be linked to ATP mimetic scaffolds. This combination poses a challenge in linker design between the two moieties, as the linker needs to introduce a 90-degree kink, which we achieved by introducing a sulfonamide moiety directly attached to the hinge binding ring system. Testing diverse hinge binding moieties and linkers we succeeded developing bivalent inhibitors with nM K_D_ assessed by ITC (Isothermal titration calorimetry) and excellent selectivity.

## RESULTS

The catalytic subunits of CK2 have unique structural features that render this kinase constitutively active and facilitate the design of selective CK2 inhibitors. Firstly, the N-terminal segment of CK2 is tightly bound to the activation loop, stabilizing an active conformation. Secondly, the DFG motif in CK2 is changed to DWG, allowing the formation of an additional hydrogen bond between the W176 and the backbone of L173 which is also contributing to the stability of the active state.^40^ Third, the αD helix has been shown to be flexible even in the apo forms of the enzyme.^41^ Brear *et al.* have used high-throughput crystallography to show that a binding site created by an outward movement of αD can be targeted by small molecular fragments and can therefore be used for the development of allosteric CK2 inhibitors.^38^

In our study, we pursued three main strategic considerations for our compound design:

1. The biphenyl-methanamine moiety served as a linchpin molecule that targets the αD out conformation (**Figure 2A**).
2. An appropriate linker length of at least 8-9 Å was employed to connect the αD ligand to an ATP site binding moiety (**Figure 2A**, ovaloid).
3. An angle of approximately 90° between the ATP binding site moiety and the linker region was introduced to position the ATP mimetic moiety of the molecule towards the hinge region (**Figure 2B**).

**Figure 2.**
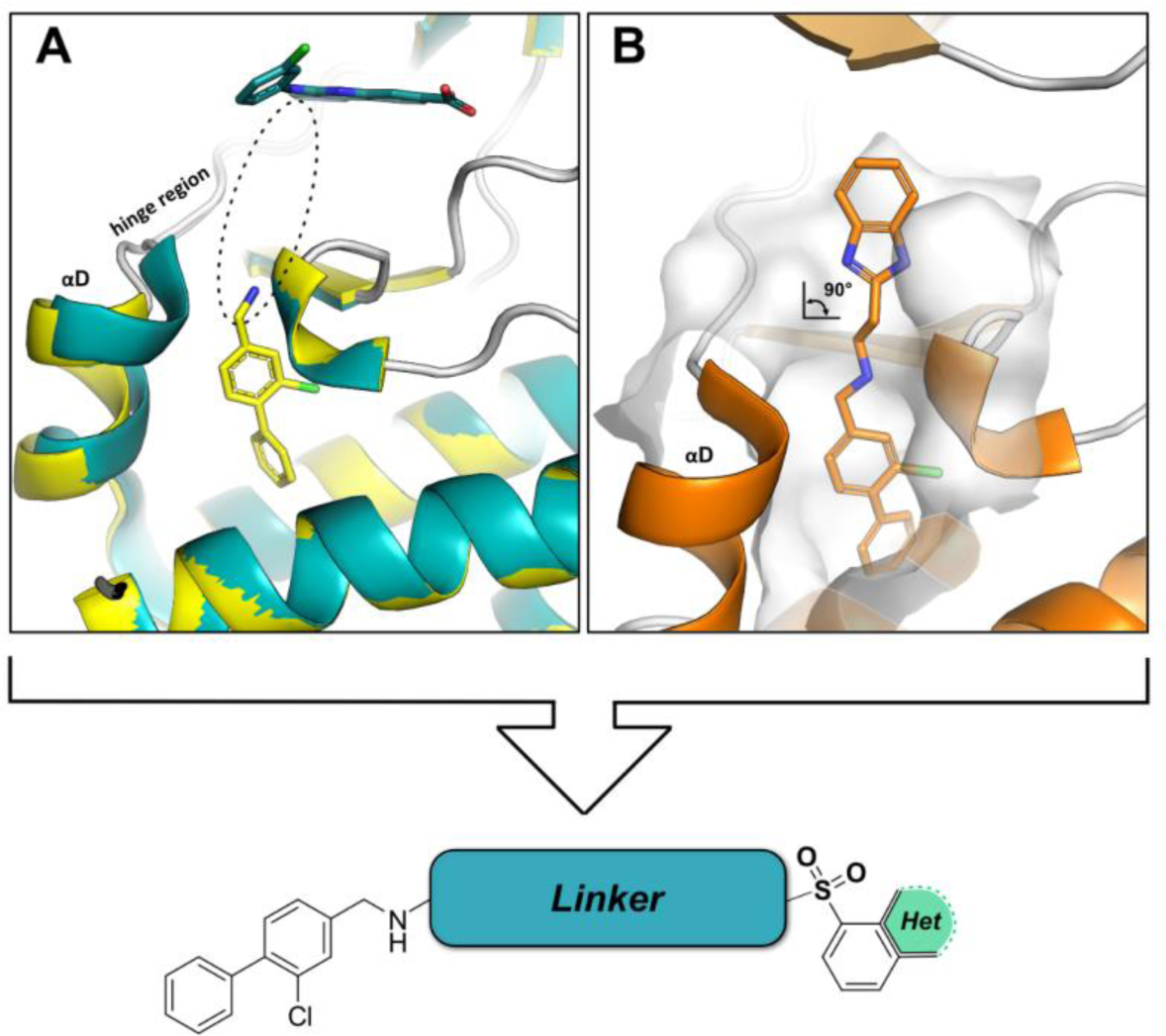
Design strategy: (A) Overlay of CK2α bound to the type-I inhibitor CX-4945 (teal, PDB 3NGA) and the αD targeting biphenyl (yellow, PDB 5CSH). Dotted ellipse indicates the position for linker design. (B) Benzimidazole motif (orange, PDB 5OUU) protruding from the αD pocket emphasizing the need of an orthogonality inducing functional group like sulfonamides and sulfones.

Compounds **26–58** were synthesized using a 5-step synthetic route as outlined in **Scheme 1**. The αD ligand was assembled by DMAP catalyzed functionalization of commercially available 3-chloro-4-hydroxybenzaldehyde **16** with triflate anhydride yielding the corresponding triflated product **17** in 86% yield on a 12 g scale.^42^ **17** underwent LiCl accelerated Suzuki Miyaura cross coupling with commercial benzene boronic acid in yields of **18** from 60-70%.^42^ Di-amino linkers of varying complexities were combined with **18** via reductive amination with yields ranging from 57–80% (**19–24**). Deprotection of the Boc group protected amines proceeded with TFA in DCM yielding the corresponding TFA salts that were used without further purification. These compounds were reacted in high dilution with different commercially available sulfonyl chlorides in THF and excess of base at ambient temperature with yields from 71-92% (**26–58**).

**Scheme 1.**
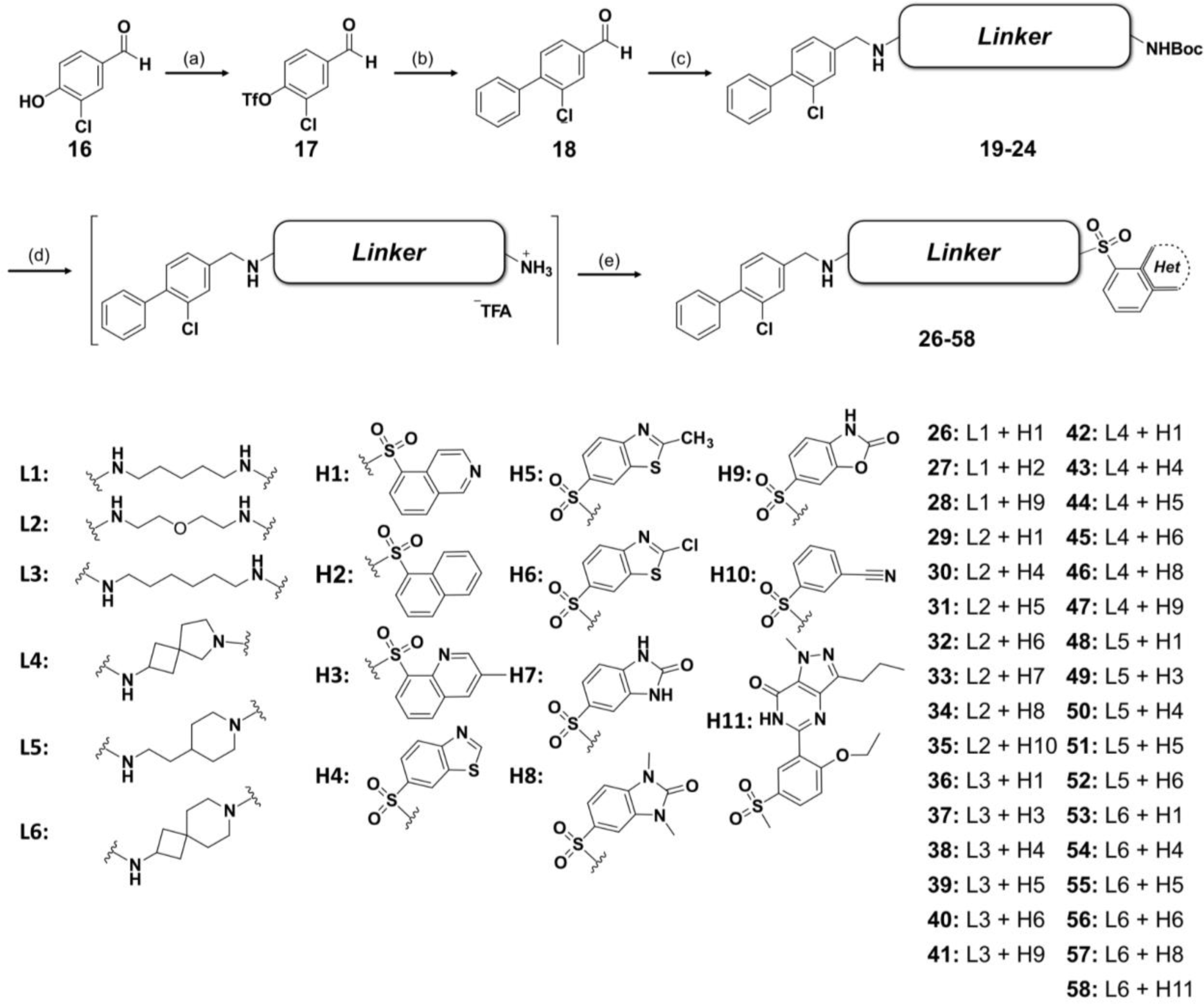
Synthesis of **26**–**58**.^a^ ^a^ Reagents and conditions: (a) Tf_2_O (1M DCM), pyridine, cat. DMAP, DCM, rt, overnight, 86%; (b) Phenyl boronic acid, Pd(PPh_3_)_4_, LiCl, DME, µW, 80 °C, 3 h, 60–70%; (c) Na[(CH₃COO)₃BH], MgSO_4_, DCM, overnight, rt, 57-80%; (d) TFA, DCM, 0 °C-rt, 1 h, quant.; (e) Sulfonyl chloride, N(Et)_3_, THF, overnight, rt; 71-92%. L = linker and H = head group.

Due to the paucity of commercially available sulfonyl chlorides directly attached to fused heterocyclic cores, we tried different reactions conditions for the *de novo* synthesis of this moiety to further expand the chemical space. Unfortunately, various attempts ranging from direct chlorosulfonylation and Li-halogen exchange to Grignard type mediated sulfinate syntheses failed on a range of heterocyclic compounds. Willis *et al.* published a milder palladium-catalyzed method for the *in situ* generation of sulfinate salts starting from brominated aromatic compounds and the SO_2_ surrogate DABSO (DABCO-Bis(sulfur dioxide)).^43^ The sulfinate salt can be reacted directly with an electrophilic F^+^ reagent such as NFSI to yield sulfonyl fluorides, which are suitable for chromatographic purification due to their higher chemical stability.^44^ Thus we applied this chemistry and synthesized sulfonyl fluoride **60** by reacting 6-bromo-3-methyl-3*H*-imidazo[4,5-*b*]pyridine (**59**) with DABSO, Pd(amphos)_2_Cl_2_, dicyclohexylamine and isopropanol serving both as solvent and reducing agent under microwave irradiation (46% isolated yield).^44^ **60** was then coupled with deprotected compound **19** that was synthesized according to Scheme 1 yielding the final product **61** in 72% overall yield (Scheme 2). However, it must be noted that this approach still has a limited substrate scope in terms of heterocyclic scaffolds, which restricts the synthesis of further sulfonyl fluorides.

**Scheme 2.**
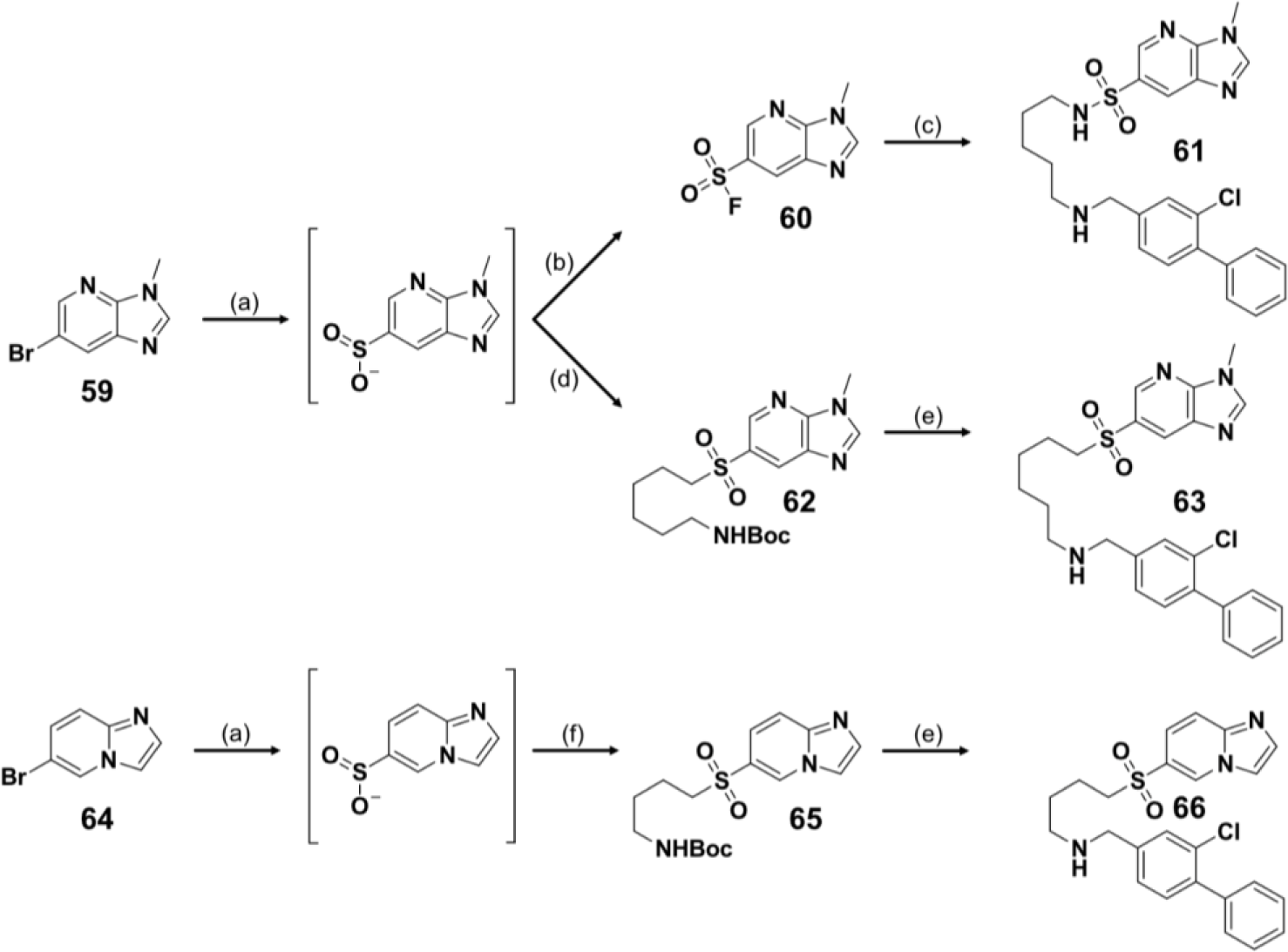
Synthesis of **61**, **63** and **66**.^a^ ^a^ Reagents and conditions: (a) DABSO, Pd(amphos)_2_Cl_2_, Cy_2_NME, *i*-PrOH, 110 °C, 1 h, µW; (b.1) NFSI, 3 h, rt, 46%; (c) **19** (deprotected), DMSO, 18 h, 110 °C, µW, 72%; (d) *tert*-butyl (6-bromohexyl)carbamate, DMF, 2 h 100 °C, µW, 40%; (e) TFA, DCM, 0 °C-rt, 1 h, quant.; **18**, Na[(CH₃COO)₃BH], MgSO_4_, DCM, overnight, rt, 25% (**63**) and 23% (**66**); (f) *tert*-butyl (4-bromobutyl)carbamate DMF, 2 h 100 °C, µW, 14%.

The first generation of compounds contained a simple alkyl chain with 4 to 5 carbon atoms, flanked by the αD-binding moiety and the fused heterocycle (henceforth referred to as the ‘head group’, H), which was designed to reach the ATP binding site. The binding of **26** (pentyl linker) to both CK2 isoforms was verified by differential scanning fluorimetry (DSF) and the proposed binding mode was confirmed by X-ray crystallography (**Figure 3**, left panel).

**Figure 3.**
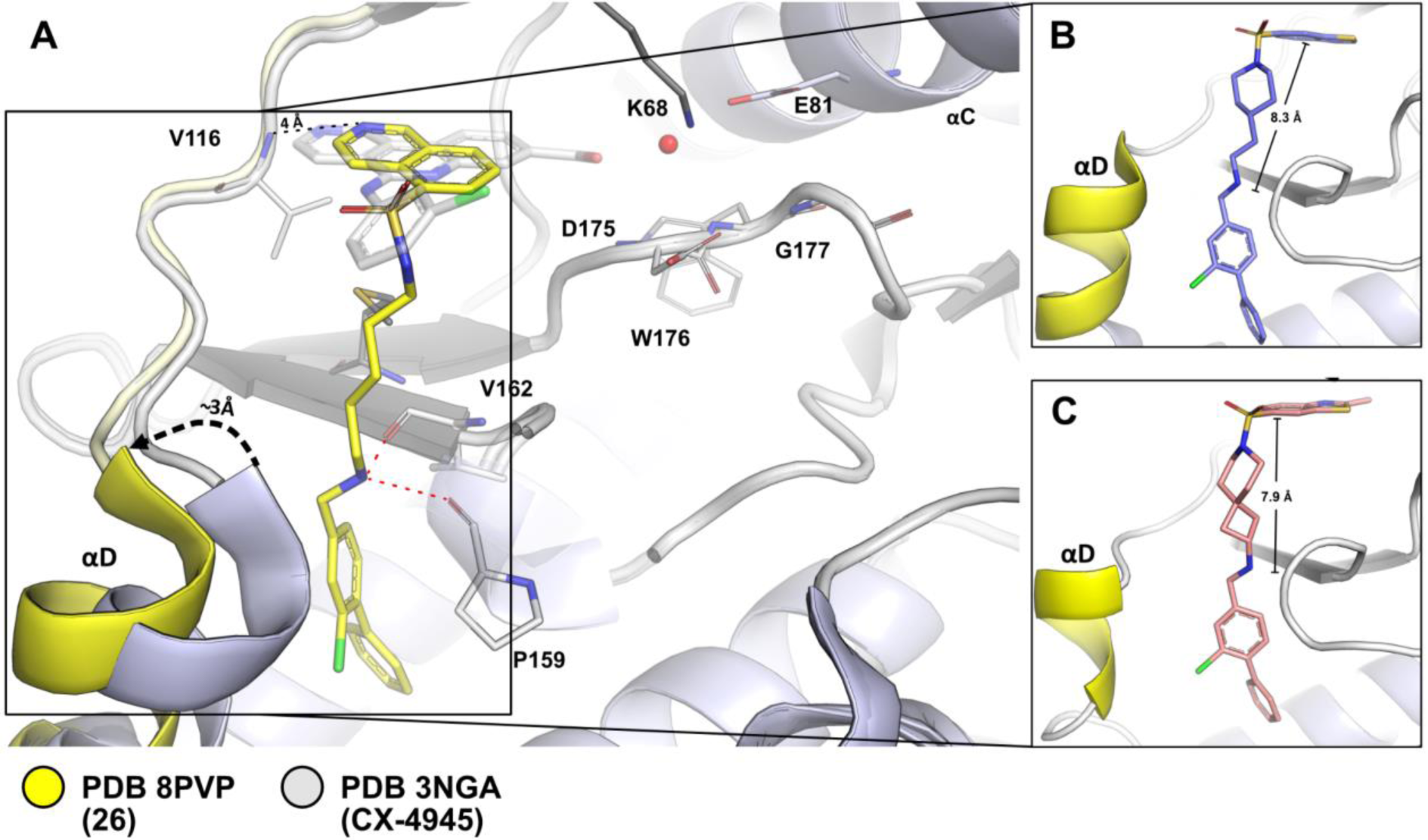
Proof of principle and comparison of linkers: (A) **26** crystallized with CK2α (yellow) overlayed with CX-4945 (grey). Interactions of **26** between P159 and V162 are shown in red. The basic nitrogen of **26** is in ca. 4 Å distance to V116 which hinders the formation of a hydrogen bond. The black arrow indicates the shift of the αD helix from the *in* to the *out* conformation. (B) **50** bearing a piperidine-4-ethanamine moiety. (C) **55** showing that further rigidification to a 7-azaspiro[3.5]nonane is tolerated.

Remarkably only the compounds carrying a linker motif and a hinge binding moiety showed significant shifts in the DSF assay, in contrast to the allosteric biphenyl fragment (**S1**) alone (**Table S3**).

The co-crystal structure analysis of **26** and CK2α shows that the combination of a pentyl linker connected to both the biphenyl and a quinoline successfully induces the desired binding mode (**Figure 3**, left side).

Based on the first DSF results, we considered the linker to be the key component of our inhibitor design. A more detailed analysis of the conformation of the aliphatic linker inspired us to use both a rigidification strategy to stabilize this conformation and to test whether the sulfonamide H-donor interferes with binding, leading to compounds **48** and **53**. **48** contains a piperidine-4-ethanamine ring, which rigidizes the linker in the upper part of the molecule and simultaneously masks the N-H of the sulfonamide. The main feature of compound **53** is a 7-azaspiro[3.5]-nonane linker designed so that the linker is pre-organized at both ends. Derivatives of **48** and **53** were co-crystallized on CK2α to additionally test whether the modifications made still address the ATP binding site (**Figure 3**, right side showing **50** and **55**). Both DSF and X-Ray structural analysis demonstrated that the new linking strategy and the introduction of different binding moieties at the ATP pocket were promising and that there was no apparent loss of binding (**Table 1**). ITC data of **26** and **48** supported the DSF results revealing K_D_ values of and 316 nM and 263 nM, respectively (**Figure 5**). Based on these results, we extended our SAR with different linkers in combination with several ATP binding site moieties and tested them using DSF as an initial assay system (**Table 1**). Aliphatic linkers of varying complexity were used to investigate different conformations of this moiety, replacing the more labile linear amide linkers that has been used in previous reports.^23,42^ All aliphatic linkers were tolerated with linkers with more carbon atoms performing better than short ones. Hexyl linkers, used in compounds **36–41,** boosted ΔTm shifts by 1-4 K compared to butyl- (**25**) or pentyl linker (**26–28, 61, 72**). Cmpd **26 (Figure 4A)** and **25 (Figure 4B)** differed in terms of linker length (pentyl vs butyl) and in the orientation of the isoquinoline ring in the ATP binding site. In conjunction with longer linkers, the isoquinoline ring was more aligned with the hinge region, while the shorter butyl linker was more aligned towards the back pocket of the kinase. Additionally, the quinoline moiety of **25** was tilted out of the plane of both **26** and **CX-4945** (**Figure S1**) which might explain the overall lower affinity of this compound since a surface complemental, coplanar orientation of the ATP site binding moiety was detrimental to be perfectly sandwiched between the N and C lobe of the kinase. Surprisingly the substitution of the C3 to an oxygen atom in the pentyl linker series rendered molecules largely inactive (**29–35)**. Polyethyleneglycol (PEG) linkers are known to adopt twisted, meander-like conformations^45^, possibly resulting in the αD ligand and/or head group being pushed out of their corresponding pocket.

**Figure 4.**
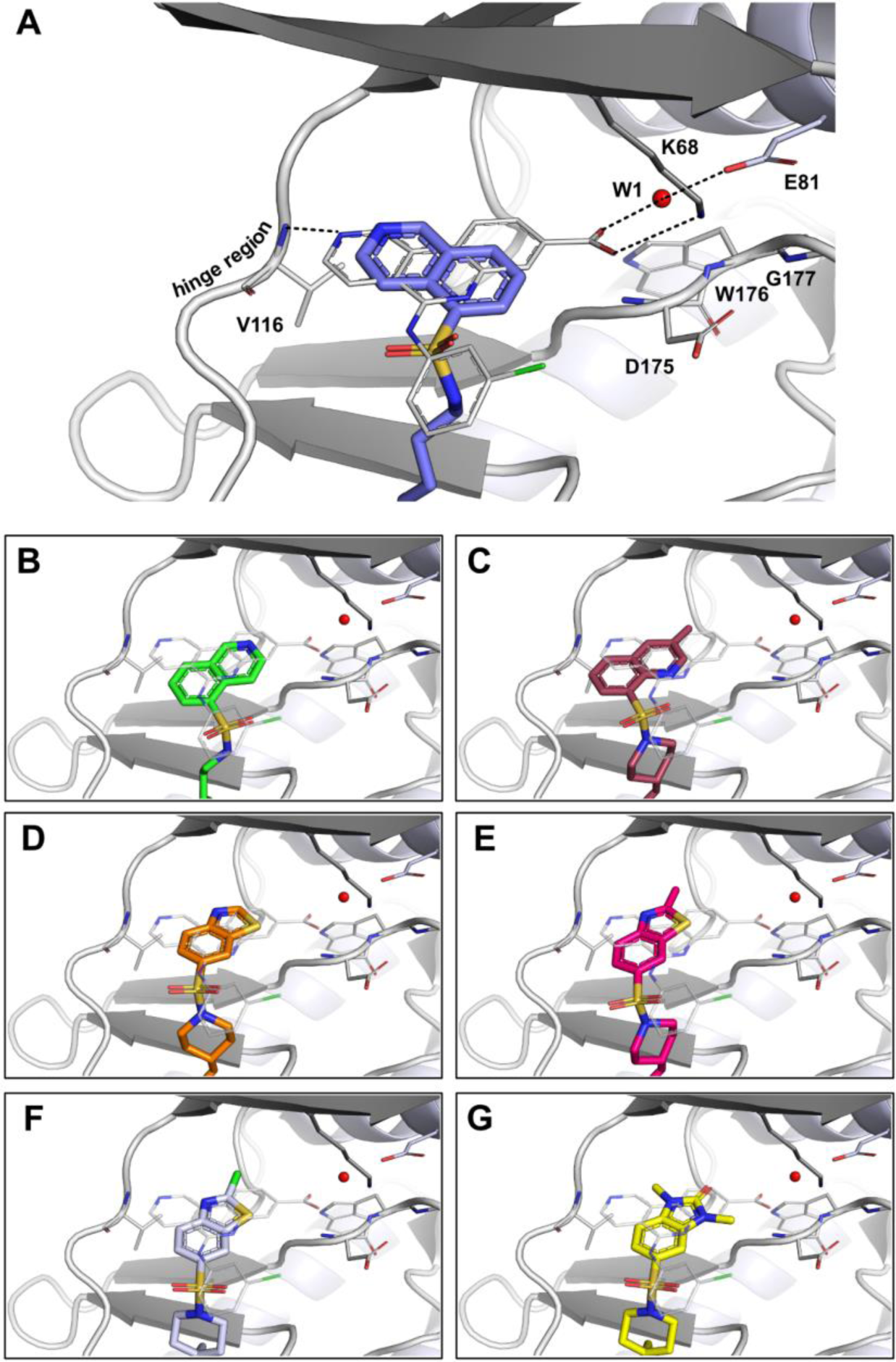
Overview of different ATP site binding moieties: (A) Overlay of **26** (PDB 8PVP) and CX-4945 (PDB 3NGA) as a reference compound for ATP site targeting scaffolds. Polar interactions of CX-4945 are shown as dotted lines. (B, C) Annulated 6 membered rings of the quinoline type bound to CK2α (**25** PDB 9EQ0, **49** PDB 9EPW). A pentyl linker attached to the quinoline orients the basic nitrogen towards the hinge region (A) as opposed to (B) bearing a butyl linker. 3-methylquinoline pointing towards the back of the kinase (C). (D–F) Benzothiazole derivatives bound to CK2α (**50** PDB 8PVO**, 51** PDB 9EPZ**, 56** PDB 9EPX). The chlorine moiety in (F) can be used as a chemical handle for further derivatization. (G) The dimethylbenzimidazolone scaffold showcases the tolerance of saturated ring systems in the ATP binding region of the kinase (**57** PDB 9EPV).

**Figure 5.**
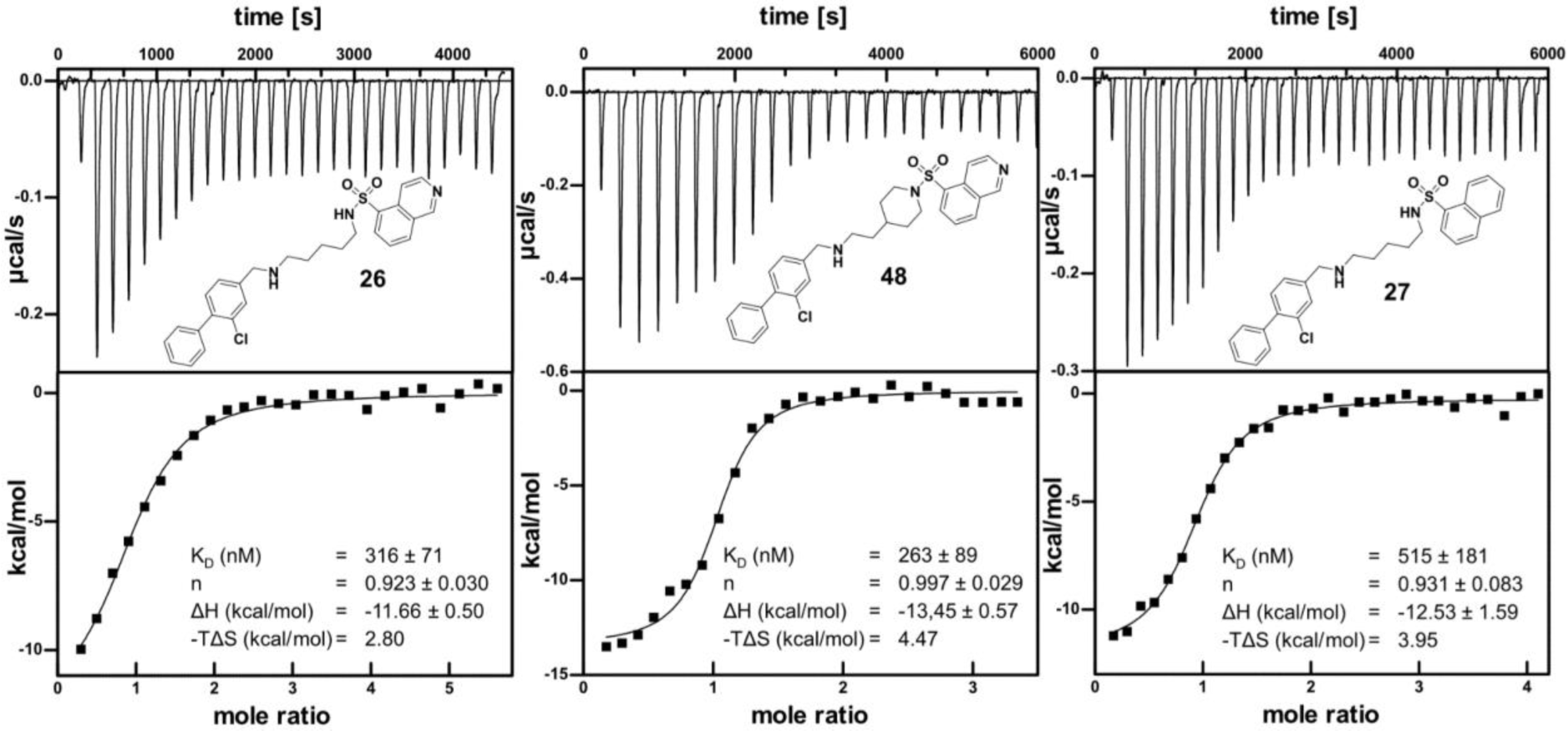
ITC measurement of **26, 27** and **48**. Rigidization and *N*-deletion strategies (**48, 27**) are investigated in ITC in comparison to proof-of-principle molecule **26**.

**Table 1.** Overview of synthesized molecules 25–58, 61, 63, 65, 69 and 70 compounds sorted by linkers: corresponding ΔTm values against CK2α/α’ in °C. **14** and **15** are used as reference compounds. Measurements were performed in technical triplicates. H1-H13 represent the employed head groups.

| Cpd ID | Linker | Het | $\Delta T_m$ [°C] | | Cpd ID | Linker | Het | $\Delta T_m$ [°C] | |
| --- | --- | --- | --- | --- | --- | --- | --- | --- | --- |
| | | | CK2α | CK2$\alpha'$ | | | | CK2α | CK2$\alpha'$ |
| 25 |  | H1 | 5.1 | 3 | 42 |  | H1 | 7.0 | 3.9 |
| 26 |  | H1 | 6.1 | 2.7 | 43 |  | H4 | 7.9 | 4 |
| 27 |  | H2 | 6.0 | 3.2 | 44 |  | H5 | 8.0 | 3.9 |
| 72 |  | H4 | 8.6 | 4 | 45 |  | H6 | 6.9 | 3.5 |
| 61 |  | H13 | 2.7 | 1.2 | 46 |  | H8 | 6.0 | 3 |
| 28 |  | H9 | 8.1 | 4.2 | 47 |  | H9 | 8.9 | 3.8 |
| 29 |  | H1 | 2.2 | 1.1 | 48 |  | H1 | 7.0 | 5 |
| 30 |  | H4 | 4.0 | 1.3 | 49 |  | H3 | 6.9 | 3.6 |
| 31 |  | H5 | 2.6 | 0.6 | 50 |  | H4 | 6.2 | 5 |
| 32 |  | H6 | 3.6 | 1.4 | 51 |  | H5 | 6.0 | 3 |
| 33 |  | H7 | 1.0 | 0.5 | 52 |  | H6 | 8.2 | 4.2 |
| 34 |  | H8 | 1.8 | 0.5 | 53 |  | H1 | 6.4 | 4.7 |
| 35 |  | H10 | 1.2 | 1 | 54 |  | H4 | 5.8 | 5.2 |
| 36 |  | H1 | 10 | 8.7 | 55 |  | H5 | 4.8 | 3.9 |
| 37 |  | H3 | 7.1 | 3 | 56 |  | H6 | 5.4 | 3.7 |
| 38 |  | H4 | 9.0 | 5.1 | 57 |  | H8 | 6.7 | 4.1 |
| 39 |  | H5 | 8.6 | 3.9 | 58 |  | H11 | 2.0 | 0.5 |
| 40 |  | H6 | 8.7 | 4.3 | 73 |  | H4 | 6.0 | 3.2 |
| 41 |  | H9 | 9.9 | 6.3 |  |  |  |  |  |
| 66 |  | H12 | 4.3 | 0.8 | proCAM4066 (14)<br>R = Me |  |  | 4.5 | 0.8 |
| 63 |  | H13 | 6.7 | 2.8 | CAM4066 (15)<br>R = H |  |  | 8.6 | 4.2 |

The spectrum of more rigid linkers included the aforementioned piperidine-4-ethanamine (**48– 52**), 7-azaspiro[3.5]nonane (**53–58**) as well as a 6-azaspiro[3.4]octane-2-amine (**42–47**) linker designed to evaluate different ring strains and to optimize the length of the original pentyl linker.

The 6-azaspiro[3.4]octane-2-amine linker was chosen to leave the linear architecture of the molecules and thus conveying a certain degree of directionality of the head group. These linkers were tolerated in all head groups examined, with values similar to those of the pentyl linker series.

In addition to masking the N-H donor moiety of the sulfonamide group by making it tertiary, we also wanted to formally remove the nitrogen. This was done by replacing the nitrogen of the sulfonamide group with a carbon atom, thus converting it into a sulfone group. To achieve this transformation, we used the protocol of Willis *et al.* but instead of using F^+^ as the electrophile, we took *N*-Boc protected bromo alkyl linkers and reacted them with the corresponding sulfinate salt, resulting in compounds **63** and **66** (Scheme 2**, bottom).** Direct comparison of compounds **61** and **63** shows that simple deletion of the nitrogen atom unexpectedly boosts the ΔTm shift from 2,7 to 6,7 K rendering the 3-methyl-imidazopyridine head group interesting for future studies. The shorter butyl linker (**66**) with the imidazo[1,2-*a*]pyridine moiety showed significantly smaller ΔTm shifts than compound (**63**).

Numerous head groups were investigated, and the binding mode was analysed by X-Ray crystallography (**Figure 4**). All compounds share a bicyclic 6-6 (**Figure 4 A–C**) or a 5-6 (**D–G**) annulated heterocyclic ring system, which was linked to a corresponding linker moiety via an orthogonality enforcing sulfone group (not shown) or a sulfonamide group. All head groups extend into the ATP binding site assuming different orientations of the annulated ring systems. **49** (**Figure 4D**) bears a 3 methylquinoline head group, suggesting that the orientation of the quinoline nitrogen atom was not important for binding (ΔTm shift of 7.0 and 6.9 K. respectively, **Table 1**). 5-6 fused heterocyclic ring systems of the benzothiazole type (**Figure 4 D–F**) showed a similar binding mode to **26** and **49.** Due to the contracted 5 ring unit, functional groups attached to the 2-position of the benzothiazole ring were preferentially oriented towards the gatekeeper residue of the kinase (**Figure 4E, F**). **56** bears a chlorine residue that gives the opportunity for organometallic coupling reactions that can be used to explore the back pocket of the kinase (**Figure 4F**). In addition to the aromatic rings, we also investigated fused 5-6 heterocycles with saturated 5-membered rings. **57** contained a 1,3-dimethyl-benzimidazolone moiety gave rise to a similar DSF shift compared to the aromatic counterparts and a similar binding mode compared to the aforementioned ATP mimetic groups **(Figure 4G)**. The transition from traditional aromatic heterocyclic systems that have been almost exclusively used in the development of kinase inhibitors^46,47^ to saturated ring systems could not only yield more favorable selectivity profiles, but may also improve physicochemical and pharmacokinetic properties of inhibitors.^48,49^

Motivated by this finding, we expanded the spectrum of saturated ring systems (**28, 33, 34, 41, 46, 47, 57**). Interestingly, the benzoxazolone scaffold exhibited higher DSF shifts in combination with several linker combinations that matched or even exceeded the ΔTm shift of the reference compound **CAM4066 (15)** (see entries **28, 41, 47** in **Table 1**).

To correlate the ΔTm shifts of the compounds with binding constants in solution, we examined the binding of individual compounds using ITC (**Figure 5**). Comparison of binding constants of **26** with **48** showed that the rigid piperidine-4-ethanamine performed similarly to the more flexible analogue (K_D_ values of 263±89 nM and 316±71 respectively) with a slightly higher contribution of entropy to overall binding for **48**. These values correlated well with the DSF data that showed comparable shifts for both compounds. We were interested in the overall contribution of the nitrogen atom in isoquinoline derivative **26** to the binding and therefore replaced it with a carbon atom in compound **27**. **27** was synthesized according to Scheme 1 and evaluated in DSF and ITC. **27** performed similarly in DSF to **26** (6.0 vs 6.1 °C for CK2α) demonstrating again that the nitrogen was not involved in direct polar interaction with the kinase. This can be confirmed based on the crystal structure of **26** with CK2α that shows a distance of ≈ 4 Å between the basic nitrogen and V116, which is too far away for a hydrogen bonding interaction (**Figure 3A**). ITC revealed a binding constant of 515±181 nM which was comparable to the nitrogen containing counterpart **26.**

Next, we evaluate the selectivity of the synthesized inhibitors. We chose **48**, **51**, **55** and **57**, harbouring different head groups and linkers and evaluated them in a DSF screen with an in-house panel of 101 protein kinases. All compounds showed excellent selectivity irrespective of the chosen hinge binding moiety or linker. **48** was chosen as an example and the obtained ΔTm shifts were mapped onto the kinome. We also evaluated non-selective compounds as positive controls (**Figure 6A**). To test against a wider panel of kinases, **26** and **50** bearing a benzothiazole ring were profiled in an orthogonal tracer displacement assay.^50^ Values are shown as % inhibition again highlighting the tolerance of different ATP site binding moieties while preserving high selectivity (**Figure 6B**). Additional selectivity data for compounds **51, 55, 57** (DSF data) and **26** (tracer displacement assay) are summarized in the supplemental data file (**Table S1** and **S2**). To illustrate the broad range of kinase coverage used in this study we overlayed the different panels of selectivity assays in a Venn diagram. A total of 177 different kinases were screened with 23 kinases overlapping in the assays used (**Figure 6C**). ΔTm data, NanoBRET IC_50_ in permeabilized cells and in vitro activity values were compared for a few selected compounds and these data showed good correlation between these orthogonal assays (**Figure 6D**).^30,51^

**Figure 6.**
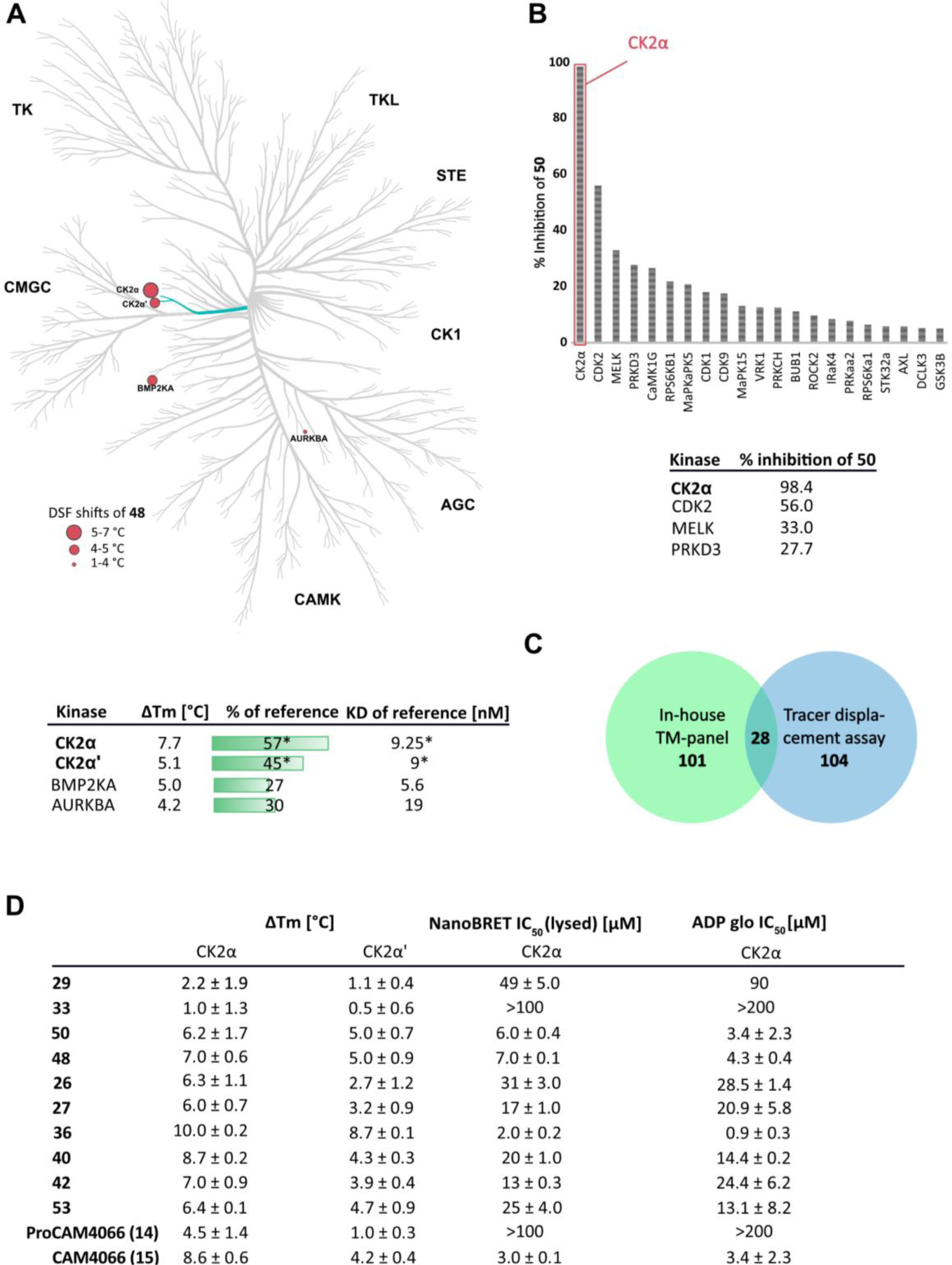
Selectivity profiling and on target evaluation: (A) Kinome tree representation of Tm shifts of **48** tested in an in-house panel of 101 protein kinases. Values are shown in °C. ΔTm values of top hits of DSF screen of **48** as absolute values and in relation to the reference compound Staurosporine. *CX-4945 was used as reference compound. (B) Structurally related cmpd **50** profiled in a panel of 104 kinases using a tracer displacement assay. Values are shown as % inhibition. Compound conc. 5µM. (C) Venn diagram representation of the two kinase panels used in this study. A total of 177 different kinases were screened with an overlap of 28 kinases. (D) Summary of tm shifts of compounds across different scaffolds in °C. Corresponding enzyme kinetic IC_50_ values for CK2α. The values were determined using a NanoBRET assay in permeabilized cells and ADP glo (*in vitro*).

The sulfonamide moiety not only ensured the correct angle between the linker and ATP binding function, but it also offered a possibility for additional derivatization by introducing tertiary sulfonamides. Derivatization at this position allowed extending the inhibitors towards the solvent region, and it can serve as a position for physicochemical optimization or as an attachment point for degrader development such as PROTACs (PROteolysis Targeting Chimeras). The accessibility of this position was confirmed by X-ray structures of **25** and **26**, in which the N-H of the sulfonamide was as expected pointing out of the orthosteric binding pocket. We decided for S_N_2 alkylation using methyl bromoacetate to generate the proof-of-concept compound **73**. In contrast to Scheme 1, the synthetic route to this target molecule was changed to avoid a loss of chemoselectivity in the alkylation step (Scheme 3).

**Scheme 3.**
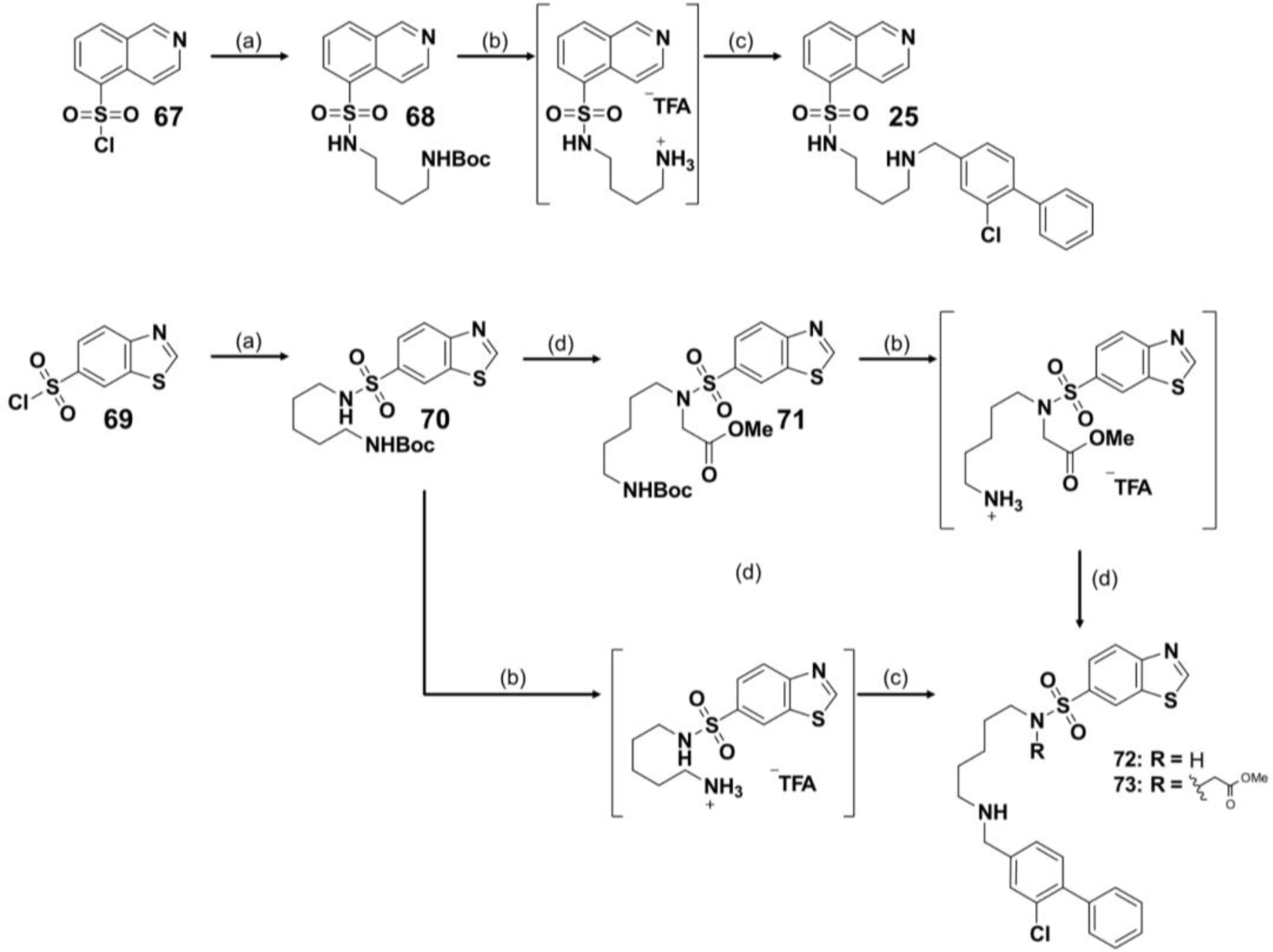
Synthesis of **25**, **72** and **73** starting from the hinge binder.^a^ ^a^ Reagents and conditions: (a) *tert*-butyl (5-aminopentyl)carbamate, *N*(Et)_3_, THF, overnight, rt, 58% (**68**) 70% (**70**); (b) TFA, DCM, 0 °C-rt, 1 h, quant.; (c) **18**, Na[(CH₃COO)₃BH], MgSO_4_, DCM, overnight, rt, 46% (**25**), 87% (**72**) and 42% (**73**); (d) Methyl bromoacetate, K_2_CO_3_, ACN, 16 h 0–60 °C, quant.

The binding of **73** to CK2α and α’ was assessed using the DSF assay, resulting however in smaller thermal shifts comparable to that of compound **72** (6.0/3.2 °C and 8.6/4 °C for α/α’ respectively). In addition, the structure of **73** in complex with CK2α was solved to elucidate the binding mode of the molecule (**Figure 7**). Encouragingly, we found that the expected orientation of the methyl ester pointed towards the triphosphate region of the kinase, while it did not significantly affect the binding mode of the biphenyl and the benzothiazole moiety in comparison to **50** (overlay of PDB 8PVO and PDB 9EQ1, not shown). In addition, the binding mode indicated that around the tertiary sulfonamide was sufficient space to extend this linker or solvent exposed moieties further and thus enable chemical modifications which can be achieved as suggested in Scheme 3 using different haloalkanes or by using the methyl ester of **73,** which can undergo further reactions such as amide couplings after hydrolysis providing a roadmap for modification of this scaffold in the future.

**Figure 7.**
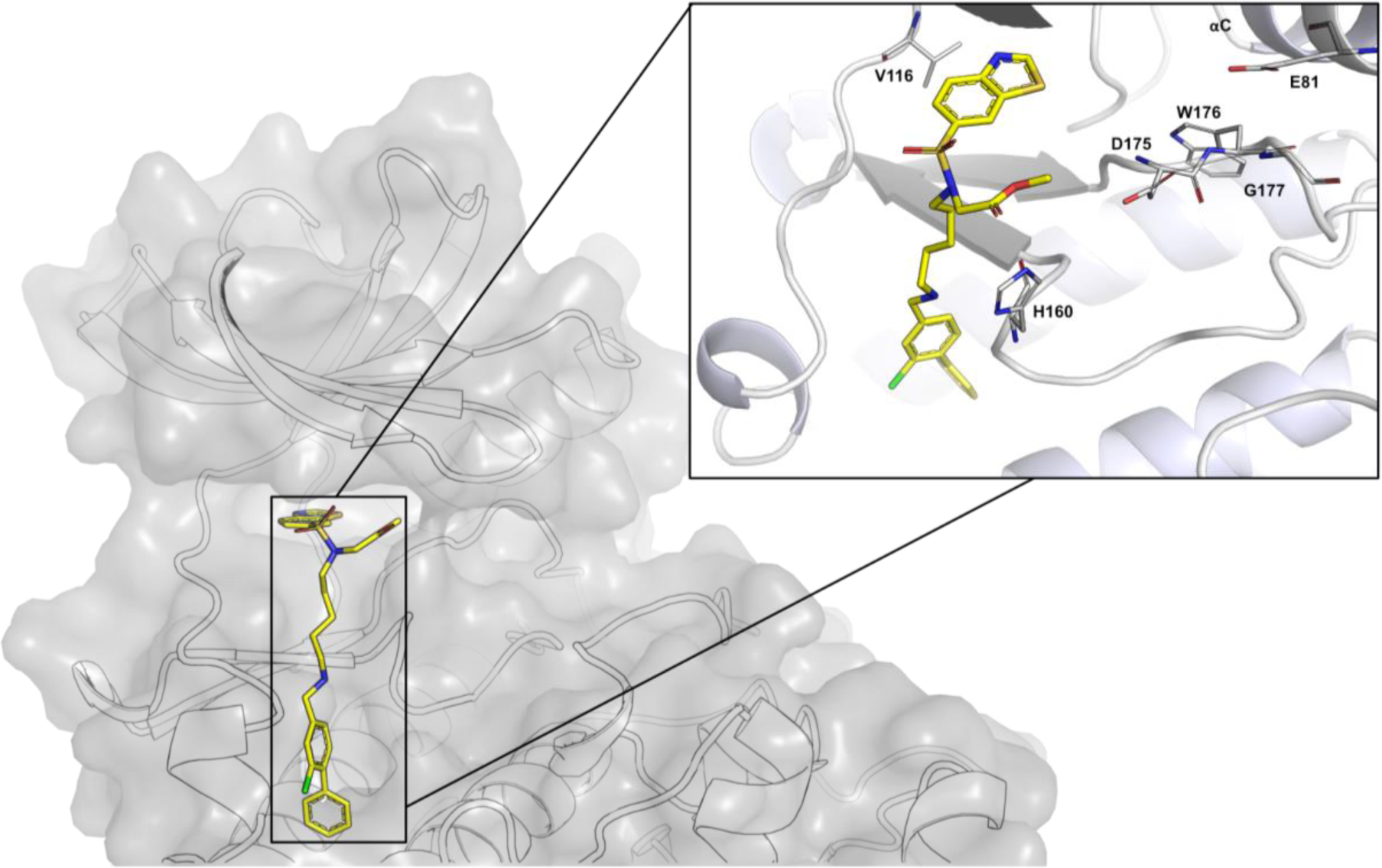
Co-crystal structure of **73** in CK2α, showcasing the third derivatization direction harnessing the sulfonamide moiety. The methyl ester chemical handle can be further extended both in the direction of the triphosphate pocket and out to the solvent region.

## Summary

In this study we outlined a strategy for CK2 inhibitor design combining known allosteric inhibitors with ATP mimetic moieties. We chose both a sulfonamide and sulfone group as a hinge element for the construction of a series of bivalent molecules. SAR of different combinations of the linker region and the head group demonstrated that 30 of the 39 compounds showed significant binding affinity to CK2 (6-10°C in DSF assay and ITC values of 263 nM for **26** and 316 nM for **48**). Binding to the kinase was achieved despite the lack of a classical hydrogen bond to backbone amino acids of the hinge region. This is in agreement with the binding mode of some natural products such as Emodin, first synthetic molecules including TBB and potent inhibitors of CK2 like GO289.^52–54^ While omitting the hinge region can be a valid strategy to increase selectivity, as this segment is highly conserved throughout the kinome, it usually comes at the cost of potency. In future optimization efforts, both an elongation to said region and growing towards the back pocket of the kinase might be pursued to increase the affinity of the compounds. In fact, the molecules synthesized in this study offer a comprehensive set of starting points that can be derivatized by simple transformation reactions towards different areas of the kinase. Firstly, all hinge binding moieties (except for **H11**) show high selectivity and can be extended either towards the back pocket or to the hinge region. Secondly, derivatization at the tertiary sulfonamide enables targeting of less addressed regions by inhibitors like the triphosphate binding site. Shorter butyl linkers (**25** and **65**) and the incorporation of a PEG moiety were less well tolerated (**29-35**). Crystal structure analysis showed that these linkers induce an unfavorable conformation of the hinge binding moieties that weakens hydrophobic interactions in the active site cleft and thus should be avoided in future studies. Cellular potency of the series is low which can be explained by a high intracellular concentration of ATP and a tight binding of ATP to CK2 (Km ≈ 10 µM) that outcompetes the mainly hydrophobic interaction of the head group moieties.^55^ This might be overcome when combining the different targeting strategies mentioned above that can result in compounds that can effectively inhibit the kinase *in cellulo*. The biological effects of CK2 *in vivo* are not exclusively linked to the catalytic activity of the protein.^35^ Indeed, it has been shown, that subunits of the holoenzyme can independently serve as scaffolding proteins.^56^ **73** was designed to extend into the solvent region of CK2 with the methyl ester serving as an exit vector. This position can be linked to an E3 ligase and harnessed for PROTAC development. Such an approach can conceivably circumvent the problem of low potency and low activity of the compounds which are both parameters that do not necessarily correlate with the pharmacological modality of protein degradation. Additionally, this might allow to further push the biological exploration of kinase-independent functions of CK2.

## Author Information

Manuscript was drafted by FAG and edited by all authors. FAG designed compounds which were synthesized by FAG, JG and JM. AK performed ITC measurement, provided the proteins for the DSF assay and performed the X-ray crystallography and structural analyses; LE did the kinome wide DSF measurements and ADP glo assay; TALE performed NanoBRET measurements with the help of our student Fabian Göllner; Scientific supervision by SM, TH and SK.

## Acknowledgement

FAG, AK, JG, TE, LW, TH, SK are grateful for support by the Structural Genomics Consortium (SGC), a registered charity (no: 1097737) that receives funds from Bayer AG, Boehringer Ingelheim, Bristol Myers Squibb, Genentech, Genome Canada through Ontario Genomics Institute, EU/EFPIA/OICR/McGill/KTH/Diamond Innovative Medicines Initiative 2 Joint Undertaking [EUbOPEN grant 875510], Janssen, Pfizer and Takeda. We would like to thank Takeda for helping with the kinome-wide tracer displacement assay. AK and SK are grateful for support by the German Cancer Research Center DKTK and AK, SM and SK by the Frankfurt Cancer Institute (FCI). FAG would like to acknowledge support by the LOEWE project TRABITA and TH by the program ENABLE. We thank the staff at the Swiss Light Source (CH) and Diamond Light Source (UK) for assistance during X-ray data collection.

## Abbreviations

ATP: adenosine triphosphate
CDK: Cyclin-dependent kinase
CK2: casein kinase-2
CLK: Cdc2-like kinase
Cy_2_NME: *N,N*-dicyclohexylmethylamine
DABSO: DABCO(1,4-diazabicyclo[2.2. 2]octane)-Bis(sulfur dioxide)
DAPK: Death associated protein kinase
DCM: dichloromethane
DMAP: 4-dimethylaminopyridine
DME: 1,2-dimethoxyethane
DMF: *N,N*-dimethylformamide
DMSO: dimethyl sulfoxide
DSF: differential scanning fluorimetry
DYRK: dual-specificity tyrosine-regulated kinase
FLT: fms related receptor tyrosine kinase
GTP: guanosine triphosphate
HIPK: Homeodomain-interacting protein kinase
*i*-PrOH: isopropanol
ITC: isothermal titration calorimetry
MAPK: mitogen-activated protein kinase
N(Et)_3_: triethylamine
*N*-Boc: *N*-*tert*-butyloxycarbonyl
NFSI: *N*-Fluorobenzenesulfonimide
on: overnight
PDB: Protein Data Base
PEG: polyethylene glycol
PIM: Pim-1 Proto-Oncogene Serine/Threonine Kinase
PK: protein kinase
rt: room temperature
SAR: structure activity relationship
SGC: Structural Genomic Consortium
S_N_2: bimolecular nucleophilic substitution
TBK: TANK-binding kinase
THF: tetrahydrofuran
Tm shift: thermal shift
µW: microwave.

## References

(1) Röhm, S.; Krämer, A.; Knapp, S. Function, Structure and Topology of Protein Kinases. In Proteinkinase Inhibitors; Laufer, S., Ed.; Topics in Medicinal Chemistry; Springer, 2021; pp 1–24. DOI: 10.1007/7355_2020_97.

(2) Fry, A. M.; O’Regan, L.; Sabir, S. R.; Bayliss, R. Cell cycle regulation by the NEK family of protein kinases. Journal of cell science 2012, 125 (Pt 19), 4423–4433. DOI: 10.1242/jcs.111195. Published Online: Nov. 6, 2012.

(3) Fortner, A.; Chera, A.; Tanca, A.; Bucur, O. Apoptosis regulation by the tyrosine-protein kinase CSK. Frontiers in cell and developmental biology 2022, 10, 1078180. DOI: 10.3389/fcell.2022.1078180. Published Online: Dec. 12, 2022.

(4) Singh, P.; Ravanan, P.; Talwar, P. Death Associated Protein Kinase 1 (DAPK1): A Regulator of Apoptosis and Autophagy. Frontiers in molecular neuroscience 2016, 9, 46. DOI: 10.3389/fnmol.2016.00046. Published Online: Jun. 23, 2016.

(5) Hluchý, M.; Gajdušková, P.; Ruiz de Los Mozos, I.; Rájecký, M.; Kluge, M.; Berger, B.-T.; Slabá, Z.; Potěšil, D.; Weiß, E.; Ule, J.; Zdráhal, Z.; Knapp, S.; Paruch, K.; Friedel, C. C.; Blazek, D. CDK11 regulates pre-mRNA splicing by phosphorylation of SF3B1. Nature 2022, 609 (7928), 829–834. DOI: 10.1038/s41586-022-05204-z. Published Online: Sep. 14, 2022.

(6) Lonskaya, I.; Hebron, M. L.; Desforges, N. M.; Franjie, A.; Moussa, C. E.-H. Tyrosine kinase inhibition increases functional parkin-Beclin-1 interaction and enhances amyloid clearance and cognitive performance. EMBO molecular medicine 2013, 5 (8), 1247–1262. DOI: 10.1002/emmm.201302771. Published Online: Jul. 4, 2013.

(7) Laufer, S., Ed. Proteinkinase Inhibitors; Topics in Medicinal Chemistry; Springer, 2021. DOI: 10.1007/978-3-030-68180-7.

(8) Patterson, H.; Nibbs, R.; McInnes, I.; Siebert, S. Protein kinase inhibitors in the treatment of inflammatory and autoimmune diseases. Clinical and experimental immunology 2014, 176 (1), 1–10. DOI: 10.1111/cei.12248.

(9) Montenarh, M.; Götz, C. The Interactome of Protein Kinase CK2. In Protein kinase CK2; Pinna, L. A., Ed.; The Wiley-IUBMB series on biochemistry and molecular biology; Wiley-Blackwell, 2013; pp 76–116. DOI: 10.1002/9781118482490.ch2.

(10) Pinna, L. A., Ed. Protein kinase CK2; The Wiley-IUBMB series on biochemistry and molecular biology; Wiley-Blackwell, 2013. DOI: 10.1002/9781118482490.

(11) Buljan, M.; Ciuffa, R.; van Drogen, A.; Vichalkovski, A.; Mehnert, M.; Rosenberger, G.; Lee, S.; Varjosalo, M.; Pernas, L. E.; Spegg, V.; Snijder, B.; Aebersold, R.; Gstaiger, M. Kinase Interaction Network Expands Functional and Disease Roles of Human Kinases. Molecular cell 2020, 79 (3), 504–520.e9. DOI: 10.1016/j.molcel.2020.07.001. Published Online: Jul. 23, 2020.

(12) Seldin, D. C.; Leder, P. Casein kinase II alpha transgene-induced murine lymphoma: relation to theileriosis in cattle. Science (New York, N.Y.) 1995, 267 (5199), 894–897. DOI: 10.1126/science.7846532.

(13) Lashuel, H. A.; Overk, C. R.; Oueslati, A.; Masliah, E. The many faces of α-synuclein: from structure and toxicity to therapeutic target. Nature reviews. Neuroscience 2013, 14 (1), 38–48. DOI: 10.1038/nrn3406.

(14) Marshall, C. A.; McBride, J. D.; Changolkar, L.; Riddle, D. M.; Trojanowski, J. Q.; Lee, V. M.-Y. Inhibition of CK2 mitigates Alzheimer’s tau pathology by preventing NR2B synaptic mislocalization. Acta neuropathologica communications 2022, 10 (1), 30. DOI:10.1186/s40478-022-01331-w. Published Online: Mar. 4, 2022.

(15) Pastori, V.; Sangalli, E.; Coccetti, P.; Pozzi, C.; Nonnis, S.; Tedeschi, G.; Fusi, P. CK2 and GSK3 phosphorylation on S29 controls wild-type ATXN3 nuclear uptake. Biochimica et biophysica acta 2010, 1802 (7-8), 583–592. DOI: 10.1016/j.bbadis.2010.03.007. Published Online: Mar. 27, 2010.

(16) Chen, I.-Y.; Chang, S. C.; Wu, H.-Y.; Yu, T.-C.; Wei, W.-C.; Lin, S.; Chien, C.-L.; Chang, M.-F. Upregulation of the chemokine (C-C motif) ligand 2 via a severe acute respiratory syndrome coronavirus spike-ACE2 signaling pathway. Journal of virology 2010, 84 (15), 7703–7712. DOI: 10.1128/JVI.02560-09. Published Online: May. 19, 2010.

(17) Firzlaff, J. M.; Galloway, D. A.; Eisenman, R. N.; Lüscher, B. The E7 protein of human papillomavirus type 16 is phosphorylated by casein kinase II. The New biologist 1989, 1 (1), 44–53.

(18) Quezada Meza, C. P.; Ruzzene, M. Protein Kinase CK2 and SARS-CoV-2: An Expected Interplay Story. Kinases and Phosphatases 2023, 1 (2), 141–150. DOI: 10.3390/kinasesphosphatases1020009.

(19) Yang, X.; Dickmander, R. J.; Bayati, A.; Taft-Benz, S. A.; Smith, J. L.; Wells, C. I.; Madden, E. A.; Brown, J. W.; Lenarcic, E. M.; Yount, B. L.; Chang, E.; Axtman, A. D.; Baric, R. S.; Heise, M. T.; McPherson, P. S.; Moorman, N. J.; Willson, T. M. Host Kinase CSNK2 is a Target for Inhibition of Pathogenic SARS-like β-Coronaviruses. ACS chemical biology 2022, 17 (7), 1937–1950. DOI: 10.1021/acschembio.2c00378. Published Online: Jun. 19, 2022.

(20) Augustine, S. A. J.; Kleshchenko, Y. Y.; Nde, P. N.; Pratap, S.; Ager, E. A.; Burns, J. M.; Lima, M. F.; Villalta, F. Molecular cloning of a Trypanosoma cruzi cell surface casein kinase II substrate, Tc-1, involved in cellular infection. Infection and immunity 2006, 74 (7), 3922–3929. DOI: 10.1128/IAI.00045-06.

(21) ole-MoiYoi, O. K. Casein kinase II in theileriosis. Science (New York, N.Y.) 1995, 267 (5199), 834–836. DOI: 10.1126/science.7846527.

(22) Krämer, A.; Kurz, C. G.; Berger, B.-T.; Celik, I. E.; Tjaden, A.; Greco, F. A.; Knapp, S.; Hanke, T. Optimization of pyrazolo1,5-apyrimidines lead to the identification of a highly selective casein kinase 2 inhibitor. European journal of medicinal chemistry 2020, 208, 112770. DOI: 10.1016/j.ejmech.2020.112770. Published Online: Aug. 23, 2020.

(23) Lindenblatt, D.; Applegate, V.; Nickelsen, A.; Klußmann, M.; Neundorf, I.; Götz, C.; Jose, J.; Niefind, K. Molecular Plasticity of Crystalline CK2α’ Leads to KN2, a Bivalent Inhibitor of Protein Kinase CK2 with Extraordinary Selectivity. Journal of medicinal chemistry 2022, 65 (2), 1302–1312. DOI: 10.1021/acs.jmedchem.1c00063. Published Online: Jul. 29, 2021.

(24) Iegre, J.; Atkinson, E. L.; Brear, P. D.; Cooper, B. M.; Hyvönen, M.; Spring, D. R. Chemical probes targeting the kinase CK2: a journey outside the catalytic box. Organic & biomolecular chemistry 2021, 19 (20), 4380–4396. DOI: 10.1039/d1ob00257k.

(25) Kufareva, I.; Bestgen, B.; Brear, P.; Prudent, R.; Laudet, B.; Moucadel, V.; Ettaoussi, M.; Sautel, C. F.; Krimm, I.; Engel, M.; Filhol, O.; Le Borgne, M.; Lomberget, T.; Cochet, C.; Abagyan, R. Discovery of holoenzyme-disrupting chemicals as substrate-selective CK2 inhibitors. Scientific reports 2019, 9 (1), 15893. DOI: 10.1038/s41598-019-52141-5. Published Online: Nov. 4, 2019.

(26) Borgo, C.; D’Amore, C.; Sarno, S.; Salvi, M.; Ruzzene, M. Protein kinase CK2: a potential therapeutic target for diverse human diseases. Signal transduction and targeted therapy 2021, 6 (1), 183. DOI: 10.1038/s41392-021-00567-7. Published Online: May. 17, 2021.

(27) ClinicalTrials.gov.

(28) Vahter, J.; Viht, K.; Uri, A.; Enkvist, E. Oligo-aspartic acid conjugates with benzoc2,6naphthyridine-8-carboxylic acid scaffold as picomolar inhibitors of CK2. Bioorganic & medicinal chemistry 2017, 25 (7), 2277–2284. DOI: 10.1016/j.bmc.2017.02.055. Published Online: Feb. 28, 2017.

(29) Pierre, F.; Chua, P. C.; O’Brien, S. E.; Siddiqui-Jain, A.; Bourbon, P.; Haddach, M.; Michaux, J.; Nagasawa, J.; Schwaebe, M. K.; Stefan, E.; Vialettes, A.; Whitten, J. P.; Chen, T. K.; Darjania, L.; Stansfield, R.; Bliesath, J.; Drygin, D.; Ho, C.; Omori, M.; Proffitt, C.; Streiner, N.; Rice, W. G.; Ryckman, D. M.; Anderes, K. Pre-clinical characterization of CX-4945, a potent and selective small molecule inhibitor of CK2 for the treatment of cancer. Molecular and cellular biochemistry 2011, 356 (1-2), 37–43. DOI: 10.1007/s11010-011-0956-5. Published Online: Jul. 14, 2011.

(30) Robers, M. B.; Friedman-Ohana, R.; Huber, K. V. M.; Kilpatrick, L.; Vasta, J. D.; Berger, B.-T.; Chaudhry, C.; Hill, S.; Müller, S.; Knapp, S.; Wood, K. V. Quantifying Target Occupancy of Small Molecules Within Living Cells. Annual review of biochemistry 2020, 89, 557–581. DOI: 10.1146/annurev-biochem-011420-092302. Published Online: Mar. 24, 2020.

(31) Wells, C. I.; Drewry, D. H.; Pickett, J. E.; Tjaden, A.; Krämer, A.; Müller, S.; Gyenis, L.; Menyhart, D.; Litchfield, D. W.; Knapp, S.; Axtman, A. D. Development of a potent and selective chemical probe for the pleiotropic kinase CK2. Cell chemical biology 2021, 28 (4), 546–558.e10. DOI: 10.1016/j.chembiol.2020.12.013. Published Online: Jan. 22, 2021.

(32) Hu, R.; Xu, H.; Jia, P.; Zhao, Z. KinaseMD: kinase mutations and drug response database. Nucleic acids research 2021, 49 (D1), D552–D561. DOI: 10.1093/nar/gkaa945.

(33) Zhou, Y.; Xiang, S.; Yang, F.; Lu, X. Targeting Gatekeeper Mutations for Kinase Drug Discovery. Journal of medicinal chemistry 2022, 65 (23), 15540–15558. DOI: 10.1021/acs.jmedchem.2c01361. Published Online: Nov. 17, 2022.

(34) Lyczek, A.; Berger, B.-T.; Rangwala, A. M.; Paung, Y.; Tom, J.; Philipose, H.; Guo, J.; Albanese, S. K.; Robers, M. B.; Knapp, S.; Chodera, J. D.; Seeliger, M. A. Mutation in Abl kinase with altered drug-binding kinetics indicates a novel mechanism of imatinib resistance. Proceedings of the National Academy of Sciences of the United States of America 2021, 118(46). DOI: 10.1073/pnas.2111451118.

(35) Salvi, M.; Borgo, C.; Pinna, L. A.; Ruzzene, M. Targeting CK2 in cancer: a valuable strategy or a waste of time? Cell death discovery 2021, 7 (1), 325. DOI: 10.1038/s41420-021-00717-4. Published Online: Oct. 29, 2021.

(36) Niefind, K.; Guerra, B.; Ermakowa, I.; Issinger, O. G. Crystal structure of human protein kinase CK2: insights into basic properties of the CK2 holoenzyme. The EMBO journal 2001, 20 (19), 5320–5331. DOI: 10.1093/emboj/20.19.5320.

(37) Fusco, C. de; Brear, P.; Iegre, J.; Georgiou, K. H.; Sore, H. F.; Hyvönen, M.; Spring, D. R. A fragment-based approach leading to the discovery of a novel binding site and the selective CK2 inhibitor CAM4066. Bioorganic & medicinal chemistry 2017, 25 (13), 3471–3482. DOI: 10.1016/j.bmc.2017.04.037. Published Online: Apr. 30, 2017.

(38) Brear, P.; Fusco, C. de; Hadje Georgiou, K.; Francis-Newton, N. J.; Stubbs, C. J.; Sore, H. F.; Venkitaraman, A. R.; Abell, C.; Spring, D. R.; Hyvönen, M. Specific inhibition of CK2α from an anchor outside the active site. Chemical science 2016, 7 (11), 6839–6845. DOI:10.1039/c6sc02335e. Published Online: Jul. 12, 2016.

(39) Bancet, A.; Frem, R.; Jeanneret, F.; Mularoni, A.; Bazelle, P.; Roelants, C.; Delcros, J.-G.; Guichou, J.-F.; Pillet, C.; Coste, I.; Renno, T.; Battail, C.; Cochet, C.; Lomberget, T.; Filhol, O.; Krimm, I. Cancer selective cell death induction by a bivalent CK2 inhibitor targeting the ATP site and the allosteric αD pocket. iScience 2024, 27 (2), 108903. DOI:10.1016/j.isci.2024.108903. Published Online: Jan. 12, 2024.

(40) Litchfield, D. W. Protein kinase CK2: structure, regulation and role in cellular decisions of life and death.

(41) Battistutta, R.; Ranchio, A.; Papinutto, E. Crystal structure of the apo-form of human CK2 alpha at pH 8.5, 2011. DOI: 10.2210/pdb3q04/pdb.

(42) Iegre, J.; Brear, P.; Fusco, C. de; Yoshida, M.; Mitchell, S. L.; Rossmann, M.; Carro, L.; Sore, H. F.; Hyvönen, M.; Spring, D. R. Second-generation CK2α inhibitors targeting the αD pocket. Chemical science 2018, 9 (11), 3041–3049. DOI: 10.1039/c7sc05122k. Published Online: Feb. 20, 2018.

(43) Woolven, H.; González-Rodríguez, C.; Marco, I.; Thompson, A. L.; Willis, M. C. DABCO-bis(sulfur dioxide), DABSO, as a convenient source of sulfur dioxide for organic synthesis: utility in sulfonamide and sulfamide preparation. Organic letters 2011, 13 (18), 4876–4878. DOI: 10.1021/ol201957n. Published Online: Aug. 25, 2011.

(44) Davies, A. T.; Curto, J. M.; Bagley, S. W.; Willis, M. C. One-pot palladium-catalyzed synthesis of sulfonyl fluorides from aryl bromides. Chemical science 2017, 8 (2), 1233–1237. DOI: 10.1039/c6sc03924c. Published Online: Oct. 11, 2016.

(45) Gadd, M. S.; Testa, A.; Lucas, X.; Chan, K.-H.; Chen, W.; Lamont, D. J.; Zengerle, M.; Ciulli, A. Structural basis of PROTAC cooperative recognition for selective protein degradation. Nature chemical biology 2017, 13 (5), 514–521. DOI: 10.1038/nchembio.2329. Published Online: Mar. 13, 2017.

(46) Musumeci, F.; Cianciusi, A.; D’Agostino, I.; Grossi, G.; Carbone, A.; Schenone, S. Synthetic Heterocyclic Derivatives as Kinase Inhibitors Tested for the Treatment of Neuroblastoma. Molecules (Basel, Switzerland) 2021, 26 (23). DOI:10.3390/molecules26237069. Published Online: Nov. 23, 2021.

(47) Taruneshwar Jha, K.; Shome, A.; Chahat; Chawla, P. A. Recent advances in nitrogen-containing heterocyclic compounds as receptor tyrosine kinase inhibitors for the treatment of cancer: Biological activity and structural activity relationship. Bioorganic chemistry 2023, 138, 106680. DOI: 10.1016/j.bioorg.2023.106680. Published Online: Jun. 17, 2023.

(48) Monteleone, S.; Fuchs, J. E.; Liedl, K. R. Molecular Connectivity Predefines Polypharmacology: Aliphatic Rings, Chirality, and sp3 Centers Enhance Target Selectivity. Frontiers in pharmacology 2017, 8, 552. DOI: 10.3389/fphar.2017.00552. Published Online: Aug. 28, 2017.

(49) Shearer, J.; Castro, J. L.; Lawson, A. D. G.; MacCoss, M.; Taylor, R. D. Rings in Clinical Trials and Drugs: Present and Future. Journal of medicinal chemistry 2022, 65 (13), 8699– 8712. DOI: 10.1021/acs.jmedchem.2c00473. Published Online: Jun. 22, 2022.

(50) Hirozane, Y.; Toyofuku, M.; Yogo, T.; Tanaka, Y.; Sameshima, T.; Miyahisa, I.; Yoshikawa, M. Structure-based rational design of staurosporine-based fluorescent probe with broad-ranging kinase affinity for kinase panel application. Bioorganic & medicinal chemistry letters 2019, 29 (21), 126641. DOI: 10.1016/j.bmcl.2019.126641. Published Online: Sep. 7, 2019.

(51) Vasta, J. D.; Corona, C. R.; Wilkinson, J.; Zimprich, C. A.; Hartnett, J. R.; Ingold, M. R.; Zimmerman, K.; Machleidt, T.; Kirkland, T. A.; Huwiler, K. G.; Ohana, R. F.; Slater, M.; Otto, P.; Cong, M.; Wells, C. I.; Berger, B.-T.; Hanke, T.; Glas, C.; Ding, K.; Drewry, D. H.; Huber, K. V. M.; Willson, T. M.; Knapp, S.; Müller, S.; Meisenheimer, P. L.; Fan, F.; Wood, K. V.; Robers, M. B. Quantitative, Wide-Spectrum Kinase Profiling in Live Cells for Assessing the Effect of Cellular ATP on Target Engagement. Cell chemical biology 2018, 25 (2), 206–214.e11. DOI: 10.1016/j.chembiol.2017.10.010. Published Online: Nov. 22, 2017.

(52) Raaf, J.; Klopffleisch, K.; Issinger, O.-G.; Niefind, K. The catalytic subunit of human protein kinase CK2 structurally deviates from its maize homologue in complex with the nucleotide competitive inhibitor emodin. Journal of molecular biology 2008, 377 (1), 1–8. DOI:10.1016/j.jmb.2008.01.008. Published Online: Jan. 11, 2008.

(53) Battistutta, R.; Moliner, E. de; Sarno, S.; Zanotti, G.; Pinna, L. A. Structural features underlying selective inhibition of protein kinase CK2 by ATP site-directed tetrabromo-2-benzotriazole. Protein science : a publication of the Protein Society 2001, 10 (11), 2200– 2206. DOI: 10.1110/ps.19601.

(54) Oshima, T.; Niwa, Y.; Kuwata, K.; Srivastava, A.; Hyoda, T.; Tsuchiya, Y.; Kumagai, M.; Tsuyuguchi, M.; Tamaru, T.; Sugiyama, A.; Ono, N.; Zolboot, N.; Aikawa, Y.; Oishi, S.; Nonami, A.; Arai, F.; Hagihara, S.; Yamaguchi, J.; Tama, F.; Kunisaki, Y.; Yagita, K.; Ikeda, M.; Kinoshita, T.; Kay, S. A.; Itami, K.; Hirota, T. Cell-based screen identifies a new potent and highly selective CK2 inhibitor for modulation of circadian rhythms and cancer cell growth. Science advances 2019, 5 (1), eaau9060. DOI: 10.1126/sciadv.aau9060. Published Online: Jan. 23, 2019.

(55) Sarno, S.; Ghisellini, P.; Pinna, L. A. Unique activation mechanism of protein kinase CK2. The N-terminal segment is essential for constitutive activity of the catalytic subunit but not of the holoenzyme. The Journal of biological chemistry 2002, 277 (25), 22509–22514. DOI:10.1074/jbc.M200486200. Published Online: Apr. 15, 2002.

(56) Zhou, B.; Ritt, D. A.; Morrison, D. K.; Der, C. J.; Cox, A. D. Protein Kinase CK2α Maintains Extracellular Signal-regulated Kinase (ERK) Activity in a CK2α Kinase-independent Manner to Promote Resistance to Inhibitors of RAF and MEK but Not ERK in BRAF Mutant Melanoma. The Journal of biological chemistry 2016, 291 (34), 17804–17815. DOI:10.1074/jbc.M115.712885. Published Online: May. 17, 2016.

