## Supporting Information for "Synthesis and evaluation of chemical linchpins for highly selective CK2α targeting"

#### Table of contents:

**Table S1**: Selectivity data of **26** from kinome wide tracer displacement screen 2

**Table S2**: Selectivity of compounds **27, 48, 50, 51, 55, 57** measured in in-house Tm assay 3

**Table S3:** Fragment study using Tm 4

**Figure S1:** Overlay of **25** and **CX-4945** 4

**Experimental section** 5

**Analytical data of compounds** 52

**Data collection and refinement stastitics** 208

**References** 389

**Table S1**: Selectivity data of **26** from kinome wide tracer displacement screen

| **Kinase** | **% Inhibition** | **Kinase** | **% Inhibition** | **Kinase** | **% Inhibition** | **Kinase** | **% Inhibition** |
|---|---|---|---|---|---|---|---|
| CSNK2a1 | 92 | PRKaa2 | 3 | aXL | 0 | CLK3 | 0 |
| MELK | 53 | aBL1 | 3 | CSNK1G1 | 0 | CHEK1 | 0 |
| ROCK2 | 39 | PRKG2 | 3 | CDK2 | 0 | MaP2K2 | 0 |
| PRKD3 | 35 | WEE1 | 3 | PLK1 | 0 | TLK1 | 0 |
| MaPK15 | 24 | aaK1 | 2 | MaPK12 | 0 | CDK11a | 0 |
| RPS6KB1 | 12 | DYRK1a | 2 | MaP3K14 | 0 | MaPK1 | 0 |
| MaPK8 | 11 | PTK6 | 2 | MaPK13 | 0 | CDK12 | 0 |
| NUaK1 | 11 | DCLK3 | 2 | EIF2aK2 | 0 | CDK16 | 0 |
| LRRK2 | 10 | TNK2 | 2 | CDK4 | 0 | PINK1 | 0 |
| PaK6 | 10 | ITK | 1 | BTK | 0 | INSR | 0 |
| TNK1 | 9 | TYK2 | 1 | CDK1 | 0 | CaSK | 0 |
| SLK | 8 | EPHB4 | 1 | CaMK2G | 0 | SgK110 | 0 |
| TSSK3 | 8 | FES | 0 | STK33 | 0 | BLK | 0 |
| PRKCH | 7 | TaOK3 | 0 | GSK3B | 0 | ULK3 | 0 |
| MaPKaPK5 | 7 | CDK9 | 0 | YES1 | 0 | STK32a | 0 |
| VRK1 | 6 | SgK085 | 0 | CSK | 0 | MST1R | 0 |
| DYRK4 | 6 | MaP3K15 | 0 | FGFR4 | 0 | BRaF | 0 |
| MaPK10 | 6 | ZaP70 | 0 | HIPK1 | 0 | GaK | 0 |
| CaMK1G | 6 | JaK2 | 0 | CDK11B | 0 | MaPK6 | 0 |
| FLT3 | 6 | MaPK14 | 0 | aKT1 | 0 | OXSR1 | 0 |
| NEK1 | 5 | MaRK4 | 0 | BRSK1 | 0 | STK3 | 0 |
| FGFR3 | 5 | TESK1 | 0 | BUB1 | 0 | CaMK2a | 0 |
| MaPK9 | 5 | IRaK4 | 0 | CHEK2 | 0 | FGR | 0 |
| TXK | 5 | RPS6Ka1 | 0 | aLK | 0 | | |
| SGK3 | 5 | JaK3 | 0 | STRaDa | 0 | | |
| TaOK2 | 4 | PI3KCD | 0 | CDK17 | 0 | | |
| LIMK1 | 4 | CaMKK2 | 0 | PSKH1 | 0 | | |

**Table S2**: Selectivity of compounds **27, 48, 50, 51, 55, 57** measured in in-house Tm assay

| 27 | | 48 | | 50 | | 51 | | 55 | | 57 | |
|---|---|---|---|---|---|---|---|---|---|---|---|
| Kinase | Δ Tm [°C] | Kinase | Δ Tm [°C] | Kinase | Δ Tm [°C] | Kinase | Δ Tm [°C] | Kinase | Δ Tm [°C] | Kinase | Δ Tm [°C] |
| CSNK2A1A | 7.49 | CSNK2A1A | 7.66 | CSNK2A1A | 7.31 | CSNK2A1A | 4.56 | CSNK2A1A | 8.51 | CSNK2A1A | 6.92 |
| CSNK2A2A | 3.01 | STK17AA | 5.34 | CSNK2A2A | 5.27 | CSNK2A2A | 2.46 | CSNK2A2A | 4.82 | CSNK2A2A | 4.06 |
| WNK1A | 2.31 | CSNK2A2A | 5.12 | ULK3A | 4.14 | TAF1A | 1.15 | TAF1A | 1.79 | BRPF1B | 2.61 |
| TAF1A | 1.84 | BMP2KA | 5.05 | AURKBA | 1.36 | PIM3A | 0.38 | NEK1A | 1.05 | TAF1A | 0.64 |
| AURKBA | 1.30 | STK6A | 4.82 | MAPK15HSD | 1.34 | NEK1A | 0.21 | ABL1A | 0.66 | MELKA | 0.47 |
| ABL1A | 0.95 | AURKBA | 4.24 | MELKA | 1.18 | STK6A | 0.19 | WNK1A | 0.56 | CASKA | 0.36 |
| CSNK1DA | 0.82 | MELKA | 3.85 | CHEK2A | 0.98 | MAPK13A | 0.18 | ULK3A | 0.15 | EPHA4A | 0.26 |
| STK6A | 0.73 | MAPK15HSD | 3.42 | STK17AA | 0.75 | ABL1A | 0.13 | EPHA4A | 0.10 | ABL1A | 0.19 |
| BRAFA | 0.63 | ULK3A | 2.45 | TAF1A | 0.74 | CASKA | 0.12 | CASKA | 0.09 | STK6A | 0.16 |
| MAPK13A | 0.57 | RPS6KA5A | 1.46 | DAPK3A | 0.71 | PAK4A | 0.11 | STK6A | 0.06 | WNK1A | 0.14 |
| MAP3K5A | 0.51 | FLT1A | 1.11 | ABL1A | 0.69 | RPS6KA1A | 0.08 | MAPK13A | 0.05 | FESA | 0.13 |
| ULK1A | 0.47 | ABL1A | 1.06 | CASKA | 0.56 | ULK1A | 0.08 | AURKBA | 0.02 | MAPK13A | 0.11 |
| FESA | 0.36 | CSNK1EA | 0.89 | STK6A | 0.47 | STK17AA | 0.08 | MARK3A | 0.00 | NEK1A | 0.03 |
| EPHA4A | 0.34 | DAPK3A | 0.82 | BMP2KA | 0.35 | FLT1A | 0.03 | NEK2A | 0.00 | CSNK1DA | -0.01 |
| SRPK1A | 0.27 | CLK1A | 0.77 | MAPK13A | 0.22 | DYRK1AA | -0.04 | OSR1A | 0.00 | CDKL1A | -0.02 |
| MAPK10A | 0.27 | MAPK13A | 0.69 | FESA | 0.18 | MELKA | -0.04 | CSNK1EA | -0.16 | RPS6KA1A | -0.04 |
| BMXA | 0.25 | AAK1A | 0.65 | GSG2A | 0.13 | EPHA4A | -0.08 | FGFR2A | -0.18 | MARK4A | -0.04 |
| BMP2KA | 0.22 | WNK1A | 0.60 | PAK4A | 0.12 | MAPK1A | -0.10 | TTKA | -0.18 | BMPR2A | -0.06 |
| ULK3A | 0.18 | CSNK1DA | 0.49 | PIM3A | 0.12 | ULK3A | -0.11 | GAKA | -0.19 | AURKBA | -0.10 |
| MAPK8B | 0.18 | STK10A | 0.38 | WNK1A | 0.11 | EPHB1A | -0.11 | MAP3K5A | -0.21 | DAPK1A | -0.11 |
| RPS6KA1A | 0.12 | TAF1A | 0.37 | EPHA4A | 0.10 | MARK3A | -0.12 | SRCA | -0.23 | SRCA | -0.11 |
| DAPK1A | 0.10 | CAMK1GA | 0.34 | CSNK1DA | 0.09 | GAKA | -0.12 | RPS6KA1A | -0.25 | DYRK1AA | -0.13 |
| PCTK1A | 0.10 | FESA | 0.34 | CLK1A | 0.07 | BMPR2A | -0.13 | MAP2K6A | -0.29 | EPHB1A | -0.14 |
| PAK1A | 0.03 | DAPK1A | 0.29 | FLT1A | 0.06 | WNK1A | -0.15 | BMPR2A | -0.33 | MAP3K5A | -0.15 |
| CAMK2BA | 0.00 | CASKA | 0.25 | NEK1A | 0.04 | MAPK15HSD | -0.15 | CAMK2DA | -0.36 | CSNK1EA | -0.16 |
| GAKA | 0.00 | SRCA | 0.24 | DAPK1A | 0.02 | DAPK1A | -0.15 | ULK1A | -0.36 | ULK3A | -0.16 |
| MARK3A | 0.00 | MAP3K5A | 0.18 | BMXA | -0.02 | CSNK1DA | -0.15 | EPHB1A | -0.38 | TTKA | -0.17 |
| NEK1A | 0.00 | NEK1A | 0.16 | CSNK1EA | -0.03 | BRAFA | -0.17 | BMP2KA | -0.43 | PAK4A | -0.18 |
| OSR1A | 0.00 | ULK1A | 0.15 | TTKA | -0.05 | DYRK2A | -0.19 | DMPK1A | -0.43 | BMP2KA | -0.23 |
| CASKA | -0.01 | BMPR2A | 0.06 | CAMK2BA | -0.06 | SRCA | -0.19 | BRAFA | -0.44 | FGFR2A | -0.26 |
| SRCA | -0.02 | EPHA4A | 0.06 | PAK1A | -0.08 | MAP2K6A | -0.21 | AKT3A | -0.44 | GSG2A | -0.27 |
| BMPR2A | -0.03 | GPRK5A | 0.00 | MAPK10A | -0.09 | MAP3K5A | -0.22 | FLT1A | -0.44 | BRAFA | -0.28 |
| CSNK1EA | -0.03 | PLK4A | -0.05 | RPS6KA1A | -0.11 | STK39A | -0.22 | DCAMKL1A | -0.47 | MAP2K6A | -0.28 |
| FLT1A | -0.11 | MAPK10A | -0.07 | PLK4A | -0.12 | AKT3A | -0.22 | FGFR1B | -0.47 | FLT1A | -0.29 |
| EPHB1A | -0.11 | BRAFA | -0.09 | MAPK1A | -0.13 | CDC42BPAA | -0.23 | CSNK1DA | -0.49 | MAP2K1A | -0.34 |
| MAP2K6A | -0.12 | RPS6KA1A | -0.12 | RPS6KA5A | -0.14 | DMPK1A | -0.24 | SRPK1A | -0.50 | STK17AA | -0.35 |
| MAPK1A | -0.25 | STK4A | -0.17 | DMPK1A | -0.16 | FGFR2A | -0.24 | PCTK1A | -0.53 | MAPK1A | -0.36 |
| STK17AA | -0.25 | EPHB1A | -0.20 | MAP3K5A | -0.17 | BMP2KA | -0.24 | GPRK5A | -0.54 | ULK1A | -0.37 |
| PIM3A | -0.26 | MAP2K1A | -0.21 | EPHB1A | -0.22 | TTKA | -0.25 | MAPK1A | -0.54 | SRPK1A | -0.37 |
| MAP2K4A | -0.26 | BMXA | -0.23 | DYRK1AA | -0.25 | CDKL1A | -0.25 | MELKA | -0.55 | MAPK10A | -0.37 |
| FGFR1B | -0.30 | FGFR2A | -0.30 | SRCA | -0.29 | TIF1A | -0.29 | CLK1A | -0.58 | DMPK1A | -0.38 |
| MAP2K1A | -0.32 | DMPK1A | -0.30 | BRAFA | -0.30 | FGFR1B | -0.29 | MAPK10A | -0.61 | CAMK2BA | -0.40 |
| AAK1A | -0.33 | EPHA7A | -0.32 | FGFR2A | -0.33 | MAP2K1A | -0.31 | CAMK2BA | -0.69 | VRK1A | -0.40 |
| FGFR2A | -0.34 | FGFR1B | -0.34 | FGFR1B | -0.35 | CSNK1EA | -0.33 | CLK3A | -0.73 | FGFR1B | -0.43 |
| STK17BA | -0.35 | MARK4A | -0.35 | MSSK1A | -0.35 | MAPKAPK2A | -0.33 | AAK1A | -0.73 | BMXA | -0.45 |
| DMPK1A | -0.36 | CHEK2A | -0.37 | ULK1A | -0.35 | MAPK14A | -0.36 | PAK4A | -0.75 | PIM1A | -0.45 |
| EPHA2A | -0.37 | TTKA | -0.39 | OSR1A | -0.36 | NQO2A | -0.37 | MAPK8B | -0.75 | HIPK2HSF | -0.48 |
| DYRK2A | -0.38 | CAMK1DA | -0.40 | SRPK1A | -0.36 | PLK4A | -0.37 | MAP2K4A | -0.79 | MAPKAPK2A | -0.50 |
| CDKL1A | -0.41 | MAPK1A | -0.40 | MAP2K1A | -0.37 | SRPK1A | -0.40 | EPHA2A | -0.80 | MAPK15HSD | -0.51 |
| CLK1A | -0.41 | AKT3A | -0.42 | MAPK9A | -0.37 | MAP2K4A | -0.41 | CHEK2A | -0.80 | STK4A | -0.52 |
| BRPF1B | -0.45 | SRPK1A | -0.42 | MAP2K6A | -0.39 | EPHA7A | -0.41 | MAP2K1A | -0.80 | DYRK2A | -0.52 |
| CLK3A | -0.46 | BRPF1B | -0.42 | CDKL1A | -0.41 | CAMK1GA | -0.41 | DYRK1AA | -0.81 | CAMK1GA | -0.53 |
| DYRK1AA | -0.47 | PIM3A | -0.44 | BRPF1B | -0.43 | CLK3A | -0.42 | STK39A | -0.84 | STK39A | -0.54 |
| MELKA | -0.58 | MAP2K6A | -0.49 | DYRK2A | -0.44 | EPHA5A | -0.44 | VRK1A | -0.85 | MSSK1A | -0.57 |
| MAPK14A | -0.61 | EPHA2A | -0.49 | CLK3A | -0.44 | HIPK2HSF | -0.46 | MSSK1A | -0.87 | PCTK1A | -0.57 |
| PAK4A | -0.62 | CAMK2BA | -0.53 | BMPR2A | -0.46 | BRPF1B | -0.46 | MERTKA | -0.88 | AKT3A | -0.58 |
| AKT3A | -0.70 | DYRK2A | -0.54 | PCTK1A | -0.48 | BMXA | -0.46 | EPHA7A | -0.89 | CLK1A | -0.58 |
| MAPK15HSD | -0.73 | VRK1A | -0.54 | EPHA5A | -0.48 | GPRK5A | -0.47 | NQO2A | -0.91 | EPHA2A | -0.59 |
| DCAMKL1A | -0.73 | CLK3A | -0.55 | CAMK1GA | -0.49 | MAPK8B | -0.47 | CDKL1A | -0.91 | CHEK2A | -0.60 |
| CAMK1DA | -0.74 | PCTK1A | -0.58 | CAMK2DA | -0.51 | MAPK10A | -0.48 | PIM3A | -0.94 | MAPK8B | -0.62 |
| EPHA5A | -0.77 | NQO2A | -0.58 | VRK1A | -0.54 | MSSK1A | -0.49 | DAPK1A | -0.98 | MAP2K4A | -0.64 |
| CHEK2A | -0.78 | PIM1A | -0.59 | EPHA2A | -0.56 | VRK1A | -0.49 | DAPK3A | -1.01 | CLK3A | -0.64 |
| VRK1A | -0.82 | CDKL1A | -0.60 | MAPK14A | -0.57 | PKMYT1A | -0.50 | BRPF1B | -1.02 | DAPK3A | -0.65 |
| MAPKAPK2A | -0.83 | MAPK8B | -0.60 | PKMYT1A | -0.58 | CAMK2DA | -0.50 | FESA | -1.03 | PLK4A | -0.68 |
| PLK4A | -0.83 | PAK4A | -0.61 | NEK7A | -0.58 | CHEK2A | -0.52 | MAPK15HSD | -1.04 | EPHA7A | -0.69 |
| EPHA7A | -0.89 | CAMK2DA | -0.64 | AKT3A | -0.59 | STK17BA | -0.52 | HIPK2HSF | -1.04 | NQO2A | -0.69 |
| PIM1A | -0.92 | PHKG2A | -0.65 | AAK1A | -0.60 | PIM1A | -0.53 | PKMYT1A | -1.04 | CDC42BPAA | -0.71 |
| STK3A | -0.94 | MAP2K4A | -0.67 | EPHA7A | -0.68 | AURKBA | -0.54 | STK4A | -1.10 | PIM3A | -0.72 |
| NQO2A | -0.95 | MARK3A | -0.73 | MAP2K4A | -0.75 | PCTK1A | -0.54 | MAPKAPK2A | -1.11 | GPRK5A | -0.74 |
| MSSK1A | -1.04 | EPHA5A | -0.82 | NQO2A | -0.75 | CLK1A | -0.60 | PIM1A | -1.15 | TIF1A | -0.75 |
| HIPK2HSF | -1.09 | STK17BA | -0.86 | MAPK8B | -0.76 | STK4A | -0.64 | STK17AA | -1.15 | EPHA5A | -0.78 |
| MERTKA | -1.11 | GAKA | -0.88 | STK4A | -0.76 | AAK1A | -0.64 | STK3A | -1.17 | AAK1A | -0.78 |
| STK39A | -1.13 | MSSK1A | -0.88 | GAKA | -0.77 | MARK4A | -0.68 | CAMK1GA | -1.17 | MAPK14A | -0.78 |
| CAMK2DA | -1.15 | DCAMKL1A | -0.90 | MAPKAPK2A | -0.79 | CAMK4A | -0.68 | NEK7A | -1.17 | GAKA | -0.82 |
| TTKA | -1.16 | MAPK1A | -0.91 | PIM1A | -0.84 | SLKA | -0.68 | EPHA5A | -1.18 | CAMK2DA | -0.84 |
| PKMYT1A | -1.27 | HIPK2HSF | -0.93 | HIPK2HSF | -0.85 | MAPK9A | -0.71 | CAMK4A | -1.24 | PAK1A | -0.86 |
| CAMK1GA | -1.31 | STK39A | -0.94 | STK3A | -0.89 | EPHA2A | -0.73 | TIF1A | -1.25 | SLKA | -0.90 |
| PHKG2A | -1.35 | DYRK1AA | -0.94 | STK17BA | -0.92 | CDK2A | -0.84 | CDC42BPAA | -1.28 | MARK3A | -0.91 |
| NEK7A | -1.41 | MAPKAPK2A | -1.00 | STK39A | -0.93 | STK38LA | -0.84 | MARK4A | -1.31 | STK3A | -0.96 |
| STK38LA | -1.44 | OSR1A | -1.13 | GPRK5A | -0.95 | PAK1A | -0.85 | RPS6KA5A | -1.32 | MERTKA | -0.97 |
| CAMK4A | -1.45 | MAPK14A | -1.14 | MERTKA | -0.95 | FESA | -0.92 | MAPK14A | -1.34 | PHKG2A | -1.07 |
| CDK2A | -1.47 | STK3A | -1.17 | MARK4A | -0.99 | MERTKA | -0.94 | PLK4A | -1.35 | CDK2A | -1.09 |
| SLKA | -1.51 | MERTKA | -1.24 | DCAMKL1A | -1.02 | PHKG2A | -0.96 | DYRK2A | -1.45 | STK38LA | -1.10 |
| STK4A | -1.55 | MAP2K7A | -1.25 | CDK2A | -1.03 | CAMK1DA | -0.98 | BMXA | -1.55 | STK17BA | -1.10 |
| TIF1A | -1.59 | MAPK9A | -1.26 | TIF1A | -1.17 | MAP2K7A | -1.04 | SLKA | -1.56 | CAMK4A | -1.13 |
| RPS6KA5A | -1.62 | GSG2A | -1.34 | PHKG2A | -1.23 | STK3A | -1.06 | PHKG2A | -1.57 | MAP2K7A | -1.14 |
| GPRK5A | -1.64 | CAMK4A | -1.42 | MAP2K7A | -1.23 | GSG2A | -1.07 | STK17BA | -1.59 | MAPK9A | -1.20 |
| STK10A | -1.67 | CDK2A | -1.54 | CAMK1DA | -1.26 | DAPK3A | -1.13 | PAK1A | -1.60 | PKMYT1A | -1.21 |
| NEK2A | -1.69 | TLK1A | -1.61 | MARK3A | -1.35 | DCAMKL1A | -1.17 | CDK2A | -1.79 | DCAMKL1A | -1.22 |
| DAPK3A | -1.79 | PKMYT1A | -1.64 | CAMK4A | -1.42 | NEK7A | -1.33 | STK38LA | -1.82 | CAMK1DA | -1.39 |
| TLK1A | -1.81 | SLKA | -1.69 | SLKA | -1.46 | TLK1A | -1.46 | MAPK9A | -1.97 | TLK1A | -1.42 |
| MAPK9A | -2.00 | EPHB3A | -1.75 | STK38LA | -1.48 | OSR1A | -1.51 | MAP2K7A | -2.00 | NEK7A | -1.43 |
| MAP2K7A | -2.04 | CDC42BPAA | -1.79 | TLK1A | -1.49 | RPS6KA5A | -1.68 | TLK1A | -2.02 | OSR1A | -1.53 |
| CDC42BPAA | -2.17 | NEK7A | -1.99 | NEK2A | -1.80 | MST4A | -1.71 | EPHB3A | -2.15 | MST4A | -1.55 |
| EPHB3A | -2.30 | STK38LA | -2.00 | STK10A | -1.85 | CAMK2BA | -1.88 | STK10A | -2.79 | EPHB3A | -2.12 |
| MARK4A | -2.73 | TIF1A | -2.31 | CDC42BPAA | -1.88 | STK10A | -2.05 | MST4A | -2.98 | NEK2A | -2.33 |
| MST4A | -3.13 | MST4A | -2.82 | MST4A | -1.94 | EPHB3A | -2.37 | MST3A | -3.19 | STK10A | -2.46 |
| MST3A | -3.25 | MST3A | -2.84 | EPHB3A | -2.14 | MST3A | -2.52 | CAMK1DA | -3.27 | MST3A | -2.48 |
| GSG2A | -3.53 | NEK2A | -3.55 | MST3A | -2.56 | NEK2A | -2.52 | GSG2A | -3.31 | RPS6KA5A | -2.66 |

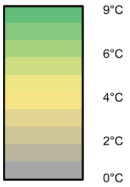

**Table S3:** Fragment study using DSF assay

| | **S1** | **S2** | **S3** | **S4** | **26** |
|---|---|---|---|---|---|
| | 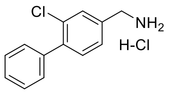 | 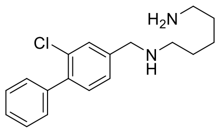 | 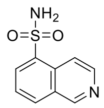 | 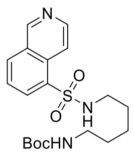 | 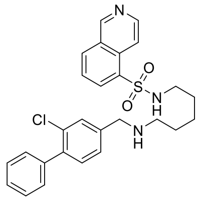 |
| CK2α (°C) | 0.64 | 0.5 | 0.42 | 0.28 | 6.66 |
| CK2α’ (°C) | 0.04 | 0 | 0 | 0 | 3.12 |

**Figure S1:** Overlay of **25** and **CX-4945**

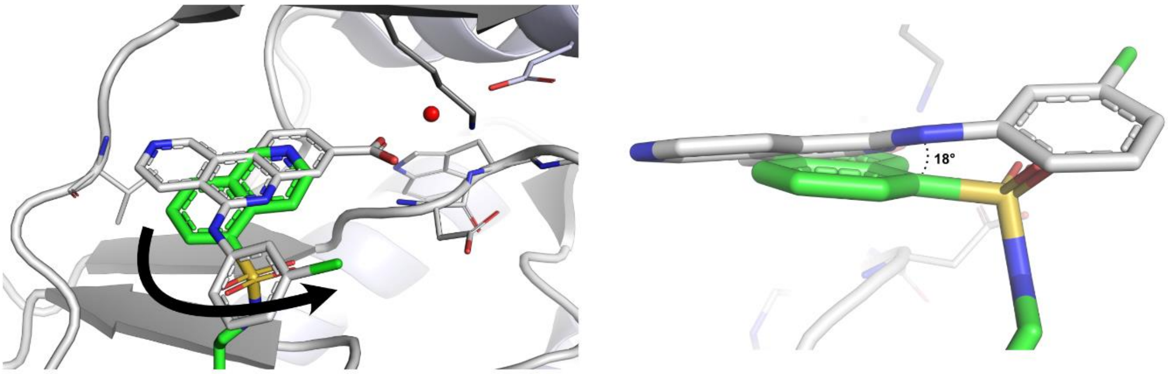

**Figure S1 :** Overlay of **25** (green, PDB 9EQ0) and **CX-4945** (white, PDB 3NGA). **25** bears a butyl linker which tilts the quinoline moiety 18° out of the plain of **CX-4945**.

**Experimental section**

**Differential Scanning Fluorimetry Assay:** Recombinant protein kinase domains with a concentration of 2 µM were mixed with a 10 µM compound solution in DMSO, using a final buffer consisting of 20 mM HEPES, pH 7.5, and 500 mM NaCl. SYPRO Orange (5000×, Invitrogen) was added as a fluorescence probe (1 µl per mL). Subsequently, temperature-dependent protein unfolding profiles were measured, using the QuantStudio™ 5 realtime PCR machine (Thermo Fisher). Excitation and emission filters were set to 465 nm and 590 nm. The temperature was raised with a step rate of 3°C per minute. Data points were analysed with the internal software (Thermal Shift SoftwareTM Version 1.4, Thermo Fisher) using the Boltzmann equation to determine the inflection point of the transition curve. Differences in melting temperature are given as $\Delta T_m$ values in °C. Measurements were performed in duplicates.

**Isothermal titration calorimetry (ITC):** The dissociation constants ($K_D$) of the compounds were assessed using isothermal calorimetry. Both the sample cell and injection syringe were pre-equilibrated with gel filtration buffer (25 mM HEPES at pH 7.5, 500 mM NaCl, and 0.5 mM TCEP). The sample cell was loaded with a solution of the compound (at concentrations ranging from 10-20 µM in gel filtration buffer), while the injection syringe contained the protein solution (ranging from 80-120 µM in gel filtration buffer). Following device equilibration (Nano ITC, TA Instruments), protein was incrementally injected over 30 injections, each comprising 8 µL (except a first injection of 4 µL). Analysis of the resulting data involved defining baseline and integration points to ascertain binding heats. Subsequently, data fitting to the Boltzmann equation enabled the determination of thermodynamic parameters.

**Kinome-Wide Selectivity Profile (Tracer displacement assay)**: Compounds **26** and **30** was tested at a concentration of 5 µM against a panel of 104 kinases in an *in vitro* tracer displacement assay as described previously (Figure 6).[1]

**Kinome-Wide Selectivity Profile (Tm panel)**: Recombinant protein kinase domains with a concentration of 2 µM were mixed with a 10 µM compound solution in DMSO, 20 mM HEPES, pH 7.5, and 500 mM NaCl. SYPRO Orange (5000×, Invitrogen) was added as a fluorescence probe (1 µl per mL). Subsequently, temperature-dependent protein unfolding profiles were measured, using the QuantStudio™ 5 realtime PCR machine (Thermo Fisher, Waltham, MA, USA). Excitation and emission filters were set to 465 and 590 nm. The temperature was raised with a step rate of 3 °C per minute. Data points were analyzed with the internal software (Thermal Shift SoftwareTM Version 1.4, Thermo Fisher) using the Boltzmann equation to determine the inflection point of the transition curve. Differences in melting temperature are given as ΔTm values in °C. Measurements were performed in duplicates.

**ADP glo assay:** The ADP Glo assay was carried out in concordance with the ADP-GloTM Kinase Assay Protocol (Promega). Recombinant protein kinase domain CK2 was seeded in a 384 well plate at a concentration of 200 nM with addition as 1 µM casein (Merck, C8654-500G) as substrate in Kinase Reaction Buffer (50 mM TRIS pH 7.5, 20 mM $MgCl_2$, 0.1% Glycerin, 0.01% Triton X100). Inhibitors of interest were titrated in wells between a range of 7.5 nM and 200 µM, respective negative and positive controls were also included. 20µM of Ultra-Pure ATP (Promega) was added to the wells to initiate the kinase reaction. All reactions were performed in a 5 µl volume at 37°C and incubated for 3 hours. After incubation, 5µl of ADP-GloTM Reagent (Promega) was added to each well and incubated at room temperature for 40 minutes to stop the kinase reaction and deplete unconsumed ATP. 10 µl of Kinase Detection Reagent (Promega) was added to each well and incubated at room temperature for 45 minutes to convert ADP to ATP and introduce luciferase and luciferin to detect ATP. Luminescence of each well was recorded using the PHERAstar plate reader (BMG Labtech, Ortenberg, Germany). Data were analysed using Graphpad Prism 9 software with a normalized response vs log(inhibitor) model with the following equation: Y=100/(1+10^((X-LogIC50))) to determine $IC_{50}$s of the tested compounds.

**NanoBRET$^{TM}$ assay**: The NanoBRET Assay was conducted following the methodology described in the literature.[2–4] Briefly, HEK293T cells were transfected with the CSNK2A1-

NanoLuc Fusion Vector (Promega, NV2981). This vector is used to express full-length CSNK2A1 kinase fused C-terminal to NanoLuc Luciferase. After trypsinization and resuspension in Opti-MEM without phenol red (Life Technologies), cells were seeded into white 384-well plates (Greiner 784075) at a density of 2 × 10^5 cells/mL. They were then transfected using FuGENE HD (Promega, E2312). Proteins were allowed to express for 20 h at 37 °C/5% CO2. Using an ECHO acoustic dispenser (Labcyte), the protein-transfected cells were exposed to increasing compound concentrations and NanoBRET K10 Tracer (Promega, TracerDB ID: T000008) at the Tracer KD concentration taken from TracerDB (tracerdb.org). To establish equilibrium, the system was incubated for 2 h at 37 °C/5% CO2. In accordance with the manufacturer's protocol, NanoBRET NanoGlo Substrate (Promega, N1573) was added, and filtered luminescence was measured using the PHERAstar plate reader (BMG Labtech) equipped with a luminescence filter pair (450 nm BP filter for the donor and 610 nm LP filter for the acceptor). Subsequently, competitive displacement data were analyzed and graphed using GraphPad Prism 10, employing a normalized 3-parameter curve fit with the following equation: $Y = 100/(1 + 10^{(X - \log IC_{50})})$. For the generation of the permeabilized data, the transfected cells were permeabilized using digitonin (0.05 µg/µL). After an equilibration period of 10 min, luminescence measurements were taken, and the displacement data were analyzed as described previously.

**Protein Expression and Purification:** Expression and purification were performed as described previously.[5–7] Briefly, transformed BL21(DE3) cells were grown in Terrific Broth medium containing 50 mg/mL kanamycin. Protein expression was induced at an $OD_{600}$ of 3 by using 0.5 mM isopropyl-thio-galactopyranoside (IPTG) at 18°C for 12 hours. Cells expressing $His_6$-tagged CK2α were lysed in lysis buffer containing 50 mM HEPES pH 7.5, 500 mM NaCl, 25 mM imidazole, 5% glycerol, and 0.5 mM Tris(2 carboxyethyl)phosphine (TCEP) by sonication. After centrifugation, the supernatant was loaded onto a Nickel-Sepharose column equilibrated with 30 mL lysis buffer. The column was washed with 60 mL lysis buffer. Proteins were eluted by an imidazole step gradient using 50, 100, 200, 300 mM imidazole concentrations. Fractions containing protein were pooled together and dialyzed overnight using 1L of final buffer (25 mM HEPES pH 7.5, 500 mM NaCl, 0.5 mM TCEP) at 4°C. Additionally, TEV protease was added (protein:TEV 1:20 molar ratio) to remove the tag. The following day the protein solution was loaded onto Nickel-Sepharose column beads again to remove the TEV protease and cleaved Tag. The combined flow through fraction and the wash fraction (25 mM imidazole) containing the protein were combined and concentrated to approximately 4-5 mL and loaded onto Superdex 75 16/60 Hi-Load gel filtration column equilibrated with final buffer. The protein was concentrated to approximately 9 mg/mL.

**Crystallization:** CK2α was crystallized using the sitting-drop vapour diffusion method by mixing protein (9 mg/mL) and well solutions in 2:1, 1:1, and 1:2 ratios. The reservoir solution contained 0.2 M ammonia sulphate, 0.1 M bis-tris pH 5.5 and between 23-26% (v/v) PEG 3350. The reservoir solution supplemented with 20% ethylene glycol was used as cryoprotectant for crystals. Crystals were stored in liquid nitrogen until measurement.

**Data collection, structure solution and refinement:** Diffraction data were collected either at beamline X06SA and X06DA (Villigen, CH) or Diamond Light Source I03 (Didcot, UK) at a wavelength of 1.0 Å at 100 K. Data were processed using XDS[8] and scaled with aimless[9]. The PDB structure with the accession code 6Z83[5] was used as an initial search MR model using the program MOLREP[10]. The dictionary files for the ligands were generated by the Grade Web Server (https://grade.globalphasing.org). The final model was built manually using Coot[11] and refined with REFMAC5[12], which is a part of the CCP4 suite[11]. Data collection and refinement statistics are summarized on pages S208-S388.

**Chemistry:** The synthetic routes of compounds will be outlined in the following and the analytical data can be found on pages S51-S207. All commercial chemicals were purchased from BLD Pharm, TCI, Merck, Enamine and Activate Scientific, with a purity ≥ 95% and were used without further purification. The solvents with an analytical grade were obtained from

VWR Chemicals, Honeywell and Merck and all dry solvents Acros Organics and Thermofischer. All reactions were carried out under an argon atmosphere. The thin layer chromatography was done with silica gel on aluminium foils (60 Å pore diameter) obtained from Macherey-Nagel and visualized with ultraviolet light (λ = 254 and 365 nm). The purification of the compounds was performed by flash chromatography using puriFlash XS 420 device with a UV-VIS multiwave detector (200−400 nm) from Interchim. Pre-packed normal-phase PF-SIHP and pre-packed PF-C18HP reverse phase columns with particle sizes of 15 and 30 µm (Interchim) were used for purification. DMSO was evaporated using a SpeedVac Concentrator (ThermoFischer Scientific, Model#: SPD121P-230) coupled to a Savant RVT5105 Refrigerated Vapor Trap (ThermoFischer Scientific) and a VLP80 vacuum pump (ThermoScientific). Nuclear magnetic resonance spectroscopy (NMR) was performed with DPX250, AV300, AV400, AV500, AV600 MHz spectrometers from Bruker. Chemical shifts (δ) are reported in parts per million (ppm). DMSO-$d_6$ and methylene chloride-$d_2$ were used as a solvent, and the spectra were calibrated to the solvent signal: 2.50 ppm ($^1$H NMR) or 39.52 ppm ($^{13}$C NMR) for DMSO-$d_6$, 7.26 ppm ($^1$H NMR) or 54.00 ppm ($^{13}$C NMR) for methylene chloride-$d_2$. Coupling constants (*J*) were reported in hertz (Hz) and multiplicities were designated as followed: s (singlet), d (doublet), dd (doublet of doublet), t (triplet), dt (doublet of triplets), td (triplet of doublets), ddd (doublet of doublet of doublet), q (quartet), m (multiplet). Mass spectra were measured on a Surveyor MSQ device from ThermoFisher measuring in the positive- or negative-ion mode. Final compounds were additionally characterized by HRMS using a MALDI LTQ Orbitrap XL from ThermoScientific and a microOTOF-Q ESI source. The purity of the final compounds was determined by HPLC using an Agilent 1260 Infinity II device with a 1260 DAD HS detector (G7117C; 254 nm, 280 nm, 310 nm) and a LC/MSD device (G6125B, ESI pos. 100-1000). The compounds were analyzed on a Poroshell 120 EC-C18 (Agilent, 3 x 150 mm, 2.7 µm) reversed phase column using 0.1% formic acid in water (A) and 0.1% formic acid in acetonitrile (B) as a mobile phase. The following gradient was used: 0 min 5% B - 2 min 5% B - 8 min 98% B - 10 min 98% B (flow rate of 0.5

mL/min). UV-detection was performed at 254, 280 and 310 nm and all compounds used for further biological characterization showed a purity ≥95%.

**Synthesis of 2-chloro-4-formylphenyltrifluoromethanesulfonate 17**

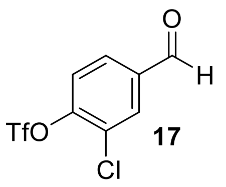

**17**

3-chloro-4-hydroxybenzaldehyde **16** (12.0 g, 76.64 mmol, 1 eq.) and DMAP (911 mg, 6.86 mmol, 0,09 eq.) were charged into a round bottom flask. 250 mL of dry DCM were added through a septum. The mixture was cooled to 0°C in an ice bath and dry pyridine (18.5 mL, 230 mmol, 3 eq.) were added. The flask was cooled in an ice bath and 1M trifluoroacetic anhydride (TFAA) solution in DCM (154 mL, 154 mmol, 2 eq.) were added dropwise through a dropping funnel. The solution was warmed to ambient temperature and stirred over 16 hours under argon atmosphere. The solvent was evaporated in vacuo. The residue was solved in EtOAc, 1 M aq. HCl were added to the flask and stirred vigorously for 5 min. After transfer into a funnel the organic phase was washed with 1x 1M aq. HCl, 3x sat. $NaHCO_3$-solution and 2x with brine. The EtOAc-phase was dried over $MgSO_4$, filtrated and evaporated in vacuo. The crude product was purified by silica column chromatography (Hexanes/EtOAc 4:1). The product was dried overnight using a high vacuum pump to obtain a pale yellow, oily substance that can be stored for up to one year in - 20°C (19,0 g, 86%). $^1$H NMR (250 MHz, DMSO-$d_6$) δ 10.03 (s, 1H), 8.29 (d, 1H), 8.06 (dd, 1H), 7.91 (d, 12H). $^{13}$C NMR (126MHz, DMSO-$d_6$) δ 190.93, 148.21, 137.07, 132.16, 130.10, 126.97, 124.53, 116.78. MS-ESI m/z $[M+H]^+$: calcd 288.2, found 288.91.

*Note: After addition of pyridine the mixture becomes clear.*

**Synthesis of 2-chloro-[1,1'-biphenyl]-4-carbaldehyde 18**

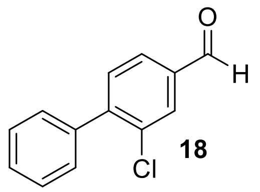

**18**

2-chloro-4-formylphenyltrifluormethansulfonate **17** (500 mg, 1.73 mmol, 1 eq,), phenylboronic acid (316 mg, 2.6 mmol, 1.5 eq.), lithium chloride (132 mg, 3.12 mmol, 1.8 eq.), potassium phosphate (1.1 g, 5.2 mmol, 3 eq.) and tetrakis(triphenylphosphine)palladium (18 mg, 0.016 mmol, 0.1 eq.) were charged in a microwave vial and were suspended in DME (10 mL). The suspension was degassed with argon under sonication. The vial was heated in a microwave oven to 80 °C for 3 h. The crude mixture was then filtrated over celite and diluted with EtOAc. The organic phase was washed with $NaHCO_3$-solution (3x50 mL), Brine (2x50 mL), dried over $MgSO_4$ and the solvent was removed under reduced pressure. The residue was purified by flash chromatography (100% Heptane) to afford the title compound was afford as white solid (255 mg, 68%).$^{1}$H NMR (300 MHz, DMSO-$d_6$) δ = 10.03 (s, 1H, H-8), 8.05 (d, J = 1.6 Hz, 1H, H-4), 7.92 (dd, J = 7.8, 1.6 Hz, 1H, H-2), 7.61 (d, J = 7.8 Hz, 1H, H, H-5), 7.53 – 7.40 (m, 5H, H-10, 11, 12, 13, 14). $^{13}$C NMR (126 MHz, DMSO-$d_6$) δ 191.87, 145.29, 137.73, 136.63, 132.40, 132.29, 130.81, 129.07, 128.50, 128.40, 127.93. MS-ESI m/z $[M+H]^+$: calcd 216.66, found 217.05.

*Note: Older batches of tetrakis(triphenylphosphine)palladium (orange colour) performed with better overall conversion and yields than fresh ones.*

*The product does not always readily crystallize- it may take at least overnight.*

**General procedure A for reductive amination**

**18** (1 eq.) and $MgSO_4$ were charged in a flask and suspended in dry DCM. After addition of the corresponding amino linker (1.3 eq.) the mixture was stirred over 1 h under argon atmosphere at room temperature. Sodium triacetoxyborhydride (STAB) (2.5 eq.) was added and the suspension was stirred overnight. Solvents were evaporated, diluted with EtOAc and washed 2x with sat. $NaHCO_3$, 2x with water and 1x with brine. The organic phase was dried over $MgSO_4$ and purified using flash chromatography using DCM/MeOH as an eluent.

**General procedure B for *N*-Boc deprotection**

*N*-Boc protected amines were solved in dry DCM. The flask was cooled in an ice bath and trifluoroacetic acid (TFA) in an excess was added dropwise. The solution stirred for 2h at room temperature. Solvents were evaporated until complete dryness and used without further purification in the next step.

**General procedure C for sulfonamide coupling**

Deprotected amines were solved in dry THF (≈ 0.01 M) and triethylamine (3 eq.) was added to the solution. The corresponding sulfonyl chloride was added and stirred under argon atmosphere overnight. Solvents were evaporated, diluted with EtOAc and washed 2x with water and 1x with brine. The organic phase was dried over $MgSO_4$ and purified using flash chromatography using DCM/MeOH as an eluent.

*Note: In most cases, after ≈ 1h, the solution became slightly blurry indicating the formation of product as it is scarcely soluble in THF. This in addition to the high dilution rendered the reaction chemoselective for the terminal amine.*

**General procedure D for the synthesis of sulfinate salts [13]**

Brominated fused heterocycles (1 eq.), DABSO (DABCO-Bis(sulfur dioxide)) (1 eq.), dicyclohexylmethylamine (3 eq.), Pd(amphos)$_2$Cl$_2$ (0.05 eq.) were charged into a microwave vial and suspended in dry isopropanol. The reaction was heated at 110°C for 1h. The *in situ* formed sulfinate salt was treated with the corresponding electrophile.

**Synthesis of tert-butyl (5-(((2-chloro-[1,1'-biphenyl]-4yl)methyl)amino)pentyl)carbamate 19**

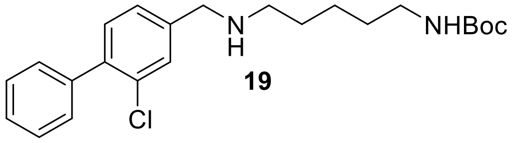

**19**

**19** was synthesized according to **General Procedure A** using **18** (154 mg, 0.7 mmol, 1 eq.) and *tert*-butyl (5-aminopentyl)carbamate (210 mg, 1.04 mmol, 1.5 eq.) in dry DCM (3 mL). (200 mg, 72%) $^{1}$H NMR (500 MHz, DMSO-d$_6$) δ 7.52 – 7.31 (m, 8H), 6.75 (t, J = 5.8 Hz, 1H), 3.71 (s, 2H), 2.89 (q, J = 6.6 Hz, 2H), 2.47 (t, J = 7.0 Hz, 2H), 1.49 – 1.38 (m, 3H), 1.36 (s, 9H), 1.35 – 1.22 (m, 3H). $^{13}$C NMR (126 MHz, DMSO-d$_6$) δ 155.57, 142.87, 138.73, 137.77, 131.12, 130.95, 129.23, 128.90, 128.18, 127.57, 126.92, 77.27, 52.00, 48.62, 29.44, 29.21, 28.27, 24.13. MS-ESI m/z [M+H]+: calcd 403.20, found 403.19.

**Synthesis of tert-butyl(2-(2-(((2-chloro-[1,1'-biphenyl]-4yl)methyl)amino)etoxy)ethyl)carbamate 20**

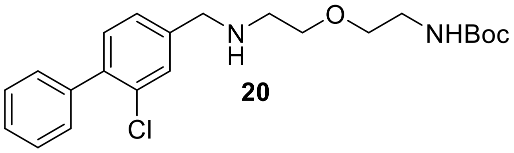

**20**

**20** was synthesized according to **General Procedure A** using **18** (250 mg, 1.15 mmol, 1 eq.) and *tert*-butyl (2-(2-aminoethoxy)ethyl)carbamate (236 mg, 1.5 mmol, 1 eq.) in dry DCM (10 mL). (225 mg, 48%) $^{1}$H NMR (400 MHz, DMSO-d$_6$) δ 7.52 (d, J = 1.5 Hz, 1H), 7.49 – 7.32 (m, 7H), 6.85 – 6.80 (m, 1H), 3.75 (s, 2H), 3.47 (t, J = 5.6 Hz, 2H), 3.37 (t, J = 5.9 Hz, 2H), 3.08 (q, J = 5.9 Hz, 2H), 2.64 (t, J = 5.6 Hz, 2H), 1.36 (s, 9H). $^{13}$C NMR (101 MHz, DMSO-d$_6$) δ 156.11, 143.10, 139.20, 138.36, 131.64, 131.47, 129.70, 129.44, 128.67, 128.07, 127.45, 78.03, 70.31, 69.63, 52.32, 48.50, 40.64, 28.70. MS-ESI m/z [M+H]+: calcd 405.94, found 405.21.

**Synthesis of tert-butyl (6-(((2-chloro-[1,1'-biphenyl]-4-yl)methyl)amino)hexyl)carbamate 21**

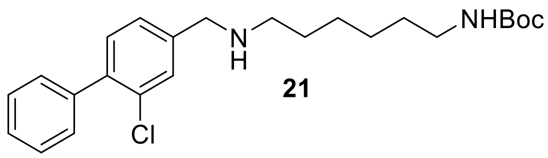

21

**21** was synthesized according to **General Procedure A** using **18** (200 mg, 0.92 mmol, 1 eq.) and *tert*-butyl (5-aminohexyl)carbamate (302 mg, 1.11 mmol, 1 eq.) in dry DCM (10 mL). (234 mg, 61%) $^1$H NMR (250 MHz, DMSO-$d_6$) δ 7.54 (d, J = 1.4 Hz, 1H), 7.48 – 7.39 (m, 5H), 7.37 – 7.31 (m, 2H), 6.74 (t, J = 5.3 Hz, 1H), 3.76 (s, 2H), 2.89 (q, J = 6.5 Hz, 2H), 1.43 (q, J = 7.3 Hz, 3H), 1.36 (d, J = 0.9 Hz, 8H), 1.35 – 1.14 (m, 7H). MS-ESI m/z [M+H]+: calcd 417.99, found 417.20.

**Synthesis of tert-butyl 4-(2-(((2-chloro-[1,1'-biphenyl]-4-yl)methyl)amino)ethyl)piperidine-1-carboxylate 22**

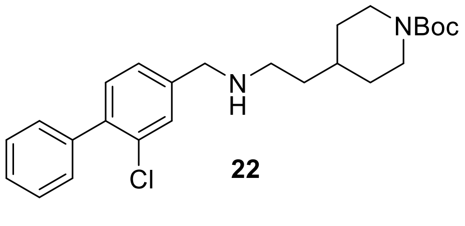

22

**22** was synthesized according to **General Procedure A** using **18** (430 mg, 1.98 mmol, 1 eq.) and 4-(aminoethyl)-1-N-boc-piperidine (570 mg, 2.5 mmol, 1.3 eq.) in dry DCM (13 mL) (406 mg, 57%). $^1$H NMR (500 MHz, Methylene Chloride-$d_2$) δ 7.47 (dt, J = 1.4, 0.7 Hz, 1H), 7.46 – 7.42 (m, 4H), 7.41 – 7.35 (m, 1H), 7.30 (dd, J = 1.5, 1.0 Hz, 2H), 4.04 (d, J = 12.2 Hz, 2H), 3.83 – 3.75 (m, 2H), 2.66 (d, J = 7.2 Hz, 2H), 1.65 (s, 2H), 1.53 (dddd, J = 12.5, 11.2, 4.7, 2.3 Hz, 3H), 1.46 (s, 2H), 1.43 (s, 10H), 1.16 – 1.01 (m, 2H). $^{13}$C NMR (126MHz, Methylene Chloride-$d_2$) δ 155.21, 142.62, 139.92, 139.36, 132.62, 131.79, 130.03, 129.87, 128.60, 128.08, 127.20, 79.35, 47.18, 37.43, 34.52, 30.62, 28.73. MS-ESI m/z [M+H]+: calcd 429.00, found 429.58.

**Synthesis of tert-butyl 2-(((2-chloro-[1,1'-biphenyl]-4-yl)methyl)amino)-7-azaspiro[3.5]-nonane-7-carboxylate 23**

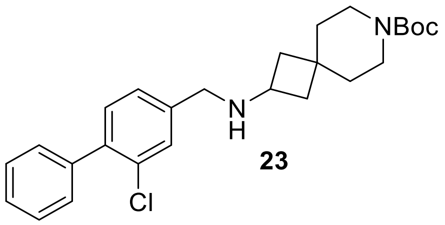

**23**

**23** was synthesized according to **General Procedure A** using **18** (150 mg, 0.69 mmol, 1 eq.) and *tert*-butyl 2-amino-7-azaspiro[3.5]nonane-7-carboxylate (200 mg, 0.83 mmol, 1.2 eq.) in dry DCM (10 mL). (210 mg, 69%) $^1$H NMR (400 MHz, DMSO-$d_6$) δ 7.54 (d, J = 1.5 Hz, 1H), 7.47 – 7.33 (m, 7H), 3.69 (s, 2H), 3.29 (s, 1H), 3.24 (dd, J = 7.0, 3.5 Hz, 2H), 3.19 (t, J = 5.7 Hz, 2H), 2.06 (ddd, J = 10.1, 7.7, 2.6 Hz, 2H), 1.59 – 1.52 (m, 2H), 1.46 – 1.41 (m, 4H), 1.38 (s, 9H). $^{13}$C NMR (101 MHz, DMSO-$d_6$) δ 153.96, 138.61, 138.16, 131.18, 130.96, 129.27, 129.20, 128.20, 127.64, 127.29, 78.40, 48.86, 47.32, 38.76, 31.49, 28.08. MS-ESI m/z [M+H]+: calcd 442.01, found 441.06.

**Synthesis of *tert*-butyl 2-(((2-chloro-[1,1'-biphenyl]-4-yl)methyl)amino)-6-azaspiro[3.4]-octane-6-carboxylate 24**

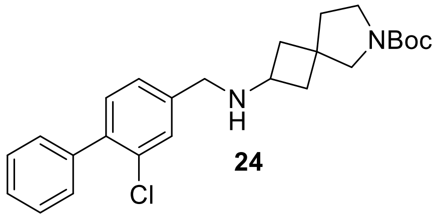

**24**

**24** was synthesized according to **General Procedure A** using **18** (230 mg, 1.06 mmol, 1 eq.) and *tert*-butyl 2-amino-6-azaspiro[3.4]octane-6-carboxylate (264 mg, 1.17 mmol, 1.1 eq.) in dry DCM (10 mL). (170 mg, %) $^1$H NMR (400 MHz, DMSO-$d_6$) δ 7.53 – 7.32 (m, 8H), 3.65 (s, 2H),

3.21 – 3.12 (m, 5H), 2.10 (tdd, J = 9.9, 7.4, 2.7 Hz, 2H), 1.74 (dt, J = 19.0, 9.8 Hz, 4H), 1.38 (s, 9H). $^{13}$C NMR (101 MHz, DMSO-$d_6$) δ 153.68, 142.47, 138.69, 137.89, 131.12, 130.93, 129.22, 129.05, 129.02, 128.19, 127.59, 127.07, 127.05, 78.06, 57.81, 57.40, 49.16, 47.92, 47.65, 47.58, 44.59, 44.36, 44.31, 44.10, 38.97, 28.18. MS-ESI m/z [M+H]+: calcd 427.99, found 427.15.

**Synthesis of 3-methyl-3*H*-imidazo[4,5-*b*]pyridine-6-sulfonyl fluoride 60**

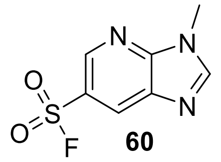

**60**

**60** was synthesized according to **General Procedure D** using 6-bromo-3-methyl-3*H*-imidazo[4,5-*b*]pyridine **59** (83 mg, 0.4 mmol, 1 eq.), DABSO (96 mg, 0.4 mmol, 1 eq.), dicyclohexylmethylamine (270 µL, 1.2 mmol, 3 eq.), Pd(amphos)$_2$Cl$_2$ (14.2 mg , 0.02 mmol, 0.05 eq.). After heating in the microwave, the reaction was cooled to 0°C using an ice bath and NFSI (*N*-Fluorosuccinimide, 189 mg, 0.6 mmol, 1.5 eq.) was added to the mixture and stirred 3h at ambient temperature. The solvent was evaporated, the crude was washed 3x with water, 1x with brine and dried over $Na_2SO_4$. The organic phase was evaporated and purified using flash chromatography (Heptane/EtOAc 2:1 as eluent) yielding the title compound as a white solid (61 mg, 71 %). $^1$H NMR (400 MHz, DMSO-$d_6$) δ 9.07 (dd, J = 2.2, 1.1 Hz, 1H), 8.92 – 8.87 (d, 1H), 8.82 (s, 1H), 3.93 (s, 3H). $^{13}$C NMR (101 MHz, DMSO-$d_6$) δ 151.34, 150.62, 143.42, 133.95, 128.14, 122.37, 122.13, 30.07. $^{19}$F NMR (377 MHz, DMSO-$d_6$) δ -73.41. MS-ESI m/z [M+H]+: calcd 216.20, found 216.00.

*Note: Addition of the NFSI resulted in an exothermic reaction (microwave vial became warm) transitioning from a grey suspension to a grey-purple colour after ~15 min.*

**Synthesis of *tert*-butyl (6-((3-methyl-3*H*-imidazo[4,5-*b*]pyridin-6-yl)sulfonyl)hexyl)carbamate 62**

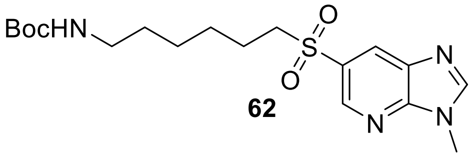

**62**

**62** was synthesized according to **General Procedure D** using 6-bromo-3-methyl-3*H*-imidazo[4,5-*b*]pyridine (200 mg, 0.94 mmol, 1 eq.), DABSO (227 mg, 0.94 mmol, 1 eq.), dicyclohexylmethylamine (912 µL, 2.83 mmol, 3 eq.), Pd(amphos)$_2$Cl$_2$ (34 mg , 0.05 mmol, 0.05 eq.). After heating in the microwave *tert*-butyl (6-bromohexyl)carbamate (502 mg, 1.9 mmol, 2 eq.) solved in 1 mL of dry DMF was added to the mixture and heated in the microwave for 1h at 120°C. The solvent was evaporated, the crude was washed 3x with water, 1x with brine and dried over $Na_2SO_4$. The organic phase was evaporated and purified using flash chromatography (Heptane/EtOAc 2:1 as eluent) yielding the title compound as a white solid (75 mg, 21 %). $^1$H NMR (400 MHz, DMSO-d$_6$) δ 8.83 (t, J = 1.9 Hz, 1H), 8.70 – 8.66 (m, 1H), 8.53 (t, J = 1.8 Hz, 1H), 6.70 (t, J = 5.5 Hz, 1H), 3.91 (d, J = 1.6 Hz, 3H), 3.46 – 3.28 (m, 4H), 2.84 (q, J = 6.5 Hz, 2H), 1.53 (h, J = 6.8, 6.0 Hz, 2H), 1.35 – 1.32 (m, 9H), 1.27 (d, J = 7.2 Hz, 2H), 1.17 (td, J = 8.7, 4.2 Hz, 2H). $^{13}$C NMR (101 MHz, DMSO-d$_6$) δ 155.56, 149.95, 149.27, 143.14, 133.88, 129.65, 127.26, 77.28, 55.33, 55.31, 39.60, 29.86, 29.07, 28.25, 28.21, 27.08, 25.63, 22.35. MS-ESI m/z [M+H]+: calcd 397.50, found 297.10 (*N*-Boc group fragmentation) .

**Synthesis of *tert*-butyl (4-(imidazo[1,2-*a*]pyridin-6-ylsulfonyl)butyl)carbamate 65**

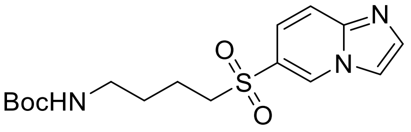

**65**

**65** was synthesized according to **General Procedure D** using 6-bromoimidazo[1,2-a]pyridine (79 mg, 0.4 mmol, 1 eq.), DABSO (96 mg, 0.4 mmol, 1 eq.), dicyclohexylmethylamine (270µL, 1.2 mmol, 3 eq.), Pd(amphos)$_2$Cl$_2$ (14.2 mg , 0.02 mmol, 0.05 eq.). After heating in the microwave *tert*-butyl (4-bromobutyl)carbamate (202 mg, 0.8 mmol, 2 eq.) solved in 500 µL of dry DMF was added to the mixture and heated in the microwave for 1h at 120°C. The solvent was evaporated, the crude was washed 3x with water, 1x with brine and dried over $Na_2SO_4$. The organic phase was evaporated and purified using flash chromatography (Heptane/EtOAc 2:1 as eluent) yielding the title compound as a white solid (20 mg, 14%). $^1$H NMR (400 MHz, DMSO-$d_6$) δ 9.26 (dd, J = 2.0, 1.0 Hz, 1H), 8.19 (s, 1H), 7.76 (dd, J = 5.4, 4.2 Hz, 2H), 7.53 (dd, J = 9.5, 1.9 Hz, 1H), 6.79 (t, J = 5.9 Hz, 1H), 3.44 – 3.37 (m, 2H), 2.88 (q, J = 6.3 Hz, 2H), 1.62 – 1.52 (m, 2H), 1.44 (p, J = 6.8 Hz, 2H), 1.30 (s, 9H). $^{13}$C NMR (101 MHz, DMSO-$d_6$) δ 155.62, 144.31, 135.34, 130.66, 124.74, 121.32, 117.38, 115.43, 77.42, 54.46, 38.93, 28.17, 27.98, 19.78.MS-ESI m/z [M+H]+: calcd 354.44, found 354.15.

**Synthesis of *tert*-butyl (4-(isoquinoline-5-sulfonamido)butyl)carbamate 68**

**68**

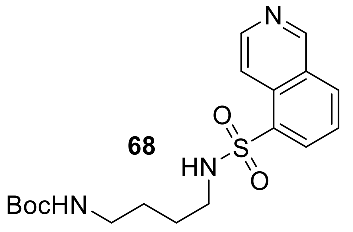

5-Isoquinolinsulfonylchloride (44 mg, 0.19 mmol, 1 eq) and *N*-Boc-1,4-butanediamine (54 mg, 0.29 mmol, 1.5 eq) were dissolved in dry DCM (1 mL). Triethylamine (133 µL, 0.96 mmol, 5 eq.) was added to the solution and stirred for 3 h at room temperature under argon atmosphere. The reaction mixture was concentrated and the residue diluted in DCM. The organic phase was washed with water (2x100 mL) and brine (2x100 mL). The organic phase was dried over $MgSO_4$ and the solvent was removed under reduced pressure. The residue was purified by flash chromatography (DCM/MeOH. 30:1) affording the title compound as yellow oil (42 mg, 1.49 mmol, 58 %). $^1$H NMR (500 MHz, Methylene Chloride-$d_2$) δ 9.37 – 9.33 (m, 1H), 8.65 (dd, J = 6.2, 1.5 Hz, 1H), 8.44 – 8.38 (m, 2H), 8.25 – 8.20 (m, 1H), 7.71 (ddd, J = 8.7, 7.4, 1.5 Hz, 1H), 5.67 (s, 1H), 4.62 (s, 1H), 2.95 (dt, J = 28.3, 6.8 Hz, 4H), 1.41 (s, 2H), 1.40 (s, 9H), 1.37 (s, 3H), 1.29 – 1.21 (m, 1H). $^{13}$C NMR (126 MHz, Methylene Chloride-$d_2$) δ 153.92, 145.64, 134.03,

133.77, 126.51, 117.69, 79.56, 54.43, 54.22, 54.00, 53.78, 53.57, 43.46, 40.17, 28.65, 27.90, 26.87. MS-ESI m/z [M+Na]+: calcd 402.48, found 402.08

**Synthesis of *tert*-butyl (5-(benzo[*d*]thiazole-6-sulfonamido)pentyl)carbamate 70**

**70**

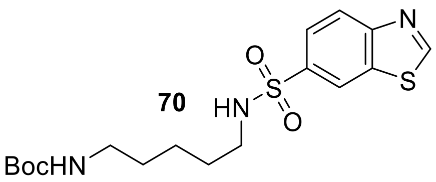

Benzo[*d*]thiazole-6-sulfonyl chloride **69** (500 mg, 2.14 mmol, 1 eq) and *N*-Boc-1,5-pentanediamine (476 mg, 2.35 mmol, 1.1 eq) were dissolved in dry THF (40 mL). Triethylamine (895 µL, 6.42 mmol, 3 eq.) was added to the solution and stirred for 3 h at room temperature under argon atmosphere. The reaction mixture was concentrated and the residue diluted in DCM. The organic phase was washed with water (2x100 mL) and brine (2x100 mL). The organic phase was dried over $MgSO_4$ and the solvent was removed under reduced pressure. The residue was purified by flash chromatography (DCM/MeOH. 30:1) affording the title compound as yellow oil (597 mg, 1.49 mmol, 70 %). $^1$H-NMR (400 MHz, DMSO-d$_6$) δ = 9.61 (s, 1H, H-8), 8.68 (d, *J* = 1.8 Hz, 1H, H-3), 8.27 (d, *J* = 8.6 Hz, 1H, H-1), 7.91 (dd, *J* = 8.6, 1.9 Hz, 1H, H-6), 7.67 (t, *J* = 5.7 Hz, 1H,N H-23), 6.71 (t, *J* = 5.7 Hz, 1H, NH-17), 2.78 (dq, *J* = 29.1, 6.6 Hz, 4H, H-22,18), 1.35 (s, 12H, H-13,21,24,26), 1.30 – 1.12 (m, 4H, H-19,20), 2.29 (s, 1H). $^{13}$C NMR (126 MHz, DMSO-d$_6$) δ = 160.48 (C-18), 155.53 (C-16), 154.94 (C-5), 137.45 (C-2), 134.09 (C-4), 128.90 (C-1), 128.20, 124.29, 124.17, 123.67 (C-6), 122.12 (C-3), 77.31 (C-14), 42.55 (C-22), 28.92 (C-12), 28.66 (C-19), 28.26 (C-13,25,26), 23.31 (C-20). MS-ESI m/z [M+H]+: calcd 399.54, found 400.05.

**Synthesis of Methyl *N*-(benzo[*d*]thiazol-6-ylsulfonyl)-*N*-(5-((*tert*-butoxycarbonyl)amino)-pentyl)glycinate 71**

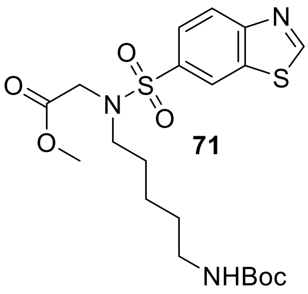

**71**

**70** (461 mg, 1.15 mmol, 1 eq.) was dissolved in dry ACN (15 mL) and cooled to 0 °C. Methyl bromoacetate (397 mg, 2.6 mmol, 2.25 eq.) and potassium carbonate (175 mg, 1.27 mmol, 1.1 eq.) were added to the reaction mixture. The reaction mixture was allowed to warm to room temperature and then heated to 60 °C for 16 h. TLC control showed not complete conversion of starting material and additional methylbromacetate (176 mg, 1.15 mmol, 1.2 eq.) was added to the solution and stirred for 3h. The suspension was diluted in EtOAc, washed with water (2x 100 mL) and brine (2x100 mL). The organic phase was dried over $MgSO_4$ and the solvent was removed under reduced pressure. The title compound crystalized as a yellow solid (542 mg, quant.). $^1$H-NMR (400 MHz, DMSO-d$_6$) δ = 9.63 (s, 1H, H-8), 8.77 (d, J = 1.9 Hz, 1H, H-3), 8.25 (d, J = 8.6 Hz, 1H, H-1), 7.93 (dd, J = 8.6, 1.9 Hz, 1H, H-6), 6.73 (s, 1H, NH-17), 4.11 (s, 2H, H-27), 3.54 (s, 3H, H-31), 3.17 (t, J = 7.4 Hz, 2H, H-22), 2.83 (q, J = 6.6 Hz, 2H, H-18), 1.43 (p, J = 7.5 Hz, 2H, H-21), 1.36 (s, 9H, H-13,25,26), 1.34 – 1.23 (m, 2H, H-19), 1.22 – 1.11 (m, 2H, H-20). $^{13}$C NMR (101 MHz, DMSO-d$_6$) δ = 169.43 (C-28), 160.89 (C-8), 155.56 (C-16), 155.23 (C-5), 136.14 (C-2), 134.25 (C-4), 124.62 (C-1), 124.57 (C-3), 123.56 C-6), 122.73, 77.32 (C-14), 51.88 (C-31), 51.83 (C-27), 48.5 (C-22)1, 48.17 (C-18), 28.98 (C-29), 28.26 (C-13,25,26), 27.11, 23.14 (C-20). MS-ESI m/z [M+Na]+: calcd 494.95, found 494.15.

**Synthesis of *N*-(4-(((2-chloro-[1,1'-biphenyl]-4-yl)methyl)amino)butyl)isoquinoline-5-sulfonamide 25**

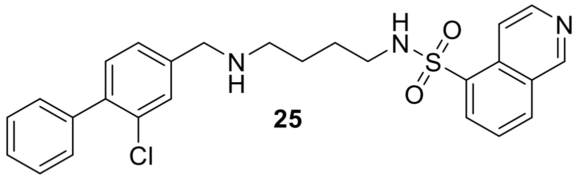

**25**

**68** (51 mg, 0.13 mmol, 1eq.) was deprotected using **General Procedure B**. The crude product was reacted with **18** (20 mg, 0.09 mmol, 1 eq.) according to **General Procedure A** (20 mg, 46%). $^{1}$H NMR (500 MHz, Methylene Chloride-$d_2$) δ 10.14 (d, J = 0.9 Hz, 1H), 8.67 (d, J = 5.7 Hz, 1H), 8.27 (dd, J = 7.3, 1.2 Hz, 1H), 8.05 (dt, J = 8.4, 1.1 Hz, 1H), 7.80 (dd, J = 5.7, 1.0 Hz, 1H), 7.76 (dd, J = 8.3, 7.3 Hz, 1H), 7.48 – 7.26 (m, 8H), 3.79 (s, 2H), 2.94 (t, J = 6.1 Hz, 2H), 2.54 (t, J = 6.2 Hz, 2H), 1.54 – 1.42 (m, 4H). $^{13}$C NMR (126 MHz, Methylene Chloride-$d_2$) δ 149.86, 144.18, 132.74, 131.97, 130.86, 130.22, 130.00, 129.50, 128.59, 128.12, 127.57, 121.72, 54.43, 54.22, 54.00, 53.78, 53.57, 53.12, 48.87, 43.65, 28.54, 27.94. MS-ESI m/z [M + H]+: calcd 480.23, found 480.02. HRMS m/z [M + H]+: calcd 479.14343, found 480.14920.

**Synthesis of *N*-(5-(((2-chloro-[1,1'-biphenyl]-4-yl)methyl)amino)pentyl)isoquinoline-5-sulfonamide 26**

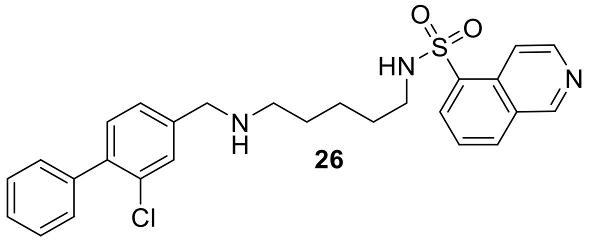

**26**

**19** (38 mg, 0.09 mmol, 1eq.) was deprotected using **General Procedure B**. The crude product was reacted with 5-Isoquinolinsulfonylchloride (22 mg, 0.09 mmol, 1 eq.) according to **General Procedure A** (38 mg, 86%). $^{1}$H NMR (500 MHz, Methylene Chloride-$d_2$) δ 9.35 (s, 1H), 8.66 (d, J = 6.0 Hz, 1H), 8.45 – 8.35 (m, 2H), 8.22 (d, J = 8.2 Hz, 1H), 7.71 (t, J = 7.8 Hz, 1H), 7.47 – 7.40 (m, 5H), 7.38 (ddd, J = 8.9, 5.6, 3.3 Hz, 1H), 7.31 – 7.25 (m, 2H), 3.76 (s, 2H), 2.92 (t, J = 6.9 Hz, 2H), 2.51 (t, J = 7.0 Hz, 2H), 1.46 – 1.31 (m, 6H). $^{13}$C NMR (126 MHz, Methylene Chloride-

$d_2$) δ 153.96, 145.69, 134.04, 133.78, 131.86, 130.04, 129.99, 128.60, 128.12, 127.37, 126.51, 117.63, 54.43, 54.22, 54.00, 49.13, 43.68, 30.25, 29.50, 24.49. MS-ESI m/z [M + H]+: calcd 494.16, found 494.24. HRMS m/z [M + H]+: calcd 494.15908, found 494.16558.

**Synthesis of *N*-(5-(((2-chloro-[1,1'-biphenyl]-4-yl)methyl)amino)pentyl)isoquinoline-5-sulfonamide 27**

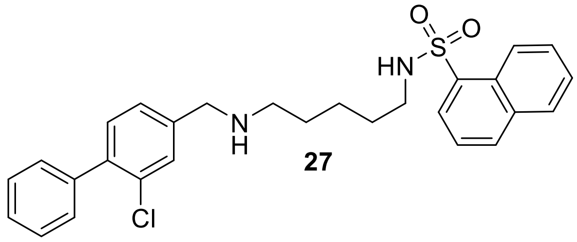

**27**

**19** (518 mg, 1.28 mmol, 1 eq.) was deprotected using **General Procedure B**. The crude product was reacted with naphthalene-1-sulfonylchloride (233 mg, 1.03 mmol, 0.8 eq.) according to **General Procedure A** (138 mg, 27%). $^1$H NMR (600 MHz, DMSO-$d_6$) δ 8.67 (dd, J = 8.5, 1.1 Hz, 1H), 8.22 – 8.18 (m, 1H), 8.12 (dd, J = 7.3, 1.3 Hz, 1H), 8.09 – 8.05 (m, 1H), 7.90 (s, 1H), 7.70 (ddd, J = 8.5, 6.8, 1.5 Hz, 1H), 7.67 – 7.60 (m, 2H), 7.48 – 7.44 (m, 3H), 7.43 – 7.38 (m, 3H), 7.35 – 7.30 (m, 2H), 3.65 (s, 2H), 2.78 (t, J = 7.0 Hz, 2H), 2.29 (t, J = 7.1 Hz, 2H), 1.24 (dq, J = 25.3, 7.4 Hz, 4H), 1.15 – 1.07 (m, 2H). $^{13}$C NMR (151 MHz, DMSO-$d_6$) δ 142.47, 138.67, 137.81, 135.85, 133.82, 133.50, 131.05, 130.91, 129.14, 128.88, 128.81, 128.28, 128.11, 127.64, 127.57, 127.51, 126.87, 126.70, 124.71, 124.42, 51.80, 48.26, 42.28, 40.06, 28.87, 28.69, 23.62. MS-ESI m/z [M + H]+: calcd 493.16, found 493.13. HRMS m/z [M + H]+: calcd 493.1638 found 493.1693.

**Synthesis of *N*-(5-(((2-chloro-[1,1'-biphenyl]-4-yl)methyl)amino)pentyl)benzo[*d*]thiazole-6-sulfonamide TFA salt 72**

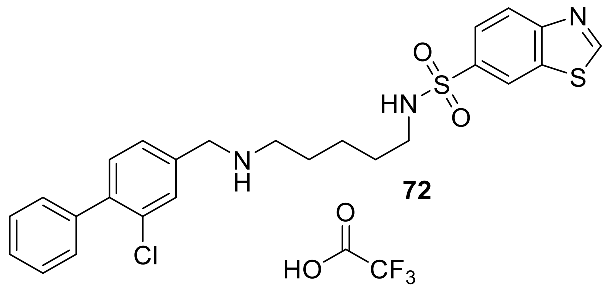

**72**

**19** (50 mg, 0.12 mmol, 1 eq.) was deprotected using **General Procedure B**. The crude product was reacted with benzo[d]thiazole-6-sulfonyl chloride (25 mg, 0.11 mmol, 1 eq.) according to **General Procedure A** (48 mg, 87%). $^1$H-NMR (400 MHz, DMSO-d$_6$) δ = 9.61 (s, 1H, H-29), 8.69 (d, J = 1.9 Hz, 1H, H-24), 8.27 (d, J = 8.6 Hz, 1H, H-22), 7.92 (dd, J = 8.6, 1.9 Hz, 1H, H-27), 7.69 (bs, 1H, NH-15), 7.51 – 7.35 (m, 9H, H-3,4,7,9,10,11,12,13, NH-21), 7.32 (s, 2H, H-1), 2.77 (t, J = 6.9 Hz, 2H, H-16), 2.39 (t, J = 6.9 Hz, 2H, H-20), 2.07 (s, 2H), 1.44 – 1.15 (m, 6H, H-17,18,19). $^{13}$C NMR (101 MHz, DMSO-d$_6$) δ = 160.47, 154.94, 142.67, 138.71, 137.80, 137.54, 134.09, 131.12, 130.95, 129.22, 128.90, 128.17, 127.57, 126.92, 124.18, 123.67, 122.11, 51.94, 48.43, 42.60, 28.93, 23.85. MS-ESI m/z [M + H]+: calcd 500.07, found 500.10. HRMS m/z [M + H]+: calcd 500.1155 found 500.1210.

**Synthesis of *N*-(5-(((2-chloro-[1,1'-biphenyl]-4-yl)methyl)amino)pentyl)-3-methyl-3*H*-imidazo[4,5-*b*]pyridine-6-sulfonamide 61**

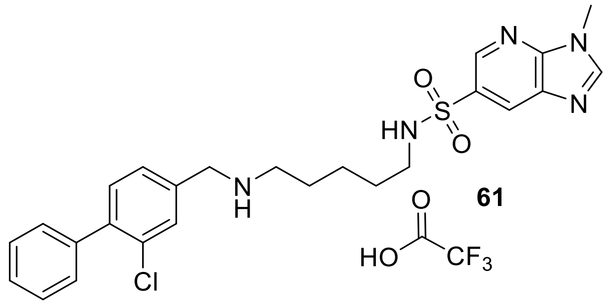

**61**

**61** was synthesized based on [13]. **19** (75 mg, 0.19 mmol, 1eq.) was deprotected using **General Procedure B**. The crude was solved in dry DMSO (3mL) and transferred into a microwave vial. **60** (40 mg, 0.19, 1 eq.) and Triethylamine (77 µL, 0.58 mmol, 3 eq.) were added and the reaction was heated to 100°C for 10h in a microwave oven. DMSO was evaporated using a SpeedVac.

The crude was solved in DCM and washed 2x with water and 2x with brine. Solvents were evaporated and purified using reverse phase flash chromatography ($H_2O$/ACN as eluents). The product was not completely pure after the first round of chromatography and product was further subjected to reverse phase flash chromatography ($H_2O$/ACN as eluents with addition of 0.2% of TFA) yielding the title compound as a TFA salt (17 mg, 18%). MS-ESI m/z [M + H]+: calcd 499.04, found 498.20. HRMS m/z [M + H]+: calcd 498.1652 found 498.1709. $^1$H NMR (500 MHz, DMSO-$d_6$) δ 8.89 (s, 2H), 8.79 (d, J = 1.5 Hz, 1H), 8.68 (s, 1H), 8.42 (s, 1H), 7.75 (s, 1H), 7.72 (s, 1H), 7.56 – 7.45 (m, 4H), 7.45 – 7.39 (m, 3H), 4.18 (d, J = 5.6 Hz, 2H), 3.89 (s, 3H), 2.95 – 2.83 (m, 2H), 2.76 (q, J = 6.6 Hz, 2H), 1.58 (q, J = 7.7 Hz, 2H), 1.48 – 1.38 (m, 3H), 1.30 (q, J = 7.9 Hz, 3H). $^{13}$C NMR (126 MHz, DMSO-$d_6$) δ 150.53 – 147.67 (m), 142.52, 140.76, 138.54, 134.56 – 134.07 (m), 133.75, 132.17, 131.81, 131.69, 131.60, 129.61, 129.54, 128.81, 128.51, 126.22, 49.50, 47.11, 42.74, 30.32, 28.86, 25.36, 23.48.

**Synthesis of *N*-(5-(((2-chloro-[1,1'-biphenyl]-4-yl)methyl)amino)pentyl)-2-oxo-2,3-dihydrobenzo[*d*]oxazole-6-sulfonamide 28**

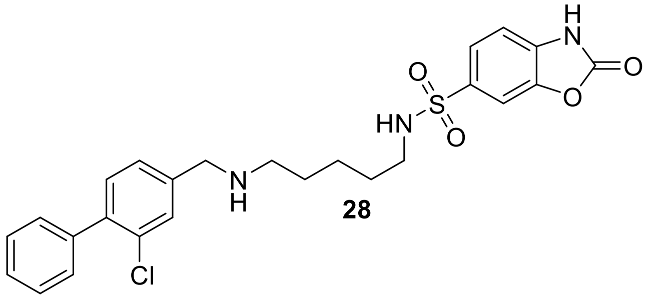

**28**

**19** (50 mg, 0.12 mmol, 1eq.) was deprotected using **General Procedure B**. The crude product was reacted with 2-oxo-2,3-dihydrobenzo[*d*]oxazole-6-sulfonyl chloride (28 mg, 0.12 mmol, 1 eq.) according to **General Procedure A** (20 mg, 34%). $^1$H-NMR (400 MHz, DMSO-$d_6$) δ = 9.61 (s, 1H, H-29), 8.69 (d, J = 1.9 Hz, 1H, H-24), 8.27 (d, J = 8.6 Hz, 1H, H-22), 7.92 (dd, J = 8.6, 1.9 Hz, 1H, H-27), 7.69 (bs, 1H, NH-15), 7.51 – 7.35 (m, 9H, H-3,4,7,9,10,11,12,13, NH-21), 7.32 (s, 2H, H-1), 2.77 (t, J = 6.9 Hz, 2H, H-16), 2.39 (t, J = 6.9 Hz, 2H, H-20), 2.07 (s, 2H), 1.44 – 1.15 (m, 6H, H-17,18,19). $^{13}$C-NMR (101 MHz, DMSO-$d_6$) δ = 160.47 (C-29), 154.94 (C-26), 142.67

(C-2), 138.71 (C-8), 137.80 (C-5), 137.54 (C-23), 134.09 (C-6), 131.12 (C-25), 130.95 (C-7), 129.22 (C-10,12), 128.90 (C-22), 128.17 (C-9,11,13), 127.57 (C-4), 126.92 (C-3), 124.18 (C-27), 123.67 (C-24), 122.11, 51.94 (C-1), 48.43 (C-20), 42.60 (C-16), 28.93 (C-17,19), 23.85 (C-18).. MS-ESI m/z [M + H]+: calcd 500.07, found 500.10. HRMS m/z [M + H]+: calcd 500.1155, found 500.1210

**Synthesis of N-(2-(2-(((2-chloro-[1,1'-biphenyl]-4-yl)methyl)amino)ethoxy)ethyl)isoquinoline-5-sulfonamide 29**

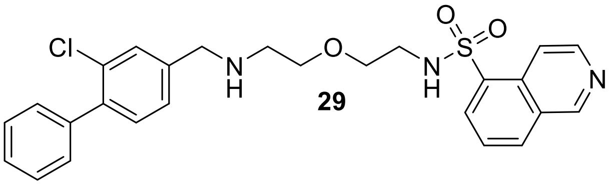

**29**

**20** (31 mg, 0.08 mmol, 1 eq.) was deprotected using **General Procedure B**. The crude product was reacted with 5-isoquinolinesulfonylchloride (18 mg, 0.08 mmol, 1 eq.) according to **General Procedure A** (32 mg, 82%). $^1$H NMR (400 MHz, DMSO-$d_6$) δ 9.46 (s, 1H), 8.68 (d, J = 6.1 Hz, 1H), 8.48 – 8.31 (m, 3H), 7.80 (t, J = 7.8 Hz, 1H), 7.51 – 7.38 (m, 6H), 7.32 (d, J = 1.6 Hz, 2H), 3.67 (s, 2H), 3.26 (t, J = 5.5 Hz, 2H), 3.22 (t, J = 5.6 Hz, 2H), 3.01 (t, J = 5.5 Hz, 2H), 2.42 (t, J = 5.6 Hz, 2H). $^{13}$C NMR (101 MHz, DMSO-$d_6$) δ 153.31, 144.50, 142.45, 138.70, 137.89, 135.27, 133.24, 132.17, 131.17, 130.99, 130.42, 129.22, 128.95, 128.67, 128.19, 127.60, 126.95, 126.35, 117.32, 69.70, 68.70, 51.71, 47.70, 42.33. MS-ESI m/z [M + H]+: calcd 497.02, found 496.20. HRMS m/z [M + H]+: calcd 496.1383, found 494.1454.

**Synthesis of *N*-(2-(2-(((2-chloro-[1,1'-biphenyl]-4-yl)methyl)amino)ethoxy)ethyl)benzo-[*d*]thiazole-6-sulfonamide 30**

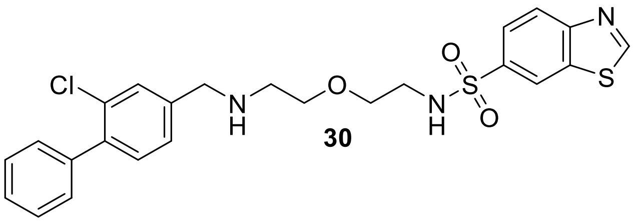

**30**

**20** (31 mg, 0.08 mmol, 1eq.) was deprotected using **General Procedure B**. The crude product was reacted with benzo[*d*]thiazole-6-sulfonyl chloride (17 mg, 0.07 mmol, 0.95 eq.) according to **General Procedure A** (25 mg, 71%). $^{1}$H NMR (500 MHz, DMSO-d$_6$) δ 9.61 (s, 1H), 8.70 (d, J = 1.9 Hz, 1H), 8.26 (d, J = 8.5 Hz, 1H), 7.94 (dd, J = 8.6, 1.9 Hz, 1H), 7.50 – 7.38 (m, 6H), 7.33 (d, J = 1.5 Hz, 2H), 3.71 (s, 2H), 3.36 (q, J = 3.2 Hz, 4H), 2.99 (t, J = 5.6 Hz, 2H), 2.56 (t, J = 5.6 Hz, 2H). $^{13}$C NMR (126 MHz, DMSO-d$_6$) δ 160.48, 154.96, 142.38, 138.69, 137.92, 137.72, 134.06, 131.17, 131.00, 129.22, 128.98, 128.19, 127.59, 126.99, 124.20, 123.64, 122.06, 69.75, 68.78, 51.72, 47.82, 42.56. MS-ESI m/z [M + H]+: calcd 503.04, found 502.05. HRMS m/z [M + H]+: calcd 502.0947, found 502.1009.

**Synthesis of *N*-(2-(2-(((2-chloro-[1,1'-biphenyl]-4-yl)methyl)amino)ethoxy)ethyl)-2-methylbenzo[*d*]thiazole-6-sulfonamide 31**

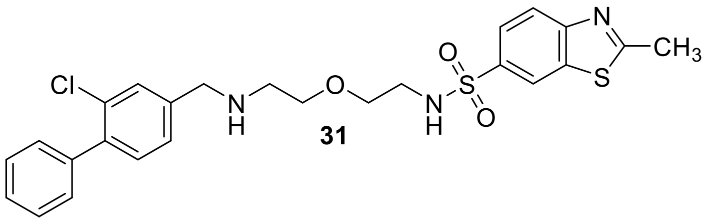

**31**

**20** (31 mg, 0.08 mmol, 1 eq.) was deprotected using **General Procedure B**. The crude product was reacted with 2-methylbenzo[*d*]thiazole-6-sulfonyl chloride (17 mg, 0.07 mmol, 0.95 eq.) according to **General Procedure A** (29 mg, 81%). $^{1}$H NMR (400 MHz, DMSO-d$_6$) δ 8.55 (d, J = 1.9 Hz, 1H), 8.07 (d, J = 8.6 Hz, 1H), 7.88 (dd, J = 8.6, 1.8 Hz, 1H), 7.50 (s, 1H), 7.49 – 7.37 (m, 5H), 7.33 (d, J = 1.4 Hz, 2H), 3.35 (t, J = 5.2 Hz, 4H), 2.98 (t, J = 5.6 Hz, 2H), 2.84 (d, J = 1.3 Hz, 3H), 2.56 (t, J = 5.6 Hz, 2H). $^{13}$C NMR (101 MHz, DMSO-d$_6$) δ 171.63, 154.99, 142.42, 138.69, 137.90, 136.93, 135.56, 131.17, 130.99, 129.22, 128.97, 128.18, 127.59, 126.97,

124.14, 122.40, 121.42, 69.76, 68.76, 51.73, 47.84, 42.55, 20.01. MS-ESI m/z [M + H]+: calcd 517.07, found 516.05. HRMS m/z [M + H]+: calcd 516,1104 found 516.1168.

**Synthesis of 2-chloro-*N*-(2-(2-(((2-chloro-[1,1'-biphenyl]-4-yl)methyl)amino)ethoxy)ethyl)benzo[*d*]thiazole-6-sulfonamide 32**

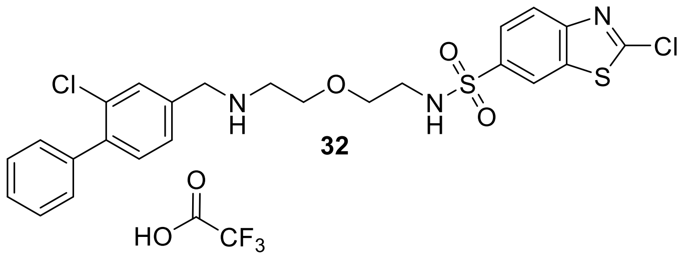

**32**

**20** (31 mg, 0.08 mmol, 1 eq.) was deprotected using **General Procedure B**. The crude product was reacted with 2-chlorobenzo[*d*]thiazole-6-sulfonyl chloride (20 mg, 0.07 mmol, 1 eq.) according to **General Procedure A** (30 mg, 81%). $^1$H NMR (400 MHz, DMSO-d$_6$) δ 8.93 (s, 2H), 8.67 (d, J = 1.9 Hz, 1H), 8.17 (d, J = 8.6 Hz, 1H), 7.94 (dd, J = 8.6, 1.9 Hz, 1H), 7.85 (t, J = 5.8 Hz, 1H), 7.73 (d, J = 1.6 Hz, 1H), 7.47 (dtd, J = 25.1, 7.4, 1.6 Hz, 8H), 4.25 – 4.18 (m, 2H), 3.61 (t, J = 5.1 Hz, 2H), 3.47 (s, 2H), 3.11 (d, J = 5.8 Hz, 2H), 3.02 (q, J = 5.8 Hz, 2H). $^{13}$C NMR (101 MHz, DMSO-d$_6$) δ 157.71, 153.04, 140.84, 138.52, 138.27, 136.70, 133.51, 132.19, 131.87, 131.83, 129.72, 129.61, 128.84, 128.55, 125.41, 123.64, 122.53, 69.41, 65.77, 49.57, 46.49, 42.65. MS-ESI m/z [M + H]+: calcd 537.49, found 536.05 HRMS m/z [M + H]+: calcd 536.0558, found 536.0628.

**Synthesis of *N*-(2-(2-(((2-chloro-[1,1'-biphenyl]-4-yl)methyl)amino)ethoxy)ethyl)-2-oxo-2,3-dihydro-1*H*-benzo[*d*]imidazole-5-sulfonamide TFA salt 33**

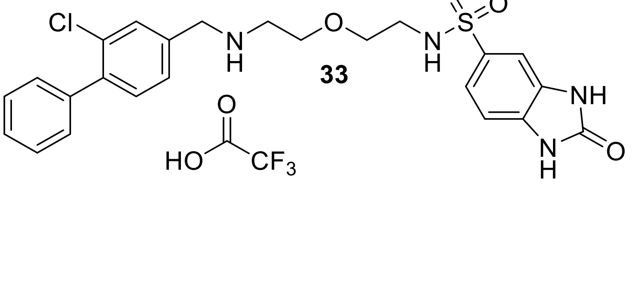

**33**

**20** (31 mg, 0.08 mmol, 1eq.) was deprotected using **General Procedure B**. The crude product was reacted with 2-oxo-2,3-dihydro-1H-benzo[d]imidazole-5-sulfonyl chloride (16 mg, 0.07 mmol, 0.95 eq.) according to **General Procedure A**. The product was not completely pure after the first round of chromatography and product was further subjected to reverse phase flash chromatography ($H_2O$/ACN as eluents with addition of 0.2% of TFA) yielding the title compound as a TFA salt (20 mg, 57%). $^1H$ NMR (400 MHz, DMSO-$d_6$) δ 11.14 (s, 1H), 11.07 (s, 1H), 9.72 (s, 1H), 9.24 (s, 2H), 7.78 (d, J = 1.7 Hz, 1H), 7.56 – 7.40 (m, 8H), 7.34 (d, J = 1.8 Hz, 1H), 7.09 (d, J = 8.2 Hz, 1H), 4.23 (s, 2H), 3.63 (t, J = 5.1 Hz, 2H), 3.06 (dd, J = 7.4, 4.0 Hz, 4H), 2.91 (q, J = 5.8 Hz, 2H). $^{13}C$ NMR (101 MHz, DMSO-$d_6$) δ 155.39, 140.30, 138.12, 133.11, 132.15, 131.66, 131.49, 131.34, 129.73, 129.33, 129.17, 128.36, 128.05, 119.99, 108.30, 106.72, 68.93, 65.27, 49.01, 45.59, 45.57, 42.18.MS-ESI m/z [M + H]+: calcd 501.99, found 501.05. HRMS m/z [M + H]+: calcd 501.1285, found 501.1353.

**Synthesis of *N*-(2-(2-(((2-chloro-[1,1'-biphenyl]-4-yl)methyl)amino)ethoxy)ethyl)-1,3-dimethyl-2-oxo-2,3-dihydro-1*H*-benzo[*d*]imidazole-5-sulfonamide TFA salt 34**

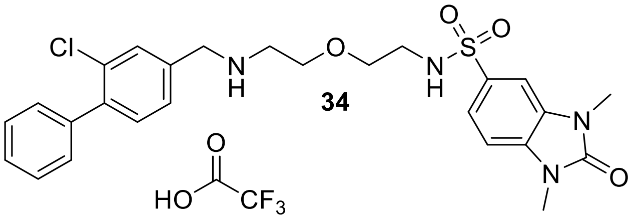

34

**20** (31 mg, 0.08 mmol, 1 eq.) was deprotected using **General Procedure B**. The crude product was reacted with 1,3-dimethyl-2-oxo-2,3-dihydro-1H-benzo[d]imidazole-5-sulfonyl chloride (18 mg, 0.07 mmol, 0.95 eq.) according to **General Procedure A**. The product was not completely pure after the first round of chromatography and product was further subjected to reverse phase flash chromatography ($H_2O$/ACN as eluents with addition of 0.2% of TFA) yielding the title compound as a TFA salt (33 mg, 89%). $^1H$ NMR (400 MHz, DMSO-$d_6$) δ 9.08 (s, 2H), 7.74 (d, J = 1.7 Hz, 1H), 7.57 – 7.40 (m, 10H), 7.34 – 7.28 (m, 1H), 4.22 (d, J = 5.6 Hz, 2H), 3.63 (t, J = 5.0 Hz, 2H), 3.45 (d, J = 11.5 Hz, 2H), 3.37 (s, 3H), 3.36 (s, 3H), 3.12 (p, J = 5.1 Hz, 2H), 2.95

(q, J = 5.8 Hz, 2H), 2.07 (s, 2H). [13]C NMR (101 MHz, DMSO-d$_6$) δ 140.35, 138.09, 133.08, 132.73, 131.69, 131.42, 131.36, 129.70, 129.25, 129.15, 128.35, 128.06, 120.16, 107.54, 105.93, 69.00, 65.33, 49.09, 46.00, 42.18, 27.24, 27.18, 1.15. MS-ESI m/z [M + H]+: calcd 530.05, found 529.15. HRMS m/z [M + H]+: calcd 529.1598, found 529.1654.

**Synthesis of *N*-(2-(2-(((2-chloro-[1,1'-biphenyl]-4-yl)methyl)amino)ethoxy)ethyl)-3-cyanobenzenesulfonamide 35**

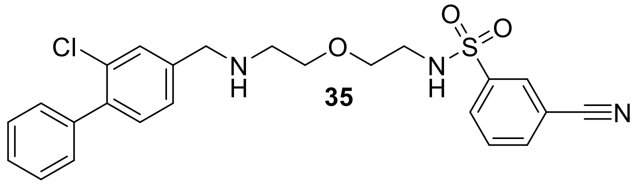

**35**

**20** (31 mg, 0.08 mmol, 1eq.) was deprotected using **General Procedure B**. The crude product was reacted with 3-cyanobenzenesulfonyl chloride (14 mg, 0.07 mmol, 0.95 eq.) according to **General Procedure A** (30 mg, 91%). [1]H NMR (400 MHz, DMSO-d$_6$) δ 8.22 (t, J = 1.7 Hz, 1H), 8.11 (dd, J = 8.0, 1.7 Hz, 2H), 7.81 (t, J = 7.9 Hz, 1H), 7.58 (d, J = 1.5 Hz, 1H), 7.51 – 7.36 (m, 7H), 3.87 (s, 2H), 3.44 (t, J = 5.5 Hz, 2H), 3.38 (t, J = 5.5 Hz, 2H), 3.01 (t, J = 5.5 Hz, 2H), 2.72 (t, J = 5.5 Hz, 2H). [13]C NMR (101 MHz, DMSO-d$_6$) δ 142.15, 138.56, 138.53, 135.94, 131.32, 131.10, 130.91, 130.71, 130.02, 129.62, 129.20, 128.23, 127.72, 127.59, 117.63, 112.40, 68.81, 68.61, 51.06, 47.34, 42.45.MS-ESI m/z [M + H]+: calcd 470.98, found 470.10. HRMS m/z [M + H]+: calcd 470.1227, found 494.1292.

**Synthesis of *N*-(6-(((2-chloro-[1,1'-biphenyl]-4-yl)methyl)amino)hexyl)isoquinoline-5-sulfonamide 36**

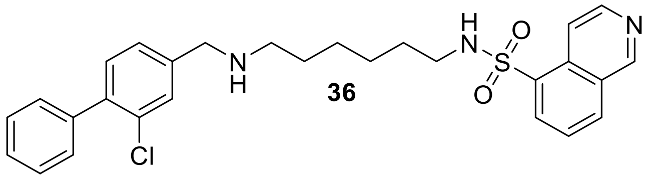

**36**

**21** (36 mg, 0.09 mmol, 1 eq.) was deprotected using **General Procedure B**. The crude product was reacted with 5-Isoquinolinsulfonylchloride (23 mg, 0.09 mmol, 1 eq.) according to **General Procedure A** (41 mg, 90%). $^1$H NMR (400 MHz, DMSO-$d_6$) δ 9.47 (s, 1H), 8.69 (d, J = 6.1 Hz, 1H), 8.43 (dd, J = 12.1, 7.2 Hz, 2H), 8.33 (d, J = 7.3 Hz, 1H), 8.06 (s, 1H), 7.82 (t, J = 7.7 Hz, 1H), 7.53 (s, 1H), 7.51 – 7.40 (m, 5H), 7.36 (s, 2H), 3.76 (s, 2H), 2.78 (t, J = 7.0 Hz, 2H), 2.42 (d, J = 7.3 Hz, 2H), 1.24 (d, J = 7.1 Hz, 4H), 1.06 (q, J = 3.5 Hz, 4H). $^{13}$C NMR (101 MHz, DMSO-$d_6$) δ 153.38, 144.51, 138.60, 138.24, 135.05, 133.29, 132.36, 131.24, 131.02, 130.39, 129.31, 129.21, 128.22, 127.68, 127.32, 126.41, 117.25, 51.42, 48.11, 42.20, 28.93, 28.54, 26.03, 25.73. MS-ESI m/z [M + H]+: calcd 509.078, found 508.15. HRMS m/z [M + H]+: calcd 508.1747, found 508.1808.

**Synthesis of *N*-(6-(((2-chloro-[1,1'-biphenyl]-4-yl)methyl)amino)hexyl)-3-methylquinoline-8-sulfonamide 37**

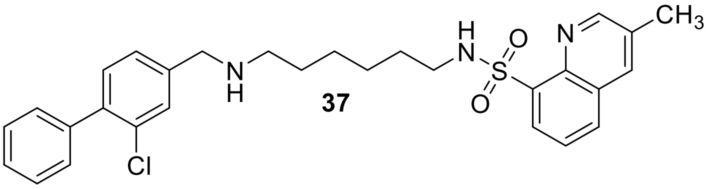

**21** (36 mg, 0.09 mmol, 1 eq.) was deprotected using **General Procedure B**. The crude product was reacted with 3-methylquinoline-8-sulfonyl chloride (22 mg, 0.09 mmol, 1 eq.) according to **General Procedure A** (39 mg, 83%). $^1$H NMR (400 MHz, DMSO-$d_6$) δ 8.94 (d, J = 2.3 Hz, 1H), 8.31 (dd, J = 2.3, 1.2 Hz, 1H), 8.21 (ddd, J = 12.6, 7.8, 1.5 Hz, 2H), 7.71 (dd, J = 8.2, 7.3 Hz, 1H), 7.56 (d, J = 1.3 Hz, 1H), 7.49 – 7.37 (m, 7H), 7.13 (s, 1H), 3.81 (s, 2H), 2.78 (q, J = 6.5 Hz, 2H), 2.54 (s, 3H), 1.32 – 1.26 (m, 4H), 1.24 (s, 3H), 1.10 (h, J = 4.3, 3.3 Hz, 4H). $^{13}$C NMR (101 MHz, DMSO-$d_6$) δ 153.50, 141.43, 139.02, 138.90, 136.80, 135.85, 133.34, 132.48, 131.75, 130.08, 129.96, 129.68, 128.82, 128.71, 128.19, 127.96, 126.26, 51.68, 48.48, 43.26, 29.38, 28.74, 26.50, 26.30, 22.57, 18.66. MS-ESI m/z [M + H]+: calcd 523.10, found 522.15. HRMS m/z [M + H]+: calcd 522.1903, found 522.1969.

**Synthesis of *N*-(6-(((2-chloro-[1,1'-biphenyl]-4-yl)methyl)amino)hexyl)benzo[*d*]thiazole-6-sulfonamide 38**

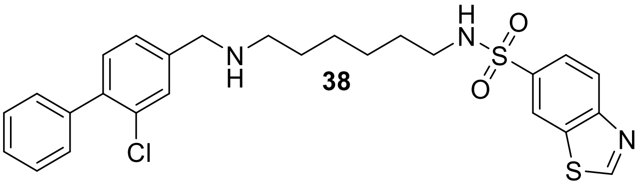

**38**

**21** (36 mg, 0.09 mmol, 1 eq.) was deprotected using **General Procedure B**. The crude product was reacted with benzo[d]thiazole-6-sulfonyl chloride (19 mg, 0.09 mmol, 1 eq.) according to **General Procedure A** (40 mg, 87%). $^{1}$H NMR (400 MHz, DMSO-$d_6$) δ 9.61 (s, 1H), 8.69 (d, J = 1.8 Hz, 1H), 8.27 (d, J = 8.6 Hz, 1H), 7.92 (dd, J = 8.6, 1.9 Hz, 1H), 7.71 (s, 1H), 7.60 (d, J = 1.5 Hz, 1H), 7.49 – 7.42 (m, 4H), 7.41 – 7.37 (m, 3H), 3.92 (s, 2H), 2.77 (t, J = 7.1 Hz, 2H), 2.64 (t, J = 7.4 Hz, 2H), 1.43 (td, J = 14.2, 7.1 Hz, 3H), 1.36 (d, J = 6.8 Hz, 3H), 1.21 (q, J = 3.4 Hz, 4H). $^{13}$C NMR (101 MHz, DMSO-$d_6$) δ 160.50, 154.95, 138.93, 138.42, 137.53, 134.10, 131.38, 131.13, 129.98, 129.18, 129.18, 128.25, 127.92, 127.78, 124.18, 123.68, 122.11, 50.61, 47.67, 42.50, 28.92, 27.48, 25.92, 25.78. MS-ESI m/z [M + H]+: calcd 515.09, found 514.10. HRMS m/z [M + H]+: calcd 514.1311, found 514.1379.

**Synthesis of *N*-(6-(((2-chloro-[1,1'-biphenyl]-4-yl)methyl)amino)hexyl)-2-methylbenzo-[*d*]thiazole-6-sulfonamide 39**

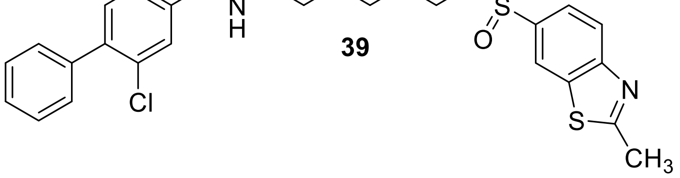

**39**

**21** (36 mg, 0.09 mmol, 1 eq.) was deprotected using **General Procedure B**. The crude product was reacted with 2-methylbenzo[*d*]thiazole-6-sulfonyl chloride (20 mg, 0.09 mmol, 1 eq.) according to **General Procedure A** (42 mg, 88%). $^{1}$H NMR (400 MHz, DMSO-d$_6$) δ 8.54 (d, J = 1.8 Hz, 1H), 8.08 (d, J = 8.6 Hz, 1H), 7.85 (dd, J = 8.6, 1.9 Hz, 1H), 7.63 (s, 1H), 7.51 (d, J = 1.3 Hz, 1H), 7.47 – 7.39 (m, 5H), 7.34 (s, 2H), 3.72 (s, 2H), 2.84 (s, 3H), 2.75 (t, J = 7.1 Hz, 2H), 2.45 (t, J = 7.1 Hz, 2H), 1.34 (q, J = 6.9 Hz, 4H), 1.19 (d, J = 3.9 Hz, 4H). $^{13}$C NMR (101 MHz, DMSO-d$_6$) δ 171.61, 154.97, 142.02, 138.66, 137.98, 136.78, 135.58, 131.16, 130.99, 129.21, 129.07, 128.19, 127.61, 127.08, 124.13, 122.43, 121.46, 51.73, 48.36, 42.53, 29.01, 28.96, 26.24, 25.95, 20.00. MS-ESI m/z [M + H]+: calcd 529.12, found 528.10. HRMS m/z [M + H]+: calcd 528.1468, found 528.1539.

**Synthesis of 2-chloro-*N*-(6-(((2-chloro-[1,1'-biphenyl]-4-yl)methyl)amino)hexyl)benzo-[*d*]thiazole-6-sulfonamide 40**

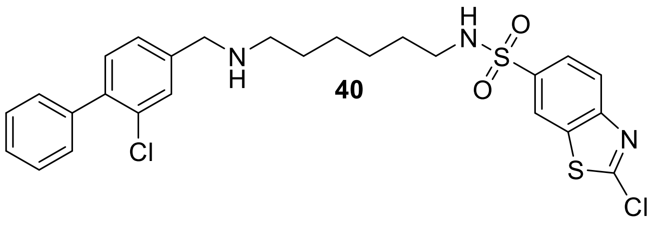

**21** (36 mg, 0.09 mmol, 1 eq.) was deprotected using **General Procedure B**. The crude product was reacted with 2-chlorobenzo[*d*]thiazole-6-sulfonyl chloride (24 mg, 0.09 mmol, 1 eq.) according to **General Procedure A** (5 mg, 10%). $^{1}$H NMR (400 MHz, DMSO-d$_6$) δ 8.66 (d, J = 1.8 Hz, 1H), 8.16 (d, J = 8.6 Hz, 1H), 7.93 (dd, J = 8.6, 1.9 Hz, 1H), 7.78 (d, J = 5.9 Hz, 1H), 7.72 (s, 1H), 7.58 – 7.32 (m, 8H), 4.10 (s, 2H), 2.79 (dt, J = 19.7, 6.9 Hz, 4H), 1.54 (s, 2H), 1.43 – 1.31 (m, 2H), 1.23 (p, J = 4.4, 3.8 Hz, 4H). $^{13}$C NMR (101 MHz, DMSO-d$_6$) δ 152.95, 140.70, 138.57, 138.42, 136.67, 133.89, 132.15, 131.79, 131.75, 129.62, 128.82, 128.73, 128.51, 125.44, 123.59, 122.51, 49.53, 47.20, 42.93, 29.30, 26.02, 25.96, 25.82. MS-ESI m/z [M + H]+: calcd 549.54, found 548.10. HRMS m/z [M + H]+: calcd 548.0921, found 548.1002.

**Synthesis of *N*-(6-(((2-chloro-[1,1'-biphenyl]-4-yl)methyl)amino)hexyl)-2-oxo-2,3-dihydrobenzo[*d*]oxazole-6-sulfonamide 41**

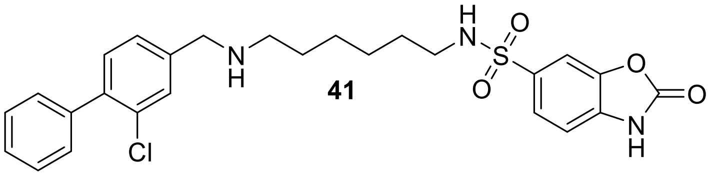

**21** (36 mg, 0.09 mmol, 1 eq.) was deprotected using **General Procedure B**. The crude product was reacted with 2-oxo-2,3-dihydrobenzo[d]oxazole-6-sulfonyl chloride (21 mg, 0.09 mmol, 1 eq.) according to **General Procedure A** (38 mg, 86%). $^{1}H$ NMR (400 MHz, DMSO-$d_6$) δ 7.68 (d, J = 1.6 Hz, 1H), 7.60 – 7.53 (m, 2H), 7.52 – 7.39 (m, 8H), 7.25 (d, J = 8.1 Hz, 1H), 4.05 (s, 2H), 2.73 (dt, J = 26.5, 7.1 Hz, 4H), 1.51 (p, J = 7.3 Hz, 2H), 1.34 (q, J = 6.8 Hz, 2H), 1.21 (p, J = 3.8 Hz, 4H). $^{13}C$ NMR (101 MHz, DMSO-$d_6$) δ 155.41, 143.35, 139.52, 138.28, 136.14, 135.83, 133.21, 131.52, 131.21, 130.59, 129.17, 128.51, 128.30, 127.90, 122.87, 109.91, 107.33, 49.93, 47.26, 42.41, 28.75, 26.52, 25.73, 25.69. MS-ESI m/z [M + H]+: calcd 515.04, found 514.20. HRMS m/z [M + H]+: calcd 514.1489, found 514.1554.

**Synthesis of *N*-((2-chloro-[1,1'-biphenyl]-4-yl)methyl)-4-(imidazo[1,2-*a*]pyridin-6-ylsulfonyl)butan-1-amine FA salt 66**

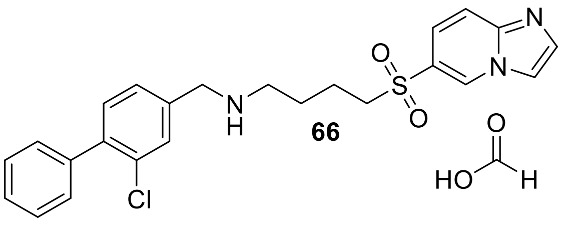

**66** was synthesized based on. [13] **65** (20 mg, 0.08 mmol, 1 eq.) was solved in 1.5 mL of dry DCM and deprotected using 5 mL 4M HCl in Dioxane. After 1h 300 µL of triethylamine were added to the solution and stirred 5 min. All solvents were evaporated, the crude was solved in dry DCM (5 mL) and transferred into a 50 mL round bottom flask. **18** (17 mg, 0.08, 1 eq.) and sodium triacetoxyborohydride (42 mg, 0.2 mmol, 2.5 eq.) were added and the reaction was stirred over night at room temperature. Sat. $NaHCO_3$ was added and stirred rigorously for 5 min. The mixture was transferred into a separatory funnel and washed 2x with water and 2x with brine and dried over $MgSO_4$. Solvents were evaporated and the crude purified using reverse phase flash chromatography ($H_2O$/ACN as eluents). The product was not completely pure after the first round of chromatography and product was further subjected to reverse phase flash chromatography ($H_2O$/ACN as eluents with addition of 0.1% of formic acid) yielding the title compound as a FA salt (4 mg, 23%). $^1$H NMR (400 MHz, DMSO-$d_6$) δ 9.28 (s, 1H), 8.26 (s, 1H), 8.19 (s, 1H), 7.77 (d, J = 8.9 Hz, 2H), 7.59 – 7.37 (m, 7H), 7.32 (s, 2H), 3.73 (d, J = 2.9 Hz, 2H), 3.42 (t, J = 7.8 Hz, 3H), 2.55 (d, J = 6.9 Hz, 2H), 1.68 (t, J = 7.9 Hz, 2H), 1.54 (p, J = 7.1 Hz, 2H). $^{13}$C NMR (101 MHz, DMSO-$d_6$) δ 138.61, 138.19, 135.36, 131.21, 131.01, 130.69, 129.27, 129.21, 128.22, 127.66, 127.25, 124.77, 121.34, 117.40, 115.44, 54.67, 51.32, 47.42, 27.12, 20.25. MS-ESI m/z [M + H]+: calcd 454.98, found 454.15. HRMS m/z [M + H]+: calcd 454.1278, found 454.1336.

**Synthesis of *N*-((2-chloro-[1,1'-biphenyl]-4-yl)methyl)-6-((3-methyl-3H-imidazo[4,5-*b*]pyridin-6-yl)sulfonyl)hexan-1-amine TFA salt 63**

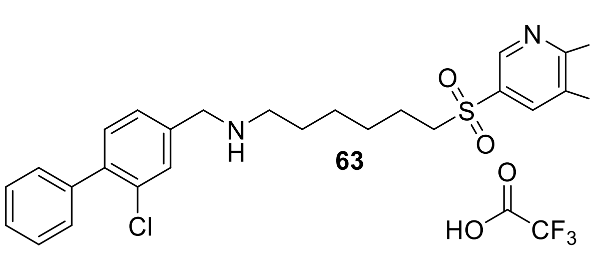

**63**

**62** (35 mg, 0.09 mmol, 1 eq.) was deprotected using **General Procedure B**. All solvents were evaporated, solved in DCM and 0.5 mL of triethylamine were added and stirred for 5 min. All solvents were evaporated again $MgSO_4$ (21 mg, 0.176 mmol, 2 eq.), sodium triacetoxyborohydride (37 mg, 0.176mmol, 2 eq.) and **18** (19 mg, 0.08 mmol, 1 eq.) were added to the flask. Dry DCM (5 mL) was added and the reaction was stirred for 16 h at ambient temperature. Solvents were evaporated and the crude purified using reverse phase flash chromatography ($H_2O$/ACN as eluents). The product was not completely pure after the first round of chromatog-raphy and product was further subjected to reverse phase flash chromatography ($H_2O$/ACN as eluents with addition of 0.2% of TFA) yielding the title compound as a TFA salt (11 mg, 25%). $^1$H NMR (400 MHz, DMSO-$d_6$) δ 8.98 (s, 2H), 8.85 (d, J = 2.0 Hz, 1H), 8.72 (s, 1H), 8.55 (d, J = 2.0 Hz, 1H), 7.73 (d, J = 1.6 Hz, 1H), 7.53 – 7.40 (m, 7H), 4.19 (t, J = 5.8 Hz, 2H), 3.91 (s, 3H), 3.48 – 3.39 (m, 2H), 2.91 (dq, J = 11.7, 6.4 Hz, 2H), 1.57 (qd, J = 7.5, 3.9 Hz, 4H), 1.37 – 1.25 (m, 4H). $^{13}$C NMR (101 MHz, DMSO-$d_6$) δ 150.00, 149.38, 143.16, 140.27, 138.09, 133.86, 133.31, 131.69, 131.35, 131.24, 129.68, 129.14, 129.08, 128.33, 128.03, 127.27, 55.19, 49.04, 46.64, 29.90, 26.86, 25.29, 25.07, 22.16. MS-ESI m/z [M + H]+: calcd 498.05, found 497.15. HRMS m/z [M + H]+: calcd 497.1699, found 497.1764.

**Synthesis of *N*-((2-chloro-[1,1'-biphenyl]-4-yl)methyl)-2-(1-(isoquinolin-5-ylsulfonyl)piperidin-4-yl)ethan-1-amine TFA salt 48**

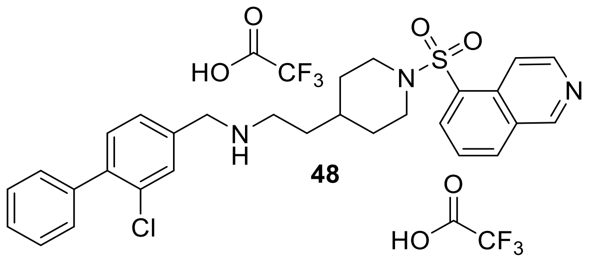

**48**

**22** (29 mg, 0.07 mmol, 1 eq.) was deprotected using **General Procedure B**. The crude product was reacted with 5-isoquinolinesulfonylchloride (18 mg, 0.07 mmol, 1 eq.) according to **General Procedure A**. The product was not completely pure after the first round of chromatography and product was further subjected to reverse phase flash chromatography ($H_2O$/ACN as eluents with addition of 0.2% of TFA) yielding the title compound as a TFA salt (23 mg, 63%). $^1$H NMR (400 MHz, DMSO-$d_6$) δ 9.50 (s, 1H), 8.98 (s, 2H), 8.70 (d, J = 6.1 Hz, 1H), 8.50 (dt, J = 8.2, 1.1 Hz, 1H), 8.46 (d, J = 6.1 Hz, 1H), 8.36 (dd, J = 7.4, 1.2 Hz, 1H), 7.88 (dd, J = 8.2, 7.4 Hz, 1H), 7.70 (d, J = 1.6 Hz, 1H), 7.52 – 7.38 (m, 7H), 4.17 (s, 2H), 3.75 (dt, J = 12.9, 3.6 Hz, 2H), 2.93 (t, J = 7.8 Hz, 2H), 2.44 (td, J = 12.1, 2.5 Hz, 2H), 1.73 – 1.62 (m, 2H), 1.51 (q, J = 7.1 Hz, 2H), 1.39 – 1.29 (m, 1H), 1.10 (qd, J = 12.3, 4.0 Hz, 2H). $^{13}$C NMR (101 MHz, DMSO-$d_6$) δ 153.57,

144.87, 140.22, 138.06, 134.22, 134.07, 133.28, 131.65, 131.53, 131.31, 131.19, 131.04, 129.12, 129.03, 128.31, 128.01, 126.56, 117.18, 48.99, 45.43, 44.42, 31.59, 31.34, 30.75.MS-ESI m/z [M + H]+: calcd 521.09, found 519.94. HRMS m/z [M + H]+: calcd 520.1747, found 520.1797.

**Synthesis of 2-(1-(benzo[*d*]thiazol-6-ylsulfonyl)piperidin-4-yl)-*N*-((2-chloro-[1,1'-biphenyl]-4-yl)methyl)ethan-1-amine 50**

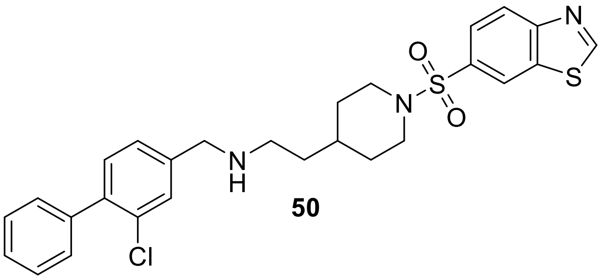

**50**

**22** (378 mg, 0.9 mmol, 2 eq.) was deprotected using **General Procedure B**. The crude product was reacted with Then benzo[*d*]thiazole-6-sulfonyl chloride (105 mg, 0.45 mmol, 1 eq.) according to **General Procedure A** . The product was recrystallized from EtOH (169 mg, 71%). $^{1}$H NMR (500 MHz, DMSO-$d_6$) δ 9.66 (s, 2H), 8.71 (d, J = 1.8 Hz, 1H), 8.29 (d, J = 8.6 Hz, 1H), 7.84 (dd, J = 8.6, 1.8 Hz, 2H), 7.63 (dq, J = 7.9, 1.3 Hz, 1H), 7.47 (tt, J = 6.9, 1.2 Hz, 2H), 7.45 – 7.37 (m, 4H), 4.11 (s, 2H), 3.67 (dt, J = 12.1, 3.4 Hz, 2H), 2.85 (t, J = 7.8 Hz, 2H), 2.22 (td, J = 11.9, 2.5 Hz, 2H), 1.74 – 1.68 (m, 2H), 1.61 (q, J = 7.3 Hz, 2H), 1.32 (tq, J = 10.7, 3.7 Hz, 1H), 1.23 – 1.12 (m, 2H). $^{13}$C NMR (101 MHz, DMSO-$d_6$): δ 161.17, 155.40, 140.00, 138.16, 134.49, 133.45, 132.27, 131.43, 131.14, 129.30, 129.13, 128.29, 127.94, 124.99, 123.61, 123.26, 48.73, 45.99, 44.17, 31.59, 31.21, 30.62 MS-ESI m/z [M + H]+: calcd 527.11, found 525.97. HRMS m/z [M + H]+: calcd 526.1311, found 526.1374.

**Synthesis of *N*-((2-chloro-[1,1'-biphenyl]-4-yl)methyl)-2-(1-((3-methylquinolin-8-yl)sulfonyl)piperidin-4-yl)ethan-1-amine 49**

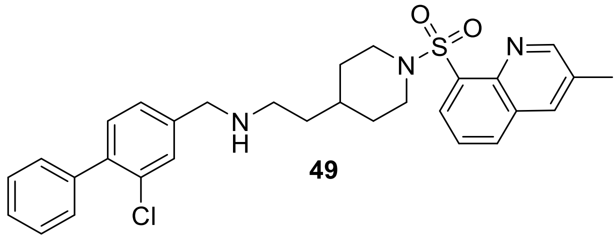

**49**

**22** (29 mg, 0.07 mmol, 1 eq.) was deprotected using **General Procedure B**. The crude product was reacted with 3-methylquinoline-8-sulfonyl chloride (17 mg, 0.07 mmol, 1 eq.) according to **General Procedure A** (19 mg, 54%). $^{1}$H NMR (400 MHz, DMSO-$d_6$) δ 8.92 (d, J = 2.3 Hz, 1H), 8.31 – 8.26 (m, 1H), 8.20 (dd, J = 8.3, 1.4 Hz, 1H), 7.74 – 7.68 (m, 2H), 7.53 – 7.38 (m, 7H), 4.18 (s, 2H), 3.88 (dt, J = 13.0, 3.6 Hz, 2H), 2.95 (t, J = 7.8 Hz, 2H), 2.66 (td, J = 12.4, 2.5 Hz, 2H), 2.52 (d, J = 1.0 Hz, 3H), 1.71 – 1.60 (m, 2H), 1.53 (q, J = 7.2 Hz, 2H), 1.41 – 1.30 (m, 1H), 1.12 (tt, J = 11.9, 6.0 Hz, 2H). $^{13}$C NMR (101 MHz, DMSO-$d_6$) δ 153.00, 141.60, 140.24, 138.05, 135.97, 135.11, 133.28, 133.25, 131.85, 131.67, 131.55, 131.31, 131.20, 129.12, 129.05, 128.61, 128.32, 128.02, 125.77, 49.03, 45.93, 44.53, 32.04, 31.59, 31.39, 18.08. MS-ESI m/z [M + H]+: calcd 535.12, found 534.02. HRMS m/z [M + H]+: calcd 534.1904, found 534.1952.

**Synthesis of *N*-((2-chloro-[1,1'-biphenyl]-4-yl)methyl)-2-(1-((2-methylbenzo[*d*]thiazol-6-yl)sulfonyl)piperidin-4-yl)ethan-1-amine 51**

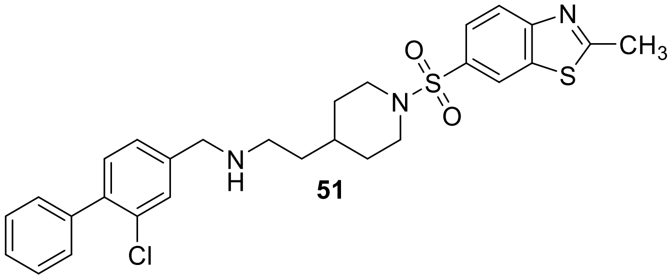

**51**

**22** (29 mg, 0.07 mmol, 1 eq.) was deprotected using **General Procedure B**. The crude product was reacted with with 2-methylbenzo[*d*]thiazole-6-sulfonyl chloride (17 mg, 0.07 mmol, 1 eq.) according to **General Procedure A** (23 mg, 61%). $^{1}$H NMR (400 MHz, DMSO-$d_6$) δ 8.56 (d, J = 1.8 Hz, 1H), 8.11 (d, J = 8.5 Hz, 1H), 7.79 (dd, J = 8.6, 1.9 Hz, 1H), 7.67 (d, J = 1.5 Hz, 1H),

7.51 – 7.38 (m, 7H), 4.10 (s, 2H), 3.66 (dt, J = 12.0, 3.6 Hz, 2H), 2.86 (s, 3H), 2.21 (td, J = 12.0, 11.5, 2.5 Hz, 2H), 1.76 – 1.61 (m, 2H), 1.49 (q, J = 7.3 Hz, 2H), 1.36 – 1.26 (m, 1H), 1.18 (qd, J = 11.8, 3.8 Hz, 3H). $^{13}$C NMR (101 MHz, DMSO-$d_6$) δ 172.32, 155.46, 139.88, 138.14, 135.98, 131.58, 131.36, 131.25, 130.86, 129.12, 128.73, 128.30, 127.95, 124.99, 122.59, 122.35, 49.46, 46.02, 44.68, 31.96, 31.61, 30.57, 20.05. MS-ESI m/z [M + H]+: calcd 541.14, found 540.10. HRMS m/z [M + H]+: calcd 540.1468, found 540.1535.

**Synthesis of *N*-((2-chloro-[1,1'-biphenyl]-4-yl)methyl)-2-(1-((2-chlorobenzo[*d*]thiazol-6-yl)sulfonyl)piperidin-4-yl)ethan-1-amine TFA salt 52**

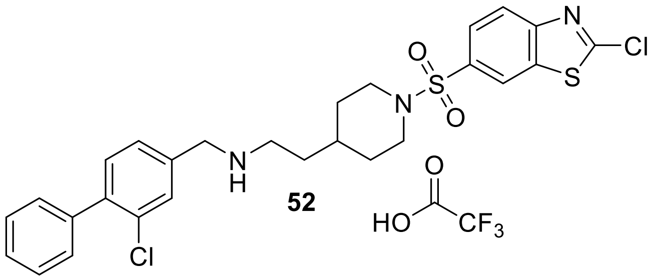

**52**

**22** (68 mg, 0.16 mmol, 1 eq.) was deprotected using **General Procedure B**. The crude product was reacted with 2-chlorobenzo[*d*]thiazole-6-sulfonyl chloride (43 mg, 0.16 mmol, 1 eq.) according to **General Procedure A**. The product was not completely pure after the first round of chromatography and product was further subjected to reverse phase flash chromatography ($H_2O$/ACN as eluents with addition of 0.1% of TFA) yielding the title compound as a TFA salt (52 mg, 89%). $^1$H NMR (400 MHz, DMSO-$d_6$) δ 9.33 (s, 2H), 8.67 (d, J = 1.9 Hz, 1H), 8.18 (d, J = 8.6 Hz, 1H), 7.86 (dd, J = 8.6, 1.9 Hz, 1H), 7.80 (d, J = 1.7 Hz, 1H), 7.58 (dd, J = 8.0, 1.7 Hz, 1H), 7.50 – 7.38 (m, 6H), 4.14 (s, 2H), 3.72 – 3.60 (m, 2H), 2.89 (s, 2H), 2.24 (td, J = 11.8, 2.5 Hz, 2H), 1.76 – 1.67 (m, 2H), 1.58 (q, J = 7.4 Hz, 2H), 1.32 (d, J = 10.8 Hz, 1H), 1.18 (qd, J = 11.8, 3.7 Hz, 2H). $^{13}$C NMR (101 MHz, DMSO-$d_6$) δ 157.67, 152.93, 140.13, 138.10, 136.55, 133.35, 132.73, 131.55, 131.35, 131.22, 129.21, 129.12, 128.31, 127.99, 125.82, 123.06,

123.02, 48.84, 45.96, 44.29, 31.55, 31.26, 30.56. MS-ESI m/z [M + H]+: calcd 561.55, found 560.10. HRMS m/z [M + H]+: calcd 560.0922, found 560.0980.

**Synthesis of *N*-((2-chloro-[1,1'-biphenyl]-4-yl)methyl)-7-(isoquinolin-5-ylsulfonyl)-7-aza-spiro[3.5]nonan-2-amine 53**

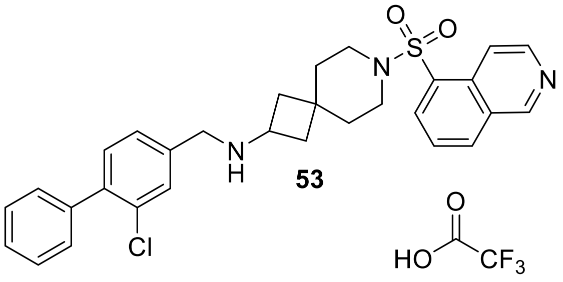

53

**23** (30 mg, 0.07 mmol, 1 eq.) was deprotected using **General Procedure B**. The crude product was reacted with 5-Isoquinolinsulfonylchloride (18 mg, 0.07 mmol, 1 eq.) according to **General Procedure A**. The product was not completely pure after the first round of chromatography and product was further subjected to reverse phase flash chromatography ($H_2O$/ACN as eluents with addition of 0.2% of TFA) yielding the title compound as a TFA salt (18 mg, 49%). $^1$H NMR (400 MHz, DMSO-$d_6$) δ 9.52 (s, 1H), 9.24 (s, 2H), 8.71 (d, J = 6.1 Hz, 1H), 8.51 (d, J = 8.2 Hz, 1H), 8.43 (d, J = 6.1 Hz, 1H), 8.37 (dd, J = 7.4, 1.2 Hz, 1H), 7.91 – 7.86 (m, 1H), 7.70 (d, J = 1.8 Hz, 1H), 7.51 – 7.38 (m, 7H), 4.02 (t, J = 5.6 Hz, 2H), 3.10 (t, J = 5.3 Hz, 2H), 3.04 (t, J = 5.4 Hz, 2H), 2.04 (dd, J = 11.6, 8.9 Hz, 2H), 1.88 (t, J = 10.1 Hz, 2H), 1.59 (dt, J = 11.6, 5.5 Hz, 4H), 1.27 – 1.22 (m, 1H). $^{13}$C NMR (101 MHz, DMSO-$d_6$) δ 153.55, 144.79, 140.28, 138.05, 134.28, 134.06, 133.16, 131.85, 131.34, 131.13, 129.13, 129.00, 128.35, 128.06, 126.65, 117.23, 46.70, 42.46, 37.54, 34.50, 31.44. MS-ESI m/z [M + H]+: calcd 533.09, found 532.15. HRMS m/z [M + H]+: calcd 532.1747, found 532.1804.

**Synthesis of 7-(benzo[*d*]thiazol-6-ylsulfonyl)-*N*-((2-chloro-[1,1'-biphenyl]-4-yl)methyl)-7-azaspiro[3.5]nonan-2-amine TFA salt 54**

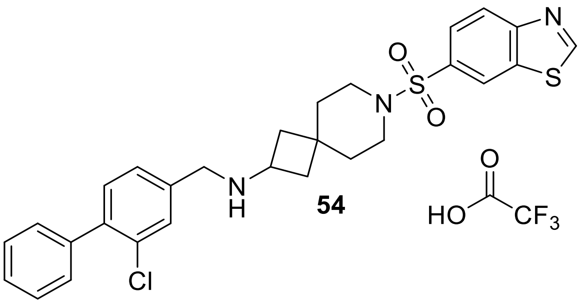

**54**

**23** (38 mg, 0.09 mmol, 1 eq.) was deprotected using **General Procedure B**. The crude product was reacted with benzo[*d*]thiazole-6-sulfonyl chloride (20 mg, 0.09 mmol, 1 eq.) according to **General Procedure A**. The product was not completely pure after the first round of chromatography and product was further subjected to reverse phase flash chromatography ($H_2O$/ACN as eluents with addition of 0.2% of TFA) yielding the title compound as a TFA salt (4 mg, 8%). $^1H$ NMR (400 MHz, DMSO-$d_6$) δ 9.66 (s, 1H), 9.23 (s, 2H), 8.72 (dd, J = 1.9, 0.6 Hz, 1H), 8.31 (dd, J = 8.6, 0.6 Hz, 1H), 7.86 (dd, J = 8.6, 1.9 Hz, 1H), 7.68 (t, J = 1.0 Hz, 1H), 7.51 – 7.38 (m, 8H), 7.28 – 7.11 (m, 1H), 4.01 (s, 2H), 3.67 (h, J = 7.9 Hz, 1H), 2.95 (t, J = 5.4 Hz, 2H), 2.88 (t, J = 5.6 Hz, 2H), 2.09 – 1.96 (m, 2H), 1.91 – 1.81 (m, 2H), 1.64 (dt, J = 11.1, 5.5 Hz, 4H). $^{13}C$ NMR (101 MHz, DMSO-$d_6$) δ 161.14, 155.41, 140.25, 138.02, 134.50, 133.15, 132.49, 131.69, 131.32, 131.11, 129.10, 128.97, 128.90, 128.32, 128.20, 128.02, 124.93, 123.66, 123.26, 46.65, 46.25, 43.05, 42.81, 37.31, 34.73, 34.30, 31.27. MS-ESI m/z [M + H]+: calcd 539.12, found 538.00. HRMS m/z [M + H]+: calcd 538.1312, found 538.1382.

**Synthesis of *N*-((2-chloro-[1,1'-biphenyl]-4-yl)methyl)-7-((2-methylbenzo[*d*]thiazol-6-yl)sulfonyl)-7-azaspiro[3.5]nonan-2-amine TFA salt 55**

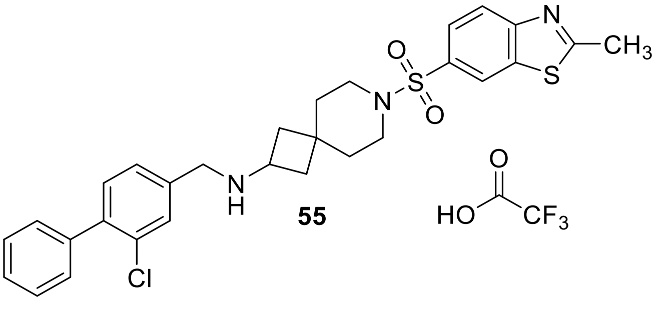

**55**

**23** (40 mg, 0.09 mmol, 1 eq.) was deprotected using **General Procedure B**. The crude product was reacted with 2-methylbenzo[*d*]thiazole-6-sulfonyl chloride (23 mg, 0.09 mmol, 1 eq.) according to **General Procedure A**. The product was not completely pure after the first round of chromatography and product was further subjected to reverse phase flash chromatography (H2O/ACN as eluents with addition of 0.2% of TFA) yielding the title compound as a TFA salt (23 mg, 45%). $^1$H NMR (400 MHz, DMSO-$d_6$) δ 9.33 (s, 2H), 8.57 (d, J = 1.8 Hz, 1H), 8.11 (d, J = 8.6 Hz, 1H), 7.80 (dd, J = 8.6, 1.9 Hz, 1H), 7.69 (d, J = 1.5 Hz, 1H), 7.51 – 7.37 (m, 7H), 4.02 (s, 2H), 3.67 (t, J = 8.1 Hz, 1H), 2.92 (t, J = 5.4 Hz, 2H), 2.07 – 1.96 (m, 2H), 1.92 – 1.83 (m, 2H), 1.64 (dt, J = 10.8, 5.4 Hz, 4H). $^{13}$C NMR (101 MHz, DMSO-$d_6$) δ 172.35, 140.23, 138.04, 135.98, 133.16, 131.67, 131.61, 131.32, 131.12, 129.11, 128.98, 128.31, 128.01, 124.93, 122.57, 122.41, 46.64, 46.24, 43.05, 42.81, 37.26, 34.69, 34.27, 31.27, 20.05. MS-ESI m/z [M + H]+: calcd 553.15, found 551.92. HRMS m/z [M + H]+: calcd 552.14680, found 552.1534.

**Synthesis of *N*-((2-chloro-[1,1'-biphenyl]-4-yl)methyl)-7-((2-chlorobenzo[*d*]thiazol-6-yl)-sulfonyl)-7-azaspiro[3.5]nonan-2-amine 56**

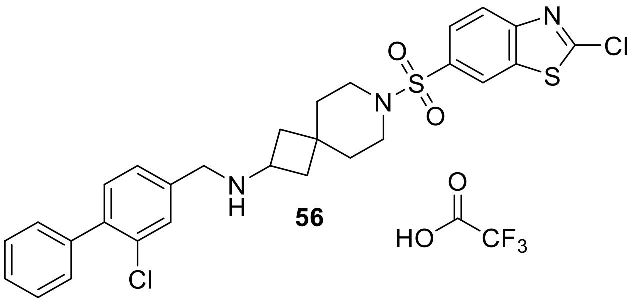

**56**

**23** (30 mg, 0.07 mmol, 1 eq.) was deprotected using **General Procedure B**. The crude product was reacted with 2-chlorobenzo[*d*]thiazole-6-sulfonyl chloride (18 mg, 0.07 mmol, 1 eq.) according to **General Procedure A**. The product was not completely pure after the first round of chromatography and product was further subjected to reverse phase flash chromatography (H2O/ACN as eluents with addition of 0.2% of TFA) yielding the title compound as a TFA salt (36 mg, 95%). $^1$H NMR (400 MHz, DMSO-$d_6$) δ 9.35 (s, 2H), 8.67 (d, J = 1.9 Hz, 1H), 8.18 (d, J

= 8.6 Hz, 1H), 7.90 – 7.84 (m, 1H), 7.70 (d, J = 1.5 Hz, 1H), 7.49 – 7.38 (m, 7H), 4.01 (s, 2H), 3.67 (p, J = 8.2 Hz, 1H), 2.94 (t, J = 5.4 Hz, 2H), 2.87 (d, J = 6.2 Hz, 2H), 1.64 (q, J = 5.9 Hz, 4H). $^{13}$C NMR (101 MHz, DMSO-$d_6$) δ 140.20, 138.05, 136.55, 133.26, 132.98, 131.65, 131.31, 131.12, 129.11, 128.99, 128.31, 128.01, 125.74, 123.06, 123.02, 46.66, 46.26, 43.03, 42.80, 37.25, 34.72, 34.30, 31.28. MS-ESI m/z [M + H]+: calcd 573.56, found 571.95. HRMS m/z [M + H]+: calcd 572.0922, found 572.1077.

**Synthesis of 5-((2-(((2-chloro-[1,1'-biphenyl]-4-yl)methyl)amino)-7-azaspiro[3.5]nonan-7-yl)sulfonyl)-1,3-dimethyl-1,3-dihydro-2*H*-benzo[*d*]imidazol-2-one 57**

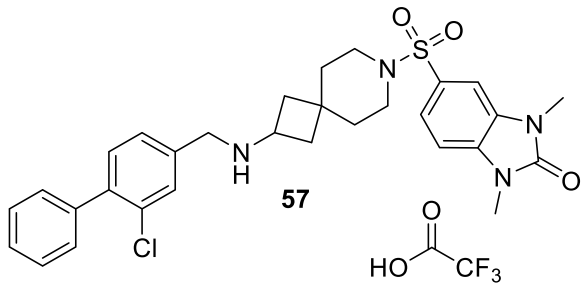

**57**

**23** (30 mg, 0.07 mmol, 1 eq.) was deprotected using **General Procedure B**. The crude product was reacted with 1,3-dimethyl-2-oxo-2,3-dihydro-1H-benzo[*d*]imidazole-5-sulfonyl chloride (17 mg, 0.07 mmol, 1 eq.) according to **General Procedure A**. The product was not completely pure after the first round of chromatography and product was further subjected to reverse phase flash chromatography (H2O/ACN as eluents with addition of 0.2% of TFA) yielding the title compound as a TFA salt (34 mg, 90%). $^{1}$H NMR (400 MHz, DMSO-$d_6$) δ 9.17 (s, 2H), 7.68 (d, J = 1.5 Hz, 1H), 7.52 – 7.41 (m, 6H), 7.45 – 7.36 (m, 3H), 7.37 (d, J = 8.2 Hz, 1H), 4.00 (s, 2H), 3.64 (p, J = 8.0 Hz, 1H), 3.41 (s, 3H), 2.88 (t, J = 5.5 Hz, 2H), 2.81 (t, J = 5.6 Hz, 2H), 2.00 (td, J = 9.3, 8.2, 5.3 Hz, 2H), 1.90 – 1.81 (m, 2H), 1.63 (dt, J = 10.6, 5.0 Hz, 4H). $^{13}$C NMR (101 MHz, DMSO-$d_6$) δ 154.07, 140.07, 138.09, 133.81, 133.18, 131.63, 131.30, 130.98, 129.79, 129.12, 128.85, 128.30, 127.98, 127.64, 121.13, 107.62, 106.76, 46.82, 46.34, 43.07, 42.82, 37.37, 35.02, 34.40, 31.27, 27.29, 27.26.MS-ESI m/z [M + H]+: calcd 566.13, found 565.02. HRMS m/z [M + H]+: calcd 565.1962, found 565.2015.

**Synthesis of 5-(5-((2-(((2-chloro-[1,1'-biphenyl]-4-yl)methyl)amino)-7-azaspiro[3.5]nonan-7-yl)sulfonyl)-2-ethoxyphenyl)-1-methyl-3-propyl-1,6-dihydro-7*H*-pyrazolo[4,3-*d*]pyrimidin-7-one TFA salt 58**

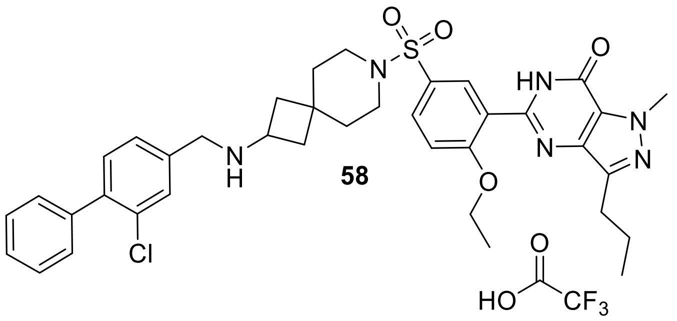

**23** (30 mg, 0.07 mmol, 1 eq.) was deprotected using **General Procedure B**. The crude product was reacted with 4-ethoxy-3-(1-methyl-7-oxo-3-propyl-6,7-dihydro-1H-pyrazolo[4,3-d]pyrimidine-5-yl)benzenesulfonyl chloride (28 mg, 0.07 mmol, 1 eq.) according to **General Procedure A**. The product was not completely pure after the first round of chromatography and product was further subjected to reverse phase flash chromatography ($H_2O$/ACN as eluents with addition of 0.2% of TFA) yielding the title compound as a TFA salt (39 mg, 81%). $^1$H NMR (400 MHz, DMSO-$d_6$) δ 12.21 (s, 1H), 9.47 (s, 1H), 7.90 (d, J = 2.6 Hz, 1H), 7.84 (dd, J = 8.8, 2.5 Hz, 1H), 7.72 (d, J = 1.6 Hz, 1H), 7.55 – 7.33 (m, 8H), 7.30 – 7.08 (m, 1H), 4.21 (q, J = 6.9 Hz, 2H), 4.16 (s, 3H), 4.05 (s, 2H), 3.73 (q, J = 8.2 Hz, 1H), 2.97 – 2.92 (m, 2H), 2.87 (d, J = 5.8 Hz, 2H), 2.78 (t, J = 7.4 Hz, 2H), 2.11 – 1.99 (m, 2H), 1.94 (dd, J = 12.5, 8.2 Hz, 2H), 1.84 – 1.73 (m, 2H), 1.65 (dt, J = 10.8, 5.4 Hz, 4H), 1.34 (t, J = 6.9 Hz, 3H), 0.94 (t, J = 7.4 Hz, 3H). $^{13}$C NMR (101 MHz, DMSO-$d_6$) δ 159.77, 148.16, 144.96, 140.26, 138.06, 137.80, 133.15, 131.67, 131.36, 131.27, 131.16, 129.84, 129.12, 129.00, 128.89, 128.31, 128.19, 128.00, 127.29, 124.41, 113.26, 64.86, 46.26, 42.96, 42.69, 37.86, 37.25, 34.72, 34.33, 31.34, 27.17, 21.66, 14.27, 13.83. MS-ESI m/z [M + H]+: calcd 716.33, found 715.20. HRMS m/z [M + H]+: calcd 715.2755, found 715.2828.

**Synthesis of *N*-((2-chloro-[1,1'-biphenyl]-4-yl)methyl)-6-(isoquinolin-8-ylsulfonyl)-6-aza-spiro[3.4]octan-2-amine 42**

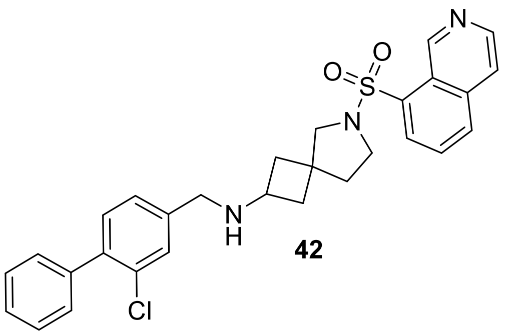

**42**

**24** (30 mg, 0.07 mmol, 1 eq.) was deprotected using **General Procedure B**. The crude product was reacted with 5-Isoquinolinsulfonylchloride (19 mg, 0.07 mmol, 1 eq.) according to **General Procedure A** (30 mg, 83%). $^{1}$H NMR (500 MHz, DMSO-$d_6$) δ 9.48 (s, 1H), 8.71 (d, J = 6.1 Hz, 1H), 8.44 (t, J = 6.1 Hz, 2H), 8.33 (d, J = 7.3 Hz, 1H), 7.83 (t, J = 7.7 Hz, 1H), 7.60 (s, 1H), 7.45 (t, J = 7.4 Hz, 2H), 7.39 (d, J = 8.0 Hz, 4H), 7.27 – 7.08 (m, 1H), 3.53 (p, J = 6.7, 6.0 Hz, 1H), 2.29 (s, 1H), 2.20 – 2.12 (m, 1H), 1.98 (p, J = 5.8 Hz, 2H), 1.93 – 1.86 (m, 2H), 1.82 (p, J = 6.0 Hz, 1H), 1.68 (dt, J = 15.5, 9.8 Hz, 2H), 1.29 – 1.17 (m, 1H). $^{13}$C NMR (126 MHz, DMSO-$d_6$) δ 153.41, 144.56, 139.31, 135.47, 133.48, 132.49, 131.41, 131.12, 130.35, 129.15, 128.61, 128.26, 127.84, 126.53, 117.20, 47.62, 47.34, 43.21, 42.26, 42.03, 30.34. MS-ESI m/z [M + H]+: calcd 519.07, found 518.10. HRMS m/z [M + H]+: calcd 518.1591, found 518.1650

**Synthesis of 6-(benzo[*d*]thiazol-6-ylsulfonyl)-*N*-((2-chloro-[1,1'-biphenyl]-4-yl)methyl)-6-azaspiro[3.4]octan-2-amine TFA salt 43**

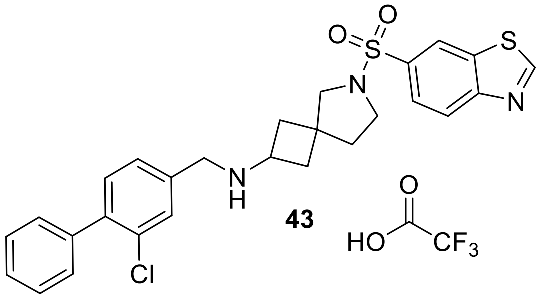

**43**

**24** (30 mg, 0.07 mmol, 1 eq.) was deprotected using **General Procedure B**. The crude product was reacted with benzo[*d*]thiazole-6-sulfonyl chloride (16 mg, 0.07 mmol, 1 eq.) according to **General Procedure A**. The product was not completely pure after the first round of chromatography and product was further subjected to reverse phase flash chromatography (H2O/ACN as eluents with addition of 0.2% of TFA) yielding the title compound as a TFA salt (29 mg, 83%). $^{1}$H NMR (400 MHz, DMSO-$d_6$) δ 9.62 (s, 1H), 8.69 (d, J = 1.8 Hz, 1H), 8.27 (d, J = 8.6 Hz, 1H), 8.06 (d, J = 8.5 Hz, 1H), 7.92 (dd, J = 8.6, 1.9 Hz, 1H), 7.64 (d, J = 1.5 Hz, 1H), 7.48 – 7.38 (m, 8H), 3.94 (s, 2H), 3.60 (h, J = 8.2 Hz, 2H), 3.45 (p, J = 8.0 Hz, 2H), 3.24 – 2.99 (m, 1H), 2.26 (ddd, J = 11.6, 7.6, 3.9 Hz, 1H), 2.09 (ddt, J = 24.7, 12.3, 4.6 Hz, 4H), 1.96 (ddd, J = 16.6, 8.3, 4.6 Hz, 2H), 1.76 (dt, J = 11.4, 7.8 Hz, 3H). $^{13}$C NMR (101 MHz, DMSO-$d_6$) δ 160.59, 155.04, 139.86, 138.34, 138.16, 134.57, 134.08, 131.58, 131.26, 130.79, 129.15, 128.67, 128.30, 127.96, 124.26, 123.65, 122.16, 47.12, 47.08, 43.32, 42.19, 42.03, 38.31, 37.74, 30.55. MS-ESI m/z [M + H]+: calcd 525.09, found 524.10. HRMS m/z [M + H]+: calcd 524.1155, found 524.1290.

**Synthesis of *N*-((2-chloro-[1,1'-biphenyl]-4-yl)methyl)-6-((2-methylbenzo[*d*]thiazol-6-yl)-sulfonyl)-6-azaspiro[3.4]octan-2-amine TFA salt 44**

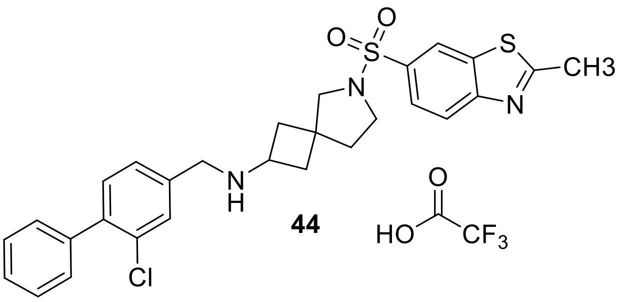

**44**

**24** (30 mg, 0.07 mmol, 1 eq.) was deprotected using **General Procedure B**. The crude product was reacted with 2-methylbenzo[*d*]thiazole-6-sulfonyl chloride (17 mg, 0.07 mmol, 1 eq.) according to **General Procedure A.** The product was not completely pure after the first round of chromatography and product was further subjected to reverse phase flash chromatography ($H_2O$/ACN as eluents with addition of 0.2% of TFA) yielding the title compound as a TFA salt (20 mg, 53%). $^1$H NMR (400 MHz, DMSO-$d_6$) δ 9.19 (s, 1H), 8.54 (d, J = 1.9 Hz, 1H), 8.08 (d, J = 8.6 Hz, 1H), 8.00 (d, J = 8.6 Hz, 1H), 7.85 (dd, J = 8.6, 1.9 Hz, 1H), 7.68 (d, J = 1.5 Hz, 1H), 7.51 – 7.37 (m, 7H), 3.99 (s, 2H), 3.67 – 3.57 (m, 1H), 3.57 – 3.51 (m, 1H), 3.09 (q, J = 7.3 Hz, 1H), 2.85 (s, 3H), 2.32 – 2.23 (m, 1H), 2.16 – 2.04 (m, 4H), 1.96 (ddd, J = 12.1, 7.4, 5.2 Hz, 1H), 1.76 (ddd, J = 11.6, 8.6, 6.2 Hz, 2H). $^{13}$C NMR (101 MHz, DMSO-$d_6$) δ 171.77, 171.77, 140.16, 138.09, 137.52, 135.56, 133.44, 131.65, 131.32, 131.05, 129.15, 128.91, 128.32, 128.01, 124.21, 122.42, 121.53, 46.90, 46.83, 43.26, 42.10, 41.99, 37.79, 37.19, 30.58, 20.03.MS-ESI m/z [M + H]+: calcd 539.12, found 538.10. HRMS m/z [M + H]+: calcd 538.1312, found 538.1372.

**Synthesis of *N*-((2-chloro-[1,1'-biphenyl]-4-yl)methyl)-6-((2-chlorobenzo[*d*]thiazol-6-yl)-sulfonyl)-6-azaspiro[3.4]octan-2-amine TFA salt 45**

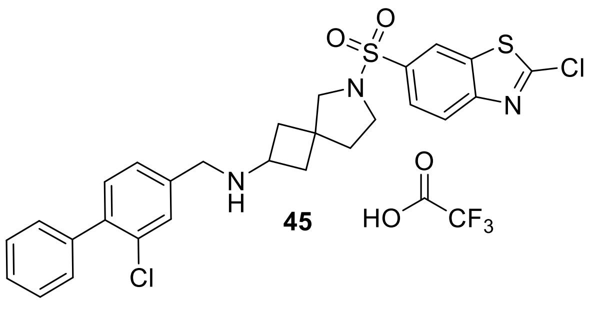

**45**

**24** (30 mg, 0.07 mmol, 1 eq.) was deprotected using **General Procedure B**. The crude product was reacted with 2-chlorobenzo[*d*]thiazole-6-sulfonyl chloride (19 mg, 0.07 mmol, 1 eq.) according to **General Procedure A**. The product was not completely pure after the first round of chromatography and product was further subjected to reverse phase flash chromatography ($H_2O$/ACN as eluents with addition of 0.2% of TFA) yielding the title compound as a TFA salt (38 mg, 86%). $^1$H NMR (400 MHz, DMSO-$d_6$) δ 8.65 (d, J = 1.9 Hz, 1H), 8.15 (d, J = 8.6 Hz, 1H), 8.09 (d, J = 8.5 Hz, 1H), 7.92 (dd, J = 8.6, 1.9 Hz, 1H), 7.66 (d, J = 1.4 Hz, 1H), 7.50 – 7.37 (m, 7H), 3.96 (s, 2H), 3.60 (p, J = 8.1 Hz, 1H), 3.49 – 3.44 (m, 1H), 3.09 (q, J = 7.3 Hz, 1H), 2.27 (ddd, J = 11.3, 7.6, 3.4 Hz, 1H), 2.13 – 2.04 (m, 3H), 1.99 (dt, J = 11.5, 4.7 Hz, 1H), 1.77 (ddd, J = 11.5, 8.7, 5.4 Hz, 2H). $^{13}$C NMR (101 MHz, DMSO-$d_6$) δ 157.16, 152.54, 139.96, 138.75, 138.12, 136.13, 134.20, 131.61, 131.28, 130.88, 129.15, 128.75, 128.31, 127.98, 125.04, 123.06, 122.05, 47.02, 43.30, 42.14, 41.98, 38.14, 37.56, 30.56. MS-ESI m/z [M + H]+: calcd 559.54, found 558.05. HRMS m/z [M + H]+: calcd 558.0765, found 558.0847.

**Synthesis of 5-((2-(((2-chloro-[1,1'-biphenyl]-4-yl)methyl)amino)-6-azaspiro[3.4]octan-6-yl)sulfonyl)-1,3-dimethyl-1,3-dihydro-2*H*-benzo[*d*]imidazol-2-one 46**

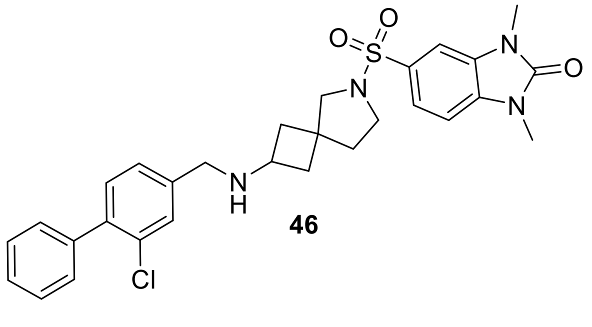

**46**

**24** (30 mg, 0.07 mmol, 1 eq.) was deprotected using **General Procedure B**. The crude product was reacted with 1,3-dimethyl-2-oxo-2,3-dihydro-1H-benzo[*d*]imidazole-5-sulfonyl chloride (18 mg, 0.07 mmol, 1 eq.) according to **General Procedure A** (38 mg, 86%). $^1$H NMR (400 MHz, DMSO-$d_6$) δ 8.89 (s, 1H), 7.75 (d, J = 8.6 Hz, 1H), 7.66 (d, J = 1.1 Hz, 1H), 7.54 – 7.50 (m, 2H), 7.49 – 7.44 (m, 4H), 7.42 – 7.39 (m, 2H), 7.32 (d, J = 8.7 Hz, 1H), 3.98 (s, 2H), 3.51 (dq, J = 19.7,

7.9 Hz, 3H), 3.39 (s, 3H), 3.37 (s, 3H), 2.28 (ddd, J = 11.5, 7.6, 3.7 Hz, 1H), 2.09 (tdd, J = 12.1, 7.5, 4.7 Hz, 4H), 1.96 (ddd, J = 12.0, 7.3, 5.1 Hz, 1H), 1.76 (ddd, J = 11.3, 8.6, 2.0 Hz, 2H). $^{13}$C NMR (101 MHz, DMSO-$d_6$) δ 140.06, 138.08, 133.84, 132.65, 131.65, 131.29, 130.93, 129.55, 129.14, 128.80, 128.32, 128.01, 120.16, 107.48, 105.93, 47.00, 43.25, 42.16, 42.05, 38.05, 37.43, 30.58, 27.25, 27.19. MS-ESI m/z [M + H]+: calcd 552.10, found 551.15. HRMS m/z [M + H]+: calcd 551.1805, found 551.1865.

**Synthesis of 6-((2-(((2-chloro-[1,1'-biphenyl]-4-yl)methyl)amino)-6-azaspiro[3.4]octan-6-yl)sulfonyl)benzo[*d*]oxazol-2(3*H*)-one 47**

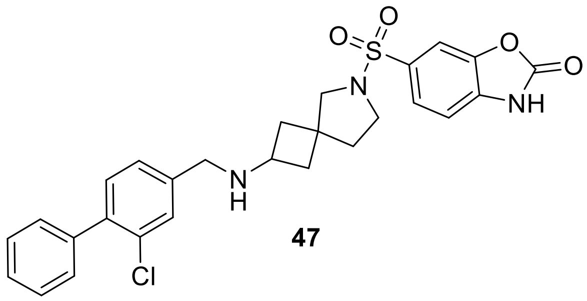

**47**

**24** (30 mg, 0.07 mmol, 1 eq.) was deprotected using **General Procedure B**. The crude product was reacted with 2-oxo-2,3-dihydrobenzo[*d*]oxazole-6-sulfonyl chloride (16 mg, 0.07 mmol, 1 eq.) according to **General Procedure A** (38 mg, 86%). $^{1}$H NMR (500 MHz, DMSO-$d_6$) δ 7.82 (d, J = 8.5 Hz, 1H), 7.59 (t, J = 9.1 Hz, 3H), 7.50 – 7.45 (m, 2H), 7.40 (d, J = 7.4 Hz, 4H), 7.24 (d, J = 7.9 Hz, 1H), 7.19 – 7.09 (m, 1H), 3.82 (s, 2H), 3.53 (d, J = 8.1 Hz, 1H), 3.27 (s, 2H), 2.28 – 2.19 (m, 1H), 2.07 (ddt, J = 18.2, 11.9, 6.3 Hz, 2H), 1.97 – 1.84 (m, 3H), 1.77 – 1.68 (m, 2H). $^{13}$C NMR (126 MHz, DMSO-$d_6$) δ 154.59, 143.00, 139.24, 138.29, 134.63, 134.59, 131.44, 131.13, 130.18, 129.15, 128.90, 128.27, 128.12, 127.85, 123.04, 109.78, 107.65, 47.80, 47.50, 43.33, 42.34, 42.09, 30.48. MS-ESI m/z [M + H]+: calcd 525.03, found 524.10. HRMS m/z [M + H]+: calcd 524.1333, found 524.1394.

**Synthesis of Methyl *N*-(benzo[*d*]thiazol-6-ylsulfonyl)-*N*-(5-(((2-chloro-[1,1'-biphenyl]-4-yl)methyl)amino)pentyl)glycinate 73**

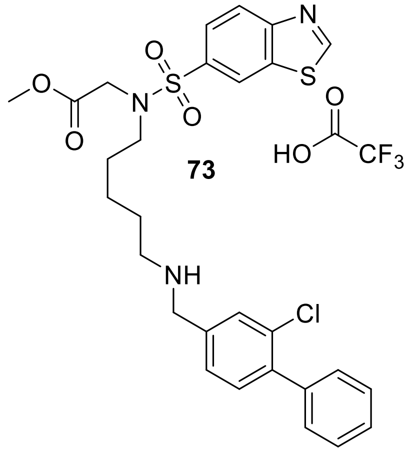

To a solution of **68** (50 mg, 0.09 mmol, 1 eq.) in DCM (1 mL) was added TFA (1 mL) at 0 °C. The reaction mixture was allowed to warm to room temperature and stirred for 1 hour. The reaction mixture was concentrated, co-evaporate two times with Toluol, quenching with triethylamine (0.5 mL) and again co-evaporated two times with toluol. The residue was dried under vacuum for 16 hours. The crude amine was solved in DCM (3 mL), **18** (21 mg, 0.09 mmol, 1 eq.) and $MgSO_4$ was added and stirred for 10 min. Sodium triacetoxyborohydride (42 mg, 0.2 mmol, 2 eq.) was added and stirred overnight. The mixture was concentrated and the residue was solved DCM. The organic phase was washed with water 2x, brine 2x, dried over $MgSO_4$ and the solvent was removed under reduced pressure. The residue was purified by rp-flash chromatography ($H_2O$/ACN). The product was not completely pure after the first round of chromatography and product was further subjected to reverse phase flash chromatography ($H_2O$/ACN as eluents with addition of 0.2% of TFA) yielding the title compound as a TFA salt (24 mg, 42%). $^1H$ NMR (400 MHz, DMSO-$d_6$) δ = 9.63 (s, 1H), 8.78 (d, J = 1.9 Hz, 1H), 8.25 (d, J = 8.6 Hz, 1H), 7.93 (dd, J = 8.7, 1.9 Hz, 1H), 7.54 (d, J = 1.3 Hz, 1H), 7.50 – 7.38 (m, 8H), 7.36 (d, J = 1.4 Hz, 2H), 4.12 (s, 2H), 3.76 (s, 2H), 3.54 (s, 3H), 3.19 (t, J = 7.3 Hz, 2H), 1.43 (dp, J = 23.3, 7.4 Hz, 6H), 1.30 – 1.14 (m, 4H). $^{13}C$ NMR (101 MHz, DMSO-$d_6$) δ = 169.46, 160.91, 155.23 , 138.60, 136.15, 134.25, 131.21, 131.00, 129.20, 128.20 C-29), 127.64, 127.25, 124.56,

123.56, 122.73, 51.83, 48.55, 48.52, 48.26, 48.17, 27.34, 23.57. MS-ESI m/z [M + H]+: calcd 573.14, found 572.20. HRMS m/z [M + H]+: calcd 572.1366 , found 572.1439.

**Synthesis of Methyl 3-(3-(3-(((2-chloro-[1,1'-biphenyl]-4-yl)methyl)amino)propanamido)-propanamido)benzoate TFA salt 14**

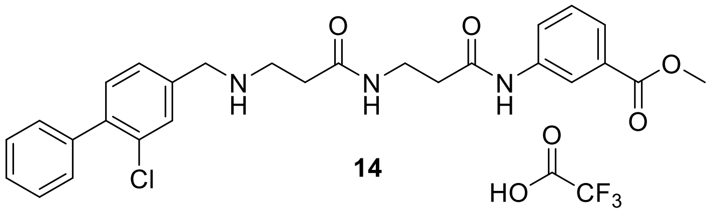

**14** was synthesized according to.[14] Additionally, purification was performed using reverse phase flash chromatography ($H_2O$/ACN as eluents with addition of 0.2% of TFA) yielding the title compound as a TFA salt (32 mg, 30% over 5 steps). $^1$H NMR (400 MHz, DMSO-$d_6$) δ 10.22 (s, 1H), 8.96 (s, 2H), 8.34 – 8.20 (m, 2H), 7.81 (ddd, J = 8.2, 2.3, 1.1 Hz, 1H), 7.74 (d, J = 1.6 Hz, 1H), 7.63 (dt, J = 7.8, 1.3 Hz, 1H), 7.53 – 7.42 (m, 8H), 4.23 (t, J = 5.5 Hz, 2H), 3.84 (s, 3H), 3.39 (q, J = 6.4 Hz, 2H), 3.16 (p, J = 6.9 Hz, 2H), 2.58 – 2.51 (m, 4H). $^{13}$C NMR (101 MHz, DMSO-$d_6$) δ 170.22, 169.56, 166.59, 140.80, 140.01, 138.53, 133.56, 132.21, 131.84, 131.75, 130.54, 129.66, 129.62, 129.60, 128.81, 128.52, 124.20, 123.97, 120.06, 52.66, 49.50, 43.32, 36.60, 35.50, 31.40. MS-ESI m/z [M + H]+: calcd 494.99, found 494.10.

**Synthesis of 3-(3-(3-(((2-chloro-[1,1'-biphenyl]-4-yl)methyl)amino)propanamido)propan-amido)benzoic acid HCl salt 15**

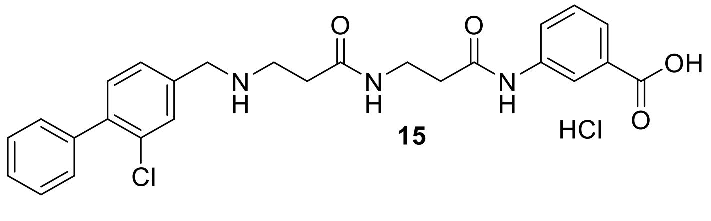

**15** was synthesized according to.[14] Additionally, purification was performed by precipitating the product as HCl salt with 4N HCl in dioxane and rinsing the product with DCM and diethyl ether (20 mg, 43% over 6 steps). $^1$H NMR (400 MHz, DMSO-$d_6$) δ 10.30 (q, J = 3.3 Hz, 1H), 8.34 (d, J = 6.8 Hz, 1H), 8.28 (q, J = 1.8 Hz, 1H), 7.85 – 7.80 (m, 2H), 7.60 (ddd, J = 8.0, 3.7, 2.2 Hz, 2H),

7.49 – 7.39 (m, 7H), 4.17 (s, 2H), 3.37 (q, J = 6.4 Hz, 2H), 3.09 (t, J = 7.3 Hz, 2H), 2.61 (td, J = 6.8, 3.7 Hz, 2H), 2.54 (t, J = 6.7 Hz, 2H). $^{13}C$ NMR (101 MHz, DMSO-$d_6$) δ 169.74, 169.19, 167.23, 140.07, 139.44, 138.16, 133.72, 131.60, 131.27, 129.17, 129.12, 128.87, 128.31, 127.98, 123.89, 123.16, 119.91, 49.01, 42.85, 36.21, 35.13, 31.21. MS-ESI m/z [M + H]+: calcd 480.96, found 480.10.

### Analytical data of compounds

#### Cmpd 25

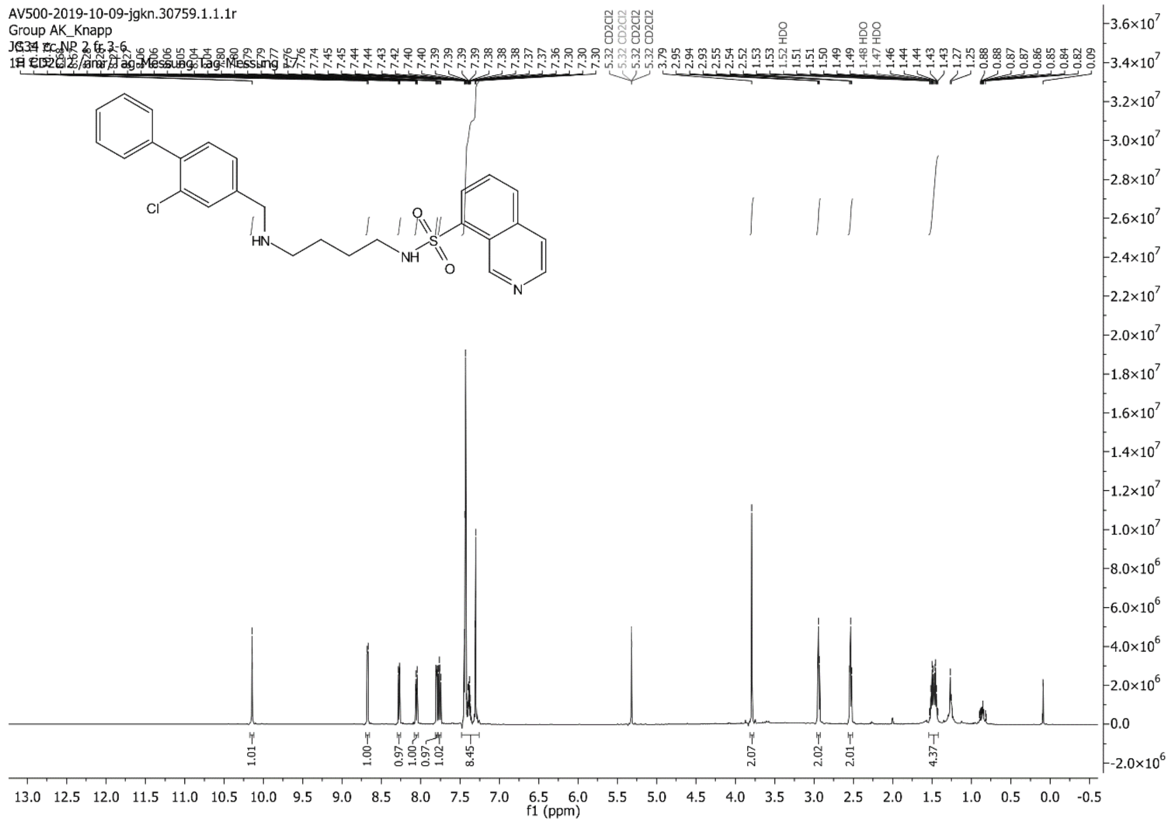

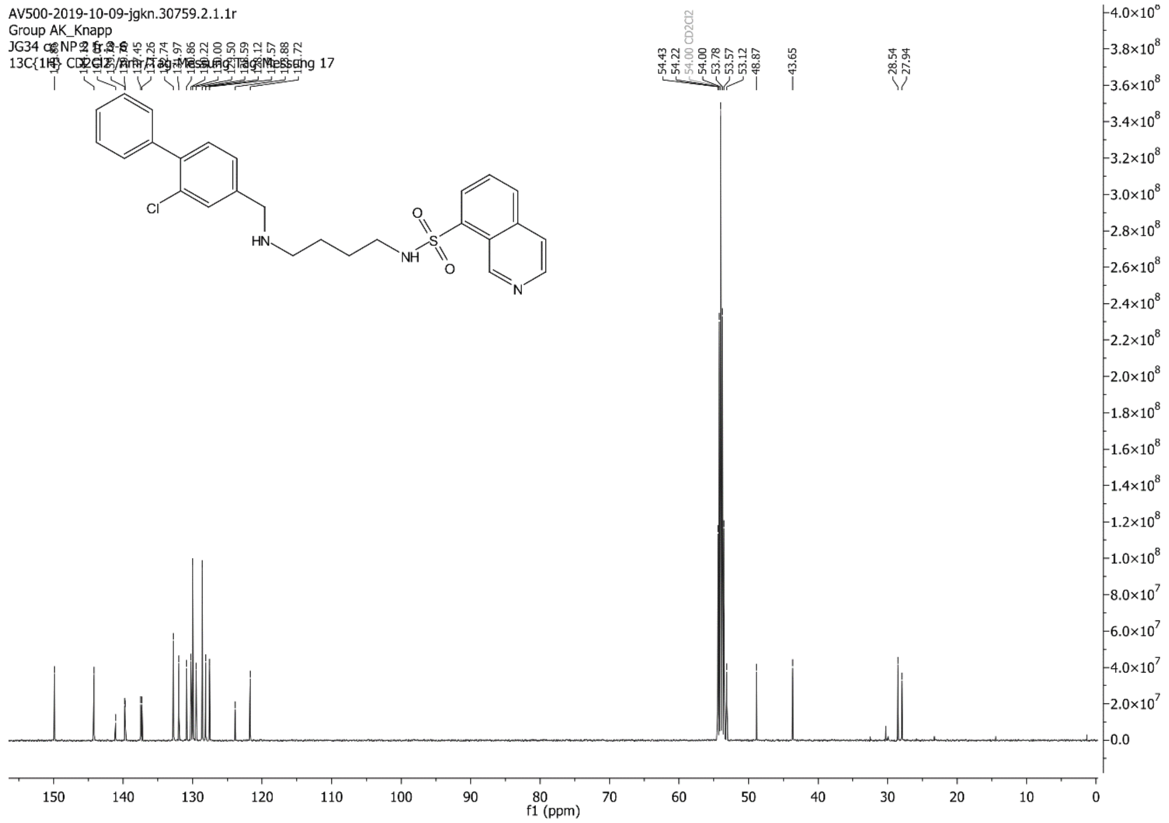

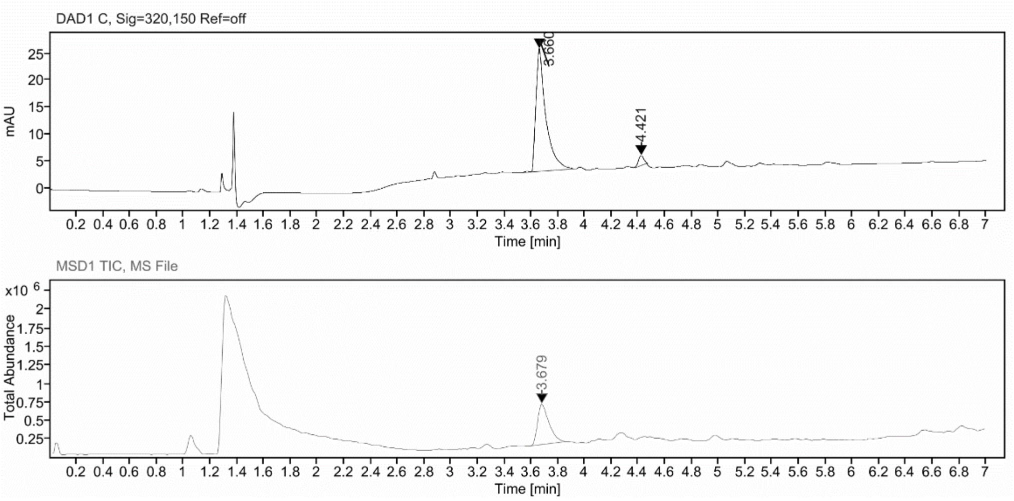

**Sample Purity**

Signal Description DAD1 C, Sig=320,150 Ref=off

| Sample Name | Name | RT | Width | Area | Area% | Height |
|---|---|---|---|---|---|---|
| FGJG12_purity | | 3.660 | 0.072 | 118.6595 | 95.80 | 22.9620 |
| FGJG12_purity | | 4.421 | 0.047 | 5.2007 | 4.20 | 1.8983 |

Max Area% 95.801
UV Signal Purity>95% Pass

**UV Apex Spectra**

**RT:**

3.660

**Sample Name:**

FGJG12_purity

**Signal Name:**

DAD1C

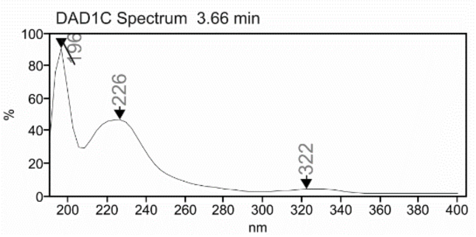

**MSD Apex Spectra**

**RT:**

3.679

**Sample Name:**

FGJG12_purity

**Signal Name:**

MSD1TIC

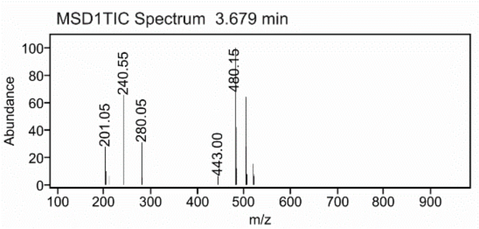

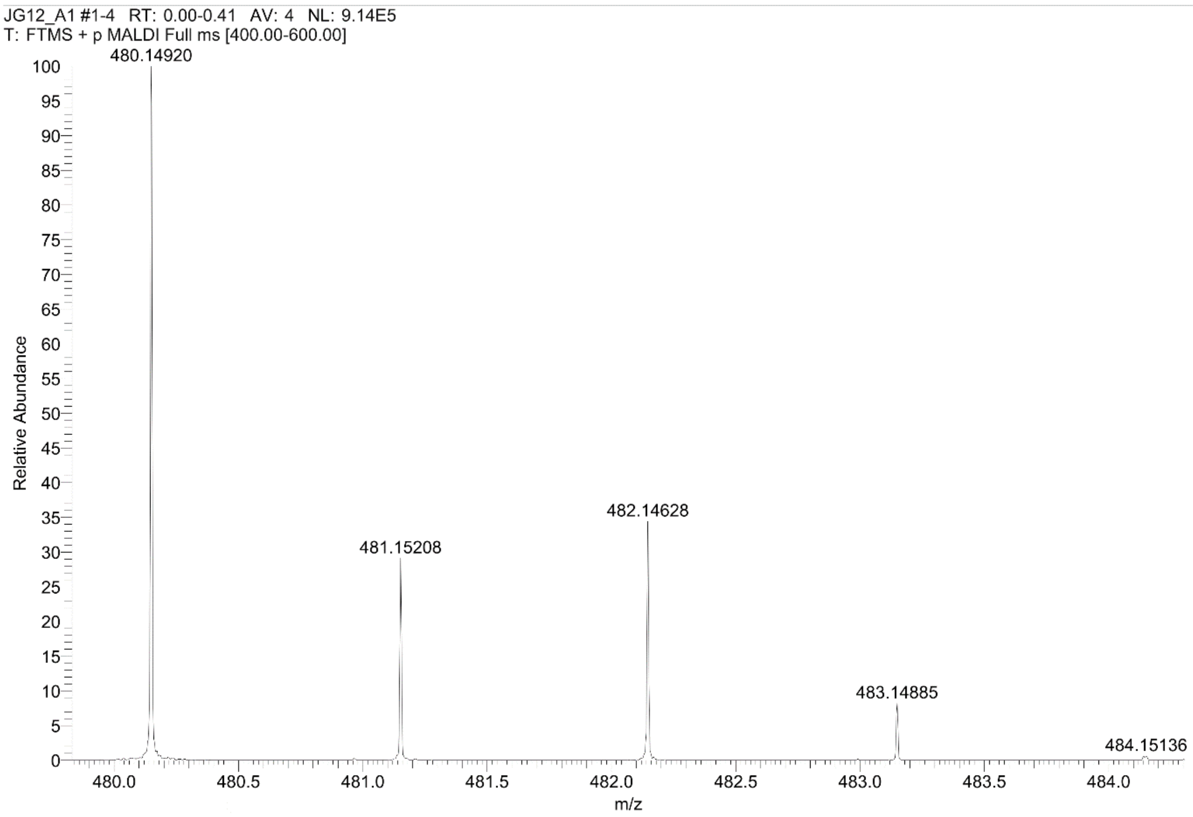
JG12_A1 #1-4 RT: 0.00-0.41 AV: 4 NL: 9.14E5
T: FTMS + p MALDI Full ms [400.00-600.00]
480.14920
481.15208
482.14628
483.14885
484.15136
Relative Abundance
m/z

#### Cmpd 26

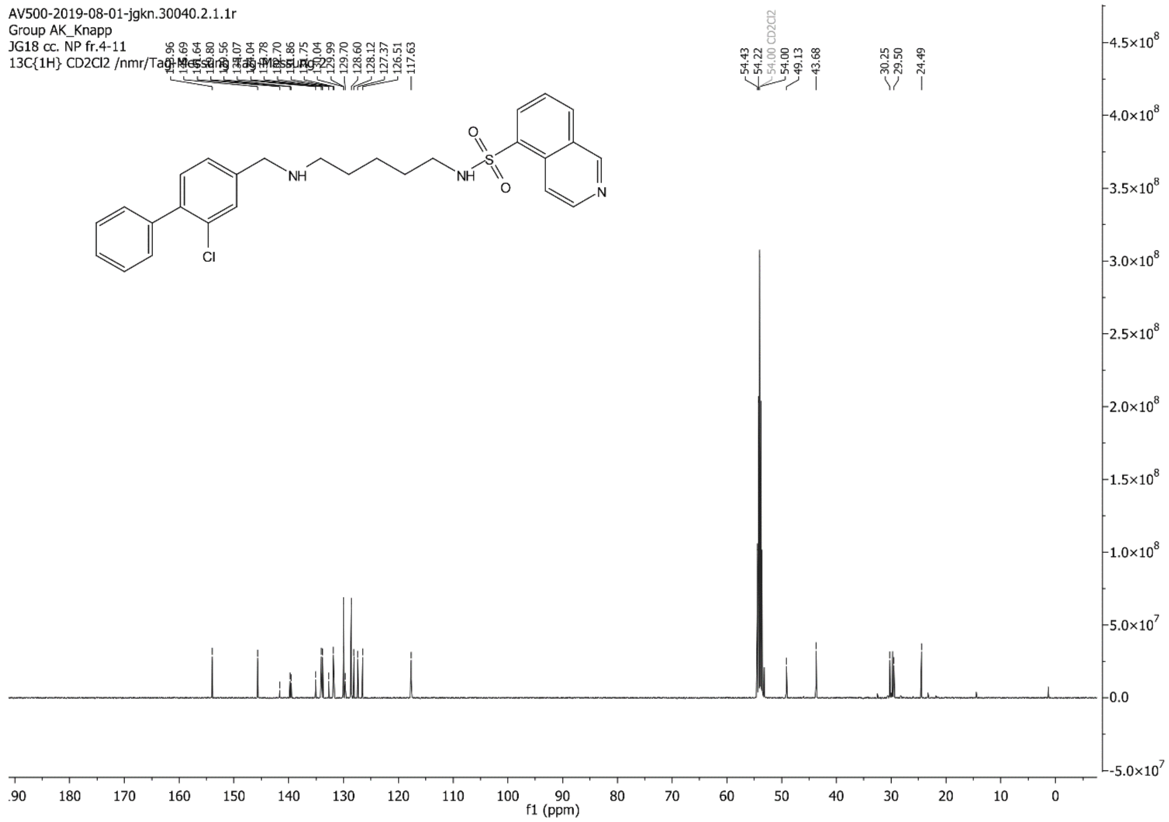

AV500-2019-08-01-jgkn.30040.2.1.1r
Group AK_Knapp
JG18 cc. NP fr.4-11
13C{1H} CD2Cl2 /nmr/Tag-Messung Tag-Messung 2
129.70
128.60
128.12
127.37
126.51
117.63
54.43
54.22
54.00 CD2Cl2
54.00
49.13
43.68
30.25
29.50
24.49
NH
NH
S
O
O
N
Cl
f1 (ppm)

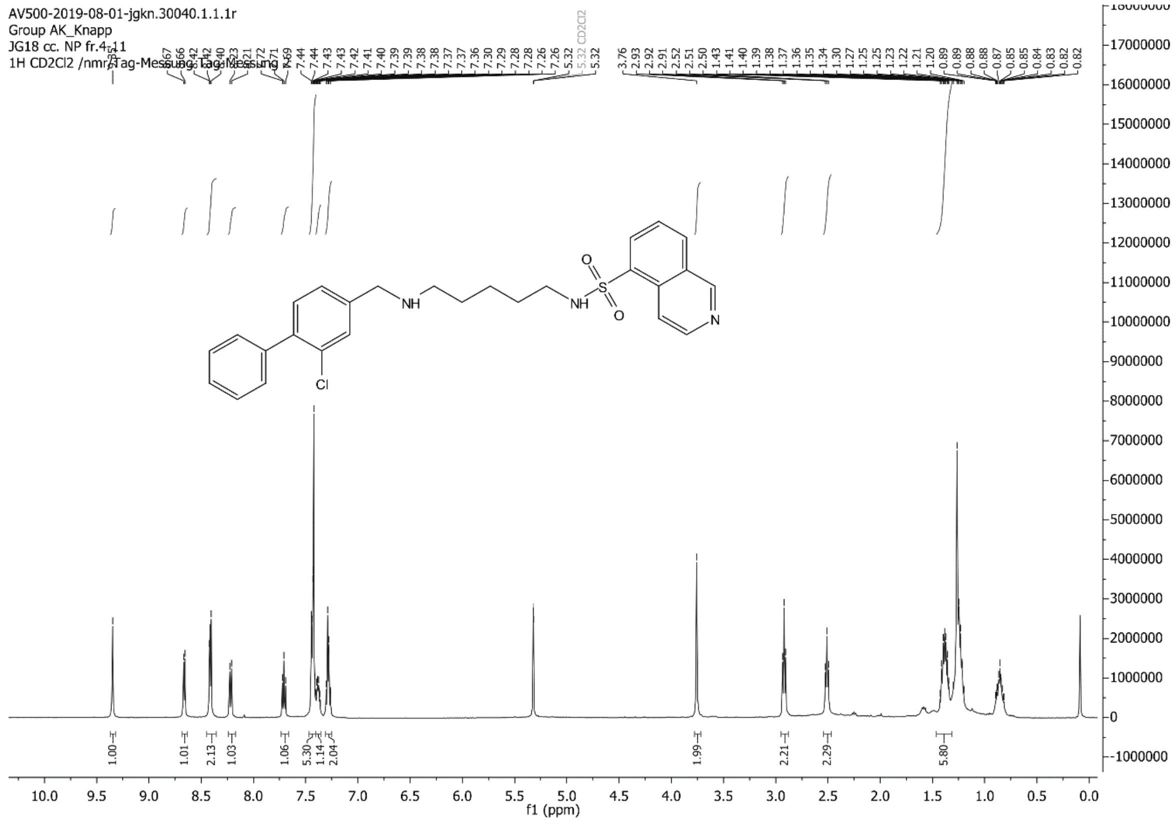

AV500-2019-08-01-jgkn.30040.1.1.1r
Group AK_Knapp
JG18 cc. NP fr.4-11
1H CD2Cl2 /nmr/Tag-Messung Tag-Messung
5.32 CD2Cl2
NH
NH
S
O
O
N
Cl
1.00
1.01
2.13
1.03
1.06
5.30
1.14
2.04
1.99
2.21
2.29
5.80
f1 (ppm)

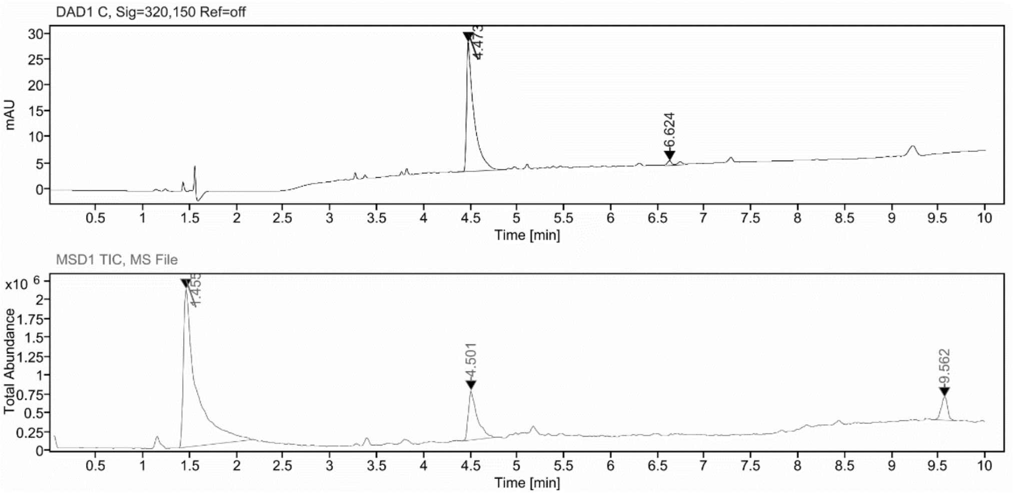

#### Sample Purity

Signal Description DAD1 C, Sig=320,150 Ref=off

| Sample Name | Name | RT | Width | Area | Area% | Height |
|---|---|---|---|---|---|---|
| FGJG18 | | 4.473 | 0.071 | 136.1407 | 96.33 | 24.9186 |
| FGJG18 | | 6.624 | 0.050 | 5.1869 | 3.67 | 0.9126 |

Max Area% 96.330
UV Signal Purity>95% Pass

#### UV Apex Spectra

**RT:**

4.473

**Sample Name:**

FGJG18

**Signal Name:**

DAD1C

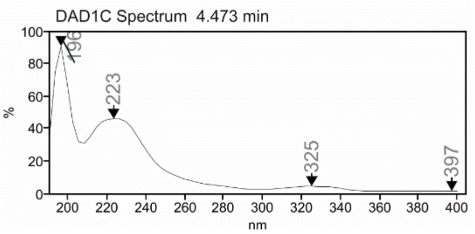

**RT:**

4.501

**Sample Name:**

FGJG18

**Signal Name:**

MSD1TIC

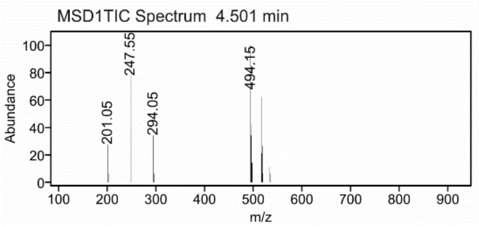

C:\User\...\Knapp\2019\190805\JG18_A2 8/5/2019 12:32:35 PM JG18 mit HCCA gemessen.

JG18_A2 #1-6 RT: 0.00-0.64 AV: 6 NL: 5.19E6
T: FTMS + p MALDI Full ms [400.00-600.00]

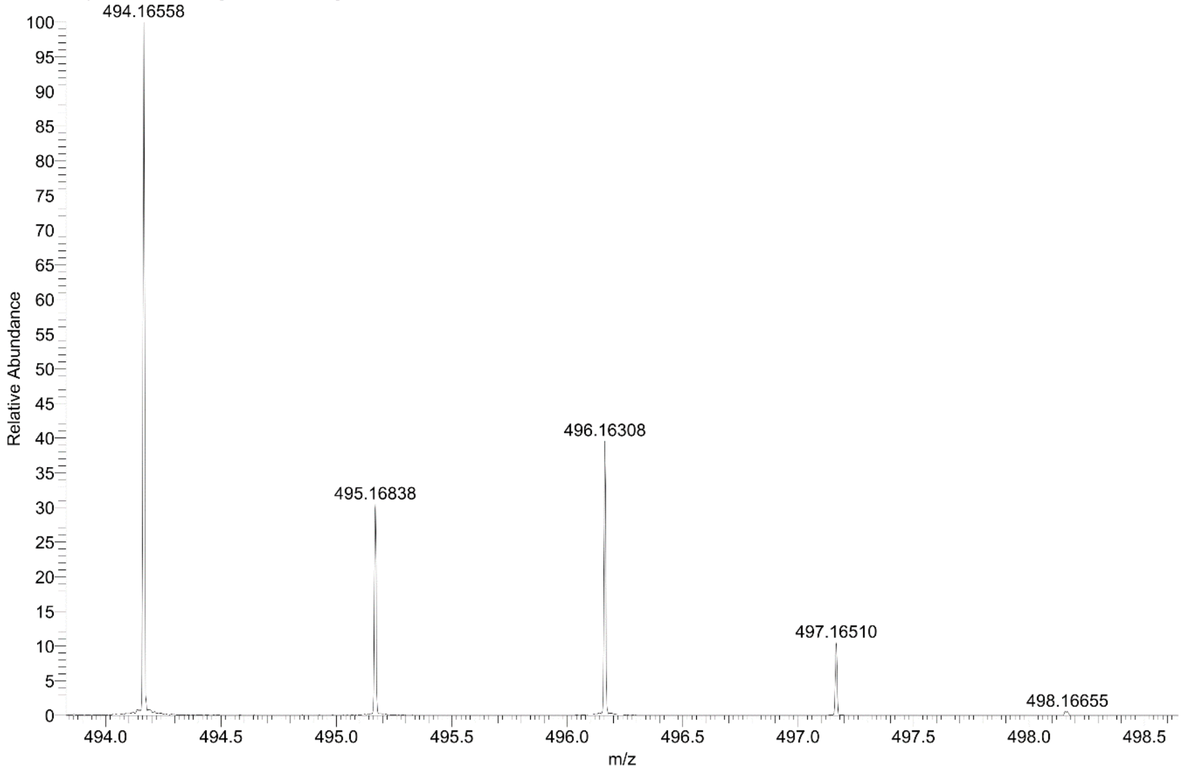

#### Cmpd 27

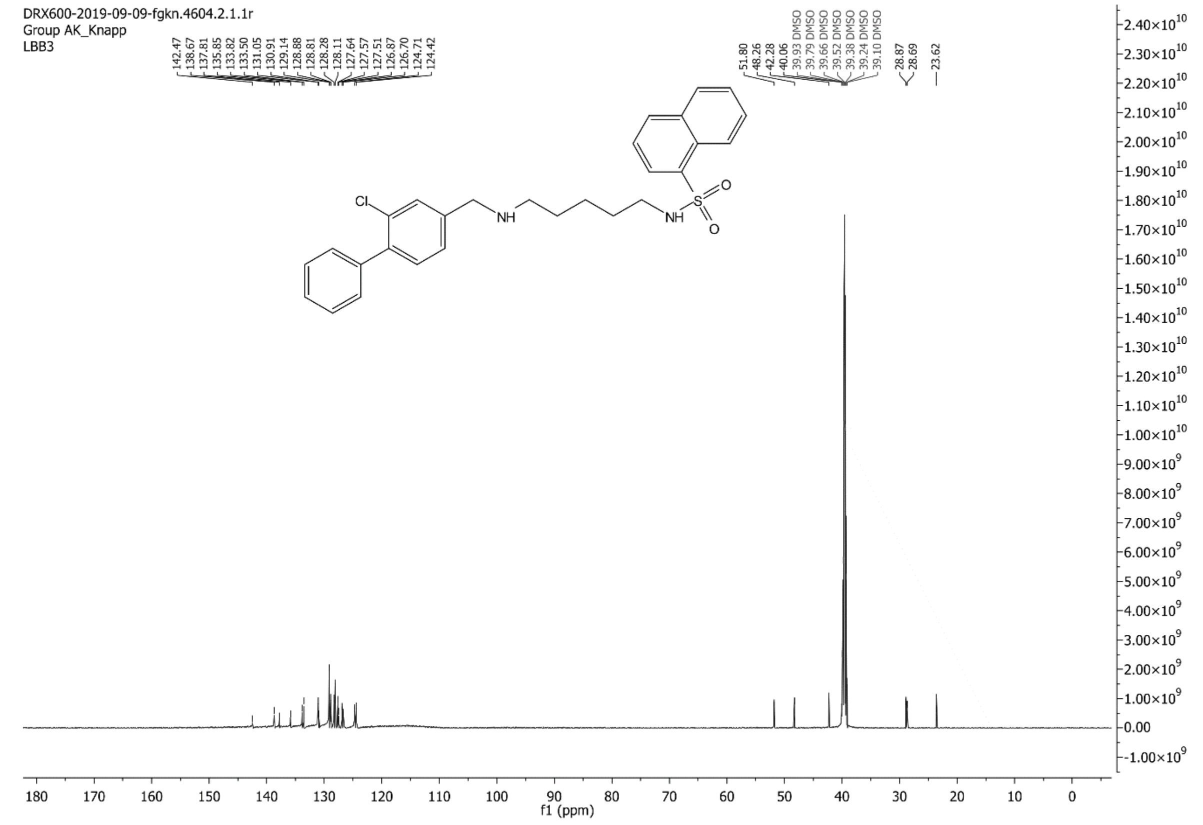
DRX600-2019-09-09-fgkn.4604.2.1.1r
Group AK_Knapp
LBB3
142.47 138.67 137.81 135.85 133.82 133.50 131.05 130.91 129.14 128.88 128.81 128.28 128.11 127.64 127.57 127.51 126.87 126.70 124.71 124.42
51.80 48.26 42.28 40.06 39.93 DMSO 39.79 DMSO 39.66 DMSO 39.52 DMSO 39.38 DMSO 39.24 DMSO 39.10 DMSO
28.87 28.69
23.62
Cl
NH
NH
S
O
O
f1 (ppm)

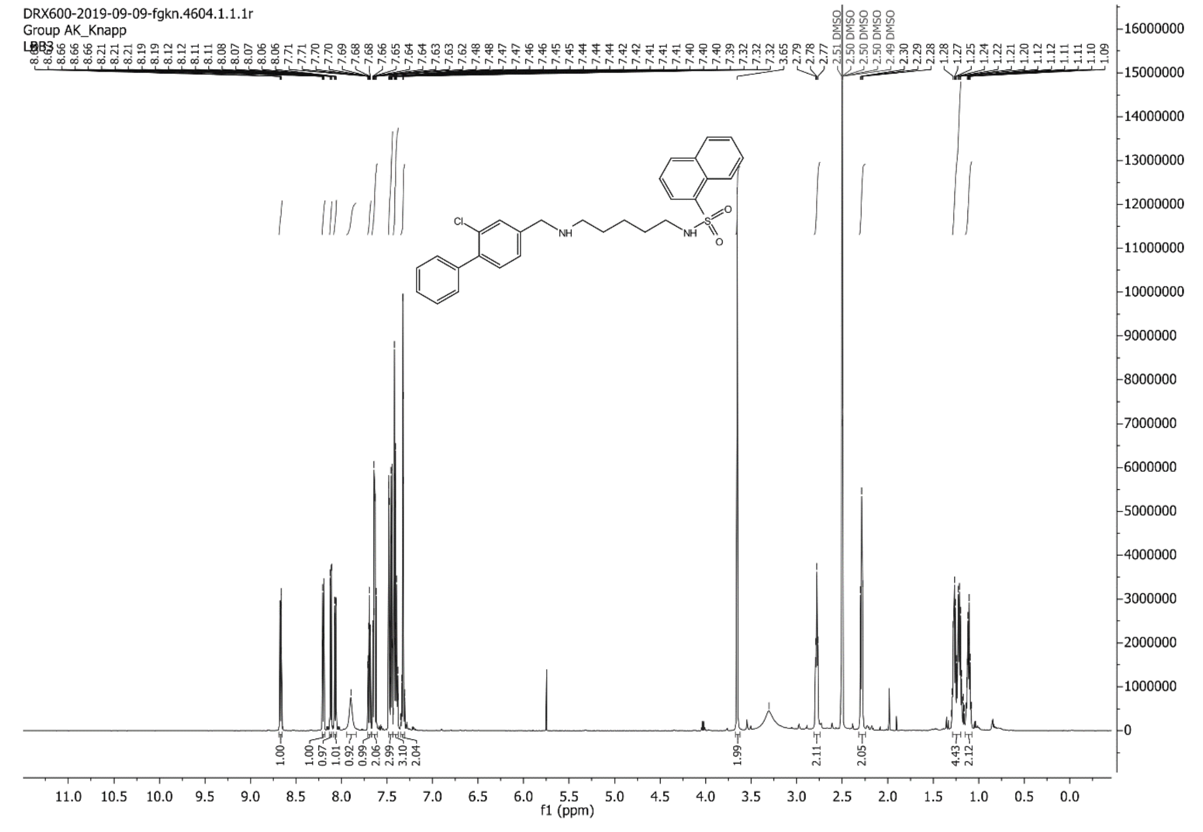
DRX600-2019-09-09-fgkn.4604.1.1.1r
Group AK_Knapp
LBB3
8.66 8.66 8.21 8.21 8.21 8.19 8.19 8.12 8.12 8.11 8.11 8.08 8.07 8.06 8.06 7.71 7.71 7.70 7.70 7.69 7.68 7.68 7.66 7.65 7.64 7.64 7.63 7.63 7.62 7.48 7.48 7.47 7.46 7.46 7.45 7.45 7.44 7.44 7.42 7.42 7.41 7.41 7.40 7.40 7.39 7.32 7.32 7.32 3.65 2.79 2.78 2.77 2.51 DMSO 2.50 DMSO 2.50 DMSO 2.49 DMSO 2.30 2.29 2.28 1.28 1.27 1.25 1.24 1.22 1.21 1.20 1.12 1.12 1.11 1.11 1.10 1.09
Cl
NH
NH
S
O
O
1.00 1.00 0.97 1.01 0.92 0.99 2.06 2.99 3.10 2.04 1.99 2.11 2.05 4.43 2.12
f1 (ppm)

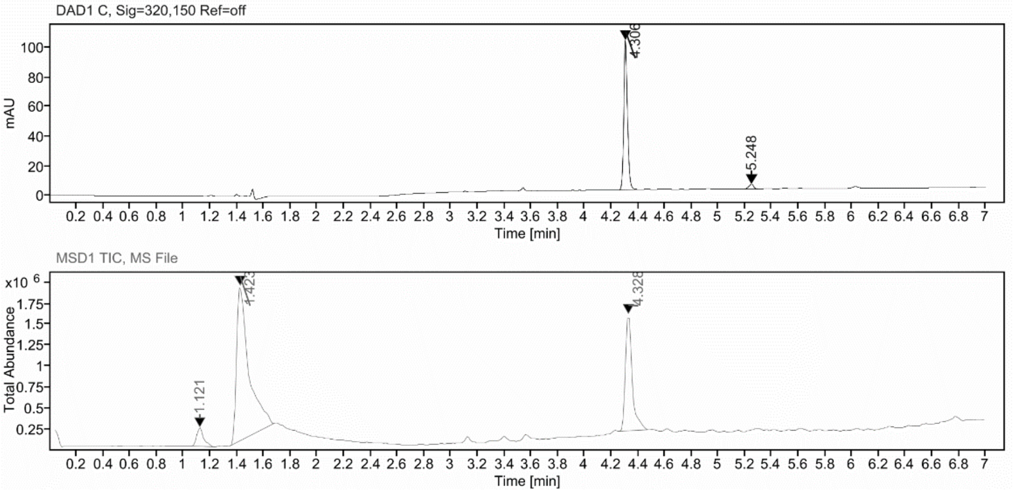

**Sample Purity**

Signal Description DAD1 C, Sig=320,150 Ref=off

| Sample Name | Name | RT | Width | Area | Area% | Height |
|---|---|---|---|---|---|---|
| FGLB3_X | | 4.306 | 0.028 | 190.0447 | 95.92 | 100.7829 |
| FGLB3_X | | 5.248 | 0.038 | 8.0868 | 4.08 | 3.2912 |

Max Area% 95.918

UV Signal Purity>95% Pass

**UV Apex Spectra**

**RT:**

4.306

**Sample Name:**

FGLB3_X

**Signal Name:**

DAD1C

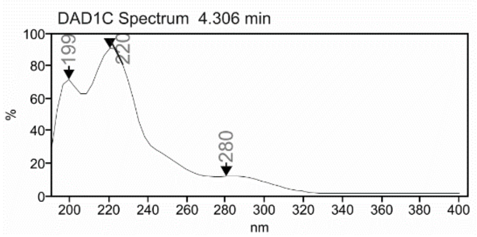

**RT:**

4.328

**Sample Name:**

FGLB3_X

**Signal Name:**

MSD1TIC

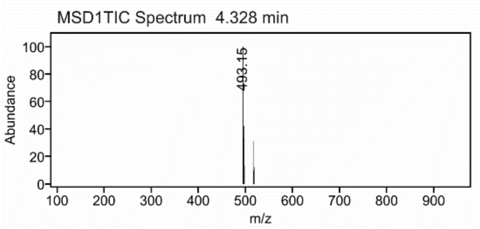

### Mass Spectrum SmartFormula Report

**Analysis Info**

| | | | |
|---|---|---|---|
| Analysis Name | D:\Data\MS_Service\Service_Kaiser\2024_Q1\20240228_SKN_FGLB3_100x_5uL-m.d | Acquisition Date | 2/28/2024 2:13:04 PM |
| Method | DirectInfusion_WithFlow_Tune_pos.m | Operator | BDAL@DE |
| Sample Name | Test1 | Instrument | micrOTOF-Q 228888.10407 |
| Comment | | | |

**Acquisition Parameter**

| | | | | | |
|---|---|---|---|---|---|
| Source Type | ESI | Ion Polarity | Positive | Set Nebulizer | 1.8 Bar |
| Focus | Active | Set Capillary | 4000 V | Set Dry Heater | 320 °C |
| Scan Begin | 50 m/z | Set End Plate Offset | -500 V | Set Dry Gas | 8.0 l/min |
| Scan End | 1600 m/z | Set Collision Cell RF | 150.0 Vpp | Set Divert Valve | Waste |

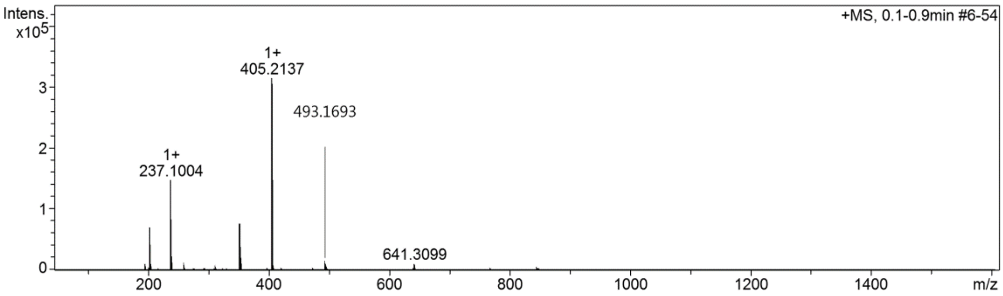

**Cmpd 28**

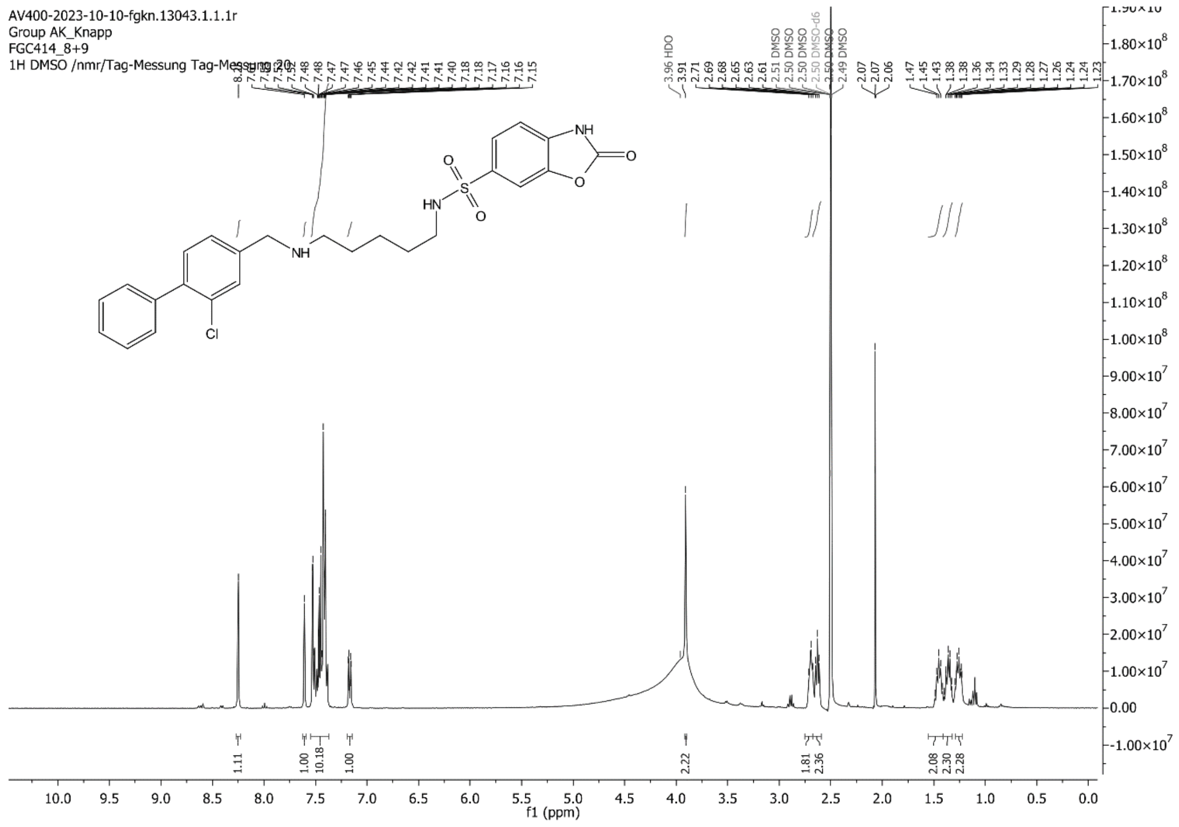

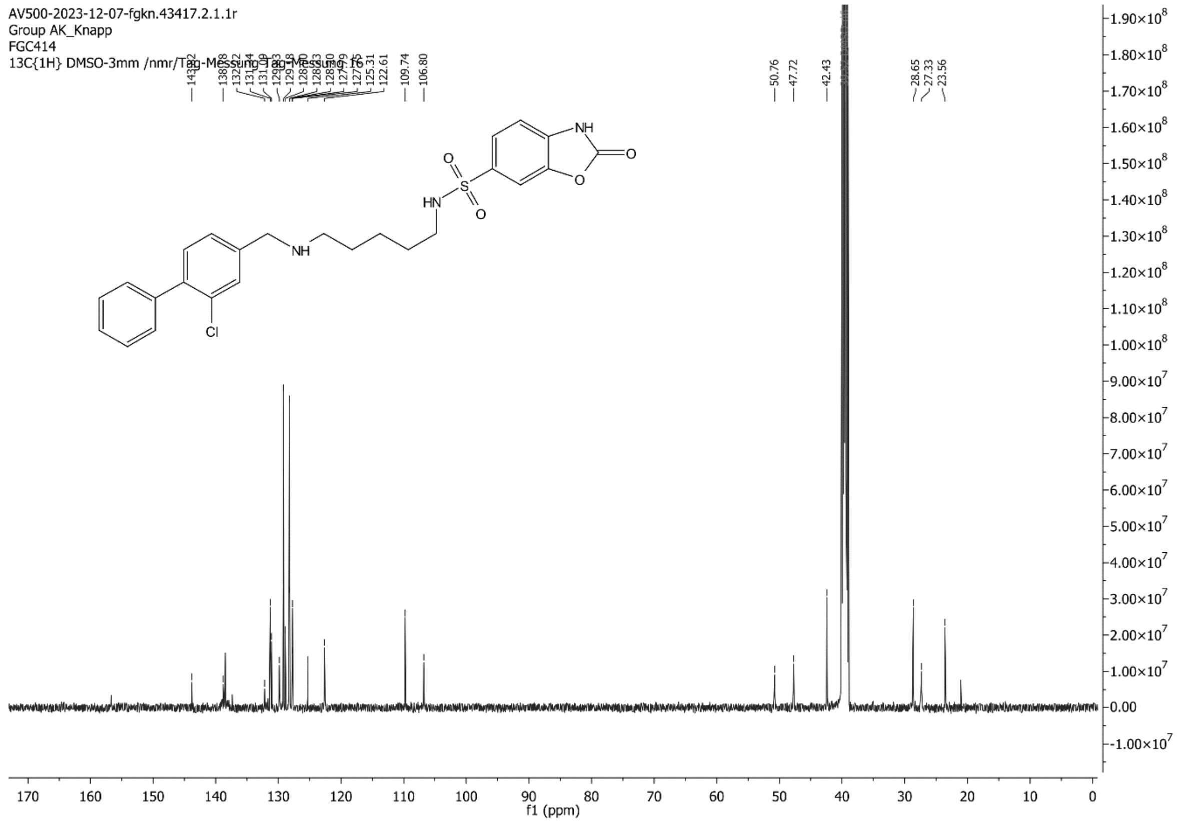

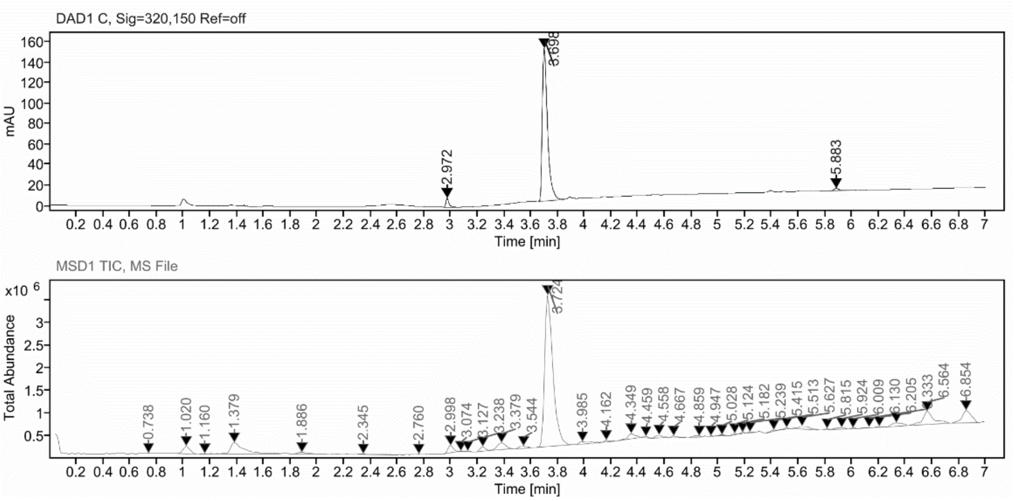

**Sample Purity**

| Signal Description | DAD1 C, Sig=320,150 Ref=off |
|---|---|

| Sample Name | Name | RT | Width | Area | Area% | Height |
|---|---|---|---|---|---|---|
| FGC414_f10 | | 2.972 | 0.021 | 13.5506 | 3.13 | 8.8338 |
| FGC414_f10 | | 3.698 | 0.041 | 412.5047 | 95.34 | 149.3519 |
| FGC414_f10 | | 5.883 | 0.038 | 6.6001 | 1.53 | 2.4245 |

| | |
|---|---|
| Max Area% | 95.343 |
| UV Signal Purity>95% | Pass |

**UV Apex Spectra**

**RT:**

2.972

**Sample Name:**

FGC414_f10

**Signal Name:**

DAD1C

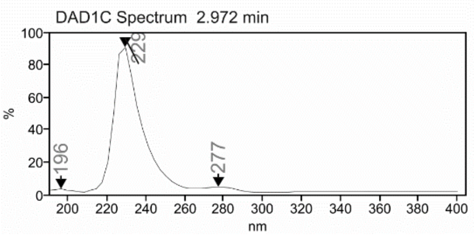

**RT:**

3.724

**Sample Name:**

FGC414_f10

**Signal Name:**

MSD1TIC

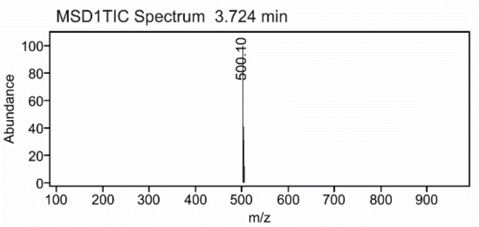

### Cmpd 29

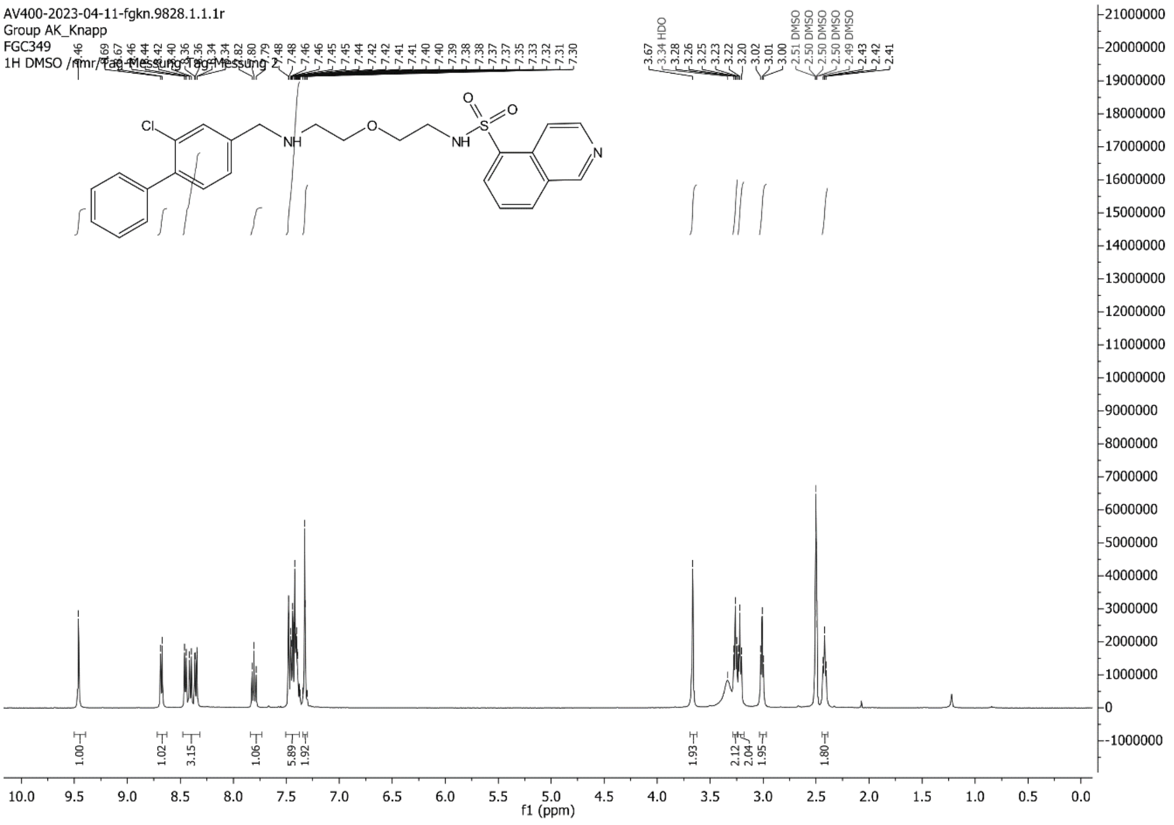
AV400-2023-04-11-fgkn.9828.1.1.1r
Group AK_Knapp
FGC349
1H DMSO /nmr/Tag-Messung Tag-Messung 2
f1 (ppm)

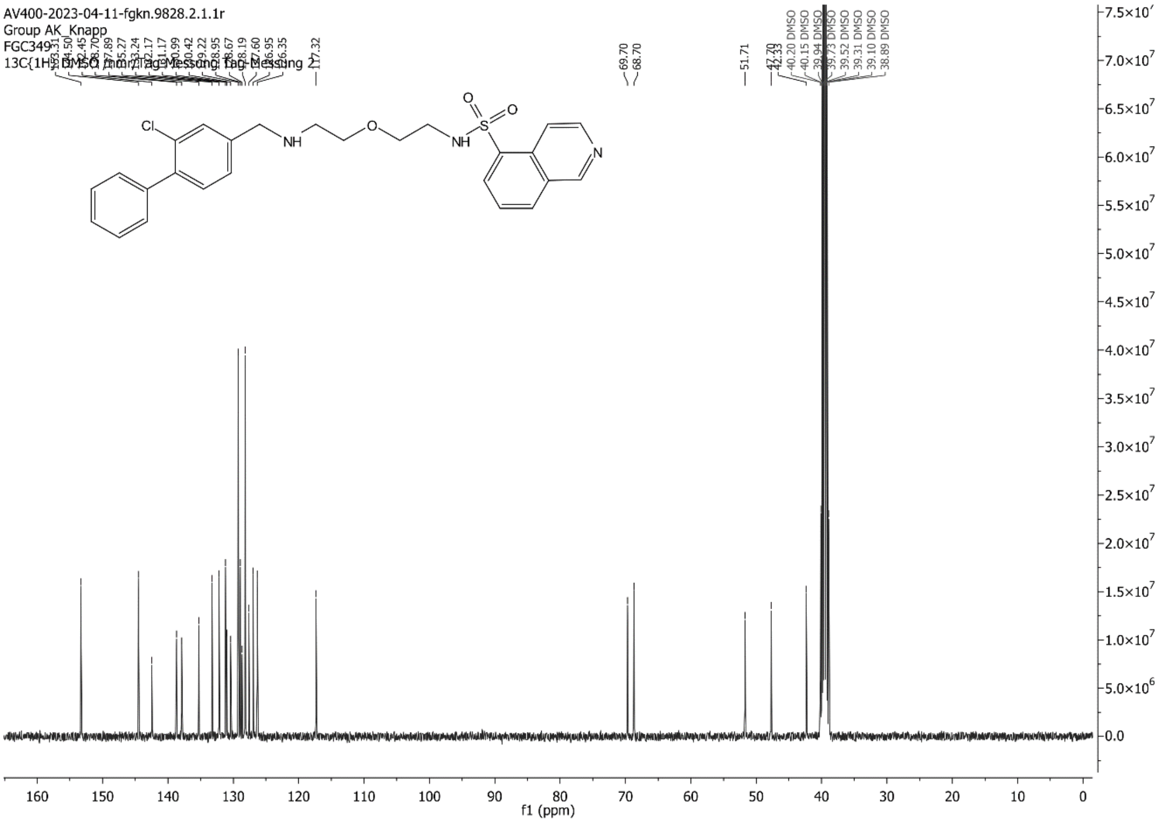
AV400-2023-04-11-fgkn.9828.2.1.1r
Group AK_Knapp
FGC349
13C{1H} DMSO /nmr/Tag-Messung Tag-Messung 2
f1 (ppm)

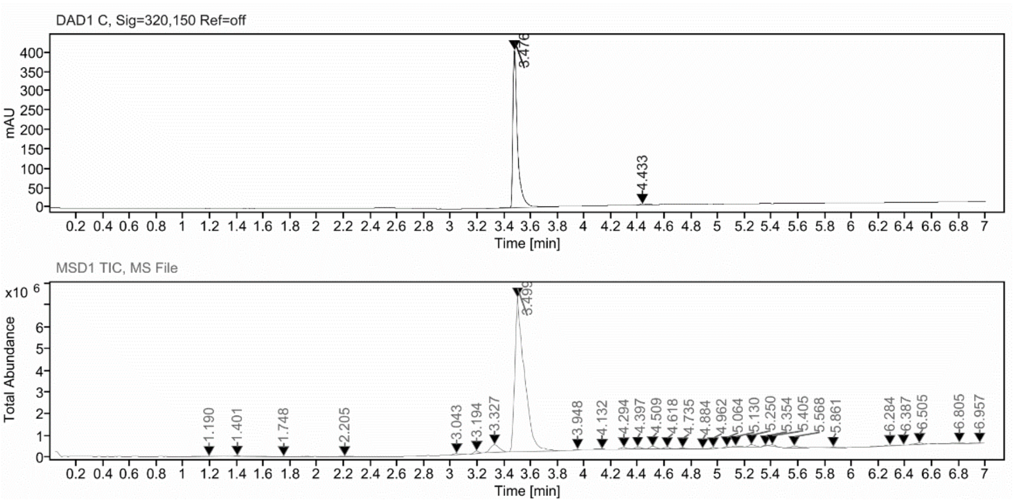

**Sample Purity**

Signal Description DAD1 C, Sig=320,150 Ref=off

| Sample Name | Name | RT | Width | Area | Area% | Height |
|---|---|---|---|---|---|---|
| FGC349_2_fr6 | | 3.476 | 0.032 | 967.1360 | 98.89 | 406.2814 |
| FGC349_2_fr6 | | 4.433 | 0.059 | 10.8585 | 1.11 | 3.0938 |

Max Area% 98.890

UV Signal Purity>95% Pass

**UV Apex Spectra**

**RT:**

3.476

**Sample Name:**

FGC349_2_fr6

**Signal Name:**

DAD1C

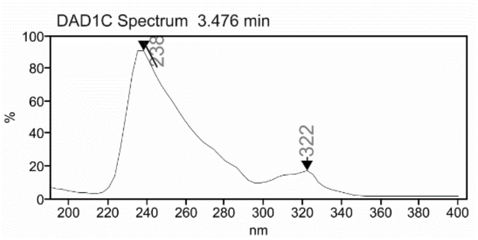

**RT:**

3.499

**Sample Name:**

FGC349_2_fr6

**Signal Name:**

MSD1TIC

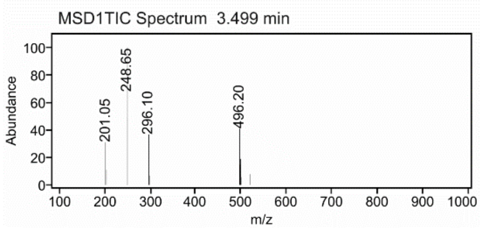

### Mass Spectrum SmartFormula Report

**Analysis Info**

| | | | |
|---|---|---|---|
| Analysis Name | D:\Data\MS_Service\Service_Kaiser\2024_Q1\20240219_SKN_FGC349_5ug-mL_5uL-m.d | Acquisition Date | 2/19/2024 2:27:02 PM |
| Method | DirectInfusion_WithFlow_Tune_pos.m | Operator | BDAL@DE |
| Sample Name | Test1 | Instrument | micrOTOF-Q 228888.10407 |
| Comment | | | |

**Acquisition Parameter**

| | | | | | |
|---|---|---|---|---|---|
| Source Type | ESI | Ion Polarity | Positive | Set Nebulizer | 1.8 Bar |
| Focus | Active | Set Capillary | 4000 V | Set Dry Heater | 220 °C |
| Scan Begin | 50 m/z | Set End Plate Offset | -500 V | Set Dry Gas | 8.0 l/min |
| Scan End | 1600 m/z | Set Collision Cell RF | 150.0 Vpp | Set Divert Valve | Waste |

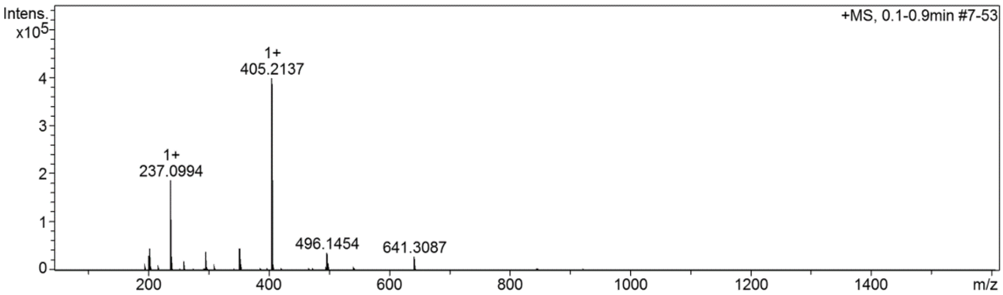

**Cmpd 30**

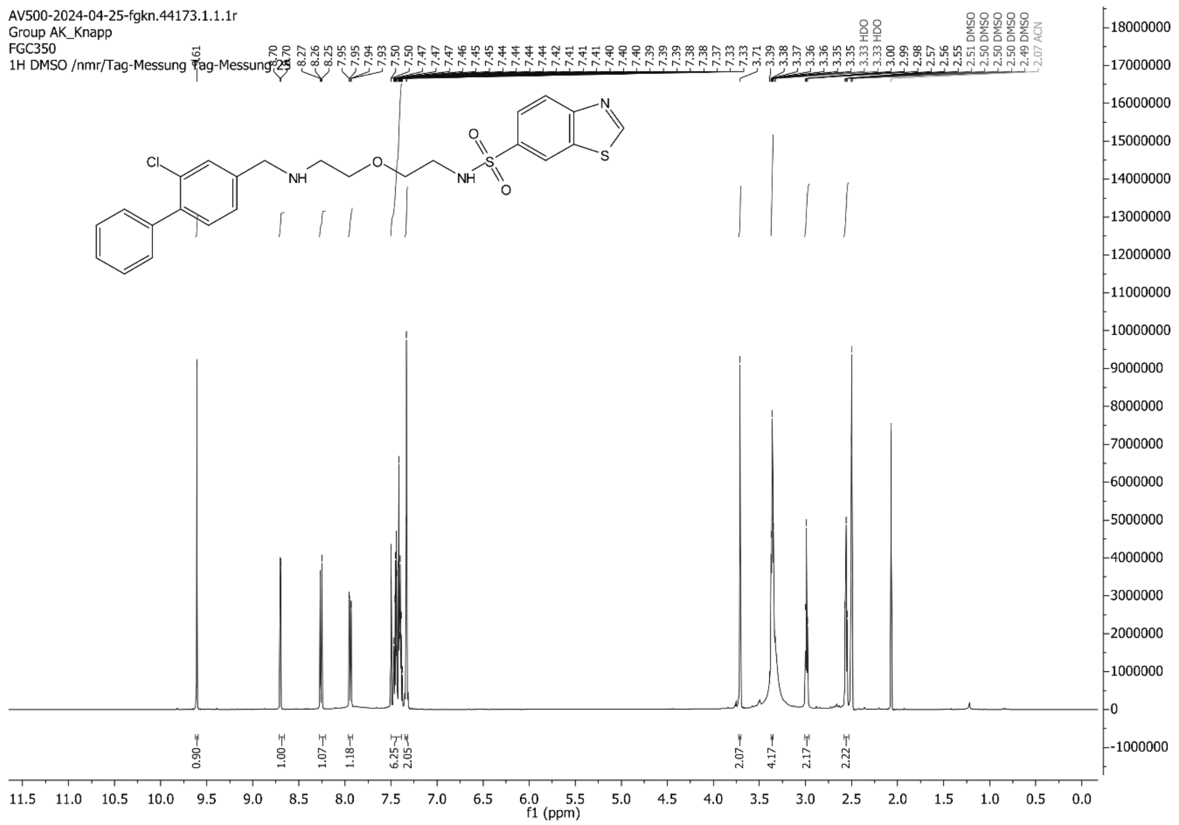

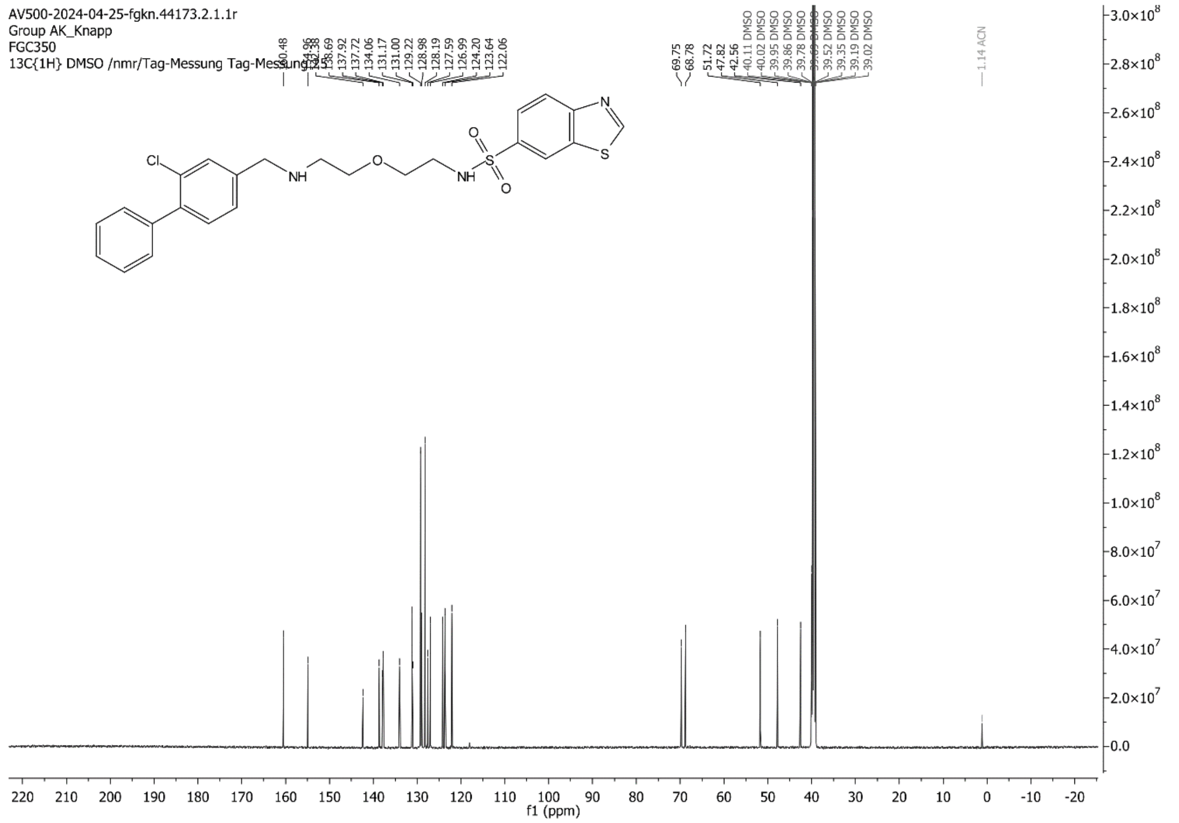

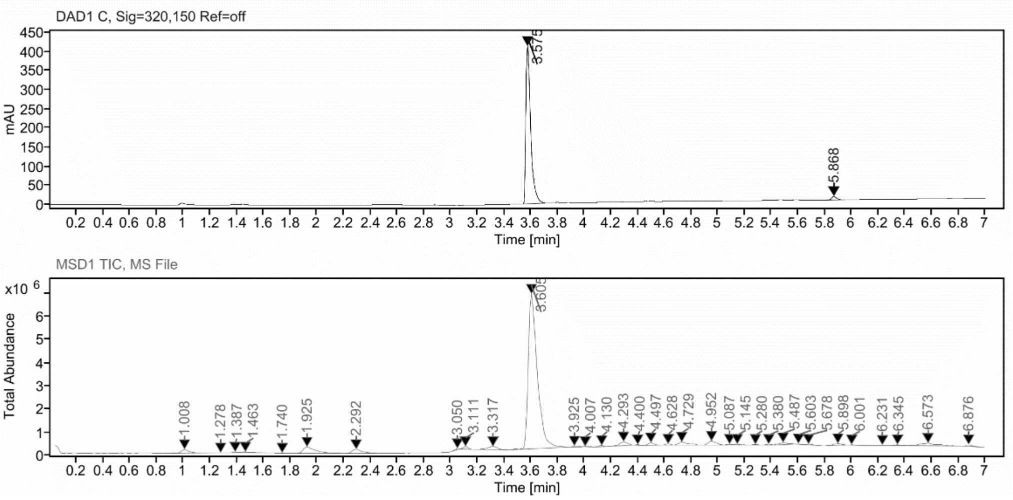

**Sample Purity**

Signal Description DAD1 C, Sig=320,150 Ref=off

| Sample Name | Name | RT | Width | Area | Area% | Height |
|---|---|---|---|---|---|---|
| FGC350_stabX | | 3.575 | 0.037 | 1032.9807 | 97.76 | 410.9646 |
| FGC350_stabX | | 5.868 | 0.038 | 23.6448 | 2.24 | 9.1461 |

Max Area% 97.762

UV Signal Purity>95% Pass

**UV Apex Spectra**

**RT:**

3.575

**Sample Name:**

FGC350_stabX

**Signal Name:**

DAD1C

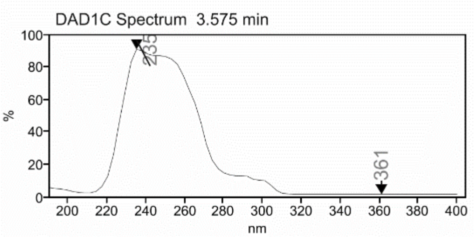

**RT:**

3.605

**Sample Name:**

FGC350_stabX

**Signal Name:**

MSD1TIC

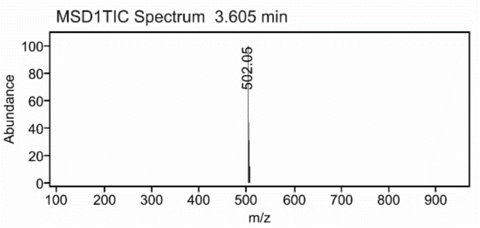

### Mass Spectrum SmartFormula Report

**Analysis Info**

| | | | |
|---|---|---|---|
| | | Acquisition Date | 2/19/2024 2:33:49 PM |
| Analysis Name | D:\Data\MS_Service\Service_Kaiser\2024_Q1\20240219_SKN_FGC350_5ug-mL_5uL-m.d | | |
| Method | DirectInfusion_WithFlow_Tune_pos.m | Operator | BDAL@DE |
| Sample Name | Test1 | Instrument | micrOTOF-Q 228888.10407 |
| Comment | | | |

**Acquisition Parameter**

| | | | | | |
|---|---|---|---|---|---|
| Source Type | ESI | Ion Polarity | Positive | Set Nebulizer | 1.8 Bar |
| Focus | Active | Set Capillary | 4000 V | Set Dry Heater | 220 °C |
| Scan Begin | 50 m/z | Set End Plate Offset | -500 V | Set Dry Gas | 8.0 l/min |
| Scan End | 1600 m/z | Set Collision Cell RF | 150.0 Vpp | Set Divert Valve | Waste |

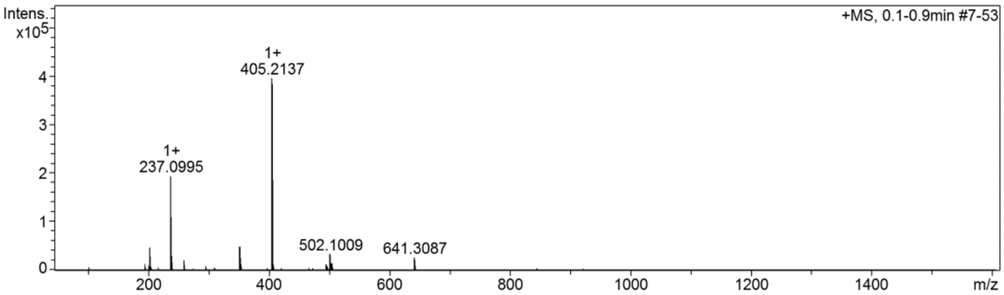

#### Cmpd 31

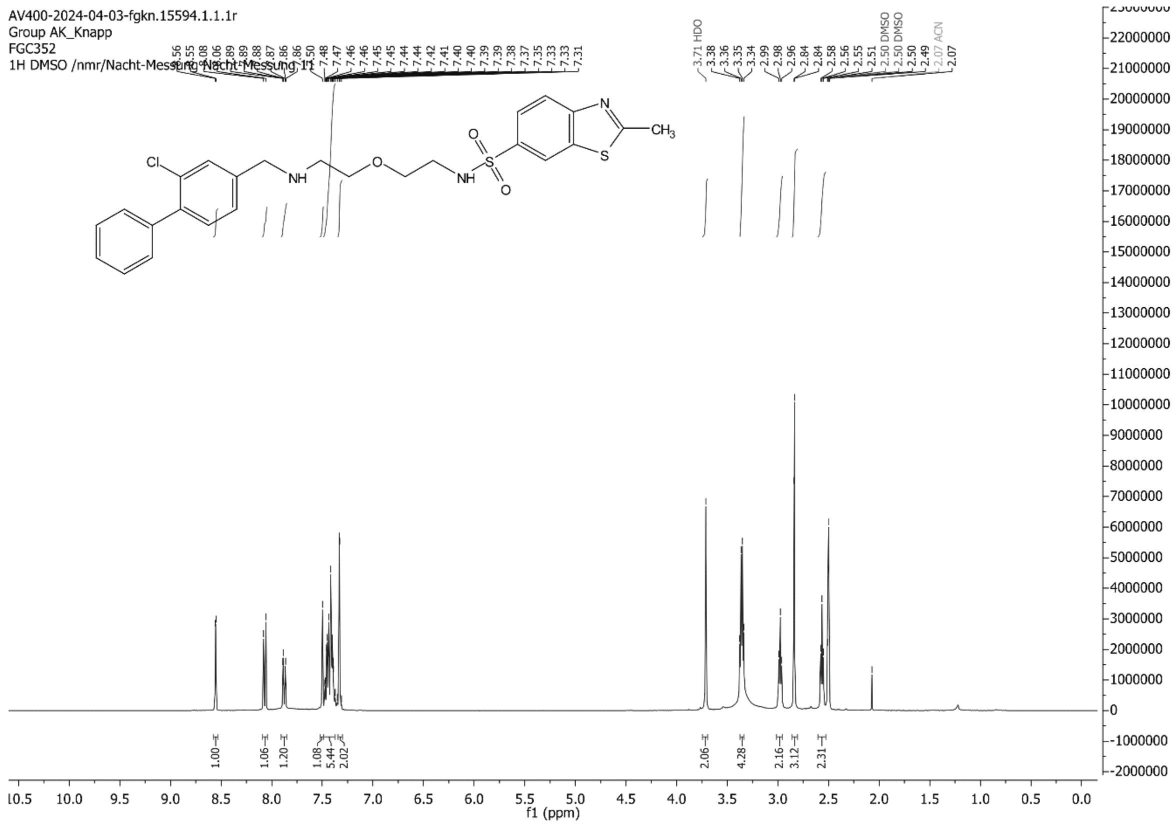

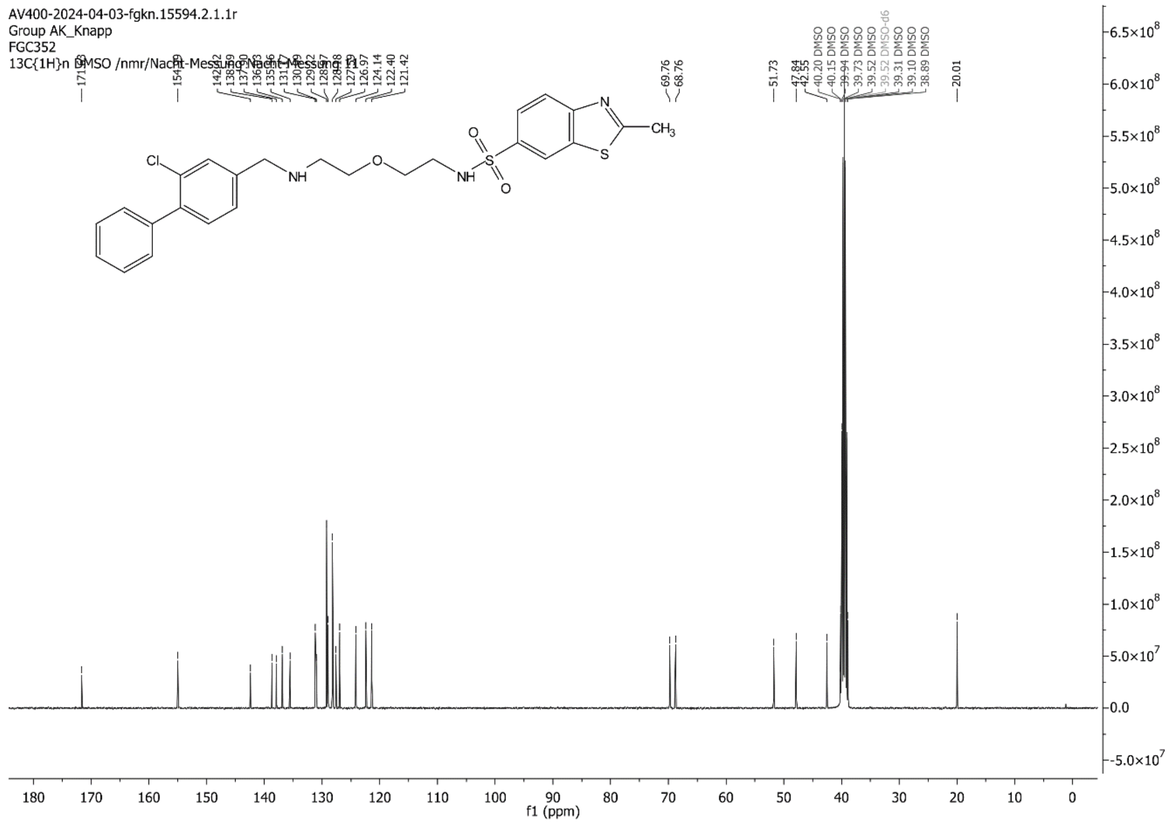

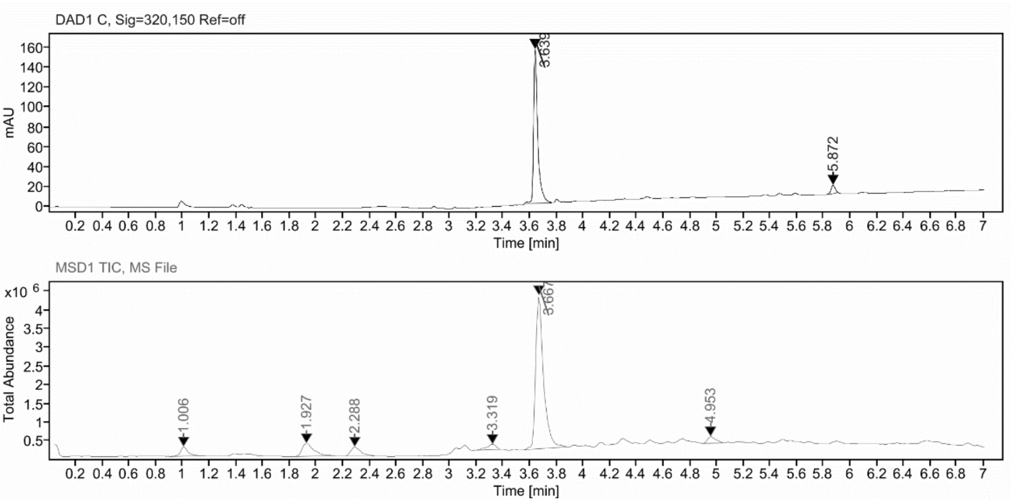

**Sample Purity**

Signal Description DAD1 C, Sig=320,150 Ref=off

| Sample Name | Name | RT | Width | Area | Area% | Height |
|---|---|---|---|---|---|---|
| FGC352_stabX | | 3.639 | 0.031 | 339.7262 | 95.07 | 155.2145 |
| FGC352_stabX | | 5.872 | 0.035 | 17.6224 | 4.93 | 8.4811 |

Max Area% 95.069

UV Signal Purity>95% Pass

**UV Apex Spectra**

**RT:**

3.639

**Sample Name:**

FGC352_stabX

**Signal Name:**

DAD1C

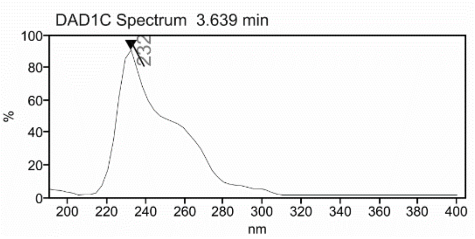

**RT:**

3.667

**Sample Name:**

FGC352_stabX

**Signal Name:**

MSD1TIC

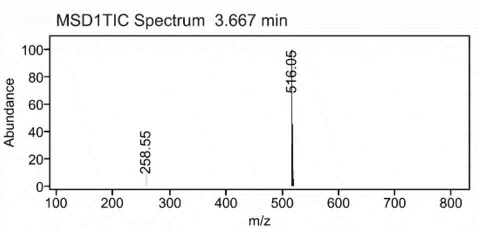

### Mass Spectrum SmartFormula Report

**Analysis Info**

| | | | |
|---|---|---|---|
| | | Acquisition Date | 2/19/2024 2:40:53 PM |
| Analysis Name | D:\Data\MS_Service\Service_Kaiser\2024_Q1\20240219_SKN_FGC352_5ug-mL_5uL-m.d | | |
| Method | DirectInfusion_WithFlow_Tune_pos.m | Operator | BDAL@DE |
| Sample Name | Test1 | Instrument | micrOTOF-Q 228888.10407 |
| Comment | | | |

**Acquisition Parameter**

| | | | | | |
|---|---|---|---|---|---|
| Source Type | ESI | Ion Polarity | Positive | Set Nebulizer | 1.8 Bar |
| Focus | Active | Set Capillary | 4000 V | Set Dry Heater | 220 °C |
| Scan Begin | 50 m/z | Set End Plate Offset | -500 V | Set Dry Gas | 8.0 l/min |
| Scan End | 1600 m/z | Set Collision Cell RF | 150.0 Vpp | Set Divert Valve | Waste |

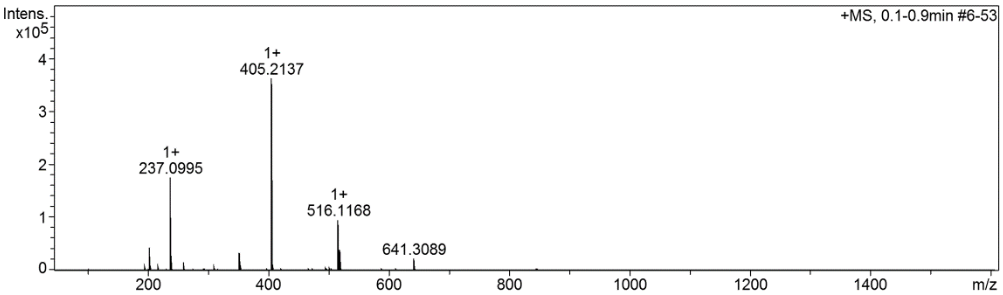

**Cmpd 32**

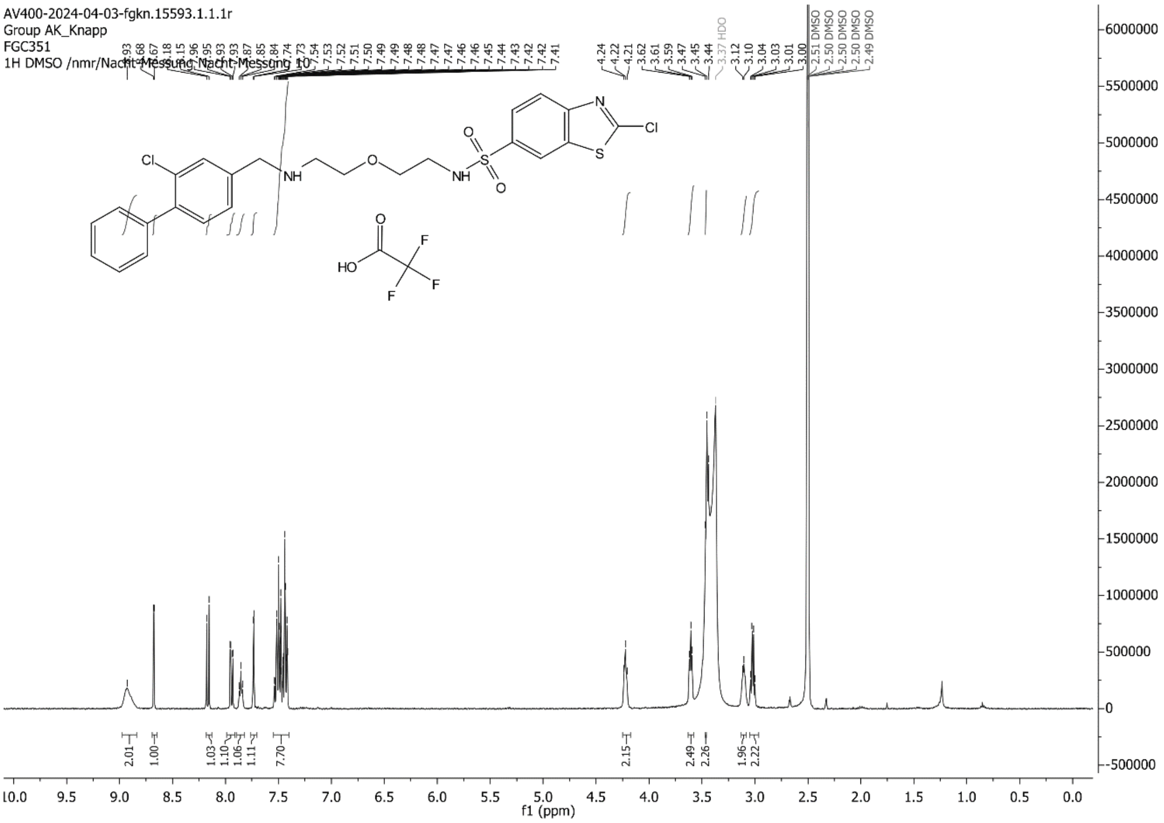

AV400-2024-04-03-fgkn.15593.1.1.1r
Group AK_Knapp
FGC351
1H DMSO /nmr/Nacht-Messung Nacht-Messung 10
f1 (ppm)

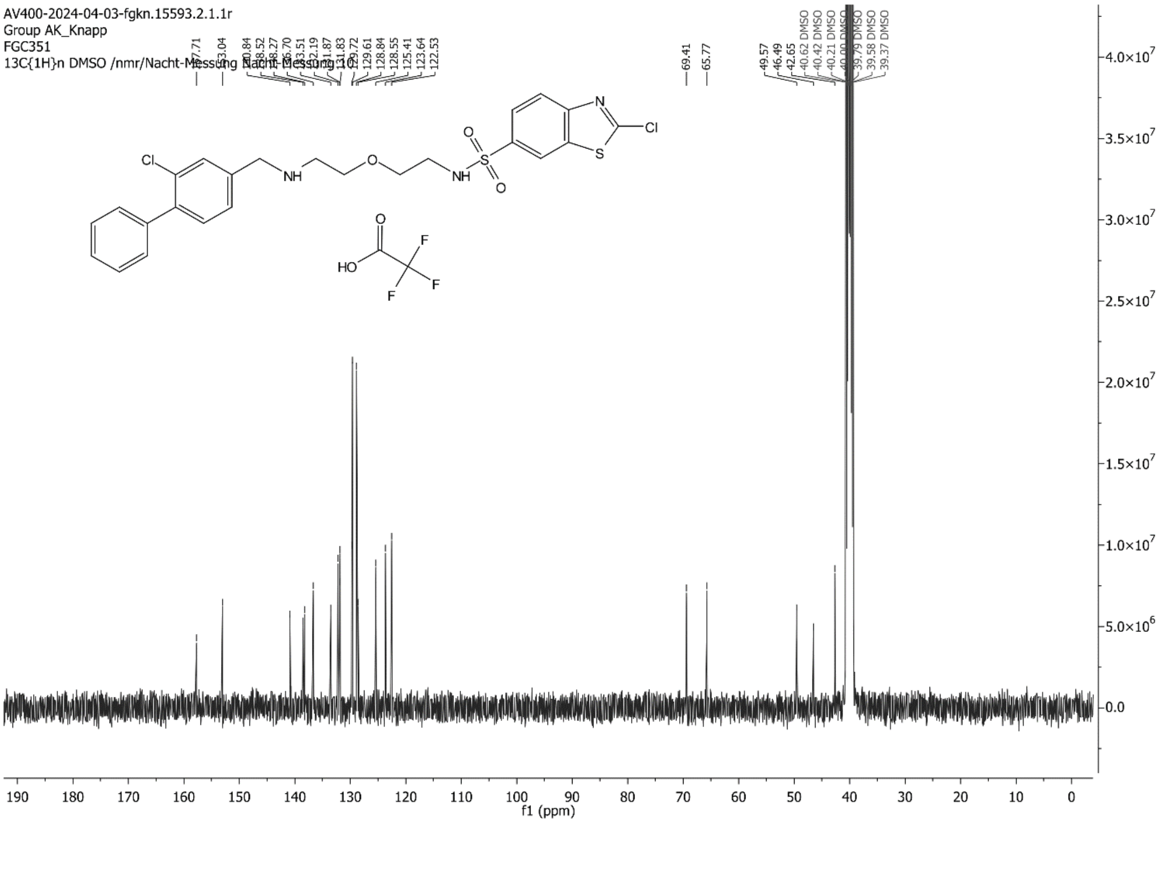

AV400-2024-04-03-fgkn.15593.2.1.1r
Group AK_Knapp
FGC351
13C{1H}n DMSO /nmr/Nacht-Messung Nacht-Messung 10
f1 (ppm)

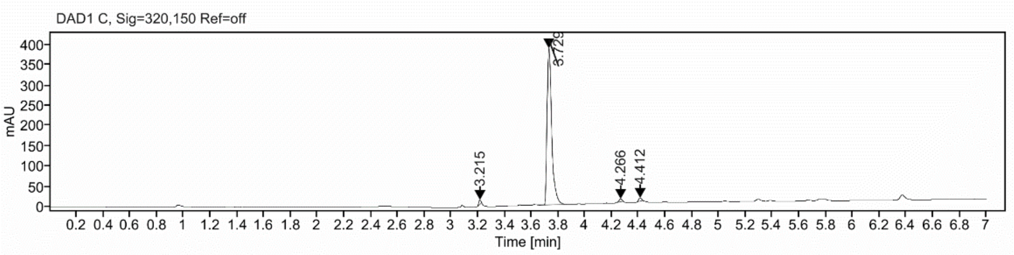

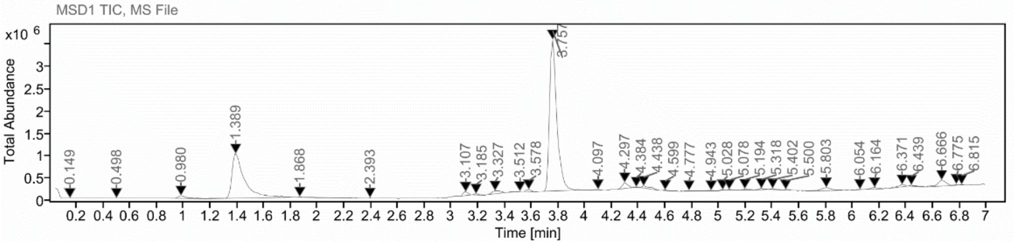

**Sample Purity**

Signal Description DAD1 C, Sig=320,150 Ref=off

| Sample Name | Name | RT | Width | Area | Area% | Height |
|---|---|---|---|---|---|---|
| FGC351 | | 3.215 | 0.018 | 10.8801 | 1.13 | 10.6211 |
| FGC351 | | 3.729 | 0.036 | 932.5604 | 96.57 | 385.7572 |
| FGC351 | | 4.266 | 0.023 | 10.0347 | 1.04 | 7.2008 |
| FGC351 | | 4.412 | 0.025 | 12.1724 | 1.26 | 8.5058 |

| | |
|---|---|
| Max Area% | 96.574 |
| UV Signal Purity>95% | Pass |

**UV Apex Spectra**

**RT:**

3.215

**Sample Name:**

FGC351

**Signal Name:**

DAD1C

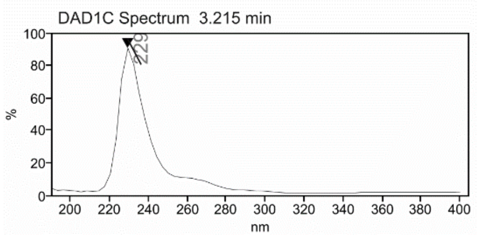

**RT:**

3.757

**Sample Name:**

FGC351

**Signal Name:**

MSD1TIC

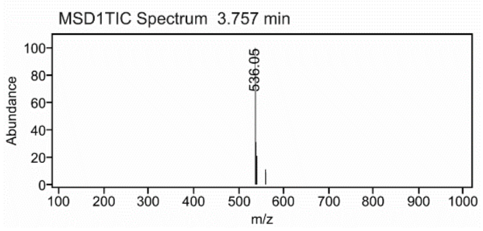

### Mass Spectrum SmartFormula Report

**Analysis Info**

| | | | |
|---|---|---|---|
| Analysis Name | D:\Data\MS_Service\Service_Kaiser\2024_Q1\20240227_SKN_FGC351_100x_5uL-m.d | Acquisition Date | 2/27/2024 2:38:13 PM |
| Method | DirectInfusion_WithFlow_Tune_pos.m | Operator | BDAL@DE |
| Sample Name | Test1 | Instrument | micrOTOF-Q 228888.10407 |
| Comment | | | |

**Acquisition Parameter**

| | | | | | |
|---|---|---|---|---|---|
| Source Type | ESI | Ion Polarity | Positive | Set Nebulizer | 1.8 Bar |
| Focus | Active | Set Capillary | 4000 V | Set Dry Heater | 320 °C |
| Scan Begin | 50 m/z | Set End Plate Offset | -500 V | Set Dry Gas | 8.0 l/min |
| Scan End | 1600 m/z | Set Collision Cell RF | 150.0 Vpp | Set Divert Valve | Waste |

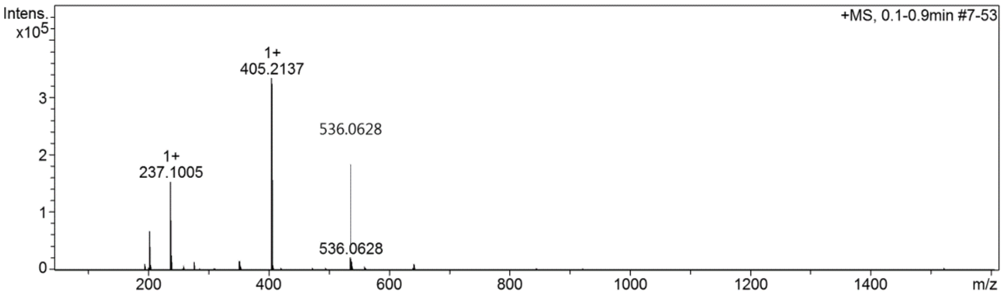

**Cmpd 33**

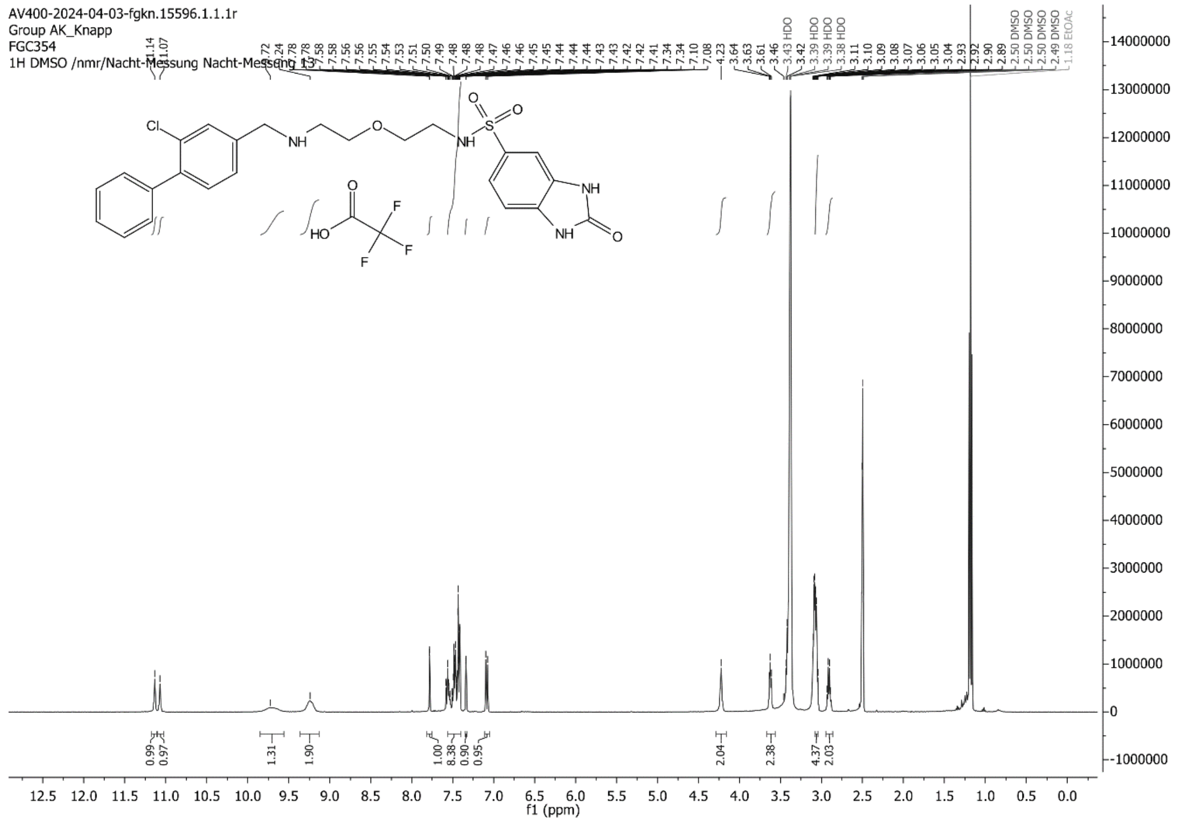

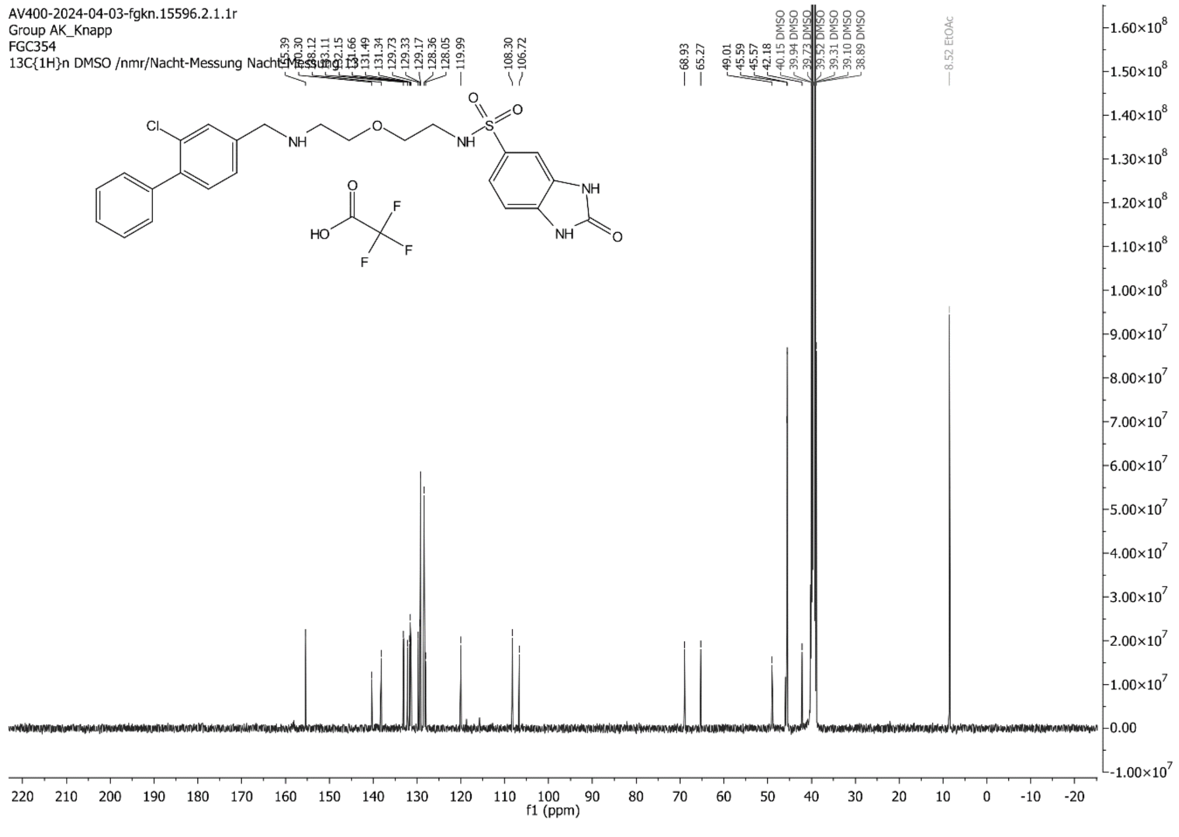

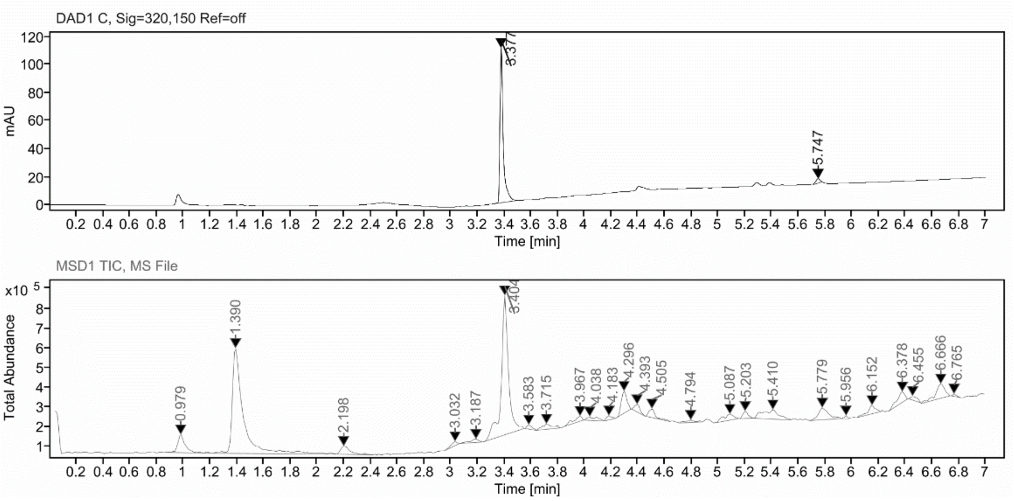

##### Sample Purity

Signal Description DAD1 C, Sig=320,150 Ref=off

| Sample Name | Name | RT | Width | Area | Area% | Height |
|---|---|---|---|---|---|---|
| FGC354 | | 3.377 | 0.022 | 186.5521 | 96.68 | 110.2966 |
| FGC354 | | 5.747 | 0.037 | 6.4074 | 3.32 | 3.0311 |

Max Area% 96.679

UV Signal Purity>95% Pass

##### UV Apex Spectra

**RT:**

3.377

**Sample Name:**

FGC354

**Signal Name:**

DAD1C

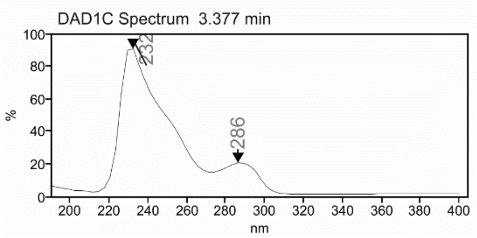

**RT:**

3.404

**Sample Name:**

FGC354

**Signal Name:**

MSD1TIC

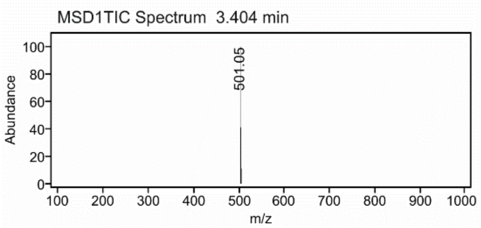

### Mass Spectrum SmartFormula Report

**Analysis Info**

| | | | |
|---|---|---|---|
| Analysis Name | D:\Data\MS_Service\Service_Kaiser\2024_Q1\20240219_SKN_FGC354_5ug-mL_5uL-m.d | Acquisition Date | 2/19/2024 2:56:15 PM |
| Method | DirectInfusion_WithFlow_Tune_pos.m | Operator | BDAL@DE |
| Sample Name | Test1 | Instrument | micrOTOF-Q 228888.10407 |
| Comment | | | |

**Acquisition Parameter**

| | | | | | |
|---|---|---|---|---|---|
| Source Type | ESI | Ion Polarity | Positive | Set Nebulizer | 1.8 Bar |
| Focus | Active | Set Capillary | 4000 V | Set Dry Heater | 220 °C |
| Scan Begin | 50 m/z | Set End Plate Offset | -500 V | Set Dry Gas | 8.0 l/min |
| Scan End | 1600 m/z | Set Collision Cell RF | 150.0 Vpp | Set Divert Valve | Waste |

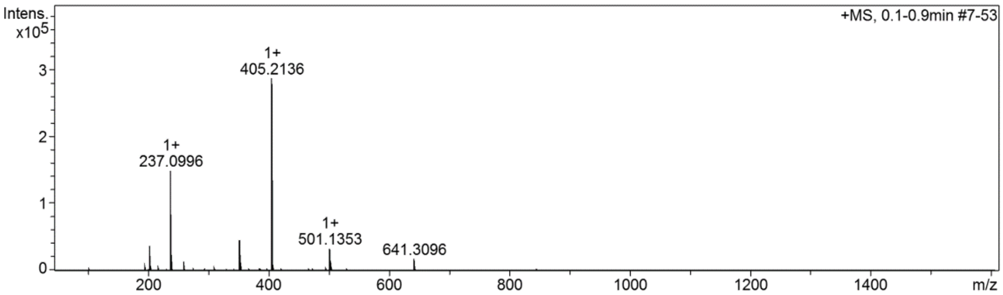

#### Cmpd 34

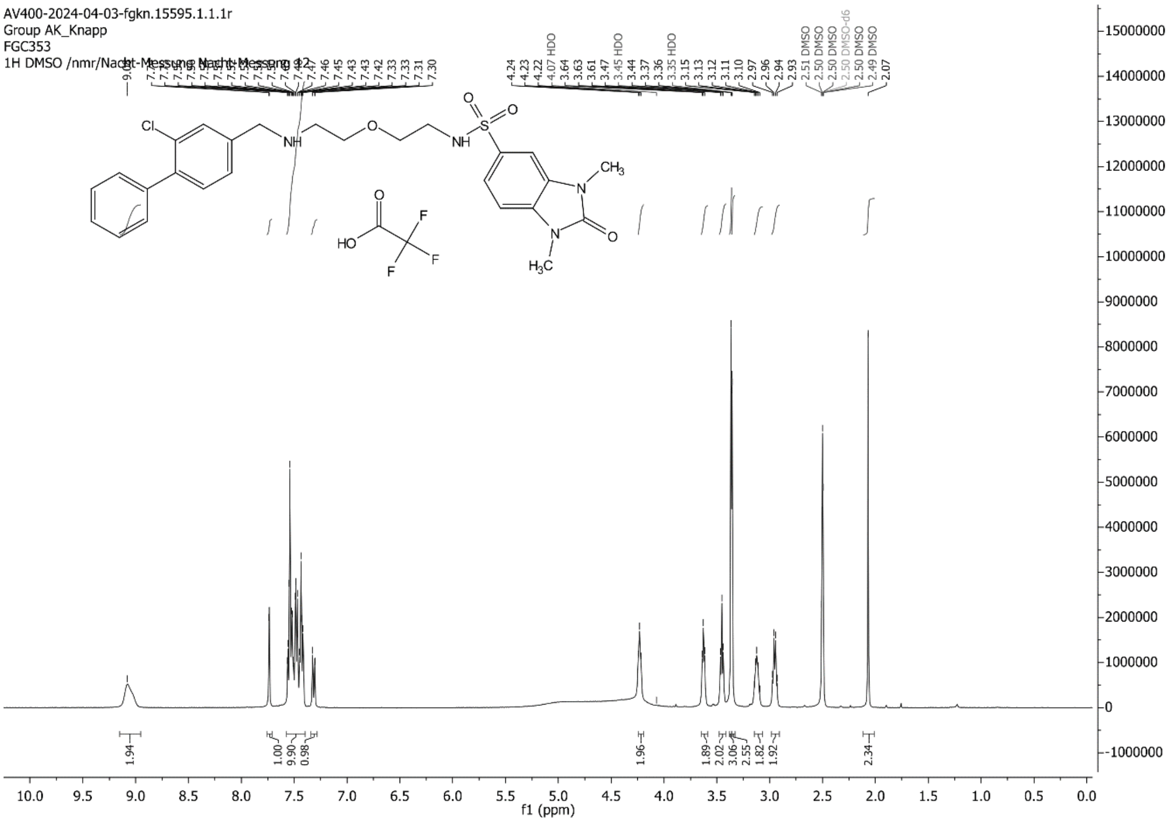
AV400-2024-04-03-fgkn.15595.1.1.1r
Group AK_Knapp
FGC353
1H DMSO /nmr/Nacht-Messung Nacht-Messung 12
f1 (ppm)

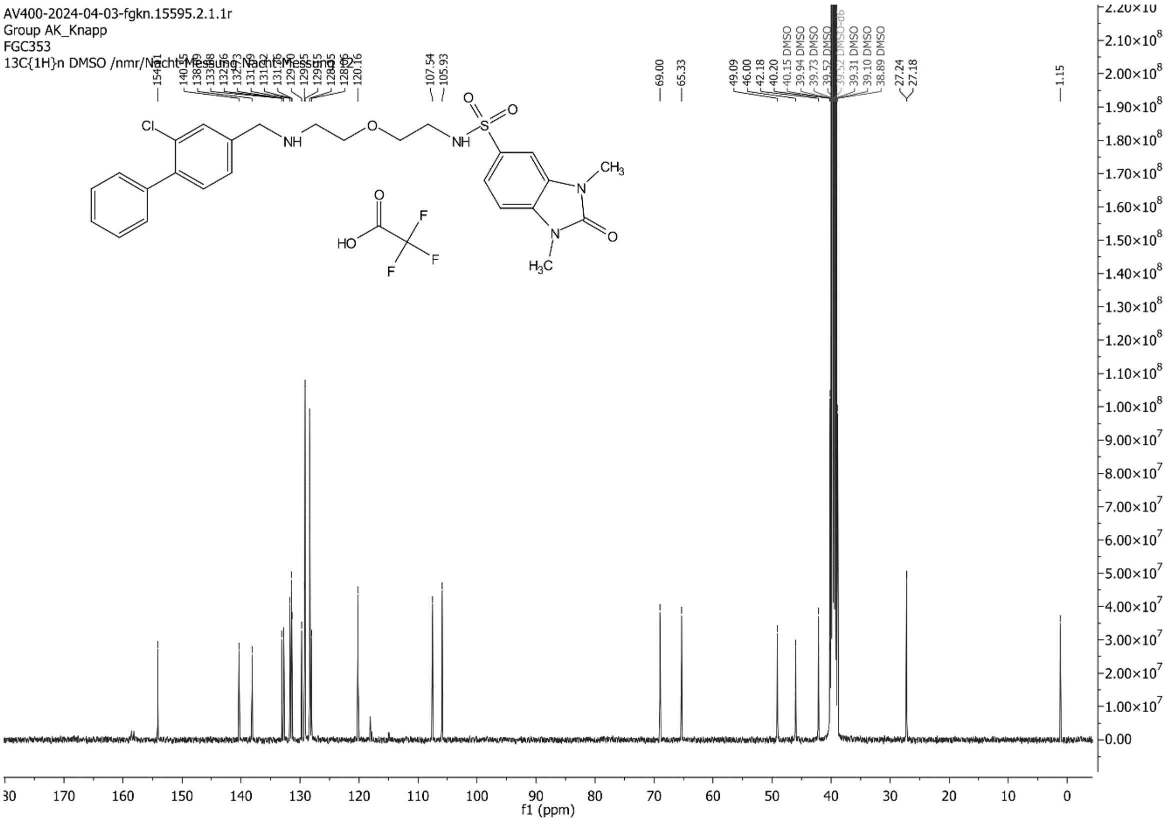
AV400-2024-04-03-fgkn.15595.2.1.1r
Group AK_Knapp
FGC353
13C{1H}n DMSO /nmr/Nacht-Messung Nacht-Messung 12
f1 (ppm)

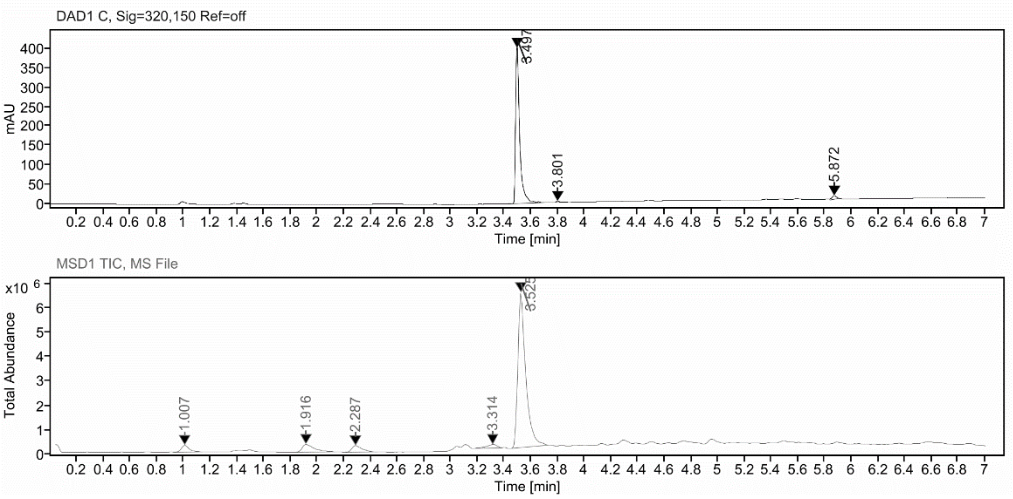

**Sample Purity**

Signal Description DAD1 C, Sig=320,150 Ref=off

| Sample Name | Name | RT | Width | Area | Area% | Height |
|---|---|---|---|---|---|---|
| FGC353_stabX | | 3.497 | 0.030 | 878.9980 | 96.73 | 400.5464 |
| FGC353_stabX | | 3.801 | 0.026 | 7.4783 | 0.82 | 3.6475 |
| FGC353_stabX | | 5.872 | 0.038 | 22.2260 | 2.45 | 9.5942 |

Max Area% 96.731

UV Signal Purity>95% Pass

**UV Apex Spectra**

**RT:**

3.497

**Sample Name:**

FGC353_stabX

**Signal Name:**

DAD1C

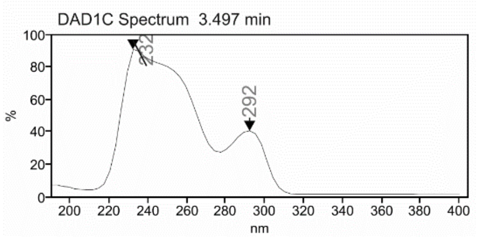

**RT:**

3.525

**Sample Name:**

FGC353_stabX

**Signal Name:**

MSD1TIC

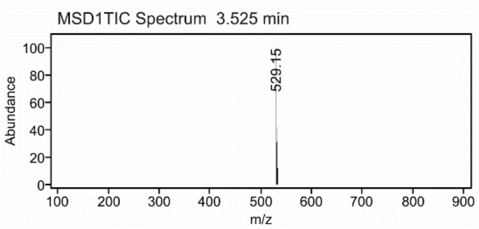

### Mass Spectrum SmartFormula Report

**Analysis Info**

| | | | |
|---|---|---|---|
| Analysis Name | D:\Data\MS_Service\Service_Kaiser\2024_Q1\20240219_SKN_FGC353_5ug-mL_5uL-m.d | Acquisition Date | 2/19/2024 2:47:47 PM |
| Method | DirectInfusion_WithFlow_Tune_pos.m | Operator | BDAL@DE |
| Sample Name | Test1 | Instrument | micrOTOF-Q 228888.10407 |
| Comment | | | |

**Acquisition Parameter**

| | | | | | |
|---|---|---|---|---|---|
| Source Type | ESI | Ion Polarity | Positive | Set Nebulizer | 1.8 Bar |
| Focus | Active | Set Capillary | 4000 V | Set Dry Heater | 220 °C |
| Scan Begin | 50 m/z | Set End Plate Offset | -500 V | Set Dry Gas | 8.0 l/min |
| Scan End | 1600 m/z | Set Collision Cell RF | 150.0 Vpp | Set Divert Valve | Waste |

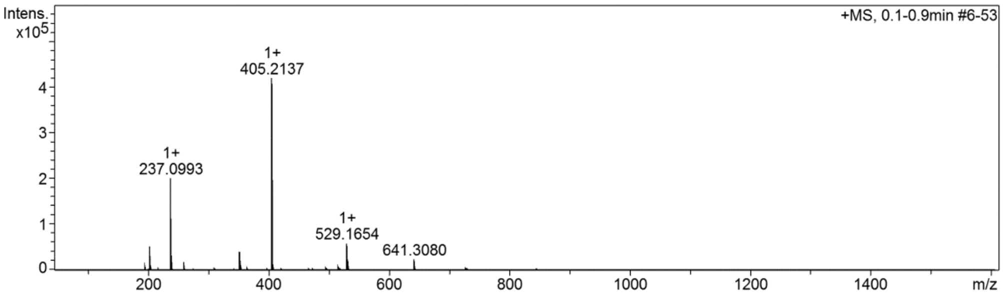

**Cmpd 35**

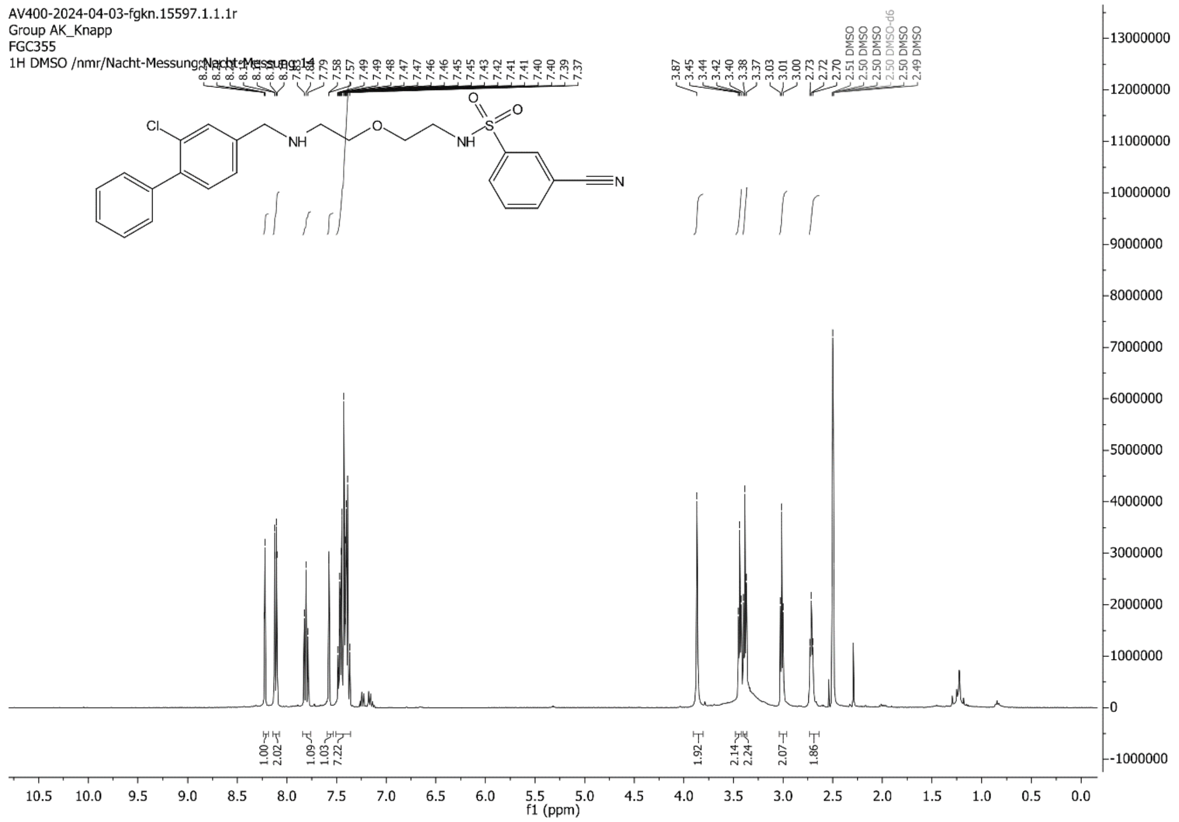
AV400-2024-04-03-fgkn.15597.1.1.1r
Group AK_Knapp
FGC355
1H DMSO /nmr/Nacht-Messung/Nacht-Messung 14
f1 (ppm)

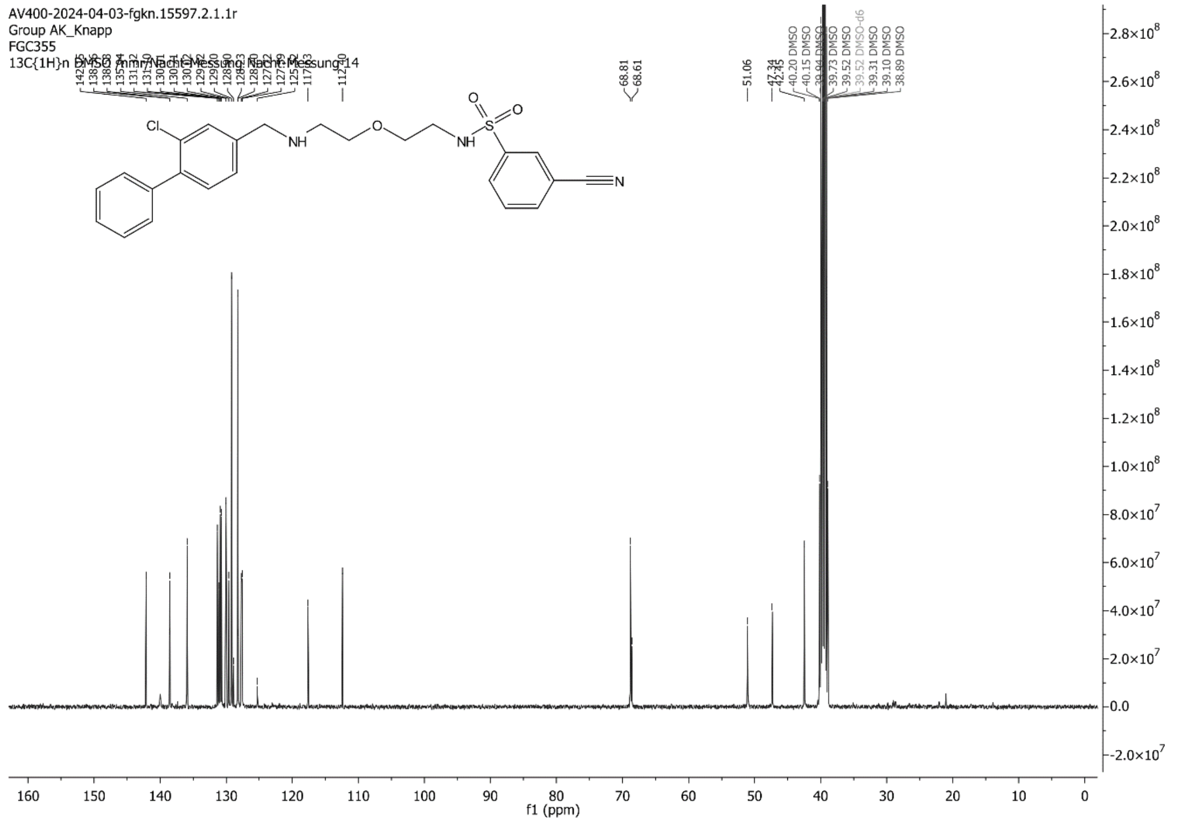
AV400-2024-04-03-fgkn.15597.2.1.1r
Group AK_Knapp
FGC355
13C{1H}n DMSO /nmr/Nacht-Messung/Nacht-Messung 14
f1 (ppm)

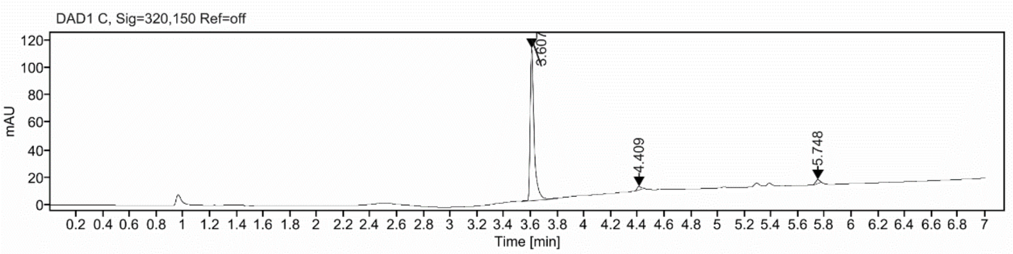

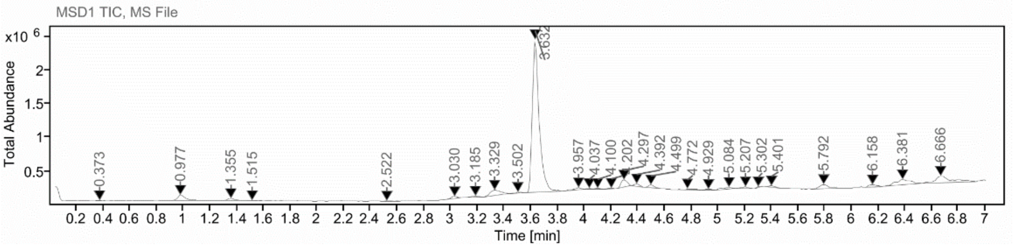

**Sample Purity**

Signal Description DAD1 C, Sig=320,150 Ref=off

| Sample Name | Name | RT | Width | Area | Area% | Height |
|---|---|---|---|---|---|---|
| FGC355 | | 3.607 | 0.028 | 228.5287 | 95.64 | 110.8743 |
| FGC355 | | 4.409 | 0.050 | 5.2438 | 2.19 | 2.5940 |
| FGC355 | | 5.748 | 0.034 | 5.1825 | 2.17 | 2.6902 |

Max Area% 95.637

UV Signal Purity>95% Pass

**UV Apex Spectra**

**RT:**

3.607

**Sample Name:**

FGC355

**Signal Name:**

DAD1C

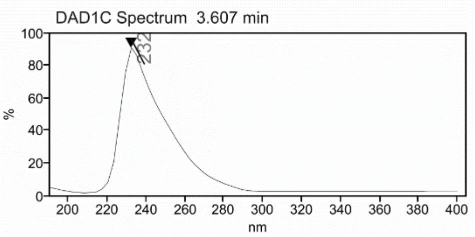

MSD1TIC

**RT:**

3.632

**Sample Name:**

FGC355

**Signal Name:**

MSD1TIC

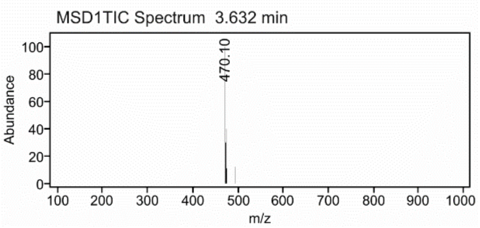

### Mass Spectrum SmartFormula Report

**Analysis Info**

Acquisition Date 2/27/2024 2:46:30 PM

Analysis Name D:\Data\MS_Service\Service_Kaiser\2024_Q1\20240227_SKN_FGC355_100x_5uL-m.d
Method DirectInfusion_WithFlow_Tune_pos.m
Sample Name Test1
Comment

Operator BDAL@DE
Instrument micrOTOF-Q 228888.10407

**Acquisition Parameter**

| | | | | | |
|---|---|---|---|---|---|
| Source Type | ESI | Ion Polarity | Positive | Set Nebulizer | 1.8 Bar |
| Focus | Active | Set Capillary | 4000 V | Set Dry Heater | 320 °C |
| Scan Begin | 50 m/z | Set End Plate Offset | -500 V | Set Dry Gas | 8.0 l/min |
| Scan End | 1600 m/z | Set Collision Cell RF | 150.0 Vpp | Set Divert Valve | Waste |

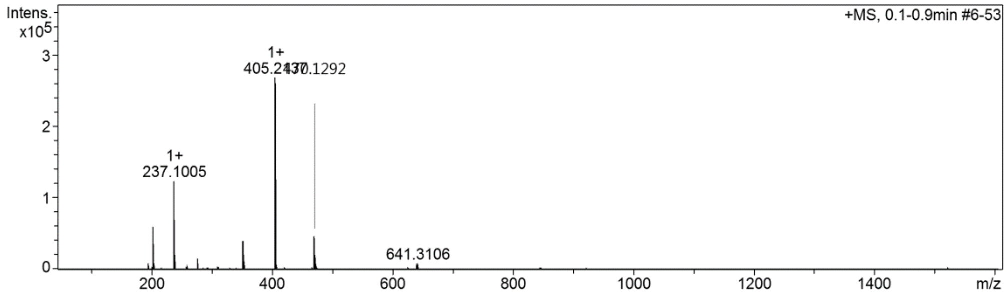

**Cmpd 36**

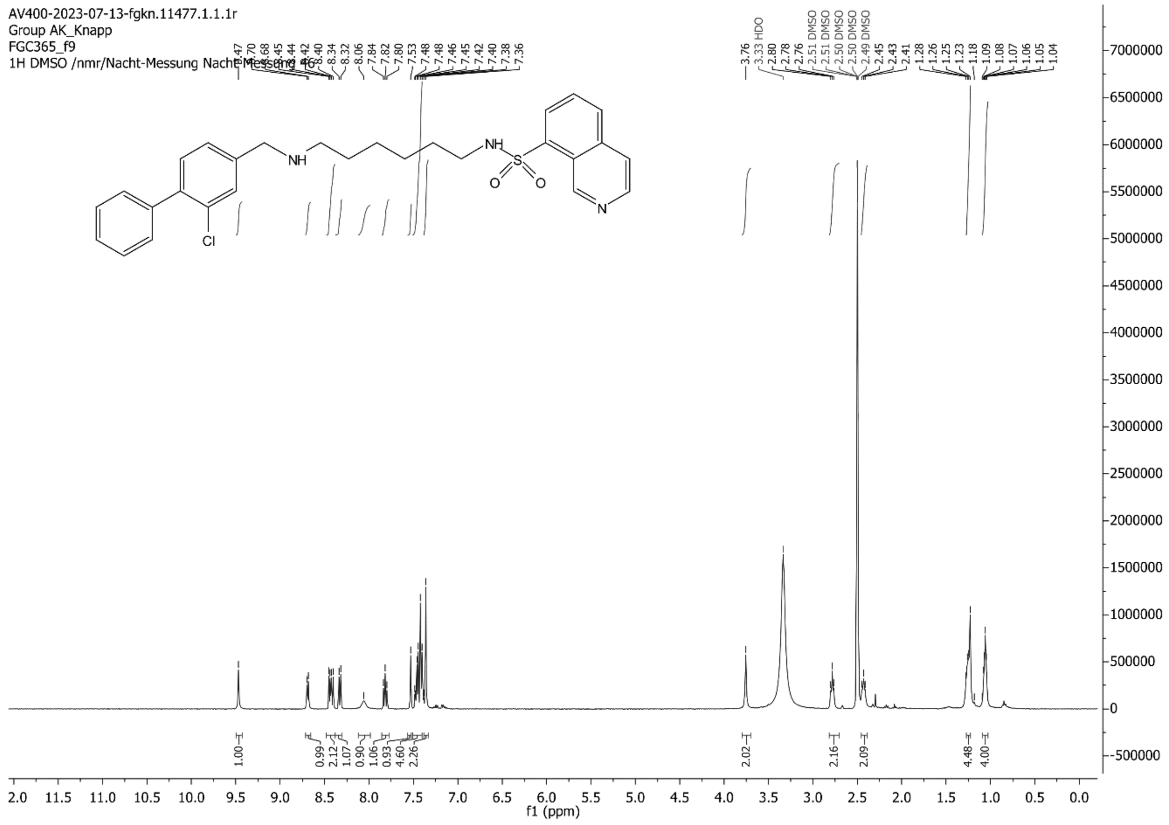
AV400-2023-07-13-fgkn.11477.1.1.1r
Group AK_Knapp
FGC365_f9
1H DMSO /nmr/Nacht-Messung Nacht-Messung 46
NH
NH
S
O
O
N
Cl
f1 (ppm)

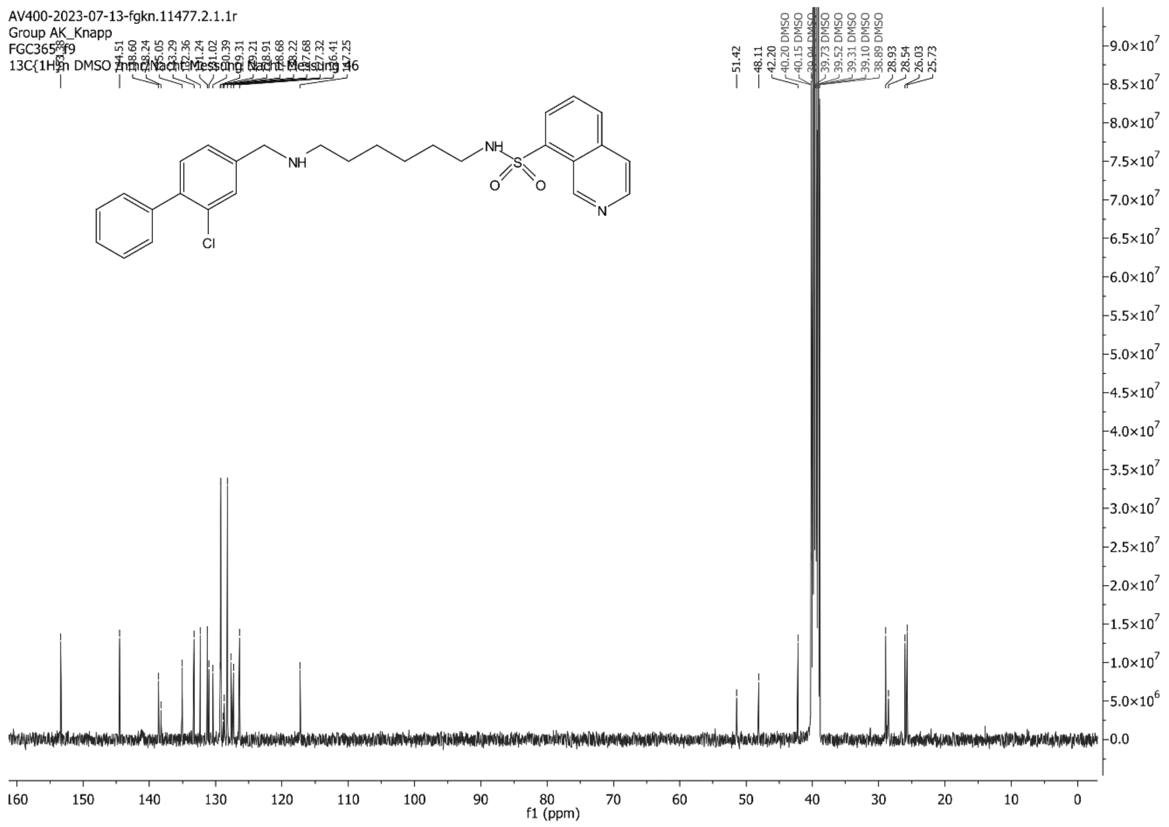
AV400-2023-07-13-fgkn.11477.2.1.1r
Group AK_Knapp
FGC365_f9
13C{1H} in DMSO /nmr/Nacht-Messung Nacht-Messung 46
NH
NH
S
O
O
N
Cl
f1 (ppm)

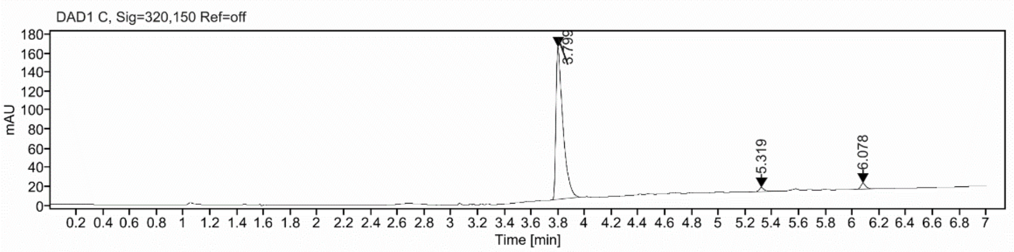

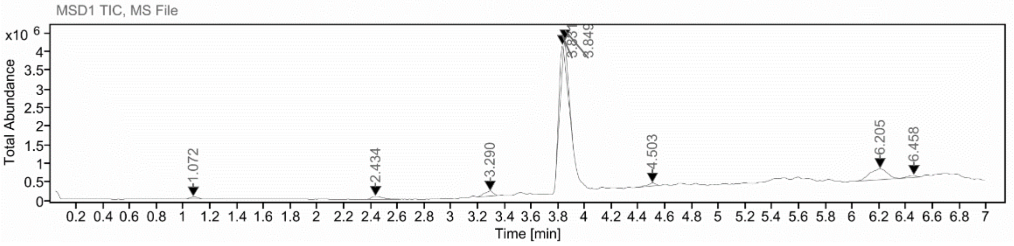

**Sample Purity**

| | |
|---|---|
| Signal Description | DAD1 C, Sig=320,150 Ref=off |

| Sample Name | Name | RT | Width | Area | Area% | Height |
|---|---|---|---|---|---|---|
| FGC365 | | 3.799 | 0.054 | 584.2643 | 95.51 | 159.6700 |
| FGC365 | | 5.319 | 0.037 | 10.8664 | 1.78 | 4.1979 |
| FGC365 | | 6.078 | 0.040 | 16.6153 | 2.72 | 6.1131 |

| | |
|---|---|
| Max Area% | 95.508 |
| UV Signal Purity>95% | Pass |

**UV Apex Spectra**

**RT:**

3.799

**Sample Name:**

FGC365

**Signal Name:**

DAD1C

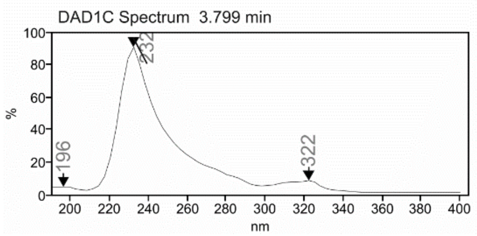

**RT:**

3.849

**Sample Name:**

FGC365

**Signal Name:**

MSD1TIC

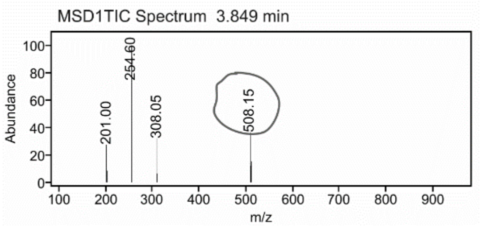

### Mass Spectrum SmartFormula Report

**Analysis Info**

Acquisition Date 2/27/2024 2:53:42 PM

Analysis Name D:\Data\MS_Service\Service_Kaiser\2024_Q1\20240227_SKN_FGC365_100x_5uL-m.d
Method DirectInfusion_WithFlow_Tune_pos.m
Sample Name Test1
Comment

Operator BDAL@DE
Instrument micrOTOF-Q 228888.10407

**Acquisition Parameter**

| | | | | | |
|---|---|---|---|---|---|
| Source Type | ESI | Ion Polarity | Positive | Set Nebulizer | 1.8 Bar |
| Focus | Active | Set Capillary | 4000 V | Set Dry Heater | 320 °C |
| Scan Begin | 50 m/z | Set End Plate Offset | -500 V | Set Dry Gas | 8.0 l/min |
| Scan End | 1600 m/z | Set Collision Cell RF | 150.0 Vpp | Set Divert Valve | Waste |

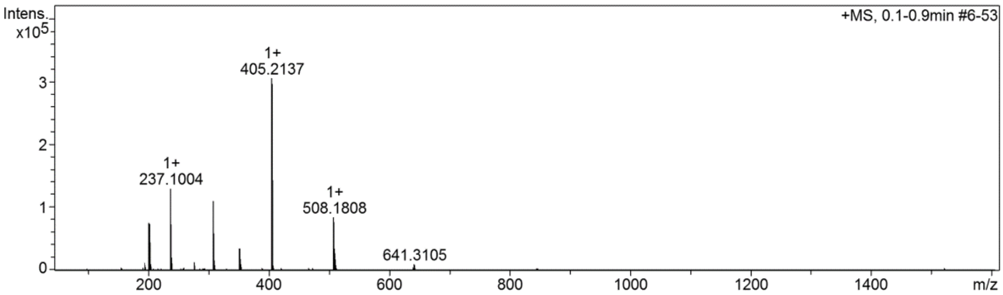

**Cmpd 37**

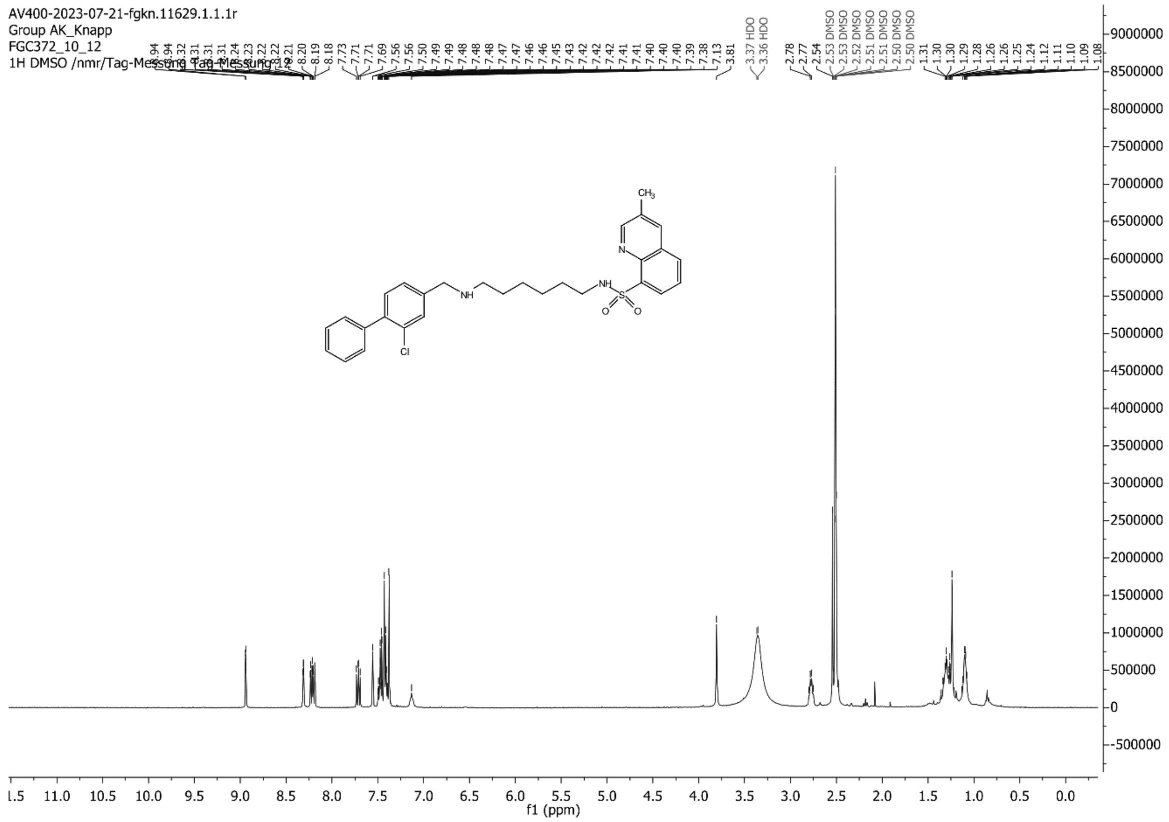

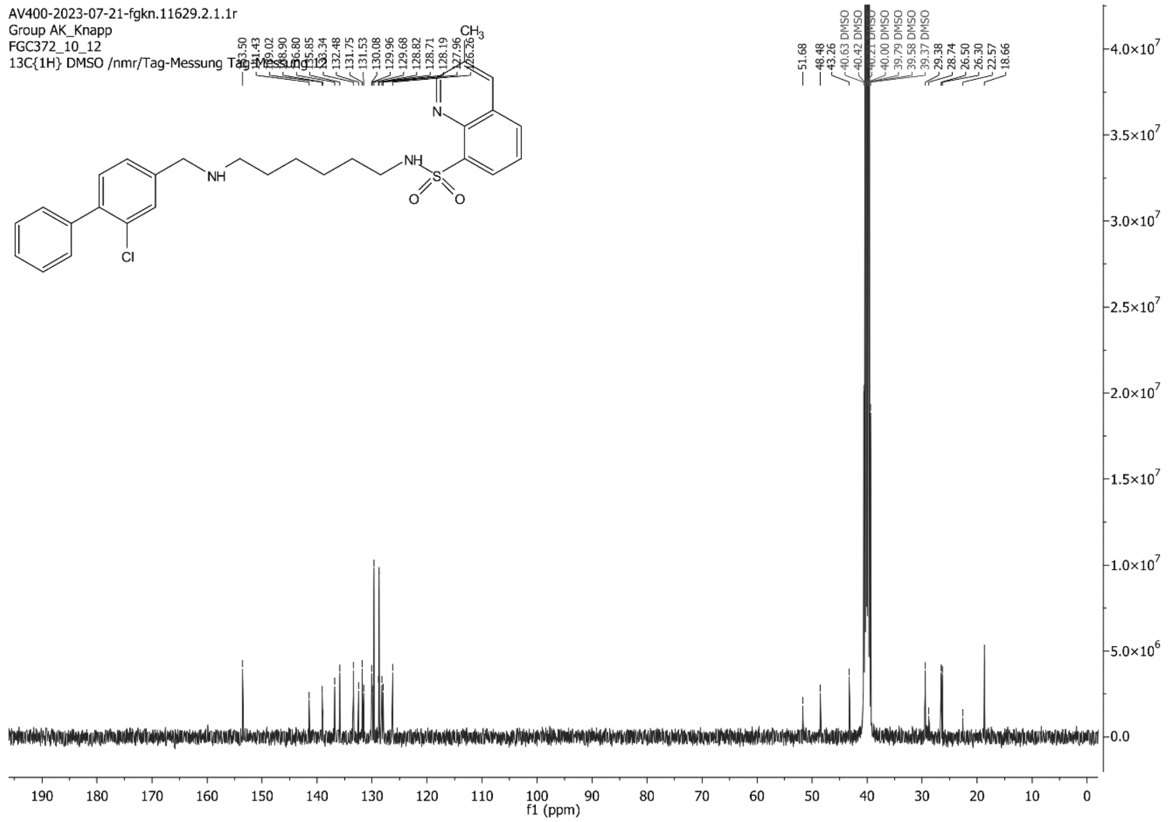

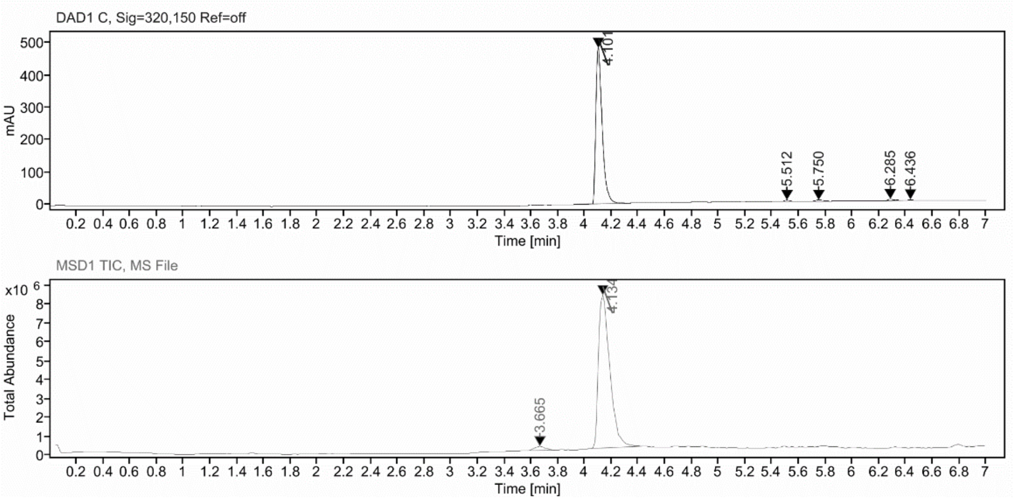

**Sample Purity**

Signal Description DAD1 C, Sig=320,150 Ref=off

| Sample Name | Name | RT | Width | Area | Area% | Height |
|---|---|---|---|---|---|---|
| FGC372 | | 4.101 | 0.050 | 1601.8835 | 96.90 | 483.7339 |
| FGC372 | | 5.512 | 0.039 | 10.7758 | 0.65 | 4.1338 |
| FGC372 | | 5.750 | 0.058 | 15.2681 | 0.92 | 4.2834 |
| FGC372 | | 6.285 | 0.045 | 17.8695 | 1.08 | 5.0284 |
| FGC372 | | 6.436 | 0.038 | 7.3130 | 0.44 | 2.9347 |

Max Area% 96.901

UV Signal Purity>95% Pass

**UV Apex Spectra**

**RT:**

4.101

**Sample Name:**

FGC372

**Signal Name:**

DAD1C

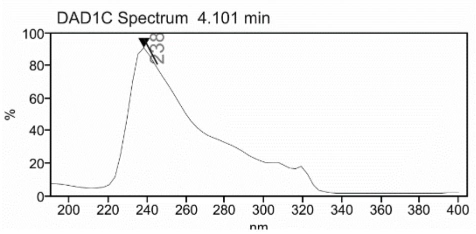

**RT:**

4.134

**Sample Name:**

FGC372

**Signal Name:**

MSD1TIC

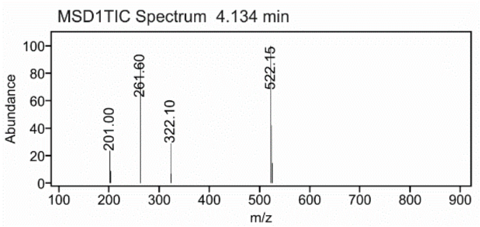

### Mass Spectrum SmartFormula Report

**Analysis Info**

| | | | |
|---|---|---|---|
| | | Acquisition Date | 2/27/2024 3:24:05 PM |
| Analysis Name | D:\Data\MS_Service\Service_Kaiser\2024_Q1\20240227_SKN_FGC372_100x_5uL-m.d | | |
| Method | DirectInfusion_WithFlow_Tune_pos.m | Operator | BDAL@DE |
| Sample Name | Test1 | Instrument | micrOTOF-Q 228888.10407 |
| Comment | | | |

**Acquisition Parameter**

| | | | | | |
|---|---|---|---|---|---|
| Source Type | ESI | Ion Polarity | Positive | Set Nebulizer | 1.8 Bar |
| Focus | Active | Set Capillary | 4000 V | Set Dry Heater | 320 °C |
| Scan Begin | 50 m/z | Set End Plate Offset | -500 V | Set Dry Gas | 8.0 l/min |
| Scan End | 1600 m/z | Set Collision Cell RF | 150.0 Vpp | Set Divert Valve | Waste |

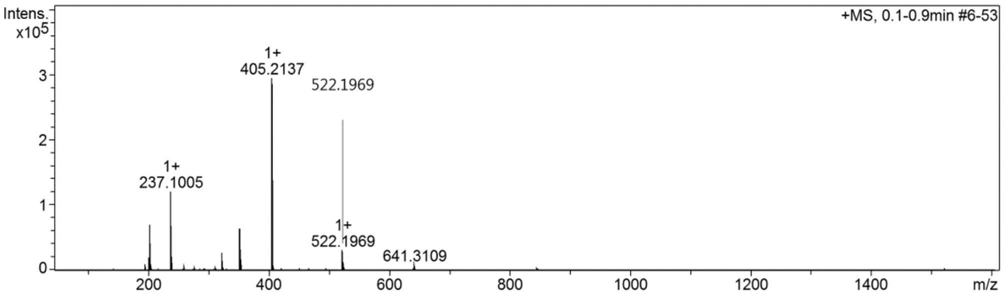

**Cmpd 38**

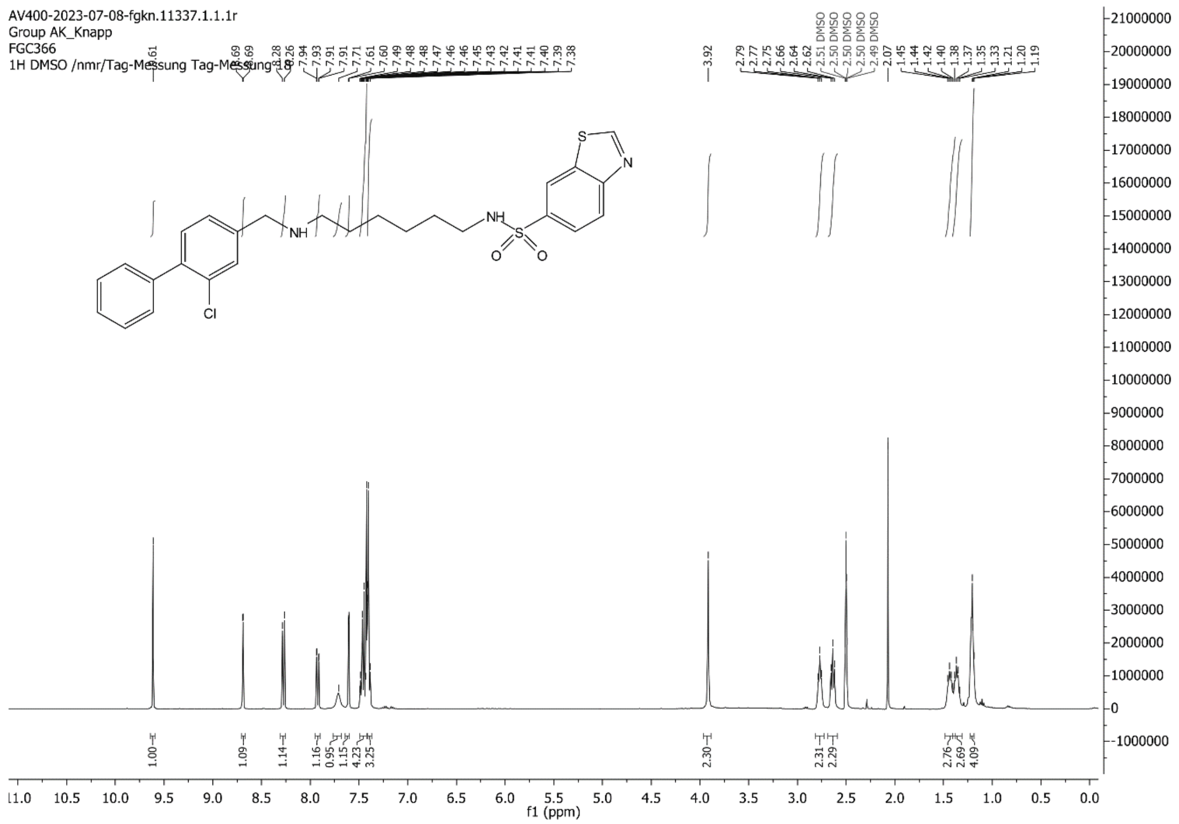
AV400-2023-07-08-fgkn.11337.1.1.1r
Group AK_Knapp
FGC366
1H DMSO /nmr/Tag-Messung Tag-Messung 18
f1 (ppm)

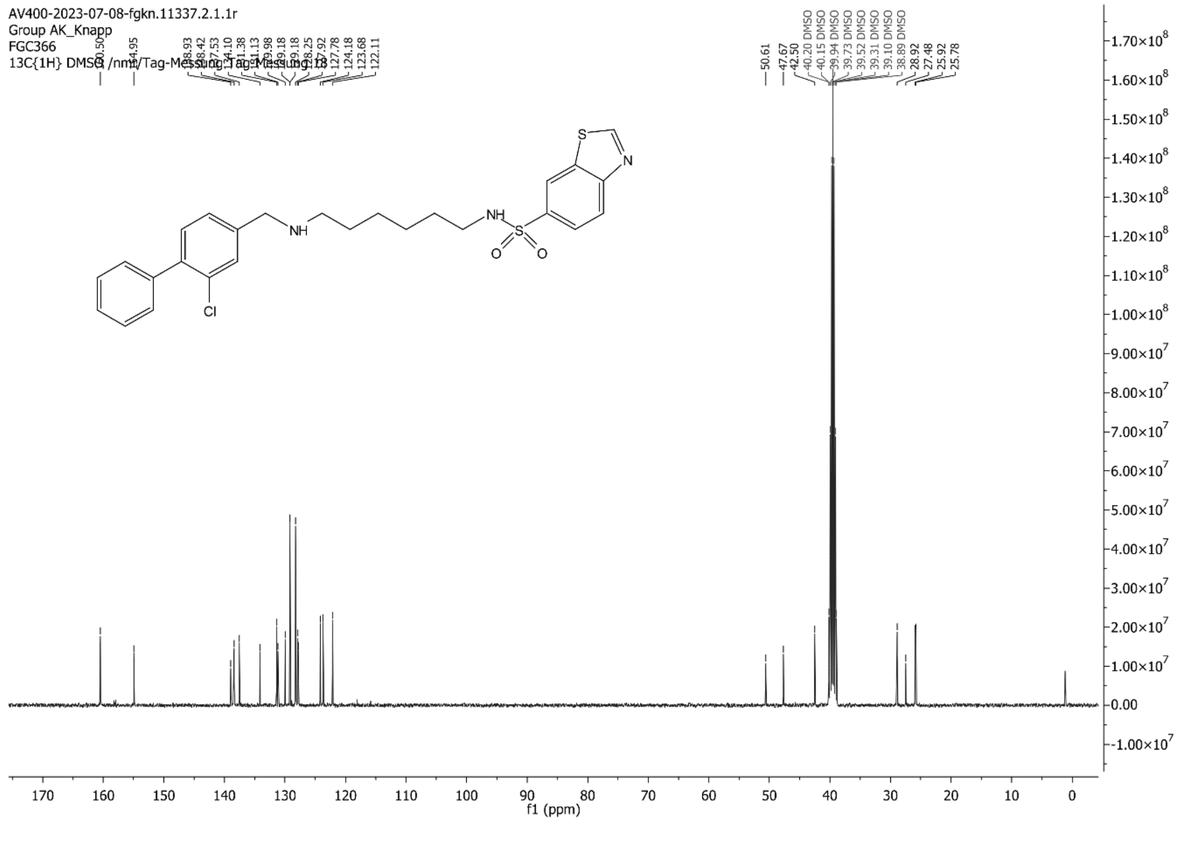
AV400-2023-07-08-fgkn.11337.2.1.1r
Group AK_Knapp
FGC366
13C{1H} DMSO /nmr/Tag-Messung Tag-Messung 18
f1 (ppm)

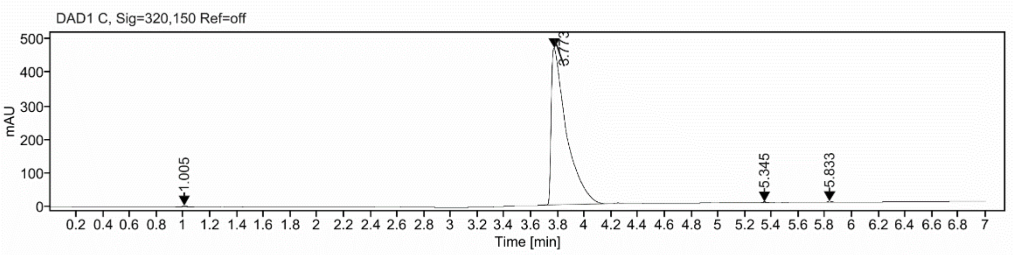

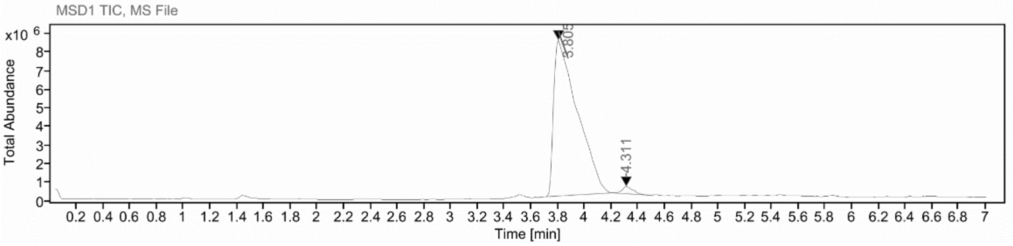

**Sample Purity**

Signal Description DAD1 C, Sig=320,150 Ref=off

| Sample Name | Name | RT | Width | Area | Area% | Height |
|---|---|---|---|---|---|---|
| FGC366_17 | | 1.005 | 0.040 | 10.0783 | 0.27 | 2.9575 |
| FGC366_17 | | 3.773 | 0.109 | 3688.9722 | 99.40 | 468.3583 |
| FGC366_17 | | 5.345 | 0.036 | 5.4639 | 0.15 | 2.3324 |
| FGC366_17 | | 5.833 | 0.037 | 6.8227 | 0.18 | 2.7630 |

| | |
|---|---|
| Max Area% | 99.397 |
| UV Signal Purity>95% | Pass |

**UV Apex Spectra**

**RT:**

3.773

**Sample Name:**

FGC366_17

**Signal Name:**

DAD1C

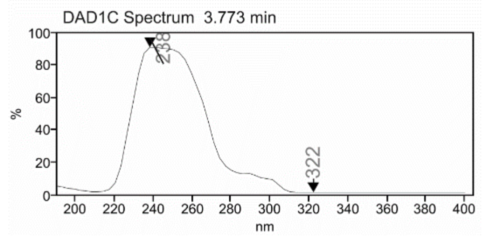

**RT:**

3.805

**Sample Name:**

FGC366_17

**Signal Name:**

MSD1TIC

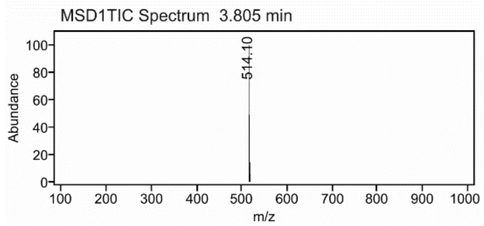

### Mass Spectrum SmartFormula Report

**Analysis Info**

| | | | |
|---|---|---|---|
| Analysis Name | D:\Data\MS_Service\Service_Kaiser\2024_Q1\20240227_SKN_FGC366_100x_5uL-m.d | Acquisition Date | 2/27/2024 3:01:02 PM |
| Method | DirectInfusion_WithFlow_Tune_pos.m | Operator | BDAL@DE |
| Sample Name | Test1 | Instrument | micrOTOF-Q 228888.10407 |
| Comment | | | |

**Acquisition Parameter**

| | | | | | |
|---|---|---|---|---|---|
| Source Type | ESI | Ion Polarity | Positive | Set Nebulizer | 1.8 Bar |
| Focus | Active | Set Capillary | 4000 V | Set Dry Heater | 320 °C |
| Scan Begin | 50 m/z | Set End Plate Offset | -500 V | Set Dry Gas | 8.0 l/min |
| Scan End | 1600 m/z | Set Collision Cell RF | 150.0 Vpp | Set Divert Valve | Waste |

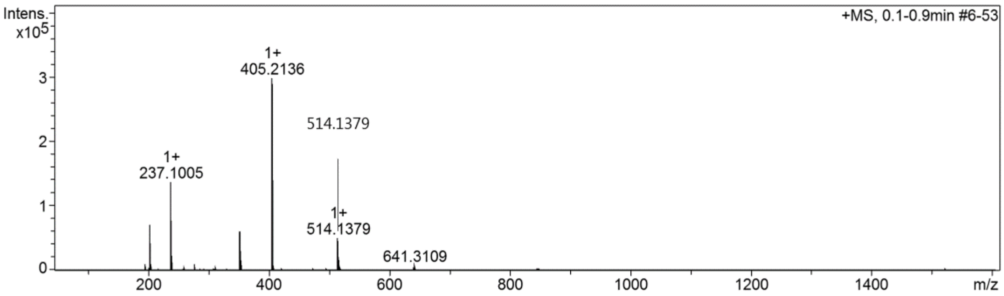

#### Cmpd 39

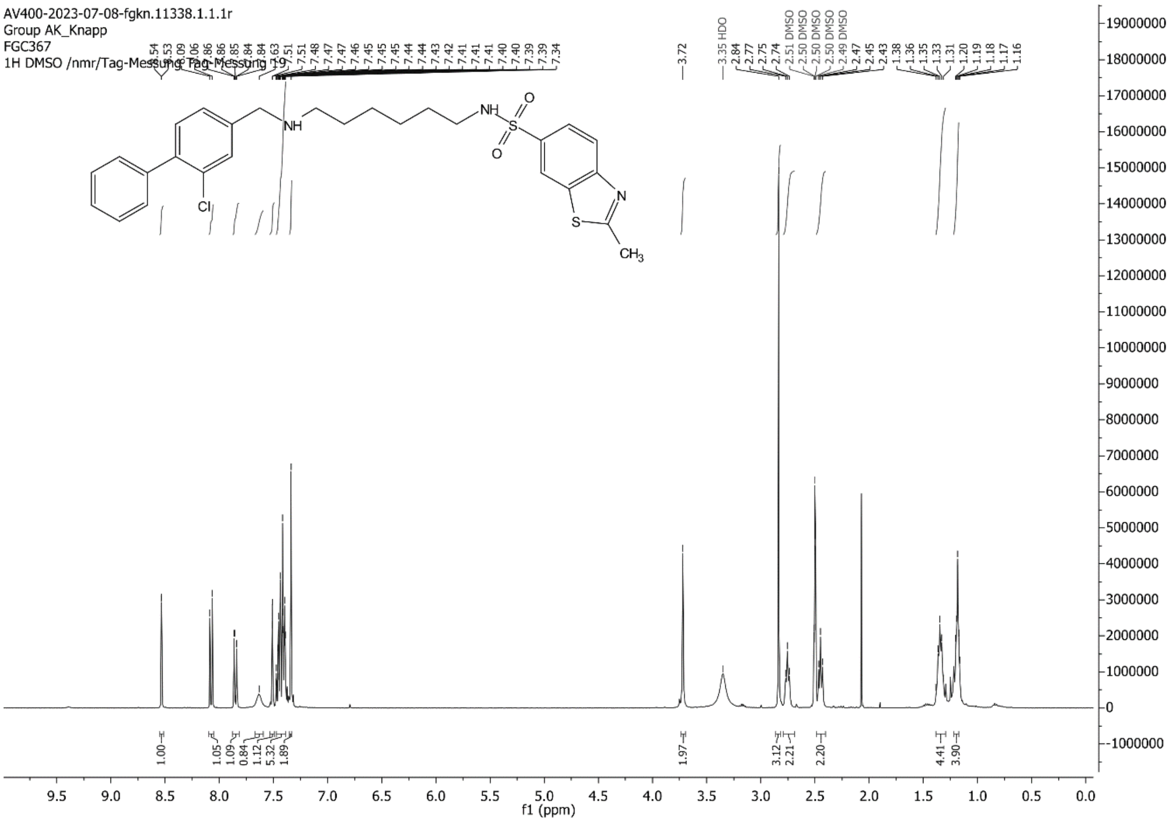

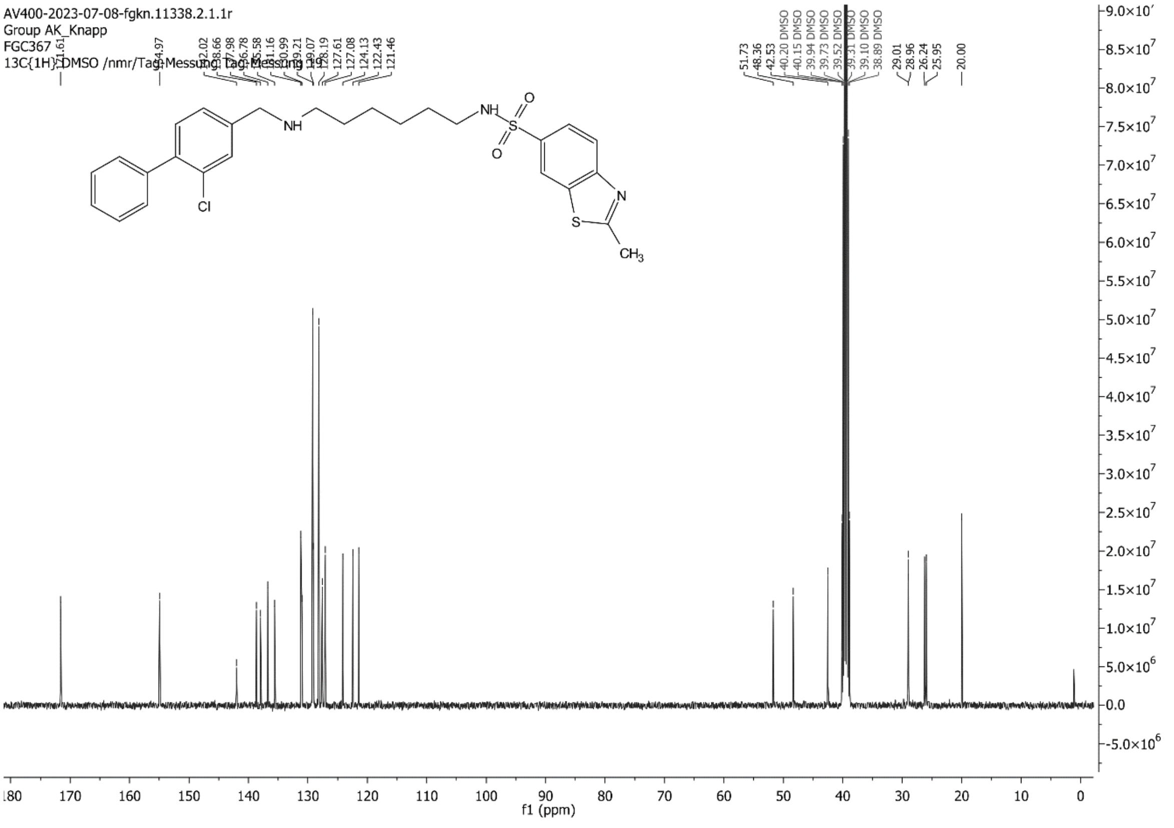

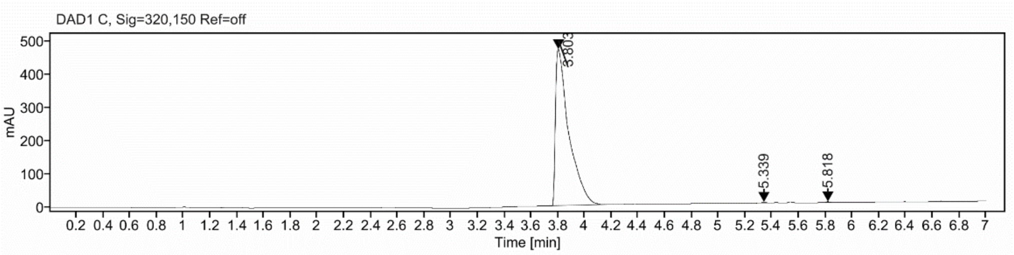

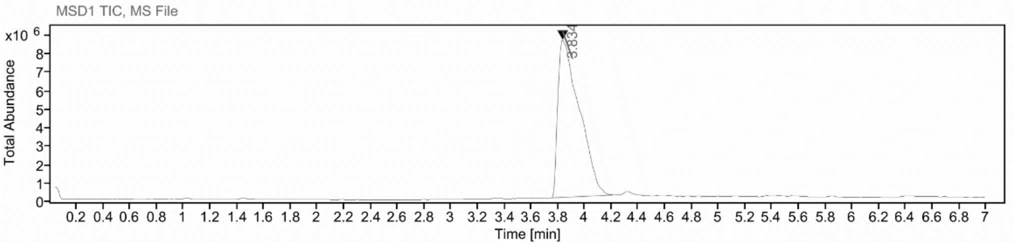

**Sample Purity**

Signal Description DAD1 C, Sig=320,150 Ref=off

| Sample Name | Name | RT | Width | Area | Area% | Height |
|---|---|---|---|---|---|---|
| FGC367_fr3 | | 3.803 | 0.094 | 3272.2114 | 99.59 | 472.2797 |
| FGC367_fr3 | | 5.339 | 0.037 | 6.1085 | 0.19 | 2.6223 |
| FGC367_fr3 | | 5.818 | 0.039 | 7.2268 | 0.22 | 2.7107 |

Max Area% 99.594

UV Signal Purity>95% Pass

**UV Apex Spectra**

**RT:**

3.803

**Sample Name:**

FGC367_fr3

**Signal Name:**

DAD1C

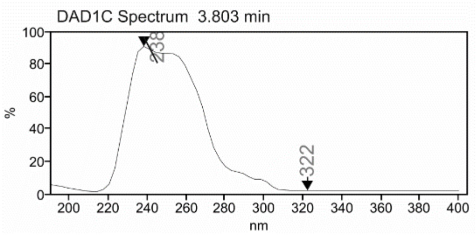

**RT:**

3.834

**Sample Name:**

FGC367_fr3

**Signal Name:**

MSD1TIC

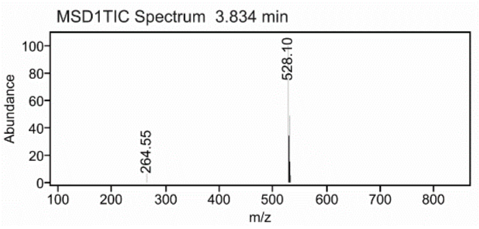

### Mass Spectrum SmartFormula Report

**Analysis Info**

| | | | |
|---|---|---|---|
| | | Acquisition Date | 2/27/2024 3:10:13 PM |
| Analysis Name | D:\Data\MS_Service\Service_Kaiser\2024_Q1\20240227_SKN_FGC367_100x_5uL-m.d | | |
| Method | DirectInfusion_WithFlow_Tune_pos.m | Operator | BDAL@DE |
| Sample Name | Test1 | Instrument | micrOTOF-Q 228888.10407 |
| Comment | | | |

**Acquisition Parameter**

| | | | | | |
|---|---|---|---|---|---|
| Source Type | ESI | Ion Polarity | Positive | Set Nebulizer | 1.8 Bar |
| Focus | Active | Set Capillary | 4000 V | Set Dry Heater | 320 °C |
| Scan Begin | 50 m/z | Set End Plate Offset | -500 V | Set Dry Gas | 8.0 l/min |
| Scan End | 1600 m/z | Set Collision Cell RF | 150.0 Vpp | Set Divert Valve | Waste |

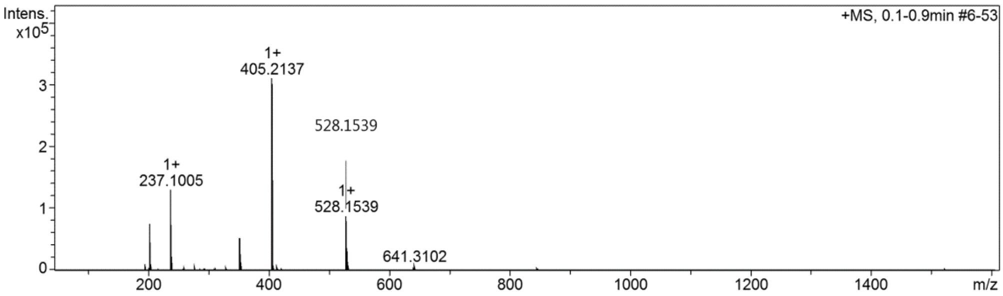

**Cmpd 40**

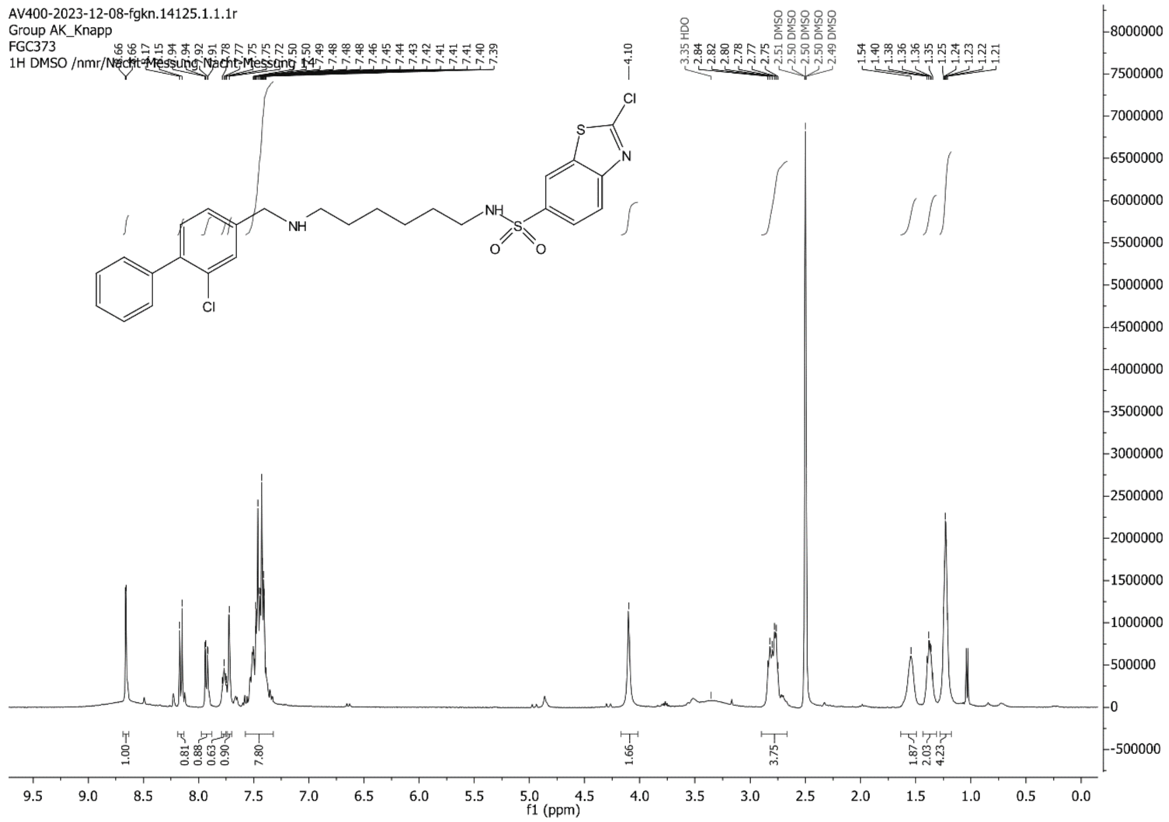
AV400-2023-12-08-fgkn.14125.1.1.1r
Group AK_Knapp
FGC373
1H DMSO /nmr/Nacht-Messung Nacht-Messung 14
Cl
S
N
NH
NH
O
S
O
Cl
f1 (ppm)

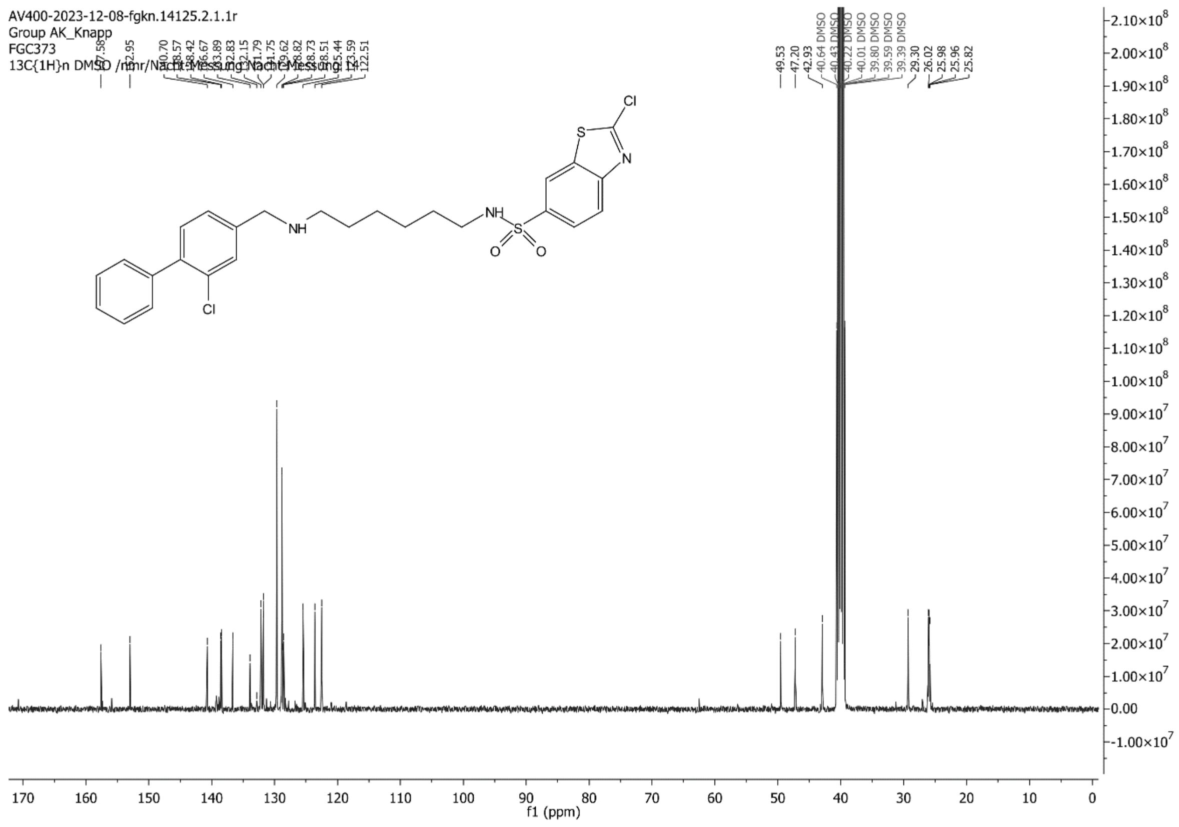
AV400-2023-12-08-fgkn.14125.2.1.1r
Group AK_Knapp
FGC373
13C{1H}n DMSO /nmr/Nacht-Messung Nacht-Messung 14
Cl
S
N
NH
NH
O
S
O
Cl
f1 (ppm)

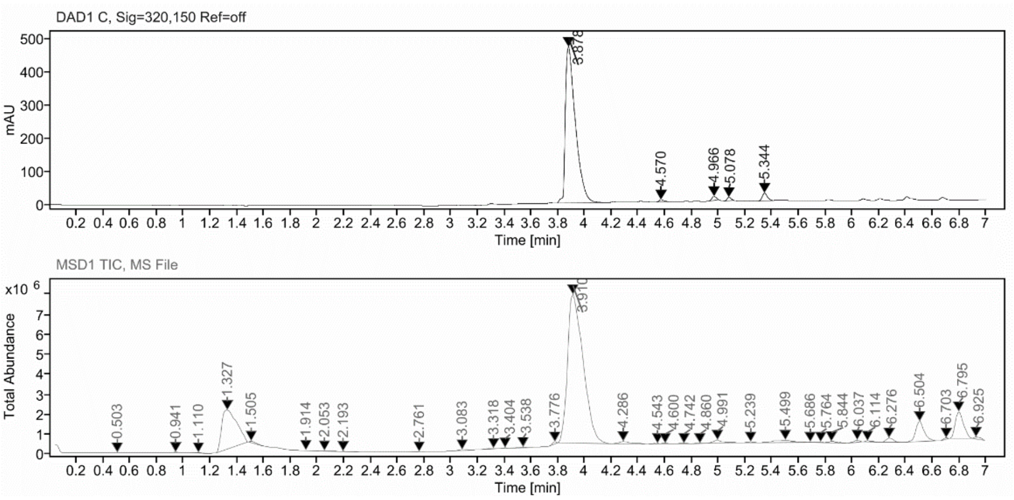

**Sample Purity**

Signal Description DAD1 C, Sig=320,150 Ref=off

| Sample Name | Name | RT | Width | Area | Area% | Height |
|---|---|---|---|---|---|---|
| FGC373_on | | 3.878 | 0.080 | 2457.3611 | 95.13 | 470.4305 |
| FGC373_on | | 4.570 | 0.039 | 15.7546 | 0.61 | 6.4380 |
| FGC373_on | | 4.966 | 0.036 | 28.0597 | 1.09 | 12.8505 |
| FGC373_on | | 5.078 | 0.033 | 20.1027 | 0.78 | 10.2239 |
| FGC373_on | | 5.344 | 0.040 | 61.7932 | 2.39 | 23.8183 |

Max Area% 95.133

UV Signal Purity>95% Pass

**UV Apex Spectra**

**RT:**

3.878

**Sample Name:**

FGC373_on

**Signal Name:**

DAD1C

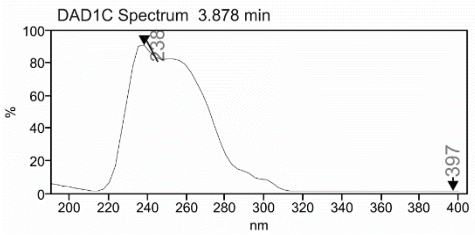

**RT:**

3.910

**Sample Name:**

FGC373_on

**Signal Name:**

MSD1TIC

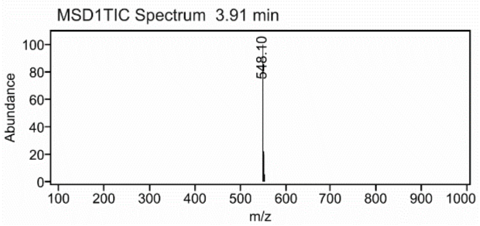

### Mass Spectrum SmartFormula Report

**Analysis Info**

| | | | |
|---|---|---|---|
| Analysis Name | D:\Data\MS_Service\Service_Kaiser\2024_Q1\20240227_SKN_FGC373_100x_5uL-m.d | Acquisition Date | 2/27/2024 3:30:33 PM |
| Method | DirectInfusion_WithFlow_Tune_pos.m | Operator | BDAL@DE |
| Sample Name | Test1 | Instrument | micrOTOF-Q 228888.10407 |
| Comment | | | |

**Acquisition Parameter**

| | | | | | |
|---|---|---|---|---|---|
| Source Type | ESI | Ion Polarity | Positive | Set Nebulizer | 1.8 Bar |
| Focus | Active | Set Capillary | 4000 V | Set Dry Heater | 320 °C |
| Scan Begin | 50 m/z | Set End Plate Offset | -500 V | Set Dry Gas | 8.0 l/min |
| Scan End | 1600 m/z | Set Collision Cell RF | 150.0 Vpp | Set Divert Valve | Waste |

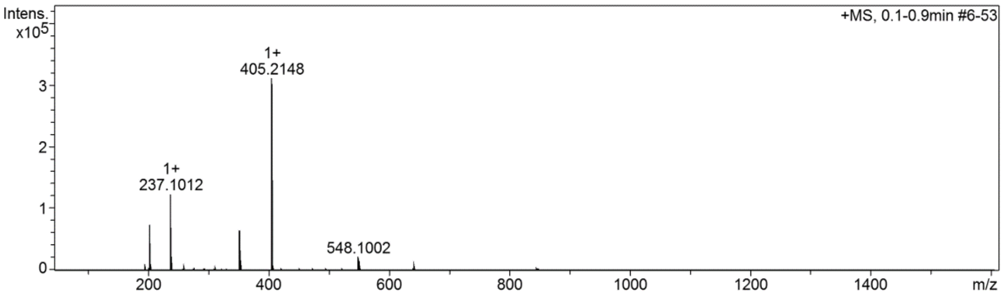

**Cmpd 41**

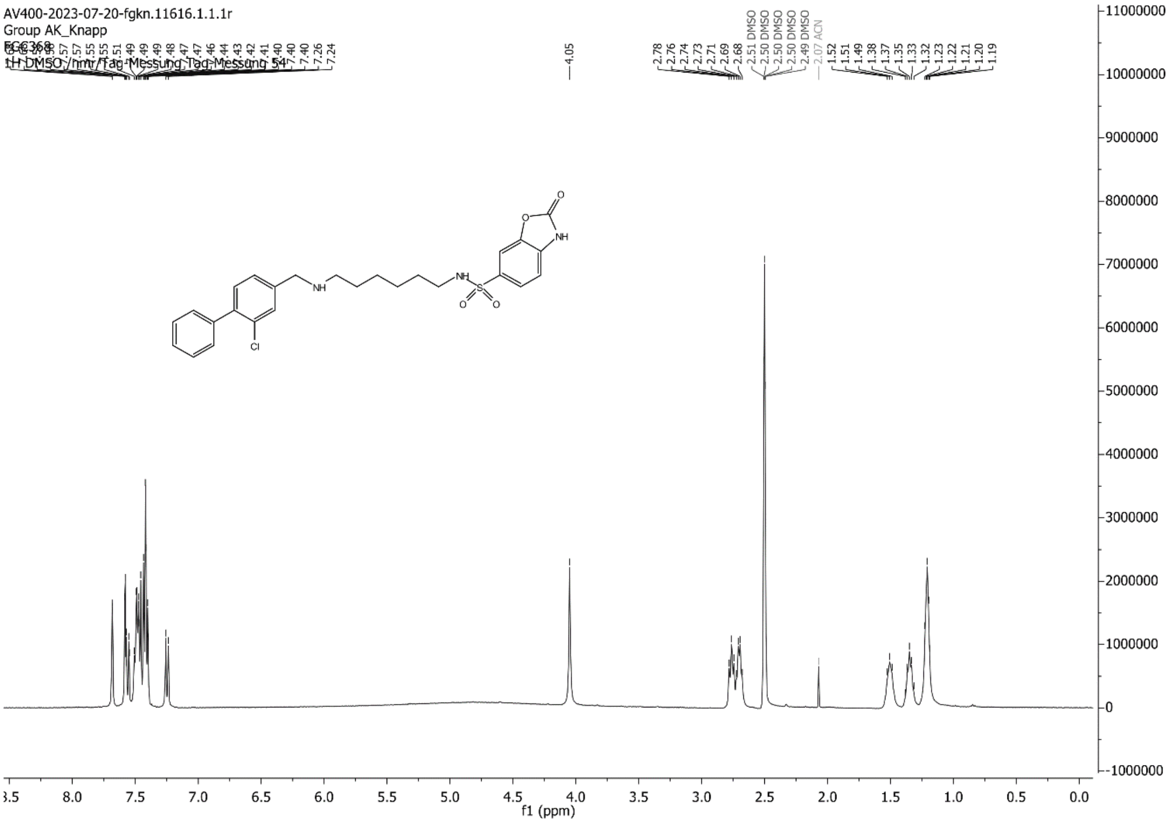

AV400-2023-07-20-fgkn.11616.1.1.1r
Group AK_Knapp
FGC368
1H DMSO /nmr/Tag-Messung Tag-Messung 54
f1 (ppm)

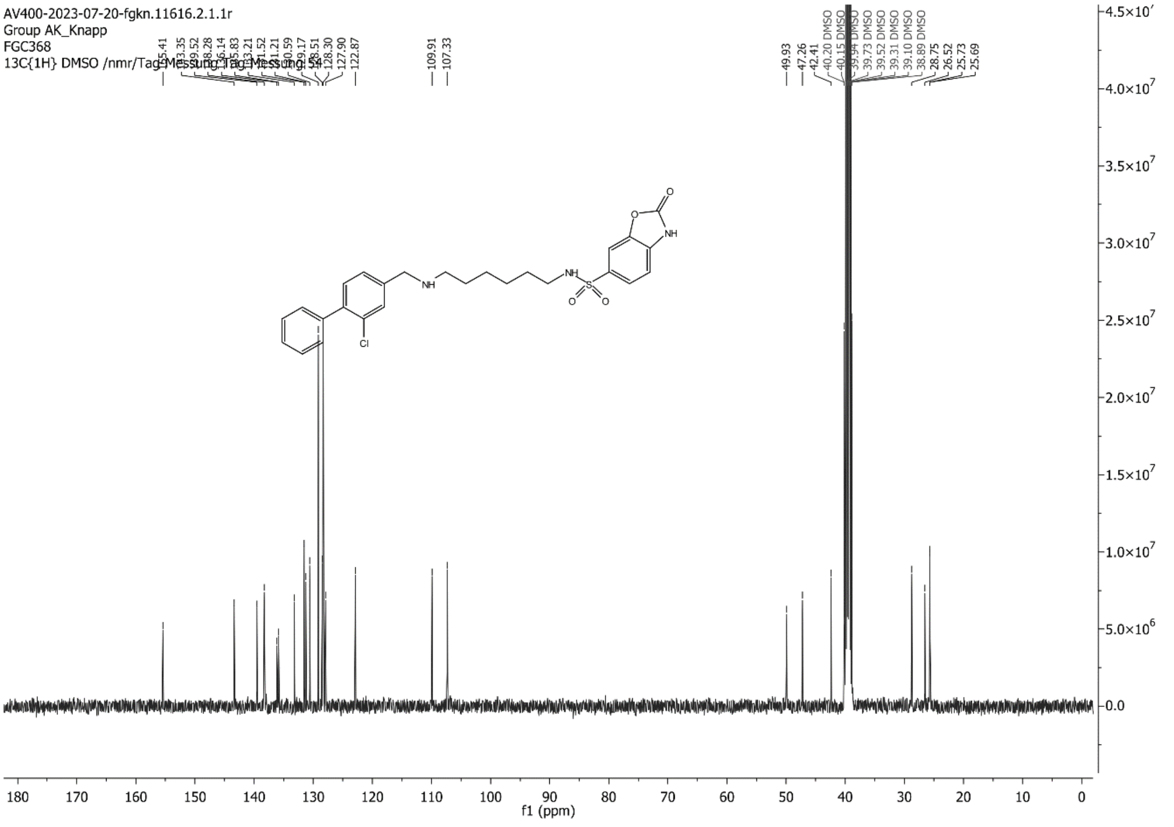

AV400-2023-07-20-fgkn.11616.2.1.1r
Group AK_Knapp
FGC368
13C{1H} DMSO /nmr/Tag-Messung Tag-Messung 54
f1 (ppm)

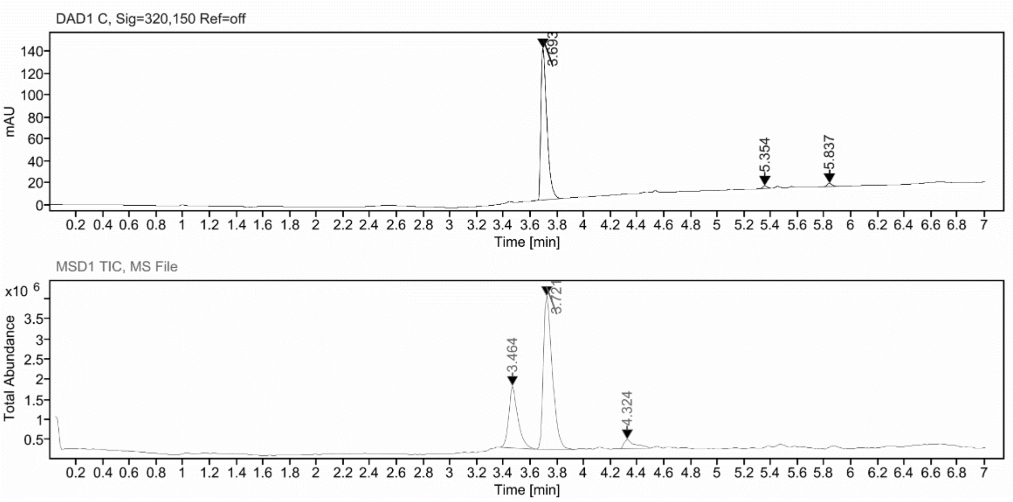

**Sample Purity**

Signal Description DAD1 C, Sig=320,150 Ref=off

| Sample Name | Name | RT | Width | Area | Area% | Height |
|---|---|---|---|---|---|---|
| FGC368_rp_2 | | 3.693 | 0.048 | 434.3026 | 96.94 | 138.6874 |
| FGC368_rp_2 | | 5.354 | 0.037 | 5.6660 | 1.26 | 2.4300 |
| FGC368_rp_2 | | 5.837 | 0.040 | 8.0507 | 1.80 | 3.0536 |

Max Area% 96.938

UV Signal Purity>95% Pass

**UV Apex Spectra**

**RT:**

3.693

**Sample Name:**

FGC368_rp_2

**Signal Name:**

DAD1C

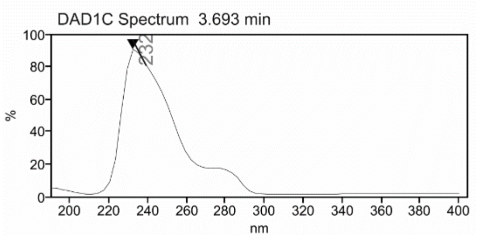

**RT:**

3.721

**Sample Name:**

FGC368_rp_2

**Signal Name:**

MSD1TIC

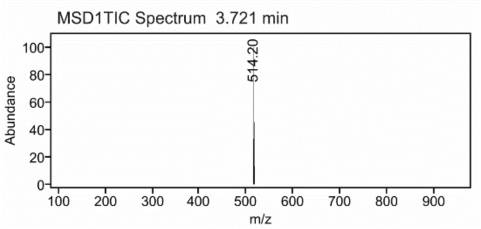

### Mass Spectrum SmartFormula Report

**Analysis Info**

| | | | |
|---|---|---|---|
| Analysis Name | D:\Data\MS_Service\Service_Kaiser\2024_Q1\20240227_SKN_FGC368_100x_5uL-m.d | Acquisition Date | 2/27/2024 3:16:40 PM |
| Method | DirectInfusion_WithFlow_Tune_pos.m | Operator | BDAL@DE |
| Sample Name | Test1 | Instrument | micrOTOF-Q 228888.10407 |
| Comment | | | |

**Acquisition Parameter**

| | | | | | |
|---|---|---|---|---|---|
| Source Type | ESI | Ion Polarity | Positive | Set Nebulizer | 1.8 Bar |
| Focus | Active | Set Capillary | 4000 V | Set Dry Heater | 320 °C |
| Scan Begin | 50 m/z | Set End Plate Offset | -500 V | Set Dry Gas | 8.0 l/min |
| Scan End | 1600 m/z | Set Collision Cell RF | 150.0 Vpp | Set Divert Valve | Waste |

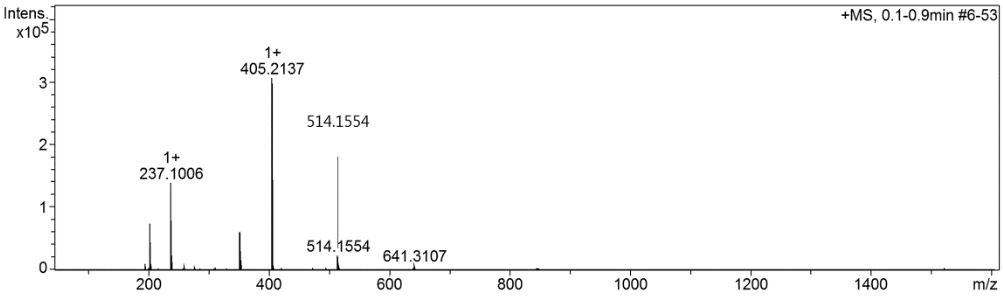

#### Cmpd 42

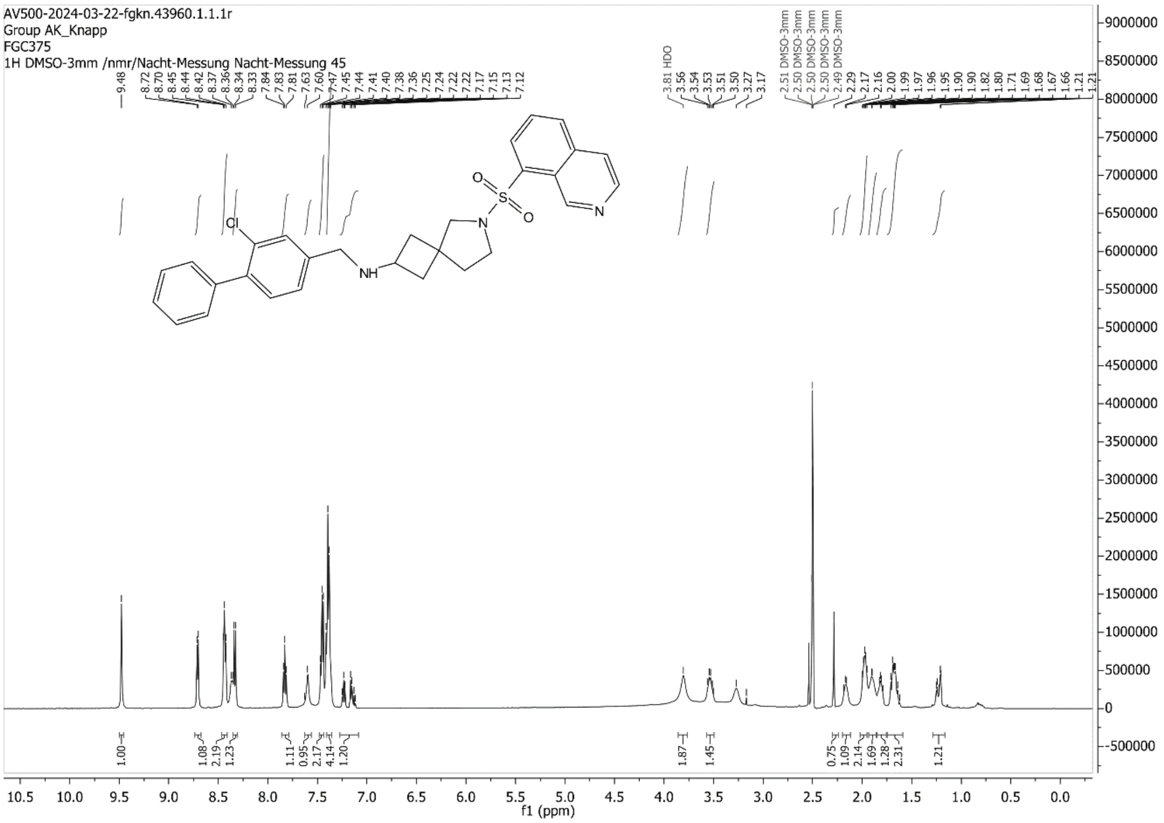

AV500-2024-03-22-fgkn.43960.1.1.1r
Group AK_Knapp
FGC375
1H DMSO-3mm /nmr/Nacht-Messung Nacht-Messung 45
9.48
8.72 8.70 8.45 8.44 8.42 8.37 8.36 8.34 8.33 7.84 7.83 7.81 7.63 7.60 7.47 7.45 7.44 7.41 7.40 7.38 7.36 7.25 7.24 7.22 7.22 7.22 7.17 7.15 7.13 7.12
3.81 HDO 3.56 3.54 3.53 3.51 3.50 3.27 3.17
2.51 DMSO-3mm 2.50 DMSO-3mm 2.50 DMSO-3mm 2.50 DMSO-3mm 2.49 DMSO-3mm 2.29 2.17 2.16 2.00 1.99 1.97 1.96 1.95 1.90 1.90 1.82 1.80 1.71 1.69 1.68 1.67 1.66 1.21
Cl
NH
N
O
S
O
N
1.00 1.08 2.19 1.23 1.11 0.95 2.17 4.14 1.20 1.87 1.45 0.75 1.09 2.14 1.69 1.28 2.31 1.21
f1 (ppm)

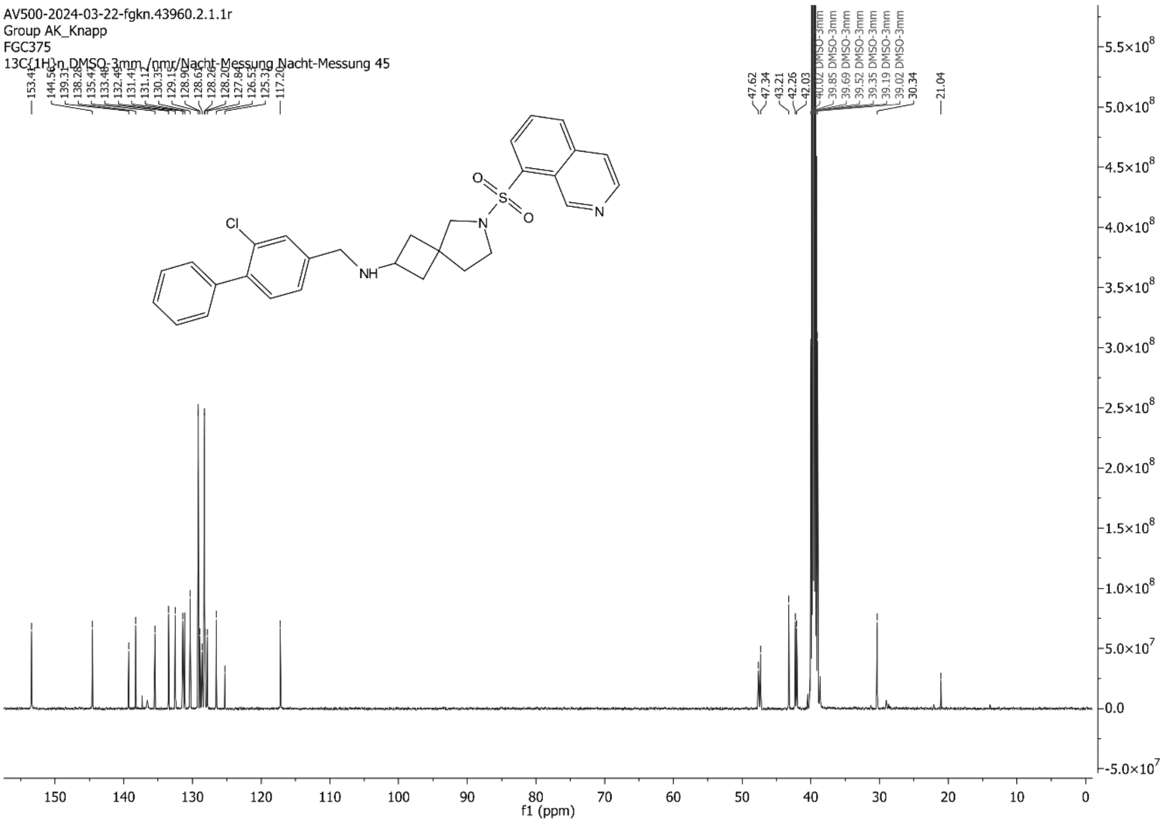

AV500-2024-03-22-fgkn.43960.2.1.1r
Group AK_Knapp
FGC375
13C{1H}n DMSO-3mm /nmr/Nacht-Messung Nacht-Messung 45
153.41 144.56 139.31 138.28 135.47 133.49 132.49 131.41 131.12 130.31 129.15 128.90 128.65 128.26 128.20 127.84 126.53 125.32 117.20
47.62 47.34 43.21 42.26 42.03 40.02 DMSO-3mm 39.85 DMSO-3mm 39.69 DMSO-3mm 39.52 DMSO-3mm 39.35 DMSO-3mm 39.19 DMSO-3mm 39.02 DMSO-3mm 30.34
21.04
Cl
NH
N
O
S
O
N
f1 (ppm)

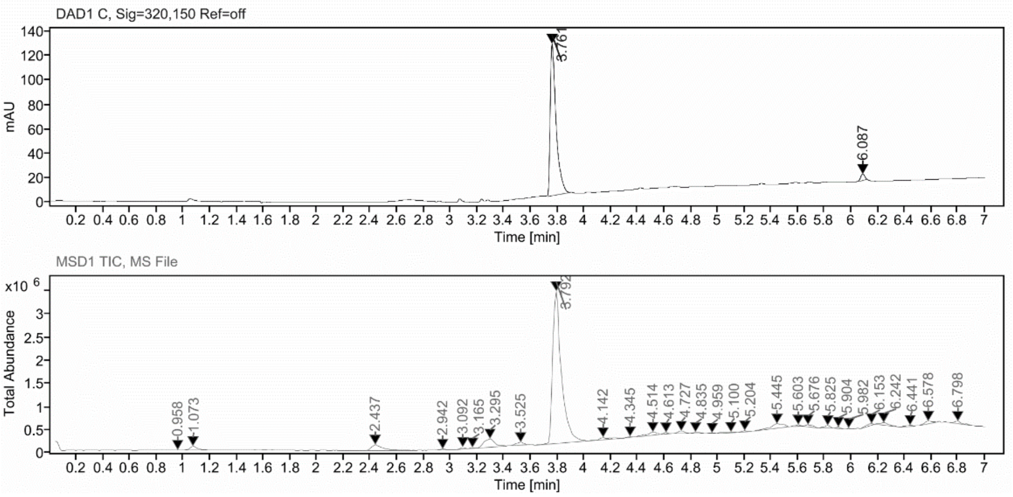

**Sample Purity**

Signal Description DAD1 C, Sig=320,150 Ref=off

| Sample Name | Name | RT | Width | Area | Area% | Height |
|---|---|---|---|---|---|---|
| FGC375 | | 3.761 | 0.044 | 378.1200 | 97.25 | 124.0509 |
| FGC375 | | 6.087 | 0.035 | 10.6874 | 2.75 | 5.3872 |

Max Area% 97.251

UV Signal Purity>95% Pass

**UV Apex Spectra**

**RT:**

3.761

**Sample Name:**

FGC375

**Signal Name:**

DAD1C

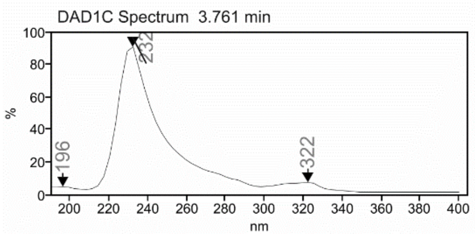

**RT:**

3.792

**Sample Name:**

FGC375

**Signal Name:**

MSD1TIC

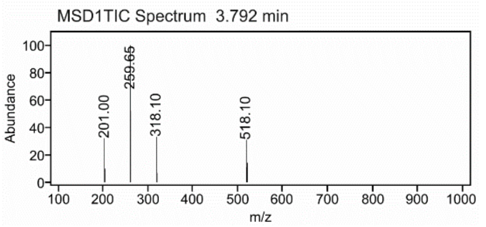

### Mass Spectrum SmartFormula Report

**Analysis Info**

| | | | |
|---|---|---|---|
| Analysis Name | D:\Data\MS_Service\Service_Kaiser\2024_Q1\20240227_SKN_FGC375_100x_5uL-m.d | Acquisition Date | 2/27/2024 3:44:20 PM |
| Method | DirectInfusion_WithFlow_Tune_pos.m | Operator | BDAL@DE |
| Sample Name | Test1 | Instrument | micrOTOF-Q 228888.10407 |
| Comment | | | |

**Acquisition Parameter**

| | | | | | |
|---|---|---|---|---|---|
| Source Type | ESI | Ion Polarity | Positive | Set Nebulizer | 1.8 Bar |
| Focus | Active | Set Capillary | 4000 V | Set Dry Heater | 320 °C |
| Scan Begin | 50 m/z | Set End Plate Offset | -500 V | Set Dry Gas | 8.0 l/min |
| Scan End | 1600 m/z | Set Collision Cell RF | 150.0 Vpp | Set Divert Valve | Waste |

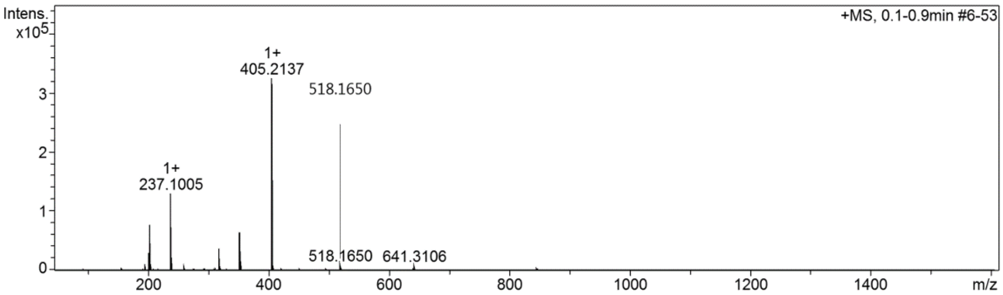

#### Cmpd 43

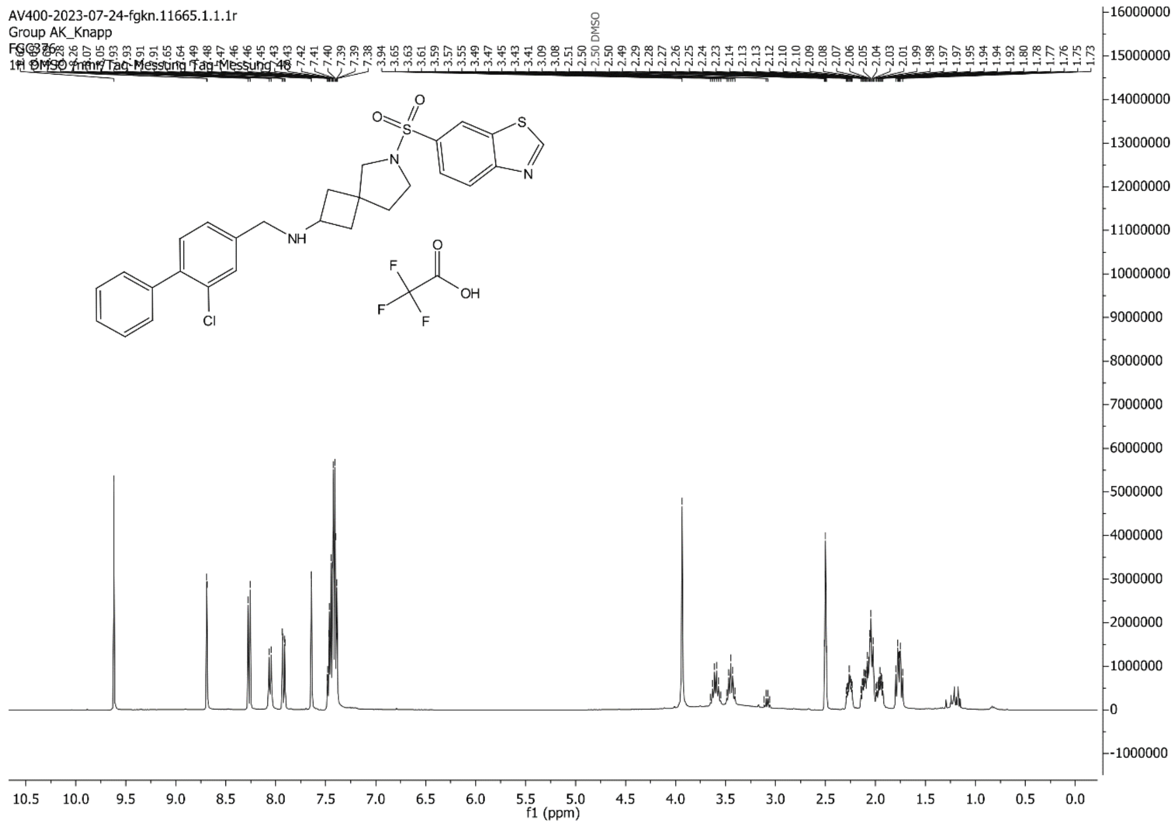

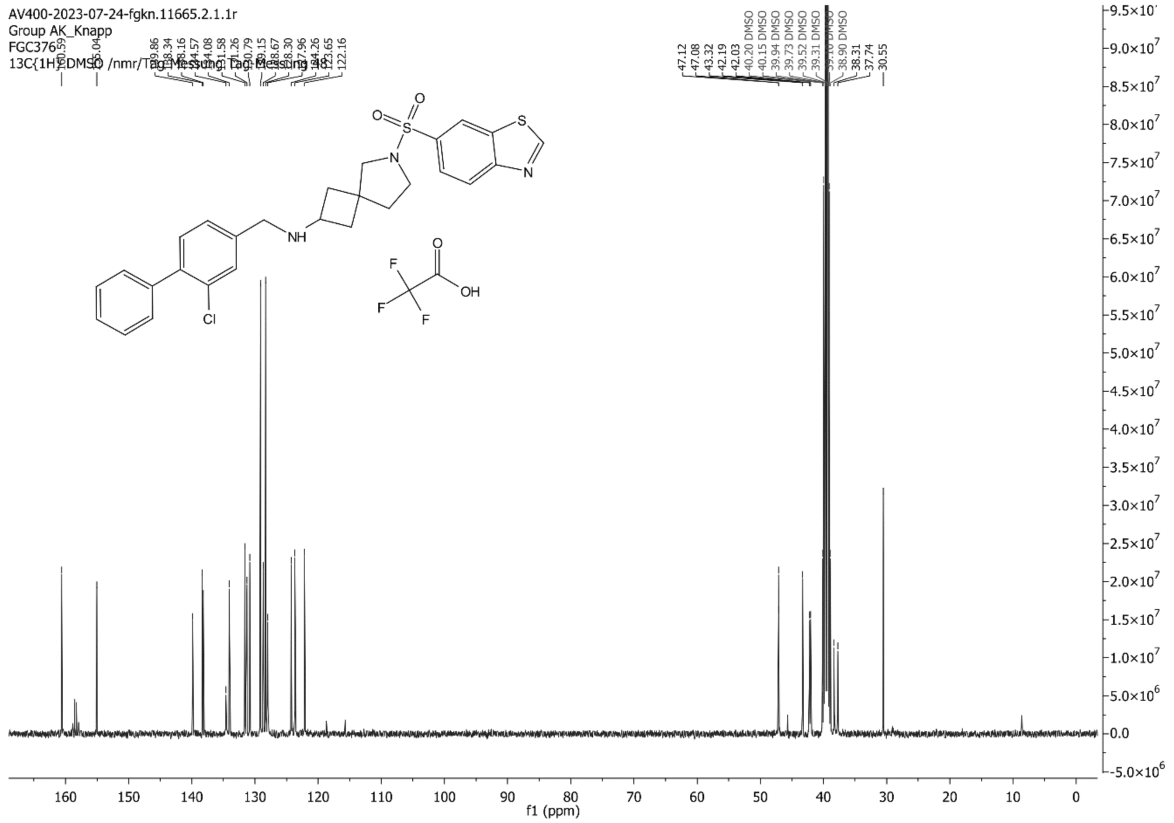

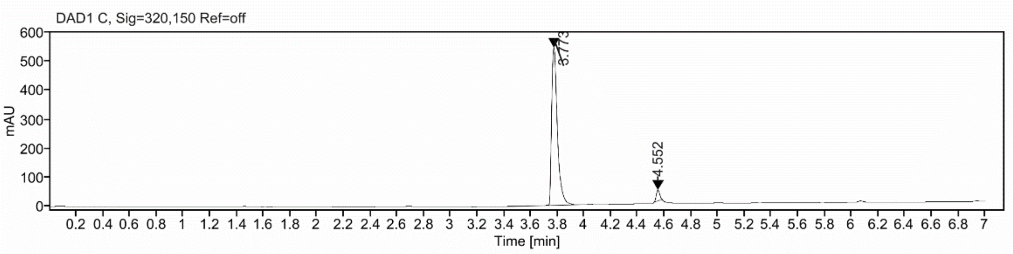

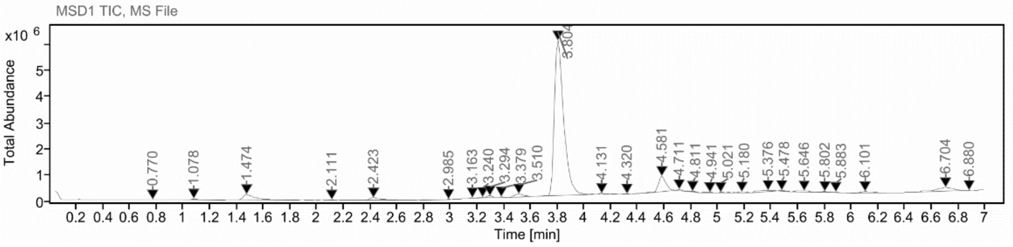

**Sample Purity**

| | |
|---|---|
| Signal Description | DAD1 C, Sig=320,150 Ref=off |

| Sample Name | Name | RT | Width | Area | Area% | Height |
|---|---|---|---|---|---|---|
| FGC376 | | 3.773 | 0.045 | 1626.5195 | 95.92 | 544.4370 |
| FGC376 | | 4.552 | 0.031 | 69.2719 | 4.08 | 37.5130 |

| | |
|---|---|
| Max Area% | 95.915 |
| UV Signal Purity>95% | Pass |

**UV Apex Spectra**

**RT:**

3.773

**Sample Name:**

FGC376

**Signal Name:**

DAD1C

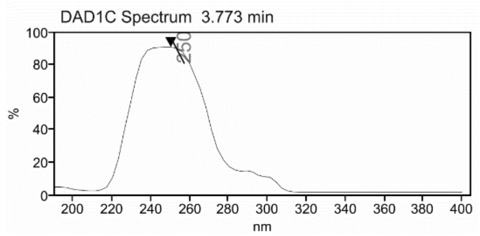

**RT:**

3.804

**Sample Name:**

FGC376

**Signal Name:**

MSD1TIC

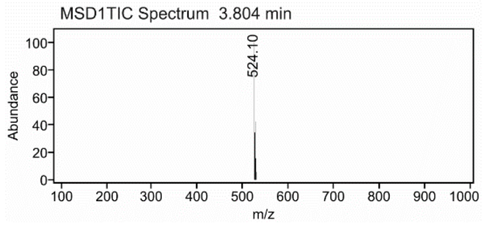

### Mass Spectrum SmartFormula Report

**Analysis Info**

| | | | |
|---|---|---|---|
| Analysis Name | D:\Data\MS_Service\Service_Kaiser\2024_Q1\20240228_SKN_FGC376_100x_5uL-m.d | Acquisition Date | 2/28/2024 12:57:41 PM |
| Method | DirectInfusion_WithFlow_Tune_pos.m | Operator | BDAL@DE |
| Sample Name | Test1 | Instrument | micrOTOF-Q 228888.10407 |
| Comment | | | |

**Acquisition Parameter**

| | | | | | |
|---|---|---|---|---|---|
| Source Type | ESI | Ion Polarity | Positive | Set Nebulizer | 1.8 Bar |
| Focus | Active | Set Capillary | 4000 V | Set Dry Heater | 320 °C |
| Scan Begin | 50 m/z | Set End Plate Offset | -500 V | Set Dry Gas | 8.0 l/min |
| Scan End | 1600 m/z | Set Collision Cell RF | 150.0 Vpp | Set Divert Valve | Waste |

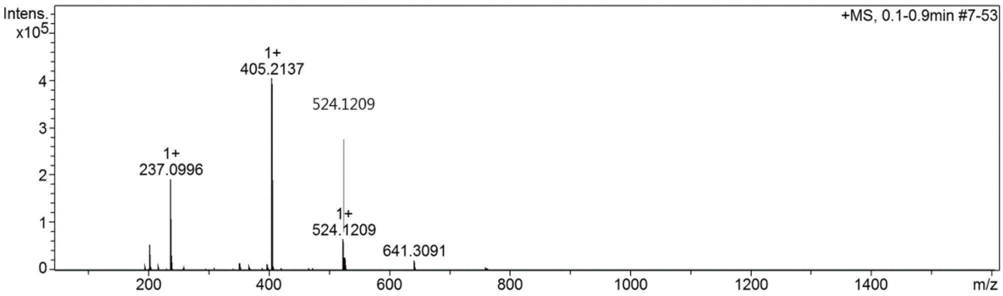

**Cmpd 44**

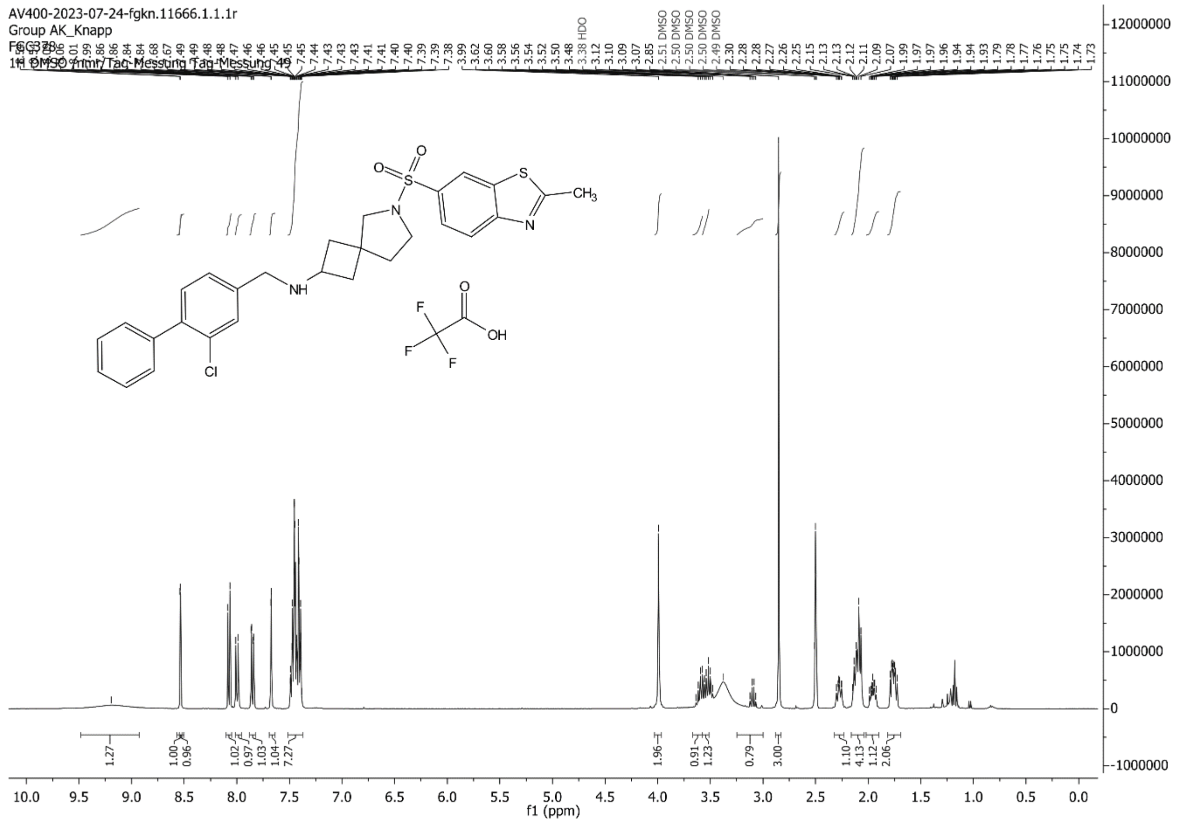
AV400-2023-07-24-fgkn.11666.1.1.1r
Group AK_Knapp
FGC378
1H DMSO /nmr/Tag-Messung Tag-Messung 49
f1 (ppm)

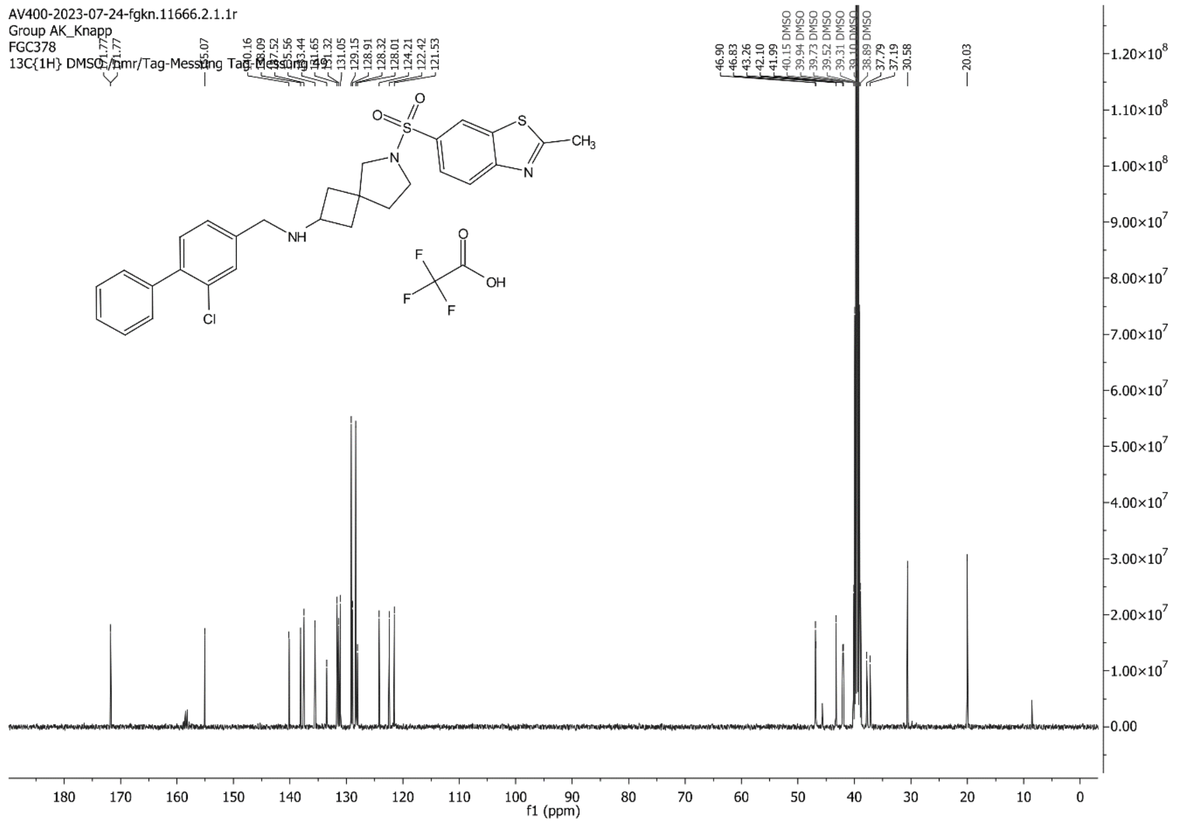
AV400-2023-07-24-fgkn.11666.2.1.1r
Group AK_Knapp
FGC378
13C{1H} DMSO /nmr/Tag-Messung Tag-Messung 49
f1 (ppm)

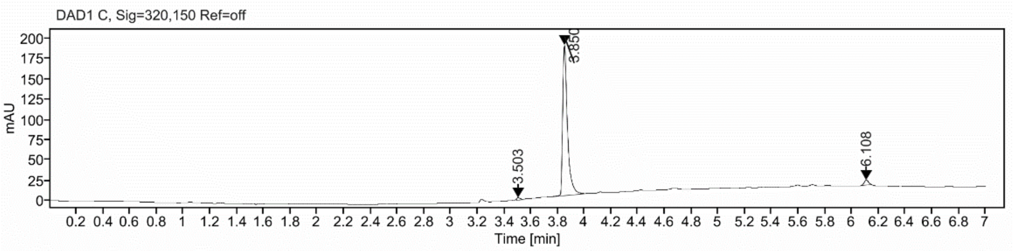

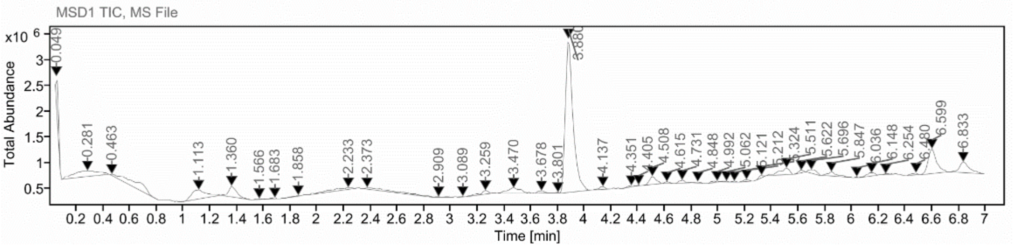

**Sample Purity**

Signal Description DAD1 C, Sig=320,150 Ref=off

| Sample Name | Name | RT | Width | Area | Area% | Height |
|---|---|---|---|---|---|---|
| FGC378 | | 3.503 | 0.037 | 5.7904 | 1.21 | 3.1417 |
| FGC378 | | 3.850 | 0.036 | 461.6666 | 96.08 | 185.2790 |
| FGC378 | | 6.108 | 0.037 | 13.0375 | 2.71 | 6.0347 |

Max Area% 96.082

UV Signal Purity>95% Pass

**UV Apex Spectra**

**RT:**

3.503

**Sample Name:**

FGC378

**Signal Name:**

DAD1C

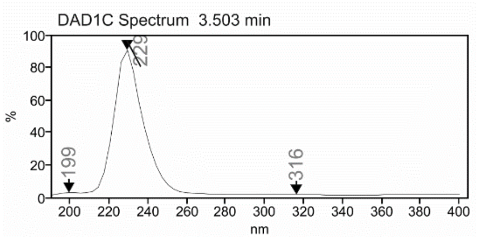

**RT:**

3.880

**Sample Name:**

FGC378

**Signal Name:**

MSD1TIC

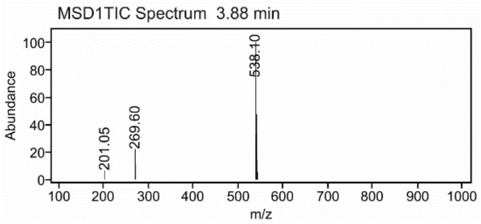

### Mass Spectrum SmartFormula Report

**Analysis Info**

| | | | |
|---|---|---|---|
| Analysis Name | D:\Data\MS_Service\Service_Kaiser\2024_Q1\20240228_SKN_FGC378_100x_5uL-m.d | Acquisition Date | 2/28/2024 1:12:12 PM |
| Method | DirectInfusion_WithFlow_Tune_pos.m | Operator | BDAL@DE |
| Sample Name | Test1 | Instrument | micrOTOF-Q 228888.10407 |
| Comment | | | |

**Acquisition Parameter**

| | | | | | |
|---|---|---|---|---|---|
| Source Type | ESI | Ion Polarity | Positive | Set Nebulizer | 1.8 Bar |
| Focus | Active | Set Capillary | 4000 V | Set Dry Heater | 320 °C |
| Scan Begin | 50 m/z | Set End Plate Offset | -500 V | Set Dry Gas | 8.0 l/min |
| Scan End | 1600 m/z | Set Collision Cell RF | 150.0 Vpp | Set Divert Valve | Waste |

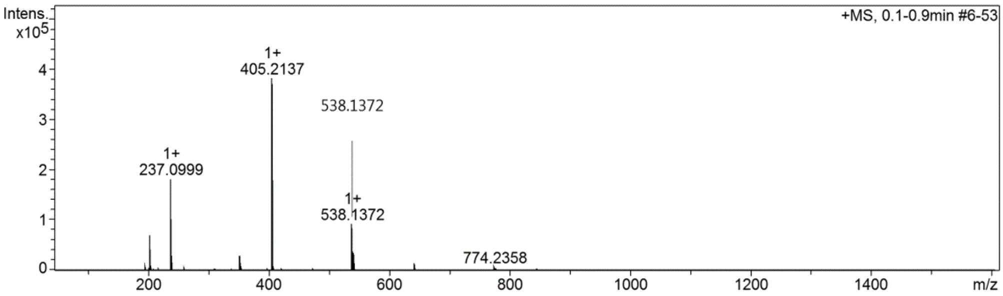

**Cmpd 45**

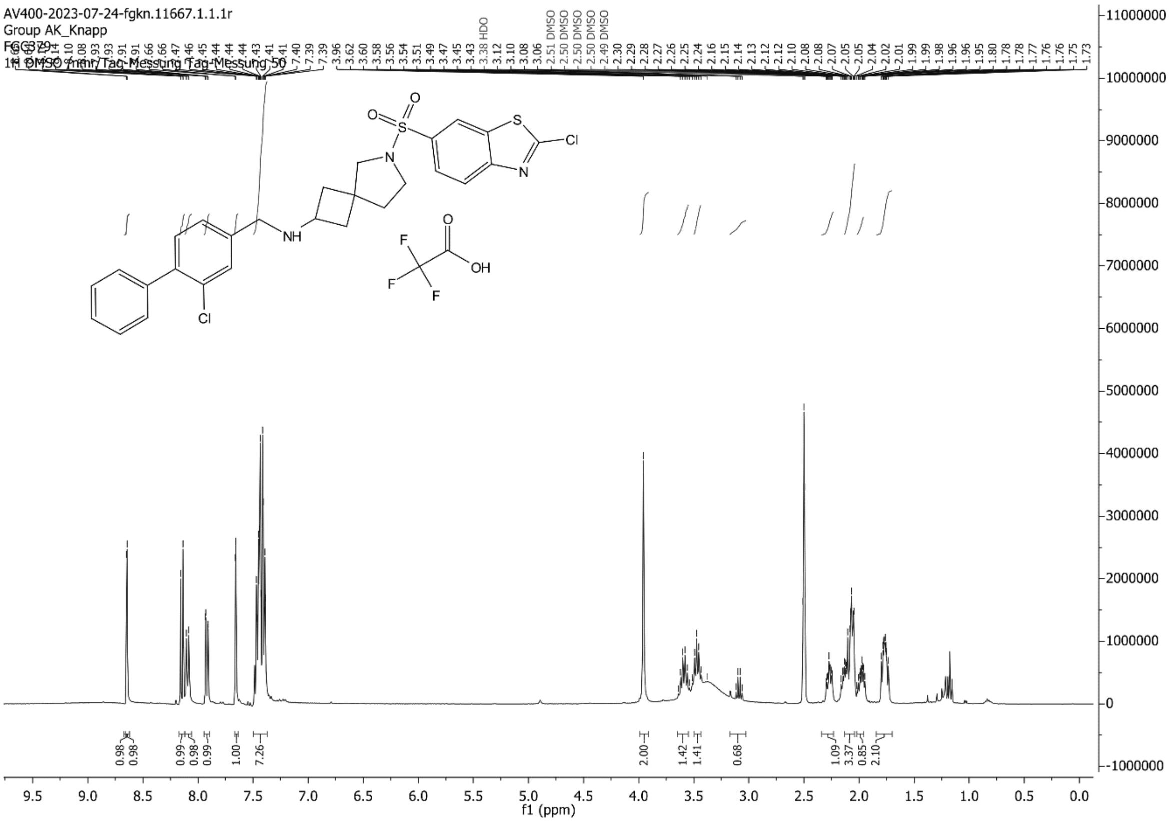

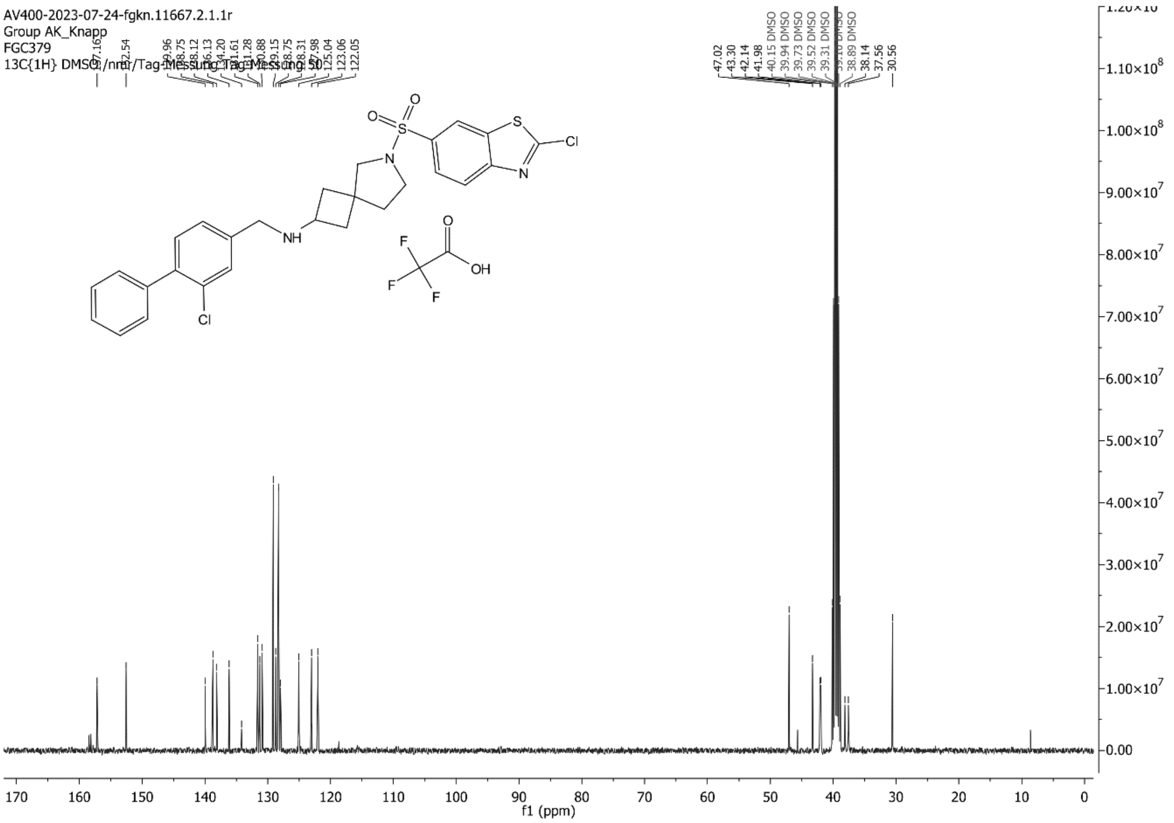

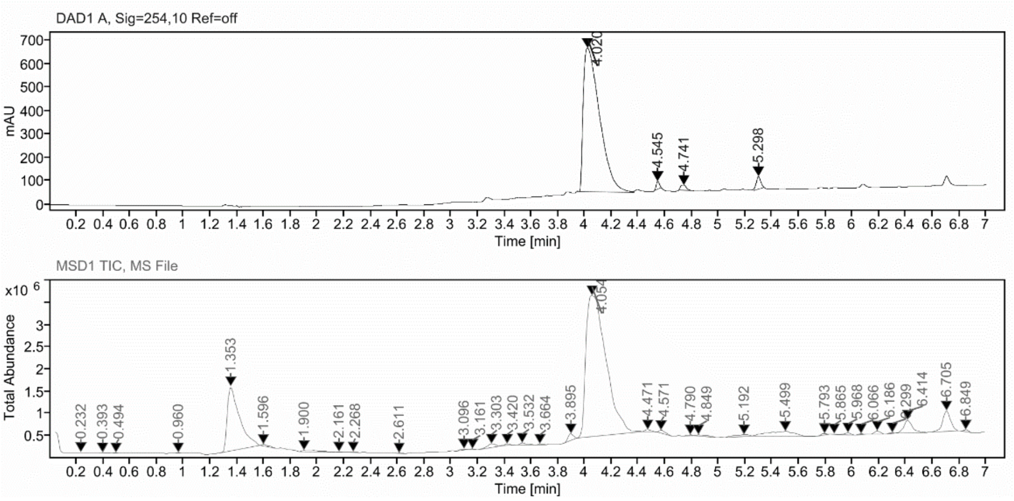

#### Sample Purity

Signal Description DAD1 A, Sig=254,10 Ref=off

| Sample Name | Name | RT | Width | Area | Area% | Height |
|---|---|---|---|---|---|---|
| FGC379_on | | 4.020 | 0.133 | 5186.1309 | 95.59 | 613.1758 |
| FGC379_on | | 4.545 | 0.028 | 53.5634 | 0.99 | 33.7198 |
| FGC379_on | | 4.741 | 0.048 | 62.4245 | 1.15 | 21.4947 |
| FGC379_on | | 5.298 | 0.038 | 123.5452 | 2.28 | 53.7307 |

Max Area% 95.585

UV Signal Purity>95% Pass

#### UV Apex Spectra

**RT:**

4.020

**Sample Name:**

FGC379_on

**Signal Name:**

DAD1A

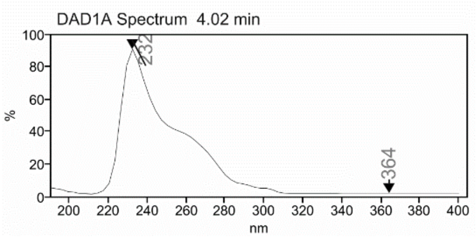

**RT:**

4.054

**Sample Name:**

FGC379_on

**Signal Name:**

MSD1TIC

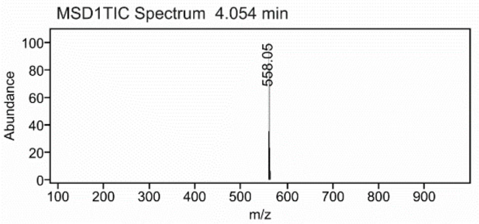

### Mass Spectrum SmartFormula Report

**Analysis Info**

| | | | |
|---|---|---|---|
| Analysis Name | D:\Data\MS_Service\Service_Kaiser\2024_Q1\20240227_SKN_FGC374_100x_5uL-m.d | Acquisition Date | 2/27/2024 3:36:42 PM |
| Method | DirectInfusion_WithFlow_Tune_pos.m | Operator | BDAL@DE |
| Sample Name | Test1 | Instrument | micrOTOF-Q 228888.10407 |
| Comment | | | |

**Acquisition Parameter**

| | | | | | |
|---|---|---|---|---|---|
| Source Type | ESI | Ion Polarity | Positive | Set Nebulizer | 1.8 Bar |
| Focus | Active | Set Capillary | 4000 V | Set Dry Heater | 320 °C |
| Scan Begin | 50 m/z | Set End Plate Offset | -500 V | Set Dry Gas | 8.0 l/min |
| Scan End | 1600 m/z | Set Collision Cell RF | 150.0 Vpp | Set Divert Valve | Waste |

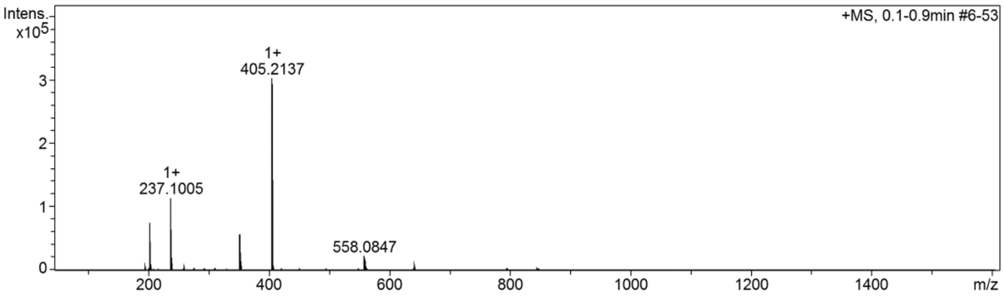

#### Cmpd 46

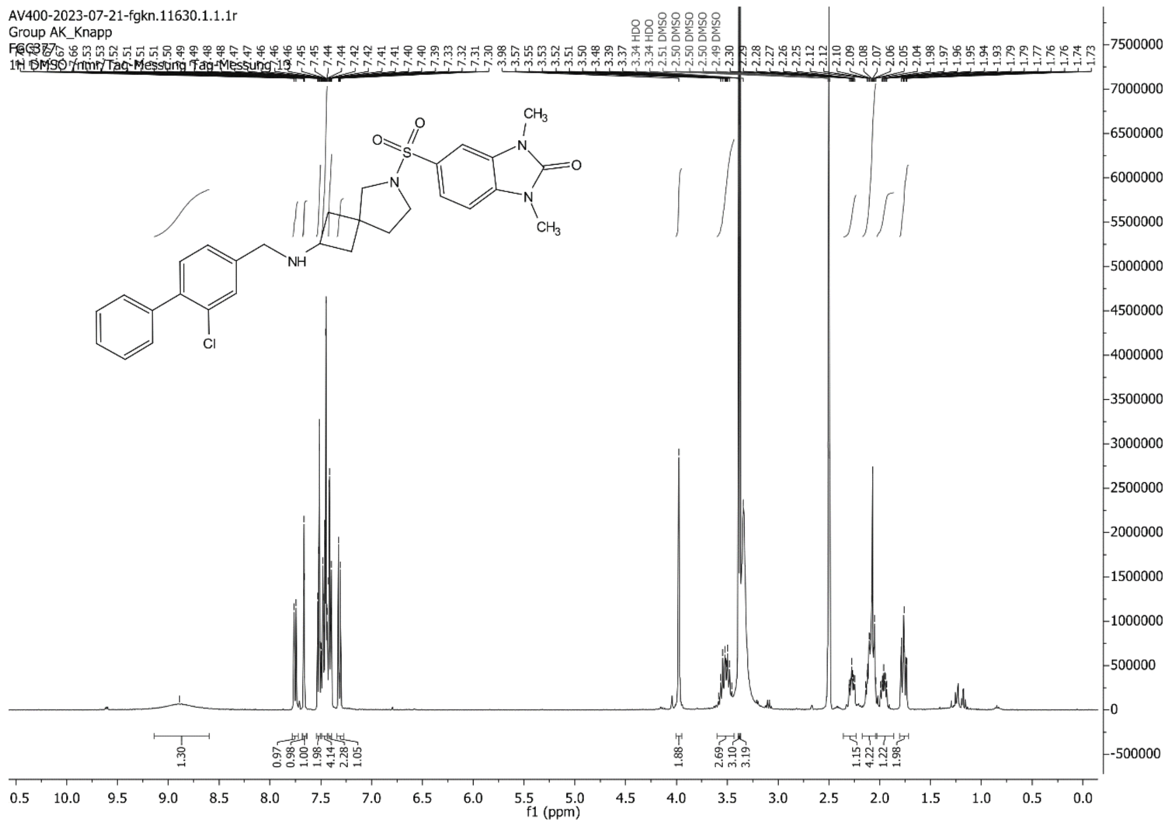

AV400-2023-07-21-fgkn.11630.1.1.1r
Group AK_Knapp
FGC377
1H DMSO /nmr/Tag-Messung Tag-Messung 13
f1 (ppm)

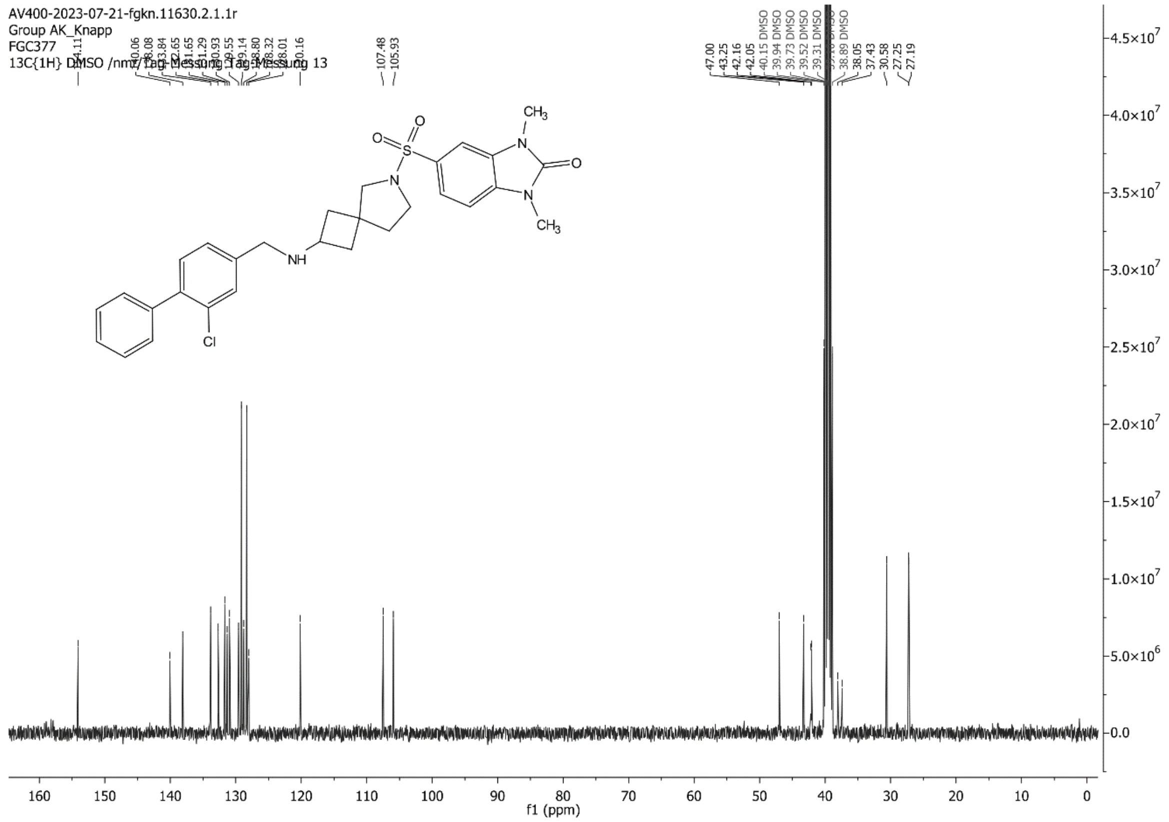

AV400-2023-07-21-fgkn.11630.2.1.1r
Group AK_Knapp
FGC377
13C{1H} DMSO /nmr/Tag-Messung Tag-Messung 13
f1 (ppm)

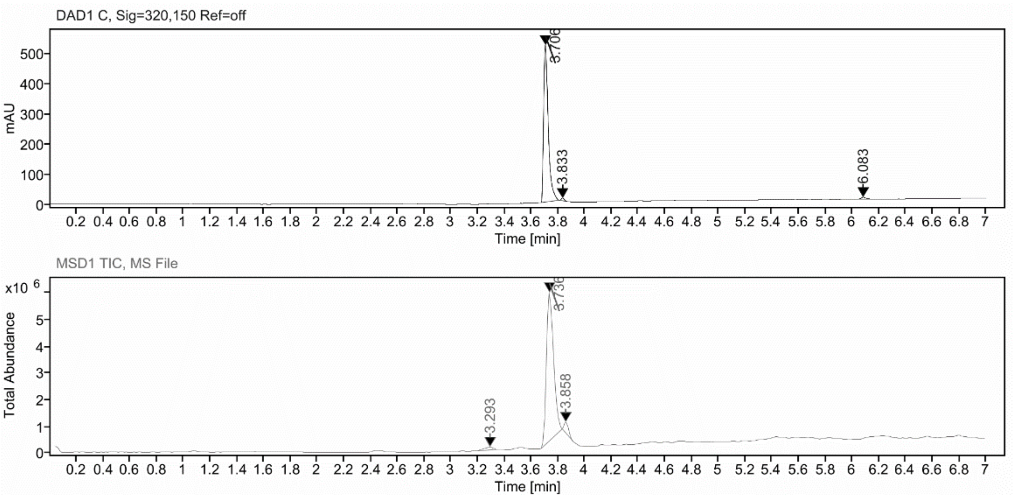

#### Sample Purity

Signal Description DAD1 C, Sig=320,150 Ref=off

| Sample Name | Name | RT | Width | Area | Area% | Height |
|---|---|---|---|---|---|---|
| FGC377 | | 3.706 | 0.036 | 1252.8505 | 97.58 | 519.3900 |
| FGC377 | | 3.833 | 0.025 | 14.3530 | 1.12 | 9.0112 |
| FGC377 | | 6.083 | 0.040 | 16.6810 | 1.30 | 6.1005 |

Max Area% 97.583

UV Signal Purity>95% Pass

#### UV Apex Spectra

**RT:**

3.706

**Sample Name:**

FGC377

**Signal Name:**

DAD1C

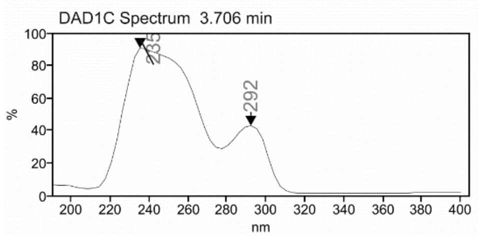

**RT:**

3.736

**Sample Name:**

FGC377

**Signal Name:**

MSD1TIC

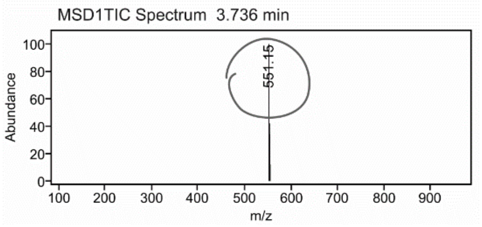

### Mass Spectrum SmartFormula Report

**Analysis Info**

| | | | |
|---|---|---|---|
| | | Acquisition Date | 2/28/2024 1:04:57 PM |
| Analysis Name | D:\Data\MS_Service\Service_Kaiser\2024_Q1\20240228_SKN_FGC377_100x_5uL-m.d | | |
| Method | DirectInfusion_WithFlow_Tune_pos.m | Operator | BDAL@DE |
| Sample Name | Test1 | Instrument | micrOTOF-Q 228888.10407 |
| Comment | | | |

**Acquisition Parameter**

| | | | | | |
|---|---|---|---|---|---|
| Source Type | ESI | Ion Polarity | Positive | Set Nebulizer | 1.8 Bar |
| Focus | Active | Set Capillary | 4000 V | Set Dry Heater | 320 °C |
| Scan Begin | 50 m/z | Set End Plate Offset | -500 V | Set Dry Gas | 8.0 l/min |
| Scan End | 1600 m/z | Set Collision Cell RF | 150.0 Vpp | Set Divert Valve | Waste |

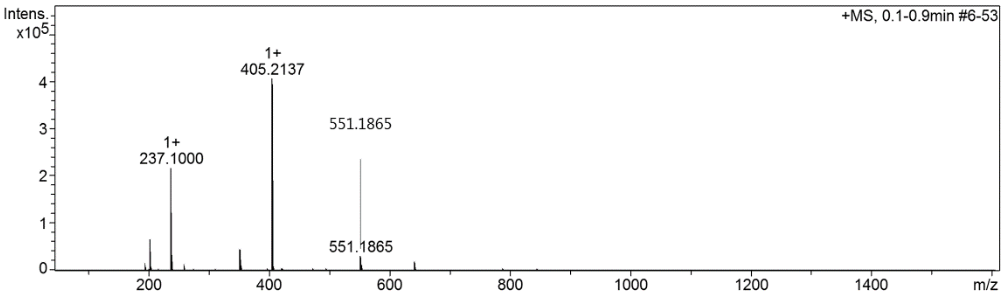

#### Cmpd 47

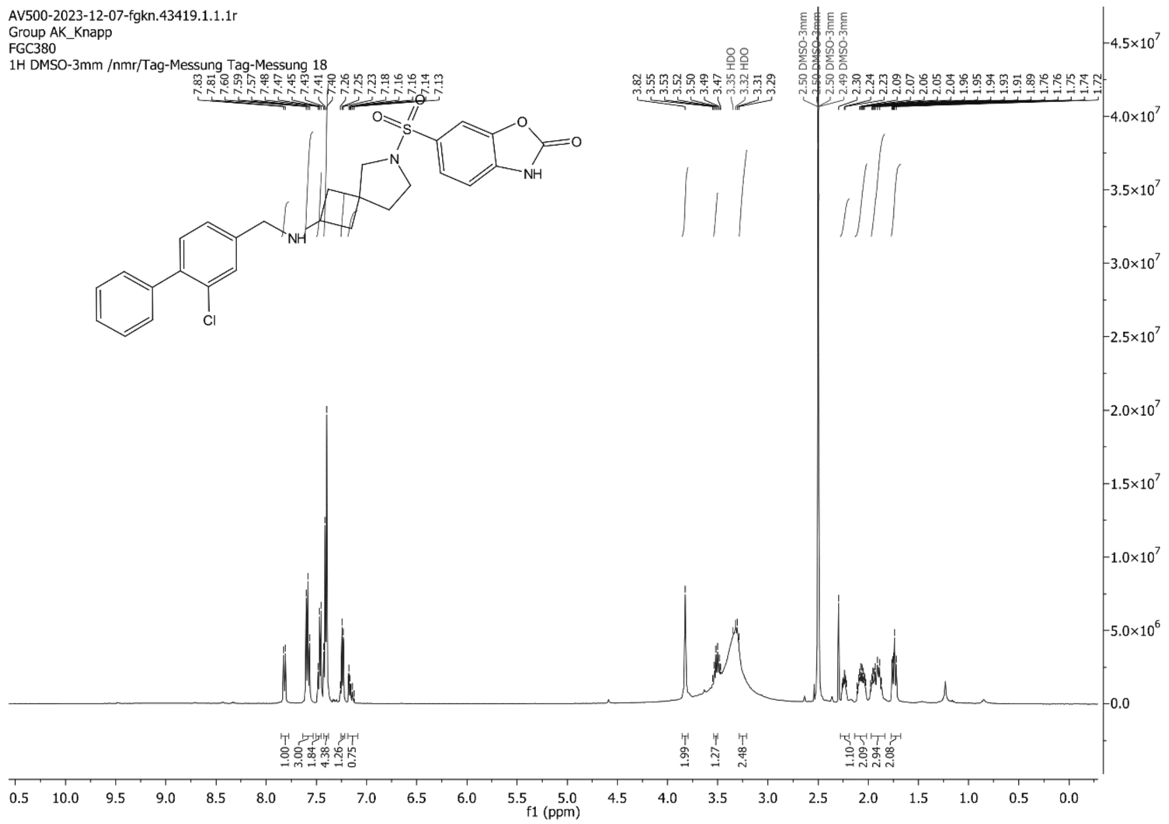
AV500-2023-12-07-fgkn.43419.1.1.1r
Group AK_Knapp
FGC380
1H DMSO-3mm /nmr/Tag-Messung Tag-Messung 18
f1 (ppm)

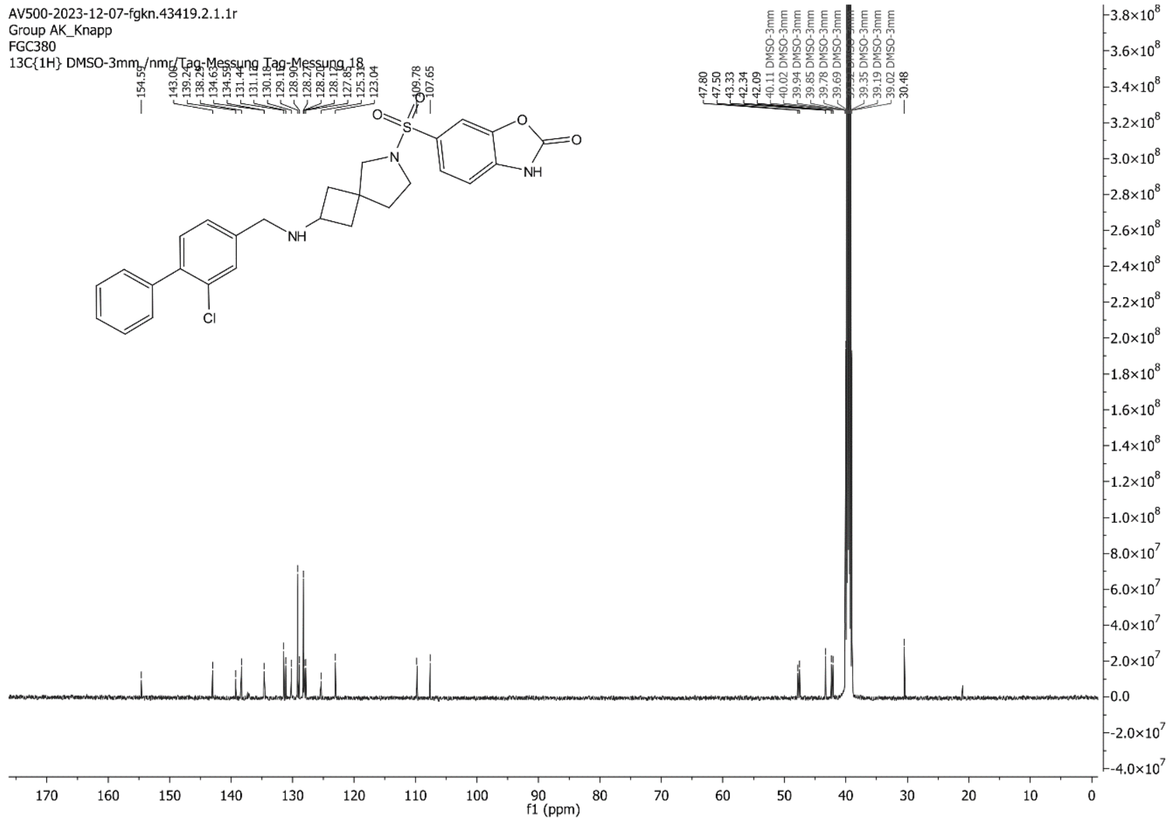
AV500-2023-12-07-fgkn.43419.2.1.1r
Group AK_Knapp
FGC380
13C{1H} DMSO-3mm /nmr/Tag-Messung Tag-Messung 18
f1 (ppm)

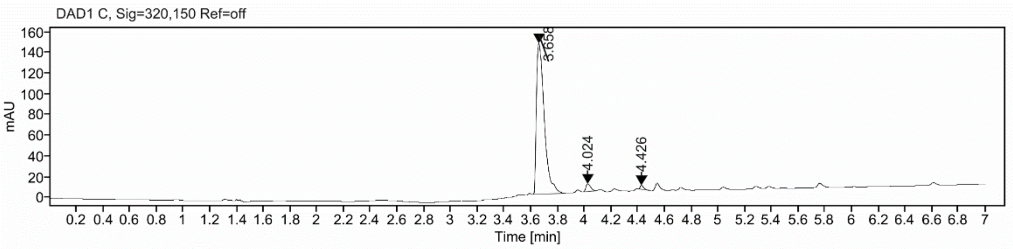

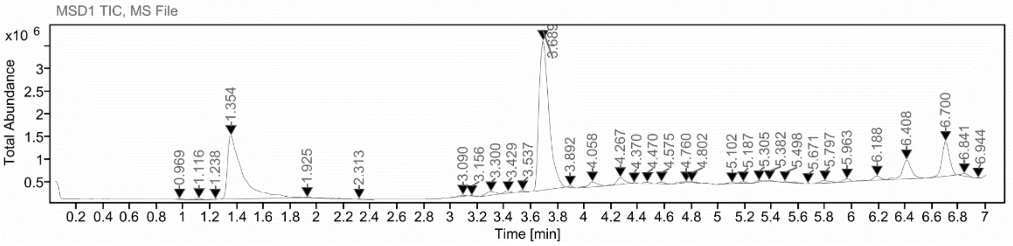

**Sample Purity**

Signal Description DAD1 C, Sig=320,150 Ref=off

| Sample Name | Name | RT | Width | Area | Area% | Height |
|---|---|---|---|---|---|---|
| FGC380_on | | 3.658 | 0.063 | 595.2827 | 96.05 | 145.5039 |
| FGC380_on | | 4.024 | 0.040 | 17.4869 | 2.82 | 7.1265 |
| FGC380_on | | 4.426 | 0.026 | 6.9897 | 1.13 | 3.8737 |

Max Area% 96.051

UV Signal Purity>95% Pass

**UV Apex Spectra**

**RT:**

3.658

**Sample Name:**

FGC380_on

**Signal Name:**

DAD1C

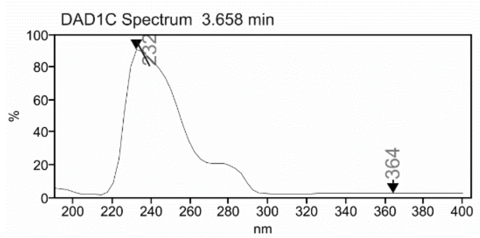

**RT:**

3.689

**Sample Name:**

FGC380_on

**Signal Name:**

MSD1TIC

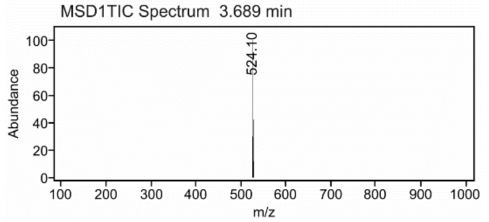

### Mass Spectrum SmartFormula Report

**Analysis Info**

| | | | |
|---|---|---|---|
| Analysis Name | D:\Data\MS_Service\Service_Kaiser\2024_Q1\20240228_SKN_FGC380_100x_5uL-m.d | Acquisition Date | 2/28/2024 1:22:22 PM |
| Method | DirectInfusion_WithFlow_Tune_pos.m | Operator | BDAL@DE |
| Sample Name | Test1 | Instrument | micrOTOF-Q 228888.10407 |
| Comment | | | |

**Acquisition Parameter**

| | | | | | |
|---|---|---|---|---|---|
| Source Type | ESI | Ion Polarity | Positive | Set Nebulizer | 1.8 Bar |
| Focus | Active | Set Capillary | 4000 V | Set Dry Heater | 320 °C |
| Scan Begin | 50 m/z | Set End Plate Offset | -500 V | Set Dry Gas | 8.0 l/min |
| Scan End | 1600 m/z | Set Collision Cell RF | 150.0 Vpp | Set Divert Valve | Waste |

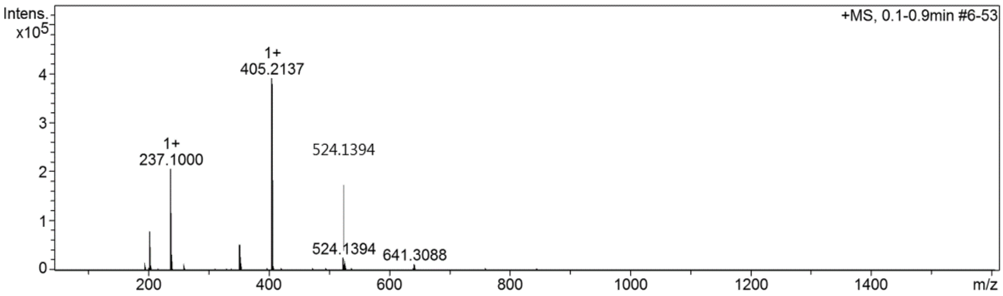

#### Cmpd 48

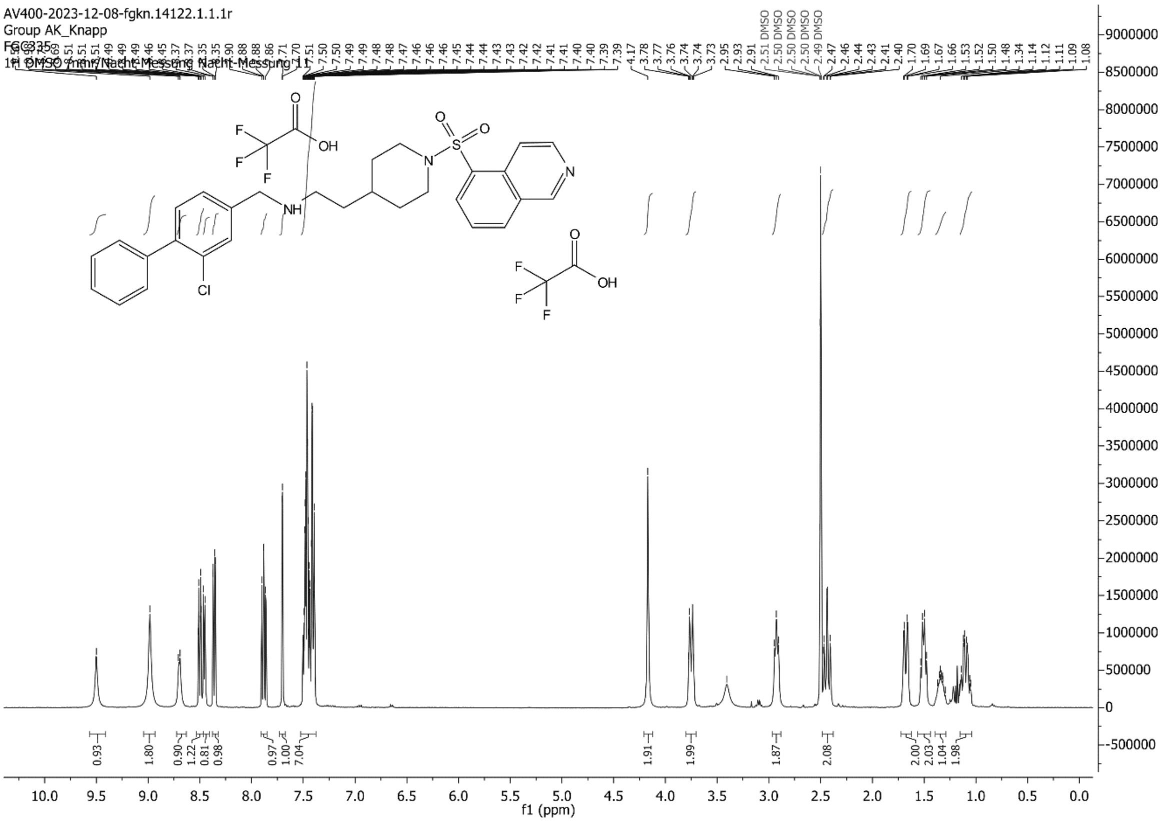

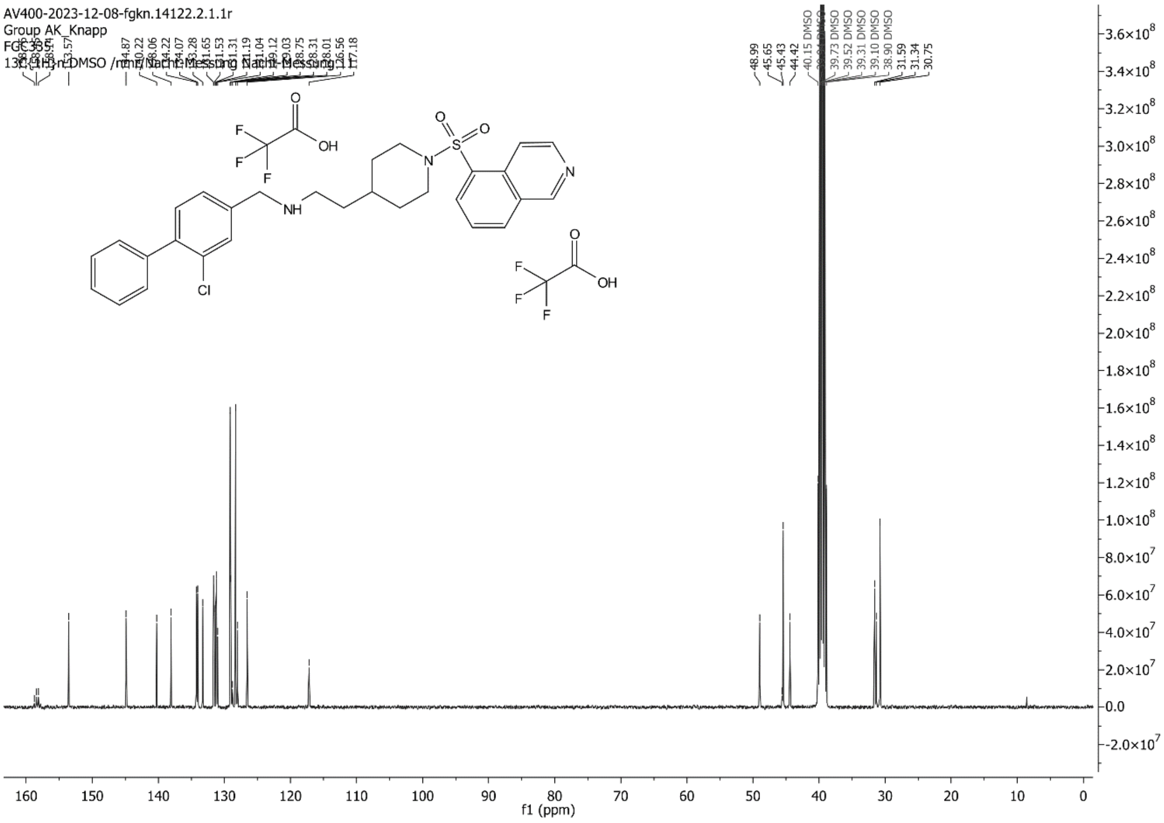

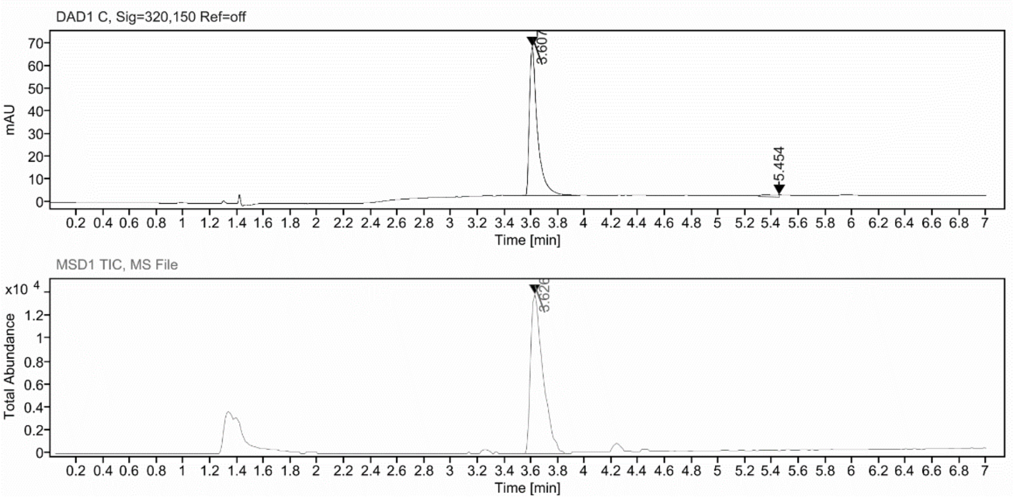

**Sample Purity**

| Signal Description | DAD1 C, Sig=320,150 Ref=off |
|---|---|

| **Sample Name** | **Name** | **RT** | **Width** | **Area** | **Area%** | **Height** |
|---|---|---|---|---|---|---|
| FGC335_purity | | 3.607 | 0.058 | 277.8574 | 97.39 | 66.0211 |
| FGC335_purity | | 5.454 | 0.157 | 7.4512 | 2.61 | 0.9509 |

| | |
|---|---|
| Max Area% | 97.388 |
| UV Signal Purity>95% | Pass |

**UV Apex Spectra**

**RT:**

3.607

**Sample Name:**

FGC335_purity

**Signal Name:**

DAD1C

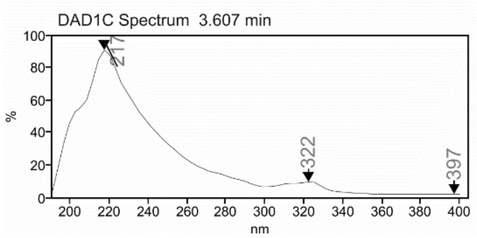

**RT:**

3.626

**Sample Name:**

FGC335_purity

**Signal Name:**

MSD1TIC

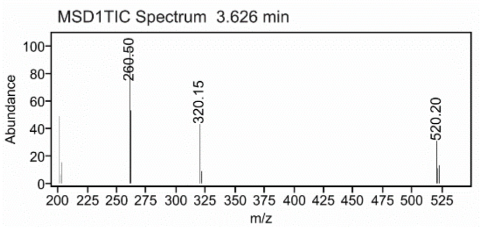

### Mass Spectrum SmartFormula Report

**Analysis Info**

| | | | |
|---|---|---|---|
| Analysis Name | D:\Data\MS_Service\Service_Kaiser\2024_Q1\20240219_SKN_FGC335_5ug-mL_5uL-m.d | Acquisition Date | 2/19/2024 2:03:00 PM |
| Method | DirectInfusion_WithFlow_Tune_pos.m | Operator | BDAL@DE |
| Sample Name | Test1 | Instrument | micrOTOF-Q 228888.10407 |
| Comment | | | |

**Acquisition Parameter**

| | | | | | |
|---|---|---|---|---|---|
| Source Type | ESI | Ion Polarity | Positive | Set Nebulizer | 1.8 Bar |
| Focus | Active | Set Capillary | 4000 V | Set Dry Heater | 220 °C |
| Scan Begin | 50 m/z | Set End Plate Offset | -500 V | Set Dry Gas | 8.0 l/min |
| Scan End | 1600 m/z | Set Collision Cell RF | 150.0 Vpp | Set Divert Valve | Waste |

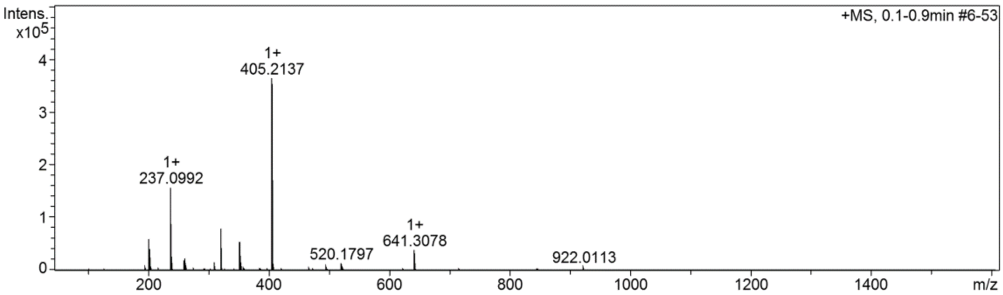

#### Cmpd 49

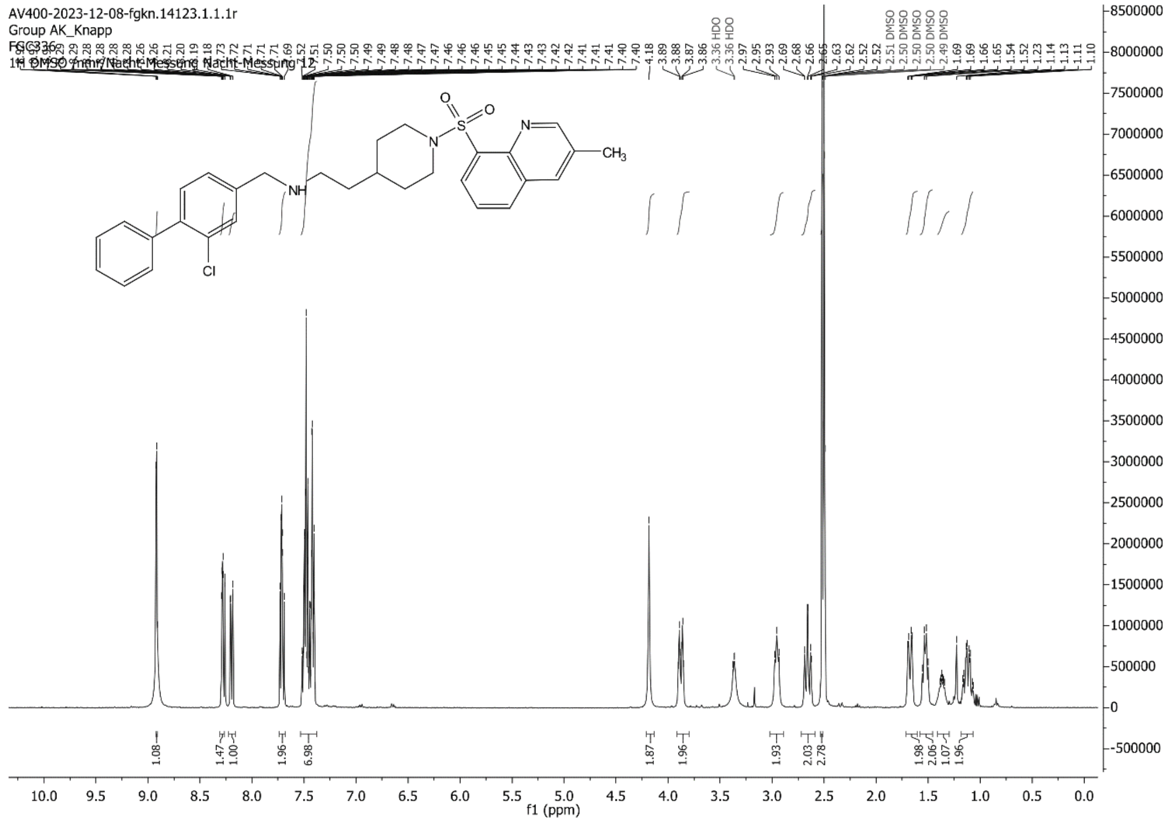
AV400-2023-12-08-fgkn.14123.1.1.1r
Group AK_Knapp
FGC3336
1H DMSO /nmr/Nacht-Messung Nacht-Messung 12
f1 (ppm)

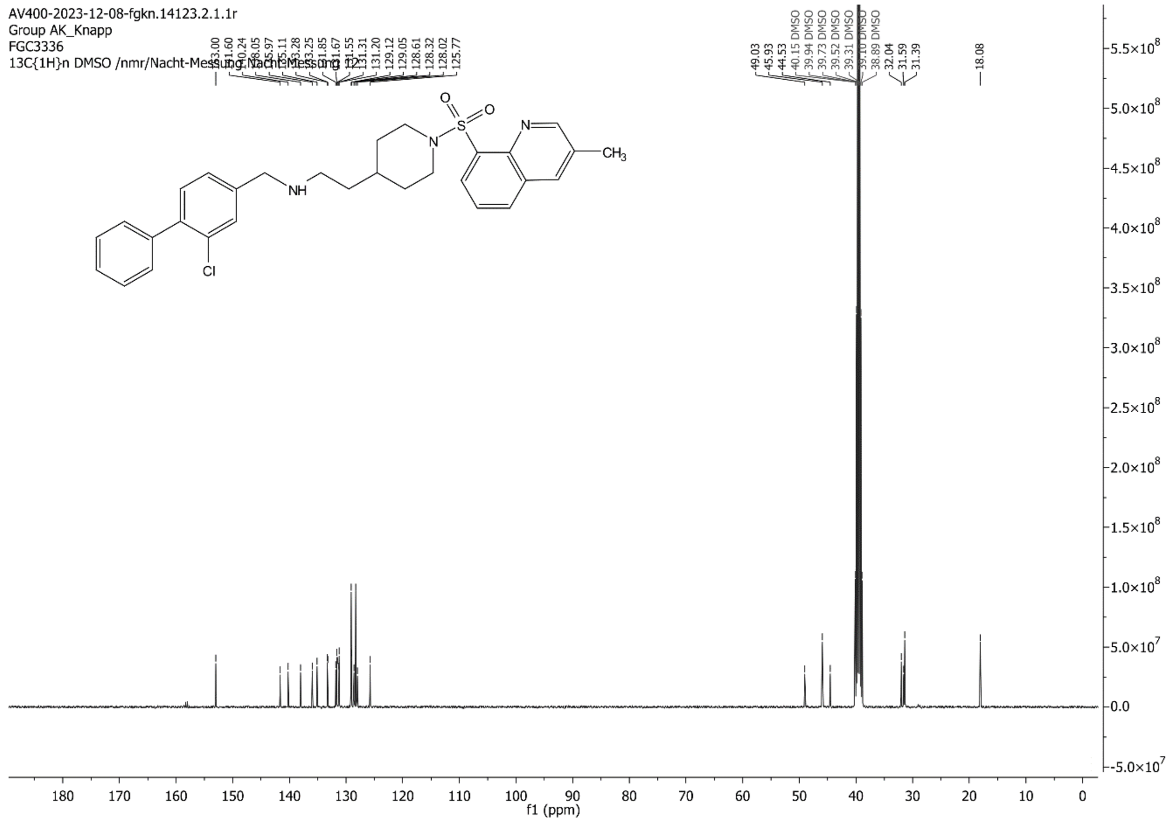
AV400-2023-12-08-fgkn.14123.2.1.1r
Group AK_Knapp
FGC3336
13C{1H}n DMSO /nmr/Nacht-Messung Nacht-Messung 12
f1 (ppm)

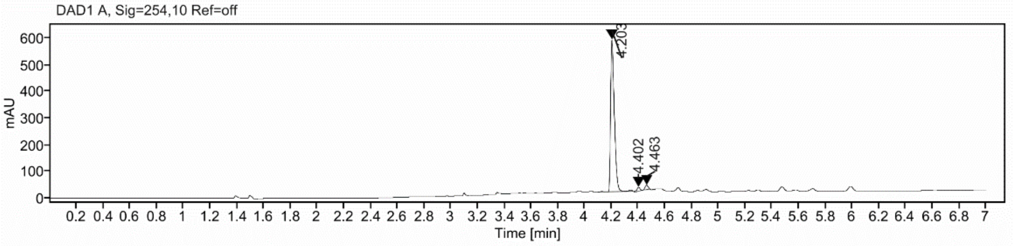

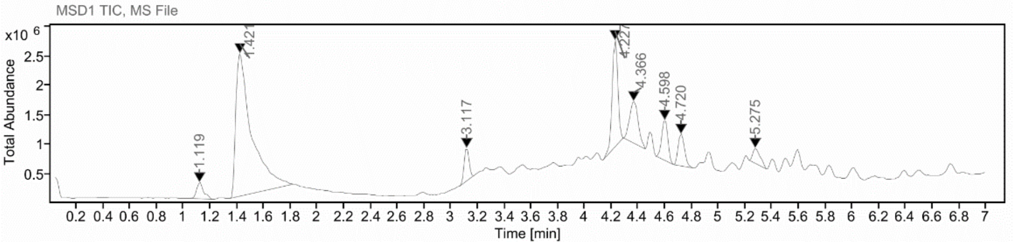

**Sample Purity**

Signal Description DAD1 A, Sig=254,10 Ref=off

| Sample Name | Name | RT | Width | Area | Area% | Height |
|---|---|---|---|---|---|---|
| FGC336_X | | 4.203 | 0.030 | 1124.2186 | 95.87 | 571.2169 |
| FGC336_X | | 4.402 | 0.025 | 24.7607 | 2.11 | 14.8312 |
| FGC336_X | | 4.463 | 0.026 | 23.7141 | 2.02 | 16.4566 |

Max Area% 95.866

UV Signal Purity>95% Pass

**UV Apex Spectra**

**RT:**

4.203

**Sample Name:**

FGC336_X

**Signal Name:**

DAD1A

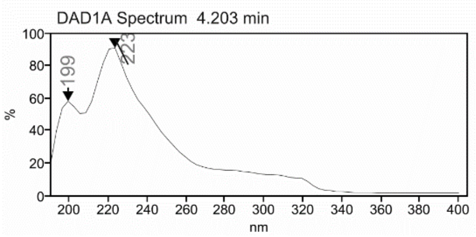

**RT:**

4.227

**Sample Name:**

FGC336_X

**Signal Name:**

MSD1TIC

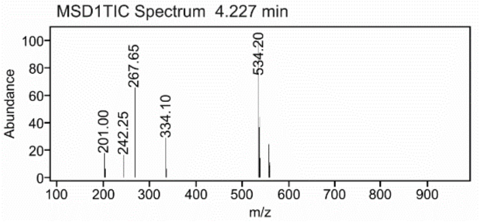

### Mass Spectrum SmartFormula Report

**Analysis Info**

| | | | |
|---|---|---|---|
| Analysis Name | D:\Data\MS_Service\Service_Kaiser\2024_Q1\20240219_SKN_FGC336_5ug-mL_5uL-m.d | Acquisition Date | 2/19/2024 2:10:47 PM |
| Method | DirectInfusion_WithFlow_Tune_pos.m | Operator | BDAL@DE |
| Sample Name | Test1 | Instrument | micrOTOF-Q 228888.10407 |
| Comment | | | |

**Acquisition Parameter**

| | | | | | |
|---|---|---|---|---|---|
| Source Type | ESI | Ion Polarity | Positive | Set Nebulizer | 1.8 Bar |
| Focus | Active | Set Capillary | 4000 V | Set Dry Heater | 220 °C |
| Scan Begin | 50 m/z | Set End Plate Offset | -500 V | Set Dry Gas | 8.0 l/min |
| Scan End | 1600 m/z | Set Collision Cell RF | 150.0 Vpp | Set Divert Valve | Waste |

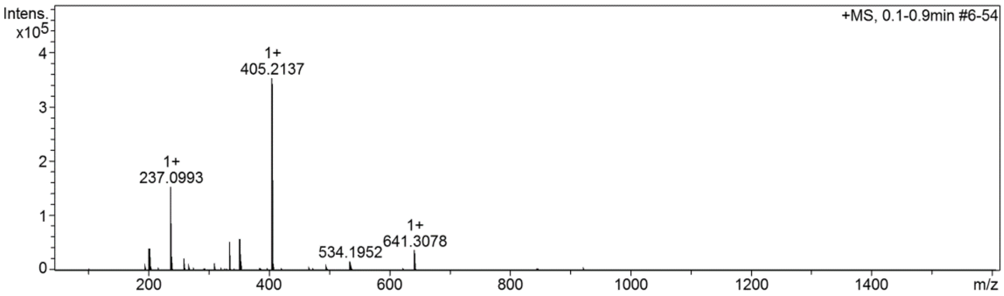

#### Cmpd 50

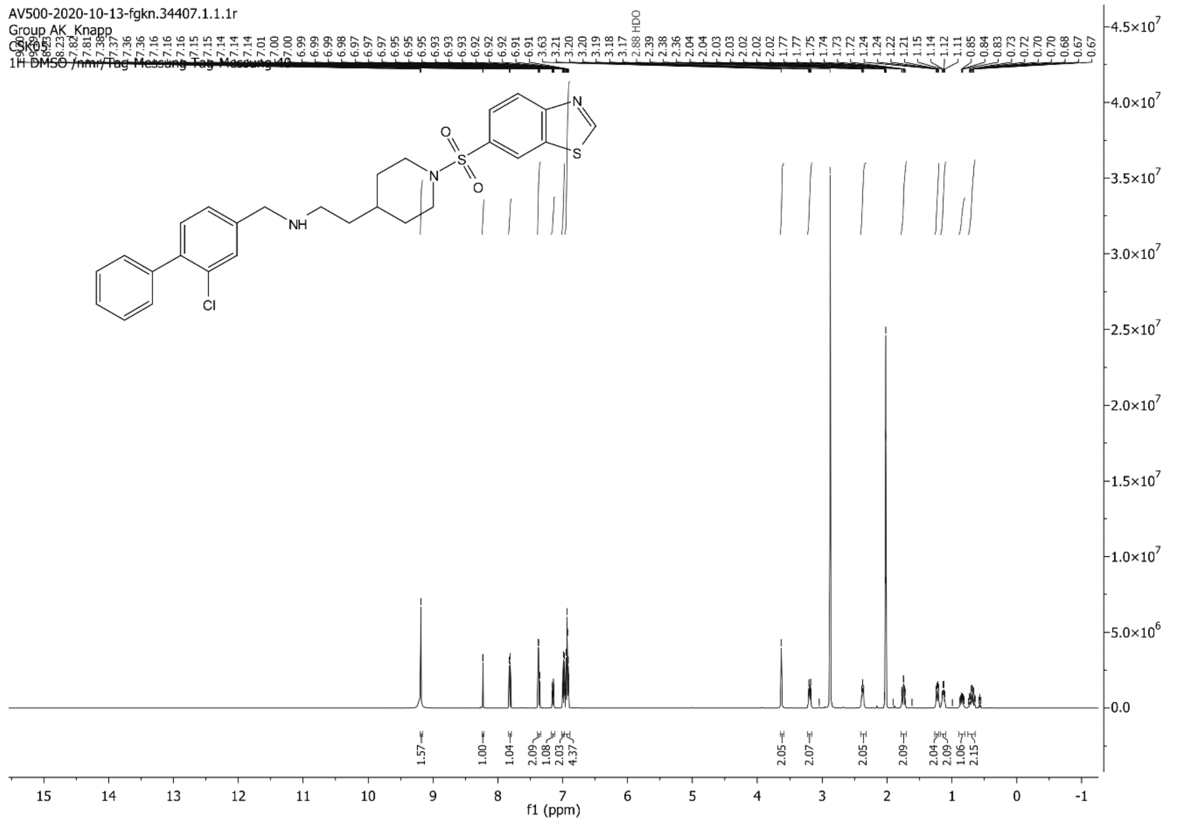
AV500-2020-10-13-fgkn.34407.1.1.1r
Group AK_Knapp
CSK05
HDO
f1 (ppm)

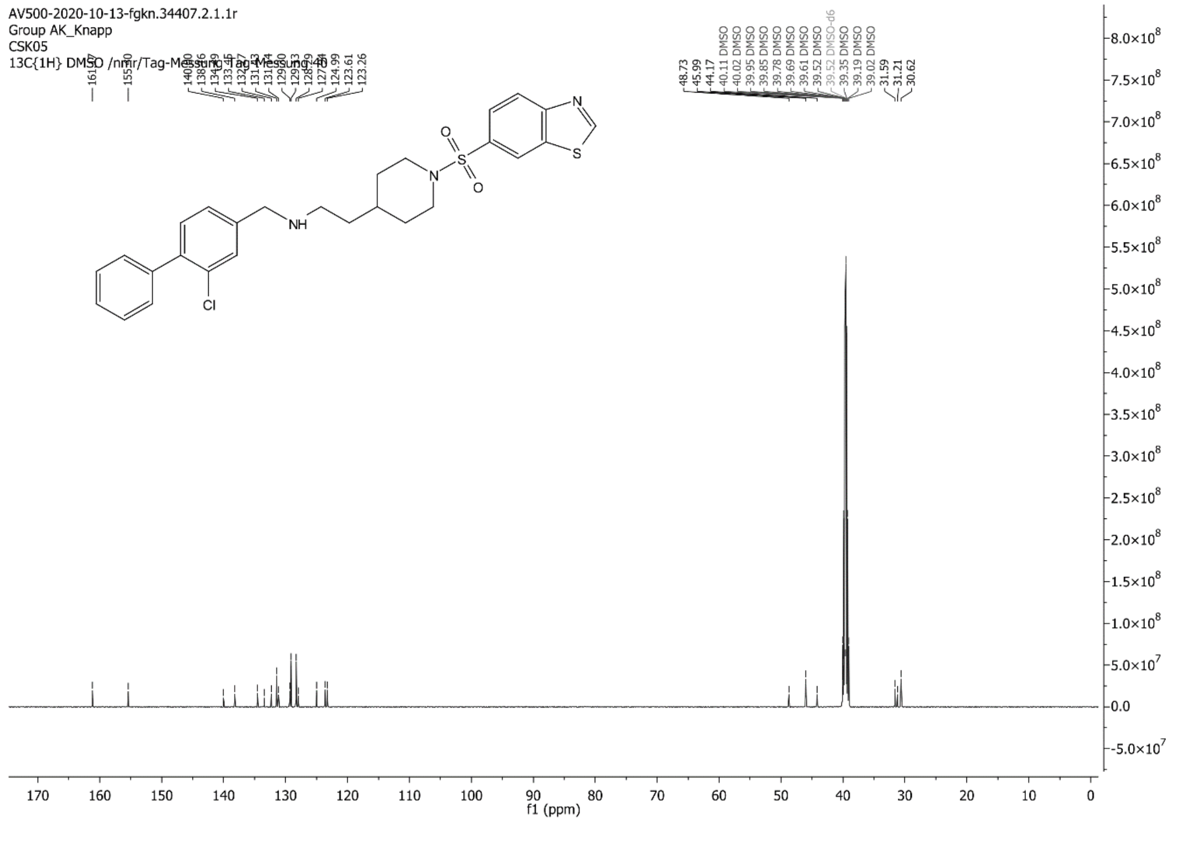
AV500-2020-10-13-fgkn.34407.2.1.1r
Group AK_Knapp
CSK05
13C{1H} DMSO /nmr/Tag-Messung Tag-Messung 40
f1 (ppm)

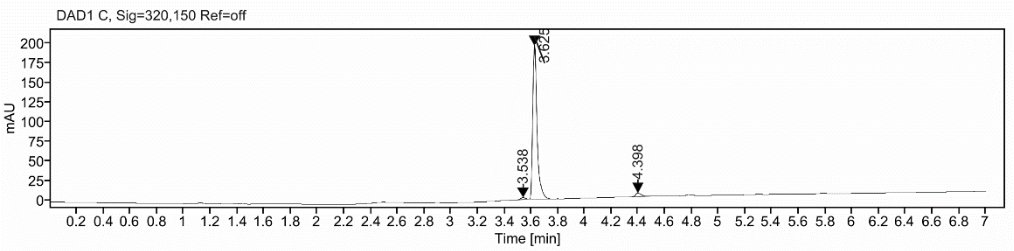

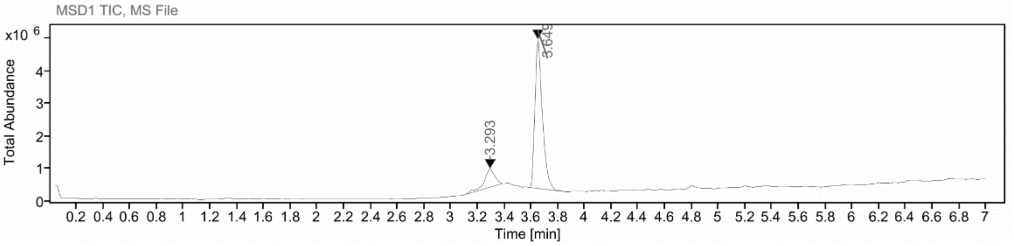

**Sample Purity**

Signal Description DAD1 C, Sig=320,150 Ref=off

| Sample Name | Name | RT | Width | Area | Area% | Height |
|---|---|---|---|---|---|---|
| FGCSK5 | | 3.538 | 0.039 | 5.2885 | 1.17 | 2.2926 |
| FGCSK5 | | 3.625 | 0.032 | 434.1948 | 95.73 | 194.9498 |
| FGCSK5 | | 4.398 | 0.046 | 14.0973 | 3.11 | 4.6828 |

Max Area% 95.726

UV Signal Purity>95% Pass

**UV Apex Spectra**

**RT:**

3.625

**Sample Name:**

FGCSK5

**Signal Name:**

DAD1C

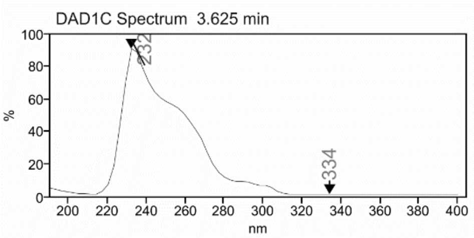

**RT:**

3.649

**Sample Name:**

FGCSK5

**Signal Name:**

MSD1TIC

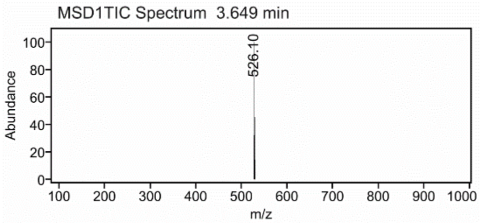

### Mass Spectrum SmartFormula Report

**Analysis Info**

Acquisition Date 2/19/2024 3:02:31 PM

| | | | |
|---|---|---|---|
| Analysis Name | D:\Data\MS_Service\Service_Kaiser\2024_Q1\20240219_SKN_FGCSK5_5ug-mL_5uL-m.d | | |
| Method | DirectInfusion_WithFlow_Tune_pos.m | Operator | BDAL@DE |
| Sample Name | Test1 | Instrument | micrOTOF-Q 228888.10407 |
| Comment | | | |

**Acquisition Parameter**

| | | | | | |
|---|---|---|---|---|---|
| Source Type | ESI | Ion Polarity | Positive | Set Nebulizer | 1.8 Bar |
| Focus | Active | Set Capillary | 4000 V | Set Dry Heater | 220 °C |
| Scan Begin | 50 m/z | Set End Plate Offset | -500 V | Set Dry Gas | 8.0 l/min |
| Scan End | 1600 m/z | Set Collision Cell RF | 150.0 Vpp | Set Divert Valve | Waste |

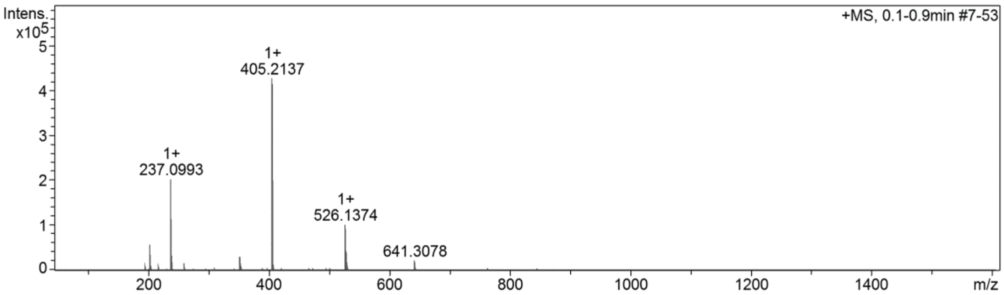

**Cmpd 51**

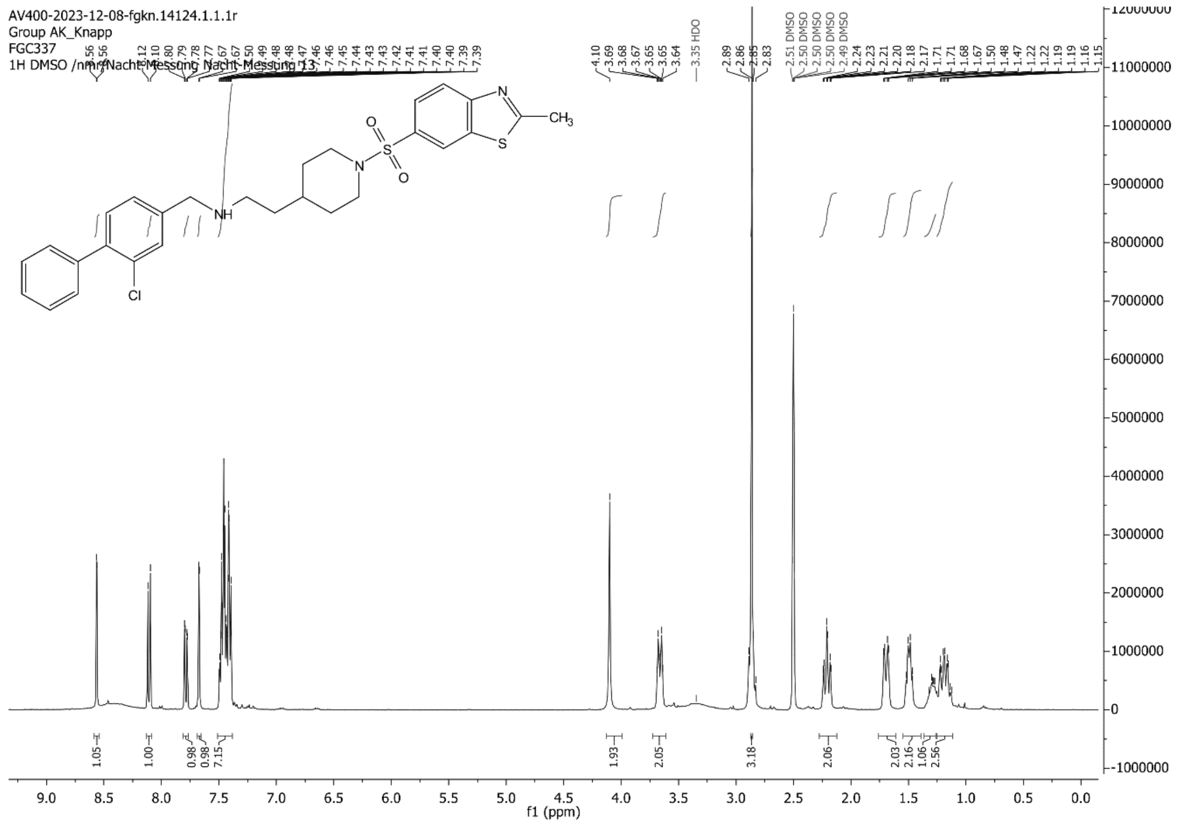
AV400-2023-12-08-fgkn.14124.1.1.1r
Group AK_Knapp
FGC337
1H DMSO /nmr/Nacht-Messung Nacht-Messung 13
f1 (ppm)
CH3
Cl
NH

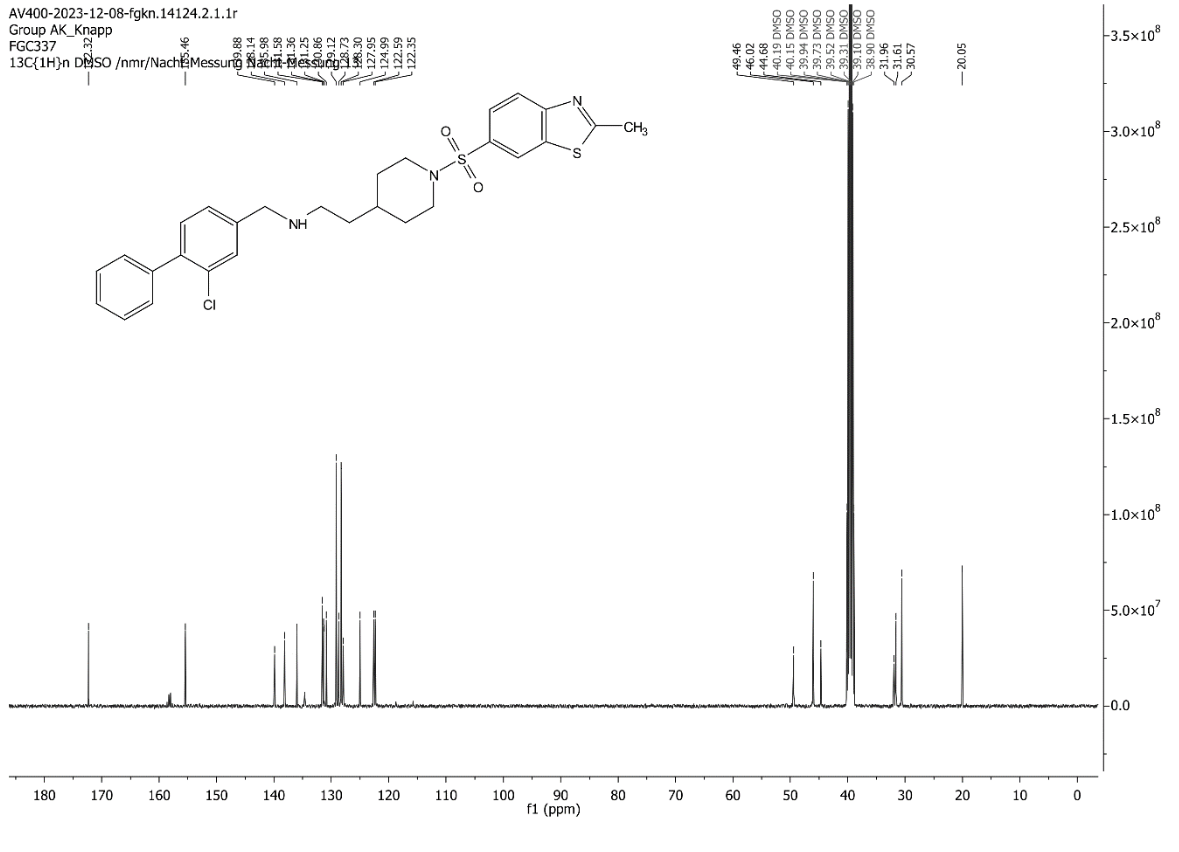
AV400-2023-12-08-fgkn.14124.2.1.1r
Group AK_Knapp
FGC337
13C{1H}n DMSO /nmr/Nacht-Messung Nacht-Messung 13
f1 (ppm)
CH3
Cl
NH

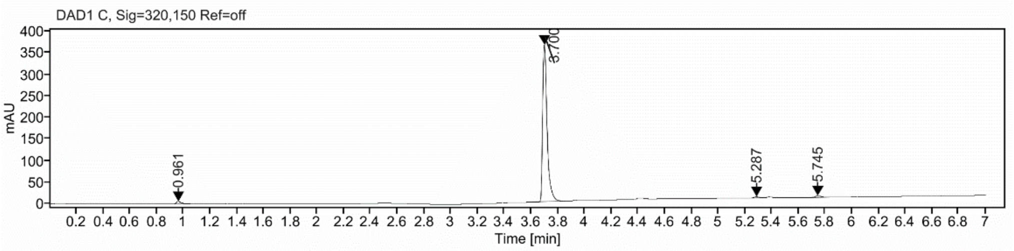

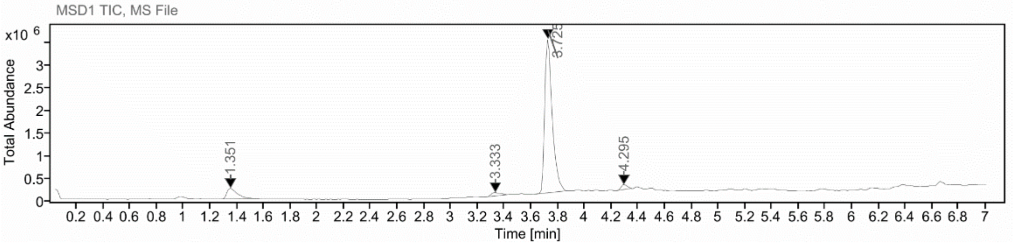

**Sample Purity**

Signal Description DAD1 C, Sig=320,150 Ref=off

| Sample Name | Name | RT | Width | Area | Area% | Height |
|---|---|---|---|---|---|---|
| FGC337 | | 0.961 | 0.037 | 18.0268 | 2.01 | 5.6408 |
| FGC337 | | 3.700 | 0.035 | 859.2681 | 96.03 | 364.4642 |
| FGC337 | | 5.287 | 0.040 | 6.3560 | 0.71 | 2.5472 |
| FGC337 | | 5.745 | 0.045 | 11.1449 | 1.25 | 3.8231 |

Max Area% 96.030

UV Signal Purity>95% Pass

**UV Apex Spectra**

**RT:**

3.700

**Sample Name:**

FGC337

**Signal Name:**

DAD1C

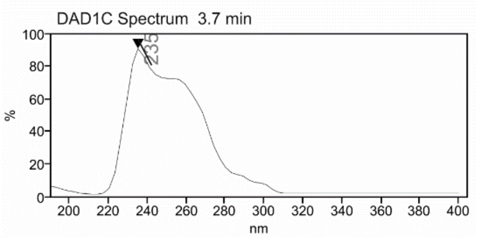

**RT:**

3.725

**Sample Name:**

FGC337

**Signal Name:**

MSD1TIC

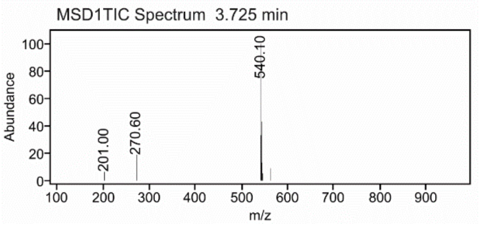

### Mass Spectrum SmartFormula Report

**Analysis Info**

| | | | |
|---|---|---|---|
| | | Acquisition Date | 2/19/2024 2:19:33 PM |
| Analysis Name | D:\Data\MS_Service\Service_Kaiser\2024_Q1\20240219_SKN_FGC337_5ug-mL_5uL-m.d | | |
| Method | DirectInfusion_WithFlow_Tune_pos.m | Operator | BDAL@DE |
| Sample Name | Test1 | Instrument | micrOTOF-Q 228888.10407 |
| Comment | | | |

**Acquisition Parameter**

| | | | | | |
|---|---|---|---|---|---|
| Source Type | ESI | Ion Polarity | Positive | Set Nebulizer | 1.8 Bar |
| Focus | Active | Set Capillary | 4000 V | Set Dry Heater | 220 °C |
| Scan Begin | 50 m/z | Set End Plate Offset | -500 V | Set Dry Gas | 8.0 l/min |
| Scan End | 1600 m/z | Set Collision Cell RF | 150.0 Vpp | Set Divert Valve | Waste |

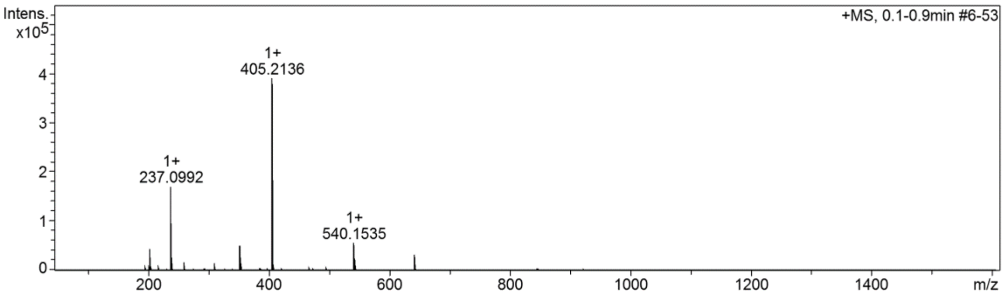

**Cmpd 52**

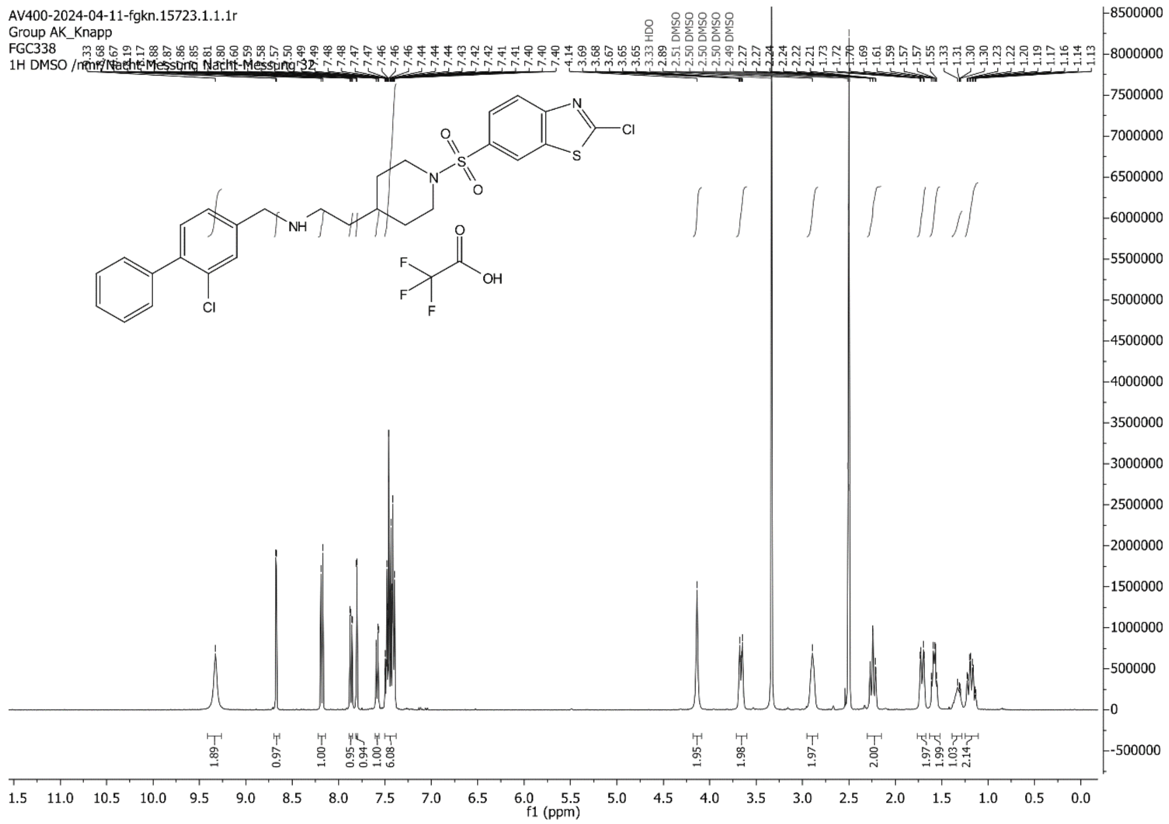
AV400-2024-04-11-fgkn.15723.1.1.1r
Group AK_Knapp
FGC338
1H DMSO /nmr/Nacht-Messung Nacht-Messung 32
f1 (ppm)

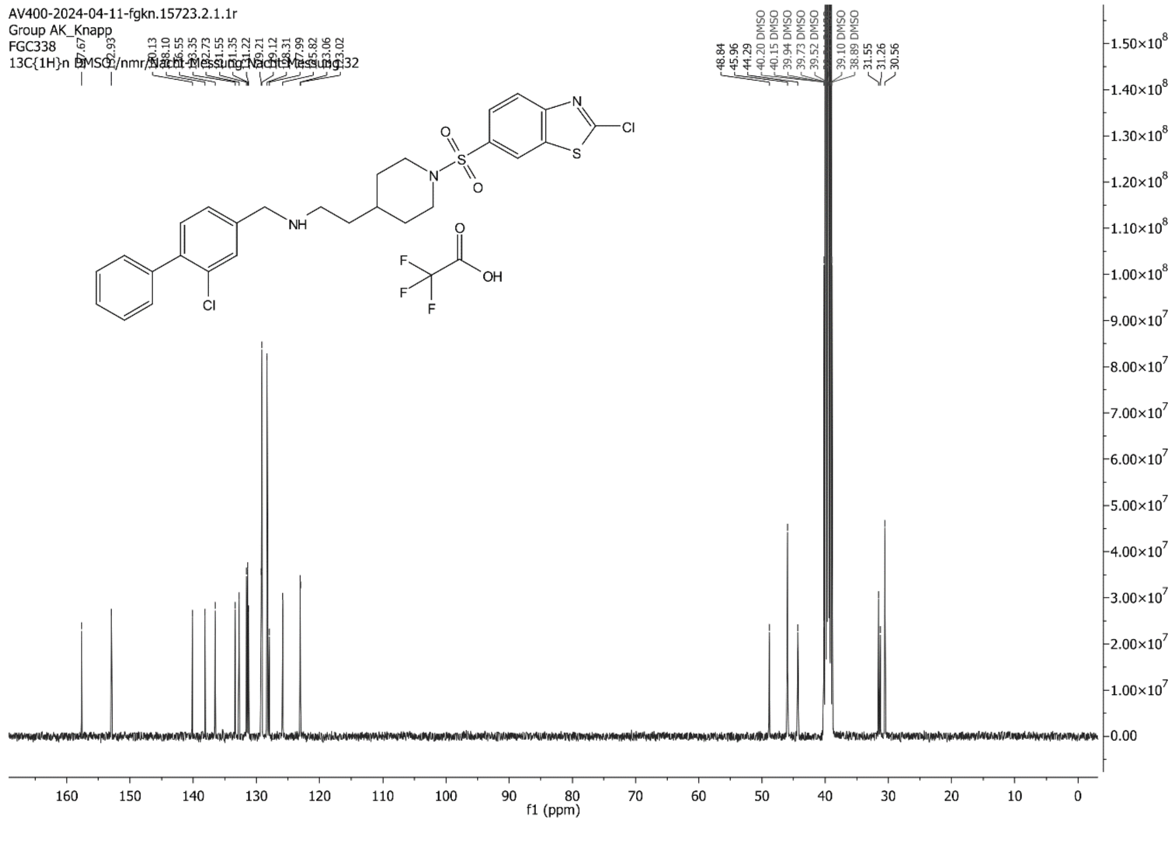
AV400-2024-04-11-fgkn.15723.2.1.1r
Group AK_Knapp
FGC338
13C{1H}n DMSO /nmr/Nacht-Messung Nacht-Messung 32
f1 (ppm)

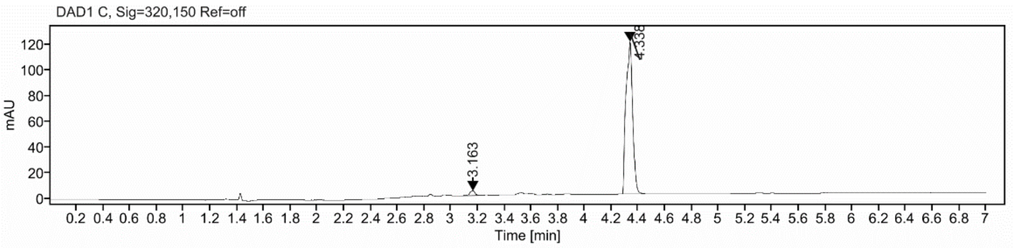

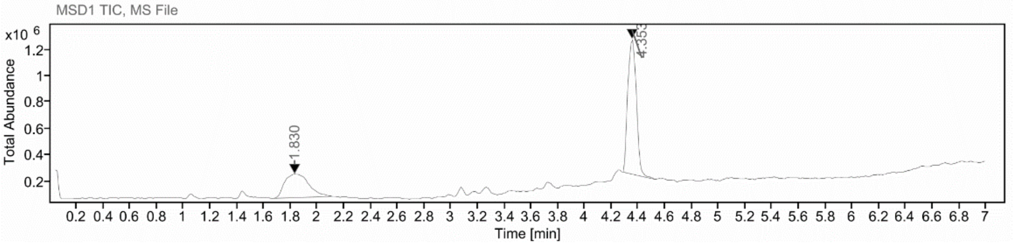

**Sample Purity**

| Signal Description | DAD1 C, Sig=320,150 Ref=off |
|---|---|

| Sample Name | Name | RT | Width | Area | Area% | Height |
|---|---|---|---|---|---|---|
| FGC338_re_f8 | | 3.163 | 0.042 | 9.8976 | 2.34 | 3.6013 |
| FGC338_re_f8 | | 4.338 | 0.060 | 413.0972 | 97.66 | 118.8883 |

| | |
|---|---|
| Max Area% | 97.660 |
| UV Signal Purity>95% | Pass |

**UV Apex Spectra**

**RT:**

4.338

**Sample Name:**

FGC338_re_f8

**Signal Name:**

DAD1C

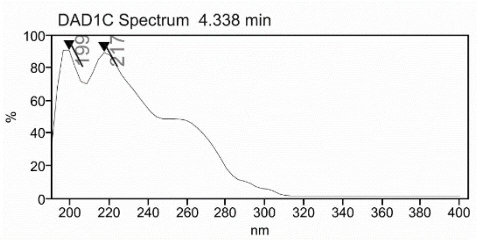

**RT:**

4.353

**Sample Name:**

FGC338_re_f8

**Signal Name:**

MSD1TIC

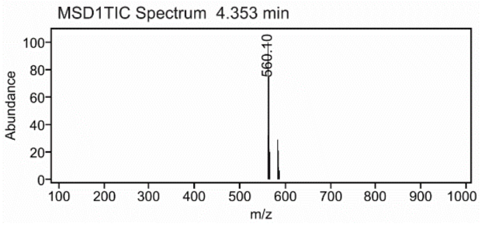

### Mass Spectrum SmartFormula Report

**Analysis Info**

| | | | |
|---|---|---|---|
| | | Acquisition Date | 2/27/2024 2:26:22 PM |
| Analysis Name | D:\Data\MS_Service\Service_Kaiser\2024_Q1\20240227_SKN_FGC338_100x_5uL-m.d | | |
| Method | DirectInfusion_WithFlow_Tune_pos.m | Operator | BDAL@DE |
| Sample Name | Test1 | Instrument | micrOTOF-Q 228888.10407 |
| Comment | | | |

**Acquisition Parameter**

| | | | | | |
|---|---|---|---|---|---|
| Source Type | ESI | Ion Polarity | Positive | Set Nebulizer | 1.8 Bar |
| Focus | Active | Set Capillary | 4000 V | Set Dry Heater | 300 °C |
| Scan Begin | 50 m/z | Set End Plate Offset | -500 V | Set Dry Gas | 8.0 l/min |
| Scan End | 1600 m/z | Set Collision Cell RF | 150.0 Vpp | Set Divert Valve | Waste |

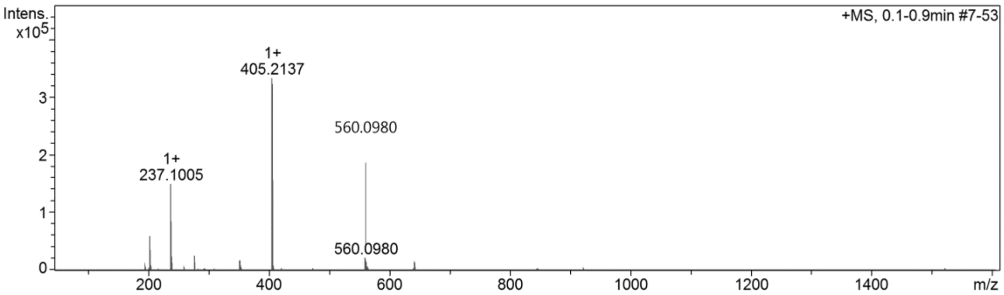

**Cmpd 53**

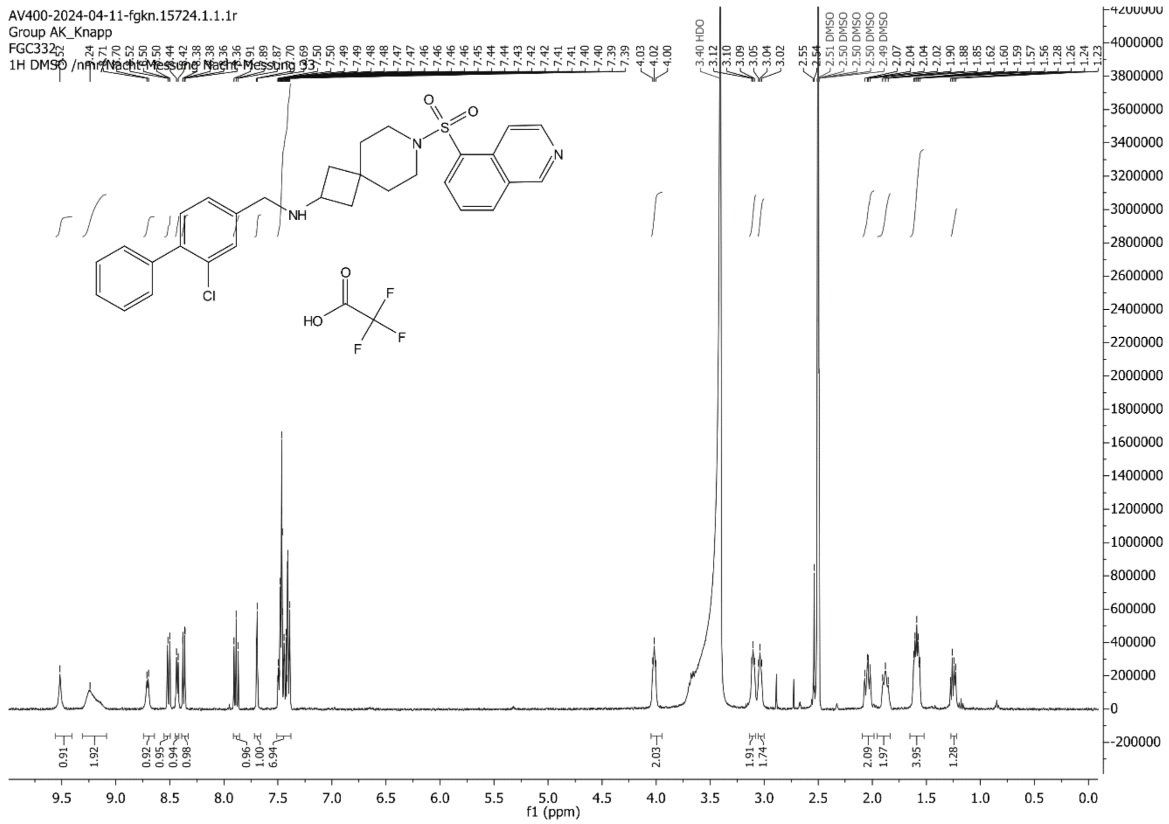

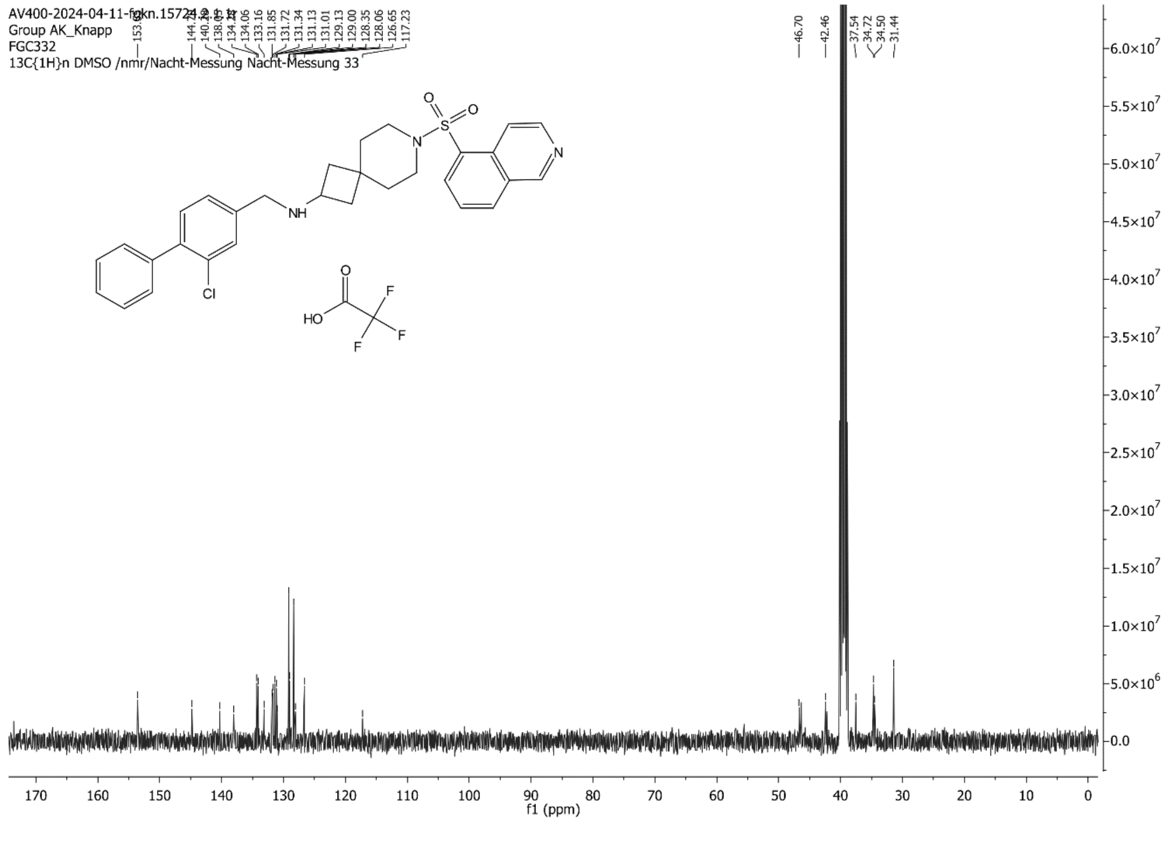

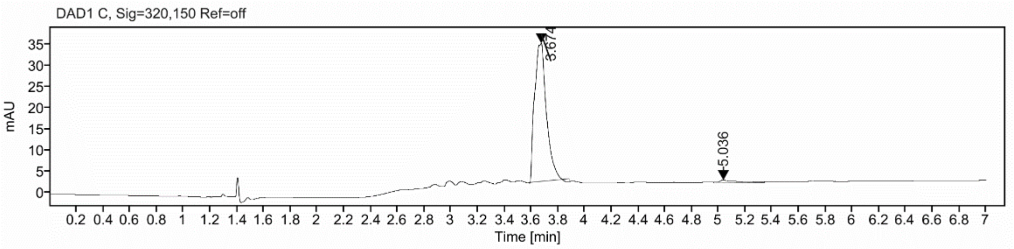

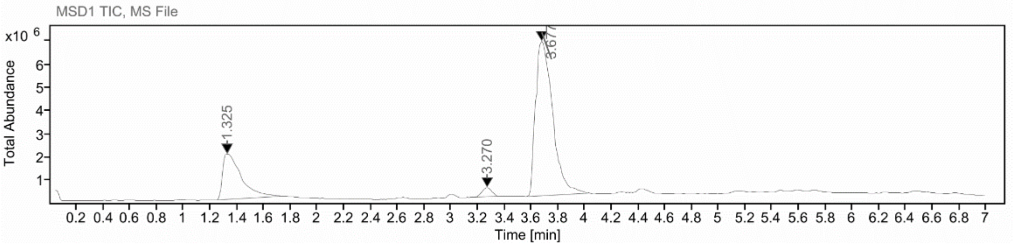

**Sample Purity**

Signal Description DAD1 C, Sig=320,150 Ref=off

| Sample Name | Name | RT | Width | Area | Area% | Height |
|---|---|---|---|---|---|---|
| FGC332 | | 3.674 | 0.099 | 199.3139 | 96.95 | 32.6430 |
| FGC332 | | 5.036 | 0.135 | 6.2690 | 3.05 | 0.6369 |

Max Area% 96.951

UV Signal Purity>95% Pass

**UV Apex Spectra**

**RT:**

3.674

**Sample Name:**

FGC332

**Signal Name:**

DAD1C

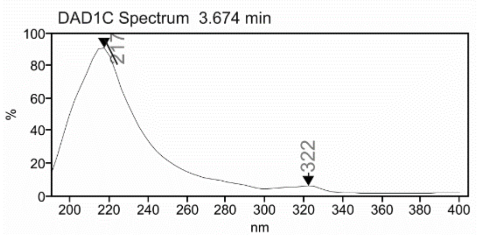

**RT:**

3.677

**Sample Name:**

FGC332

**Signal Name:**

MSD1TIC

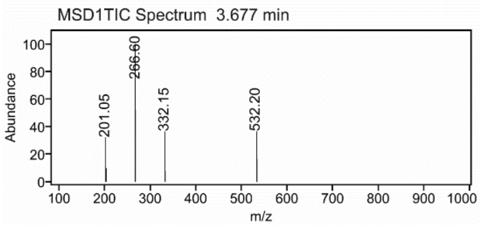

### Mass Spectrum SmartFormula Report

**Analysis Info**

| | | | |
|---|---|---|---|
| Analysis Name | D:\Data\MS_Service\2024Q1\20240319_SKN_FG_FGC332\20240319_SKN_FG_FGC332.d | Acquisition Date | 3/19/2024 4:24:10 PM |
| Method | DirectInfusion_WithFlow_Tune_pos.m | Operator | BDAL@DE |
| Sample Name | Test1 | Instrument | micrOTOF-Q 228888.10407 |
| Comment | | | |

**Acquisition Parameter**

| | | | | | |
|---|---|---|---|---|---|
| Source Type | ESI | Ion Polarity | Positive | Set Nebulizer | 1.8 Bar |
| Focus | Active | Set Capillary | 4000 V | Set Dry Heater | 300 °C |
| Scan Begin | 50 m/z | Set End Plate Offset | -500 V | Set Dry Gas | 8.0 l/min |
| Scan End | 1600 m/z | Set Collision Cell RF | 150.0 Vpp | Set Divert Valve | Waste |

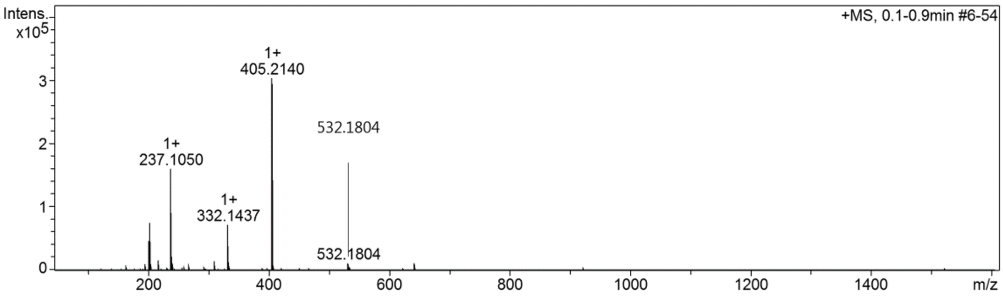

#### Cmpd 54

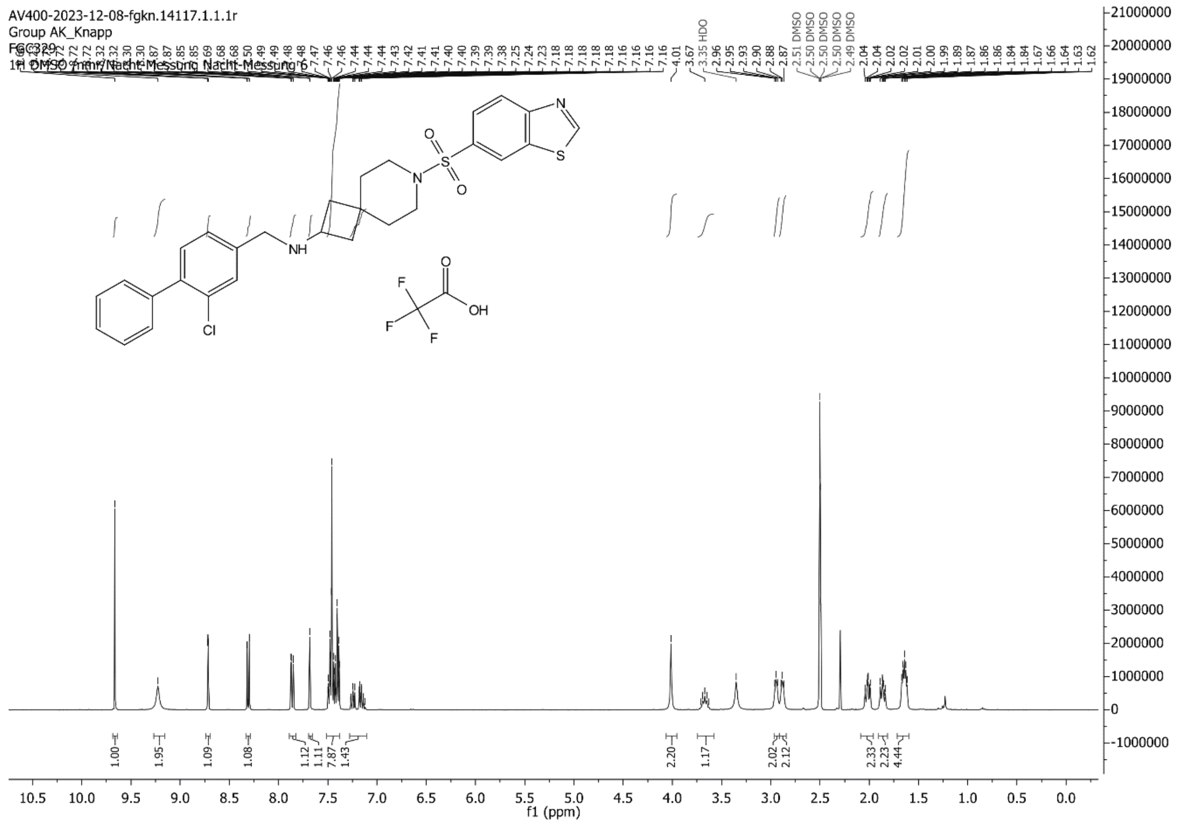
AV400-2023-12-08-fgkn.14117.1.1.1r
Group AK_Knapp
FGC329
1H DMSO /nmr/Nacht-Messung Nacht-Messung 6
NH
N
O
S
O
N
S
Cl
F
F
F
O
OH
f1 (ppm)

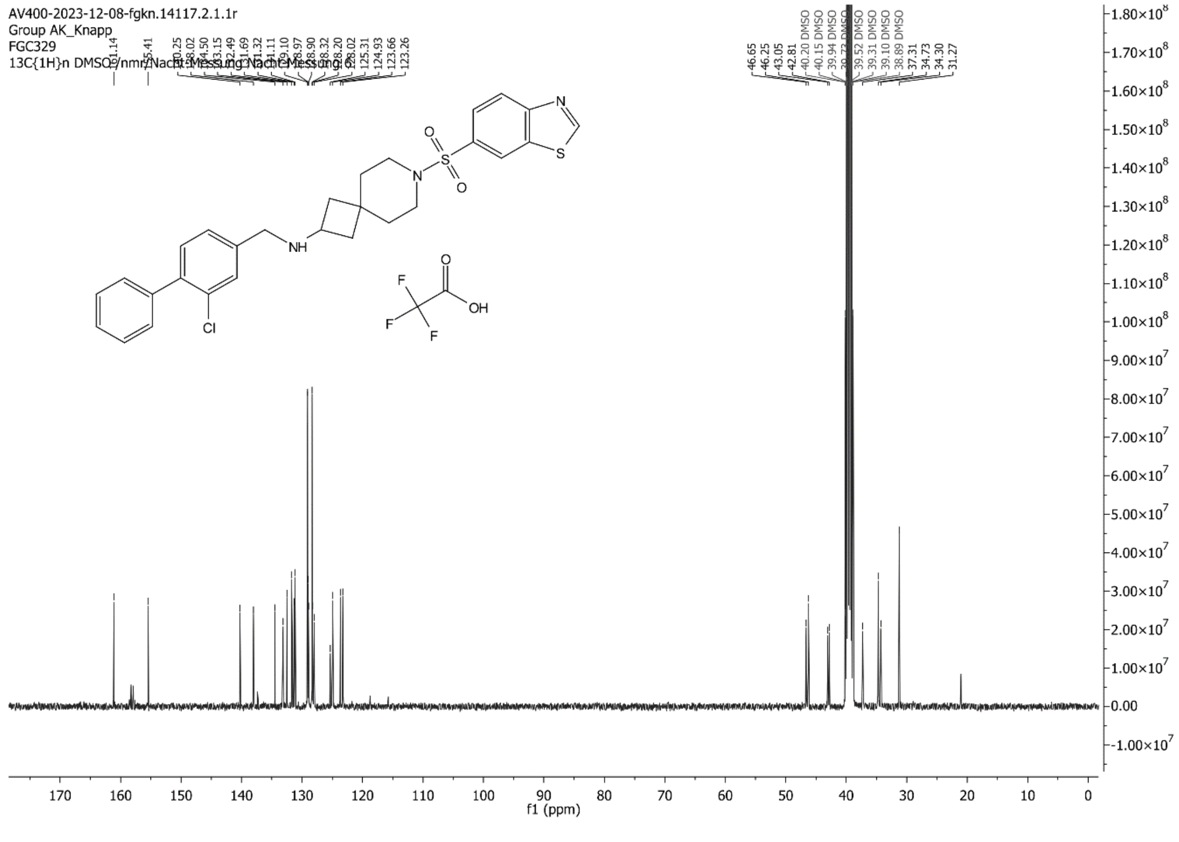
AV400-2023-12-08-fgkn.14117.2.1.1r
Group AK_Knapp
FGC329
13C{1H}n DMSO /nmr/Nacht-Messung Nacht-Messung 8
NH
N
O
S
O
N
S
Cl
F
F
F
O
OH
f1 (ppm)

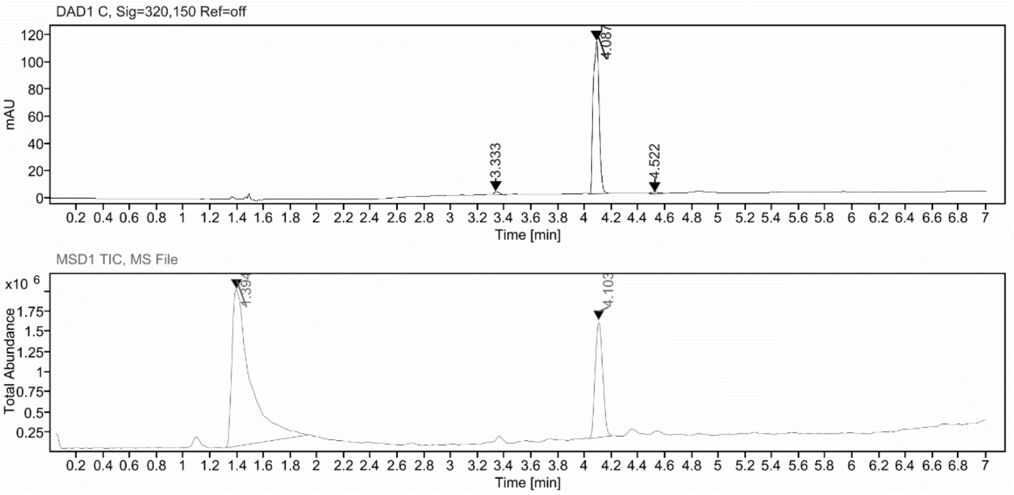

**Sample Purity**

| Signal Description | DAD1 C, Sig=320,150 Ref=off |
|---|---|

| Sample Name | Name | RT | Width | Area | Area% | Height |
|---|---|---|---|---|---|---|
| FGC329 | | 3.333 | 0.039 | 5.9687 | 1.68 | 2.1117 |
| FGC329 | | 4.087 | 0.052 | 344.9969 | 96.91 | 112.1918 |
| FGC329 | | 4.522 | 0.092 | 5.0293 | 1.41 | 0.9881 |

| | |
|---|---|
| Max Area% | 96.911 |
| UV Signal Purity>95% | Pass |

**UV Apex Spectra**

**RT:**

4.087

**Sample Name:**

FGC329

**Signal Name:**

DAD1C

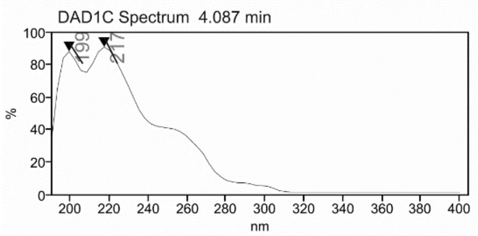

**RT:**

4.103

**Sample Name:**

FGC329

**Signal Name:**

MSD1TIC

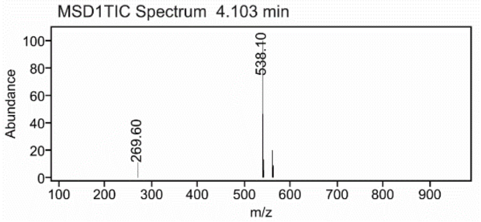

### Mass Spectrum SmartFormula Report

**Analysis Info**

| | | | |
|---|---|---|---|
| | | Acquisition Date | 2/12/2024 1:47:12 PM |
| Analysis Name | D:\Data\MS_Service\Service_Kaiser\2024_Q1\20240212_SKN_FGC329_5ug-mL_3uL-m.d | | |
| Method | DirectInfusion_WithFlow_Tune_pos.m | Operator | BDAL@DE |
| Sample Name | Test1 | Instrument | micrOTOF-Q 228888.10407 |
| Comment | | | |

**Acquisition Parameter**

| | | | | | |
|---|---|---|---|---|---|
| Source Type | ESI | Ion Polarity | Positive | Set Nebulizer | 1.8 Bar |
| Focus | Active | Set Capillary | 4000 V | Set Dry Heater | 220 °C |
| Scan Begin | 50 m/z | Set End Plate Offset | -500 V | Set Dry Gas | 8.0 l/min |
| Scan End | 1600 m/z | Set Collision Cell RF | 150.0 Vpp | Set Divert Valve | Waste |

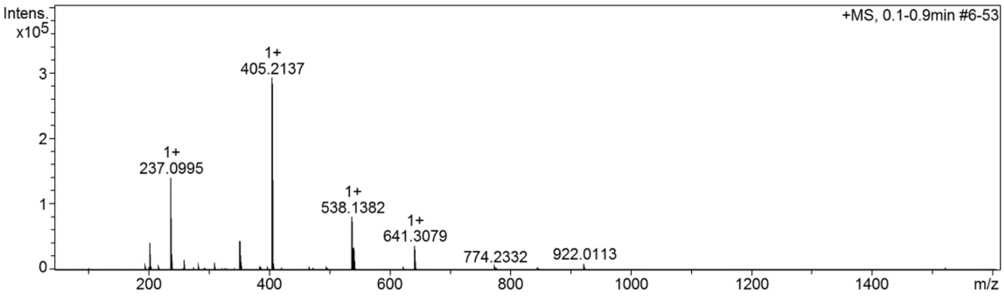

**Cmpd 55**

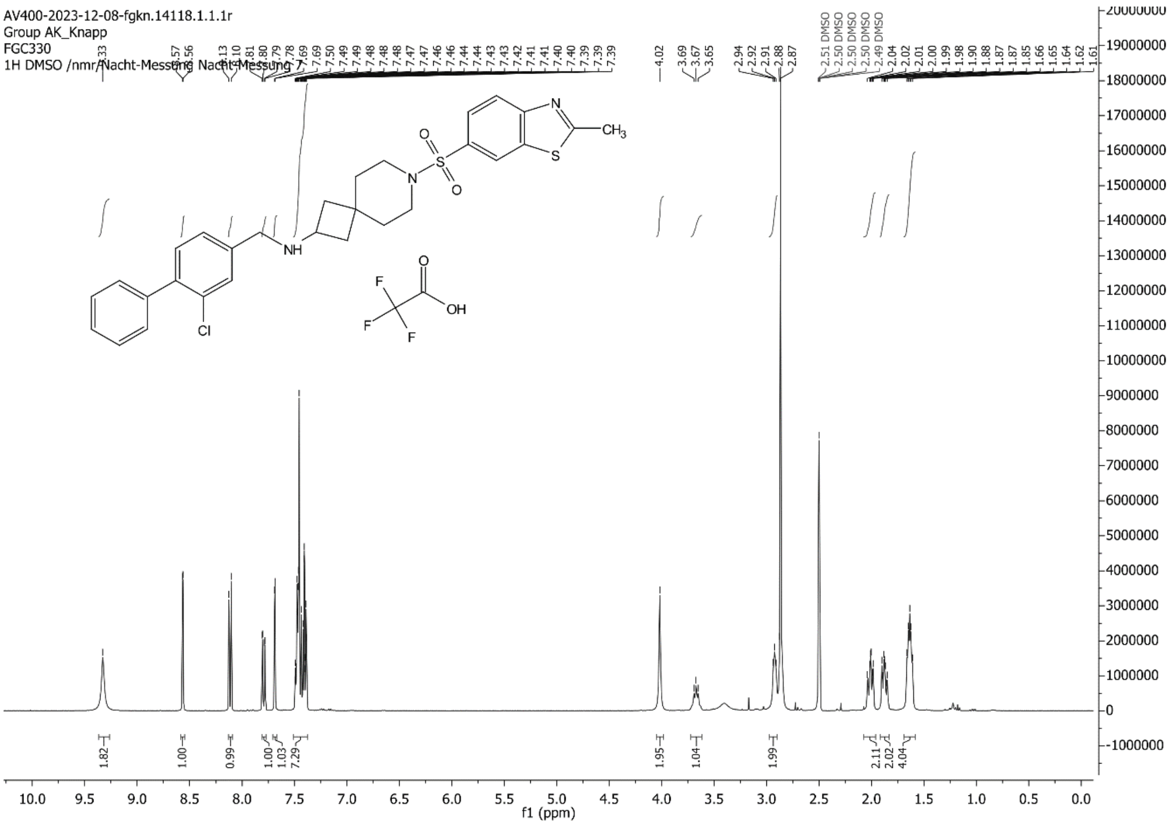

AV400-2023-12-08-fgkn.14118.1.1.1r
Group AK_Knapp
FGC330
1H DMSO /nmr/Nacht-Messung Nacht-Messung 7
f1 (ppm)

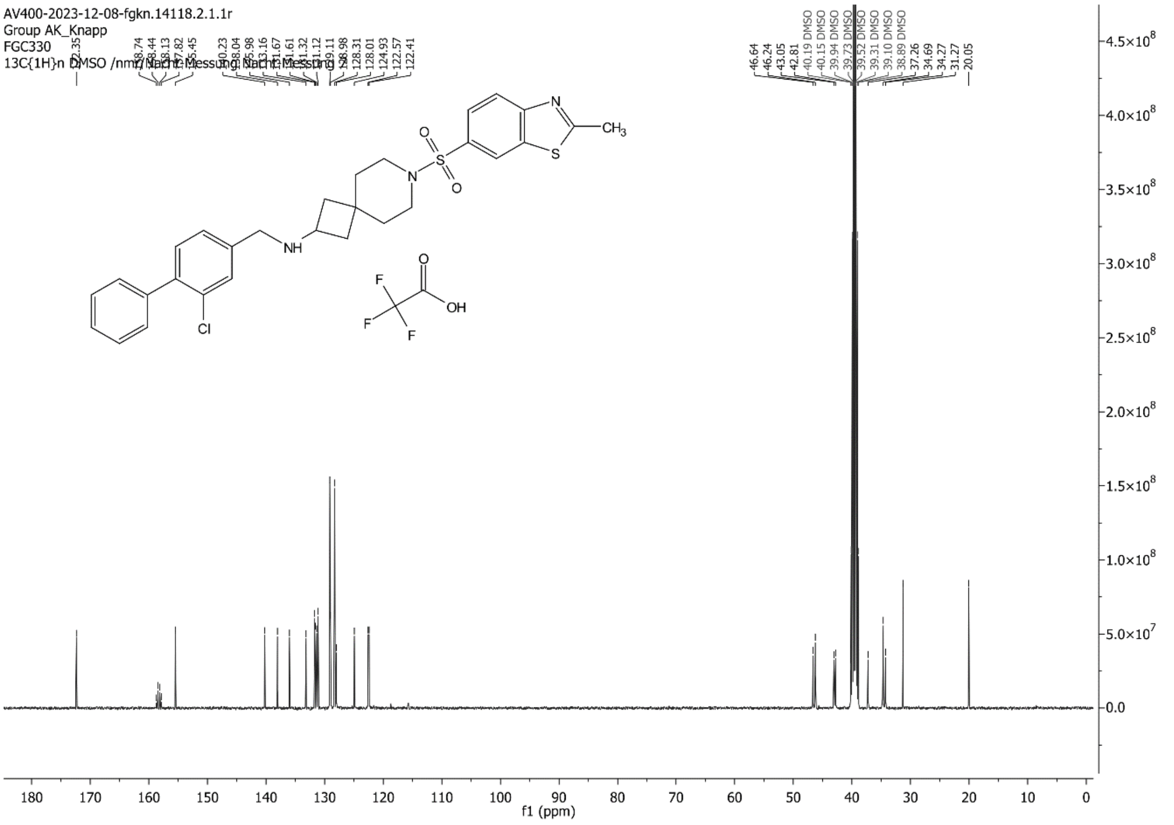

AV400-2023-12-08-fgkn.14118.2.1.1r
Group AK_Knapp
FGC330
13C{1H}n DMSO /nmr/Nacht-Messung Nacht-Messung 7
f1 (ppm)

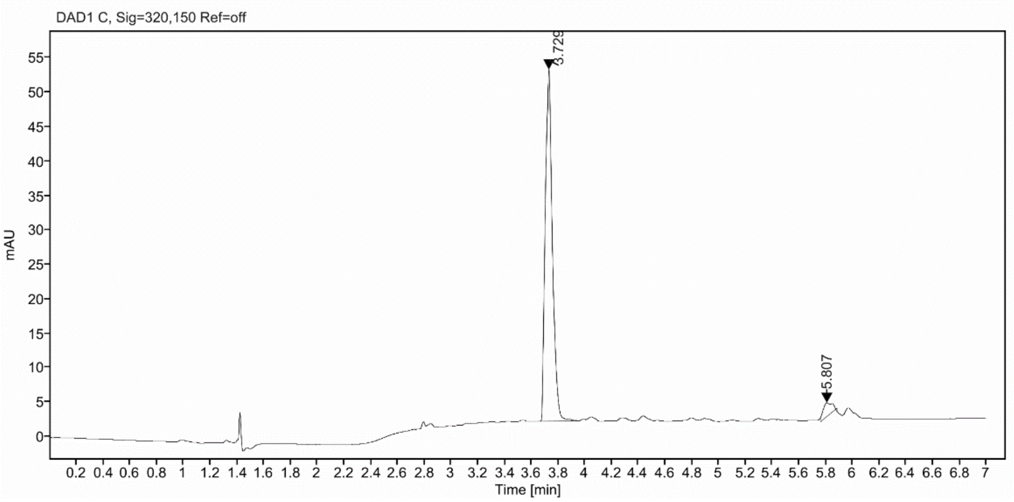

**Sample Purity**

Signal Description DAD1 C, Sig=320,150 Ref=off

| Sample Name | Name | RT | Width | Area | Area% | Height |
|---|---|---|---|---|---|---|
| FGC330_f13 | | 3.729 | 0.058 | 189.7316 | 95.49 | 51.1395 |
| FGC330_f13 | | 5.807 | 0.097 | 8.9696 | 4.51 | 1.9918 |

Max Area% 95.486

UV Signal Purity>95% Pass

**UV Apex Spectra**

**RT:**

3.729

**Sample Name:**

FGC330_f13

**Signal Name:**

DAD1C

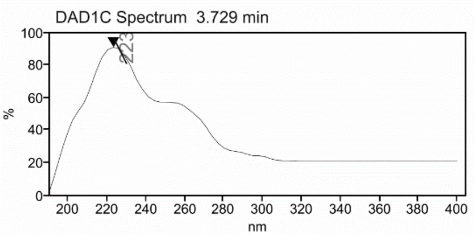

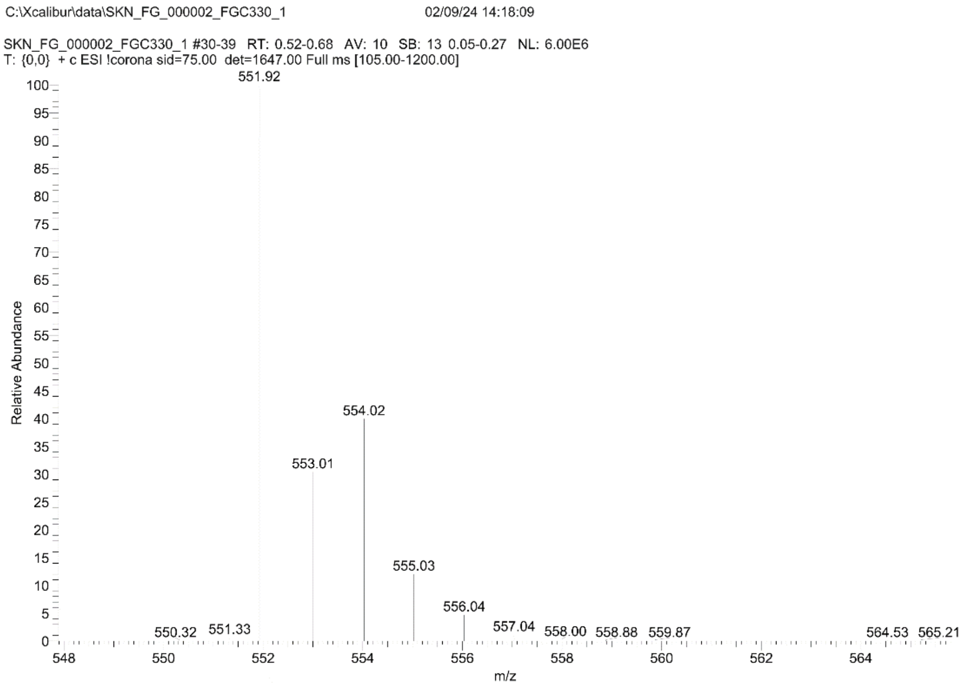
C:\Xcalibur\data\SKN_FG_000002_FGC330_1
02/09/24 14:18:09
SKN_FG_000002_FGC330_1 #30-39 RT: 0.52-0.68 AV: 10 SB: 13 0.05-0.27 NL: 6.00E6
T: {0,0} + c ESI !corona sid=75.00 det=1647.00 Full ms [105.00-1200.00]
Relative Abundance
m/z
551.92
554.02
553.01
555.03
556.04
557.04
558.00
558.88
559.87
550.32
551.33
564.53
565.21

### Mass Spectrum SmartFormula Report

**Analysis Info**

| | | | |
|---|---|---|---|
| Analysis Name | D:\Data\MS_Service\Service_Kaiser\2024_Q1\20240212_SKN_FGC330_5ug-mL_3uL-m.d | Acquisition Date | 2/12/2024 1:56:48 PM |
| Method | DirectInfusion_WithFlow_Tune_pos.m | Operator | BDAL@DE |
| Sample Name | Test1 | Instrument | micrOTOF-Q 228888.10407 |
| Comment | | | |

**Acquisition Parameter**

| | | | | | |
|---|---|---|---|---|---|
| Source Type | ESI | Ion Polarity | Positive | Set Nebulizer | 1.8 Bar |
| Focus | Active | Set Capillary | 4000 V | Set Dry Heater | 220 °C |
| Scan Begin | 50 m/z | Set End Plate Offset | -500 V | Set Dry Gas | 8.0 l/min |
| Scan End | 1600 m/z | Set Collision Cell RF | 150.0 Vpp | Set Divert Valve | Waste |

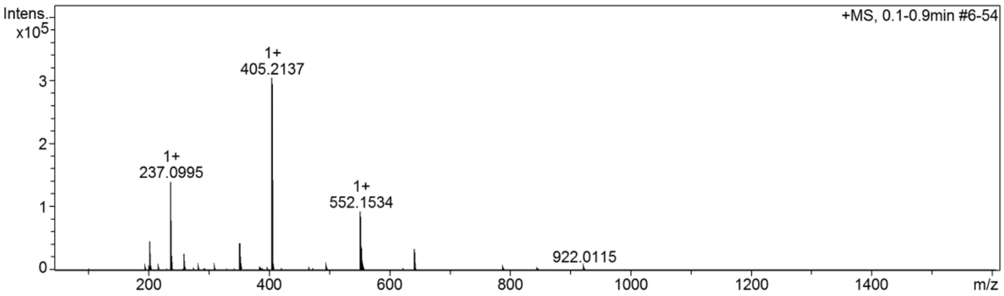

#### Cmpd 56

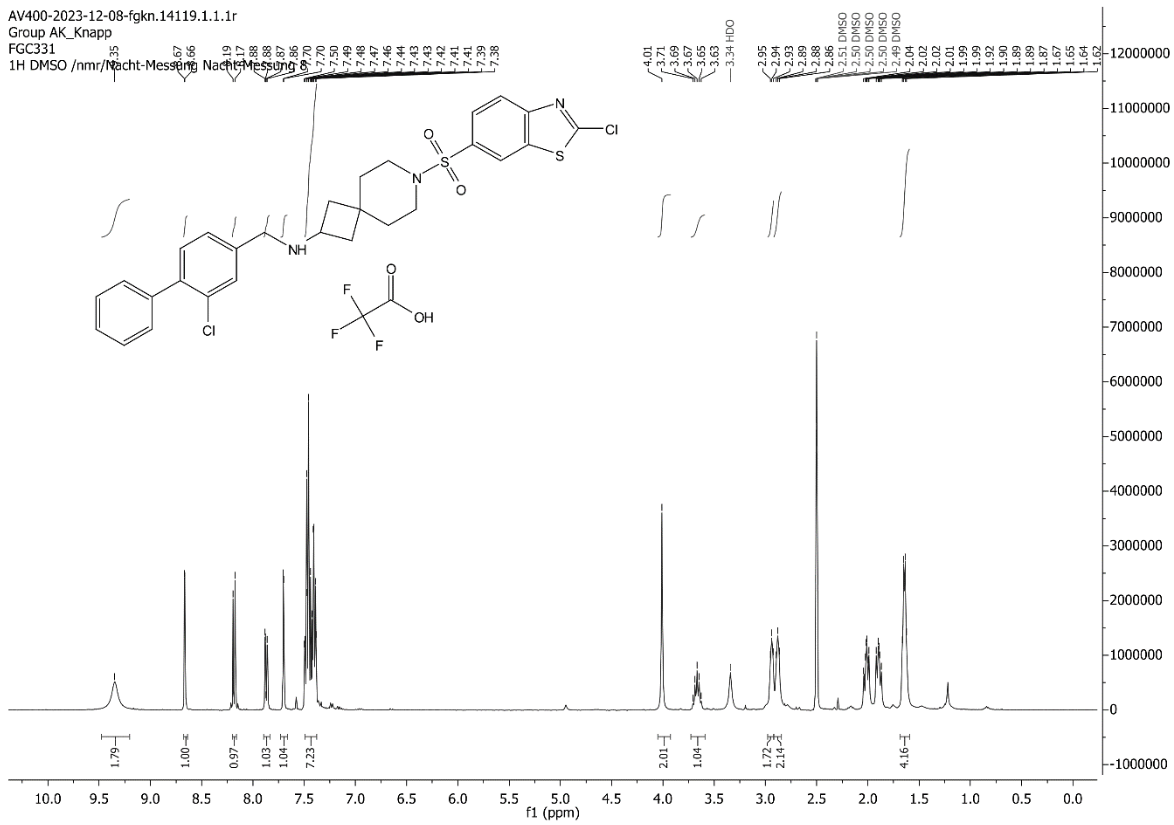
AV400-2023-12-08-fgkn.14119.1.1.1r
Group AK_Knapp
FGC331
1H DMSO /nmr/Nacht-Messung Nacht-Messung 8
f1 (ppm)

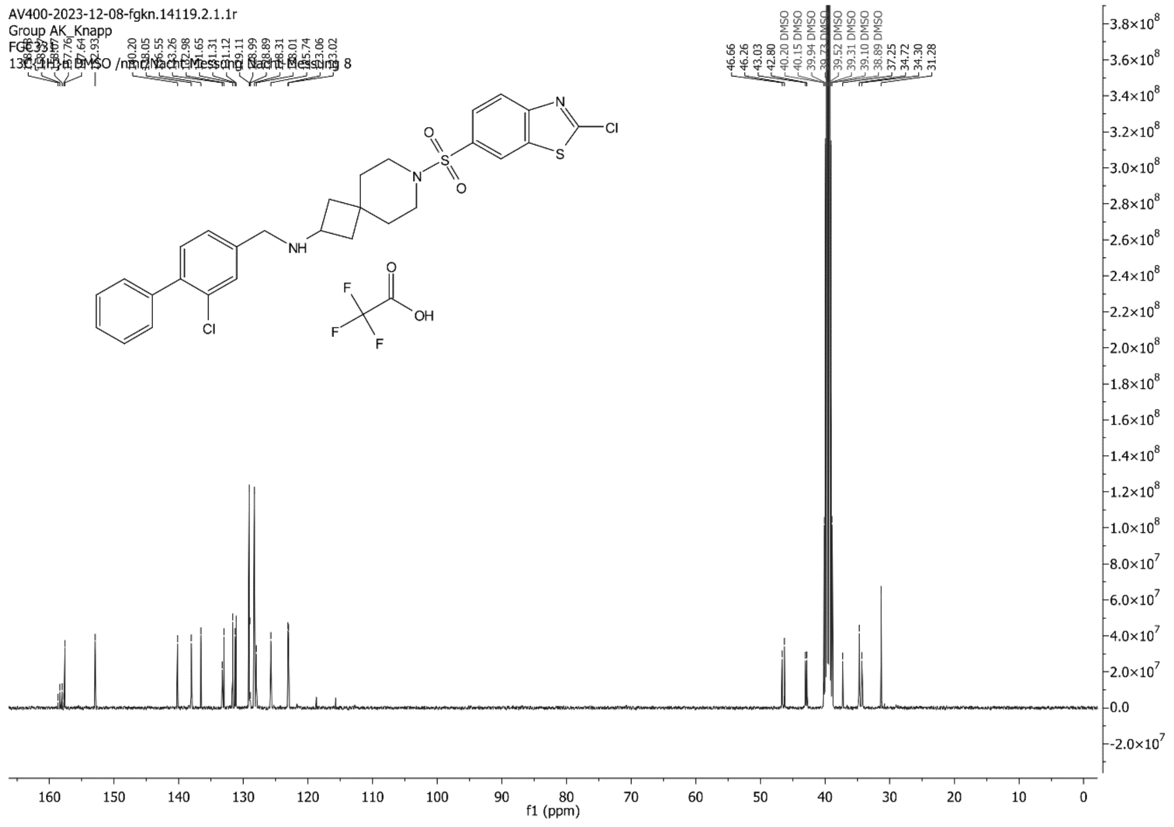
AV400-2023-12-08-fgkn.14119.2.1.1r
Group AK_Knapp
FGC331
13C{1H} DMSO /nmr/Nacht-Messung Nacht-Messung 8
f1 (ppm)

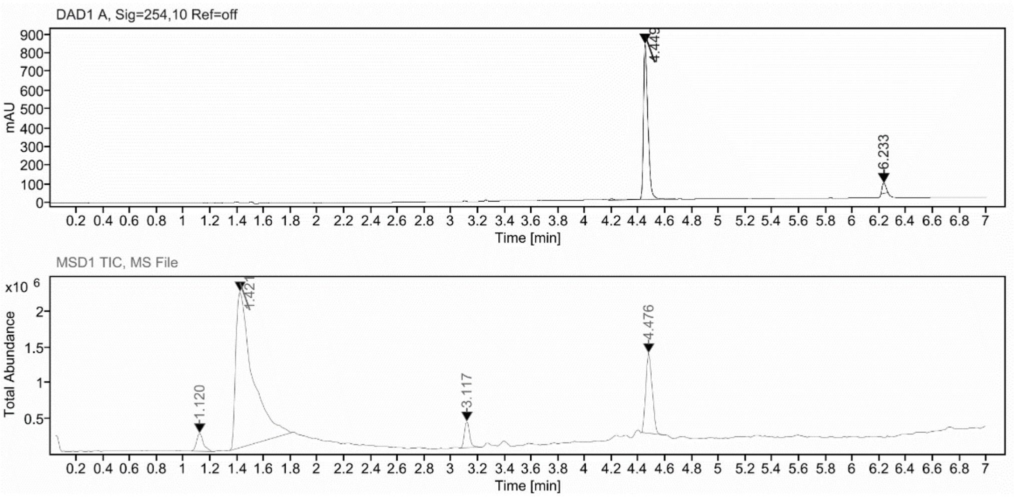

**Sample Purity**

| Signal Description | DAD1 A, Sig=254,10 Ref=off |
|---|---|

| Sample Name | Name | RT | Width | Area | Area% | Height |
|---|---|---|---|---|---|---|
| FGC331_X | | 4.449 | 0.036 | 1947.9377 | 95.04 | 830.1127 |
| FGC331_X | | 6.233 | 0.033 | 101.7257 | 4.96 | 55.5746 |

| | |
|---|---|
| Max Area% | 95.037 |
| UV Signal Purity>95% | Pass |

**UV Apex Spectra**

**RT:**

4.449

**Sample Name:**

FGC331_X

**Signal Name:**

DAD1A

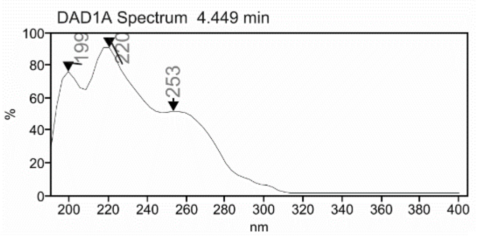

**RT:**

4.476

**Sample Name:**

FGC331_X

**Signal Name:**

MSD1TIC

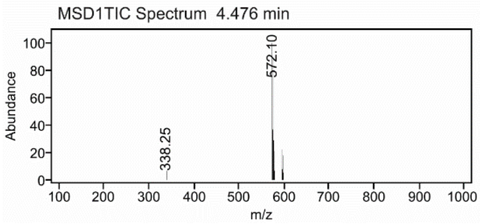

### Mass Spectrum SmartFormula Report

**Analysis Info**

| | | | |
|---|---|---|---|
| Analysis Name | D:\Data\MS_Service\Service_Kaiser\2024_Q1\20240212_SKN_FGC331_5ug-mL_3uL-m.d | Acquisition Date | 2/12/2024 2:11:49 PM |
| Method | DirectInfusion_WithFlow_Tune_pos.m | Operator | BDAL@DE |
| Sample Name | Test1 | Instrument | micrOTOF-Q 228888.10407 |
| Comment | | | |

**Acquisition Parameter**

| | | | | | |
|---|---|---|---|---|---|
| Source Type | ESI | Ion Polarity | Positive | Set Nebulizer | 1.8 Bar |
| Focus | Active | Set Capillary | 4000 V | Set Dry Heater | 220 °C |
| Scan Begin | 50 m/z | Set End Plate Offset | -500 V | Set Dry Gas | 8.0 l/min |
| Scan End | 1600 m/z | Set Collision Cell RF | 150.0 Vpp | Set Divert Valve | Waste |

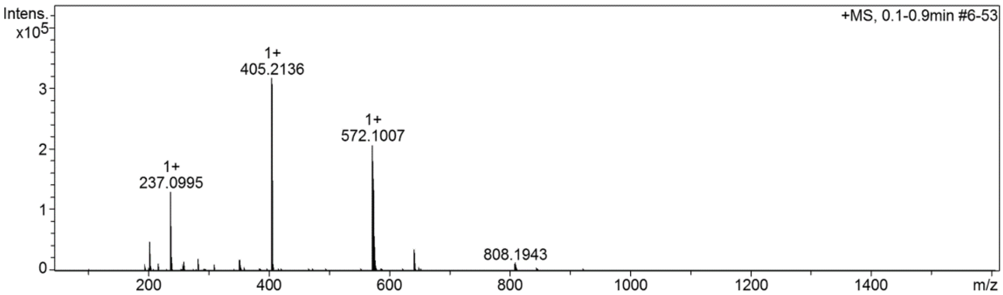

**Cmpd 57**

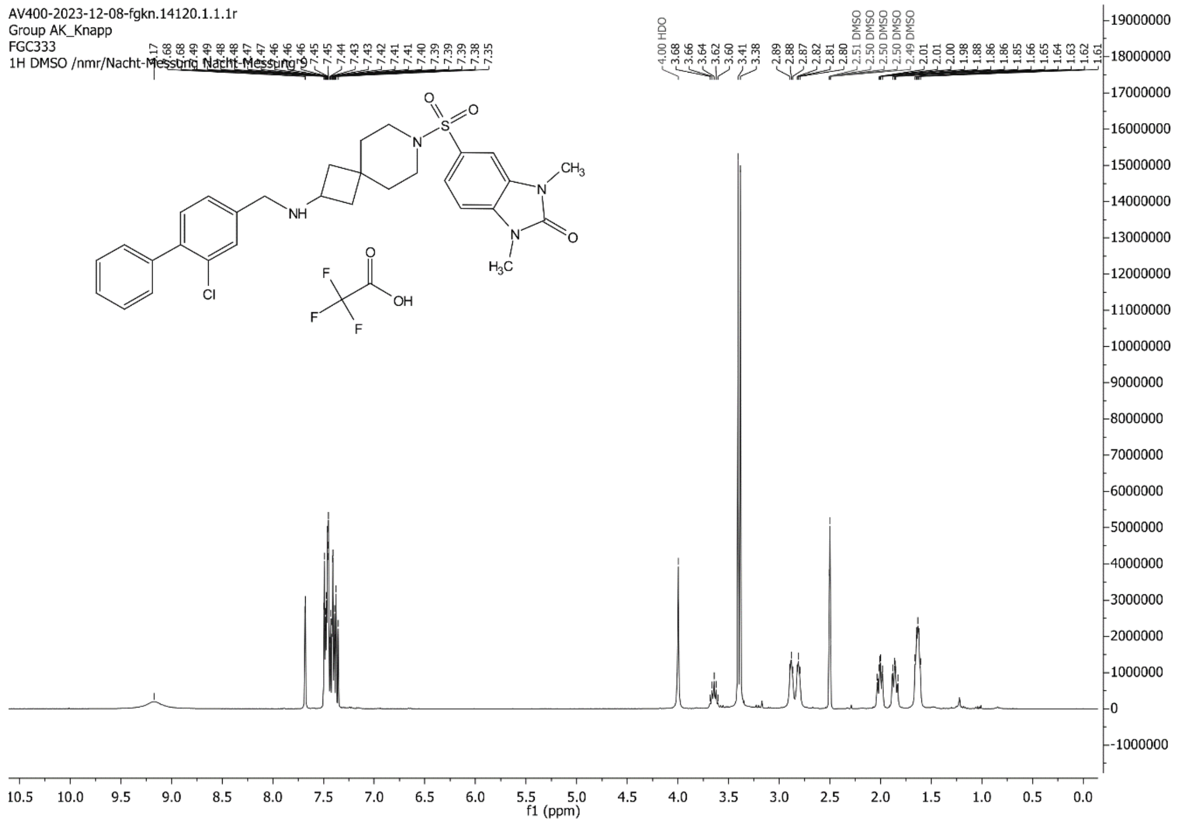
AV400-2023-12-08-fgkn.14120.1.1.1r
Group AK_Knapp
FGC333
1H DMSO /nmr/Nacht-Messung Nacht-Messung 9
f1 (ppm)

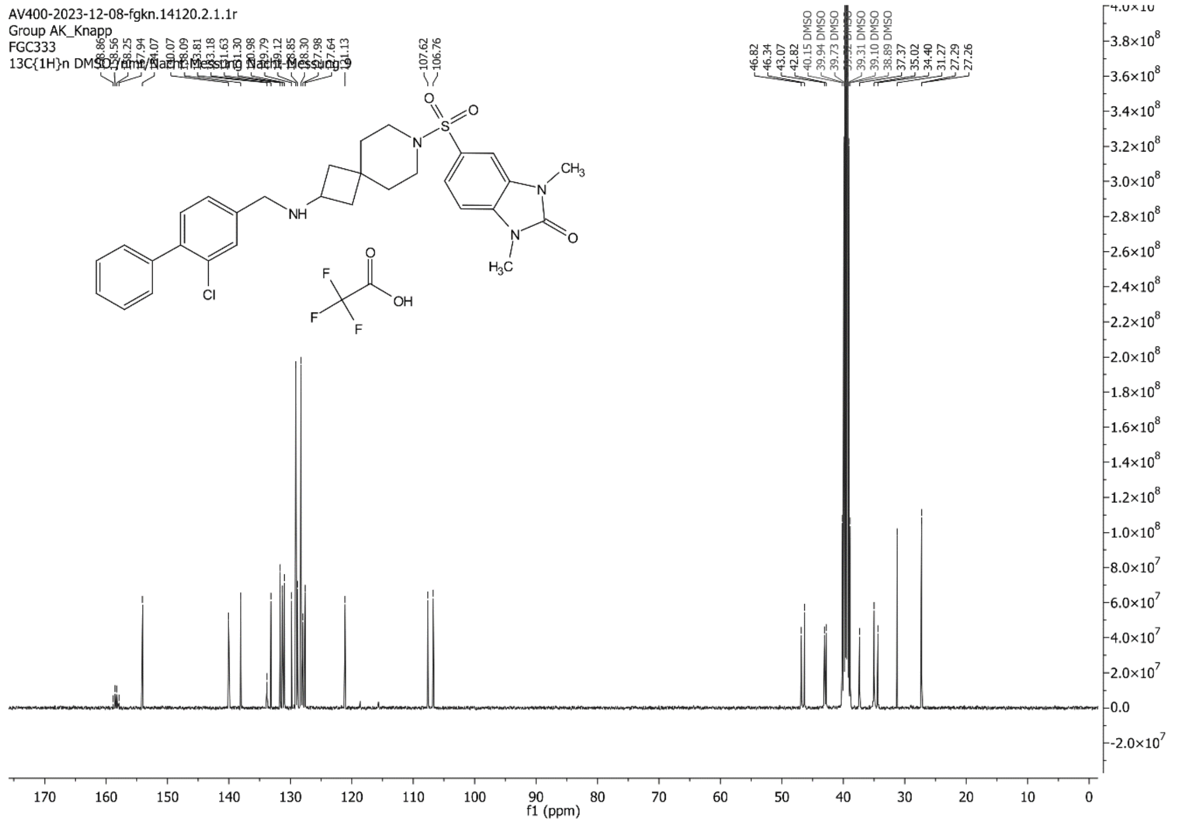
AV400-2023-12-08-fgkn.14120.2.1.1r
Group AK_Knapp
FGC333
13C{1H}n DMSO /nmr/Nacht-Messung Nacht-Messung 9
f1 (ppm)

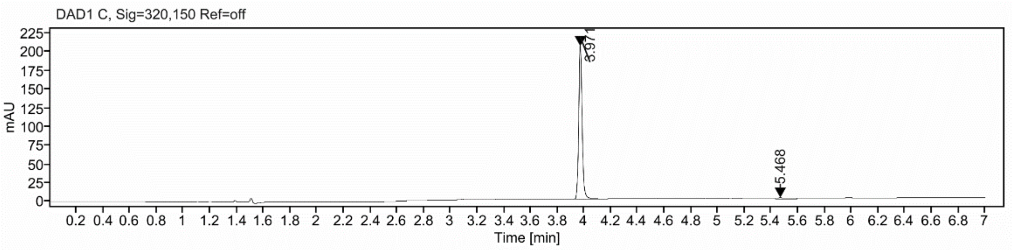

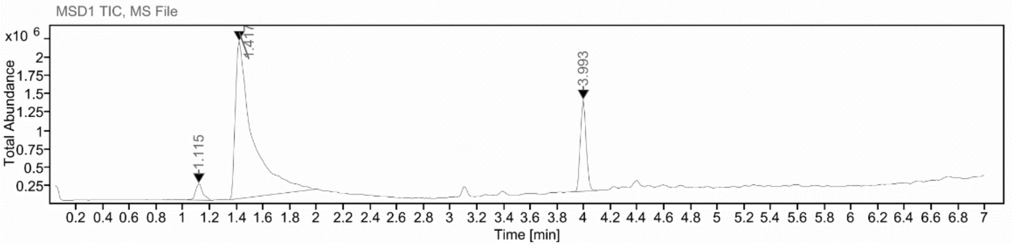

**Sample Purity**

| Signal Description | DAD1 C, Sig=320,150 Ref=off |
|---|---|

| Sample Name | Name | RT | Width | Area | Area% | Height |
|---|---|---|---|---|---|---|
| FGC333_X | | 3.971 | 0.025 | 356.2317 | 97.02 | 204.6024 |
| FGC333_X | | 5.468 | 0.160 | 10.9364 | 2.98 | 1.5322 |

| | |
|---|---|
| Max Area% | 97.021 |
| UV Signal Purity>95% | Pass |

**UV Apex Spectra**

**RT:**

3.971

**Sample Name:**

FGC333_X

**Signal Name:**

DAD1C

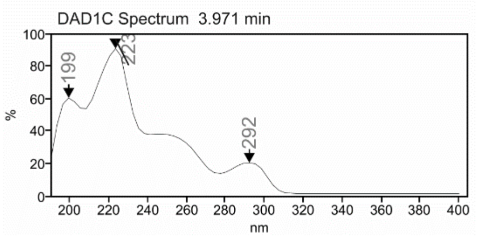

**RT:**

3.993

**Sample Name:**

FGC333_X

**Signal Name:**

MSD1TIC

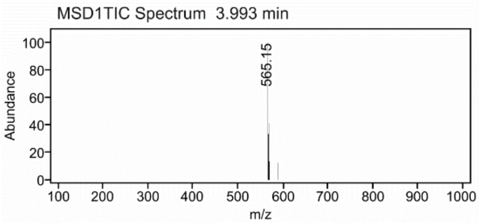

### Mass Spectrum SmartFormula Report

**Analysis Info**

| | | | |
|---|---|---|---|
| Analysis Name | D:\Data\MS_Service\Service_Kaiser\2024_Q1\20240219_SKN_FGC333_5ug-mL_5uL-m.d | Acquisition Date | 2/19/2024 1:47:36 PM |
| Method | DirectInfusion_WithFlow_Tune_pos.m | Operator | BDAL@DE |
| Sample Name | Test1 | Instrument | micrOTOF-Q 228888.10407 |
| Comment | | | |

**Acquisition Parameter**

| | | | | | |
|---|---|---|---|---|---|
| Source Type | ESI | Ion Polarity | Positive | Set Nebulizer | 1.8 Bar |
| Focus | Active | Set Capillary | 4000 V | Set Dry Heater | 220 °C |
| Scan Begin | 50 m/z | Set End Plate Offset | -500 V | Set Dry Gas | 8.0 l/min |
| Scan End | 1600 m/z | Set Collision Cell RF | 150.0 Vpp | Set Divert Valve | Waste |

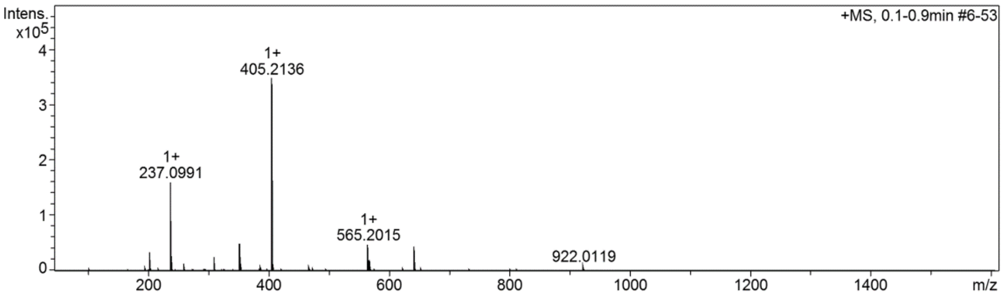

**Cmpd 58**

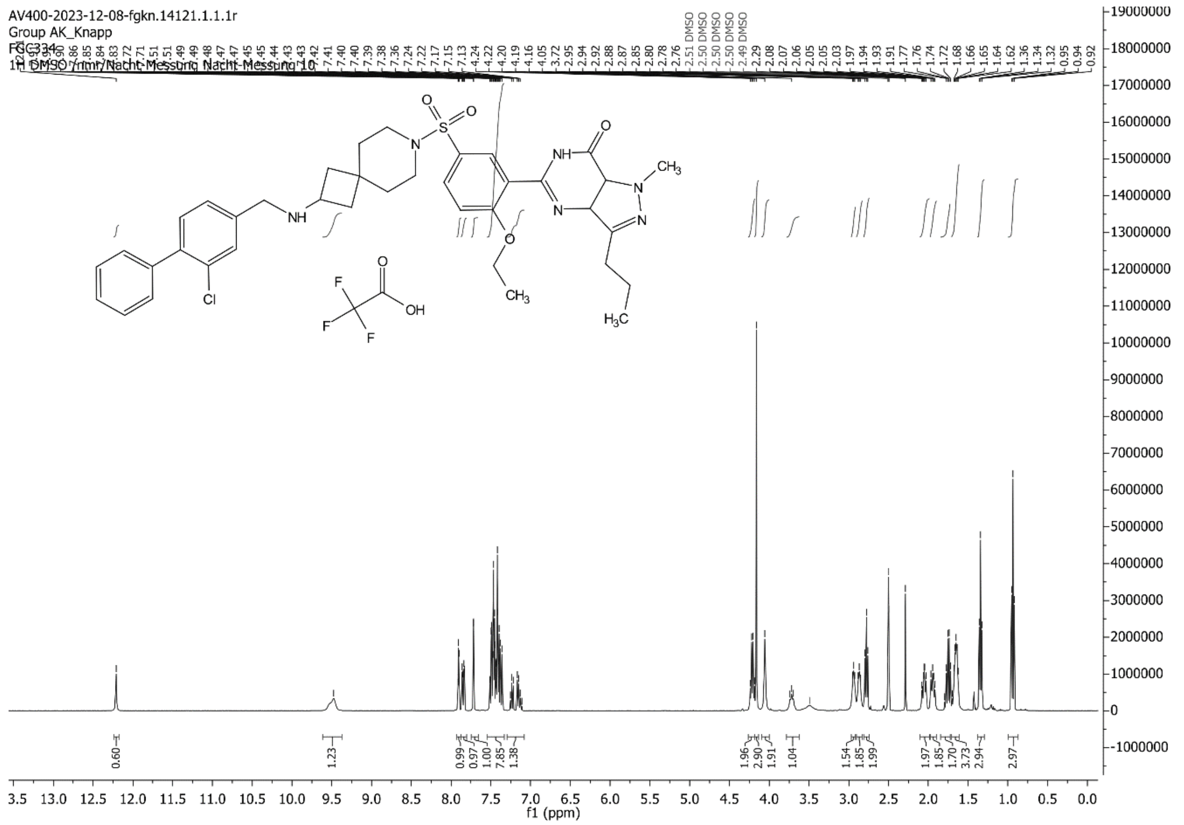

AV400-2023-12-08-fgkn.14121.1.1.1r
Group AK_Knapp
FGC334
1H DMSO /nmr/Nacht-Messung Nacht-Messung 10
f1 (ppm)

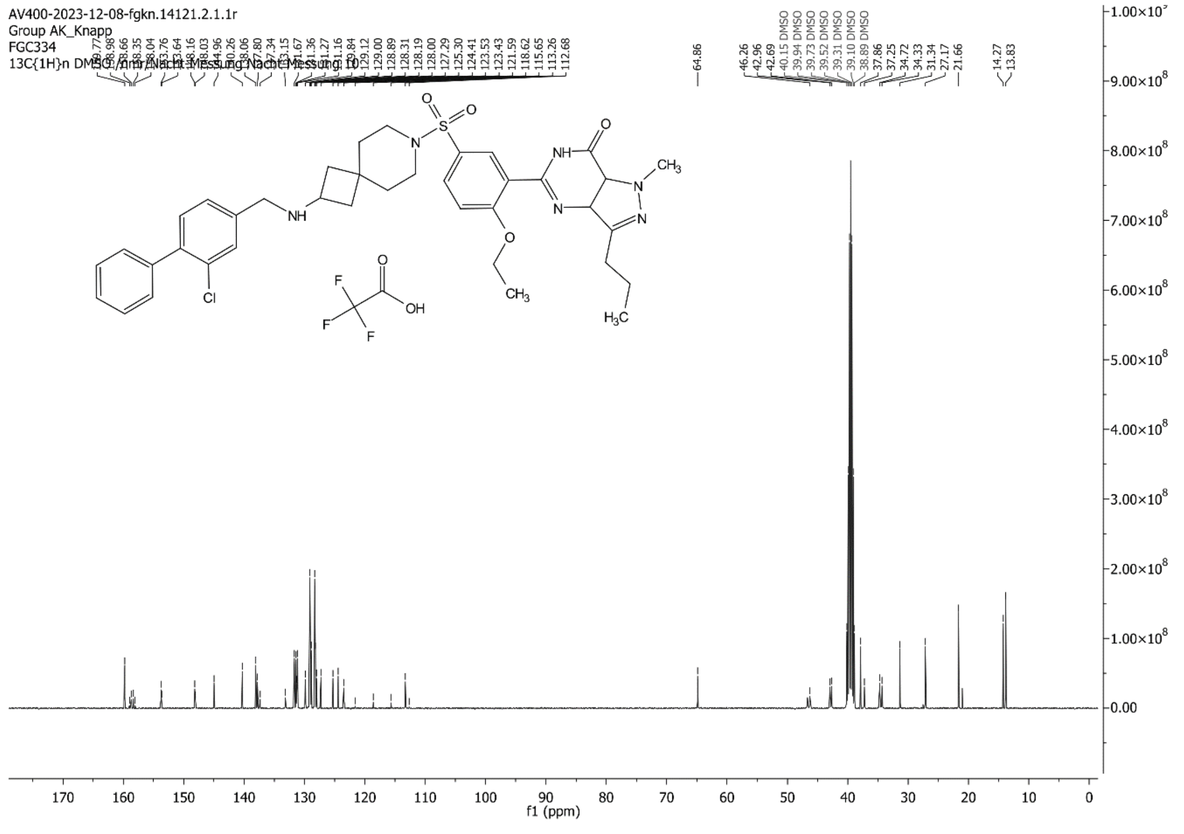

AV400-2023-12-08-fgkn.14121.2.1.1r
Group AK_Knapp
FGC334
13C{1H}n DMSO /nmr/Nacht-Messung Nacht-Messung 10
f1 (ppm)

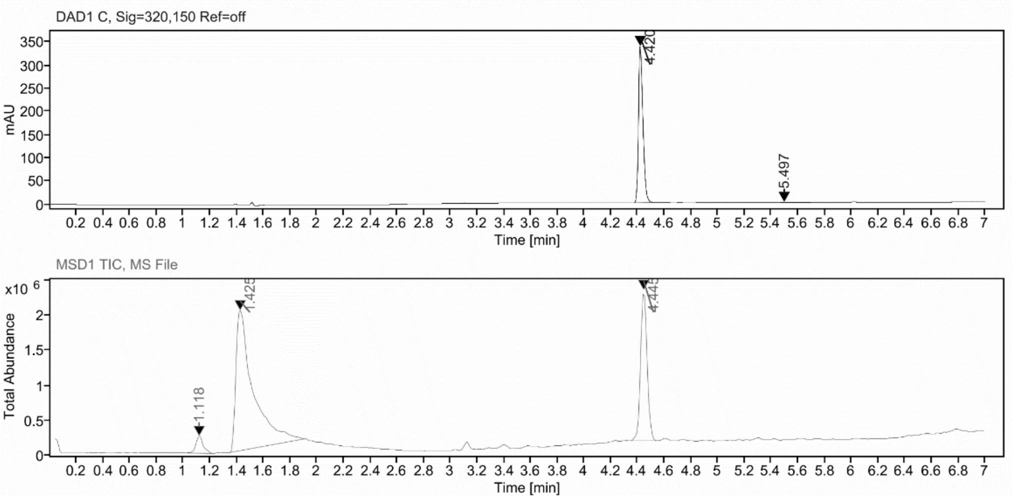

**Sample Purity**

Signal Description DAD1 C, Sig=320,150 Ref=off

| Sample Name | Name | RT | Width | Area | Area% | Height |
|---|---|---|---|---|---|---|
| FGC334_X | | 4.420 | 0.035 | 773.6117 | 98.67 | 337.7867 |
| FGC334_X | | 5.497 | 0.218 | 10.4063 | 1.33 | 1.3373 |

Max Area% 98.673

UV Signal Purity>95% Pass

**UV Apex Spectra**

**RT:**

4.420

**Sample Name:**

FGC334_X

**Signal Name:**

DAD1C

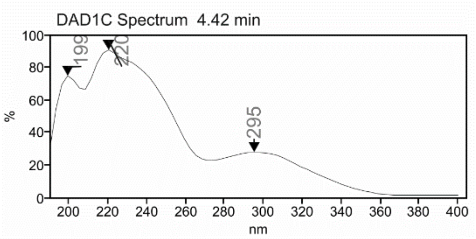

**RT:**

4.445

**Sample Name:**

FGC334_X

**Signal Name:**

MSD1TIC

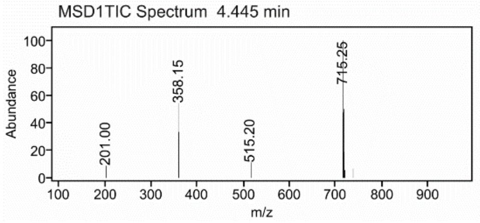

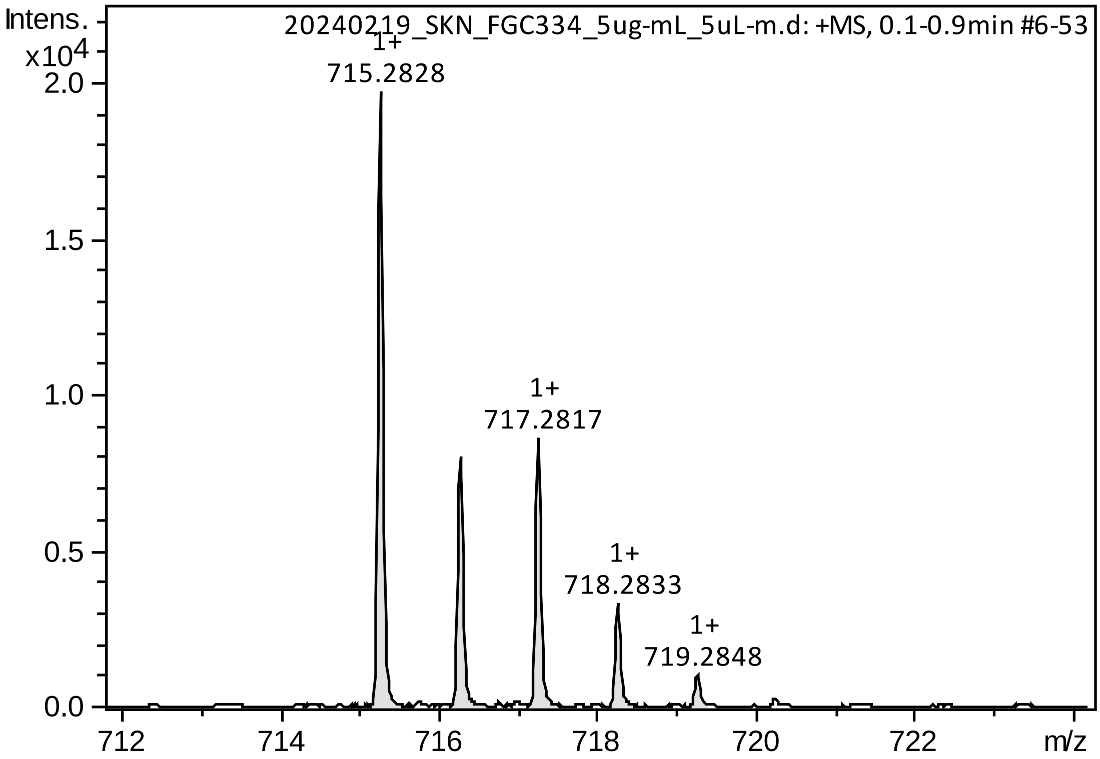

Intens.
x10^4
20240219_SKN_FGC334_5ug-mL_5uL-m.d: +MS, 0.1-0.9min #6-53
1+
715.2828
1+
717.2817
1+
718.2833
1+
719.2848
2.0
1.5
1.0
0.5
0.0
712
714
716
718
720
722
m/z

#### Cmpd 61

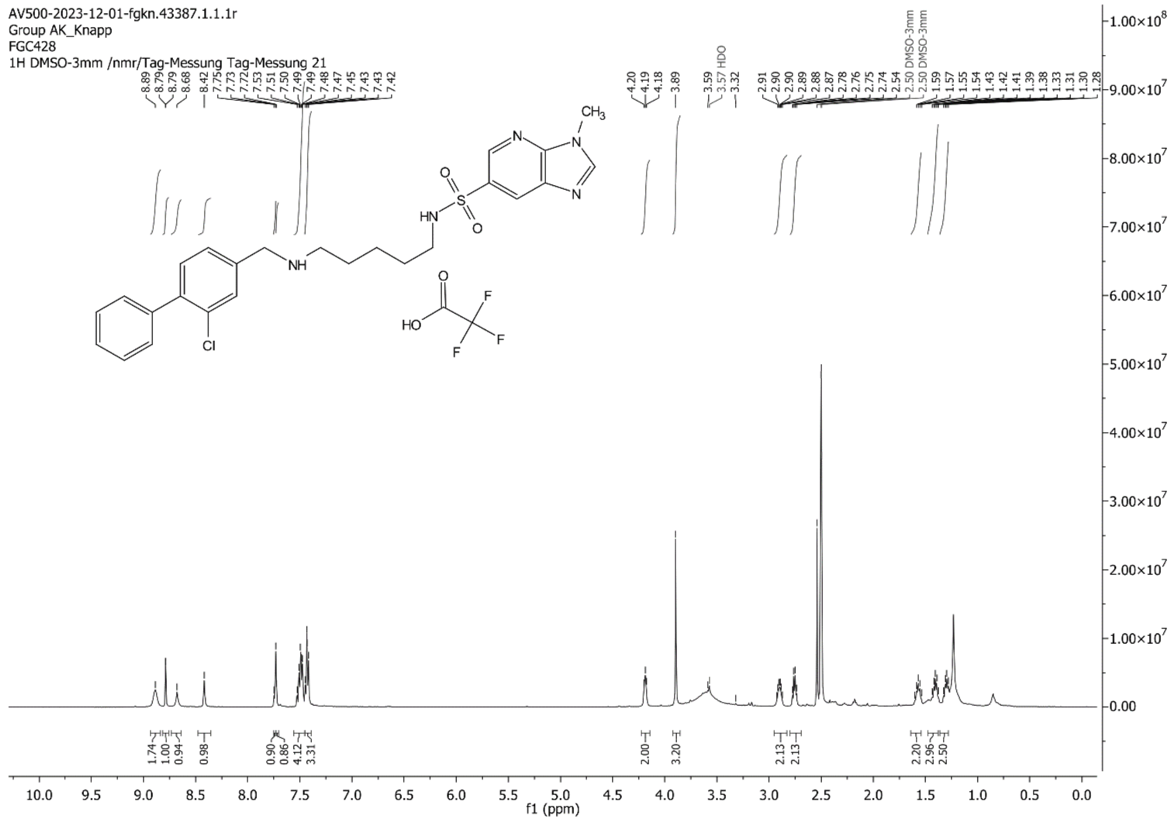
AV500-2023-12-01-fgkn.43387.1.1.1r
Group AK_Knapp
FGC428
1H DMSO-3mm /nmr/Tag-Messung Tag-Messung 21
f1 (ppm)

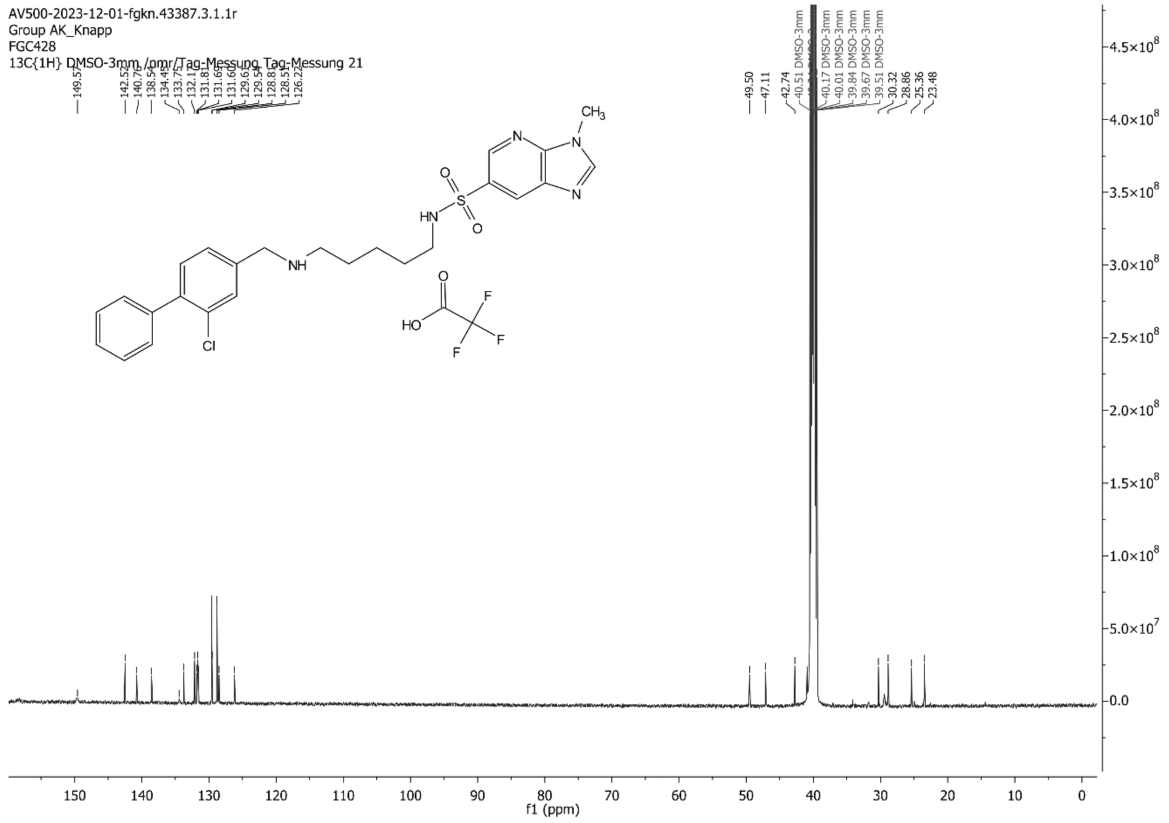
AV500-2023-12-01-fgkn.43387.3.1.1r
Group AK_Knapp
FGC428
13C{1H} DMSO-3mm /nmr/Tag-Messung Tag-Messung 21
f1 (ppm)

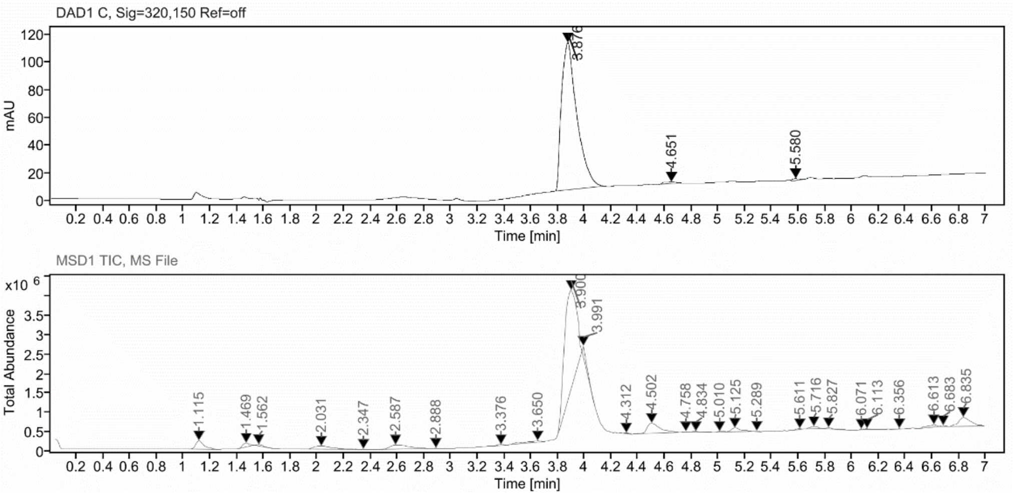

**Sample Purity**

Signal Description DAD1 C, Sig=320,150 Ref=off

| Sample Name | Name | RT | Width | Area | Area% | Height |
|---|---|---|---|---|---|---|
| FGC428_4 | | 3.876 | 0.124 | 842.1753 | 98.47 | 106.0214 |
| FGC428_4 | | 4.651 | 0.113 | 7.9924 | 0.93 | 1.2674 |
| FGC428_4 | | 5.580 | 0.109 | 5.1180 | 0.60 | 1.4536 |

Max Area% 98.467

UV Signal Purity>95% Pass

**UV Apex Spectra**

**RT:**

3.876

**Sample Name:**

FGC428_4

**Signal Name:**

DAD1C

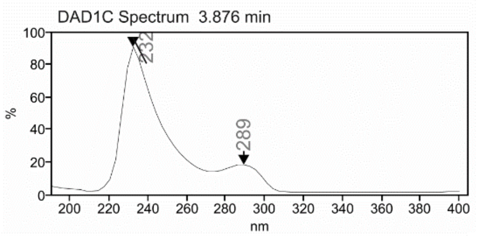

**RT:**

3.900

**Sample Name:**

FGC428_4

**Signal Name:**

MSD1TIC

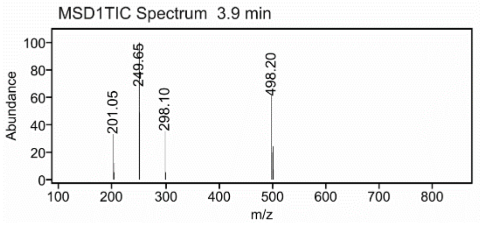

### Mass Spectrum SmartFormula Report

**Analysis Info**

| | | | |
|---|---|---|---|
| Analysis Name | D:\Data\MS_Service\Service_Kaiser\2024_Q1\20240229_SKN_FGC428_25x_5uL-m.d | Acquisition Date | 2/29/2024 3:46:49 PM |
| Method | DirectInfusion_WithFlow_Tune_pos.m | Operator | BDAL@DE |
| Sample Name | Test1 | Instrument | micrOTOF-Q 228888.10407 |
| Comment | | | |

**Acquisition Parameter**

| | | | | | |
|---|---|---|---|---|---|
| Source Type | ESI | Ion Polarity | Positive | Set Nebulizer | 1.8 Bar |
| Focus | Active | Set Capillary | 4000 V | Set Dry Heater | 320 °C |
| Scan Begin | 50 m/z | Set End Plate Offset | -500 V | Set Dry Gas | 8.0 l/min |
| Scan End | 1600 m/z | Set Collision Cell RF | 150.0 Vpp | Set Divert Valve | Waste |

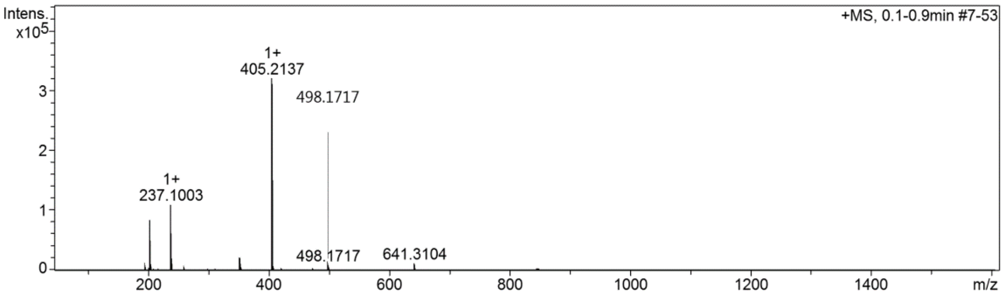

**Cmpd 63**

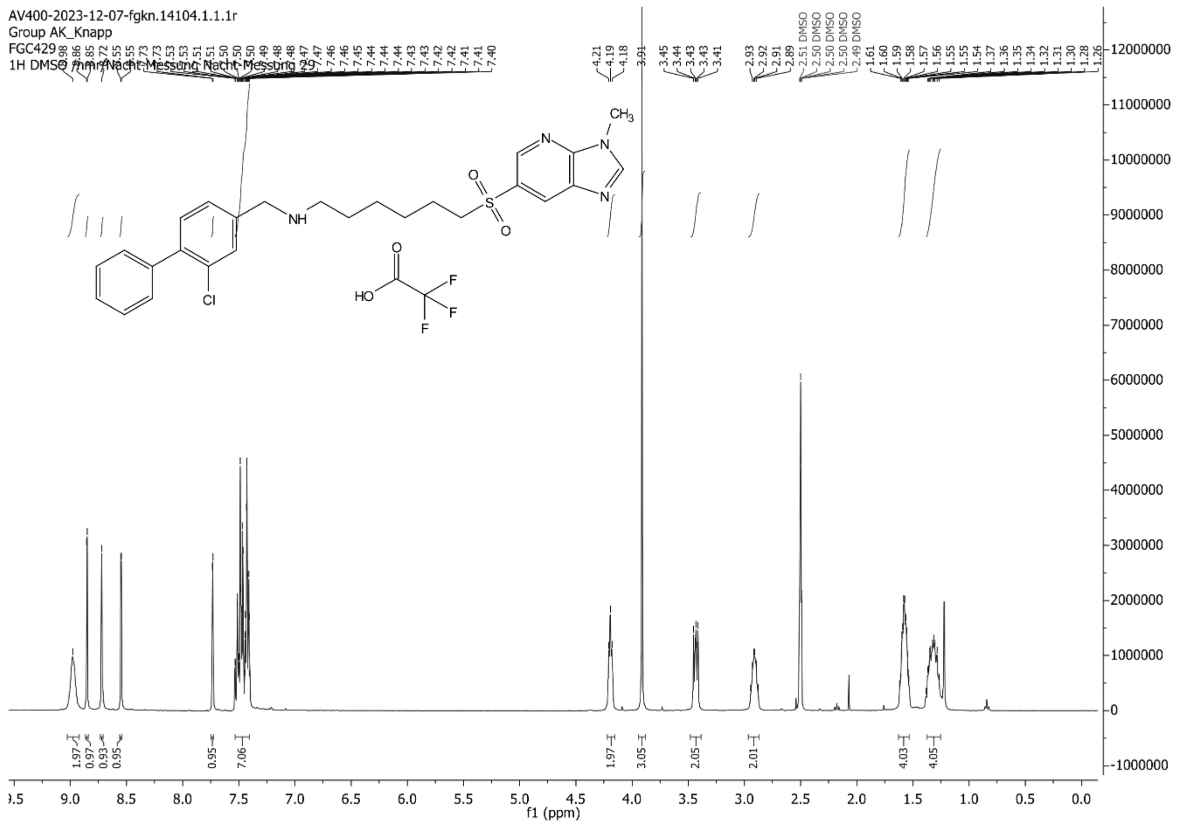

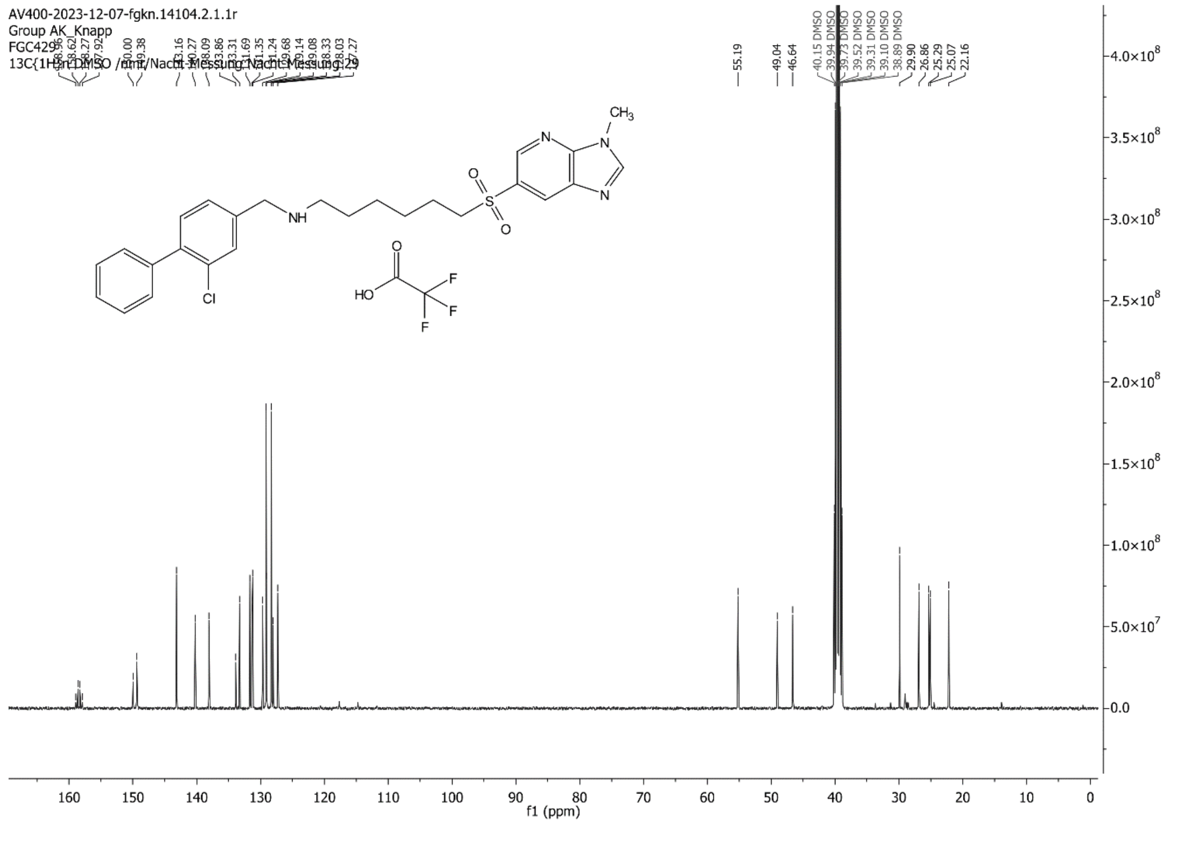

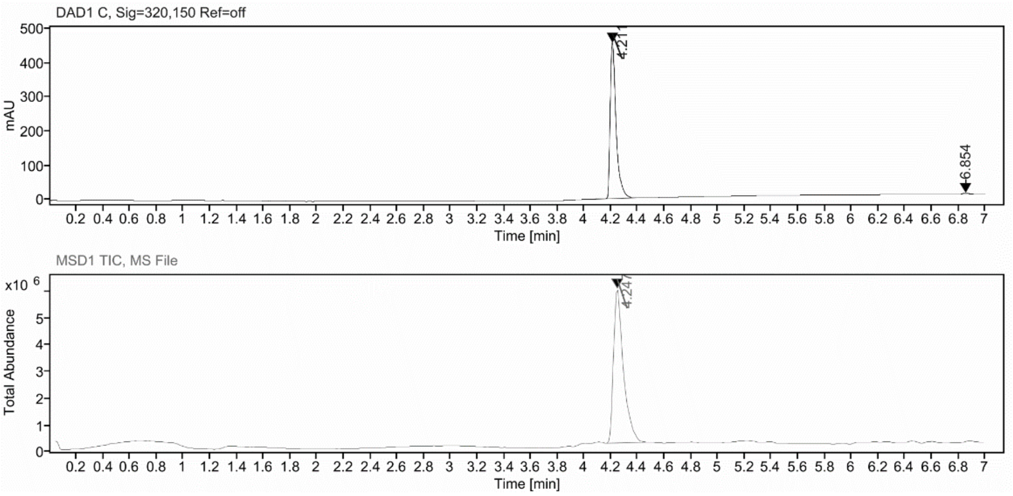

**Sample Purity**

Signal Description DAD1 C, Sig=320,150 Ref=off

| Sample Name | Name | RT | Width | Area | Area% | Height |
|---|---|---|---|---|---|---|
| FGC429_9+10 | | 4.211 | 0.045 | 1409.7264 | 98.98 | 456.5709 |
| FGC429_9+10 | | 6.854 | 0.048 | 14.5748 | 1.02 | 4.4909 |

Max Area% 98.977

UV Signal Purity>95% Pass

**UV Apex Spectra**

**RT:**

4.211

**Sample Name:**

FGC429_9+10

**Signal Name:**

DAD1C

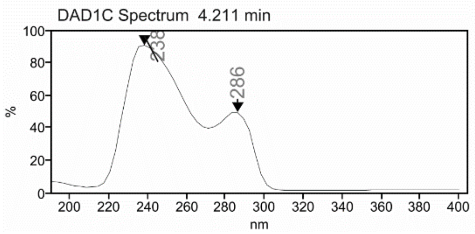

**RT:**

4.247

**Sample Name:**

FGC429_9+10

**Signal Name:**

MSD1TIC

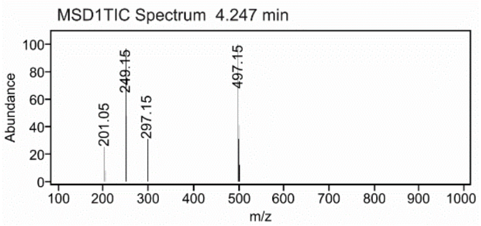

### Mass Spectrum SmartFormula Report

**Analysis Info**

| | | | |
|---|---|---|---|
| Analysis Name | D:\Data\MS_Service\Service_Kaiser\2024_Q1\20240228_SKN_FGC429_100x_5uL-m.d | Acquisition Date | 2/28/2024 1:57:45 PM |
| Method | DirectInfusion_WithFlow_Tune_pos.m | Operator | BDAL@DE |
| Sample Name | Test1 | Instrument | micrOTOF-Q 228888.10407 |
| Comment | | | |

**Acquisition Parameter**

| | | | | | |
|---|---|---|---|---|---|
| Source Type | ESI | Ion Polarity | Positive | Set Nebulizer | 1.8 Bar |
| Focus | Active | Set Capillary | 4000 V | Set Dry Heater | 320 °C |
| Scan Begin | 50 m/z | Set End Plate Offset | -500 V | Set Dry Gas | 8.0 l/min |
| Scan End | 1600 m/z | Set Collision Cell RF | 150.0 Vpp | Set Divert Valve | Waste |

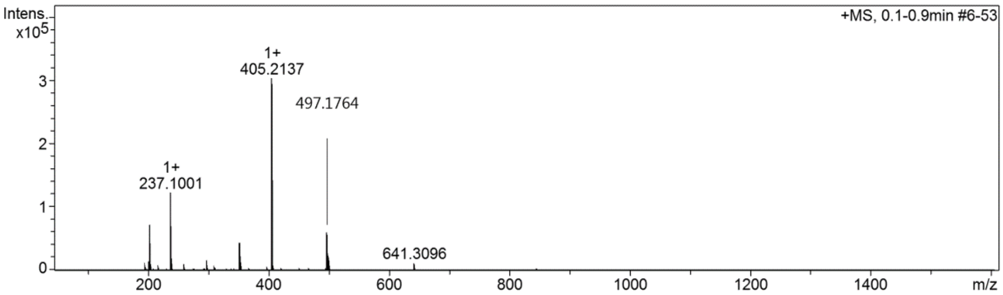

**Cmpd 66**

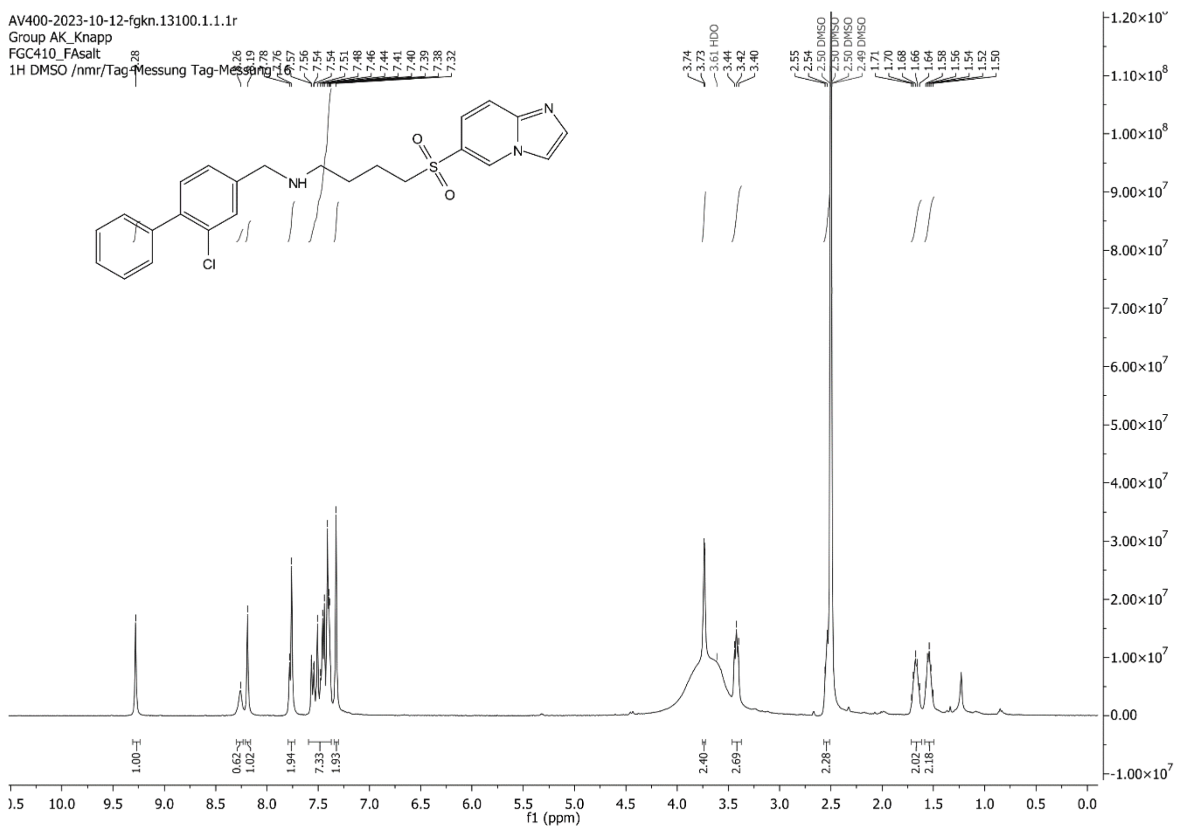
AV400-2023-10-12-fgkn.13100.1.1.1r
Group AK_Knapp
FGC410_FAsalt
1H DMSO /nmr/Tag-Messung Tag-Messung 16
f1 (ppm)

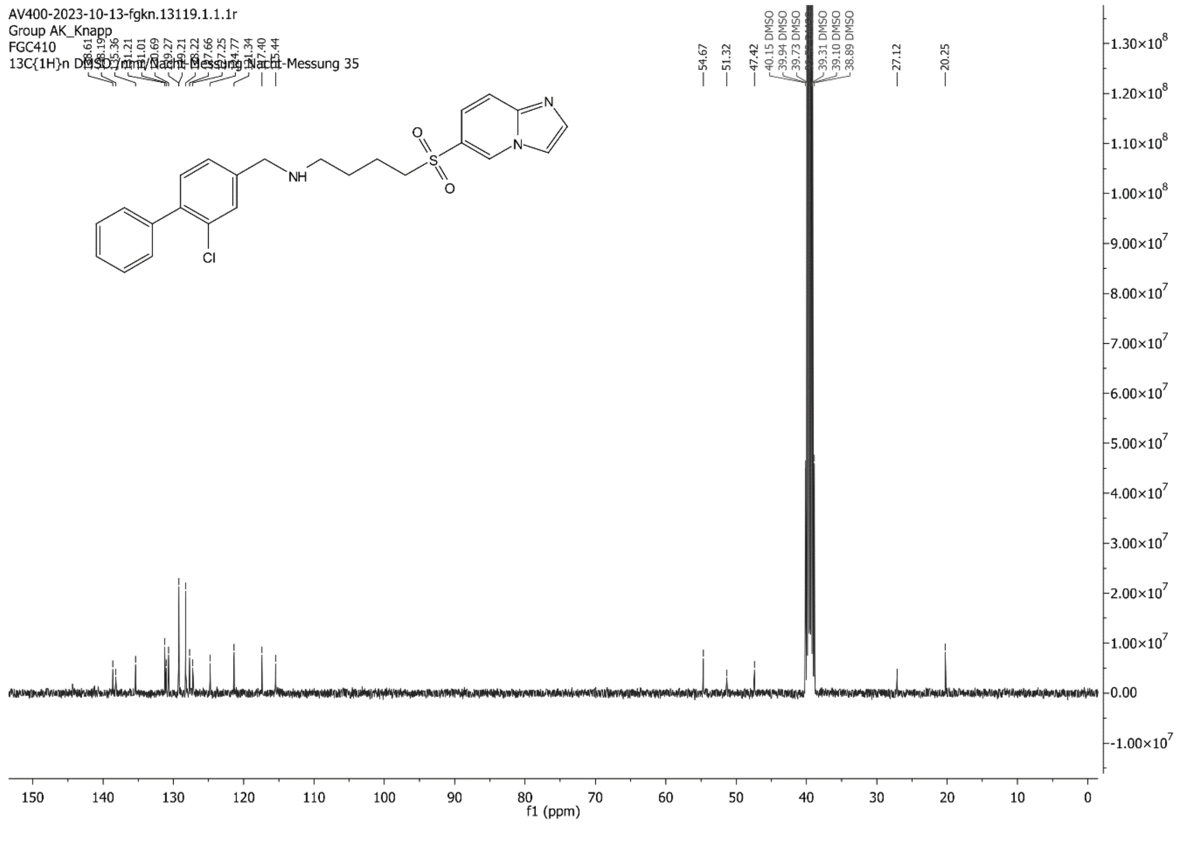
AV400-2023-10-13-fgkn.13119.1.1.1r
Group AK_Knapp
FGC410
13C{1H}n DMSO /nmr/Nacht-Messung Nacht-Messung 35
f1 (ppm)

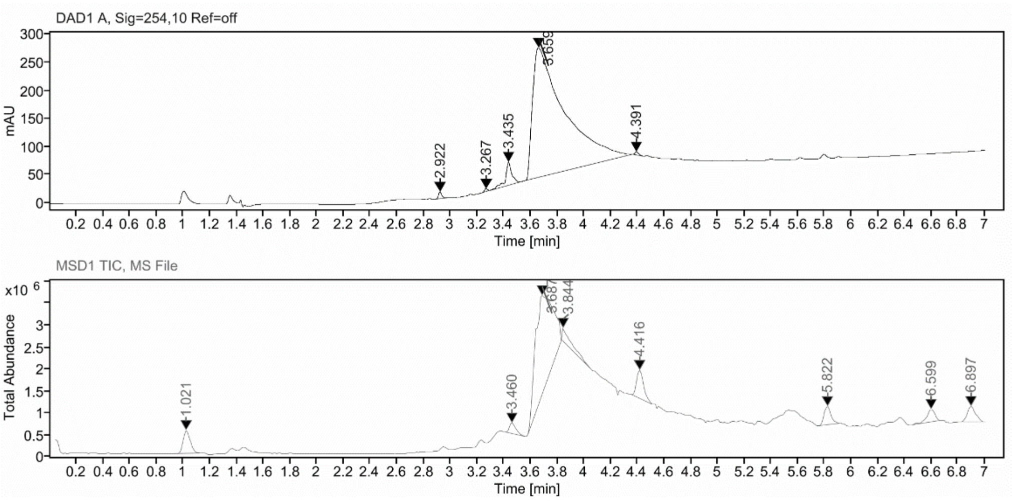

**Sample Purity**

| | |
|---|---|
| Signal Description | DAD1 A, Sig=254,10 Ref=off |

| Sample Name | Name | RT | Width | Area | Area% | Height |
|---|---|---|---|---|---|---|
| FGC410_RPmitFA_fr1 | | 2.922 | 0.024 | 20.2041 | 0.51 | 12.2740 |
| FGC410_RPmitFA_fr1 | | 3.267 | 0.028 | 7.1638 | 0.18 | 4.5115 |
| FGC410_RPmitFA_fr1 | | 3.435 | 0.040 | 130.1319 | 3.26 | 40.5206 |
| FGC410_RPmitFA_fr1 | | 3.659 | 0.246 | 3818.8171 | 95.76 | 230.5060 |
| FGC410_RPmitFA_fr1 | | 4.391 | 0.031 | 11.5576 | 0.29 | 5.5687 |

| | |
|---|---|
| Max Area% | 95.761 |
| UV Signal Purity>95% | Pass |

**UV Apex Spectra**

**RT:**

3.659

**Sample Name:**

FGC410_RPmitFA_fr1

**Signal Name:**

DAD1A

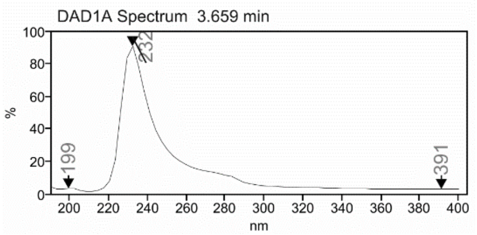

**RT:**

3.687

**Sample Name:**

FGC410_RPmitFA_fr1

**Signal Name:**

MSD1TIC

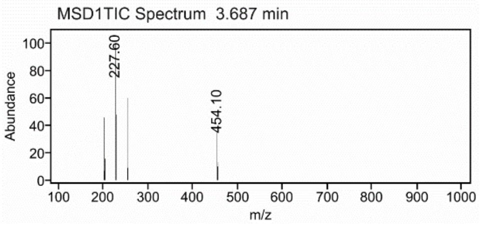

### Mass Spectrum SmartFormula Report

**Analysis Info**

| | | | |
|---|---|---|---|
| Analysis Name | D:\Data\MS_Service\Service_Kaiser\2024_Q1\20240228_SKN_FGC410_100x_5uL-m.d | Acquisition Date | 2/28/2024 1:31:13 PM |
| Method | DirectInfusion_WithFlow_Tune_pos.m | Operator | BDAL@DE |
| Sample Name | Test1 | Instrument | micrOTOF-Q 228888.10407 |
| Comment | | | |

**Acquisition Parameter**

| | | | | | |
|---|---|---|---|---|---|
| Source Type | ESI | Ion Polarity | Positive | Set Nebulizer | 1.8 Bar |
| Focus | Active | Set Capillary | 4000 V | Set Dry Heater | 320 °C |
| Scan Begin | 50 m/z | Set End Plate Offset | -500 V | Set Dry Gas | 8.0 l/min |
| Scan End | 1600 m/z | Set Collision Cell RF | 150.0 Vpp | Set Divert Valve | Waste |

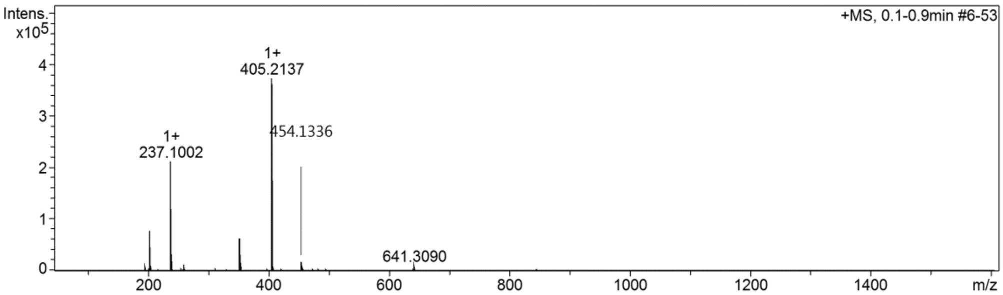

**Cmpd 72**

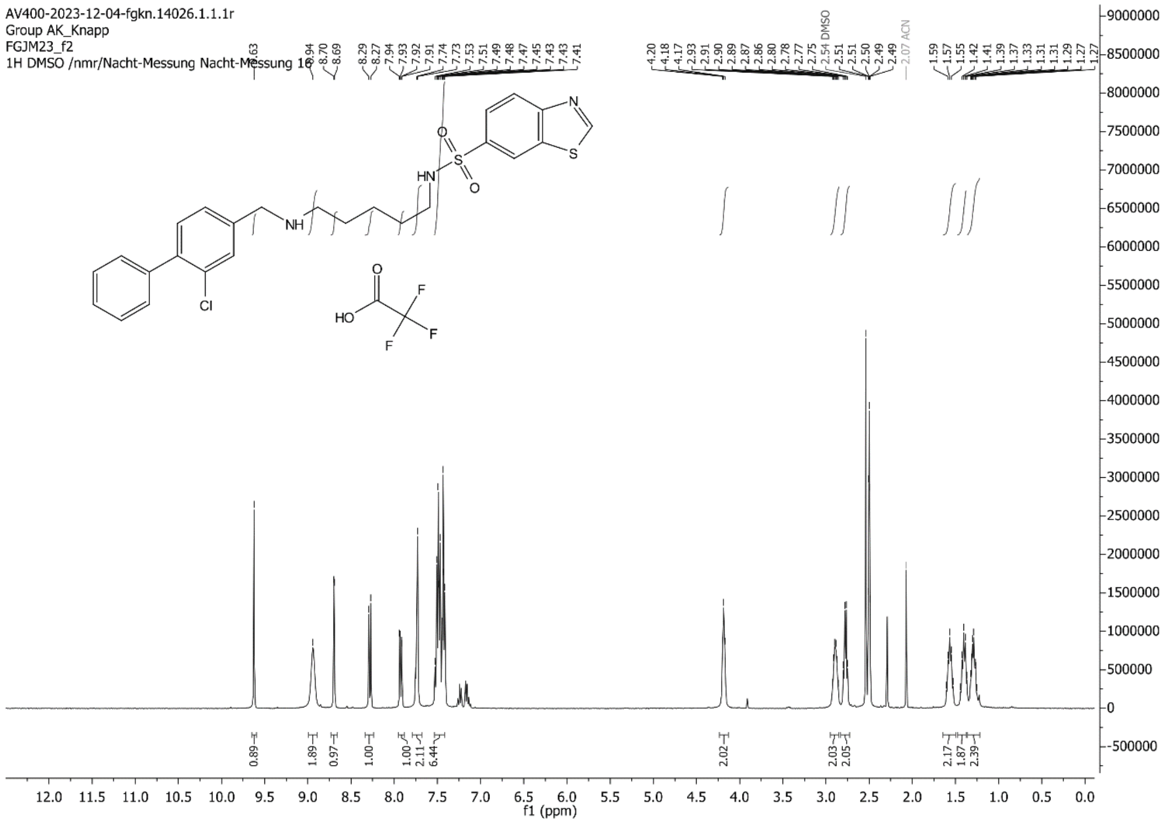
AV400-2023-12-04-fgkn.14026.1.1.1r
Group AK_Knapp
FGJM23_f2
1H DMSO /nmr/Nacht-Messung Nacht-Messung 16
f1 (ppm)

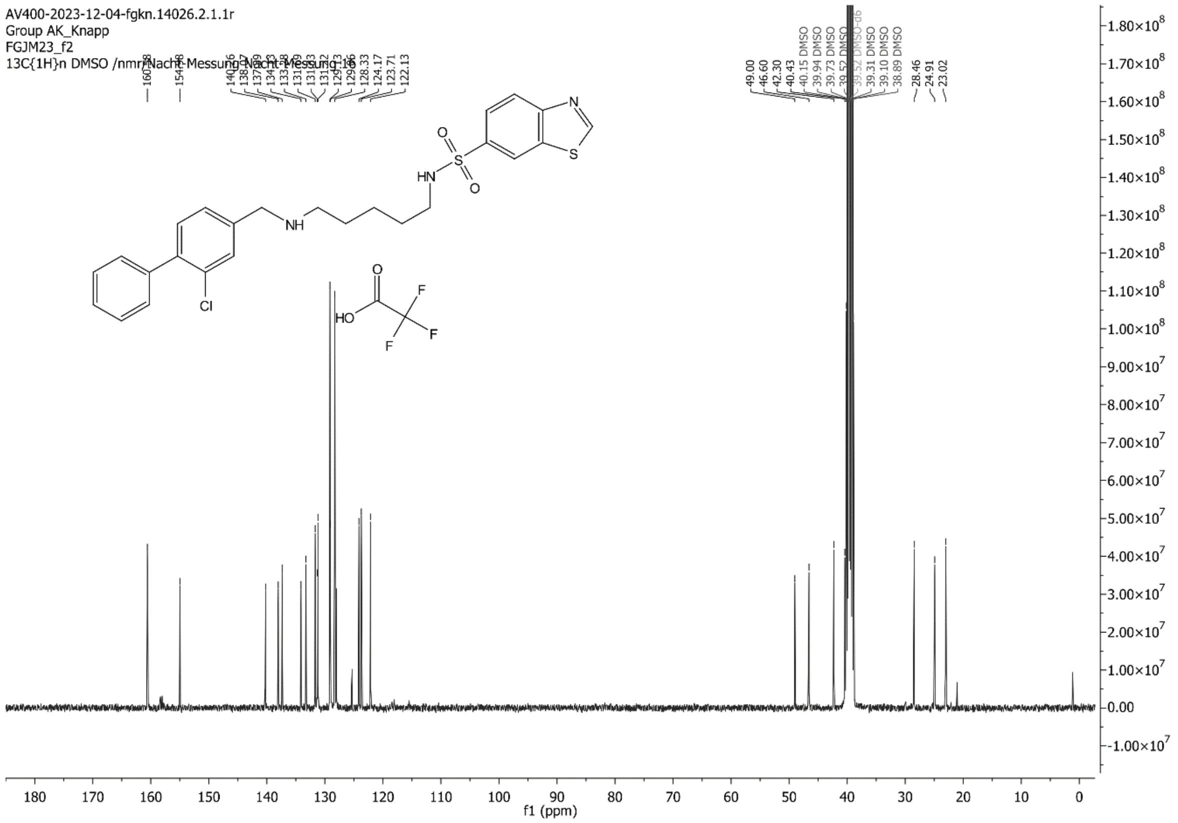
AV400-2023-12-04-fgkn.14026.2.1.1r
Group AK_Knapp
FGJM23_f2
13C{1H}n DMSO /nmr/Nacht-Messung Nacht-Messung 16
f1 (ppm)

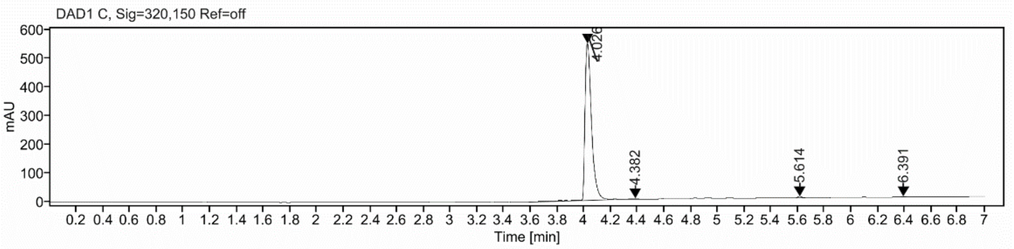

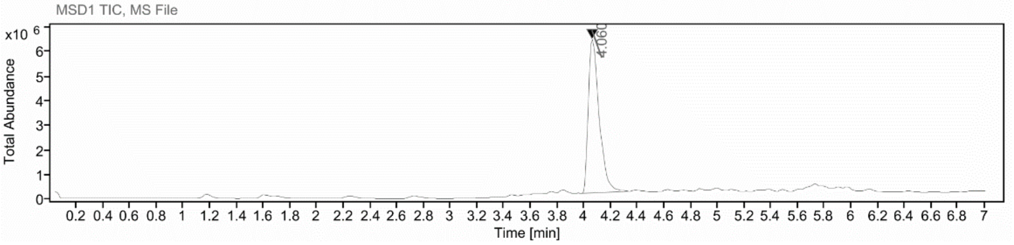

**Sample Purity**

Signal Description DAD1 C, Sig=320,150 Ref=off

| Sample Name | Name | RT | Width | Area | Area% | Height |
|---|---|---|---|---|---|---|
| FGJM23_8-12 | | 4.026 | 0.051 | 1855.5946 | 98.36 | 543.5182 |
| FGJM23_8-12 | | 4.382 | 0.043 | 9.4532 | 0.50 | 2.6585 |
| FGJM23_8-12 | | 5.614 | 0.038 | 13.1885 | 0.70 | 4.9748 |
| FGJM23_8-12 | | 6.391 | 0.043 | 8.3430 | 0.44 | 2.4638 |

Max Area% 98.358

UV Signal Purity>95% Pass

**UV Apex Spectra**

**RT:**

4.026

**Sample Name:**

FGJM23_8-12

**Signal Name:**

DAD1C

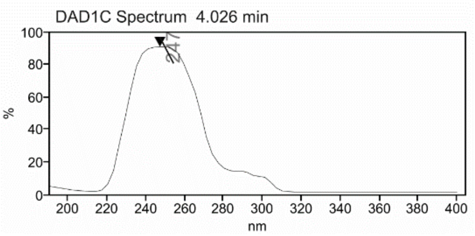

**RT:**

4.060

**Sample Name:**

FGJM23_8-12

**Signal Name:**

MSD1TIC

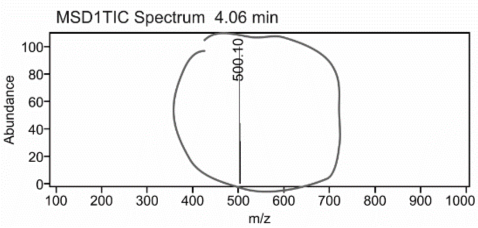

### Mass Spectrum SmartFormula Report

**Analysis Info**

| | | | |
|---|---|---|---|
| | | Acquisition Date | 2/28/2024 2:20:26 PM |
| Analysis Name | D:\Data\MS_Service\Service_Kaiser\2024_Q1\20240228_SKN_FGJM23_100x_5uL-m.d | | |
| Method | DirectInfusion_WithFlow_Tune_pos.m | Operator | BDAL@DE |
| Sample Name | Test1 | Instrument | micrOTOF-Q 228888.10407 |
| Comment | | | |

**Acquisition Parameter**

| | | | | | |
|---|---|---|---|---|---|
| Source Type | ESI | Ion Polarity | Positive | Set Nebulizer | 1.8 Bar |
| Focus | Active | Set Capillary | 4000 V | Set Dry Heater | 320 °C |
| Scan Begin | 50 m/z | Set End Plate Offset | -500 V | Set Dry Gas | 8.0 l/min |
| Scan End | 1600 m/z | Set Collision Cell RF | 150.0 Vpp | Set Divert Valve | Waste |

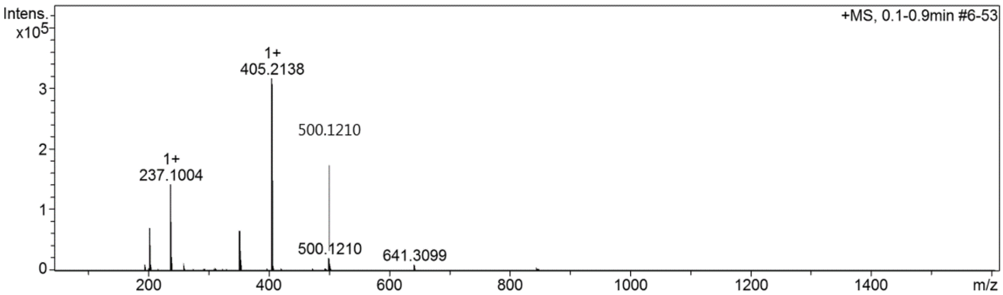

**Cmpd 73**

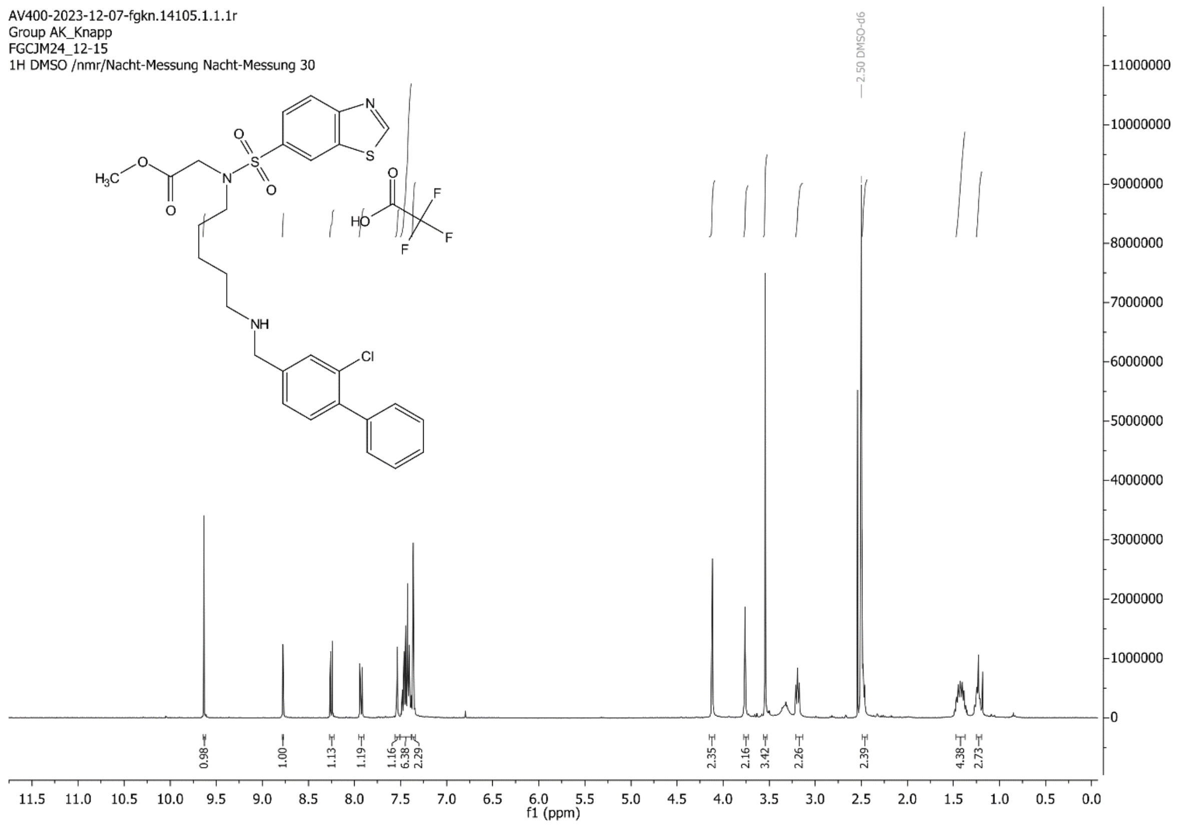

AV400-2023-12-07-fgkn.14105.1.1.1r
Group AK_Knapp
FGCJM24_12-15
1H DMSO /nmr/Nacht-Messung Nacht-Messung 30
f1 (ppm)

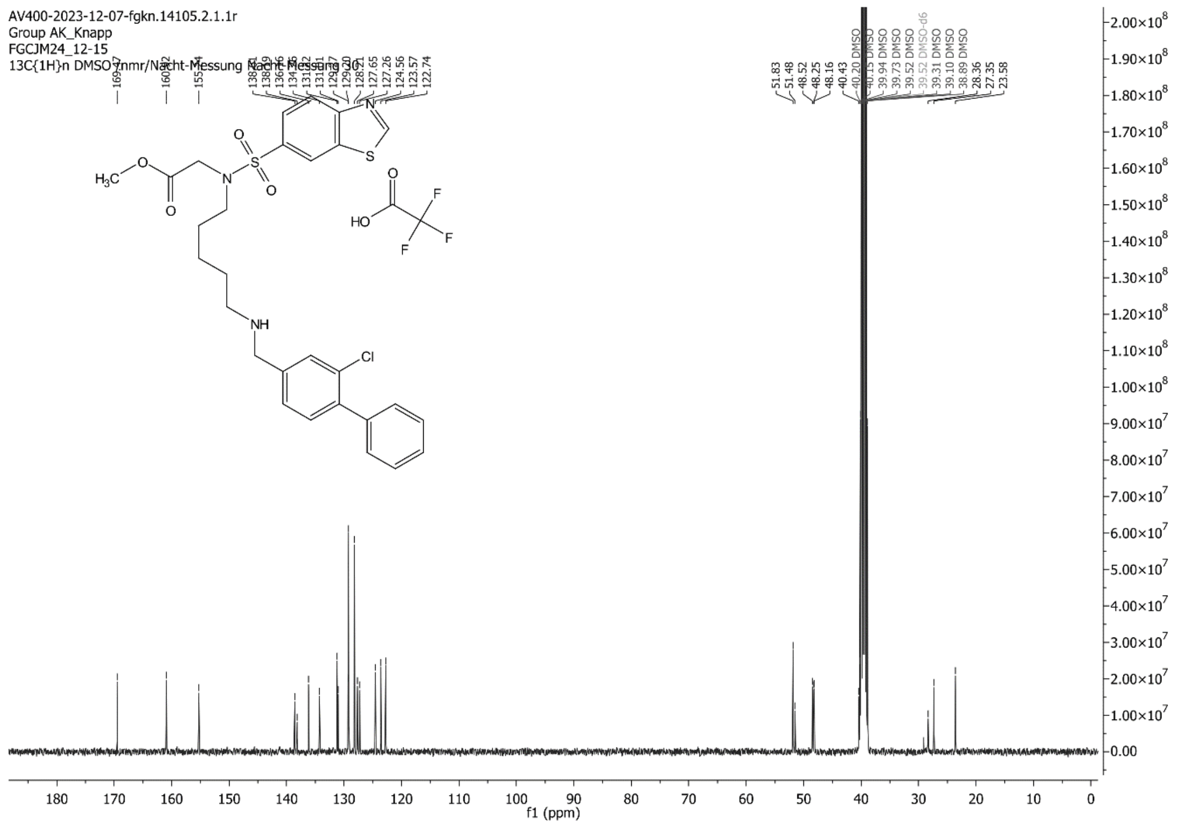

AV400-2023-12-07-fgkn.14105.2.1.1r
Group AK_Knapp
FGCJM24_12-15
13C{1H}n DMSO /nmr/Nacht-Messung Nacht-Messung 30
f1 (ppm)

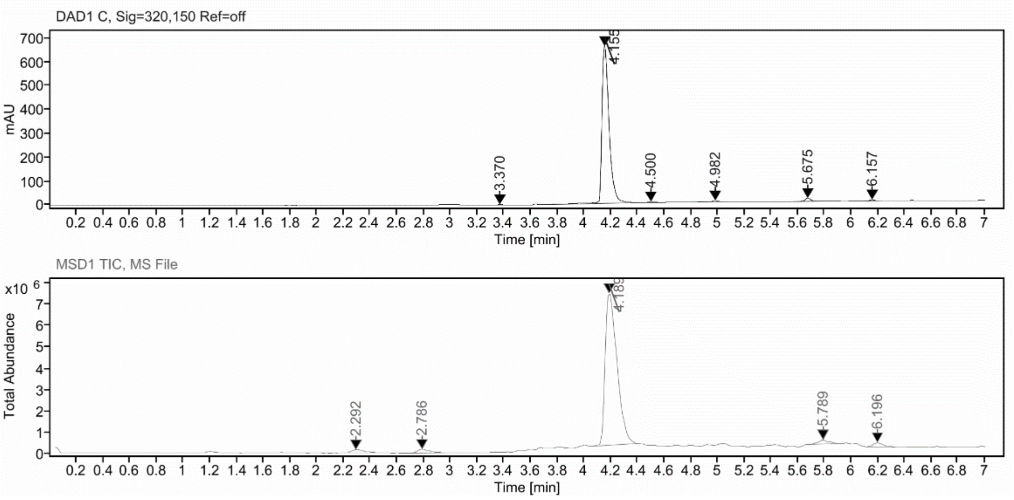

**Sample Purity**

Signal Description DAD1 C, Sig=320,150 Ref=off

| Sample Name | Name | RT | Width | Area | Area% | Height |
|---|---|---|---|---|---|---|
| FGCJM24_12_15_stock_urgent | | 3.370 | 0.034 | 6.8194 | 0.27 | 2.7176 |
| FGCJM24_12_15_stock_urgent | | 4.155 | 0.056 | 2456.9043 | 96.31 | 657.7203 |
| FGCJM24_12_15_stock_urgent | | 4.500 | 0.042 | 9.5445 | 0.37 | 3.6404 |
| FGCJM24_12_15_stock_urgent | | 4.982 | 0.050 | 23.7622 | 0.93 | 5.0894 |
| FGCJM24_12_15_stock_urgent | | 5.675 | 0.044 | 40.2743 | 1.58 | 13.6254 |
| FGCJM24_12_15_stock_urgent | | 6.157 | 0.043 | 13.8305 | 0.54 | 4.7288 |

Max Area% 96.306

UV Signal Purity>95% Pass

**UV Apex Spectra**

**RT:**

4.155

**Sample Name:**

FGCJM24_12_15_stock_urgent

**Signal Name:**

DAD1C

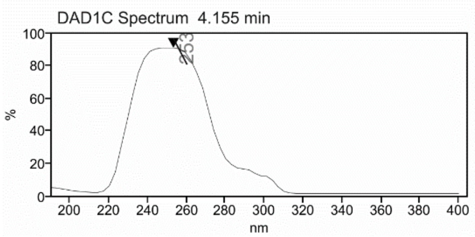

**RT:**

4.189

**Sample Name:**

FGCJM24_12_15_stock_urgent

**Signal Name:**

MSD1TIC

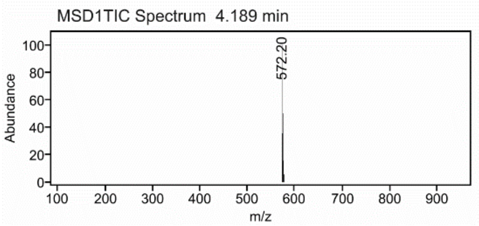

### Mass Spectrum SmartFormula Report

**Analysis Info**

| | | | |
|---|---|---|---|
| | | Acquisition Date | 2/28/2024 2:26:51 PM |
| Analysis Name | D:\Data\MS_Service\Service_Kaiser\2024_Q1\20240228_SKN_FGJM24_100x_5uL-m.d | | |
| Method | DirectInfusion_WithFlow_Tune_pos.m | Operator | BDAL@DE |
| Sample Name | Test1 | Instrument | micrOTOF-Q 228888.10407 |
| Comment | | | |

**Acquisition Parameter**

| | | | | | |
|---|---|---|---|---|---|
| Source Type | ESI | Ion Polarity | Positive | Set Nebulizer | 1.8 Bar |
| Focus | Active | Set Capillary | 4000 V | Set Dry Heater | 320 °C |
| Scan Begin | 50 m/z | Set End Plate Offset | -500 V | Set Dry Gas | 8.0 l/min |
| Scan End | 1600 m/z | Set Collision Cell RF | 150.0 Vpp | Set Divert Valve | Waste |

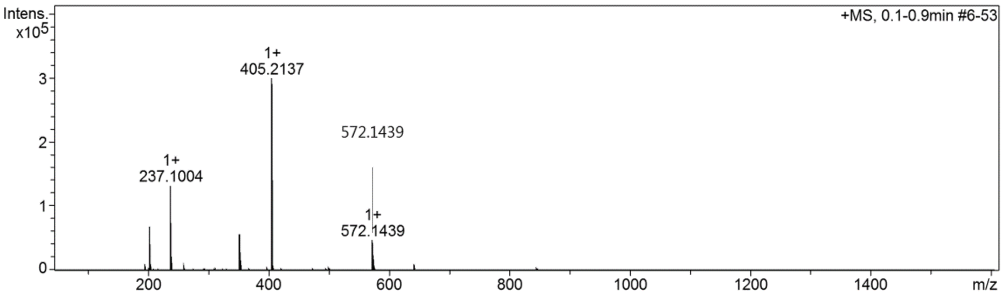

#### Cmpd 14 (ProCAM4066)[14]

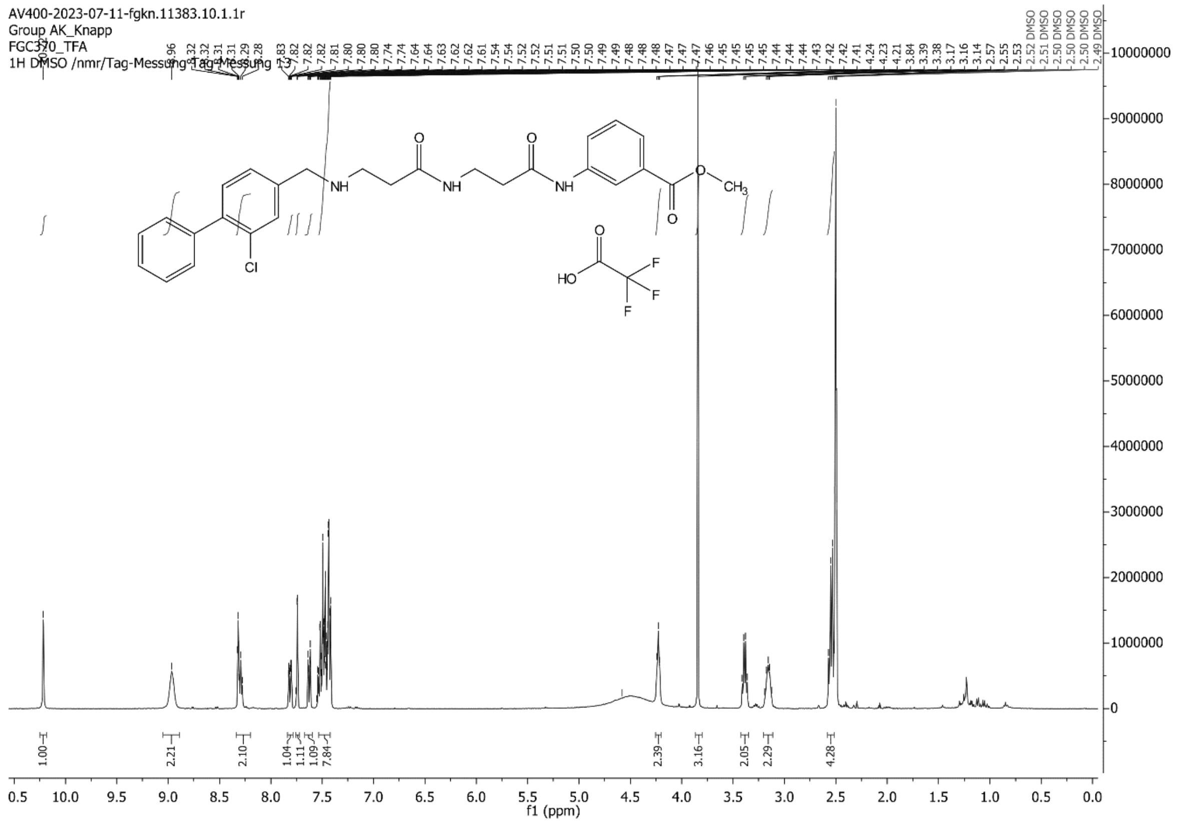

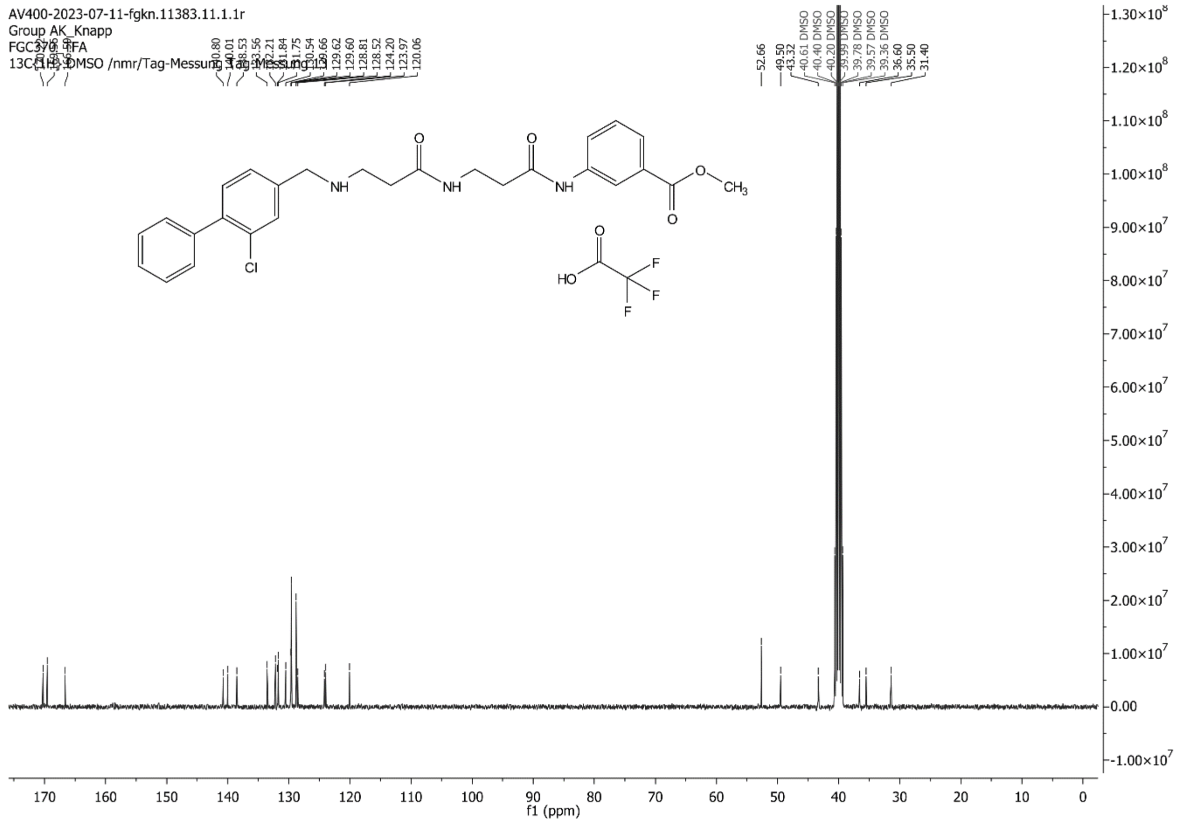

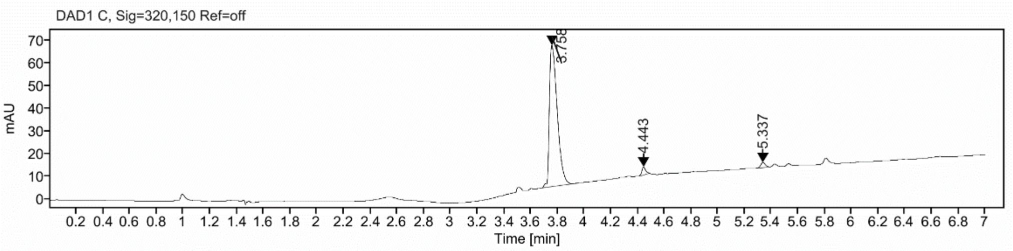

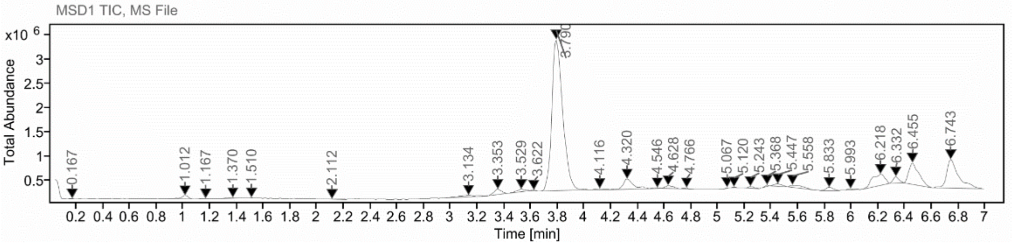

**Sample Purity**

Signal Description DAD1 C, Sig=320,150 Ref=off

| Sample Name | Name | RT | Width | Area | Area% | Height |
|---|---|---|---|---|---|---|
| FGC370 | | 3.758 | 0.065 | 256.5222 | 95.44 | 62.4564 |
| FGC370 | | 4.443 | 0.032 | 6.9990 | 2.60 | 3.5640 |
| FGC370 | | 5.337 | 0.037 | 5.2603 | 1.96 | 2.2621 |

Max Area% 95.439

UV Signal Purity>95% Pass

**UV Apex Spectra**

**RT:**

3.758

**Sample Name:**

FGC370

**Signal Name:**

DAD1C

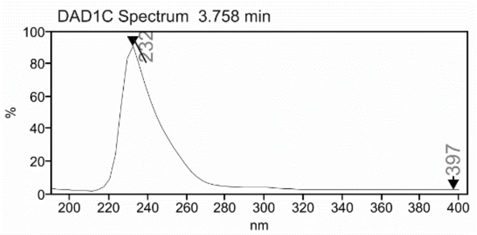

**RT:**

3.790

**Sample Name:**

FGC370

**Signal Name:**

MSD1TIC

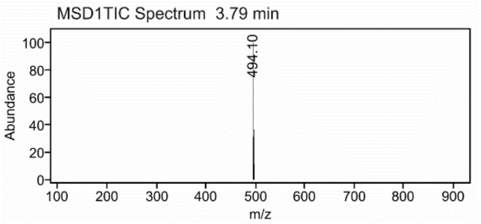

#### Cmpd 15 (CAM4066)[14]

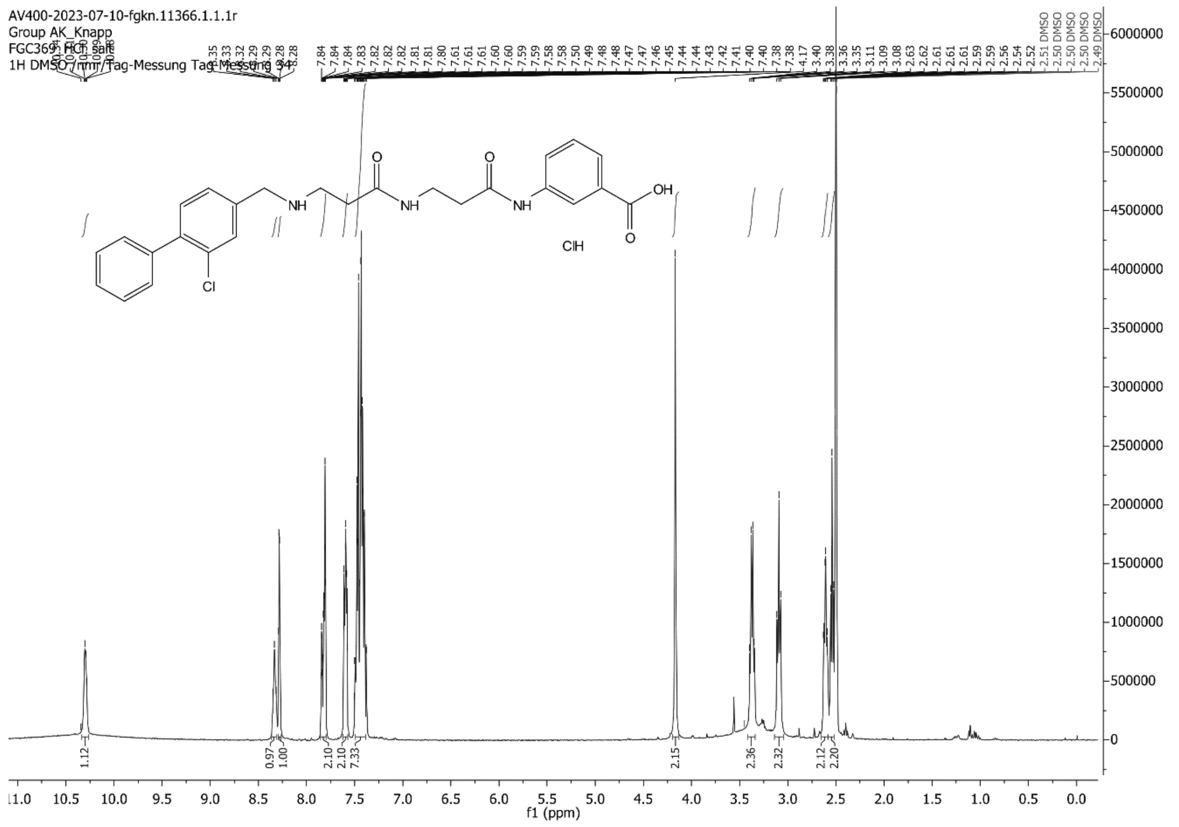

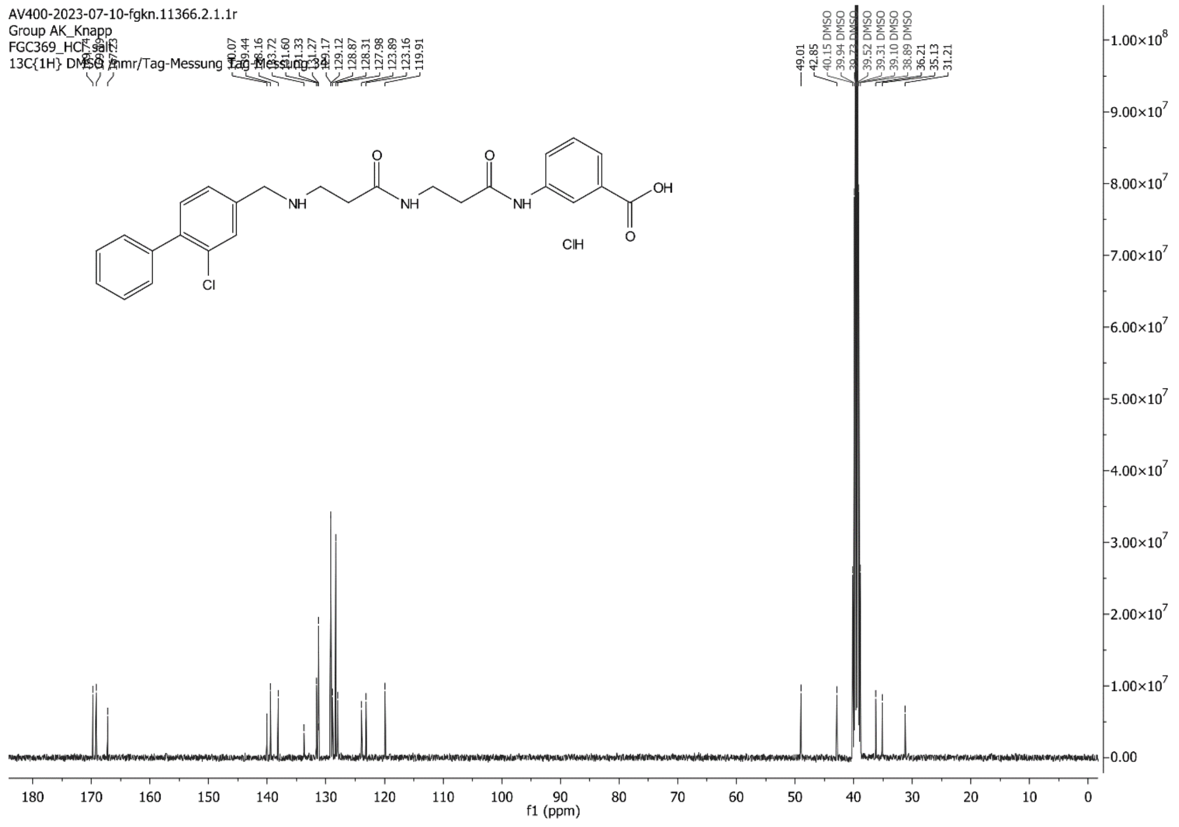

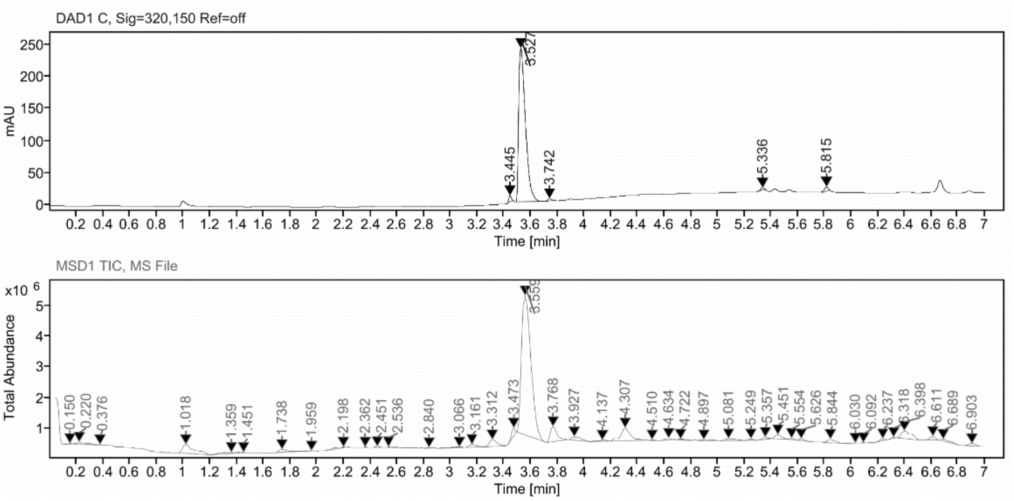

##### Sample Purity

Signal Description DAD1 C, Sig=320,150 Ref=off

| Sample Name | Name | RT | Width | Area | Area% | Height |
|---|---|---|---|---|---|---|
| FGC369_filtr | | 3.445 | 0.022 | 11.7966 | 1.36 | 9.6005 |
| FGC369_filtr | | 3.527 | 0.054 | 831.8133 | 95.72 | 238.9734 |
| FGC369_filtr | | 3.742 | 0.031 | 5.7374 | 0.66 | 3.2036 |
| FGC369_filtr | | 5.336 | 0.026 | 6.1774 | 0.71 | 4.0240 |
| FGC369_filtr | | 5.815 | 0.033 | 13.4562 | 1.55 | 6.7368 |

Max Area% 95.723

UV Signal Purity>95% Pass

##### UV Apex Spectra

**RT:**

3.527

**Sample Name:**

FGC369_filtr

**Signal Name:**

DAD1C

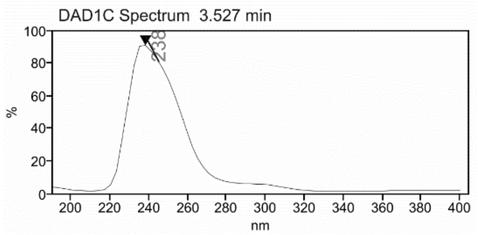

**RT:**

3.559

**Sample Name:**

FGC369_filtr

**Signal Name:**

MSD1TIC

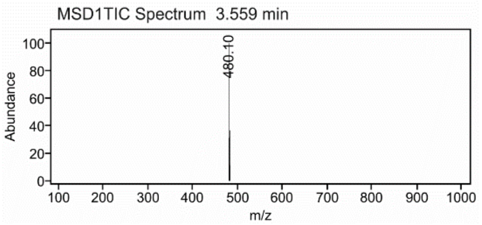

**S1**[14]

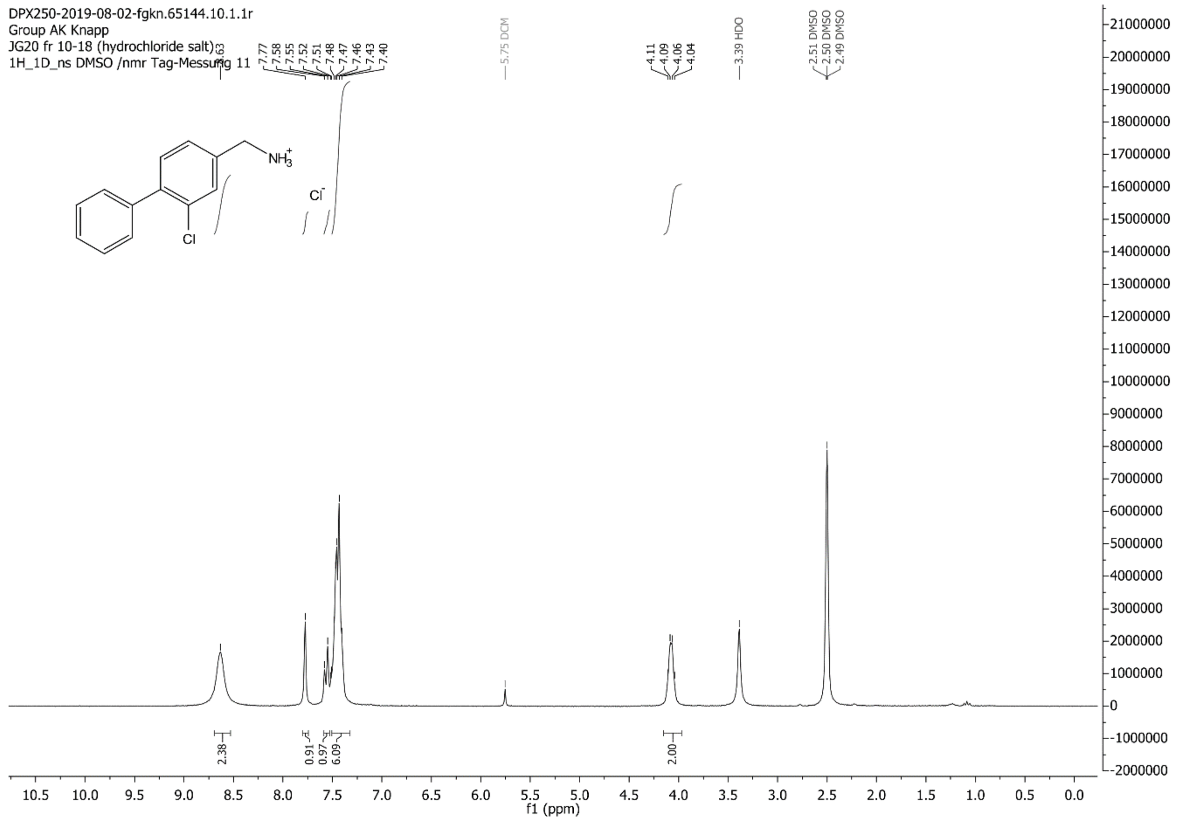

DPX250-2019-08-02-fgkn.65144.10.1.1r
Group AK Knapp
JG20 fr 10-18 (hydrochloride salt)
1H_1D_ns DMSO /nmr Tag-Messung 11
8.63
7.77
7.58
7.55
7.52
7.51
7.48
7.47
7.46
7.43
7.40
5.75 DCM
4.11
4.09
4.06
4.04
3.39 HDO
2.51 DMSO
2.50 DMSO
2.49 DMSO
NH3+
Cl-
Cl
2.38
0.91
0.97
6.09
2.00
f1 (ppm)

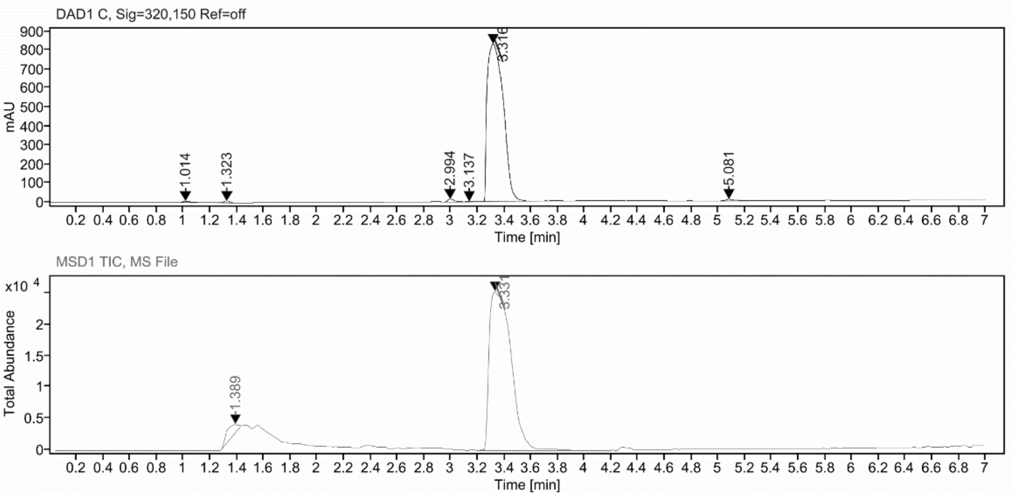

**Sample Purity**

Signal Description DAD1 C, Sig=320,150 Ref=off

| Sample Name | Name | RT | Width | Area | Area% | Height |
|---|---|---|---|---|---|---|
| FGJG20_control | | 1.014 | 0.078 | 34.5318 | 0.49 | 6.6436 |
| FGJG20_control | | 1.323 | 0.047 | 29.1153 | 0.41 | 9.2920 |
| FGJG20_control | | 2.994 | 0.050 | 46.1166 | 0.66 | 14.0670 |
| FGJG20_control | | 3.137 | 0.053 | 6.4471 | 0.09 | 2.1860 |
| FGJG20_control | | 3.316 | 0.143 | 6862.6118 | 97.79 | 828.9218 |
| FGJG20_control | | 5.081 | 0.069 | 39.0219 | 0.56 | 8.0325 |

Max Area% 97.788

UV Signal Purity>95% Pass

**UV Apex Spectra**

**RT:**

3.316

**Sample Name:**

FGJG20_control

**Signal Name:**

DAD1C

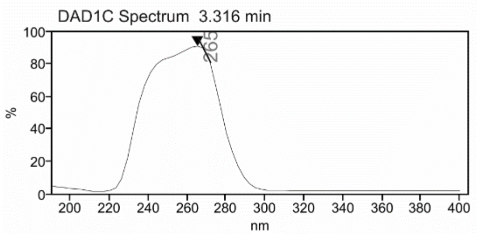

**RT:**

3.331

**Sample Name:**

FGJG20_control

**Signal Name:**

MSD1TIC

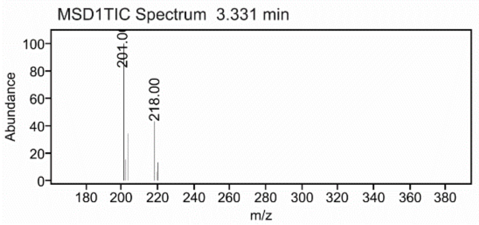

**Cmpd S1**

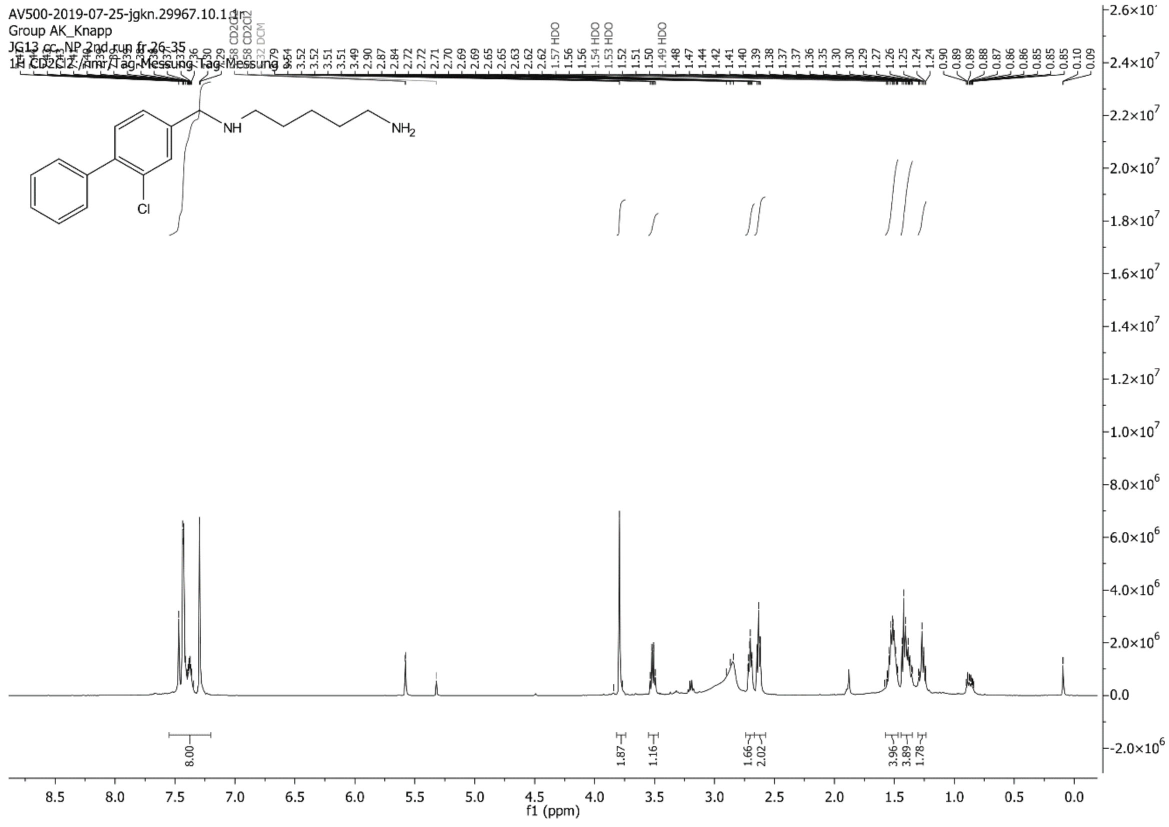
AV500-2019-07-25-jgkn.29967.10.1.1r
Group AK_Knapp
JG13 cc. NP 2nd run fr 26-35
1H CD2Cl2 /Amr/Tag-Messung Tag-Messung 3
NH
NH2
Cl
f1 (ppm)

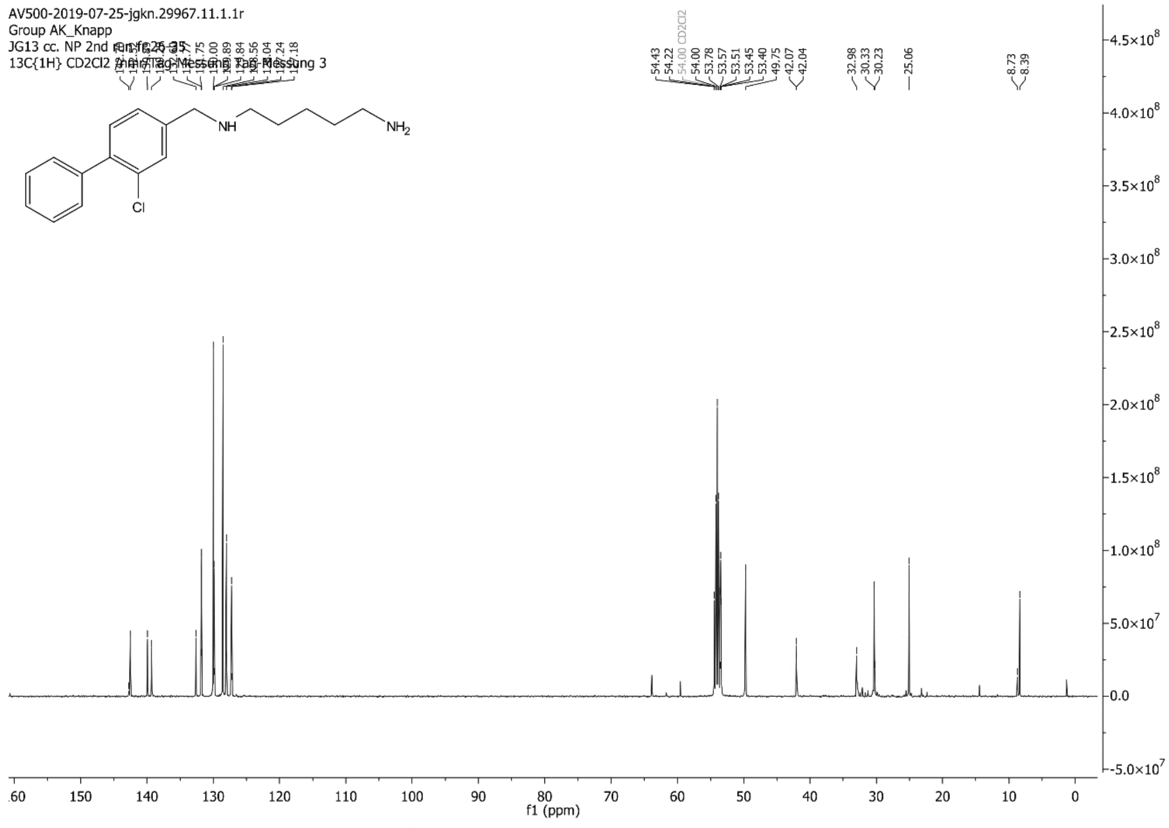
AV500-2019-07-25-jgkn.29967.11.1.1r
Group AK_Knapp
JG13 cc. NP 2nd run fr 26-35
13C{1H} CD2Cl2 /Amr/Tag-Messung Tag-Messung 3
NH
NH2
Cl
f1 (ppm)

C:\Xcalibur\data\JG13-1 7/18/2019 12:25:53 PM

JG13-1 #33-44 RT: 0.55-0.75 AV: 12 SB: 16 0.05-0.31 NL: 2.19E4
T: {0,0} + c ESI !corona sid=75.00 det=1306.00 Full ms [105.00-600.00]

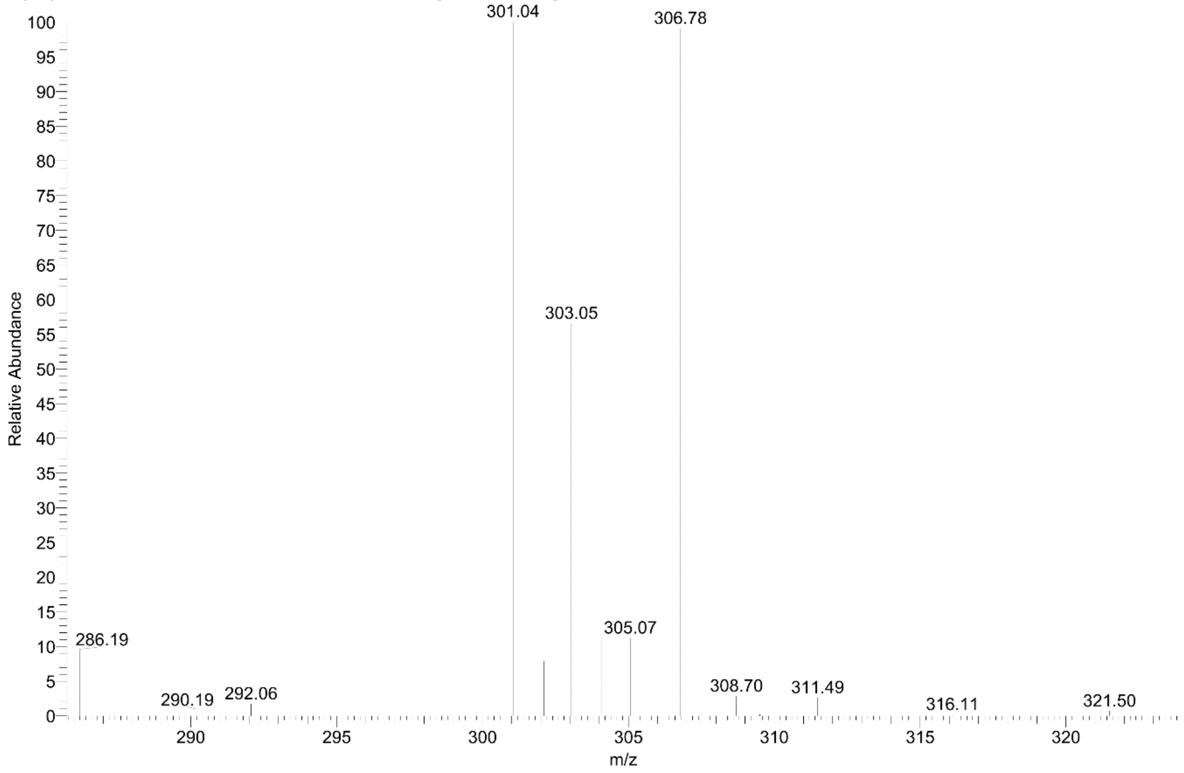

#### Cmpd S3

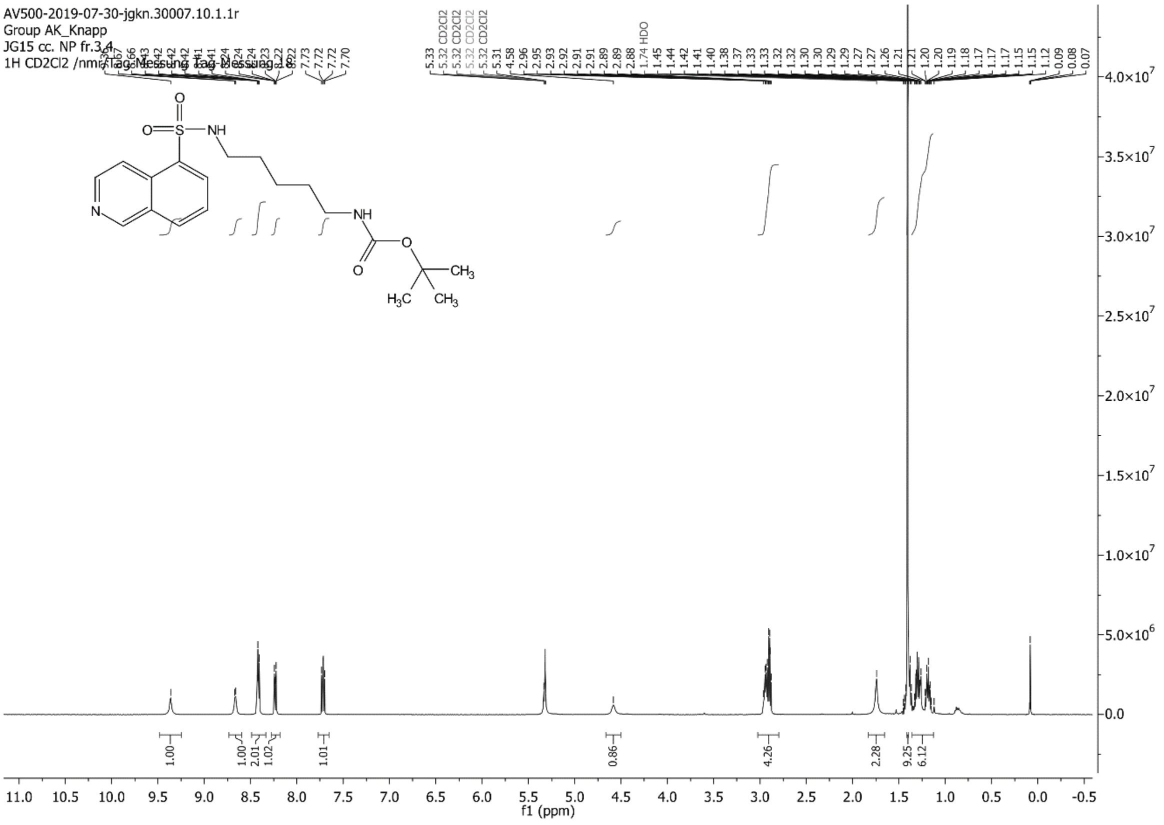

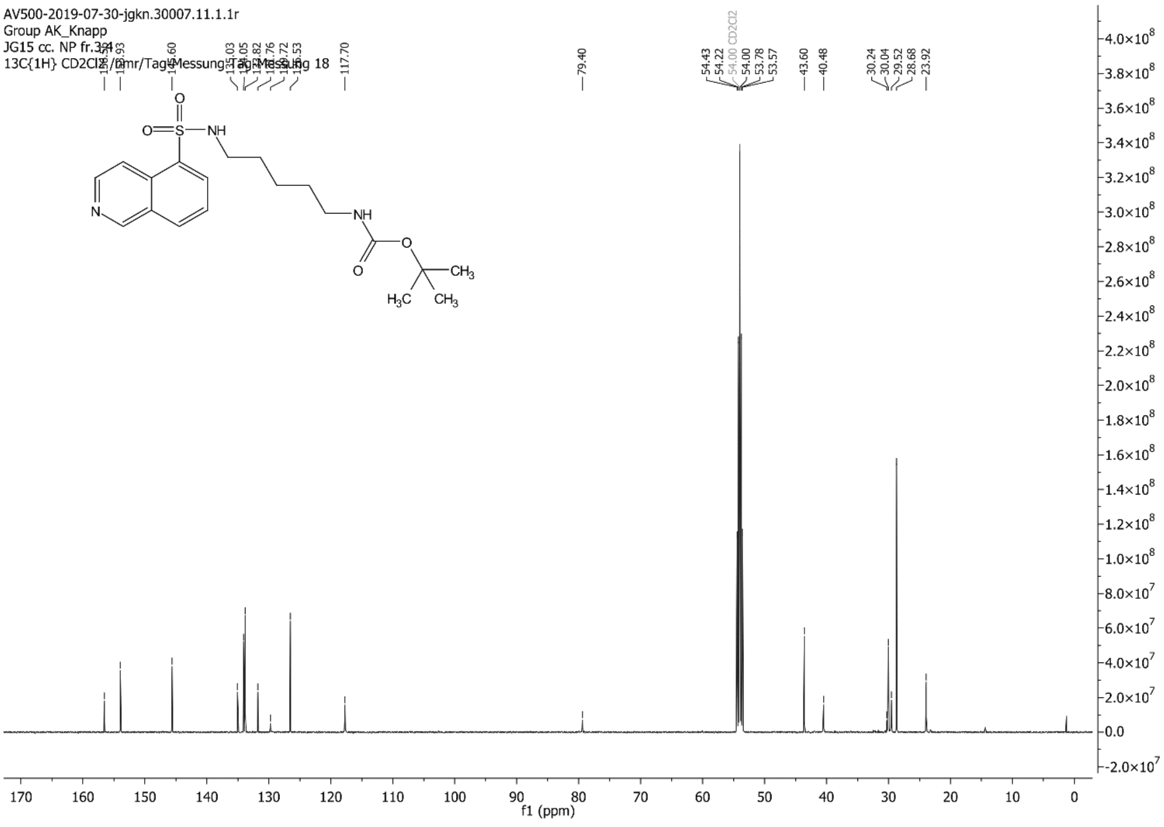

C:\Xcalibur\data\JG15-1 7/19/2019 11:45:09 AM

JG15-1 #32-43 RT: 0.54-0.74 AV: 12 SB: 13 0.07-0.28 NL: 2.69E6
T: {0,0} + c ESI !corona sid=75.00 det=1306.00 Full ms [105.00-800.00]

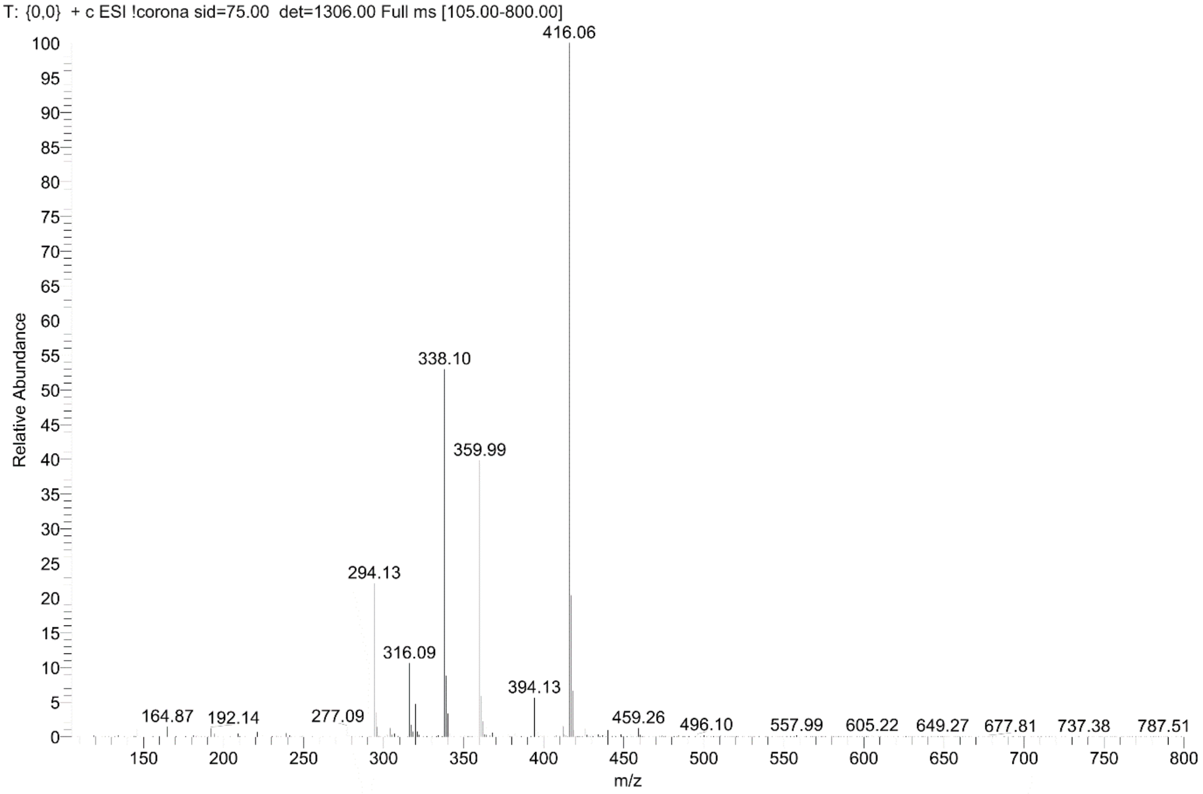

#### Intermediates Cmpd 17

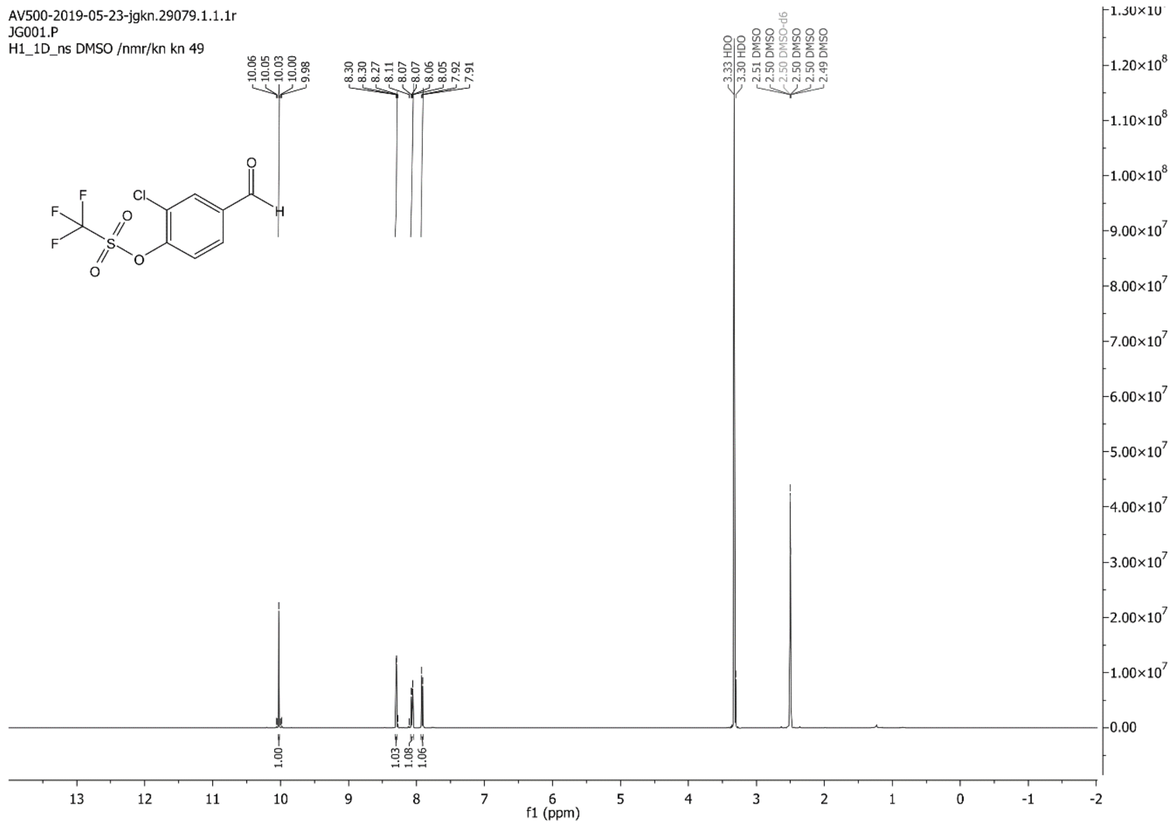

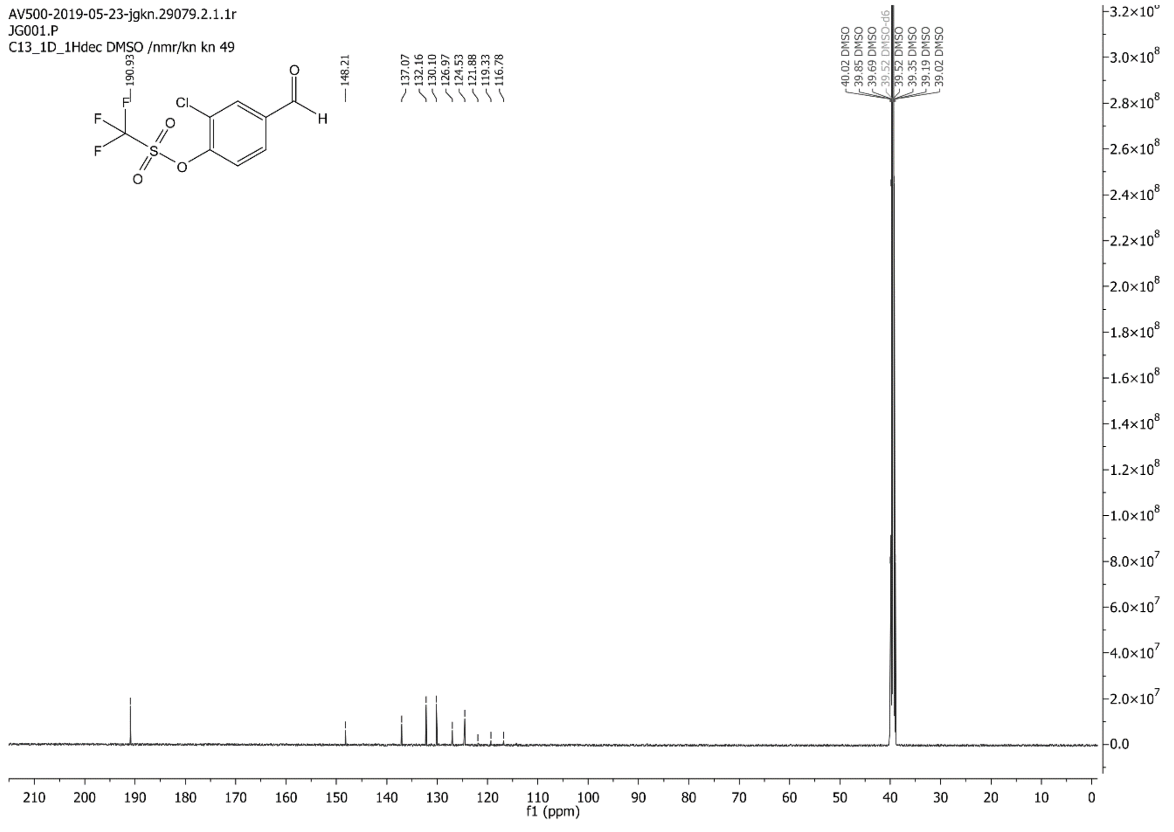

C:\Xcalibur\data\JG01-1 5/21/2019 7:21:06 AM

JG01-1 #43-44 RT: 0.73-0.75 AV: 2 SB: 9 0.16-0.29 NL: 2.44E4
T: {0,0} + c ESI !corona sid=75.00 det=1306.00 Full ms [105.00-600.00]

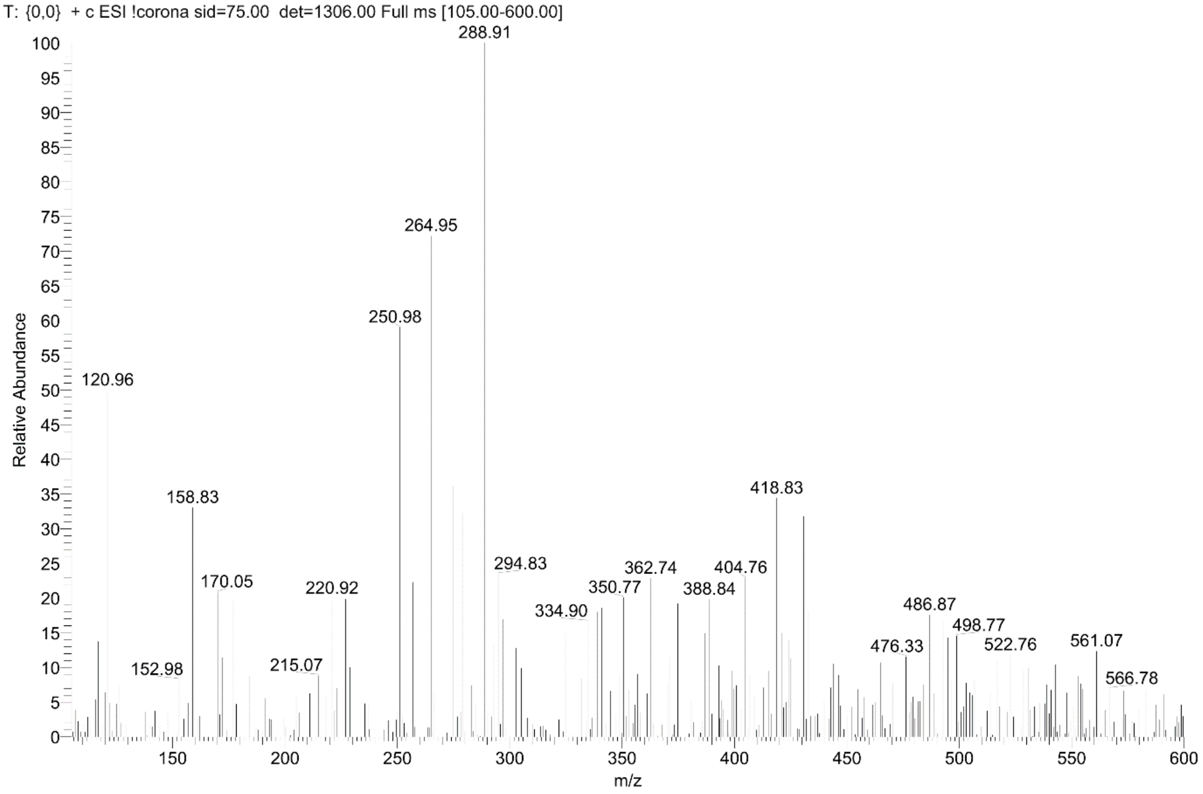

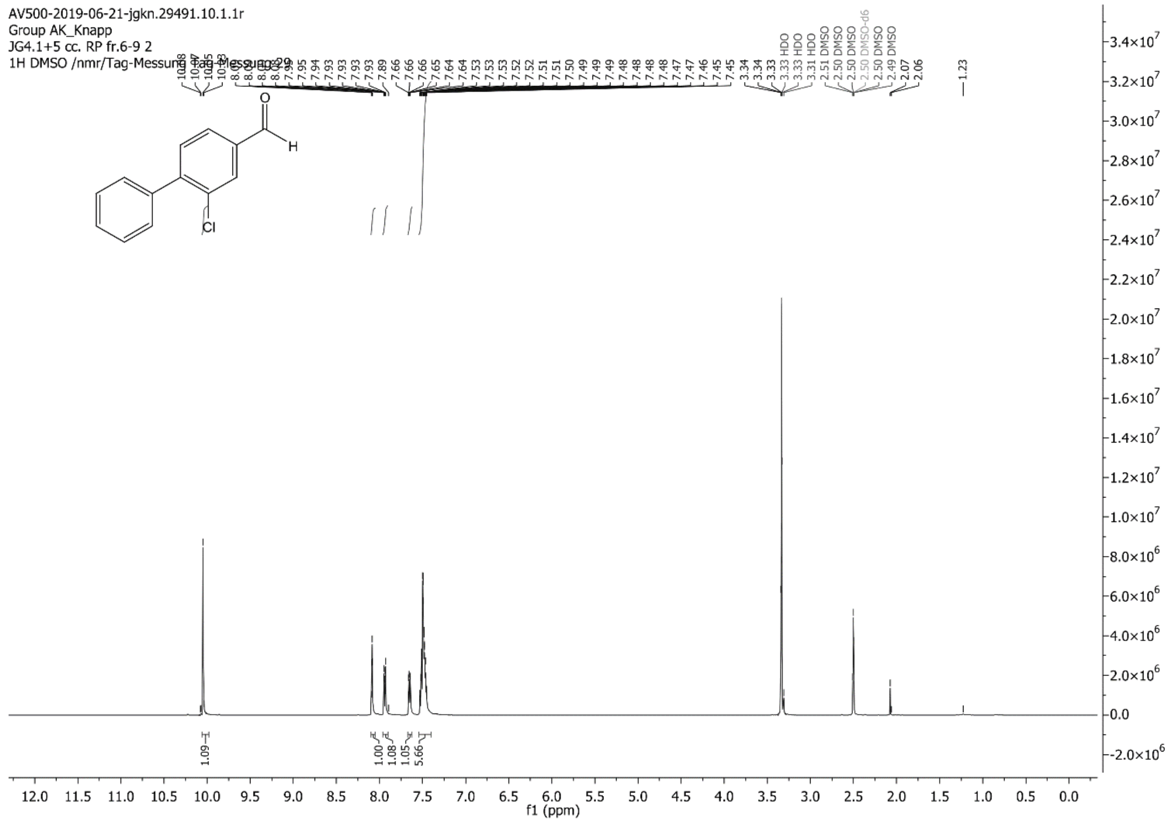
AV500-2019-06-21-jgkn.29491.10.1.1r
Group AK_Knapp
JG4.1+5 cc. RP fr.6-9 2
1H DMSO /nmr/Tag-Messung Tag-Messung 29
Cl
H
O
f1 (ppm)

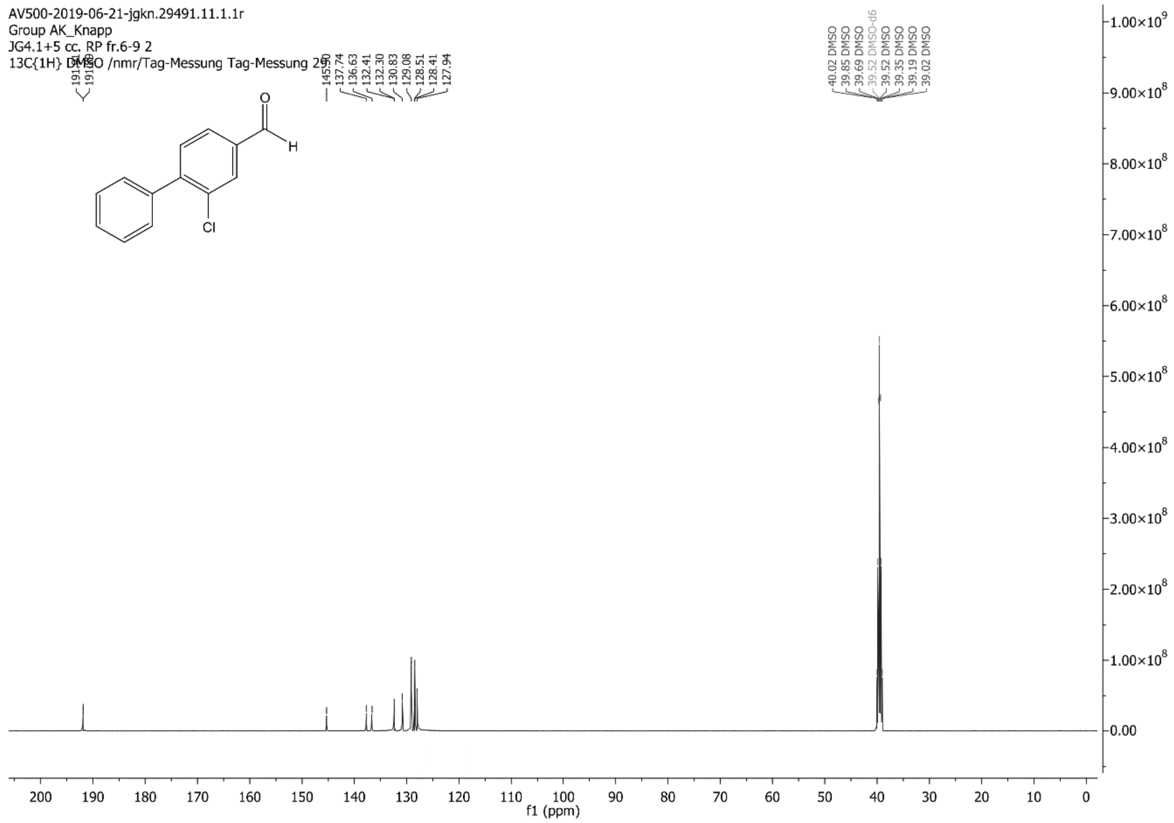
AV500-2019-06-21-jgkn.29491.11.1.1r
Group AK_Knapp
JG4.1+5 cc. RP fr.6-9 2
13C{1H} DMSO /nmr/Tag-Messung Tag-Messung 29
137.74
136.63
132.41
132.30
130.83
129.08
128.51
128.41
127.94
Cl
H
O
f1 (ppm)

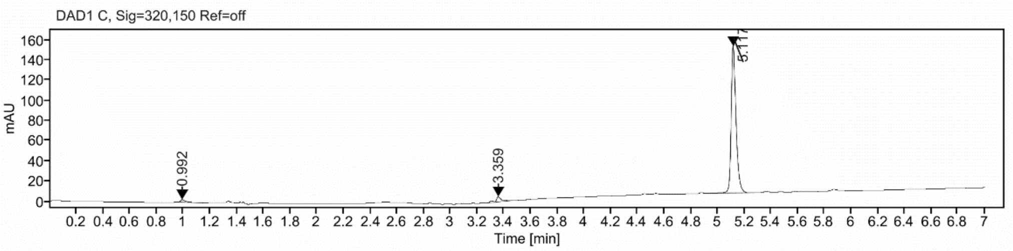

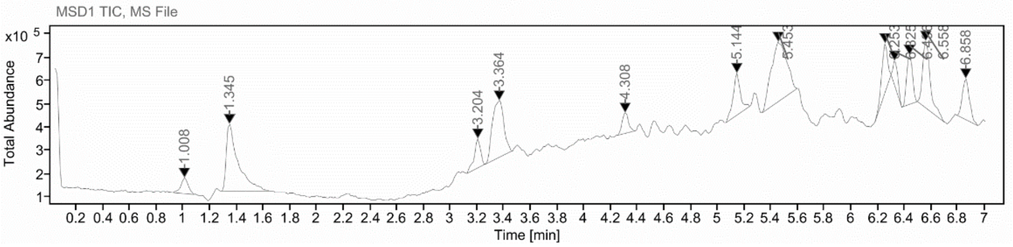

**Sample Purity**

Signal Description DAD1 C, Sig=320,150 Ref=off

| Sample Name | Name | RT | Width | Area | Area% | Height |
|---|---|---|---|---|---|---|
| FGC325_control | | 0.992 | 0.035 | 6.5760 | 1.68 | 2.3649 |
| FGC325_control | | 3.359 | 0.032 | 15.4574 | 3.96 | 4.9221 |
| FGC325_control | | 5.117 | 0.037 | 368.3724 | 94.36 | 145.1889 |

Max Area% 94.356

UV Signal Purity>95% Fail

**UV Apex Spectra**

**RT:**

5.117

**Sample Name:**

FGC325_control

**Signal Name:**

DAD1C

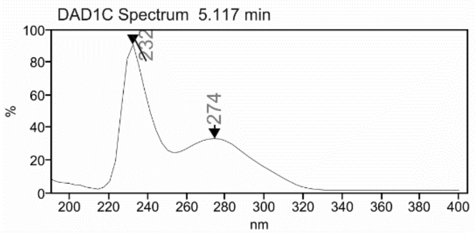

**RT:**

5.144

**Sample Name:**

FGC325_control

**Signal Name:**

MSD1TIC

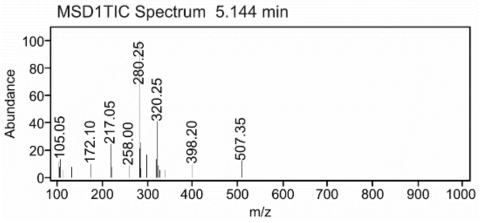

**Cmpd 19**

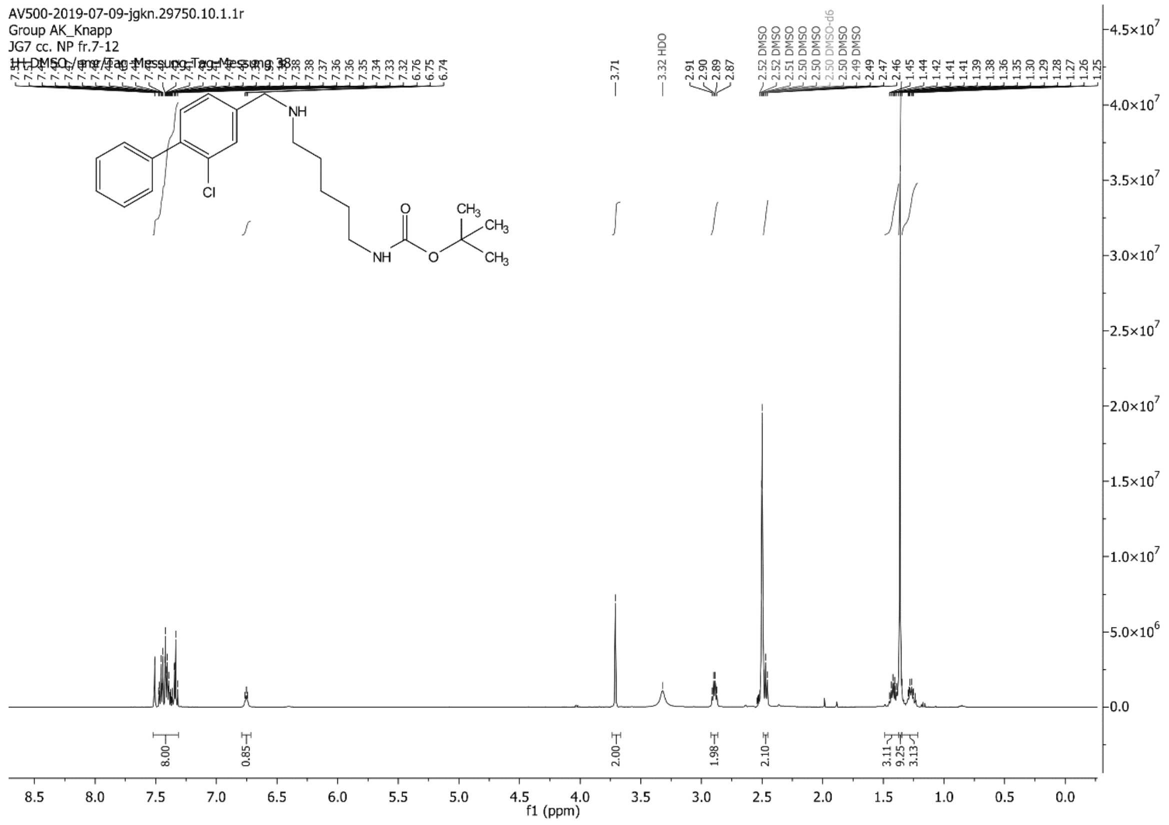
AV500-2019-07-09-jgkn.29750.10.1.1r
Group AK_Knapp
JG7 cc. NP fr.7-12
1H DMSO /nmr/Tag-Messung Tag-Messung 38
3.71
3.32 HDO
f1 (ppm)

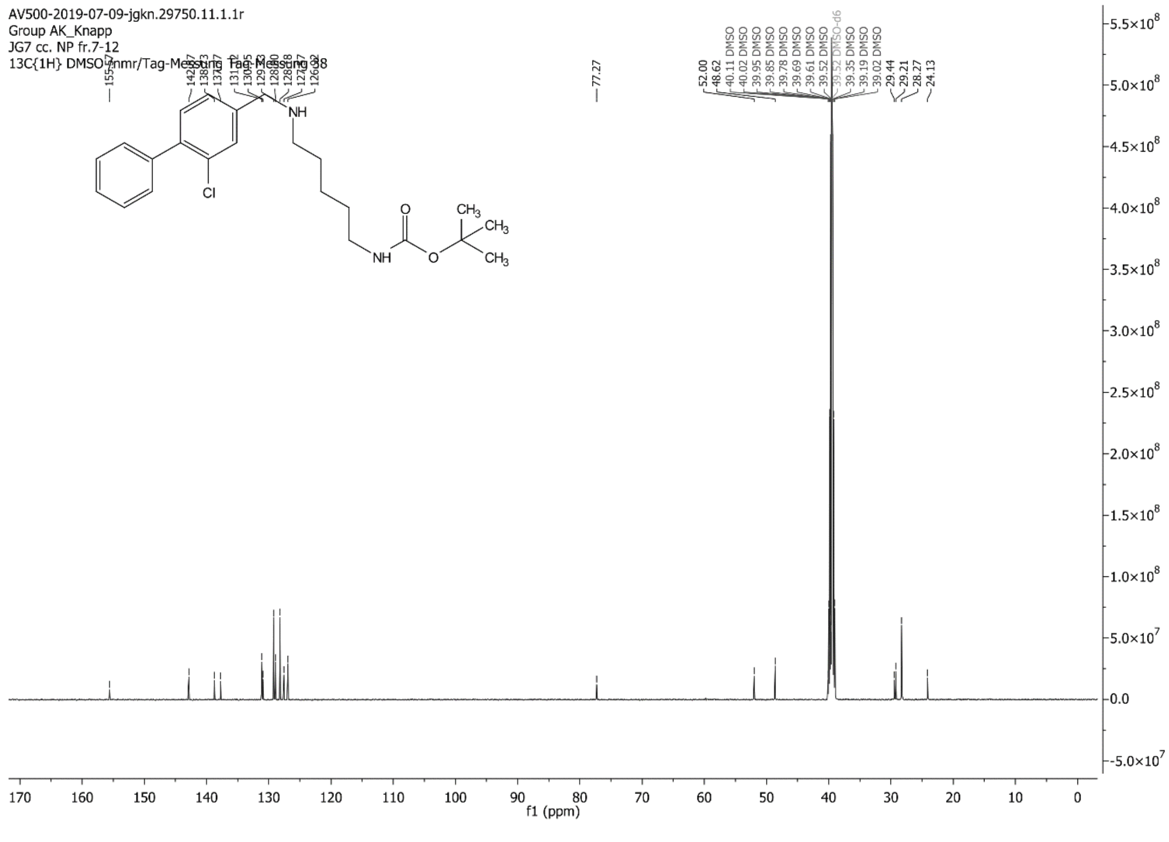
AV500-2019-07-09-jgkn.29750.11.1.1r
Group AK_Knapp
JG7 cc. NP fr.7-12
13C{1H} DMSO /nmr/Tag-Messung Tag-Messung 38
77.27
f1 (ppm)

C:\Xcalibur\data\JG7-2 7/4/2019 3:18:52 PM

JG7-2 #40-43 RT: 0.68-0.74 AV: 4 SB: 11 0.07-0.25 NL: 3.87E6
T: {0,0} + c ESI !corona sid=75.00 det=1306.00 Full ms [105.00-800.00]

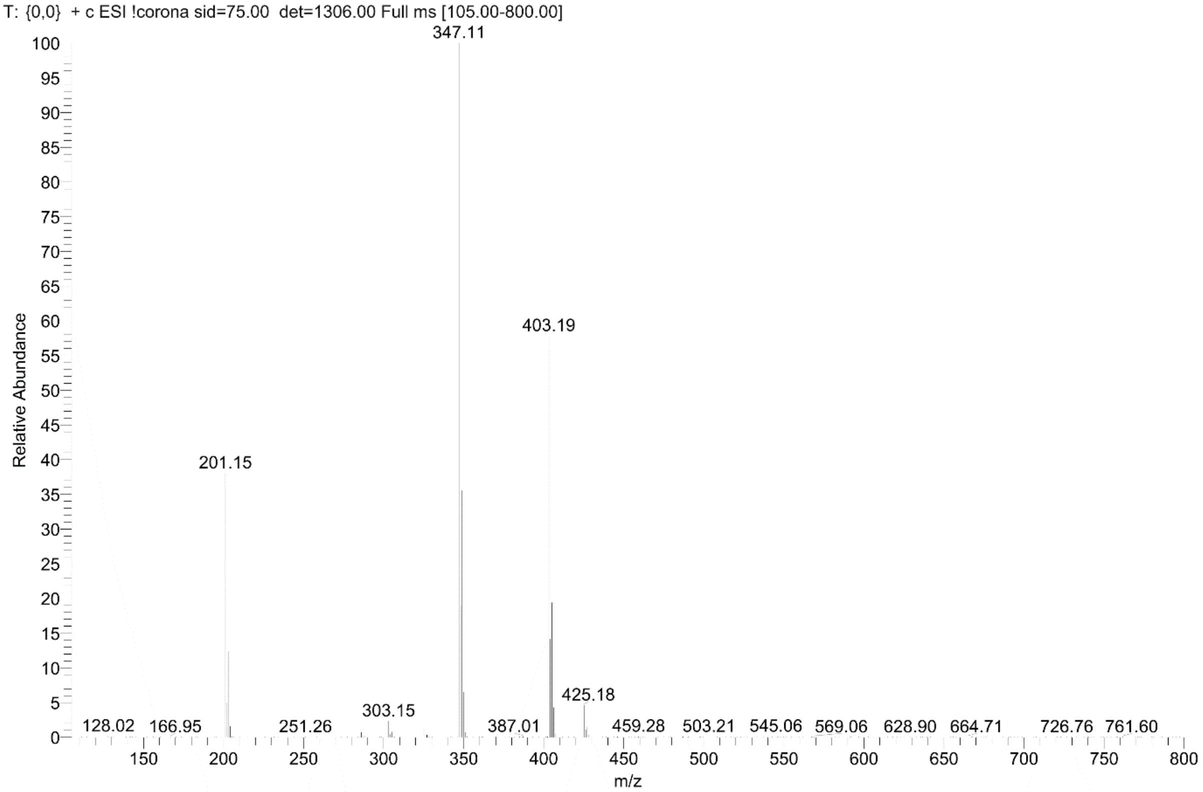

**Cmpd 20**

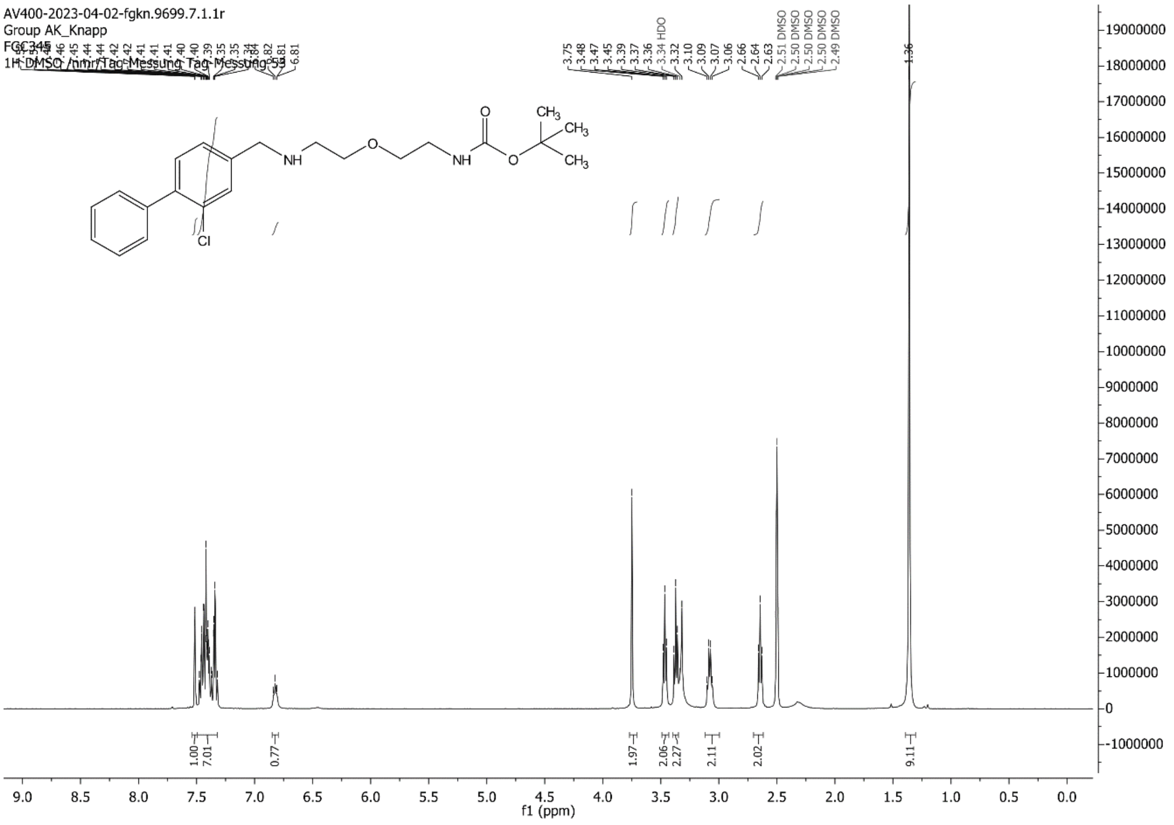

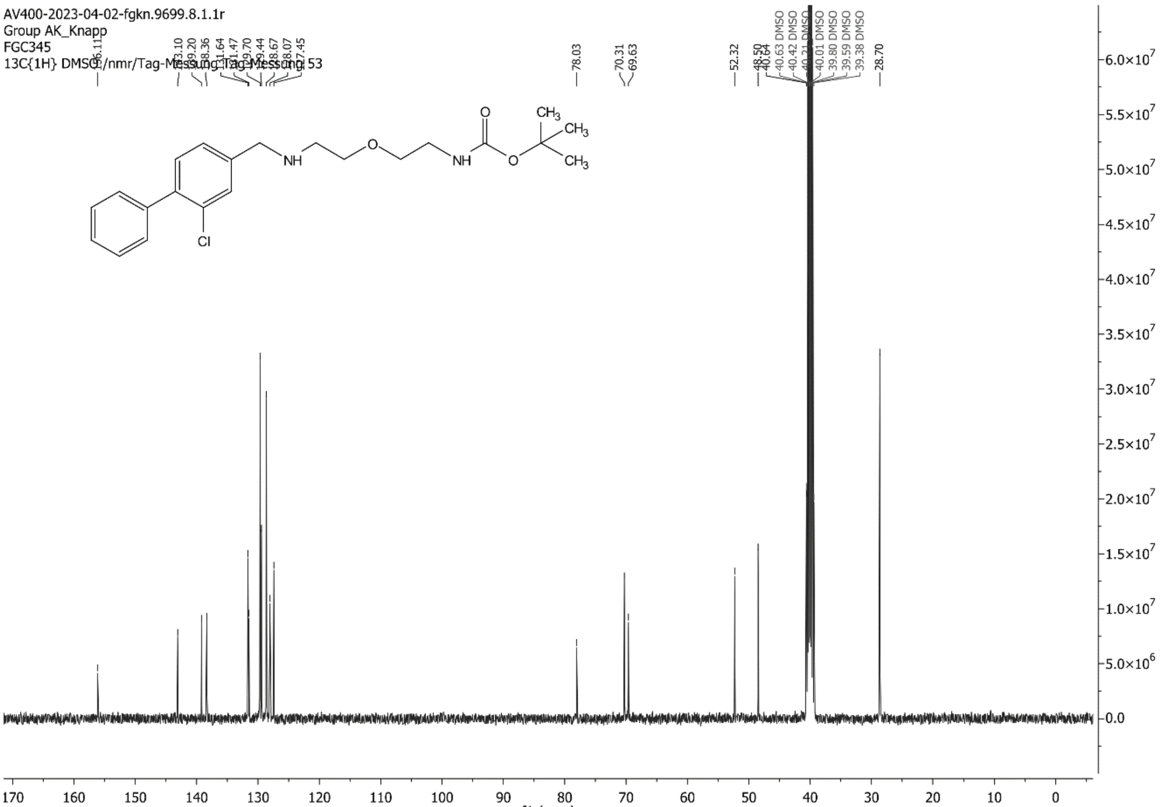

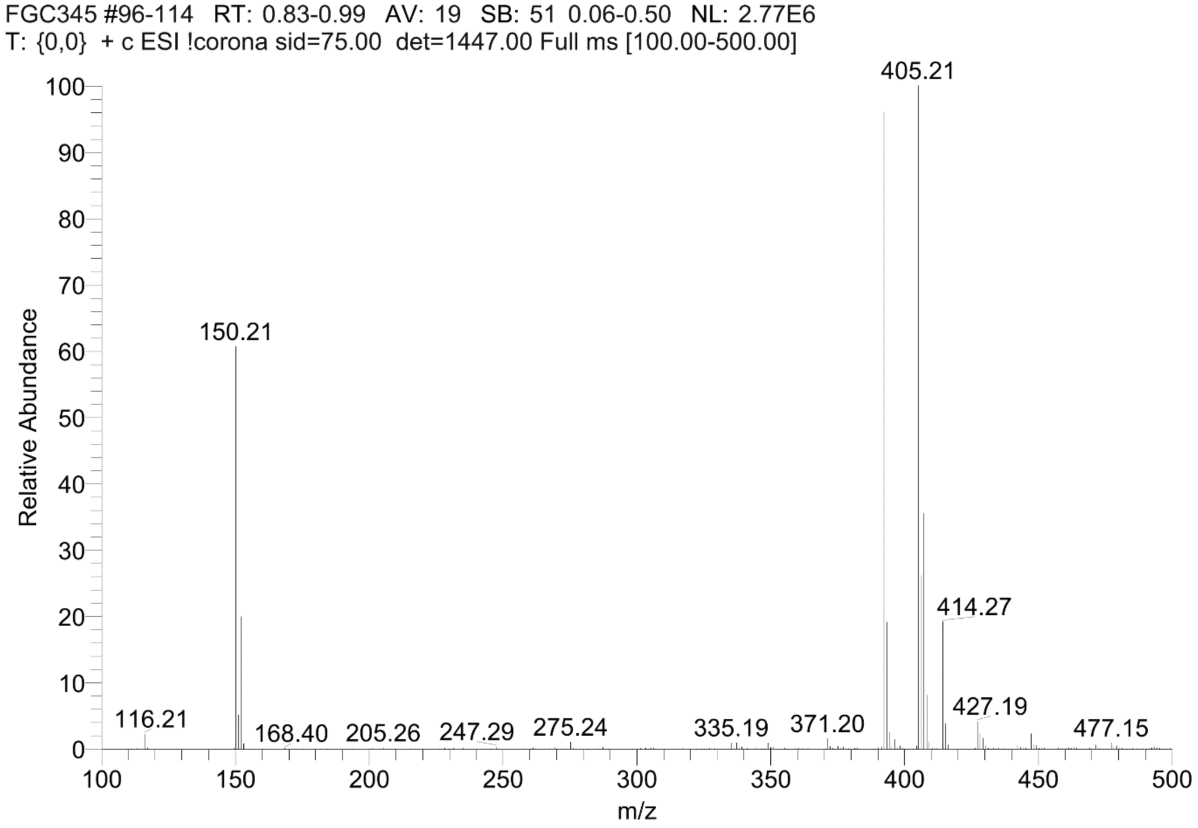
FGC345 #96-114 RT: 0.83-0.99 AV: 19 SB: 51 0.06-0.50 NL: 2.77E6
T: {0,0} + c ESI !corona sid=75.00 det=1447.00 Full ms [100.00-500.00]
405.21
150.21
414.27
116.21
168.40
205.26
247.29
275.24
335.19
371.20
427.19
477.15
Relative Abundance
m/z

**Cmpd 21**

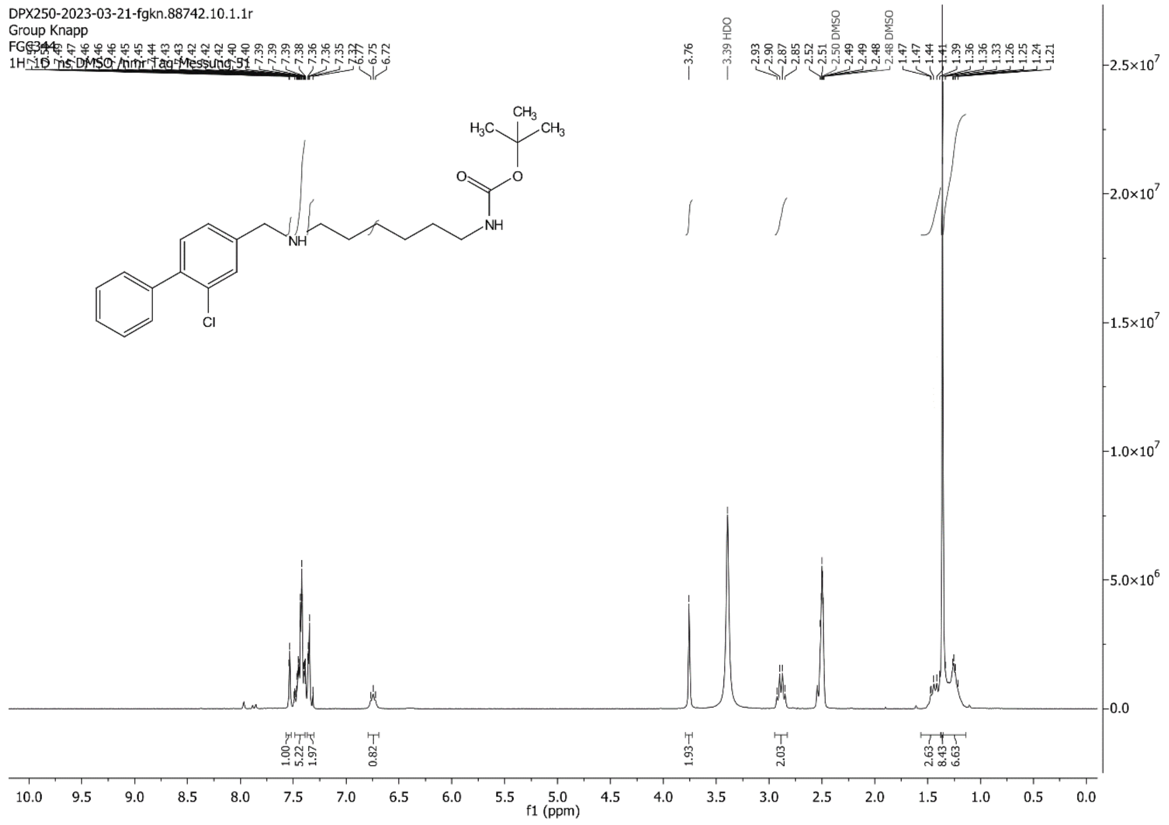
DPX250-2023-03-21-fgkn.88742.10.1.1r
Group Knapp
FGG344
1H 1D ns DMSO /nmr Tag Messung 51
3.76
3.39 HDO
2.50 DMSO
2.48 DMSO
Cl
NH
NH
O
O
$H_3C$
$CH_3$
$CH_3$
1.00
5.22
1.97
0.82
1.93
2.03
2.63
8.43
6.63
f1 (ppm)

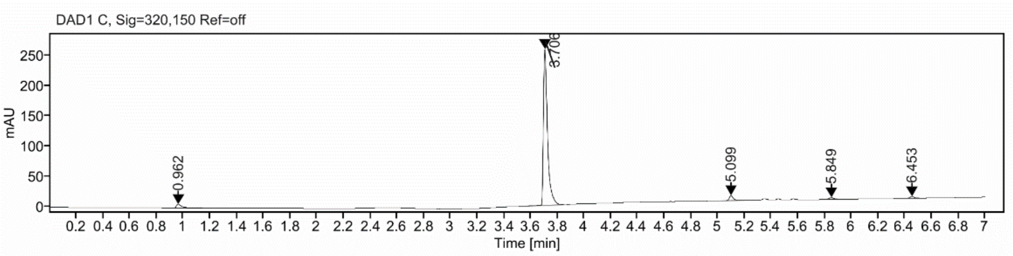

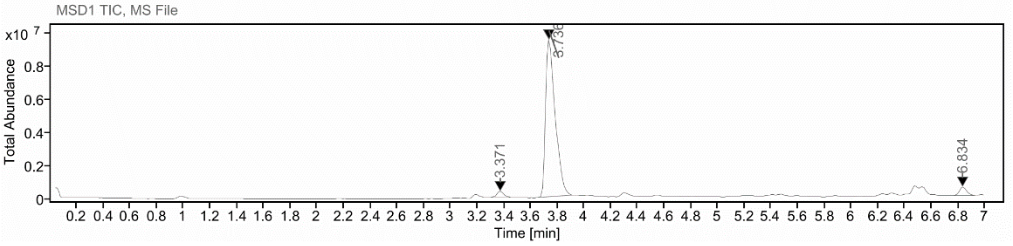

**Sample Purity**

Signal Description DAD1 C, Sig=320,150 Ref=off

| Sample Name | Name | RT | Width | Area | Area% | Height |
|---|---|---|---|---|---|---|
| FGC344 | | 0.962 | 0.037 | 20.7980 | 3.44 | 6.1086 |
| FGC344 | | 3.706 | 0.030 | 550.0280 | 90.95 | 258.0023 |
| FGC344 | | 5.099 | 0.032 | 18.8128 | 3.11 | 8.3917 |
| FGC344 | | 5.849 | 0.037 | 7.8384 | 1.30 | 3.1527 |
| FGC344 | | 6.453 | 0.036 | 7.2779 | 1.20 | 2.9818 |

Max Area% 90.951

UV Signal Purity>95% Fail

**UV Apex Spectra**

**RT:**

3.706

**Sample Name:**

FGC344

**Signal Name:**

DAD1C

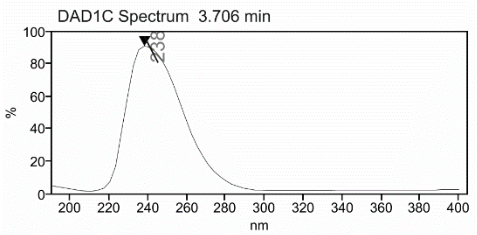

**RT:**

3.736

**Sample Name:**

FGC344

**Signal Name:**

MSD1TIC

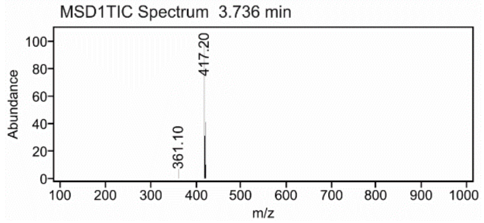

**Cmpd 22**

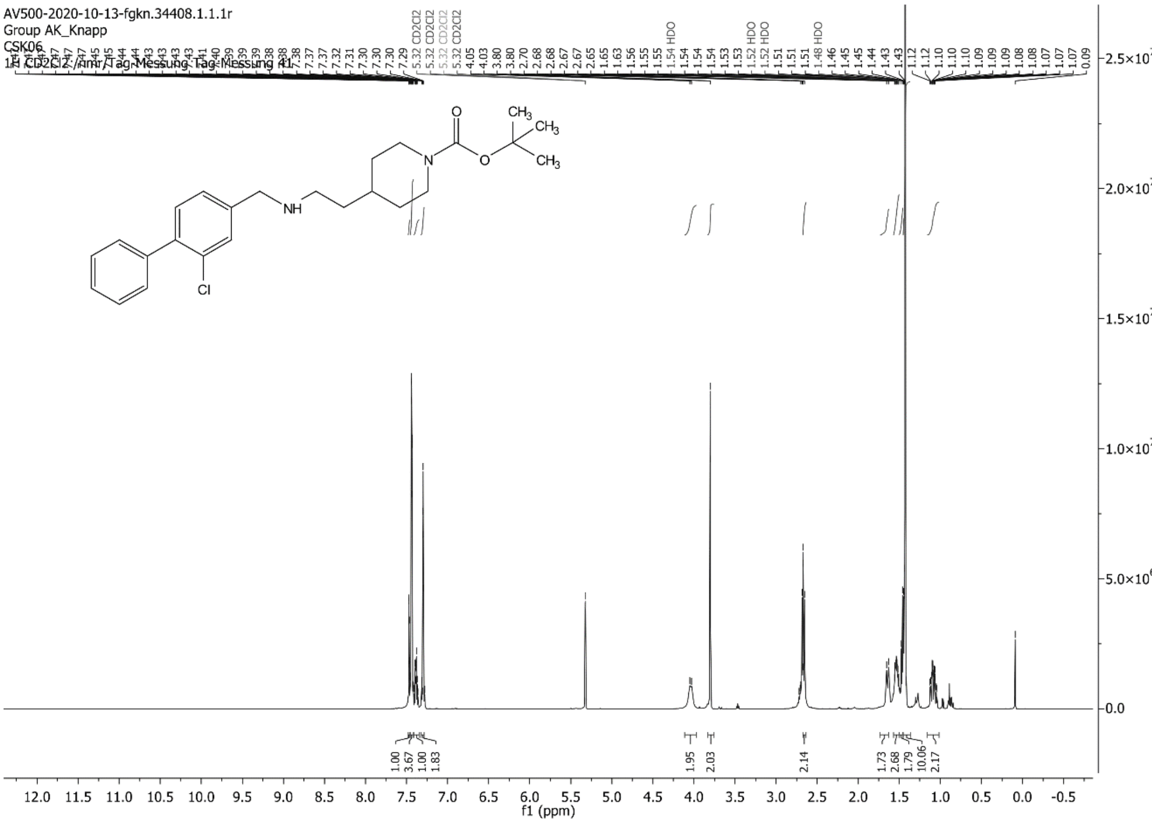

AV500-2020-10-13-fgkn.34408.1.1.1r
Group AK_Knapp
CSK06
1H CD2Cl2 /nmr/Tag-Messung Tag-Messung 41
f1 (ppm)

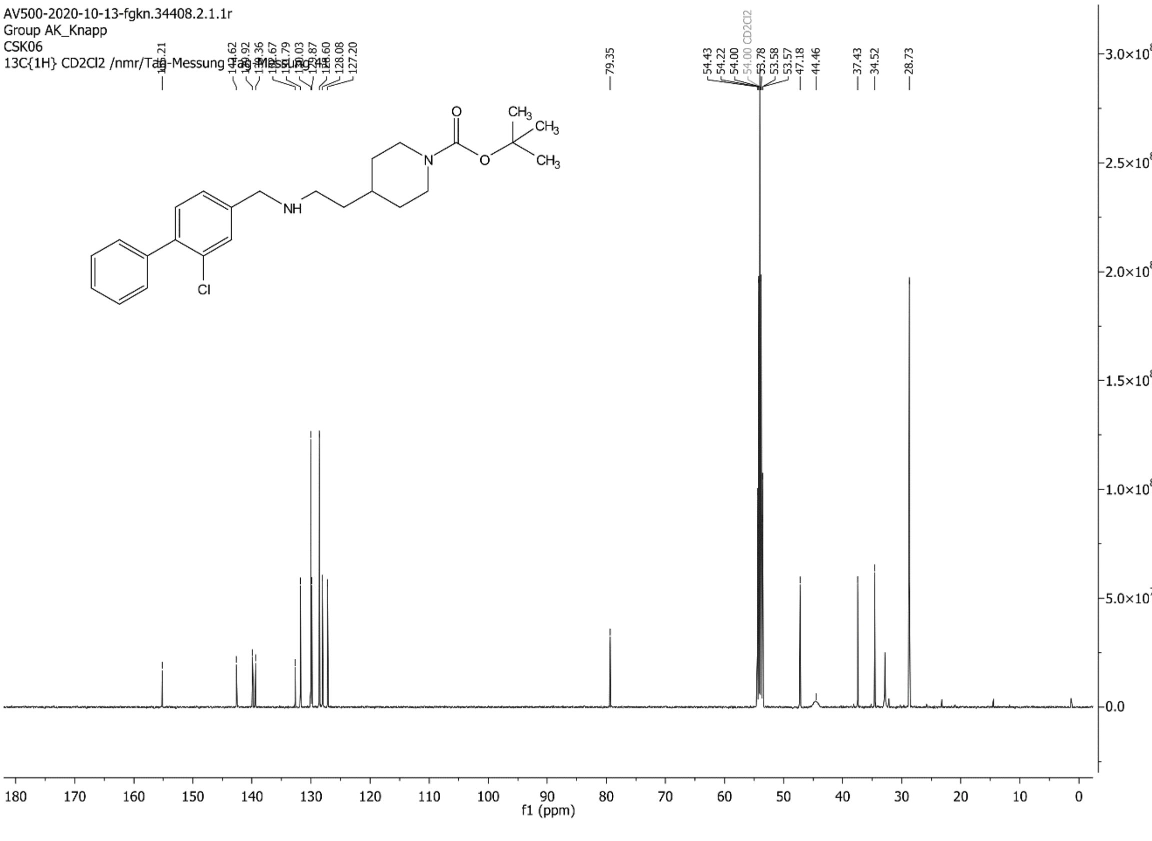

AV500-2020-10-13-fgkn.34408.2.1.1r
Group AK_Knapp
CSK06
13C{1H} CD2Cl2 /nmr/Tag-Messung Tag-Messung 41
f1 (ppm)

C:\Xcalibur\data\CK1-1 9/25/2020 10:27:39 AM

CK1-1 #38-43 RT: 0.65-0.74 AV: 6 SB: 8 0.16-0.28 NL: 5.33E6
T: {0,0} + c ESI !corona sid=75.00 det=1600.00 Full ms [105.00-800.00]

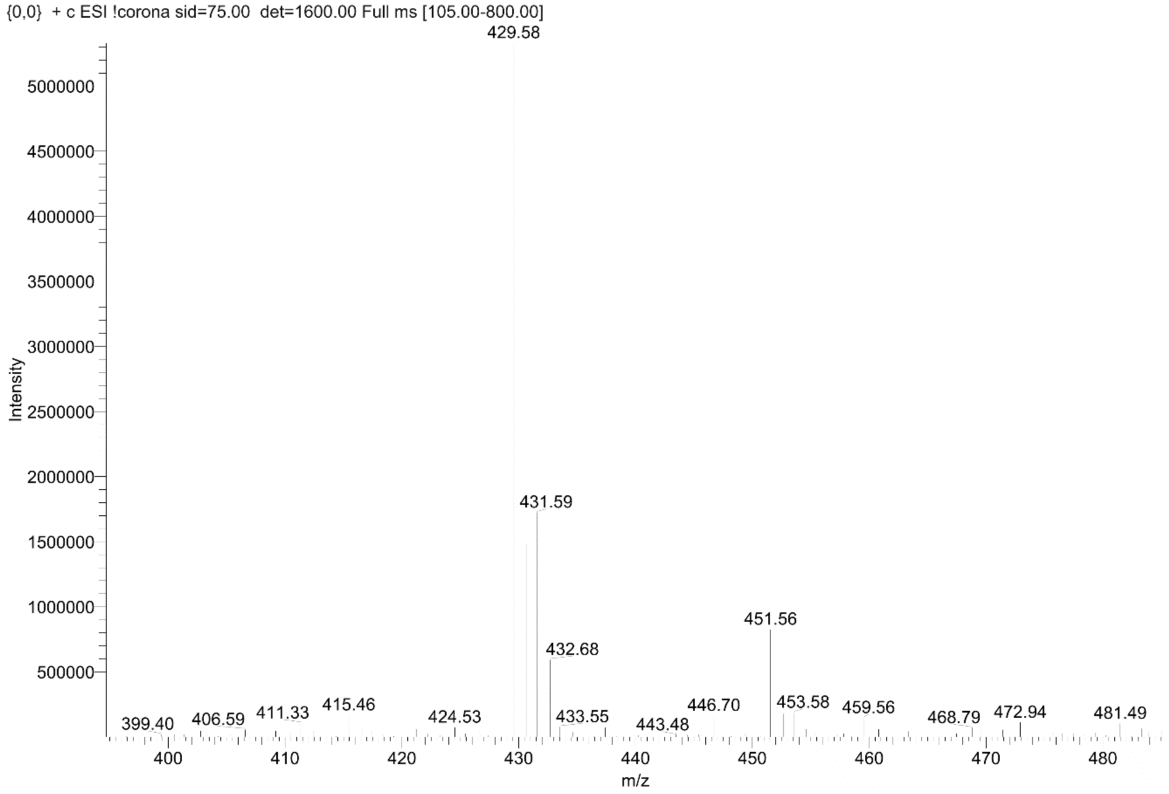

**Cmpd 23**

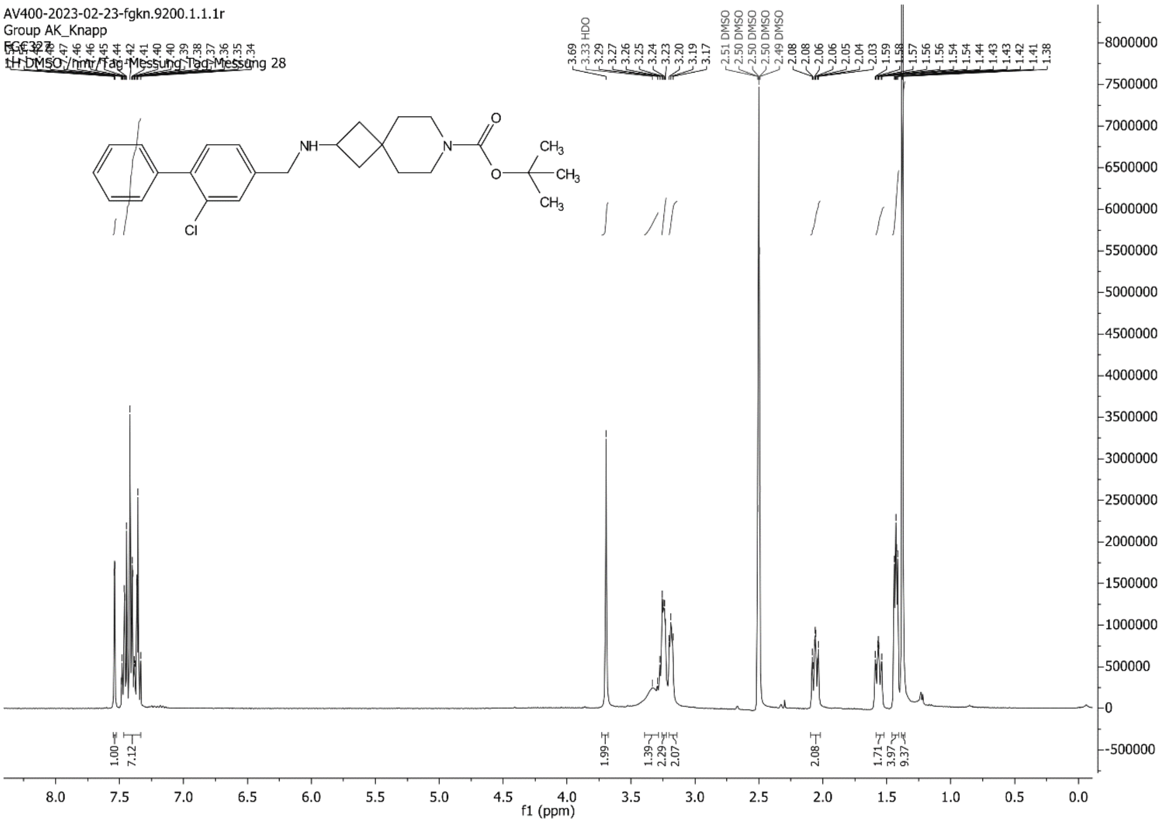

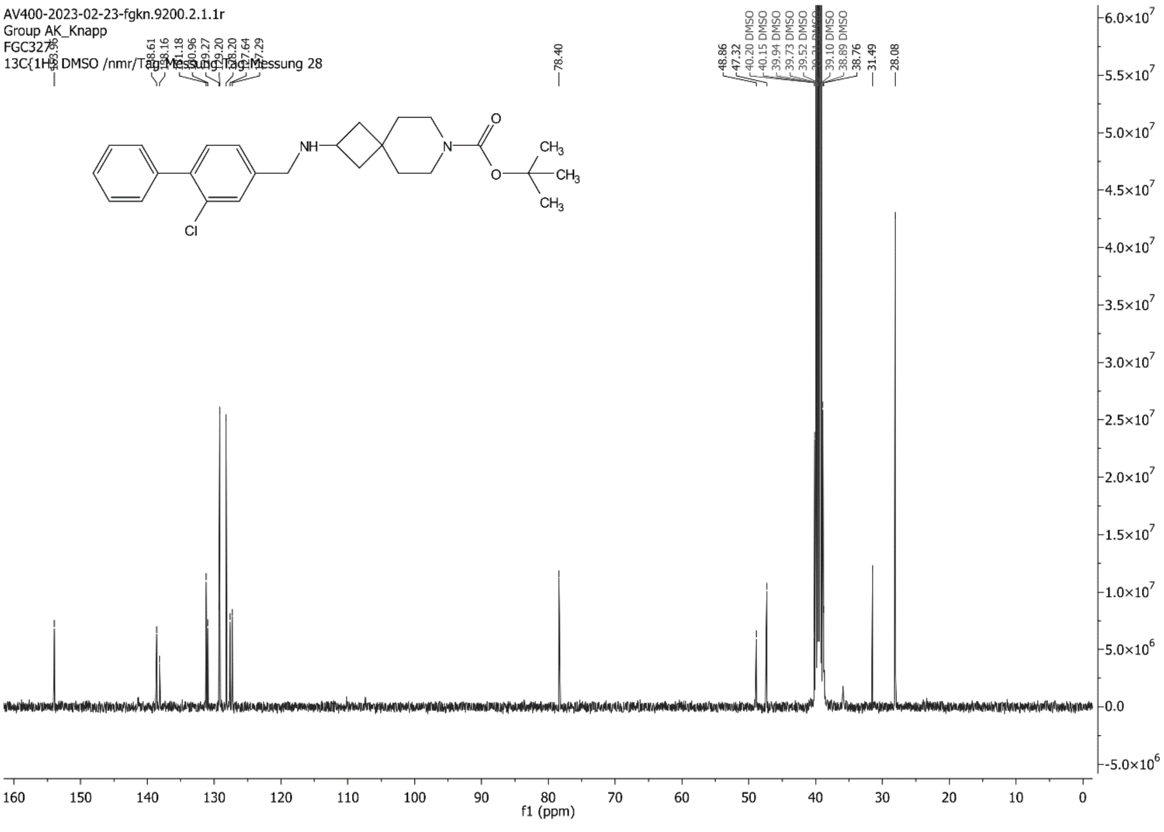

C:\Xcalibur\data\FGC327_2 02/23/23 10:16:58

FGC327_2 #35-42 RT: 0.60-0.72 AV: 8 SB: 20 0.12-0.46 NL: 2.82E6
T: {0,0} + c ESI !corona sid=75.00 det=1647.00 Full ms [105.00-800.00]

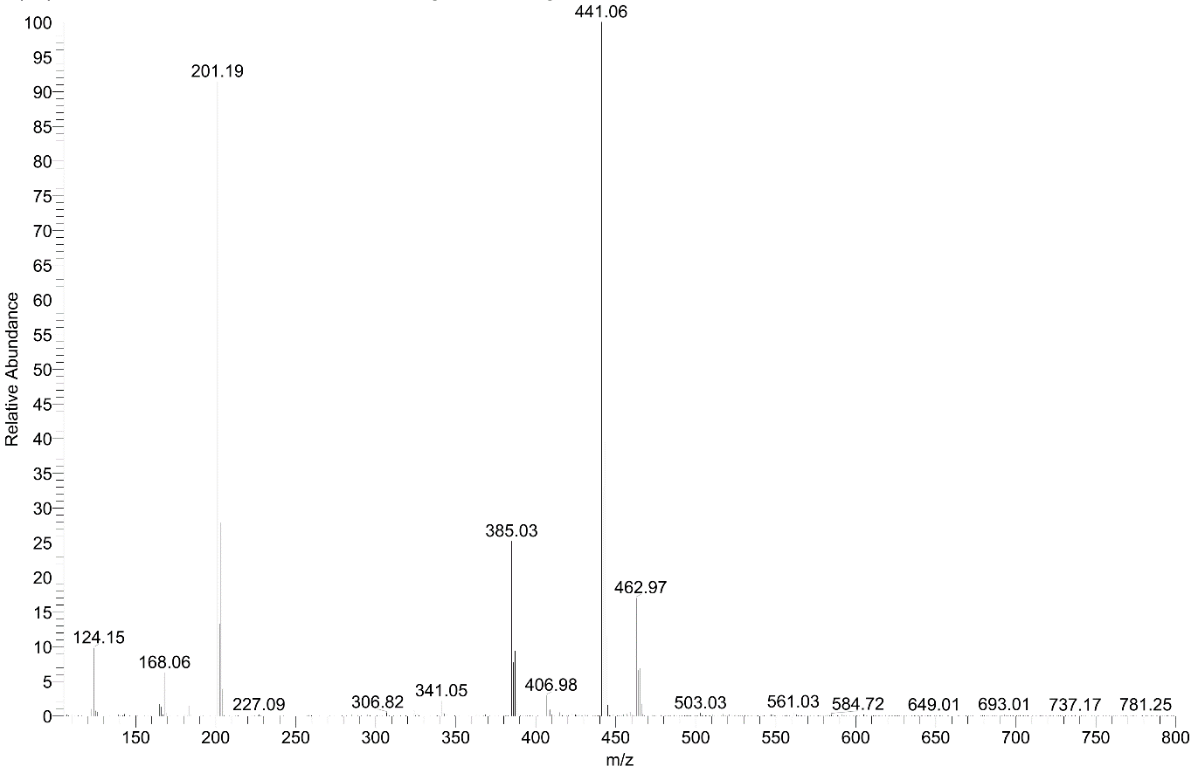

**Cmpd 24**

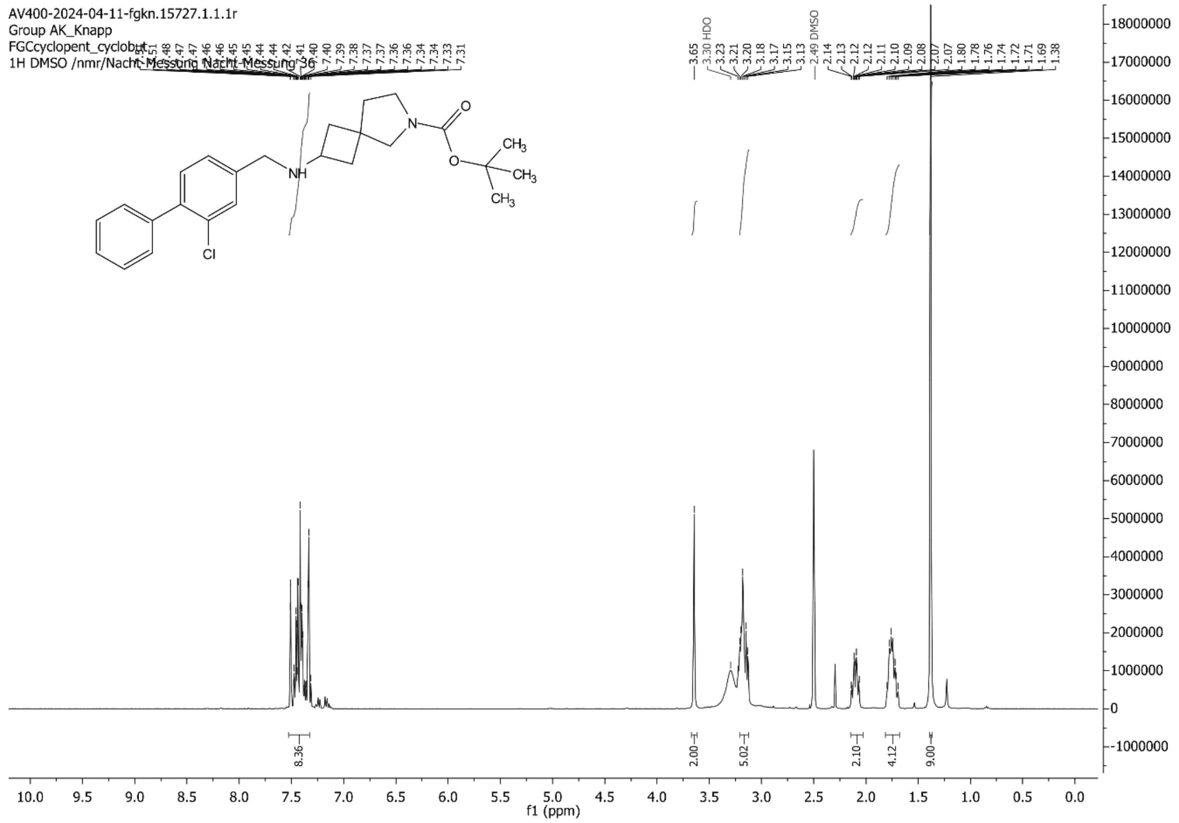

AV400-2024-04-11-fgkn.15727.1.1.1r
Group AK_Knapp
FGCcyclopent_cyclobut
1H DMSO /nmr/Nacht-Messung Nacht-Messung 36
f1 (ppm)

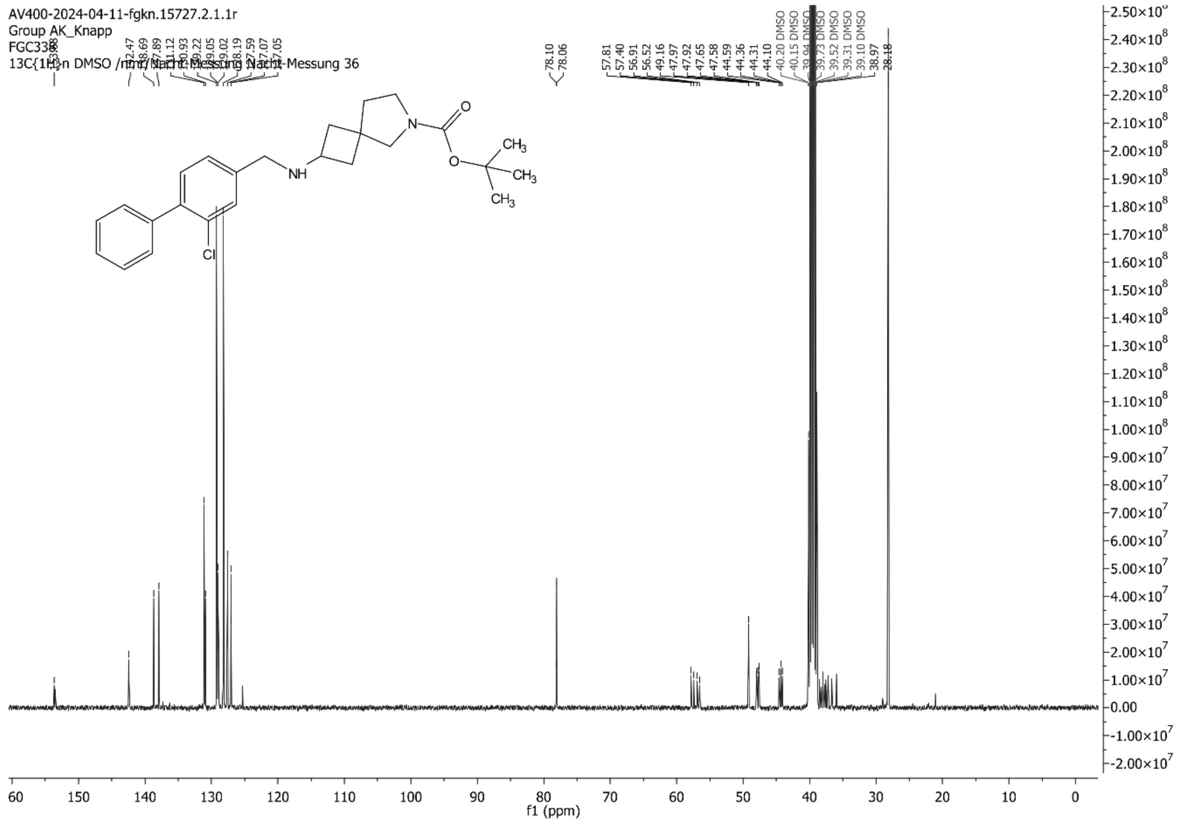

AV400-2024-04-11-fgkn.15727.2.1.1r
Group AK_Knapp
13C{1H} DMSO /nmr/Nacht-Messung Nacht-Messung 36
f1 (ppm)

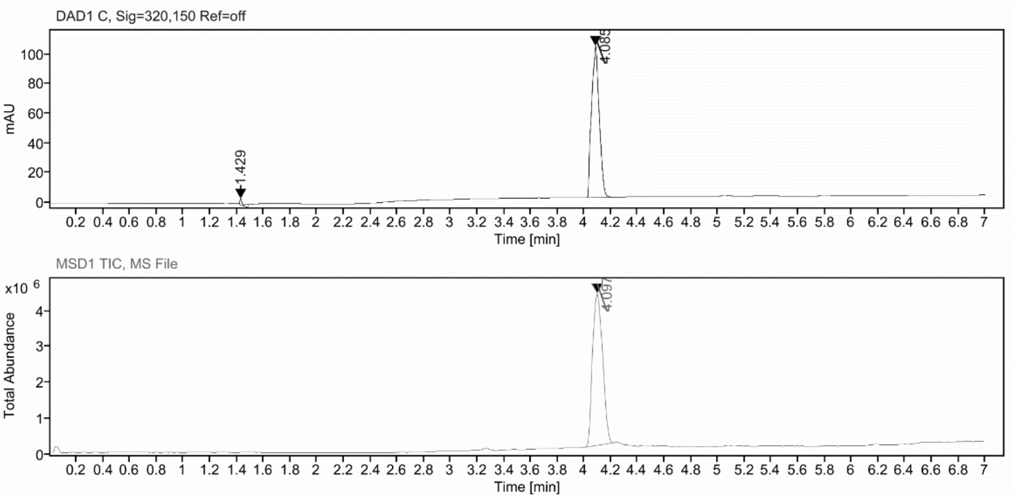

**Sample Purity**

Signal Description DAD1 C, Sig=320,150 Ref=off

| Sample Name | Name | RT | Width | Area | Area% | Height |
|---|---|---|---|---|---|---|
| FGC_cyclopent_cyclobut_13_14 | | 1.429 | 0.015 | 6.4417 | 1.56 | 4.1601 |
| FGC_cyclopent_cyclobut_13_14 | | 4.085 | 0.066 | 407.6381 | 98.44 | 102.3484 |

Max Area% 98.444

UV Signal Purity>95% Pass

**UV Apex Spectra**

**RT:**

4.085

**Sample Name:**

FGC_cyclopent_cyclobut_13_14

**Signal Name:**

DAD1C

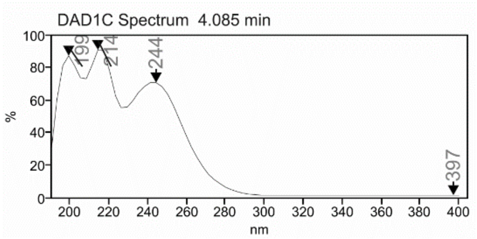

**MSD Apex Spectra**

**RT:**

4.097

**Sample Name:**

FGC_cyclopent_cyclobut_13_14

**Signal Name:**

MSD1TIC

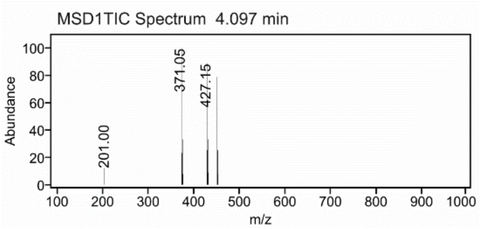

#### Cmpd 60

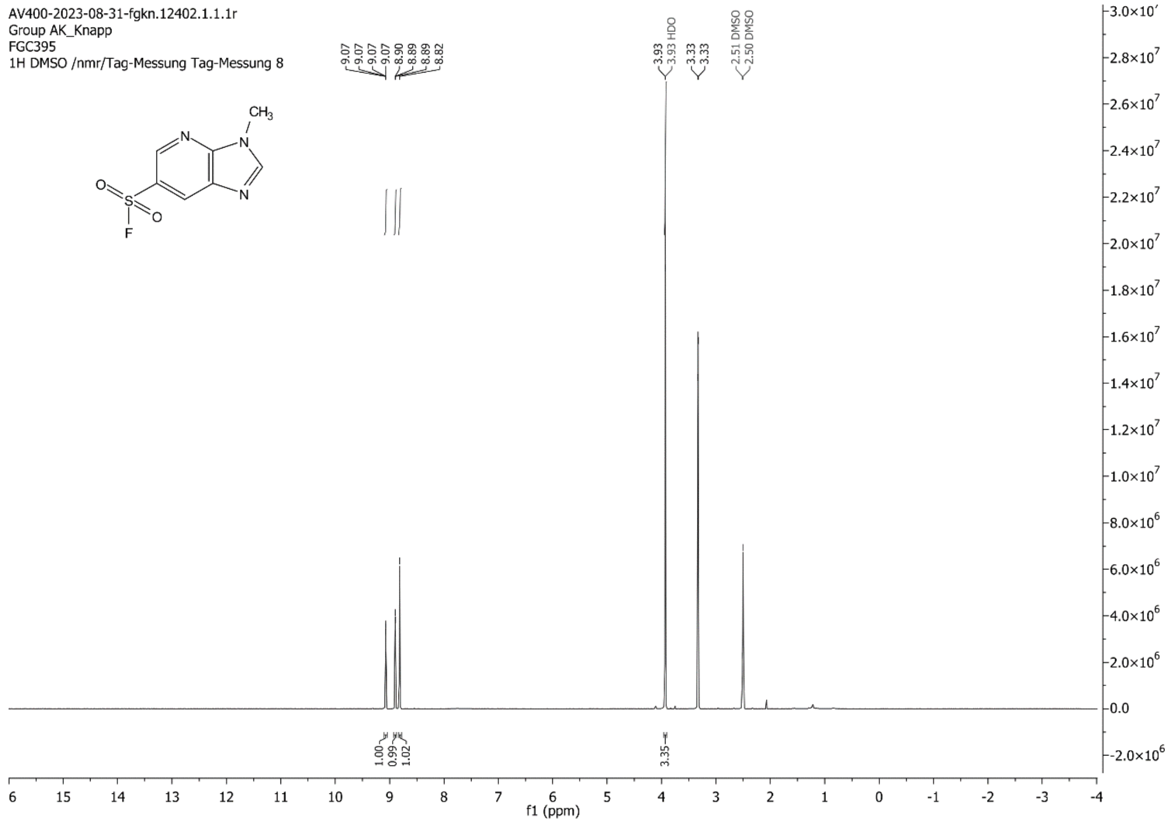
AV400-2023-08-31-fgkn.12402.1.1.1r
Group AK_Knapp
FGC395
1H DMSO /nmr/Tag-Messung Tag-Messung 8
9.07
9.07
9.07
9.07
8.90
8.89
8.89
8.82
3.93
3.93 HDO
3.33
3.33
2.51 DMSO
2.50 DMSO
CH3
N
N
N
O
S
O
F
1.00
0.99
1.02
3.35
f1 (ppm)

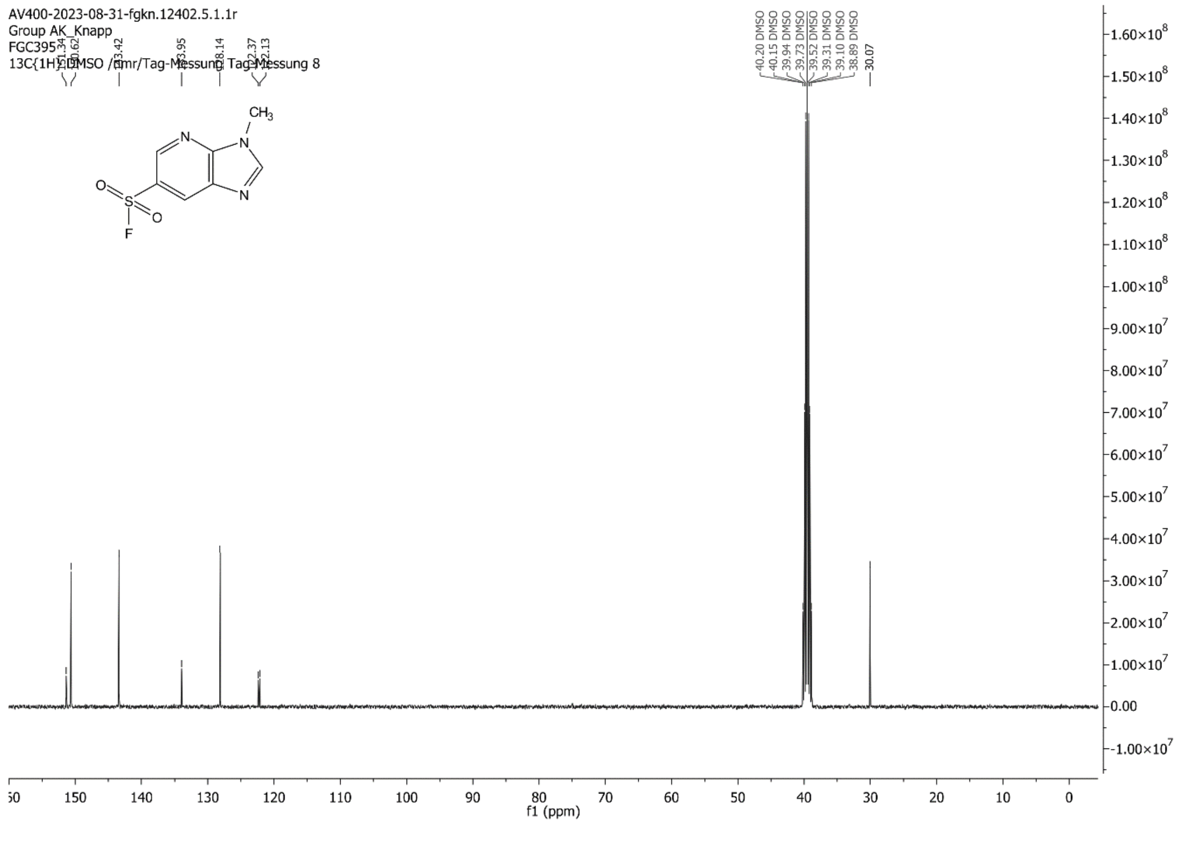
AV400-2023-08-31-fgkn.12402.5.1.1r
Group AK_Knapp
FGC395
13C{1H} DMSO /nmr/Tag-Messung Tag-Messung 8
151.34
150.62
143.42
133.95
128.14
122.37
122.13
40.20 DMSO
40.15 DMSO
39.94 DMSO
39.73 DMSO
39.52 DMSO
39.31 DMSO
39.10 DMSO
38.89 DMSO
30.07
CH3
N
N
N
O
S
O
F
f1 (ppm)

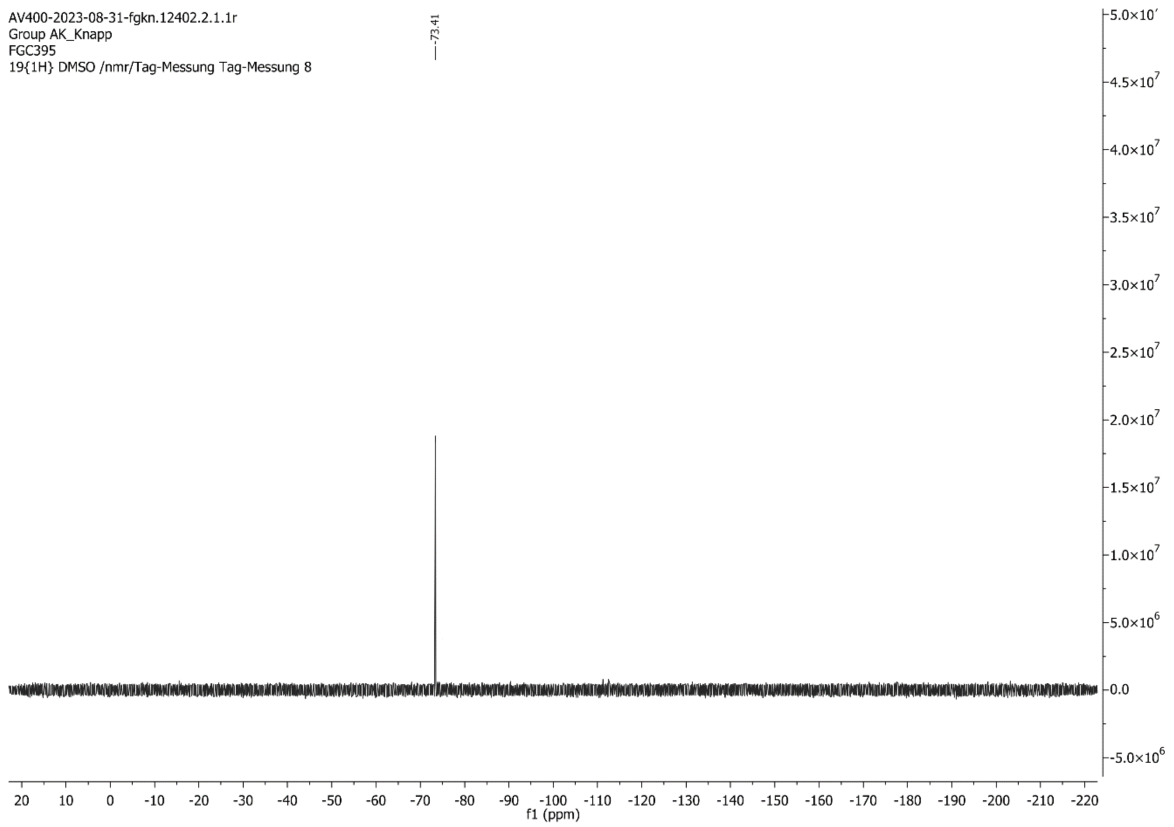
AV400-2023-08-31-fgkn.12402.2.1.1r
Group AK_Knapp
FGC395
19{1H} DMSO /nmr/Tag-Messung Tag-Messung 8
-73.41
f1 (ppm)

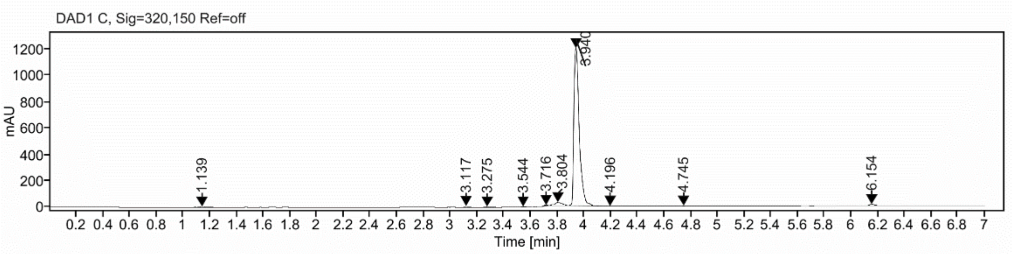

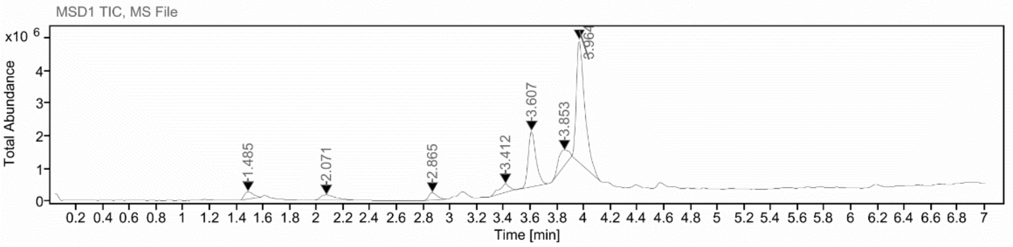

**Sample Purity**

Signal Description DAD1 C, Sig=320,150 Ref=off

| Sample Name | Name | RT | Width | Area | Area% | Height |
|---|---|---|---|---|---|---|
| FGC395_2_10_X | | 1.139 | 0.046 | 21.4046 | 0.60 | 5.9480 |
| FGC395_2_10_X | | 3.117 | 0.025 | 15.7861 | 0.44 | 8.3653 |
| FGC395_2_10_X | | 3.275 | 0.034 | 13.5879 | 0.38 | 4.6231 |
| FGC395_2_10_X | | 3.544 | 0.027 | 11.1750 | 0.31 | 3.0150 |
| FGC395_2_10_X | | 3.716 | 0.033 | 12.0629 | 0.34 | 6.5883 |
| FGC395_2_10_X | | 3.804 | 0.058 | 91.4198 | 2.57 | 23.2438 |
| FGC395_2_10_X | | 3.940 | 0.041 | 3348.3936 | 94.05 | 1200.0676 |
| FGC395_2_10_X | | 4.196 | 0.042 | 6.7412 | 0.19 | 2.4056 |
| FGC395_2_10_X | | 4.745 | 0.036 | 6.9909 | 0.20 | 2.5594 |
| FGC395_2_10_X | | 6.154 | 0.039 | 32.7617 | 0.92 | 12.1968 |

Max Area% 94.047

UV Signal Purity>95% Fail

**UV Apex Spectra**

**RT:**

3.940

**Sample Name:**

FGC395_2_10_X

**Signal Name:**

DAD1C

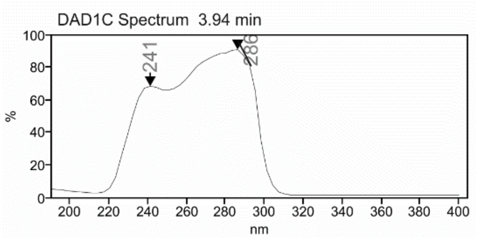

**RT:**

3.964

**Sample Name:**

FGC395_2_10_X

**Signal Name:**

MSD1TIC

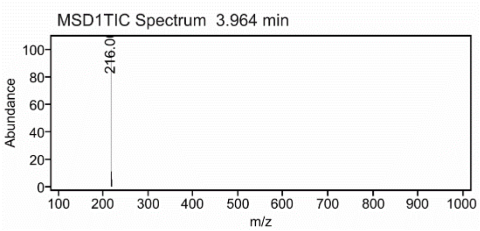

**Cmpd 62**

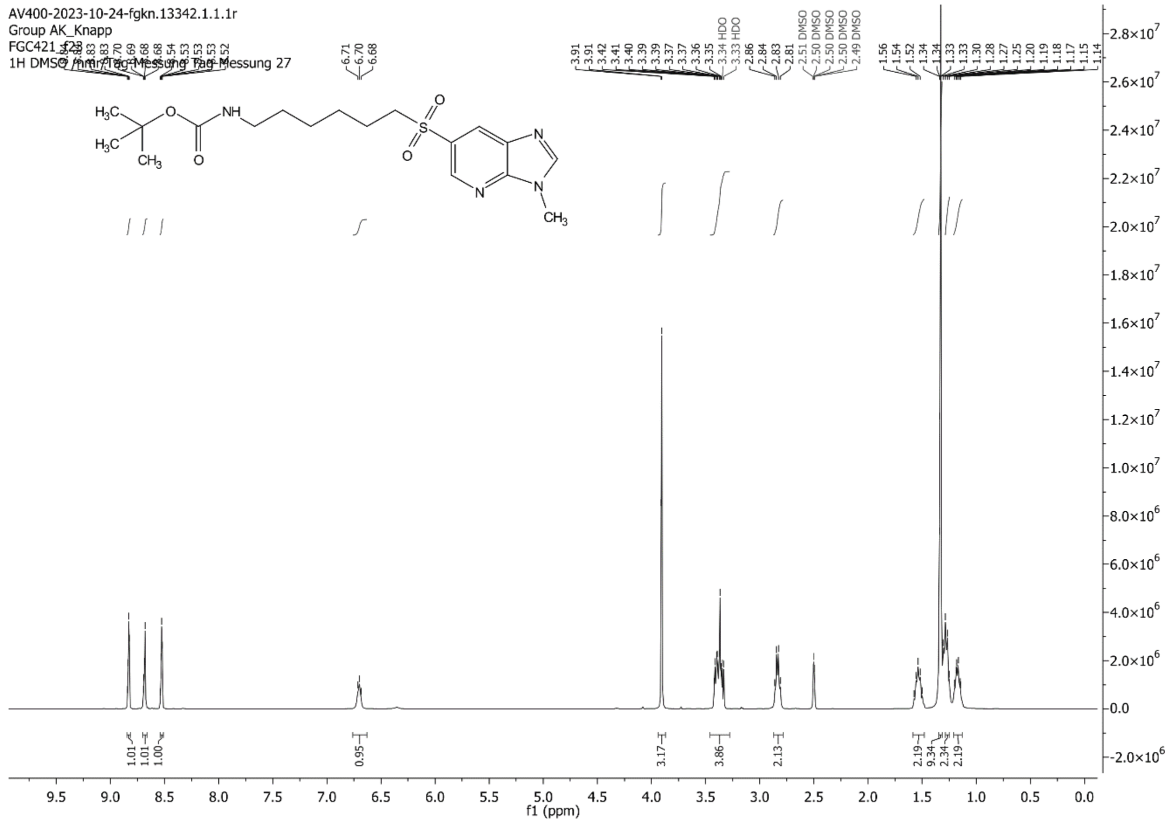

AV400-2023-10-24-fgkn.13342.1.1.1r
Group AK_Knapp
FGC421_f23
1H DMSO /nmr/Tag-Messung Tag-Messung 27
f1 (ppm)

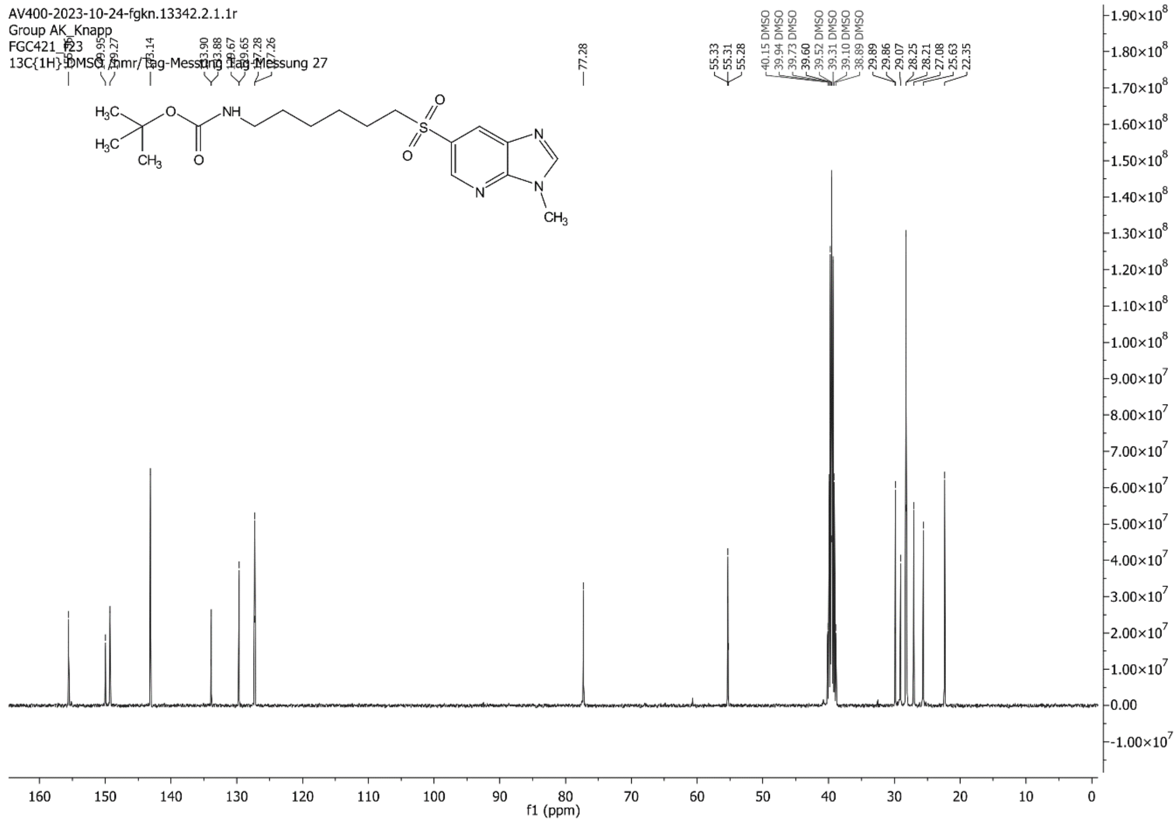

AV400-2023-10-24-fgkn.13342.2.1.1r
Group AK_Knapp
FGC421_f23
13C{1H} DMSO /nmr/Tag-Messung Tag-Messung 27
f1 (ppm)

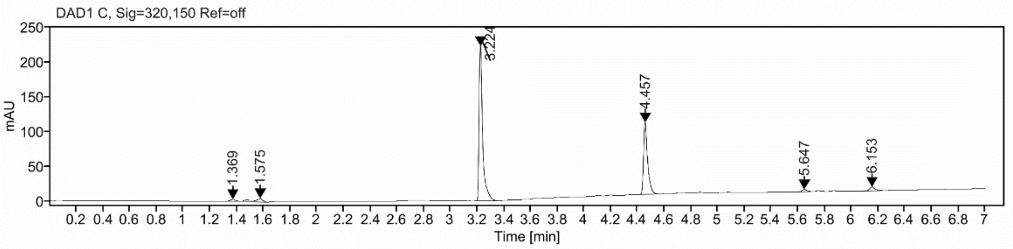

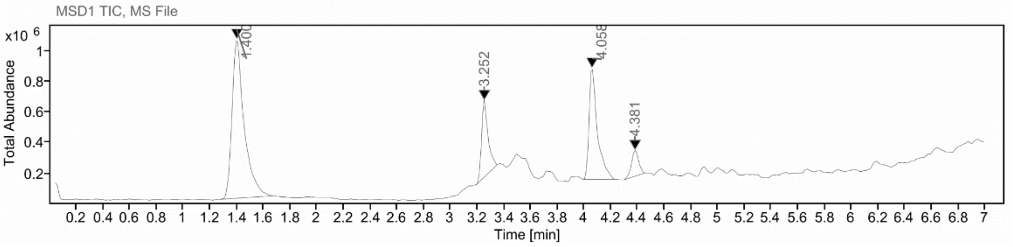

**Sample Purity**

Signal Description DAD1 C, Sig=320,150 Ref=off

| Sample Name | Name | RT | Width | Area | Area% | Height |
|---|---|---|---|---|---|---|
| FGC421_depro_eppi | | 1.369 | 0.034 | 6.9028 | 1.01 | 2.8575 |
| FGC421_depro_eppi | | 1.575 | 0.043 | 22.6987 | 3.34 | 5.0211 |
| FGC421_depro_eppi | | 3.224 | 0.024 | 407.1072 | 59.86 | 222.5836 |
| FGC421_depro_eppi | | 4.457 | 0.031 | 221.3668 | 32.55 | 103.5213 |
| FGC421_depro_eppi | | 5.647 | 0.038 | 8.0640 | 1.19 | 3.2283 |
| FGC421_depro_eppi | | 6.153 | 0.038 | 13.9749 | 2.05 | 5.2637 |

Max Area% 59.859

UV Signal Purity>95% Fail

**UV Apex Spectra**

**RT:**

3.224

**Sample Name:**

FGC421_depro_eppi

**Signal Name:**

DAD1C

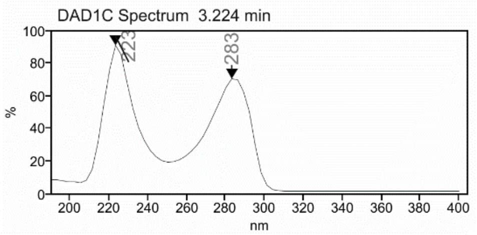

**RT:**

3.252

**Sample Name:**

FGC421_depro_eppi

**Signal Name:**

MSD1TIC

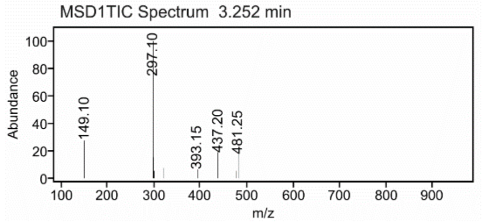

### Cmpd 65

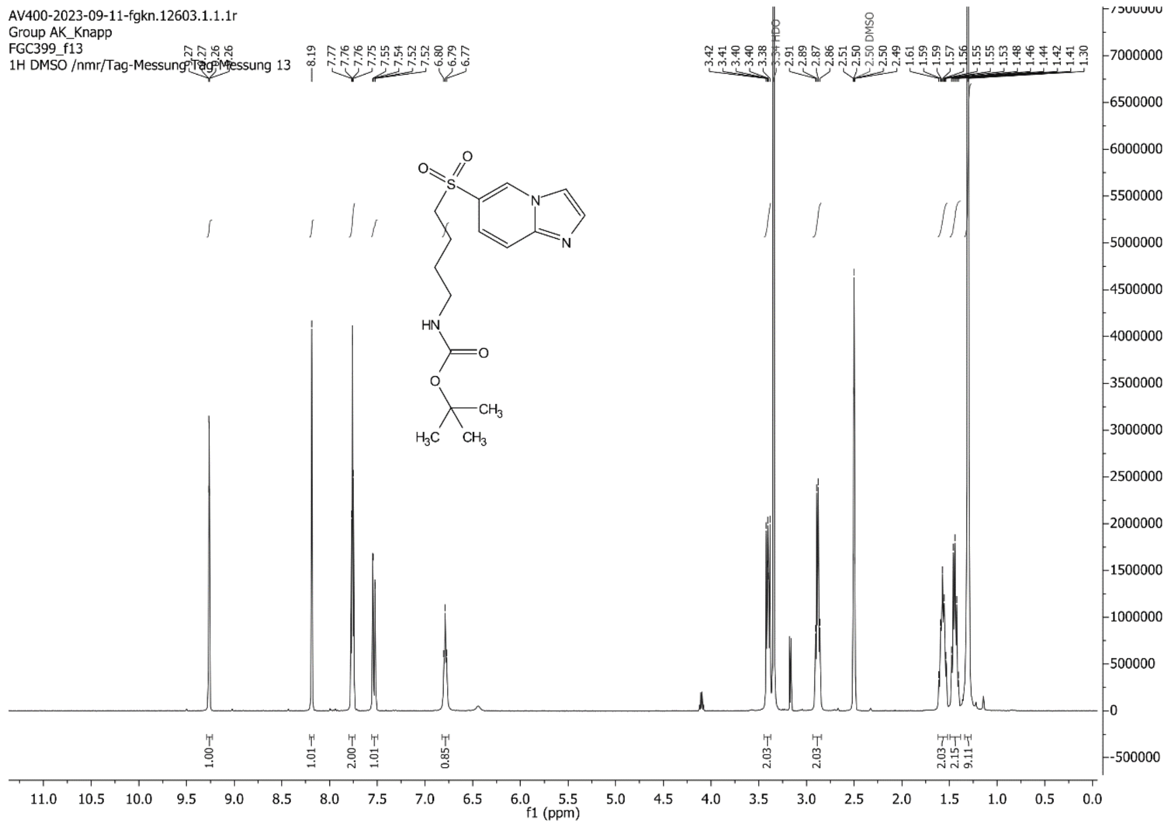
AV400-2023-09-11-fgkn.12603.1.1.1r
Group AK_Knapp
FGC399_f13
1H DMSO /nmr/Tag-Messung Tag-Messung 13
HN
O
CH3
H3C
N
f1 (ppm)

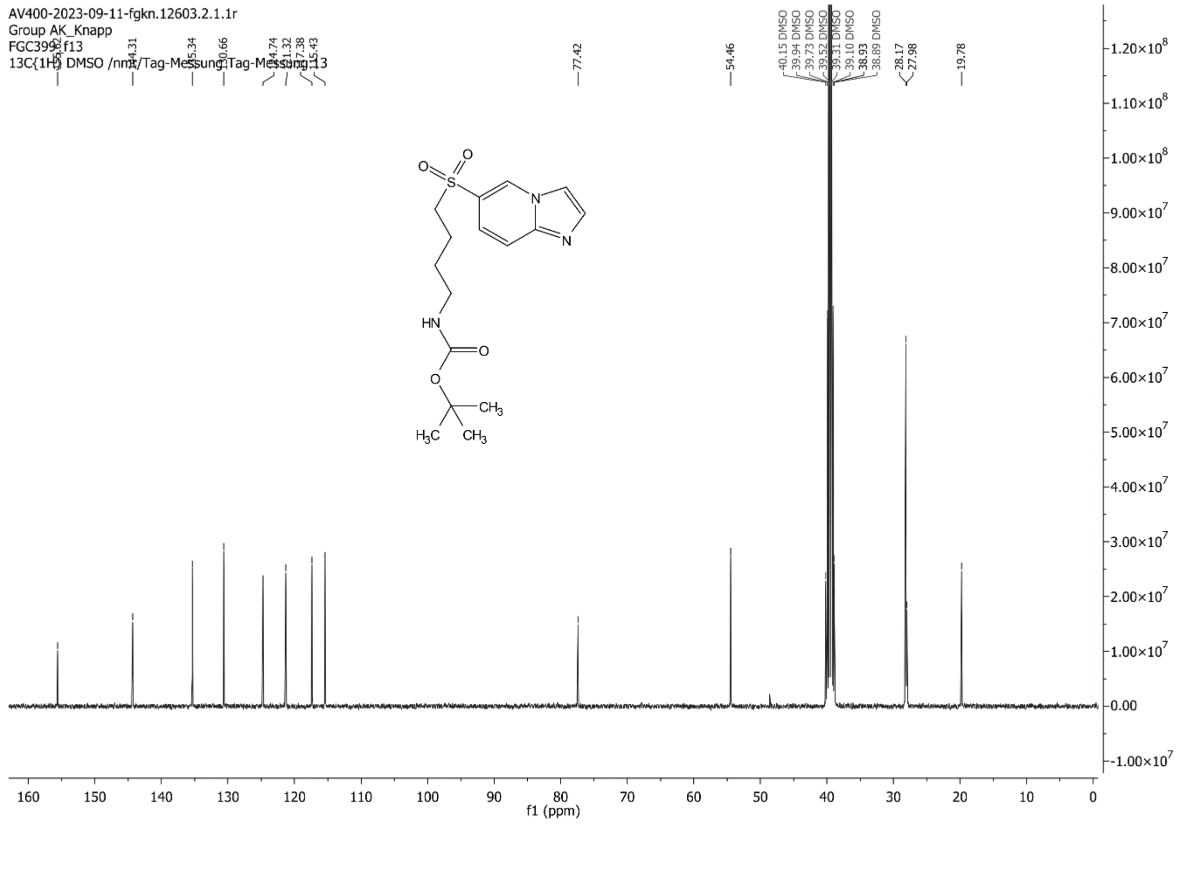
AV400-2023-09-11-fgkn.12603.2.1.1r
Group AK_Knapp
FGC399_f13
13C{1H} DMSO /nmr/Tag-Messung Tag-Messung 13
HN
O
CH3
H3C
N
f1 (ppm)

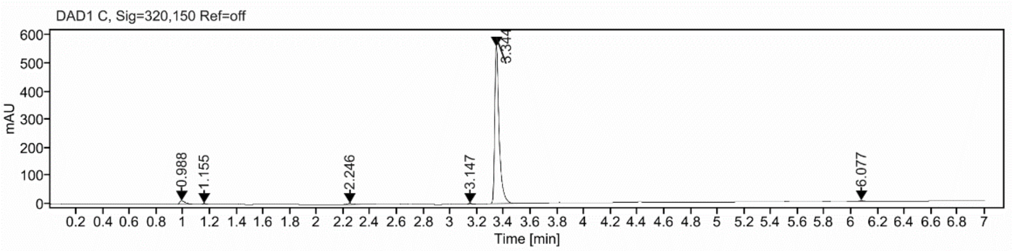

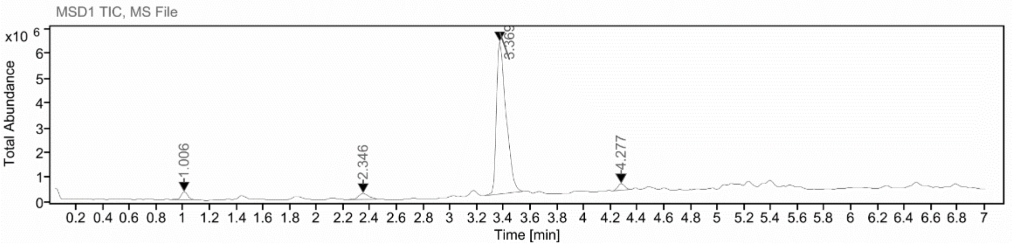

**Sample Purity**

Signal Description DAD1 C, Sig=320,150 Ref=off

| Sample Name | Name | RT | Width | Area | Area% | Height |
|---|---|---|---|---|---|---|
| FGC399_f13 | | 0.988 | 0.045 | 39.8280 | 2.88 | 12.3253 |
| FGC399_f13 | | 1.155 | 0.030 | 7.7693 | 0.56 | 2.9876 |
| FGC399_f13 | | 2.246 | 0.053 | 12.5366 | 0.91 | 3.7044 |
| FGC399_f13 | | 3.147 | 0.030 | 9.1812 | 0.66 | 4.0796 |
| FGC399_f13 | | 3.344 | 0.033 | 1306.5642 | 94.33 | 561.0756 |
| FGC399_f13 | | 6.077 | 0.040 | 9.2237 | 0.67 | 3.3784 |

Max Area% 94.330

UV Signal Purity>95% Fail

**UV Apex Spectra**

**RT:**

3.344

**Sample Name:**

FGC399_f13

**Signal Name:**

DAD1C

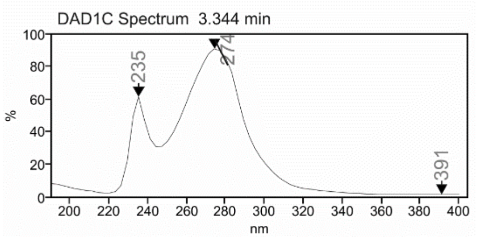

**RT:**

3.369

**Sample Name:**

FGC399_f13

**Signal Name:**

MSD1TIC

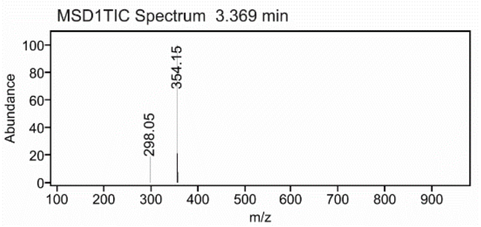

**Cmpd 68**

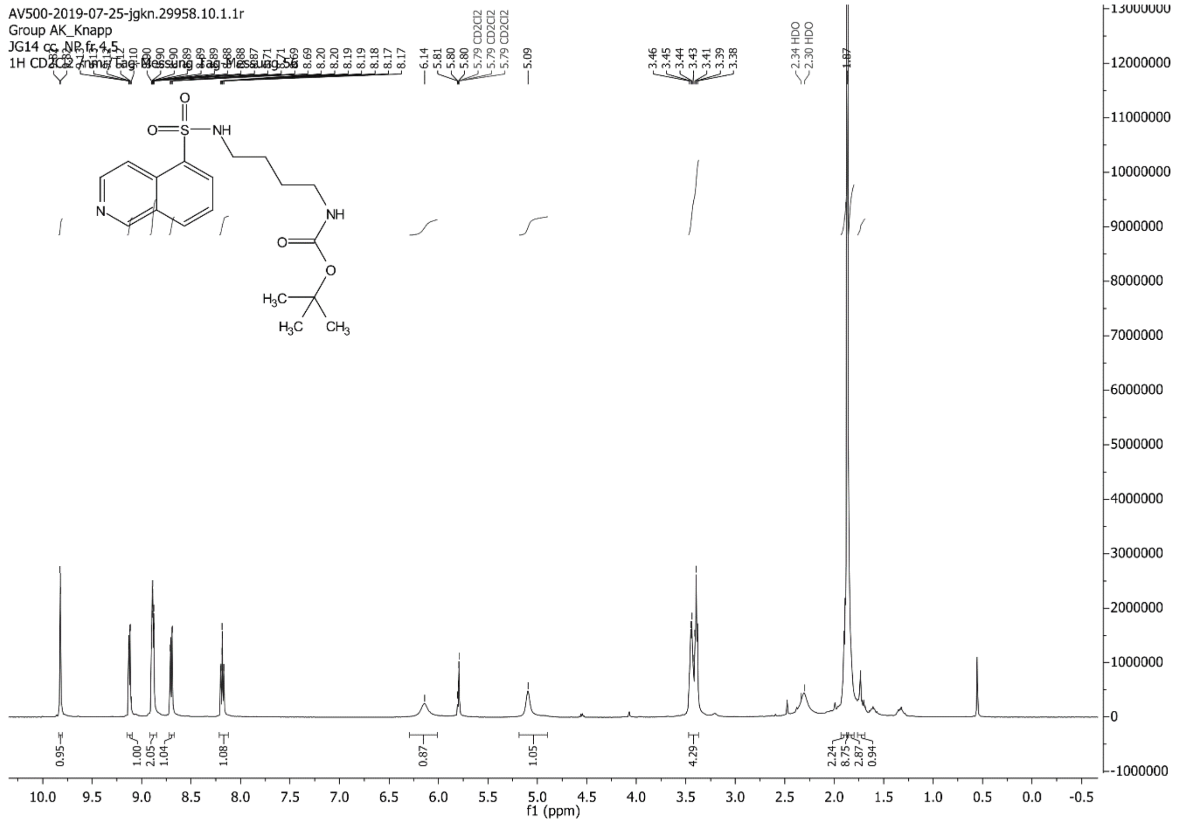

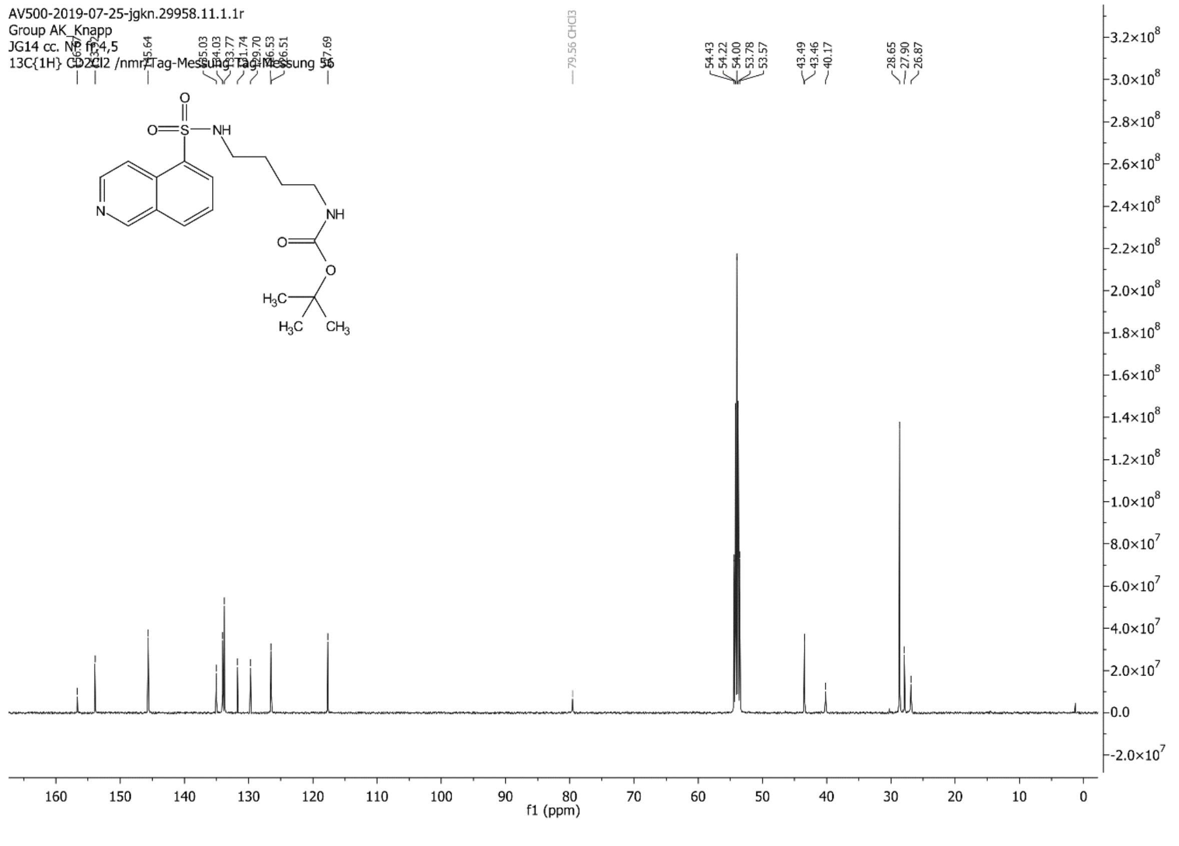

C:\Xcalibur\data\JG14-1 7/18/2019 12:22:12 PM

JG14-1 #33-44 RT: 0.56-0.75 AV: 12 SB: 10 0.05-0.21 NL: 2.42E6
T: {0,0} + c ESI !corona sid=75.00 det=1306.00 Full ms [105.00-700.00]

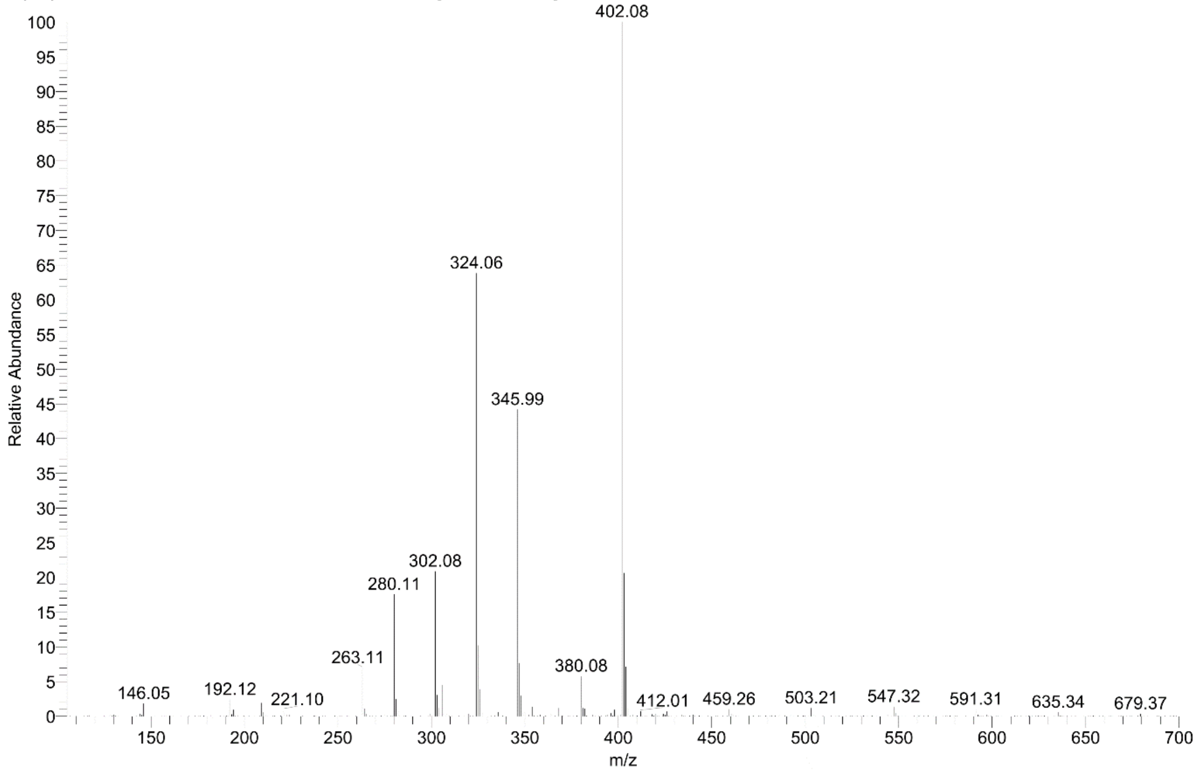

### Cmpd 70

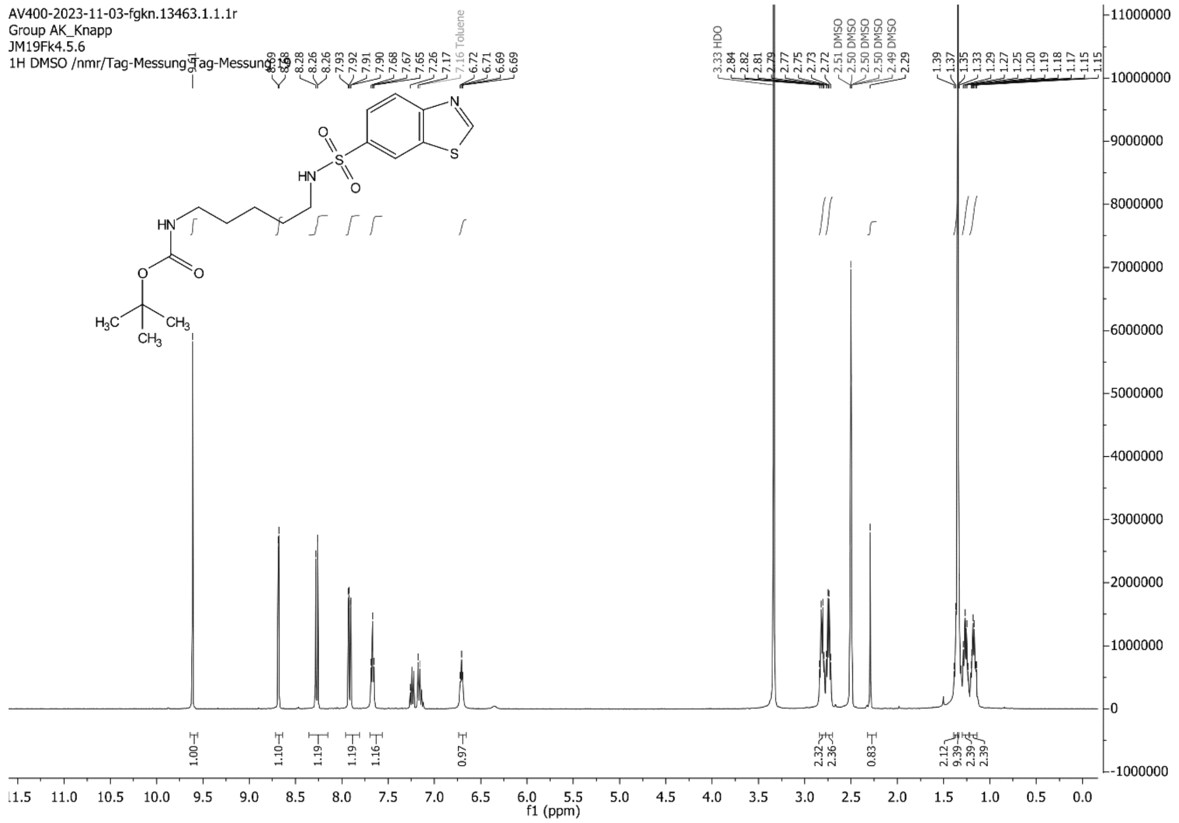
AV400-2023-11-03-fgkn.13463.1.1.1r
Group AK_Knapp
JM19Fk4.5.6
1H DMSO /nmr/Tag-Messung Tag-Messung
f1 (ppm)

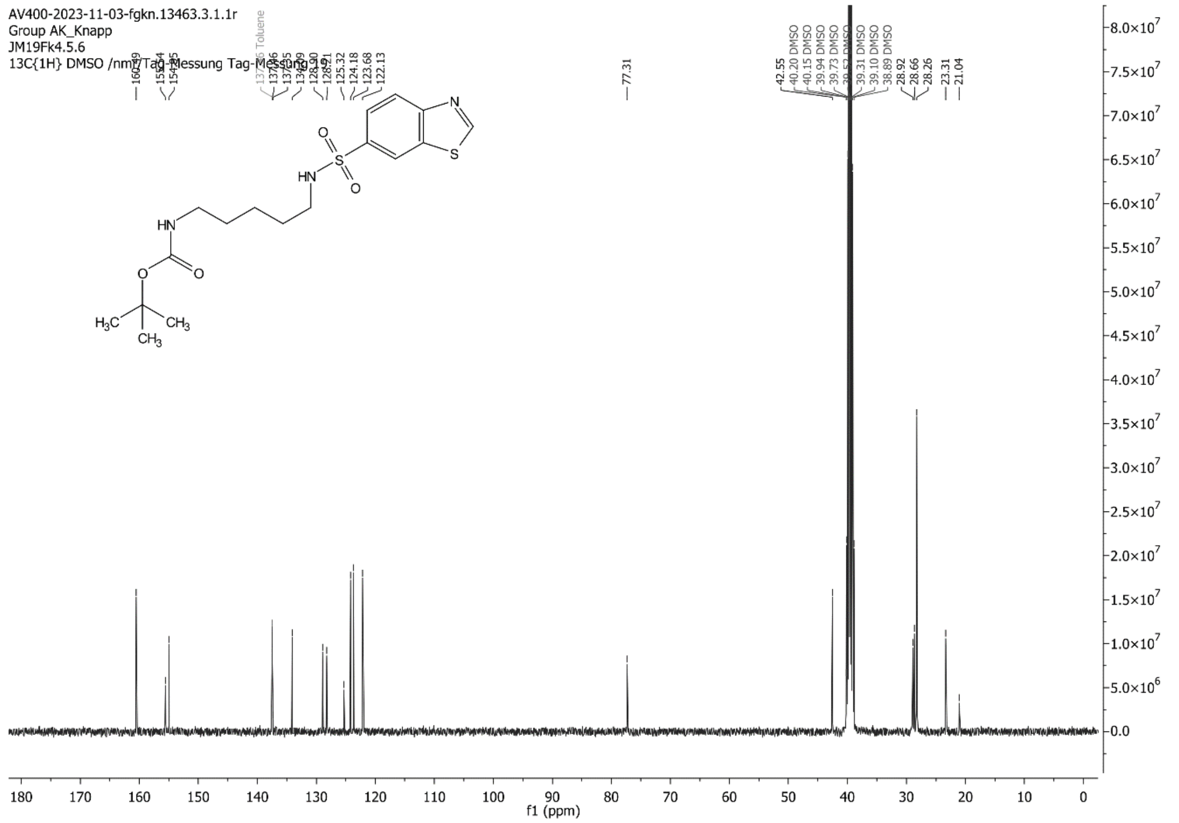
AV400-2023-11-03-fgkn.13463.3.1.1r
Group AK_Knapp
JM19Fk4.5.6
13C{1H} DMSO /nmr/Tag-Messung Tag-Messung
f1 (ppm)

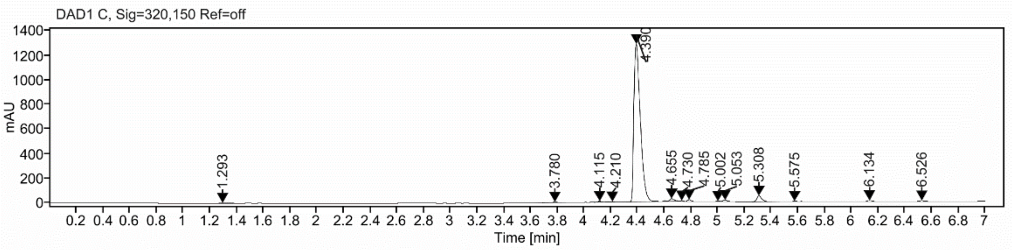

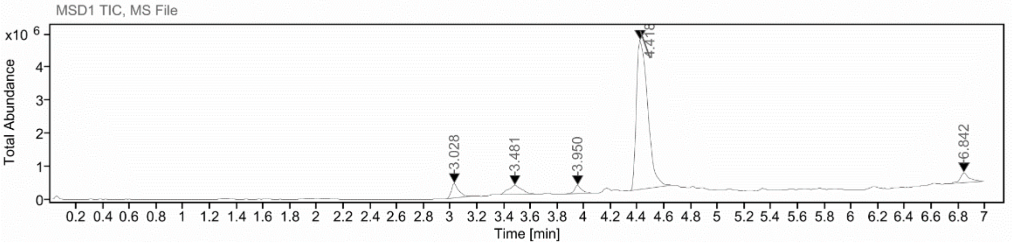

**Sample Purity**

Signal Description DAD1 C, Sig=320,150 Ref=off

| Sample Name | Name | RT | Width | Area | Area% | Height |
|---|---|---|---|---|---|---|
| FGJM19A | | 1.293 | 0.044 | 11.7689 | 0.25 | 3.6474 |
| FGJM19A | | 3.780 | 0.052 | 25.6545 | 0.55 | 5.3412 |
| FGJM19A | | 4.115 | 0.032 | 14.0655 | 0.30 | 7.2759 |
| FGJM19A | | 4.210 | 0.070 | 15.3058 | 0.33 | 4.1831 |
| FGJM19A | | 4.390 | 0.052 | 4298.2915 | 91.76 | 1283.3811 |
| FGJM19A | | 4.655 | 0.038 | 56.9764 | 1.22 | 20.5018 |
| FGJM19A | | 4.730 | 0.024 | 5.2283 | 0.11 | 3.6237 |
| FGJM19A | | 4.785 | 0.028 | 30.1596 | 0.64 | 14.5532 |
| FGJM19A | | 5.002 | 0.027 | 7.0968 | 0.15 | 4.3745 |
| FGJM19A | | 5.053 | 0.031 | 26.3982 | 0.56 | 12.2926 |
| FGJM19A | | 5.308 | 0.038 | 129.4260 | 2.76 | 48.7346 |
| FGJM19A | | 5.575 | 0.088 | 19.9836 | 0.43 | 4.2789 |
| FGJM19A | | 6.134 | 0.040 | 27.4128 | 0.59 | 9.9794 |
| FGJM19A | | 6.526 | 0.043 | 16.4251 | 0.35 | 5.8829 |

Max Area% 91.762

UV Signal Purity>95% Fail

**RT:**

4.390

**Sample Name:**

FGJM19A

**Signal Name:**

DAD1C

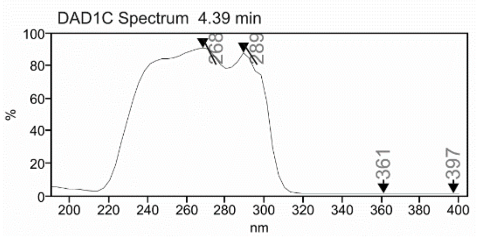

**RT:**

4.418

**Sample Name:**

FGJM19A

**Signal Name:**

MSD1TIC

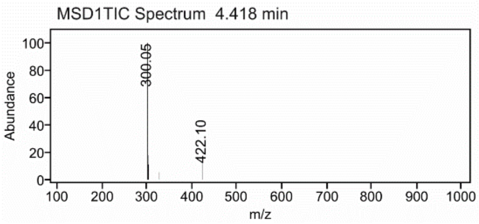

**Cmpd 71**

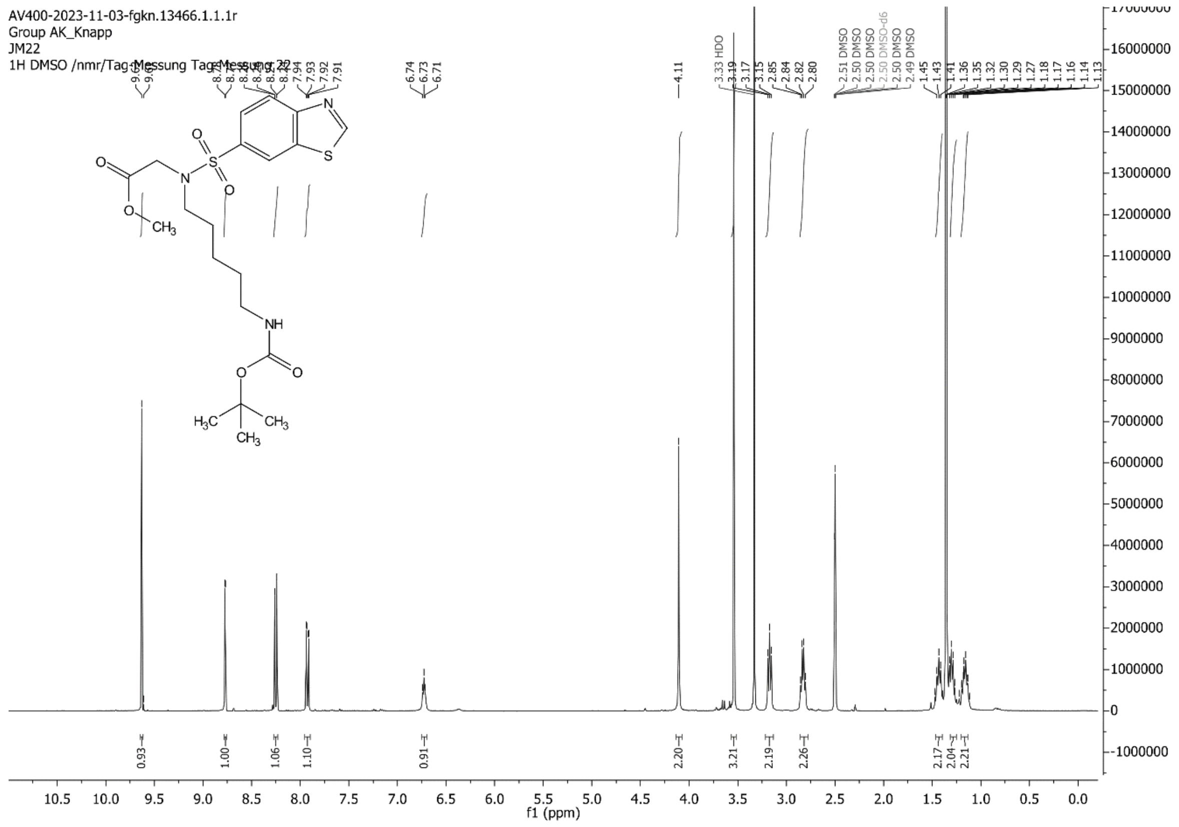
AV400-2023-11-03-fgkn.13466.1.1.1r
Group AK_Knapp
JM22
1H DMSO /nmr/Tag-Messung Tag-Messung 22
f1 (ppm)

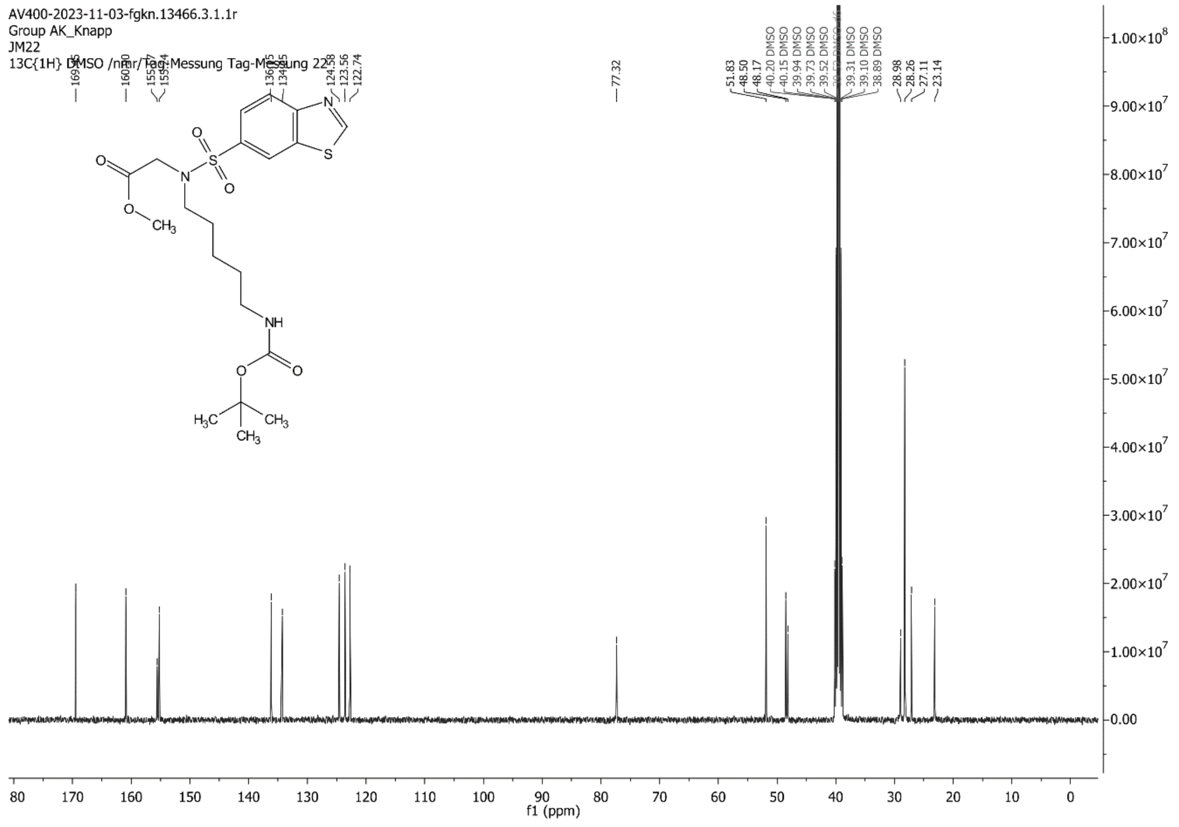
AV400-2023-11-03-fgkn.13466.3.1.1r
Group AK_Knapp
JM22
13C{1H} DMSO /nmr/Tag-Messung Tag-Messung 22
f1 (ppm)

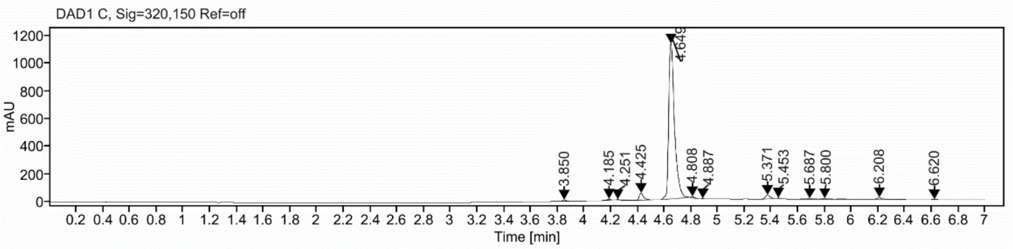

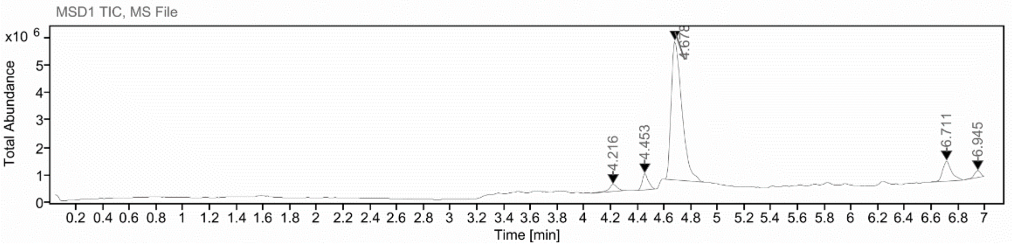

**Sample Purity**

Signal Description DAD1 C, Sig=320,150 Ref=off

| Sample Name | Name | RT | Width | Area | Area% | Height |
|---|---|---|---|---|---|---|
| FGJM22R2 | | 3.850 | 0.032 | 21.3403 | 0.57 | 10.4219 |
| FGJM22R2 | | 4.185 | 0.030 | 14.8281 | 0.40 | 8.2865 |
| FGJM22R2 | | 4.251 | 0.027 | 9.3045 | 0.25 | 4.4950 |
| FGJM22R2 | | 4.425 | 0.032 | 115.6291 | 3.09 | 51.0741 |
| FGJM22R2 | | 4.649 | 0.044 | 3392.8152 | 90.73 | 1120.2457 |
| FGJM22R2 | | 4.808 | 0.025 | 13.8270 | 0.37 | 8.3020 |
| FGJM22R2 | | 4.887 | 0.026 | 8.6800 | 0.23 | 2.2404 |
| FGJM22R2 | | 5.371 | 0.038 | 67.8904 | 1.82 | 27.9329 |
| FGJM22R2 | | 5.453 | 0.028 | 8.3818 | 0.22 | 4.7059 |
| FGJM22R2 | | 5.687 | 0.045 | 18.8720 | 0.50 | 5.2107 |
| FGJM22R2 | | 5.800 | 0.042 | 22.9293 | 0.61 | 4.7677 |
| FGJM22R2 | | 6.208 | 0.039 | 35.9986 | 0.96 | 12.7371 |
| FGJM22R2 | | 6.620 | 0.043 | 8.9683 | 0.24 | 3.2010 |

Max Area% 90.730

UV Signal Purity>95% Fail

**RT:**

4.649

**Sample Name:**

FGJM22R2

**Signal Name:**

DAD1C

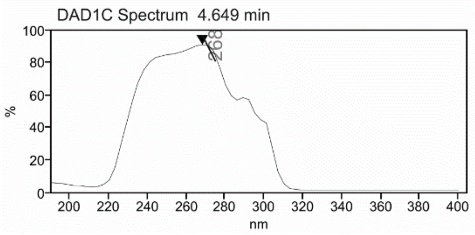

**RT:**

4.678

**Sample Name:**

FGJM22R2

**Signal Name:**

MSD1TIC

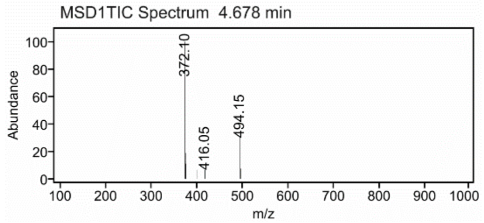

### Data collection and refinement stastitics

#### 8PVO

W O R L D W I D E PDB PROTEIN DATA BANK

### Full wwPDB X-ray Structure Validation Report (i)

Jul 20, 2023 – 02:35 pm BST

| | | |
|---|---|---|
| PDB ID | : | 8PVO |
| Title | : | Crystal structure of human Casein Kinase II subunit alpha (CK2a1) in complex with allosteric compound FG5 |
| Deposited on | : | 2023-07-18 |
| Resolution | : | 2.25 Å(reported) |

**This wwPDB validation report is for manuscript review**

For Manuscript Review

This is a Full wwPDB X-ray Structure Validation Report.

This report is produced by the wwPDB biocuration pipeline after annotation of the structure.

We welcome your comments at **
A user guide is available at
https://www.wwpdb.org/validation/2017/XrayValidationReportHelp
with specific help available everywhere you see the (i) symbol.

The types of validation reports are described at
http://www.wwpdb.org/validation/2017/FAQs#types.

The following versions of software and data (see references (i)) were used in the production of this report:

| | | |
|---|---|---|
| MolProbity | : | 4.02b-467 |
| Mogul | : | 1.8.4, CSD as541be (2020) |
| Xtriage (Phenix) | : | 1.13 |
| EDS | : | 2.34 |
| buster-report | : | 1.1.7 (2018) |
| Percentile statistics | : | 20191225.v01 (using entries in the PDB archive December 25th 2019) |
| Refmac | : | 5.8.0158 |
| CCP4 | : | 7.0.044 (Gargrove) |
| Ideal geometry (proteins) | : | Engh & Huber (2001) |

### 1 Overall quality at a glance (i)

The following experimental techniques were used to determine the structure:
*X-RAY DIFFRACTION*

The reported resolution of this entry is 2.25 Å.

Percentile scores (ranging between 0-100) for global validation metrics of the entry are shown in the following graphic. The table shows the number of entries on which the scores are based.

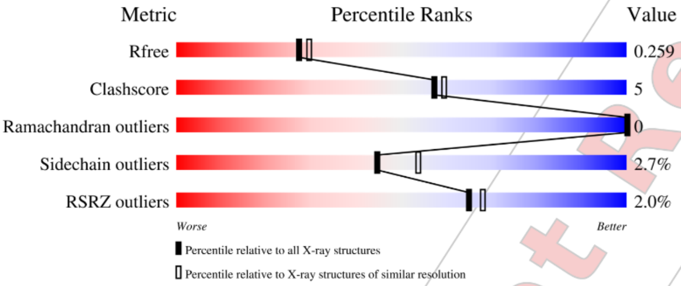

| Metric | Whole archive (#Entries) | Similar resolution (#Entries, resolution range(Å)) |
|---|---|---|
| $R_{free}$ | 130704 | 1377 (2.26-2.26) |
| Clashscore | 141614 | 1487 (2.26-2.26) |
| Ramachandran outliers | 138981 | 1449 (2.26-2.26) |
| Sidechain outliers | 138945 | 1450 (2.26-2.26) |
| RSRZ outliers | 127900 | 1356 (2.26-2.26) |

The table below summarises the geometric issues observed across the polymeric chains and their fit to the electron density. The red, orange, yellow and green segments of the lower bar indicate the fraction of residues that contain outliers for >=3, 2, 1 and 0 types of geometric quality criteria respectively. A grey segment represents the fraction of residues that are not modelled. The numeric value for each fraction is indicated below the corresponding segment, with a dot representing fractions <=5% The upper red bar (where present) indicates the fraction of residues that have poor fit to the electron density. The numeric value is given above the bar.

| Mol | Chain | Length | Quality of chain |
|---|---|---|---|
| 1 | A | 338 | 2%<br>88% 9% • • |
| 1 | B | 338 | 2%<br>84% 12% • • |

The following table lists non-polymeric compounds, carbohydrate monomers and non-standard

Ideal geometry (DNA, RNA) : Parkinson et al. (1996)
Validation Pipeline (wwPDB-VP) : 2.34

For Manuscript Review

residues in protein, DNA, RNA chains that are outliers for geometric or electron-density-fit criteria:

| Mol | Type | Chain | Res | Chirality | Geometry | Clashes | Electron density |
|---|---|---|---|---|---|---|---|
| 2 | SO4 | A | 403 | - | - | X | - |

For Manuscript Review

#### 2 Entry composition (i)

There are 6 unique types of molecules in this entry. The entry contains 5839 atoms, of which 0 are hydrogens and 0 are deuteriums.

In the tables below, the ZeroOcc column contains the number of atoms modelled with zero occupancy, the AltConf column contains the number of residues with at least one atom in alternate conformation and the Trace column contains the number of residues modelled with at most 2 atoms.

- Molecule 1 is a protein called CSNK2A1 protein.

| Mol | Chain | Residues | Atoms | | | | | ZeroOcc | AltConf | Trace |
|---|---|---|---|---|---|---|---|---|---|---|
| 1 | A | 330 | Total 2745 | C 1758 | N 479 | O 497 | S 11 | 0 | 0 | 0 |
| 1 | B | 329 | Total 2753 | C 1765 | N 479 | O 498 | S 11 | 0 | 0 | 0 |

There are 2 discrepancies between the modelled and reference sequences:

| Chain | Residue | Modelled | Actual | Comment | Reference |
|---|---|---|---|---|---|
| A | 0 | SER | - | expression tag | UNP Q5U5J2 |
| B | 0 | SER | - | expression tag | UNP Q5U5J2 |

- Molecule 2 is SULFATE ION (three-letter code: SO4) (formula: $O_4S$).

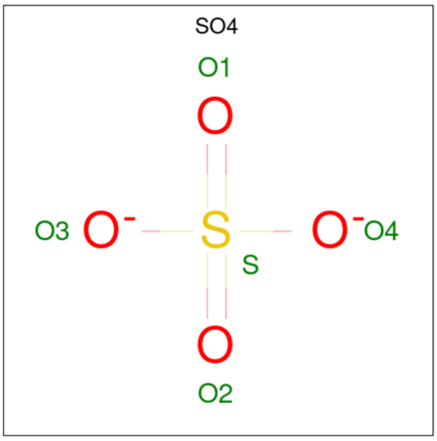

| Mol | Chain | Residues | Atoms | | | ZeroOcc | AltConf |
|---|---|---|---|---|---|---|---|
| 2 | A | 1 | Total 5 | O 4 | S 1 | 0 | 0 |

*Continued on next page...*

For Manuscript Review

*Continued from previous page...*

| Mol | Chain | Residues | Atoms | ZeroOcc | AltConf |
|---|---|---|---|---|---|
| 2 | A | 1 | Total O S<br>5 4 1 | 0 | 0 |
| 2 | A | 1 | Total O S<br>5 4 1 | 0 | 0 |
| 2 | A | 1 | Total O S<br>5 4 1 | 0 | 0 |
| 2 | A | 1 | Total O S<br>5 4 1 | 0 | 0 |
| 2 | A | 1 | Total O S<br>5 4 1 | 0 | 0 |
| 2 | A | 1 | Total O S<br>5 4 1 | 0 | 0 |
| 2 | A | 1 | Total O S<br>5 4 1 | 0 | 0 |
| 2 | A | 1 | Total O S<br>5 4 1 | 0 | 0 |
| 2 | B | 1 | Total O S<br>5 4 1 | 0 | 0 |
| 2 | B | 1 | Total O S<br>5 4 1 | 0 | 0 |
| 2 | B | 1 | Total O S<br>5 4 1 | 0 | 0 |
| 2 | B | 1 | Total O S<br>5 4 1 | 0 | 0 |
| 2 | B | 1 | Total O S<br>5 4 1 | 0 | 0 |
| 2 | B | 1 | Total O S<br>5 4 1 | 0 | 0 |
| 2 | B | 1 | Total O S<br>5 4 1 | 0 | 0 |
| 2 | B | 1 | Total O S<br>5 4 1 | 0 | 0 |
| 2 | B | 1 | Total O S<br>5 4 1 | 0 | 0 |
| 2 | B | 1 | Total O S<br>5 4 1 | 0 | 0 |
| 2 | B | 1 | Total O S<br>5 4 1 | 0 | 0 |
| 2 | B | 1 | Total O S<br>5 4 1 | 0 | 0 |

- Molecule 3 is 1,2-ETHANEDIOL (three-letter code: EDO) (formula: $C_2H_6O_2$).

For Manuscript Review

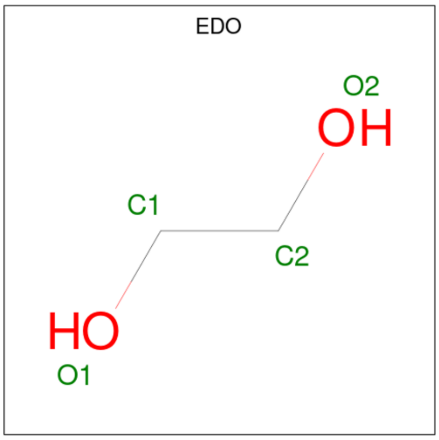

| Mol | Chain | Residues | Atoms | ZeroOcc | AltConf |
|---|---|---|---|---|---|
| 3 | A | 1 | Total C O<br>4 2 2 | 0 | 0 |
| 3 | A | 1 | Total C O<br>4 2 2 | 0 | 0 |
| 3 | A | 1 | Total C O<br>4 2 2 | 0 | 0 |
| 3 | B | 1 | Total C O<br>4 2 2 | 0 | 0 |

- Molecule 4 is 2-[1-(1,3-benzothiazol-6-ylsulfonyl)piperidin-4-yl]- {N}-[(3-chloranyl-4-pheny l-phenyl)methyl]ethanamine (three-letter code: FW3) (formula: $C_{27}H_{28}ClN_3O_2S_2$) (labeled as "Ligand of Interest" by depositor).

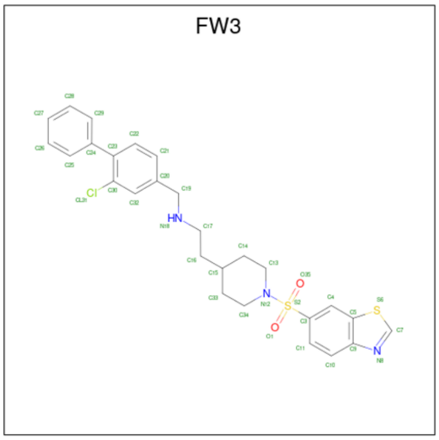

For Manuscript Review

| Mol | Chain | Residues | Atoms | | | | | | ZeroOcc | AltConf |
|---|---|---|---|---|---|---|---|---|---|---|
| 4 | A | 1 | Total<br>35 | C<br>27 | Cl<br>1 | N<br>3 | O<br>2 | S<br>2 | 0 | 0 |
| 4 | B | 1 | Total<br>35 | C<br>27 | Cl<br>1 | N<br>3 | O<br>2 | S<br>2 | 0 | 0 |

- Molecule 5 is SODIUM ION (three-letter code: NA) (formula: Na).

| Mol | Chain | Residues | Atoms | | ZeroOcc | AltConf |
|---|---|---|---|---|---|---|
| 5 | A | 1 | Total<br>1 | Na<br>1 | 0 | 0 |
| 5 | B | 1 | Total<br>1 | Na<br>1 | 0 | 0 |

- Molecule 6 is water.

| Mol | Chain | Residues | Atoms | | ZeroOcc | AltConf |
|---|---|---|---|---|---|---|
| 6 | A | 77 | Total<br>77 | O<br>77 | 0 | 0 |
| 6 | B | 71 | Total<br>71 | O<br>71 | 0 | 0 |

For Manuscript Review

There are no Ramachandran outliers to report.

##### 5.3.2 Protein sidechains (i)

In the following table, the Percentiles column shows the percent sidechain outliers of the chain as a percentile score with respect to all X-ray entries followed by that with respect to entries of similar resolution.

The Analysed column shows the number of residues for which the sidechain conformation was analysed, and the total number of residues.

| Mol | Chain | Analysed | Rotameric | Outliers | Percentiles | |
|---|---|---|---|---|---|---|
| 1 | A | 294/310 (95%) | 287 (98%) | 7 (2%) | 49 | 58 |
| 1 | B | 296/310 (96%) | 287 (97%) | 9 (3%) | 41 | 50 |
| All | All | 590/620 (95%) | 574 (97%) | 16 (3%) | 44 | 54 |

All (16) residues with a non-rotameric sidechain are listed below:

| Mol | Chain | Res | Type |
|---|---|---|---|
| 1 | A | 21 | ARG |
| 1 | A | 36 | GLN |
| 1 | A | 62 | ASN |
| 1 | A | 153 | MET |
| 1 | A | 180 | GLU |
| 1 | A | 258 | ILE |
| 1 | A | 273 | LEU |
| 1 | B | 8 | ARG |
| 1 | B | 15 | VAL |
| 1 | B | 36 | GLN |
| 1 | B | 120 | ASP |
| 1 | B | 153 | MET |
| 1 | B | 170 | LYS |
| 1 | B | 244 | ARG |
| 1 | B | 258 | ILE |
| 1 | B | 259 | ASP |

Sometimes sidechains can be flipped to improve hydrogen bonding and reduce clashes. All (12) such sidechains are listed below:

| Mol | Chain | Res | Type |
|---|---|---|---|
| 1 | A | 36 | GLN |
| 1 | A | 115 | HIS |
| 1 | A | 262 | ASN |

*Continued on next page...*

For Manuscript Review

### 3 Residue-property plots (i)

These plots are drawn for all protein, RNA, DNA and oligosaccharide chains in the entry. The first graphic for a chain summarises the proportions of the various outlier classes displayed in the second graphic. The second graphic shows the sequence view annotated by issues in geometry and electron density. Residues are color-coded according to the number of geometric quality criteria for which they contain at least one outlier: green = 0, yellow = 1, orange = 2 and red = 3 or more. A red dot above a residue indicates a poor fit to the electron density (RSRZ > 2). Stretches of 2 or more consecutive residues without any outlier are shown as a green connector. Residues present in the sample, but not in the model, are shown in grey.

• Molecule 1: CSNK2A1 protein

Chain A:

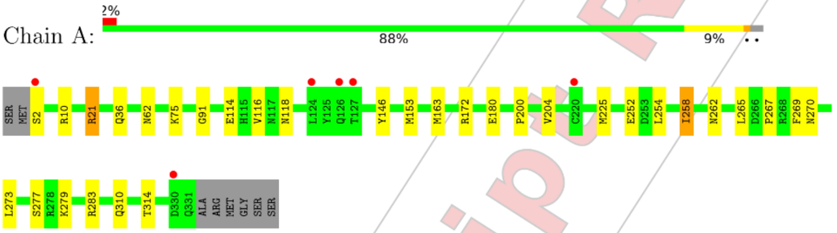

• Molecule 1: CSNK2A1 protein

Chain B:

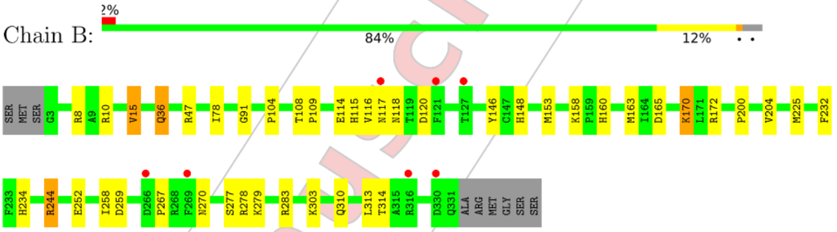

For Manuscript Review

#### 4 Data and refinement statistics (i)

| Property | Value | Source |
|---|---|---|
| Space group | P 43 21 2 | Depositor |
| Cell constants<br>a, b, c, $\alpha$, $\beta$, $\gamma$ | 127.65Å 127.65Å 123.90Å<br>90.00° 90.00° 90.00° | Depositor |
| Resolution (Å) | 45.17 – 2.25<br>45.13 – 2.25 | Depositor<br>EDS |
| % Data completeness<br>(in resolution range) | 100.0 (45.17-2.25)<br>100.0 (45.13-2.25) | Depositor<br>EDS |
| $R_{merge}$ | (Not available) | Depositor |
| $R_{sym}$ | (Not available) | Depositor |
| $< I/\sigma(I) >$ [1] | 1.88 (at 2.24Å) | Xtriage |
| Refinement program | REFMAC 5.8.0352 | Depositor |
| R, $R_{free}$ | 0.208 , 0.253<br>0.214 , 0.259 | Depositor<br>DCC |
| $R_{free}$ test set | 2399 reflections (4.89%) | wwPDB-VP |
| Wilson B-factor (Å$^2$) | 49.2 | Xtriage |
| Anisotropy | 0.452 | Xtriage |
| Bulk solvent $k_{sol}$(e/Å$^3$), $B_{sol}$(Å$^2$) | 0.36 , 40.4 | EDS |
| L-test for twinning[2] | $< \|L\| >$ = 0.50, $< L^2 >$ = 0.34 | Xtriage |
| Estimated twinning fraction | 0.000 for -h,-l,-k<br>0.000 for l,-k,h | Xtriage |
| $F_o$,$F_c$ correlation | 0.95 | EDS |
| Total number of atoms | 5839 | wwPDB-VP |
| Average B, all atoms (Å$^2$) | 57.0 | wwPDB-VP |

Xtriage's analysis on translational NCS is as follows: *The analyses of the Patterson function reveals a significant off-origin peak that is 24.03 % of the origin peak, indicating pseudo-translational symmetry. The chance of finding a peak of this or larger height randomly in a structure without pseudo-translational symmetry is equal to 4.1511e-03. The detected translational NCS is most likely also responsible for the elevated intensity ratio.*

[1] Intensities estimated from amplitudes.
[2] Theoretical values of $< |L| >$, $< L^2 >$ for acentric reflections are 0.5, 0.333 respectively for untwinned datasets, and 0.375, 0.2 for perfectly twinned datasets.

For Manuscript Review

### 5 Model quality (i)

#### 5.1 Standard geometry (i)

Bond lengths and bond angles in the following residue types are not validated in this section: NA, SO4, FW3, EDO

The Z score for a bond length (or angle) is the number of standard deviations the observed value is removed from the expected value. A bond length (or angle) with $|Z| > 5$ is considered an outlier worth inspection. RMSZ is the root-mean-square of all Z scores of the bond lengths (or angles).

| Mol | Chain | Bond lengths | | Bond angles | |
|---|---|---|---|---|---|
| | | RMSZ | #\|Z\| >5 | RMSZ | #\|Z\| >5 |
| 1 | A | 0.43 | 0/2818 | 0.79 | 0/3818 |
| 1 | B | 0.42 | 0/2827 | 0.79 | 0/3829 |
| All | All | 0.43 | 0/5645 | 0.79 | 0/7647 |

There are no bond length outliers.

There are no bond angle outliers.

There are no chirality outliers.

There are no planarity outliers.

#### 5.2 Too-close contacts (i)

In the following table, the Non-H and H(model) columns list the number of non-hydrogen atoms and hydrogen atoms in the chain respectively. The H(added) column lists the number of hydrogen atoms added and optimized by MolProbity. The Clashes column lists the number of clashes within the asymmetric unit, whereas Symm-Clashes lists symmetry-related clashes.

| Mol | Chain | Non-H | H(model) | H(added) | Clashes | Symm-Clashes |
|---|---|---|---|---|---|---|
| 1 | A | 2745 | 0 | 2652 | 25 | 0 |
| 1 | B | 2753 | 0 | 2666 | 30 | 0 |
| 2 | A | 45 | 0 | 0 | 2 | 0 |
| 2 | B | 60 | 0 | 0 | 2 | 0 |
| 3 | A | 12 | 0 | 18 | 0 | 0 |
| 3 | B | 4 | 0 | 6 | 1 | 0 |
| 4 | A | 35 | 0 | 0 | 3 | 0 |
| 4 | B | 35 | 0 | 0 | 3 | 0 |
| 5 | A | 1 | 0 | 0 | 0 | 0 |
| 5 | B | 1 | 0 | 0 | 0 | 0 |
| 6 | A | 77 | 0 | 0 | 1 | 0 |

*Continued on next page...*

For Manuscript Review

*Continued from previous page...*

| Mol | Chain | Non-H | H(model) | H(added) | Clashes | Symm-Clashes |
|---|---|---|---|---|---|---|
| 6 | B | 71 | 0 | 0 | 1 | 0 |
| All | All | 5839 | 0 | 5342 | 54 | 0 |

The all-atom clashscore is defined as the number of clashes found per 1000 atoms (including hydrogen atoms). The all-atom clashscore for this structure is 5.

All (54) close contacts within the same asymmetric unit are listed below, sorted by their clash magnitude.

| Atom-1 | Atom-2 | Interatomic distance (Å) | Clash overlap (Å) |
|---|---|---|---|
| 1:A:21:ARG:HG2 | 1:A:21:ARG:HH11 | 0.92 | 1.03 |
| 1:A:21:ARG:HG2 | 1:A:21:ARG:NH1 | 1.68 | 0.99 |
| 1:A:21:ARG:HH11 | 1:A:21:ARG:CG | 1.76 | 0.99 |
| 1:B:36:GLN:HG3 | 1:B:104:PRO:HD3 | 1.70 | 0.73 |
| 4:A:411:FW3:N8 | 6:A:501:HOH:O | 2.24 | 0.70 |
| 1:A:2:SER:HB2 | 1:A:262:ASN:HD21 | 1.57 | 0.69 |
| 1:B:10:ARG:NH1 | 1:B:314:THR:HG23 | 2.11 | 0.66 |
| 1:B:165:ASP:HB3 | 1:B:170:LYS:HG2 | 1.78 | 0.65 |
| 1:A:277:SER:HB3 | 1:B:252:GLU:HG2 | 1.80 | 0.63 |
| 1:A:267:PRO:HA | 1:A:270:ASN:ND2 | 2.16 | 0.61 |
| 1:B:148:HIS:ND1 | 3:B:414:EDO:H12 | 2.14 | 0.61 |
| 1:B:234:HIS:O | 1:B:244:ARG:NH1 | 2.34 | 0.60 |
| 1:A:225:MET:HG3 | 4:A:411:FW3:C26 | 2.34 | 0.57 |
| 1:A:10:ARG:HD2 | 1:A:310:GLN:O | 2.09 | 0.53 |
| 1:B:232:PHE:O | 2:B:410:SO4:O4 | 2.27 | 0.53 |
| 1:A:116:VAL:HG21 | 1:A:172:ARG:HG3 | 1.93 | 0.51 |
| 1:A:10:ARG:NH1 | 1:A:314:THR:HG23 | 2.26 | 0.51 |
| 1:B:278:ARG:NH1 | 2:B:409:SO4:O4 | 2.41 | 0.50 |
| 1:B:10:ARG:HD2 | 1:B:310:GLN:O | 2.12 | 0.50 |
| 1:B:78:ILE:HD13 | 1:B:109:PRO:HG2 | 1.93 | 0.50 |
| 1:B:118:ASN:HB2 | 1:B:163:MET:CE | 2.41 | 0.50 |
| 1:A:75:LYS:HE3 | 2:A:403:SO4:O1 | 2.13 | 0.49 |
| 1:A:252:GLU:HG2 | 1:B:277:SER:HB3 | 1.93 | 0.49 |
| 1:B:10:ARG:HH12 | 1:B:314:THR:HG23 | 1.76 | 0.49 |
| 1:A:75:LYS:HE2 | 2:A:403:SO4:O4 | 2.13 | 0.48 |
| 1:B:116:VAL:HG21 | 1:B:172:ARG:HG3 | 1.96 | 0.48 |
| 1:B:91:GLY:HA3 | 1:B:146:TYR:CE2 | 2.50 | 0.47 |
| 1:B:225:MET:HG2 | 4:B:413:FW3:C28 | 2.45 | 0.46 |
| 1:A:225:MET:CG | 4:A:411:FW3:C26 | 2.93 | 0.46 |
| 1:B:303:LYS:HB3 | 1:B:313:LEU:HG | 1.98 | 0.46 |
| 1:A:254:LEU:O | 1:A:258:ILE:HG23 | 2.17 | 0.45 |
| 1:A:265:LEU:HD22 | 1:A:269:PHE:CD2 | 2.51 | 0.45 |

*Continued on next page...*

For Manuscript Review

Continued from previous page...

| Atom-1 | Atom-2 | Interatomic distance (Å) | Clash overlap (Å) |
|---|---|---|---|
| 1:B:115:HIS:HE1 | 1:B:117:ASN:ND2 | 2.13 | 0.45 |
| 1:B:47:ARG:HH11 | 1:B:47:ARG:HG2 | 1.82 | 0.45 |
| 1:B:108:THR:HB | 1:B:109:PRO:CD | 2.46 | 0.45 |
| 1:B:158:LYS:HE3 | 1:B:160:HIS:HB3 | 1.99 | 0.45 |
| 1:A:279:LYS:HB3 | 1:A:283:ARG:HD3 | 1.97 | 0.44 |
| 1:B:36:GLN:CG | 1:B:104:PRO:HD3 | 2.45 | 0.44 |
| 1:B:200:PRO:O | 1:B:204:VAL:HG22 | 2.17 | 0.44 |
| 1:A:118:ASN:HB2 | 1:A:163:MET:HE2 | 1.98 | 0.44 |
| 1:A:118:ASN:HB2 | 1:A:163:MET:CE | 2.47 | 0.44 |
| 1:A:200:PRO:O | 1:A:204:VAL:HG22 | 2.18 | 0.43 |
| 1:B:225:MET:CG | 4:B:413:FW3:C28 | 2.97 | 0.43 |
| 1:B:15:VAL:HG13 | 6:B:538:HOH:O | 2.18 | 0.43 |
| 1:B:118:ASN:HB2 | 1:B:163:MET:HE3 | 2.00 | 0.43 |
| 1:A:91:GLY:HA3 | 1:A:146:TYR:CE2 | 2.55 | 0.42 |
| 1:B:279:LYS:HB3 | 1:B:283:ARG:HD3 | 2.00 | 0.42 |
| 1:B:225:MET:HG3 | 4:B:413:FW3:C27 | 2.50 | 0.41 |
| 1:A:265:LEU:HD23 | 1:A:265:LEU:HA | 1.88 | 0.40 |
| 1:A:21:ARG:NH1 | 1:A:21:ARG:CG | 2.48 | 0.40 |
| 1:A:2:SER:HB2 | 1:A:262:ASN:ND2 | 2.31 | 0.40 |
| 1:B:114:GLU:O | 1:B:114:GLU:HG3 | 2.22 | 0.40 |
| 1:A:114:GLU:O | 1:A:114:GLU:HG3 | 2.21 | 0.40 |
| 1:B:267:PRO:HA | 1:B:270:ASN:OD1 | 2.21 | 0.40 |

There are no symmetry-related clashes.

#### 5.3 Torsion angles (i)

##### 5.3.1 Protein backbone (i)

In the following table, the Percentiles column shows the percent Ramachandran outliers of the chain as a percentile score with respect to all X-ray entries followed by that with respect to entries of similar resolution.

The Analysed column shows the number of residues for which the backbone conformation was analysed, and the total number of residues.

| Mol | Chain | Analysed | Favoured | Allowed | Outliers | Percentiles | |
|---|---|---|---|---|---|---|---|
| 1 | A | 328/338 (97%) | 319 (97%) | 9 (3%) | 0 | 100 | 100 |
| 1 | B | 327/338 (97%) | 320 (98%) | 7 (2%) | 0 | 100 | 100 |
| All | All | 655/676 (97%) | 639 (98%) | 16 (2%) | 0 | 100 | 100 |

For Manuscript Review

There are no Ramachandran outliers to report.

##### 5.3.2 Protein sidechains ⓘ

In the following table, the Percentiles column shows the percent sidechain outliers of the chain as a percentile score with respect to all X-ray entries followed by that with respect to entries of similar resolution.

The Analysed column shows the number of residues for which the sidechain conformation was analysed, and the total number of residues.

| Mol | Chain | Analysed | Rotameric | Outliers | Percentiles | |
|---|---|---|---|---|---|---|
| 1 | A | 294/310 (95%) | 287 (98%) | 7 (2%) | 49 | 58 |
| 1 | B | 296/310 (96%) | 287 (97%) | 9 (3%) | 41 | 50 |
| All | All | 590/620 (95%) | 574 (97%) | 16 (3%) | 44 | 54 |

All (16) residues with a non-rotameric sidechain are listed below:

| Mol | Chain | Res | Type |
|---|---|---|---|
| 1 | A | 21 | ARG |
| 1 | A | 36 | GLN |
| 1 | A | 62 | ASN |
| 1 | A | 153 | MET |
| 1 | A | 180 | GLU |
| 1 | A | 258 | ILE |
| 1 | A | 273 | LEU |
| 1 | B | 8 | ARG |
| 1 | B | 15 | VAL |
| 1 | B | 36 | GLN |
| 1 | B | 120 | ASP |
| 1 | B | 153 | MET |
| 1 | B | 170 | LYS |
| 1 | B | 244 | ARG |
| 1 | B | 258 | ILE |
| 1 | B | 259 | ASP |

Sometimes sidechains can be flipped to improve hydrogen bonding and reduce clashes. All (12) such sidechains are listed below:

| Mol | Chain | Res | Type |
|---|---|---|---|
| 1 | A | 36 | GLN |
| 1 | A | 115 | HIS |
| 1 | A | 262 | ASN |

*Continued on next page...*

For Manuscript Review

*Continued from previous page...*

| Mol | Chain | Res | Type |
|---|---|---|---|
| 1 | A | 270 | ASN |
| 1 | A | 286 | HIS |
| 1 | A | 310 | GLN |
| 1 | A | 331 | GLN |
| 1 | B | 36 | GLN |
| 1 | B | 62 | ASN |
| 1 | B | 115 | HIS |
| 1 | B | 117 | ASN |
| 1 | B | 286 | HIS |

###### 5.3.3 RNA (i)

There are no RNA molecules in this entry.

##### 5.4 Non-standard residues in protein, DNA, RNA chains (i)

There are no non-standard protein/DNA/RNA residues in this entry.

##### 5.5 Carbohydrates (i)

There are no monosaccharides in this entry.

##### 5.6 Ligand geometry (i)

Of 29 ligands modelled in this entry, 2 are monoatomic - leaving 27 for Mogul analysis.

In the following table, the Counts columns list the number of bonds (or angles) for which Mogul statistics could be retrieved, the number of bonds (or angles) that are observed in the model and the number of bonds (or angles) that are defined in the Chemical Component Dictionary. The Link column lists molecule types, if any, to which the group is linked. The Z score for a bond length (or angle) is the number of standard deviations the observed value is removed from the expected value. A bond length (or angle) with $|Z| > 2$ is considered an outlier worth inspection. RMSZ is the root-mean-square of all Z scores of the bond lengths (or angles).

| Mol | Type | Chain | Res | Link | Bond lengths | | | Bond angles | | |
|---|---|---|---|---|---|---|---|---|---|---|
| | | | | | Counts | RMSZ | $\#\|Z\| > 2$ | Counts | RMSZ | $\#\|Z\| > 2$ |
| 3 | EDO | B | 414 | - | 3,3,3 | 0.16 | 0 | 2,2,2 | 0.44 | 0 |
| 2 | SO4 | B | 405 | - | 4,4,4 | 0.27 | 0 | 6,6,6 | 0.17 | 0 |
| 2 | SO4 | A | 402 | - | 4,4,4 | 0.28 | 0 | 6,6,6 | 0.13 | 0 |
| 2 | SO4 | B | 403 | - | 4,4,4 | 0.34 | 0 | 6,6,6 | 0.05 | 0 |
| 2 | SO4 | A | 405 | - | 4,4,4 | 0.28 | 0 | 6,6,6 | 0.10 | 0 |

For Manuscript Review

| Mol | Type | Chain | Res | Link | Bond lengths | | | Bond angles | | |
|---|---|---|---|---|---|---|---|---|---|---|
| | | | | | Counts | RMSZ | #\|Z\| > 2 | Counts | RMSZ | #\|Z\| > 2 |
| 2 | SO4 | B | 410 | - | 4,4,4 | 0.30 | 0 | 6,6,6 | 0.22 | 0 |
| 3 | EDO | A | 413 | - | 3,3,3 | 0.22 | 0 | 2,2,2 | 0.32 | 0 |
| 2 | SO4 | A | 409 | - | 4,4,4 | 0.34 | 0 | 6,6,6 | 0.26 | 0 |
| 2 | SO4 | B | 402 | - | 4,4,4 | 0.34 | 0 | 6,6,6 | 0.17 | 0 |
| 3 | EDO | A | 412 | - | 3,3,3 | 0.23 | 0 | 2,2,2 | 0.22 | 0 |
| 4 | FW3 | B | 413 | - | 37,39,39 | 0.55 | 0 | 48,55,55 | 0.79 | 1 (2%) |
| 3 | EDO | A | 408 | - | 3,3,3 | 0.37 | 0 | 2,2,2 | 0.68 | 0 |
| 2 | SO4 | B | 408 | - | 4,4,4 | 0.35 | 0 | 6,6,6 | 0.05 | 0 |
| 4 | FW3 | A | 411 | - | 37,39,39 | 0.53 | 0 | 48,55,55 | 0.94 | 2 (4%) |
| 2 | SO4 | B | 404 | - | 4,4,4 | 0.34 | 0 | 6,6,6 | 0.11 | 0 |
| 2 | SO4 | A | 404 | - | 4,4,4 | 0.34 | 0 | 6,6,6 | 0.08 | 0 |
| 2 | SO4 | A | 406 | - | 4,4,4 | 0.30 | 0 | 6,6,6 | 0.11 | 0 |
| 2 | SO4 | B | 411 | - | 4,4,4 | 0.30 | 0 | 6,6,6 | 0.28 | 0 |
| 2 | SO4 | B | 407 | - | 4,4,4 | 0.35 | 0 | 6,6,6 | 0.07 | 0 |
| 2 | SO4 | B | 412 | - | 4,4,4 | 0.37 | 0 | 6,6,6 | 0.20 | 0 |
| 2 | SO4 | A | 403 | - | 4,4,4 | 0.29 | 0 | 6,6,6 | 0.26 | 0 |
| 2 | SO4 | A | 407 | - | 4,4,4 | 0.33 | 0 | 6,6,6 | 0.09 | 0 |
| 2 | SO4 | B | 401 | - | 4,4,4 | 0.26 | 0 | 6,6,6 | 0.20 | 0 |
| 2 | SO4 | B | 409 | - | 4,4,4 | 0.37 | 0 | 6,6,6 | 0.13 | 0 |
| 2 | SO4 | A | 410 | - | 4,4,4 | 0.48 | 0 | 6,6,6 | 0.25 | 0 |
| 2 | SO4 | A | 401 | - | 4,4,4 | 0.22 | 0 | 6,6,6 | 0.28 | 0 |
| 2 | SO4 | B | 406 | - | 4,4,4 | 0.34 | 0 | 6,6,6 | 0.09 | 0 |

In the following table, the Chirals column lists the number of chiral outliers, the number of chiral centers analysed, the number of these observed in the model and the number defined in the Chemical Component Dictionary. Similar counts are reported in the Torsion and Rings columns. '-' means no outliers of that kind were identified.

| Mol | Type | Chain | Res | Link | Chirals | Torsions | Rings |
|---|---|---|---|---|---|---|---|
| 4 | FW3 | B | 413 | - | - | 9/23/33/33 | 0/5/5/5 |
| 3 | EDO | B | 414 | - | - | 0/1/1/1 | - |
| 3 | EDO | A | 408 | - | - | 1/1/1/1 | - |
| 4 | FW3 | A | 411 | - | - | 11/23/33/33 | 0/5/5/5 |
| 3 | EDO | A | 413 | - | - | 1/1/1/1 | - |
| 3 | EDO | A | 412 | - | - | 1/1/1/1 | - |

There are no bond length outliers.

All (3) bond angle outliers are listed below:

| Mol | Chain | Res | Type | Atoms | Z | Observed(°) | Ideal(°) |
|---|---|---|---|---|---|---|---|
| 4 | A | 411 | FW3 | C5-C9-N8 | 3.15 | 114.88 | 107.87 |

*Continued on next page...*

For Manuscript Review

*Continued from previous page...*

| Mol | Chain | Res | Type | Atoms | Z | Observed(°) | Ideal(°) |
|---|---|---|---|---|---|---|---|
| 4 | B | 413 | FW3 | C5-C9-N8 | 3.13 | 114.82 | 107.87 |
| 4 | A | 411 | FW3 | C3-S2-N12 | -2.13 | 104.77 | 107.30 |

There are no chirality outliers.

All (23) torsion outliers are listed below:

| Mol | Chain | Res | Type | Atoms |
|---|---|---|---|---|
| 4 | A | 411 | FW3 | C30-C23-C24-C25 |
| 4 | A | 411 | FW3 | C30-C23-C24-C29 |
| 4 | B | 413 | FW3 | C30-C23-C24-C25 |
| 4 | B | 413 | FW3 | C30-C23-C24-C29 |
| 4 | B | 413 | FW3 | C34-N12-S2-O35 |
| 4 | A | 411 | FW3 | C13-N12-S2-O1 |
| 4 | A | 411 | FW3 | C13-N12-S2-O35 |
| 4 | B | 413 | FW3 | C34-N12-S2-C3 |
| 4 | A | 411 | FW3 | C13-N12-S2-C3 |
| 4 | B | 413 | FW3 | C34-N12-S2-O1 |
| 3 | A | 408 | EDO | O1-C1-C2-O2 |
| 3 | A | 412 | EDO | O1-C1-C2-O2 |
| 4 | B | 413 | FW3 | C13-N12-S2-O35 |
| 4 | A | 411 | FW3 | C34-N12-S2-O1 |
| 4 | A | 411 | FW3 | C34-N12-S2-O35 |
| 4 | B | 413 | FW3 | C13-N12-S2-O1 |
| 4 | A | 411 | FW3 | C33-C15-C16-C17 |
| 3 | A | 413 | EDO | O1-C1-C2-O2 |
| 4 | B | 413 | FW3 | C13-N12-S2-C3 |
| 4 | A | 411 | FW3 | C34-N12-S2-C3 |
| 4 | A | 411 | FW3 | C22-C23-C24-C25 |
| 4 | A | 411 | FW3 | C22-C23-C24-C29 |
| 4 | B | 413 | FW3 | C22-C23-C24-C29 |

There are no ring outliers.

6 monomers are involved in 11 short contacts:

| Mol | Chain | Res | Type | Clashes | Symm-Clashes |
|---|---|---|---|---|---|
| 3 | B | 414 | EDO | 1 | 0 |
| 2 | B | 410 | SO4 | 1 | 0 |
| 4 | B | 413 | FW3 | 3 | 0 |
| 4 | A | 411 | FW3 | 3 | 0 |
| 2 | A | 403 | SO4 | 2 | 0 |
| 2 | B | 409 | SO4 | 1 | 0 |

For Manuscript Review

The following is a two-dimensional graphical depiction of Mogul quality analysis of bond lengths, bond angles, torsion angles, and ring geometry for all instances of the Ligand of Interest. In addition, ligands with molecular weight > 250 and outliers as shown on the validation Tables will also be included. For torsion angles, if less then 5% of the Mogul distribution of torsion angles is within 10 degrees of the torsion angle in question, then that torsion angle is considered an outlier. Any bond that is central to one or more torsion angles identified as an outlier by Mogul will be highlighted in the graph. For rings, the root-mean-square deviation (RMSD) between the ring in question and similar rings identified by Mogul is calculated over all ring torsion angles. If the average RMSD is greater than 60 degrees and the minimal RMSD between the ring in question and any Mogul-identified rings is also greater than 60 degrees, then that ring is considered an outlier. The outliers are highlighted in purple. The color gray indicates Mogul did not find sufficient equivalents in the CSD to analyse the geometry.

**Ligand FW3 B 413**

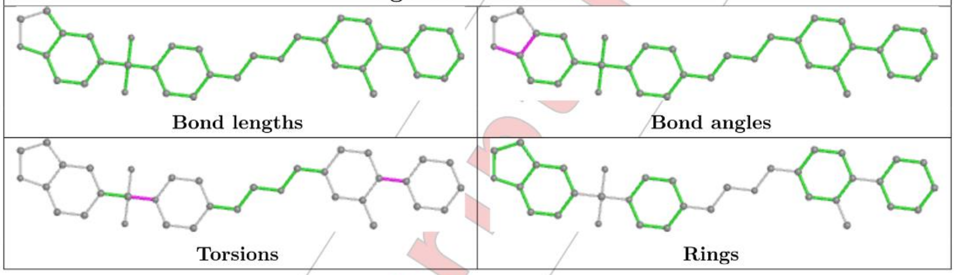

**Ligand FW3 A 411**

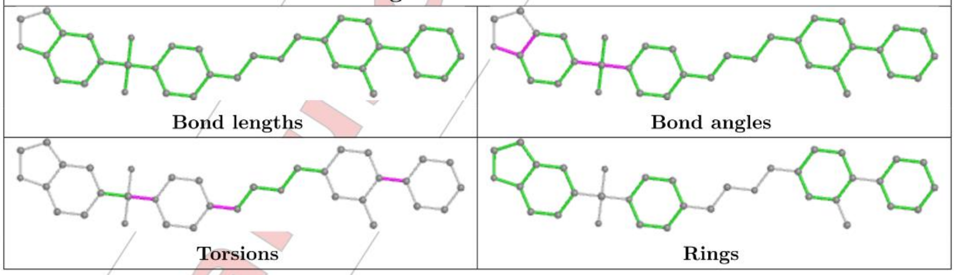

#### 5.7 Other polymers (i)

There are no such residues in this entry.

#### 5.8 Polymer linkage issues (i)

There are no chain breaks in this entry.

For Manuscript Review

### 6 Fit of model and data (i)

#### 6.1 Protein, DNA and RNA chains (i)

In the following table, the column labelled '#RSRZ> 2' contains the number (and percentage) of RSRZ outliers, followed by percent RSRZ outliers for the chain as percentile scores relative to all X-ray entries and entries of similar resolution. The OWAB column contains the minimum, median, $95^{th}$ percentile and maximum values of the occupancy-weighted average B-factor per residue. The column labelled 'Q< 0.9' lists the number of (and percentage) of residues with an average occupancy less than 0.9.

| Mol | Chain | Analysed | <RSRZ> | #RSRZ>2 | | | OWAB(Å$^2$) | Q<0.9 |
|---|---|---|---|---|---|---|---|---|
| 1 | A | 330/338 (97%) | 0.07 | 6 (1%) | 68 | 71 | 36, 53, 87, 110 | 0 |
| 1 | B | 329/338 (97%) | 0.08 | 7 (2%) | 63 | 66 | 39, 53, 91, 113 | 0 |
| All | All | 659/676 (97%) | 0.08 | 13 (1%) | 65 | 68 | 36, 53, 89, 113 | 0 |

All (13) RSRZ outliers are listed below:

| Mol | Chain | Res | Type | RSRZ |
|---|---|---|---|---|
| 1 | A | 126 | GLN | 3.6 |
| 1 | B | 127 | THR | 3.4 |
| 1 | A | 2 | SER | 3.2 |
| 1 | A | 127 | THR | 2.9 |
| 1 | A | 124 | LEU | 2.7 |
| 1 | B | 117 | ASN | 2.6 |
| 1 | A | 220 | CYS | 2.5 |
| 1 | B | 269 | PHE | 2.5 |
| 1 | B | 330 | ASP | 2.4 |
| 1 | B | 316 | ARG | 2.3 |
| 1 | A | 330 | ASP | 2.2 |
| 1 | B | 266 | ASP | 2.2 |
| 1 | B | 121 | PHE | 2.1 |

#### 6.2 Non-standard residues in protein, DNA, RNA chains (i)

There are no non-standard protein/DNA/RNA residues in this entry.

#### 6.3 Carbohydrates (i)

There are no monosaccharides in this entry.

For Manuscript Review

#### 6.4 Ligands (i)

In the following table, the Atoms column lists the number of modelled atoms in the group and the number defined in the chemical component dictionary. The B-factors column lists the minimum, median, $95^{th}$ percentile and maximum values of B factors of atoms in the group. The column labelled 'Q< 0.9' lists the number of atoms with occupancy less than 0.9.

| Mol | Type | Chain | Res | Atoms | RSCC | RSR | B-factors($Å^2$) | Q<0.9 |
|---|---|---|---|---|---|---|---|---|
| 3 | EDO | A | 412 | 4/4 | 0.62 | 0.15 | 65,75,76,79 | 0 |
| 2 | SO4 | A | 409 | 5/5 | 0.75 | 0.20 | 70,73,107,115 | 0 |
| 3 | EDO | A | 408 | 4/4 | 0.76 | 0.33 | 64,64,72,74 | 0 |
| 2 | SO4 | B | 406 | 5/5 | 0.77 | 0.15 | 96,104,115,116 | 0 |
| 3 | EDO | A | 413 | 4/4 | 0.81 | 0.36 | 56,63,79,80 | 0 |
| 2 | SO4 | B | 408 | 5/5 | 0.82 | 0.34 | 105,120,133,134 | 0 |
| 2 | SO4 | A | 407 | 5/5 | 0.86 | 0.37 | 116,118,122,132 | 0 |
| 4 | FW3 | B | 413 | 35/35 | 0.86 | 0.21 | 51,66,119,124 | 0 |
| 2 | SO4 | A | 404 | 5/5 | 0.88 | 0.20 | 90,113,128,131 | 0 |
| 2 | SO4 | B | 410 | 5/5 | 0.89 | 0.21 | 66,69,104,107 | 0 |
| 2 | SO4 | B | 411 | 5/5 | 0.89 | 0.14 | 63,71,87,89 | 0 |
| 2 | SO4 | B | 409 | 5/5 | 0.89 | 0.33 | 107,119,130,136 | 0 |
| 2 | SO4 | B | 405 | 5/5 | 0.91 | 0.17 | 71,75,88,94 | 0 |
| 2 | SO4 | A | 406 | 5/5 | 0.91 | 0.24 | 93,107,113,114 | 0 |
| 4 | FW3 | A | 411 | 35/35 | 0.92 | 0.17 | 42,67,103,138 | 0 |
| 2 | SO4 | B | 407 | 5/5 | 0.93 | 0.14 | 75,94,102,106 | 0 |
| 2 | SO4 | B | 404 | 5/5 | 0.93 | 0.13 | 89,91,104,106 | 0 |
| 2 | SO4 | B | 403 | 5/5 | 0.94 | 0.12 | 81,83,86,94 | 0 |
| 2 | SO4 | A | 405 | 5/5 | 0.95 | 0.11 | 78,86,94,95 | 0 |
| 3 | EDO | B | 414 | 4/4 | 0.95 | 0.15 | 37,51,51,52 | 0 |
| 2 | SO4 | A | 402 | 5/5 | 0.96 | 0.10 | 67,71,78,89 | 0 |
| 2 | SO4 | B | 412 | 5/5 | 0.96 | 0.12 | 65,68,75,75 | 0 |
| 2 | SO4 | A | 410 | 5/5 | 0.97 | 0.14 | 58,62,69,72 | 0 |
| 2 | SO4 | B | 402 | 5/5 | 0.97 | 0.11 | 71,73,87,89 | 0 |
| 2 | SO4 | A | 403 | 5/5 | 0.97 | 0.13 | 60,62,67,77 | 0 |
| 5 | NA | A | 414 | 1/1 | 0.97 | 0.36 | 53,53,53,53 | 0 |
| 2 | SO4 | B | 401 | 5/5 | 0.98 | 0.10 | 53,55,70,79 | 0 |
| 2 | SO4 | A | 401 | 5/5 | 0.99 | 0.10 | 54,54,57,62 | 0 |
| 5 | NA | B | 415 | 1/1 | 0.99 | 0.28 | 57,57,57,57 | 0 |

The following is a graphical depiction of the model fit to experimental electron density of all instances of the Ligand of Interest. In addition, ligands with molecular weight > 250 and outliers as shown on the geometry validation Tables will also be included. Each fit is shown from different orientation to approximate a three-dimensional view.

For Manuscript Review

**Electron density around FW3 B 413:**

$2mF_o$-$DF_c$ (at 0.7 rmsd) in gray
$mF_o$-$DF_c$ (at 3 rmsd) in purple (negative)
and green (positive)

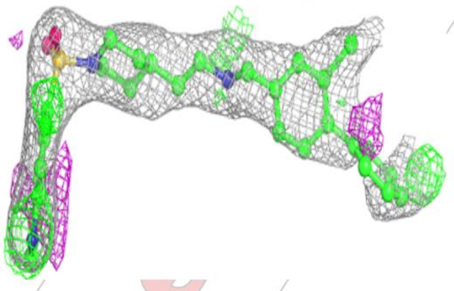

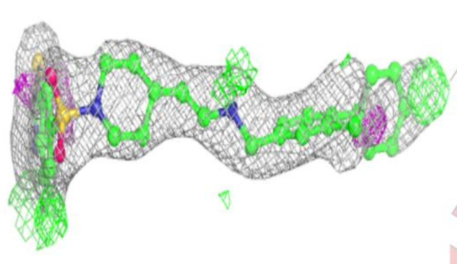

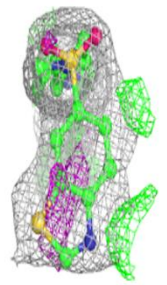

**Electron density around FW3 A 411:**

$2mF_o$-$DF_c$ (at 0.7 rmsd) in gray
$mF_o$-$DF_c$ (at 3 rmsd) in purple (negative)
and green (positive)

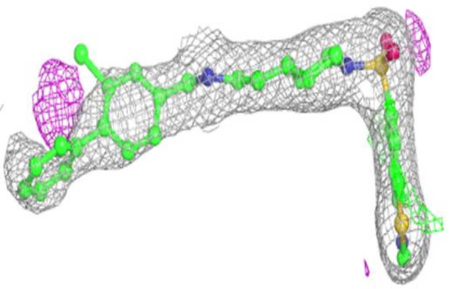

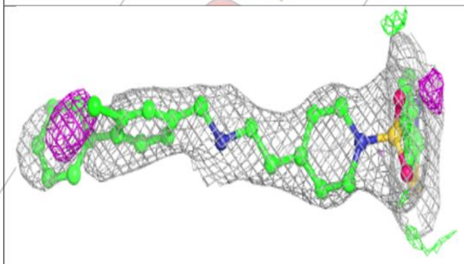

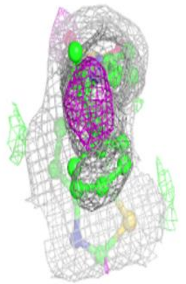

For Manuscript Review

#### 6.5 Other polymers (i)

There are no such residues in this entry.

For Manuscript Review

### Full wwPDB X-ray Structure Validation Report (i)

Jul 20, 2023 – 02:33 pm BST

| | | |
|---|---|---|
| PDB ID | : | 8PVP |
| Title | : | Crystal structure of human Casein Kinase II subunit alpha (CK2a1) in complex with allosteric compound FGJG18 |
| Deposited on | : | 2023-07-18 |
| Resolution | : | 2.60 Å(reported) |

**This wwPDB validation report is for manuscript review**

For Manuscript Review

This is a Full wwPDB X-ray Structure Validation Report.

This report is produced by the wwPDB biocuration pipeline after annotation of the structure.

We welcome your comments at **
A user guide is available at
https://www.wwpdb.org/validation/2017/XrayValidationReportHelp
with specific help available everywhere you see the (i) symbol.

The types of validation reports are described at
http://www.wwpdb.org/validation/2017/FAQs#types.

The following versions of software and data (see references (i)) were used in the production of this report:

| | | |
|---|---|---|
| MolProbity | : | 4.02b-467 |
| Mogul | : | 1.8.4, CSD as541be (2020) |
| Xtriage (Phenix) | : | 1.13 |
| EDS | : | 2.34 |
| buster-report | : | 1.1.7 (2018) |
| Percentile statistics | : | 20191225.v01 (using entries in the PDB archive December 25th 2019) |
| Refmac | : | 5.8.0158 |
| CCP4 | : | 7.0.044 (Gargrove) |
| Ideal geometry (proteins) | : | Engh & Huber (2001) |

### 1 Overall quality at a glance (i)

The following experimental techniques were used to determine the structure:
*X-RAY DIFFRACTION*

The reported resolution of this entry is 2.60 Å.

Percentile scores (ranging between 0-100) for global validation metrics of the entry are shown in the following graphic. The table shows the number of entries on which the scores are based.

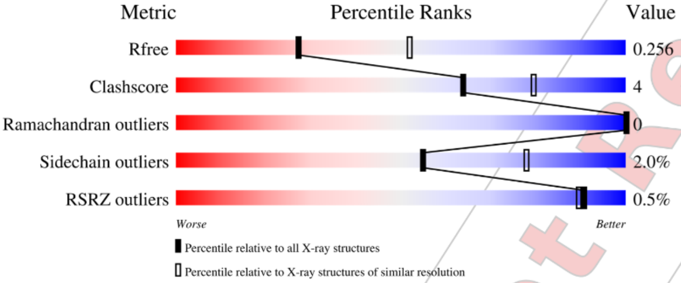

| Metric | Whole archive (#Entries) | Similar resolution (#Entries, resolution range(Å)) |
|---|---|---|
| $R_{free}$ | 130704 | 3163 (2.60-2.60) |
| Clashscore | 141614 | 3518 (2.60-2.60) |
| Ramachandran outliers | 138981 | 3455 (2.60-2.60) |
| Sidechain outliers | 138945 | 3455 (2.60-2.60) |
| RSRZ outliers | 127900 | 3104 (2.60-2.60) |

The table below summarises the geometric issues observed across the polymeric chains and their fit to the electron density. The red, orange, yellow and green segments of the lower bar indicate the fraction of residues that contain outliers for >=3, 2, 1 and 0 types of geometric quality criteria respectively. A grey segment represents the fraction of residues that are not modelled. The numeric value for each fraction is indicated below the corresponding segment, with a dot representing fractions <=5% The upper red bar (where present) indicates the fraction of residues that have poor fit to the electron density. The numeric value is given above the bar.

| Mol | Chain | Length | Quality of chain |
|---|---|---|---|
| 1 | A | 338 | 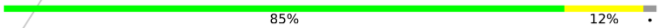  |
| 1 | B | 338 | 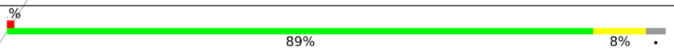  |

Ideal geometry (DNA, RNA) : Parkinson et al. (1996)
Validation Pipeline (wwPDB-VP) : 2.34

For Manuscript Review

#### 2 Entry composition (i)

There are 4 unique types of molecules in this entry. The entry contains 5499 atoms, of which 0 are hydrogens and 0 are deuteriums.

In the tables below, the ZeroOcc column contains the number of atoms modelled with zero occupancy, the AltConf column contains the number of residues with at least one atom in alternate conformation and the Trace column contains the number of residues modelled with at most 2 atoms.

- Molecule 1 is a protein called CSNK2A1 protein.

| Mol | Chain | Residues | Atoms | | | | | ZeroOcc | AltConf | Trace |
|---|---|---|---|---|---|---|---|---|---|---|
| | | | Total | C | N | O | S | | | |
| 1 | A | 330 | 2673 | 1720 | 453 | 489 | 11 | 0 | 0 | 0 |
| 1 | B | 328 | 2663 | 1710 | 460 | 482 | 11 | 0 | 0 | 0 |

There are 2 discrepancies between the modelled and reference sequences:

| Chain | Residue | Modelled | Actual | Comment | Reference |
|---|---|---|---|---|---|
| A | 0 | SER | - | expression tag | UNP Q5U5J2 |
| B | 0 | SER | - | expression tag | UNP Q5U5J2 |

- Molecule 2 is {N}-[5-[(3-chloranyl-4-phenyl-phenyl)methylamino]pentyl]isoquinoline-5 -sulfonamide (three-letter code: FWU) (formula: $C_{27}H_{28}ClN_3O_2S$) (labeled as "Ligand of Interest" by depositor).

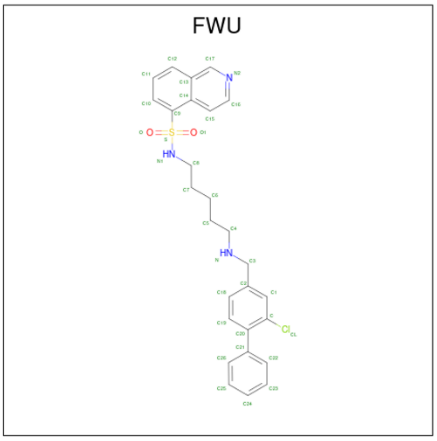

For Manuscript Review

| Mol | Chain | Residues | Atoms | ZeroOcc | AltConf |
|---|---|---|---|---|---|
| 2 | A | 1 | Total C Cl N O S<br>34 27 1 3 2 1 | 0 | 0 |
| 2 | B | 1 | Total C Cl N O S<br>34 27 1 3 2 1 | 0 | 0 |

- Molecule 3 is SULFATE ION (three-letter code: SO4) (formula: $O_4S$).

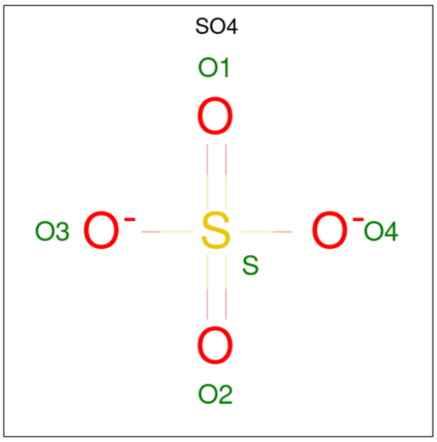

| Mol | Chain | Residues | Atoms | ZeroOcc | AltConf |
|---|---|---|---|---|---|
| 3 | A | 1 | Total O S<br>5 4 1 | 0 | 0 |
| 3 | A | 1 | Total O S<br>5 4 1 | 0 | 0 |
| 3 | A | 1 | Total O S<br>5 4 1 | 0 | 0 |
| 3 | A | 1 | Total O S<br>5 4 1 | 0 | 0 |
| 3 | A | 1 | Total O S<br>5 4 1 | 0 | 0 |
| 3 | A | 1 | Total O S<br>5 4 1 | 0 | 0 |
| 3 | B | 1 | Total O S<br>5 4 1 | 0 | 0 |
| 3 | B | 1 | Total O S<br>5 4 1 | 0 | 0 |
| 3 | B | 1 | Total O S<br>5 4 1 | 0 | 0 |
| 3 | B | 1 | Total O S<br>5 4 1 | 0 | 0 |

*Continued on next page...*

For Manuscript Review

*Continued from previous page...*

| Mol | Chain | Residues | Atoms | ZeroOcc | AltConf |
|---|---|---|---|---|---|
| 3 | B | 1 | Total O S<br>5 4 1 | 0 | 0 |
| 3 | B | 1 | Total O S<br>5 4 1 | 0 | 0 |
| 3 | B | 1 | Total O S<br>5 4 1 | 0 | 0 |

- Molecule 4 is water.

| Mol | Chain | Residues | Atoms | ZeroOcc | AltConf |
|---|---|---|---|---|---|
| 4 | A | 22 | Total O<br>22 22 | 0 | 0 |
| 4 | B | 8 | Total O<br>8 8 | 0 | 0 |

For Manuscript Review

#### 3 Residue-property plots (i)

These plots are drawn for all protein, RNA, DNA and oligosaccharide chains in the entry. The first graphic for a chain summarises the proportions of the various outlier classes displayed in the second graphic. The second graphic shows the sequence view annotated by issues in geometry and electron density. Residues are color-coded according to the number of geometric quality criteria for which they contain at least one outlier: green = 0, yellow = 1, orange = 2 and red = 3 or more. A red dot above a residue indicates a poor fit to the electron density (RSRZ > 2). Stretches of 2 or more consecutive residues without any outlier are shown as a green connector. Residues present in the sample, but not in the model, are shown in grey.

- Molecule 1: CSNK2A1 protein

Chain A:
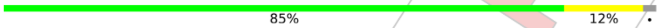

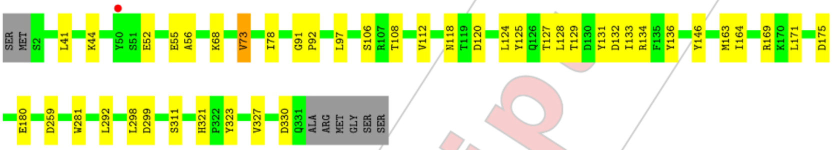

- Molecule 1: CSNK2A1 protein

Chain B:
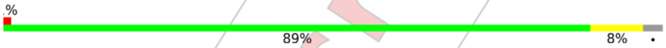

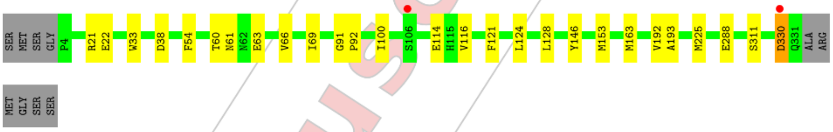

For Manuscript Review

### 4 Data and refinement statistics (i)

| Property | Value | Source |
|---|---|---|
| Space group | P 43 21 2 | Depositor |
| Cell constants<br>a, b, c, $\alpha$, $\beta$, $\gamma$ | 127.01Å 127.01Å 125.84Å<br>90.00° 90.00° 90.00° | Depositor |
| Resolution (Å) | 44.74 – 2.60<br>44.70 – 2.60 | Depositor<br>EDS |
| % Data completeness<br>(in resolution range) | 99.1 (44.74-2.60)<br>99.2 (44.70-2.60) | Depositor<br>EDS |
| $R_{merge}$ | (Not available) | Depositor |
| $R_{sym}$ | (Not available) | Depositor |
| $< I/\sigma(I) >$ [1] | 2.86 (at 2.61Å) | Xtriage |
| Refinement program | REFMAC 5.8.0352 | Depositor |
| R, $R_{free}$ | 0.222 , 0.253<br>0.229 , 0.256 | Depositor<br>DCC |
| $R_{free}$ test set | 1614 reflections (5.05%) | wwPDB-VP |
| Wilson B-factor (Å$^2$) | 49.0 | Xtriage |
| Anisotropy | 0.162 | Xtriage |
| Bulk solvent $k_{sol}$(e/Å$^3$), $B_{sol}$(Å$^2$) | 0.33 , 34.2 | EDS |
| L-test for twinning[2] | $< \|L\| >$ = 0.51, $< L^2 >$ = 0.34 | Xtriage |
| Estimated twinning fraction | 0.000 for -h,-l,-k<br>0.000 for l,-k,h | Xtriage |
| $F_o$,$F_c$ correlation | 0.93 | EDS |
| Total number of atoms | 5499 | wwPDB-VP |
| Average B, all atoms (Å$^2$) | 55.0 | wwPDB-VP |

Xtriage's analysis on translational NCS is as follows: *The analyses of the Patterson function reveals a significant off-origin peak that is 32.58 % of the origin peak, indicating pseudo-translational symmetry. The chance of finding a peak of this or larger height randomly in a structure without pseudo-translational symmetry is equal to 9.1817e-04. The detected translational NCS is most likely also responsible for the elevated intensity ratio.*

[1] Intensities estimated from amplitudes.
[2] Theoretical values of $< |L| >$, $< L^2 >$ for acentric reflections are 0.5, 0.333 respectively for untwinned datasets, and 0.375, 0.2 for perfectly twinned datasets.

For Manuscript Review

### 5 Model quality (i)

#### 5.1 Standard geometry (i)

Bond lengths and bond angles in the following residue types are not validated in this section: SO4, FWU

The Z score for a bond length (or angle) is the number of standard deviations the observed value is removed from the expected value. A bond length (or angle) with $|Z| > 5$ is considered an outlier worth inspection. RMSZ is the root-mean-square of all Z scores of the bond lengths (or angles).

| Mol | Chain | Bond lengths | | Bond angles | |
|---|---|---|---|---|---|
| | | RMSZ | $\#\|Z\| > 5$ | RMSZ | $\#\|Z\| > 5$ |
| 1 | A | 0.42 | 0/2747 | 0.78 | 0/3737 |
| 1 | B | 0.44 | 0/2735 | 0.79 | 0/3715 |
| All | All | 0.43 | 0/5482 | 0.78 | 0/7452 |

There are no bond length outliers.

There are no bond angle outliers.

There are no chirality outliers.

There are no planarity outliers.

#### 5.2 Too-close contacts (i)

In the following table, the Non-H and H(model) columns list the number of non-hydrogen atoms and hydrogen atoms in the chain respectively. The H(added) column lists the number of hydrogen atoms added and optimized by MolProbity. The Clashes column lists the number of clashes within the asymmetric unit, whereas Symm-Clashes lists symmetry-related clashes.

| Mol | Chain | Non-H | H(model) | H(added) | Clashes | Symm-Clashes |
|---|---|---|---|---|---|---|
| 1 | A | 2673 | 0 | 2507 | 25 | 0 |
| 1 | B | 2663 | 0 | 2522 | 17 | 0 |
| 2 | A | 34 | 0 | 0 | 2 | 0 |
| 2 | B | 34 | 0 | 0 | 3 | 0 |
| 3 | A | 30 | 0 | 0 | 0 | 0 |
| 3 | B | 35 | 0 | 0 | 0 | 0 |
| 4 | A | 22 | 0 | 0 | 0 | 0 |
| 4 | B | 8 | 0 | 0 | 0 | 0 |
| All | All | 5499 | 0 | 5029 | 44 | 0 |

The all-atom clashscore is defined as the number of clashes found per 1000 atoms (including hydrogen atoms). The all-atom clashscore for this structure is 4.

For Manuscript Review

All (44) close contacts within the same asymmetric unit are listed below, sorted by their clash magnitude.

| Atom-1 | Atom-2 | Interatomic distance (Å) | Clash overlap (Å) |
|---|---|---|---|
| 1:B:116:VAL:HB | 1:B:163:MET:CE | 2.32 | 0.60 |
| 1:A:118:ASN:HB2 | 1:A:163:MET:CE | 2.32 | 0.60 |
| 1:A:164:ILE:CD1 | 1:A:171:LEU:HD12 | 2.32 | 0.59 |
| 1:A:106:SER:HB3 | 1:A:108:THR:OG1 | 2.03 | 0.59 |
| 1:A:91:GLY:HA3 | 1:A:146:TYR:CE2 | 2.38 | 0.58 |
| 1:B:60:THR:OG1 | 1:B:61:ASN:OD1 | 2.20 | 0.58 |
| 1:A:118:ASN:HB2 | 1:A:163:MET:HE2 | 1.85 | 0.58 |
| 1:B:54:PHE:CE1 | 1:B:69:ILE:HD12 | 2.39 | 0.57 |
| 1:B:91:GLY:HA3 | 1:B:146:TYR:CE2 | 2.41 | 0.56 |
| 1:B:121:PHE:HA | 1:B:124:LEU:CB | 2.36 | 0.55 |
| 1:A:299:ASP:OD2 | 1:A:321:HIS:NE2 | 2.38 | 0.54 |
| 1:A:124:LEU:O | 1:A:127:THR:HG22 | 2.09 | 0.53 |
| 1:A:68:LYS:HE3 | 1:A:175:ASP:OD1 | 2.09 | 0.52 |
| 1:B:66:VAL:HG21 | 2:B:401:FWU:C16 | 2.40 | 0.51 |
| 1:A:281:TRP:HB3 | 1:A:298:LEU:HD22 | 1.93 | 0.51 |
| 1:A:134:ARG:HG2 | 1:A:323:TYR:CZ | 2.46 | 0.50 |
| 1:A:163:MET:HE3 | 2:A:401:FWU:C15 | 2.42 | 0.50 |
| 1:B:114:GLU:O | 1:B:114:GLU:HG3 | 2.12 | 0.49 |
| 1:B:33:TRP:CZ3 | 1:B:100:ILE:HG22 | 2.48 | 0.48 |
| 1:A:125:TYR:HA | 1:A:128:LEU:HD23 | 1.96 | 0.48 |
| 1:A:118:ASN:CB | 1:A:163:MET:HE2 | 2.43 | 0.47 |
| 1:A:41:LEU:HD23 | 1:A:56:ALA:HB2 | 1.96 | 0.46 |
| 1:A:281:TRP:CB | 1:A:298:LEU:HD22 | 2.45 | 0.45 |
| 1:A:44:LYS:NZ | 1:A:52:GLU:OE2 | 2.38 | 0.45 |
| 1:B:21:ARG:HG3 | 1:B:22:GLU:N | 2.32 | 0.44 |
| 1:A:129:THR:HG22 | 1:A:132:ASP:OD2 | 2.17 | 0.44 |
| 1:B:61:ASN:ND2 | 1:B:63:GLU:HB2 | 2.33 | 0.44 |
| 1:B:116:VAL:HB | 1:B:163:MET:HE1 | 1.99 | 0.44 |
| 1:A:118:ASN:CB | 1:A:163:MET:CE | 2.95 | 0.44 |
| 1:A:120:ASP:O | 1:A:124:LEU:HD22 | 2.18 | 0.43 |
| 2:A:401:FWU:C15 | 2:A:401:FWU:O1 | 2.66 | 0.43 |
| 1:B:60:THR:OG1 | 1:B:61:ASN:N | 2.52 | 0.43 |
| 1:A:133:ILE:HD12 | 1:A:292:LEU:HD22 | 2.01 | 0.42 |
| 1:A:92:PRO:HD2 | 1:A:146:TYR:CG | 2.54 | 0.41 |
| 1:A:73:VAL:CG1 | 1:A:78:ILE:HG13 | 2.50 | 0.41 |
| 1:B:92:PRO:HD2 | 1:B:146:TYR:CG | 2.55 | 0.41 |
| 1:A:136:TYR:OH | 1:A:169:ARG:HA | 2.21 | 0.41 |
| 1:B:128:LEU:HD12 | 1:B:225:MET:HE1 | 2.03 | 0.41 |
| 1:B:192:VAL:O | 1:B:193:ALA:HB3 | 2.21 | 0.41 |
| 2:B:401:FWU:C15 | 2:B:401:FWU:O1 | 2.68 | 0.40 |

*Continued on next page...*

For Manuscript Review

*Continued from previous page...*

| Atom-1 | Atom-2 | Interatomic distance (Å) | Clash overlap (Å) |
|---|---|---|---|
| 1:A:131:TYR:OH | 1:A:330:ASP:OD2 | 2.33 | 0.40 |
| 1:A:97:LEU:HD12 | 1:A:112:VAL:O | 2.21 | 0.40 |
| 1:B:225:MET:HG3 | 2:B:401:FWU:C23 | 2.52 | 0.40 |
| 1:B:330:ASP:OD1 | 1:B:330:ASP:N | 2.54 | 0.40 |

There are no symmetry-related clashes.

#### 5.3 Torsion angles (i)

##### 5.3.1 Protein backbone (i)

In the following table, the Percentiles column shows the percent Ramachandran outliers of the chain as a percentile score with respect to all X-ray entries followed by that with respect to entries of similar resolution.

The Analysed column shows the number of residues for which the backbone conformation was analysed, and the total number of residues.

| Mol | Chain | Analysed | Favoured | Allowed | Outliers | Percentiles | |
|---|---|---|---|---|---|---|---|
| 1 | A | 328/338 (97%) | 317 (97%) | 11 (3%) | 0 | 100 | 100 |
| 1 | B | 326/338 (96%) | 317 (97%) | 9 (3%) | 0 | 100 | 100 |
| All | All | 654/676 (97%) | 634 (97%) | 20 (3%) | 0 | 100 | 100 |

There are no Ramachandran outliers to report.

##### 5.3.2 Protein sidechains (i)

In the following table, the Percentiles column shows the percent sidechain outliers of the chain as a percentile score with respect to all X-ray entries followed by that with respect to entries of similar resolution.

The Analysed column shows the number of residues for which the sidechain conformation was analysed, and the total number of residues.

| Mol | Chain | Analysed | Rotameric | Outliers | Percentiles | |
|---|---|---|---|---|---|---|
| 1 | A | 277/310 (89%) | 271 (98%) | 6 (2%) | 52 | 76 |
| 1 | B | 277/310 (89%) | 272 (98%) | 5 (2%) | 59 | 80 |
| All | All | 554/620 (89%) | 543 (98%) | 11 (2%) | 55 | 78 |

For Manuscript Review

All (11) residues with a non-rotameric sidechain are listed below:

| Mol | Chain | Res | Type |
|---|---|---|---|
| 1 | A | 55 | GLU |
| 1 | A | 73 | VAL |
| 1 | A | 180 | GLU |
| 1 | A | 259 | ASP |
| 1 | A | 311 | SER |
| 1 | A | 327 | VAL |
| 1 | B | 38 | ASP |
| 1 | B | 153 | MET |
| 1 | B | 288 | GLU |
| 1 | B | 311 | SER |
| 1 | B | 330 | ASP |

Sometimes sidechains can be flipped to improve hydrogen bonding and reduce clashes. All (8) such sidechains are listed below:

| Mol | Chain | Res | Type |
|---|---|---|---|
| 1 | A | 118 | ASN |
| 1 | A | 262 | ASN |
| 1 | A | 291 | HIS |
| 1 | B | 58 | ASN |
| 1 | B | 61 | ASN |
| 1 | B | 118 | ASN |
| 1 | B | 168 | HIS |
| 1 | B | 262 | ASN |

##### 5.3.3 RNA (i)

There are no RNA molecules in this entry.

#### 5.4 Non-standard residues in protein, DNA, RNA chains (i)

There are no non-standard protein/DNA/RNA residues in this entry.

#### 5.5 Carbohydrates (i)

There are no monosaccharides in this entry.

For Manuscript Review

#### 5.6 Ligand geometry (i)

15 ligands are modelled in this entry.

In the following table, the Counts columns list the number of bonds (or angles) for which Mogul statistics could be retrieved, the number of bonds (or angles) that are observed in the model and the number of bonds (or angles) that are defined in the Chemical Component Dictionary. The Link column lists molecule types, if any, to which the group is linked. The Z score for a bond length (or angle) is the number of standard deviations the observed value is removed from the expected value. A bond length (or angle) with $|Z| > 2$ is considered an outlier worth inspection. RMSZ is the root-mean-square of all Z scores of the bond lengths (or angles).

| Mol | Type | Chain | Res | Link | Bond lengths | | | Bond angles | | |
|---|---|---|---|---|---|---|---|---|---|---|
| | | | | | Counts | RMSZ | $\#\|Z\| > 2$ | Counts | RMSZ | $\#\|Z\| > 2$ |
| 3 | SO4 | A | 406 | - | 4,4,4 | 0.26 | 0 | 6,6,6 | 0.13 | 0 |
| 3 | SO4 | A | 407 | - | 4,4,4 | 0.34 | 0 | 6,6,6 | 0.09 | 0 |
| 3 | SO4 | B | 407 | - | 4,4,4 | 0.33 | 0 | 6,6,6 | 0.12 | 0 |
| 3 | SO4 | A | 403 | - | 4,4,4 | 0.26 | 0 | 6,6,6 | 0.07 | 0 |
| 3 | SO4 | B | 403 | - | 4,4,4 | 0.30 | 0 | 6,6,6 | 0.14 | 0 |
| 3 | SO4 | A | 405 | - | 4,4,4 | 0.29 | 0 | 6,6,6 | 0.07 | 0 |
| 2 | FWU | A | 401 | - | 37,37,37 | 0.36 | 0 | 50,50,50 | 0.82 | 3 (6%) |
| 3 | SO4 | B | 402 | - | 4,4,4 | 0.32 | 0 | 6,6,6 | 0.12 | 0 |
| 3 | SO4 | A | 402 | - | 4,4,4 | 0.41 | 0 | 6,6,6 | 0.12 | 0 |
| 3 | SO4 | B | 408 | - | 4,4,4 | 0.43 | 0 | 6,6,6 | 0.06 | 0 |
| 3 | SO4 | B | 406 | - | 4,4,4 | 0.33 | 0 | 6,6,6 | 0.11 | 0 |
| 3 | SO4 | A | 404 | - | 4,4,4 | 0.30 | 0 | 6,6,6 | 0.29 | 0 |
| 3 | SO4 | B | 405 | - | 4,4,4 | 0.33 | 0 | 6,6,6 | 0.12 | 0 |
| 2 | FWU | B | 401 | - | 37,37,37 | 0.26 | 0 | 50,50,50 | 0.58 | 1 (2%) |
| 3 | SO4 | B | 404 | - | 4,4,4 | 0.30 | 0 | 6,6,6 | 0.21 | 0 |

In the following table, the Chirals column lists the number of chiral outliers, the number of chiral centers analysed, the number of these observed in the model and the number defined in the Chemical Component Dictionary. Similar counts are reported in the Torsion and Rings columns. '-' means no outliers of that kind were identified.

| Mol | Type | Chain | Res | Link | Chirals | Torsions | Rings |
|---|---|---|---|---|---|---|---|
| 2 | FWU | A | 401 | - | - | 14/22/22/22 | 0/4/4/4 |
| 2 | FWU | B | 401 | - | - | 7/22/22/22 | 0/4/4/4 |

There are no bond length outliers.

All (4) bond angle outliers are listed below:

| Mol | Chain | Res | Type | Atoms | Z | Observed(°) | Ideal(°) |
|---|---|---|---|---|---|---|---|
| 2 | A | 401 | FWU | C9-S-N1 | 2.54 | 111.41 | 106.62 |

*Continued on next page...*

For Manuscript Review

*Continued from previous page...*

| Mol | Chain | Res | Type | Atoms | Z | Observed(°) | Ideal(°) |
|---|---|---|---|---|---|---|---|
| 2 | A | 401 | FWU | O1-S-C9 | -2.31 | 104.00 | 108.08 |
| 2 | A | 401 | FWU | C21-C20-C | 2.24 | 127.31 | 123.53 |
| 2 | B | 401 | FWU | C21-C20-C | 2.05 | 126.97 | 123.53 |

There are no chirality outliers.

All (21) torsion outliers are listed below:

| Mol | Chain | Res | Type | Atoms |
|---|---|---|---|---|
| 2 | A | 401 | FWU | C-C20-C21-C22 |
| 2 | A | 401 | FWU | C-C20-C21-C26 |
| 2 | A | 401 | FWU | C14-C9-S-O1 |
| 2 | A | 401 | FWU | C8-N1-S-O |
| 2 | A | 401 | FWU | C8-N1-S-C9 |
| 2 | A | 401 | FWU | C6-C7-C8-N1 |
| 2 | B | 401 | FWU | C14-C9-S-O1 |
| 2 | A | 401 | FWU | C14-C9-S-O |
| 2 | B | 401 | FWU | C14-C9-S-O |
| 2 | A | 401 | FWU | C5-C6-C7-C8 |
| 2 | A | 401 | FWU | C14-C9-S-N1 |
| 2 | B | 401 | FWU | C14-C9-S-N1 |
| 2 | A | 401 | FWU | C5-C4-N-C3 |
| 2 | A | 401 | FWU | C8-N1-S-O1 |
| 2 | A | 401 | FWU | C4-C5-C6-C7 |
| 2 | B | 401 | FWU | C10-C9-S-O1 |
| 2 | B | 401 | FWU | C10-C9-S-O |
| 2 | B | 401 | FWU | C5-C6-C7-C8 |
| 2 | A | 401 | FWU | C19-C20-C21-C26 |
| 2 | A | 401 | FWU | C19-C20-C21-C22 |
| 2 | B | 401 | FWU | C2-C3-N-C4 |

There are no ring outliers.

2 monomers are involved in 5 short contacts:

| Mol | Chain | Res | Type | Clashes | Symm-Clashes |
|---|---|---|---|---|---|
| 2 | A | 401 | FWU | 2 | 0 |
| 2 | B | 401 | FWU | 3 | 0 |

The following is a two-dimensional graphical depiction of Mogul quality analysis of bond lengths, bond angles, torsion angles, and ring geometry for all instances of the Ligand of Interest. In addition, ligands with molecular weight > 250 and outliers as shown on the validation Tables will also be included. For torsion angles, if less then 5% of the Mogul distribution of torsion angles is within 10 degrees of the torsion angle in question, then that torsion angle is considered an outlier.

For Manuscript Review

Any bond that is central to one or more torsion angles identified as an outlier by Mogul will be highlighted in the graph. For rings, the root-mean-square deviation (RMSD) between the ring in question and similar rings identified by Mogul is calculated over all ring torsion angles. If the average RMSD is greater than 60 degrees and the minimal RMSD between the ring in question and any Mogul-identified rings is also greater than 60 degrees, then that ring is considered an outlier. The outliers are highlighted in purple. The color gray indicates Mogul did not find sufficient equivalents in the CSD to analyse the geometry.

**Ligand FWU A 401**

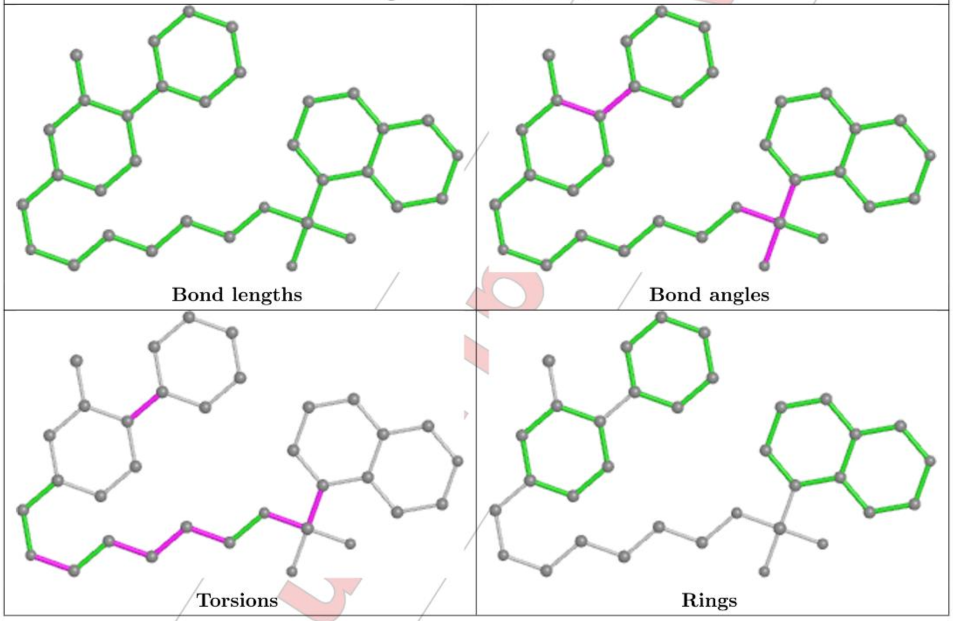

For Manuscript Review

| Ligand FWU B 401 |
|---|

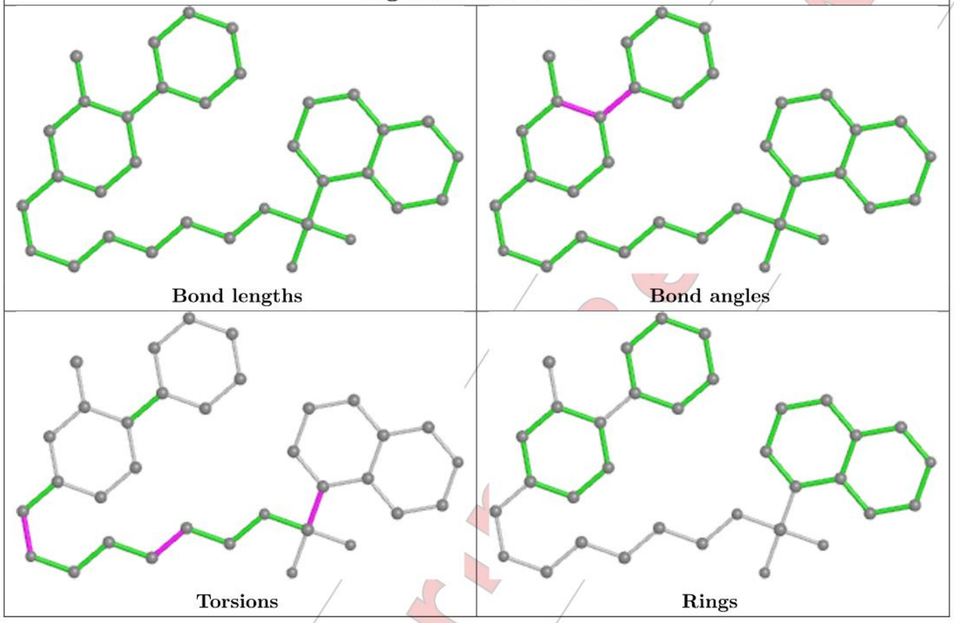

#### 5.7 Other polymers (i)

There are no such residues in this entry.

#### 5.8 Polymer linkage issues (i)

There are no chain breaks in this entry.

For Manuscript Review

### 6 Fit of model and data (i)

#### 6.1 Protein, DNA and RNA chains (i)

In the following table, the column labelled '#RSRZ> 2' contains the number (and percentage) of RSRZ outliers, followed by percent RSRZ outliers for the chain as percentile scores relative to all X-ray entries and entries of similar resolution. The OWAB column contains the minimum, median, $95^{th}$ percentile and maximum values of the occupancy-weighted average B-factor per residue. The column labelled 'Q< 0.9' lists the number of (and percentage) of residues with an average occupancy less than 0.9.

| Mol | Chain | Analysed | <RSRZ> | #RSRZ>2 | | | OWAB($Å^2$) | Q<0.9 |
|---|---|---|---|---|---|---|---|---|
| 1 | A | 330/338 (97%) | -0.21 | 1 (0%) | 94 | 93 | 29, 49, 93, 141 | 0 |
| 1 | B | 328/338 (97%) | -0.20 | 2 (0%) | 89 | 88 | 30, 50, 92, 143 | 0 |
| All | All | 658/676 (97%) | -0.21 | 3 (0%) | 91 | 89 | 29, 50, 93, 143 | 0 |

All (3) RSRZ outliers are listed below:

| Mol | Chain | Res | Type | RSRZ |
|---|---|---|---|---|
| 1 | A | 50 | TYR | 2.6 |
| 1 | B | 330 | ASP | 2.2 |
| 1 | B | 106 | SER | 2.2 |

#### 6.2 Non-standard residues in protein, DNA, RNA chains (i)

There are no non-standard protein/DNA/RNA residues in this entry.

#### 6.3 Carbohydrates (i)

There are no monosaccharides in this entry.

#### 6.4 Ligands (i)

In the following table, the Atoms column lists the number of modelled atoms in the group and the number defined in the chemical component dictionary. The B-factors column lists the minimum, median, $95^{th}$ percentile and maximum values of B factors of atoms in the group. The column labelled 'Q< 0.9' lists the number of atoms with occupancy less than 0.9.

| Mol | Type | Chain | Res | Atoms | RSCC | RSR | B-factors($Å^2$) | Q<0.9 |
|---|---|---|---|---|---|---|---|---|
| 3 | SO4 | B | 406 | 5/5 | 0.71 | 0.23 | 99,124,130,131 | 0 |
| 3 | SO4 | A | 407 | 5/5 | 0.73 | 0.25 | 103,104,120,133 | 0 |

*Continued on next page...*

For Manuscript Review

*Continued from previous page...*

| Mol | Type | Chain | Res | Atoms | RSCC | RSR | B-factors($Å^2$) | Q<0.9 |
|---|---|---|---|---|---|---|---|---|
| 3 | SO4 | A | 404 | 5/5 | 0.74 | 0.22 | 79,80,104,107 | 0 |
| 3 | SO4 | B | 407 | 5/5 | 0.80 | 0.28 | 99,102,117,124 | 0 |
| 2 | FWU | A | 401 | 34/34 | 0.86 | 0.26 | 59,75,109,122 | 0 |
| 3 | SO4 | B | 408 | 5/5 | 0.87 | 0.61 | 30,30,30,30 | 0 |
| 3 | SO4 | A | 405 | 5/5 | 0.88 | 0.20 | 74,76,87,92 | 0 |
| 2 | FWU | B | 401 | 34/34 | 0.89 | 0.27 | 51,71,93,133 | 0 |
| 3 | SO4 | A | 406 | 5/5 | 0.90 | 0.16 | 68,71,77,90 | 0 |
| 3 | SO4 | B | 404 | 5/5 | 0.91 | 0.15 | 75,85,91,96 | 0 |
| 3 | SO4 | B | 405 | 5/5 | 0.93 | 0.13 | 77,81,89,91 | 0 |
| 3 | SO4 | B | 403 | 5/5 | 0.93 | 0.15 | 69,80,92,106 | 0 |
| 3 | SO4 | A | 403 | 5/5 | 0.96 | 0.15 | 53,63,72,74 | 0 |
| 3 | SO4 | B | 402 | 5/5 | 0.96 | 0.14 | 65,67,70,85 | 0 |
| 3 | SO4 | A | 402 | 5/5 | 0.97 | 0.13 | 49,57,63,69 | 0 |

The following is a graphical depiction of the model fit to experimental electron density of all instances of the Ligand of Interest. In addition, ligands with molecular weight > 250 and outliers as shown on the geometry validation Tables will also be included. Each fit is shown from different orientation to approximate a three-dimensional view.

**Electron density around FWU A 401:**

$2mF_o$-$DF_c$ (at 0.7 rmsd) in gray
$mF_o$-$DF_c$ (at 3 rmsd) in purple (negative)
and green (positive)

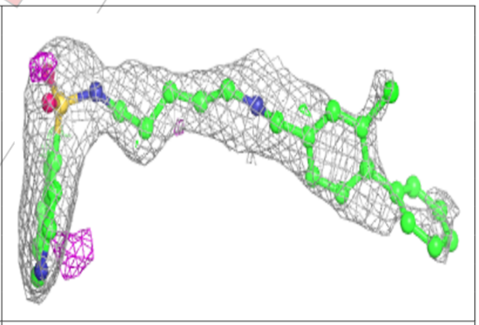

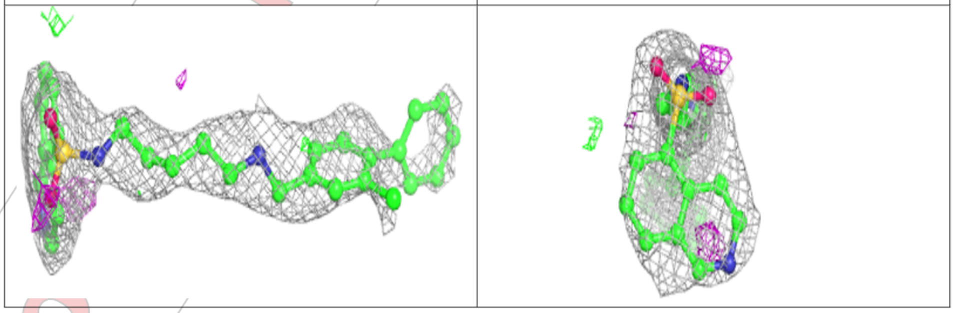

For Manuscript Review

**Electron density around FWU B 401:**

2mF$_o$-DF$_c$ (at 0.7 rmsd) in gray
mF$_o$-DF$_c$ (at 3 rmsd) in purple (negative)
and green (positive)

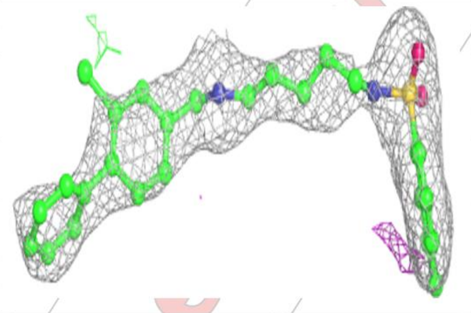

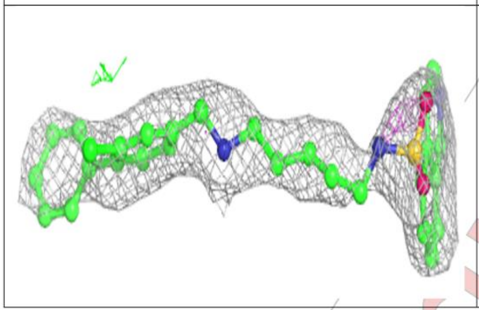

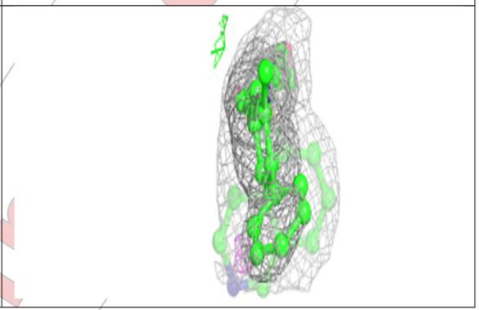

#### 6.5 Other polymers (i)

There are no such residues in this entry.

For Manuscript Review

### Full wwPDB X-ray Structure Validation Report (i)

Mar 20, 2024 – 02:48 pm GMT

| | | |
|---|---|---|
| PDB ID | : | 9EPV |
| Title | : | Crystal structure of human Casein Kinase II subunit alpha (CK2a1) in complex with allosteric ligand FGC333 |
| Deposited on | : | 2024-03-20 |
| Resolution | : | 2.30 Å(reported) |

**This wwPDB validation report is for manuscript review**

For Manuscript Review

This is a Full wwPDB X-ray Structure Validation Report.

This report is produced by the wwPDB biocuration pipeline after annotation of the structure.

We welcome your comments at **
A user guide is available at
https://www.wwpdb.org/validation/2017/XrayValidationReportHelp
with specific help available everywhere you see the (i) symbol.

The types of validation reports are described at
http://www.wwpdb.org/validation/2017/FAQs#types.

---

The following versions of software and data (see references (i)) were used in the production of this report:

| | | |
|---|---|---|
| MolProbity | : | 4.02b-467 |
| Mogul | : | 1.8.4, CSD as541be (2020) |
| Xtriage (Phenix) | : | 1.13 |
| EDS | : | 2.36 |
| buster-report | : | 1.1.7 (2018) |
| Percentile statistics | : | 20191225.v01 (using entries in the PDB archive December 25th 2019) |
| Refmac | : | 5.8.0158 |
| CCP4 | : | 7.0.044 (Gargrove) |
| Ideal geometry (proteins) | : | Engh & Huber (2001) |

### 1 Overall quality at a glance (i)

The following experimental techniques were used to determine the structure:
*X-RAY DIFFRACTION*

The reported resolution of this entry is 2.30 Å.

Percentile scores (ranging between 0-100) for global validation metrics of the entry are shown in the following graphic. The table shows the number of entries on which the scores are based.

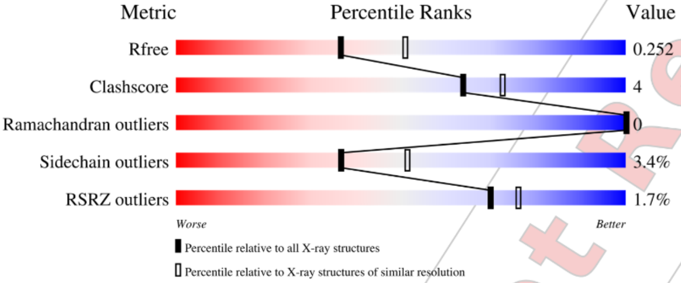

| Metric | Whole archive (#Entries) | Similar resolution (#Entries, resolution range(Å)) |
|---|---|---|
| $R_{free}$ | 130704 | 5042 (2.30-2.30) |
| Clashscore | 141614 | 5643 (2.30-2.30) |
| Ramachandran outliers | 138981 | 5575 (2.30-2.30) |
| Sidechain outliers | 138945 | 5575 (2.30-2.30) |
| RSRZ outliers | 127900 | 4938 (2.30-2.30) |

The table below summarises the geometric issues observed across the polymeric chains and their fit to the electron density. The red, orange, yellow and green segments of the lower bar indicate the fraction of residues that contain outliers for >=3, 2, 1 and 0 types of geometric quality criteria respectively. A grey segment represents the fraction of residues that are not modelled. The numeric value for each fraction is indicated below the corresponding segment, with a dot representing fractions <=5% The upper red bar (where present) indicates the fraction of residues that have poor fit to the electron density. The numeric value is given above the bar.

| Mol | Chain | Length | Quality of chain |
|---|---|---|---|
| 1 | A | 338 | % / 88% 8% • • |
| 1 | B | 338 | 2% / 87% 8% • • |

The following table lists non-polymeric compounds, carbohydrate monomers and non-standard

Ideal geometry (DNA, RNA) : Parkinson et al. (1996)
Validation Pipeline (wwPDB-VP) : 2.36

For Manuscript Review

residues in protein, DNA, RNA chains that are outliers for geometric or electron-density-fit criteria:

| Mol | Type | Chain | Res | Chirality | Geometry | Clashes | Electron density |
|---|---|---|---|---|---|---|---|
| 2 | SO4 | B | 403 | - | - | - | X |
| 3 | EDO | A | 403 | - | - | X | - |

For Manuscript Review

#### 2 Entry composition (i)

There are 5 unique types of molecules in this entry. The entry contains 5663 atoms, of which 0 are hydrogens and 0 are deuteriums.

In the tables below, the ZeroOcc column contains the number of atoms modelled with zero occupancy, the AltConf column contains the number of residues with at least one atom in alternate conformation and the Trace column contains the number of residues modelled with at most 2 atoms.

- Molecule 1 is a protein called Casein kinase II subunit alpha.

| Mol | Chain | Residues | Atoms | | | | | ZeroOcc | AltConf | Trace |
|---|---|---|---|---|---|---|---|---|---|---|
| 1 | A | 329 | Total 2667 | C 1711 | N 462 | O 483 | S 11 | 0 | 0 | 0 |
| 1 | B | 323 | Total 2645 | C 1698 | N 455 | O 481 | S 11 | 0 | 0 | 0 |

There are 2 discrepancies between the modelled and reference sequences:

| Chain | Residue | Modelled | Actual | Comment | Reference |
|---|---|---|---|---|---|
| A | 0 | SER | - | expression tag | UNP P68400 |
| B | 0 | SER | - | expression tag | UNP P68400 |

- Molecule 2 is SULFATE ION (three-letter code: SO4) (formula: $O_4S$).

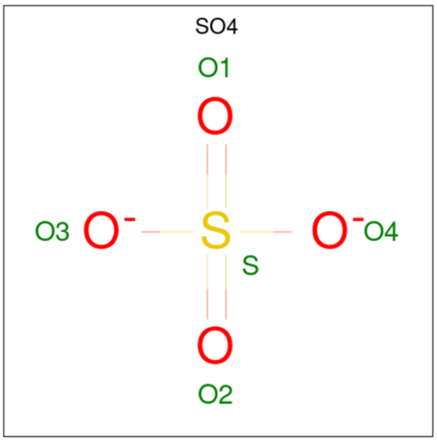

| Mol | Chain | Residues | Atoms | | | ZeroOcc | AltConf |
|---|---|---|---|---|---|---|---|
| 2 | A | 1 | Total 5 | O 4 | S 1 | 0 | 0 |

*Continued on next page...*

*Continued from previous page...*

| Mol | Chain | Residues | Atoms | ZeroOcc | AltConf |
|---|---|---|---|---|---|
| 2 | A | 1 | Total O S<br>5 4 1 | 0 | 0 |
| 2 | A | 1 | Total O S<br>5 4 1 | 0 | 0 |
| 2 | A | 1 | Total O S<br>5 4 1 | 0 | 0 |
| 2 | A | 1 | Total O S<br>5 4 1 | 0 | 0 |
| 2 | A | 1 | Total O S<br>5 4 1 | 0 | 0 |
| 2 | B | 1 | Total O S<br>5 4 1 | 0 | 0 |
| 2 | B | 1 | Total O S<br>5 4 1 | 0 | 0 |
| 2 | B | 1 | Total O S<br>5 4 1 | 0 | 0 |
| 2 | B | 1 | Total O S<br>5 4 1 | 0 | 0 |
| 2 | B | 1 | Total O S<br>5 4 1 | 0 | 0 |
| 2 | B | 1 | Total O S<br>5 4 1 | 0 | 0 |
| 2 | B | 1 | Total O S<br>5 4 1 | 0 | 0 |

- Molecule 3 is 1,2-ETHANEDIOL (three-letter code: EDO) (formula: $C_2H_6O_2$).

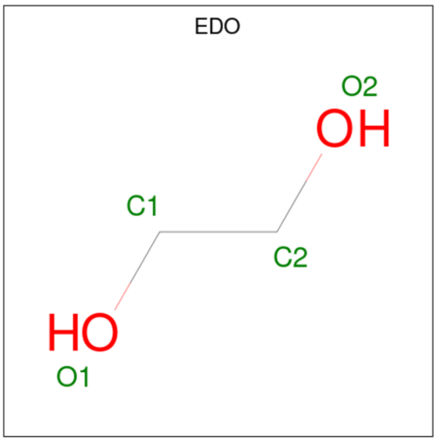

For Manuscript Review

| Mol | Chain | Residues | Atoms | ZeroOcc | AltConf |
|---|---|---|---|---|---|
| 3 | A | 1 | Total C O<br>4 2 2 | 0 | 0 |
| 3 | A | 1 | Total C O<br>4 2 2 | 0 | 0 |
| 3 | B | 1 | Total C O<br>4 2 2 | 0 | 0 |
| 3 | B | 1 | Total C O<br>4 2 2 | 0 | 0 |
| 3 | B | 1 | Total C O<br>4 2 2 | 0 | 0 |
| 3 | B | 1 | Total C O<br>4 2 2 | 0 | 0 |

- Molecule 4 is 5-[[2-[(3-chloranyl-4-phenyl-phenyl)methylamino]-7-azaspiro[3.5]nonan-7-yl]sulfonyl]-1,3-dimethyl-benzimidazol-2-one (three-letter code: A1H6H) (formula: $C_{30}H_{33}ClN_4O_3S$) (labeled as "Ligand of Interest" by depositor).

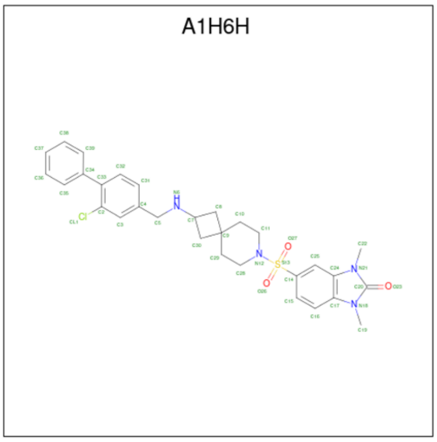

| Mol | Chain | Residues | Atoms | ZeroOcc | AltConf |
|---|---|---|---|---|---|
| 4 | A | 1 | Total C Cl N O S<br>39 30 1 4 3 1 | 0 | 0 |
| 4 | B | 1 | Total C Cl N O S<br>39 30 1 4 3 1 | 0 | 0 |

- Molecule 5 is water.

| Mol | Chain | Residues | Atoms | ZeroOcc | AltConf |
|---|---|---|---|---|---|
| 5 | A | 84 | Total O<br>84 84 | 0 | 0 |

*Continued on next page...*

For Manuscript Review

*Continued from previous page...*

| Mol | Chain | Residues | Atoms | | ZeroOcc | AltConf |
|---|---|---|---|---|---|---|
| | | | Total | O | | |
| 5 | B | 100 | 100 | 100 | 0 | 0 |

For Manuscript Review

#### 3 Residue-property plots ⓘ

These plots are drawn for all protein, RNA, DNA and oligosaccharide chains in the entry. The first graphic for a chain summarises the proportions of the various outlier classes displayed in the second graphic. The second graphic shows the sequence view annotated by issues in geometry and electron density. Residues are color-coded according to the number of geometric quality criteria for which they contain at least one outlier: green = 0, yellow = 1, orange = 2 and red = 3 or more. A red dot above a residue indicates a poor fit to the electron density (RSRZ > 2). Stretches of 2 or more consecutive residues without any outlier are shown as a green connector. Residues present in the sample, but not in the model, are shown in grey.

- Molecule 1: Casein kinase II subunit alpha

Chain A:

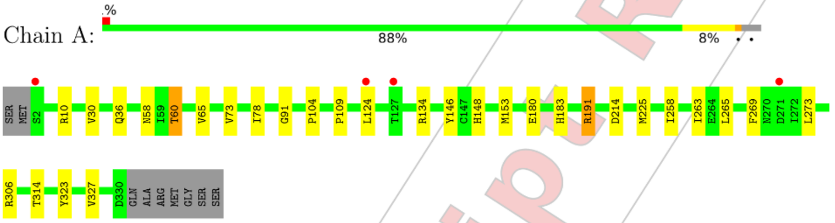

- Molecule 1: Casein kinase II subunit alpha

Chain B:

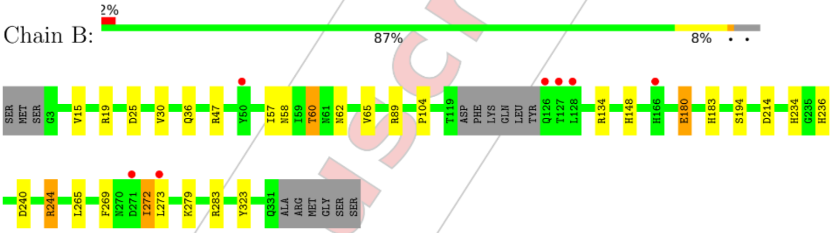

For Manuscript Review

### 4 Data and refinement statistics (i)

| Property | Value | Source |
|---|---|---|
| Space group | P 43 21 2 | Depositor |
| Cell constants<br>a, b, c, $\alpha$, $\beta$, $\gamma$ | 127.09Å 127.09Å 125.97Å<br>90.00° 90.00° 90.00° | Depositor |
| Resolution (Å) | 44.97 – 2.30<br>44.93 – 2.30 | Depositor<br>EDS |
| % Data completeness<br>(in resolution range) | 100.0 (44.97-2.30)<br>100.0 (44.93-2.30) | Depositor<br>EDS |
| $R_{merge}$ | (Not available) | Depositor |
| $R_{sym}$ | (Not available) | Depositor |
| $< I/\sigma(I) >$ [1] | 2.18 (at 2.29Å) | Xtriage |
| Refinement program | REFMAC 5.8.0419 | Depositor |
| R, $R_{free}$ | 0.215 , 0.248<br>0.221 , 0.252 | Depositor<br>DCC |
| $R_{free}$ test set | 2282 reflections (4.92%) | wwPDB-VP |
| Wilson B-factor (Å$^2$) | 35.6 | Xtriage |
| Anisotropy | 0.365 | Xtriage |
| Bulk solvent $k_{sol}$(e/Å$^3$), $B_{sol}$(Å$^2$) | 0.37 , 37.8 | EDS |
| L-test for twinning[2] | $< \|L\| > = 0.51$, $< L^2 > = 0.34$ | Xtriage |
| Estimated twinning fraction | 0.000 for -h,-l,-k<br>0.000 for l,-k,h | Xtriage |
| $F_o$,$F_c$ correlation | 0.94 | EDS |
| Total number of atoms | 5663 | wwPDB-VP |
| Average B, all atoms (Å$^2$) | 41.0 | wwPDB-VP |

Xtriage's analysis on translational NCS is as follows: *The analyses of the Patterson function reveals a significant off-origin peak that is 37.26 % of the origin peak, indicating pseudo-translational symmetry. The chance of finding a peak of this or larger height randomly in a structure without pseudo-translational symmetry is equal to 4.4081e-04. The detected translational NCS is most likely also responsible for the elevated intensity ratio.*

[1] Intensities estimated from amplitudes.
[2] Theoretical values of $< |L| >$, $< L^2 >$ for acentric reflections are 0.5, 0.333 respectively for untwinned datasets, and 0.375, 0.2 for perfectly twinned datasets.

For Manuscript Review

### 5 Model quality (i)

#### 5.1 Standard geometry (i)

Bond lengths and bond angles in the following residue types are not validated in this section: SO4, EDO, A1H6H

The Z score for a bond length (or angle) is the number of standard deviations the observed value is removed from the expected value. A bond length (or angle) with $|Z| > 5$ is considered an outlier worth inspection. RMSZ is the root-mean-square of all Z scores of the bond lengths (or angles).

| Mol | Chain | Bond lengths | | Bond angles | |
|---|---|---|---|---|---|
| | | RMSZ | $\#\|Z\|>5$ | RMSZ | $\#\|Z\|>5$ |
| 1 | A | 0.43 | 0/2739 | 0.72 | 0/3721 |
| 1 | B | 0.47 | 0/2717 | 0.75 | 0/3687 |
| All | All | 0.45 | 0/5456 | 0.74 | 0/7408 |

Chiral center outliers are detected by calculating the chiral volume of a chiral center and verifying if the center is modelled as a planar moiety or with the opposite hand.A planarity outlier is detected by checking planarity of atoms in a peptide group, atoms in a mainchain group or atoms of a sidechain that are expected to be planar.

| Mol | Chain | #Chirality outliers | #Planarity outliers |
|---|---|---|---|
| 1 | A | 0 | 3 |

There are no bond length outliers.

There are no bond angle outliers.

There are no chirality outliers.

All (3) planarity outliers are listed below:

| Mol | Chain | Res | Type | Group |
|---|---|---|---|---|
| 1 | A | 10 | ARG | Sidechain |
| 1 | A | 191 | ARG | Sidechain |
| 1 | A | 306 | ARG | Sidechain |

#### 5.2 Too-close contacts (i)

In the following table, the Non-H and H(model) columns list the number of non-hydrogen atoms and hydrogen atoms in the chain respectively. The H(added) column lists the number of hydrogen atoms added and optimized by MolProbity. The Clashes column lists the number of clashes within the asymmetric unit, whereas Symm-Clashes lists symmetry-related clashes.

For Manuscript Review

| Mol | Chain | Non-H | H(model) | H(added) | Clashes | Symm-Clashes |
|---|---|---|---|---|---|---|
| 1 | A | 2667 | 0 | 2527 | 22 | 0 |
| 1 | B | 2645 | 0 | 2502 | 21 | 0 |
| 2 | A | 30 | 0 | 0 | 1 | 0 |
| 2 | B | 35 | 0 | 0 | 0 | 0 |
| 3 | A | 8 | 0 | 12 | 4 | 0 |
| 3 | B | 16 | 0 | 24 | 4 | 0 |
| 4 | A | 39 | 0 | 0 | 1 | 0 |
| 4 | B | 39 | 0 | 0 | 0 | 0 |
| 5 | A | 84 | 0 | 0 | 2 | 0 |
| 5 | B | 100 | 0 | 0 | 4 | 0 |
| All | All | 5663 | 0 | 5065 | 44 | 0 |

The all-atom clashscore is defined as the number of clashes found per 1000 atoms (including hydrogen atoms). The all-atom clashscore for this structure is 4.

All (44) close contacts within the same asymmetric unit are listed below, sorted by their clash magnitude.

| Atom-1 | Atom-2 | Interatomic distance (Å) | Clash overlap (Å) |
|---|---|---|---|
| 1:A:258:ILE:HD13 | 1:A:265:LEU:HG | 1.48 | 0.93 |
| 1:A:58:ASN:OD1 | 1:A:60:THR:HB | 1.87 | 0.74 |
| 1:B:58:ASN:OD1 | 1:B:60:THR:HB | 1.87 | 0.73 |
| 1:B:234:HIS:O | 1:B:244:ARG:NH1 | 2.24 | 0.70 |
| 1:B:148:HIS:CD2 | 3:B:406:EDO:H12 | 2.28 | 0.68 |
| 1:B:180:GLU:HG3 | 3:B:410:EDO:H21 | 1.78 | 0.66 |
| 1:B:19:ARG:HD2 | 5:B:525:HOH:O | 1.95 | 0.66 |
| 1:A:148:HIS:CD2 | 3:A:403:EDO:H12 | 2.33 | 0.64 |
| 1:B:183:HIS:HE1 | 5:B:540:HOH:O | 1.81 | 0.62 |
| 1:B:36:GLN:HG3 | 1:B:104:PRO:HD3 | 1.82 | 0.60 |
| 1:A:78:ILE:HD13 | 1:A:109:PRO:HG2 | 1.83 | 0.59 |
| 1:B:269:PHE:O | 1:B:273:LEU:HG | 2.04 | 0.58 |
| 1:A:148:HIS:CD2 | 3:A:403:EDO:C1 | 2.86 | 0.58 |
| 1:A:183:HIS:HE1 | 5:A:519:HOH:O | 1.86 | 0.57 |
| 1:B:15:VAL:HG13 | 5:B:525:HOH:O | 2.04 | 0.55 |
| 4:A:407:A1H6H:CL1 | 4:A:407:A1H6H:C39 | 2.92 | 0.55 |
| 1:B:183:HIS:HD2 | 5:B:523:HOH:O | 1.90 | 0.55 |
| 1:B:236:HIS:ND1 | 1:B:240:ASP:OD2 | 2.39 | 0.55 |
| 1:A:269:PHE:O | 1:A:273:LEU:HG | 2.09 | 0.53 |
| 1:A:258:ILE:HG23 | 1:A:263:ILE:HB | 1.91 | 0.52 |
| 1:A:258:ILE:HD13 | 1:A:265:LEU:CG | 2.32 | 0.52 |
| 1:B:57:ILE:HD11 | 1:B:62:ASN:HD22 | 1.74 | 0.52 |
| 1:B:134:ARG:HG2 | 1:B:323:TYR:CZ | 2.44 | 0.52 |
| 1:A:73:VAL:HG23 | 1:A:78:ILE:HD11 | 1.91 | 0.51 |

Continued on next page...

For Manuscript Review

*Continued from previous page...*

| Atom-1 | Atom-2 | Interatomic distance (Å) | Clash overlap (Å) |
|---|---|---|---|
| 1:A:73:VAL:HG23 | 1:A:78:ILE:HG13 | 1.93 | 0.51 |
| 1:A:191:ARG:HD2 | 2:A:409:SO4:O3 | 2.12 | 0.50 |
| 1:A:134:ARG:HG2 | 1:A:323:TYR:CZ | 2.47 | 0.50 |
| 1:B:236:HIS:CE1 | 1:B:240:ASP:CG | 2.85 | 0.49 |
| 1:A:73:VAL:HG23 | 1:A:78:ILE:CG1 | 2.43 | 0.49 |
| 1:A:148:HIS:HD2 | 3:A:403:EDO:C1 | 2.26 | 0.49 |
| 1:A:183:HIS:HD2 | 5:A:531:HOH:O | 1.95 | 0.49 |
| 1:A:36:GLN:HG3 | 1:A:104:PRO:HD3 | 1.95 | 0.48 |
| 1:B:236:HIS:ND1 | 1:B:240:ASP:CG | 2.68 | 0.47 |
| 1:A:73:VAL:HG23 | 1:A:78:ILE:CD1 | 2.45 | 0.47 |
| 1:A:258:ILE:CD1 | 1:A:265:LEU:HG | 2.34 | 0.46 |
| 1:B:148:HIS:HE1 | 1:B:214:ASP:OD2 | 1.99 | 0.46 |
| 1:B:25:ASP:HA | 3:B:411:EDO:C1 | 2.48 | 0.43 |
| 1:B:244:ARG:HG2 | 1:B:244:ARG:HH11 | 1.83 | 0.43 |
| 1:B:272:ILE:HD12 | 1:B:272:ILE:HA | 1.92 | 0.43 |
| 1:A:148:HIS:HE1 | 1:A:214:ASP:OD2 | 2.01 | 0.43 |
| 1:A:314:THR:HA | 3:A:403:EDO:H22 | 2.01 | 0.42 |
| 1:A:91:GLY:HA3 | 1:A:146:TYR:CE2 | 2.55 | 0.42 |
| 1:B:25:ASP:HA | 3:B:411:EDO:H12 | 2.01 | 0.41 |
| 1:B:265:LEU:HD23 | 1:B:265:LEU:HA | 1.92 | 0.41 |

There are no symmetry-related clashes.

#### 5.3 Torsion angles (i)

##### 5.3.1 Protein backbone (i)

In the following table, the Percentiles column shows the percent Ramachandran outliers of the chain as a percentile score with respect to all X-ray entries followed by that with respect to entries of similar resolution.

The Analysed column shows the number of residues for which the backbone conformation was analysed, and the total number of residues.

| Mol | Chain | Analysed | Favoured | Allowed | Outliers | Percentiles | |
|---|---|---|---|---|---|---|---|
| 1 | A | 327/338 (97%) | 316 (97%) | 11 (3%) | 0 | 100 | 100 |
| 1 | B | 319/338 (94%) | 311 (98%) | 8 (2%) | 0 | 100 | 100 |
| All | All | 646/676 (96%) | 627 (97%) | 19 (3%) | 0 | 100 | 100 |

There are no Ramachandran outliers to report.

For Manuscript Review

##### 5.3.2 Protein sidechains (i)

In the following table, the Percentiles column shows the percent sidechain outliers of the chain as a percentile score with respect to all X-ray entries followed by that with respect to entries of similar resolution.

The Analysed column shows the number of residues for which the sidechain conformation was analysed, and the total number of residues.

| Mol | Chain | Analysed | Rotameric | Outliers | Percentiles | |
|---|---|---|---|---|---|---|
| 1 | A | 278/310 (90%) | 270 (97%) | 8 (3%) | 42 | 58 |
| 1 | B | 277/310 (89%) | 266 (96%) | 11 (4%) | 31 | 44 |
| All | All | 555/620 (90%) | 536 (97%) | 19 (3%) | 37 | 51 |

All (19) residues with a non-rotameric sidechain are listed below:

| Mol | Chain | Res | Type |
|---|---|---|---|
| 1 | A | 30 | VAL |
| 1 | A | 60 | THR |
| 1 | A | 65 | VAL |
| 1 | A | 124 | LEU |
| 1 | A | 153 | MET |
| 1 | A | 180 | GLU |
| 1 | A | 225 | MET |
| 1 | A | 327 | VAL |
| 1 | B | 30 | VAL |
| 1 | B | 47 | ARG |
| 1 | B | 60 | THR |
| 1 | B | 65 | VAL |
| 1 | B | 89 | ARG |
| 1 | B | 180 | GLU |
| 1 | B | 194 | SER |
| 1 | B | 244 | ARG |
| 1 | B | 272 | ILE |
| 1 | B | 279 | LYS |
| 1 | B | 283 | ARG |

Sometimes sidechains can be flipped to improve hydrogen bonding and reduce clashes. All (6) such sidechains are listed below:

| Mol | Chain | Res | Type |
|---|---|---|---|
| 1 | A | 148 | HIS |
| 1 | A | 183 | HIS |
| 1 | A | 270 | ASN |

*Continued on next page...*

For Manuscript Review

*Continued from previous page...*

| Mol | Chain | Res | Type |
|---|---|---|---|
| 1 | B | 62 | ASN |
| 1 | B | 148 | HIS |
| 1 | B | 183 | HIS |

###### 5.3.3 RNA (i)

There are no RNA molecules in this entry.

##### 5.4 Non-standard residues in protein, DNA, RNA chains (i)

There are no non-standard protein/DNA/RNA residues in this entry.

##### 5.5 Carbohydrates (i)

There are no monosaccharides in this entry.

##### 5.6 Ligand geometry (i)

21 ligands are modelled in this entry.

In the following table, the Counts columns list the number of bonds (or angles) for which Mogul statistics could be retrieved, the number of bonds (or angles) that are observed in the model and the number of bonds (or angles) that are defined in the Chemical Component Dictionary. The Link column lists molecule types, if any, to which the group is linked. The Z score for a bond length (or angle) is the number of standard deviations the observed value is removed from the expected value. A bond length (or angle) with $|Z| > 2$ is considered an outlier worth inspection. RMSZ is the root-mean-square of all Z scores of the bond lengths (or angles).

| Mol | Type | Chain | Res | Link | Bond lengths | | | Bond angles | | |
|---|---|---|---|---|---|---|---|---|---|---|
| | | | | | Counts | RMSZ | $\#\|Z\| > 2$ | Counts | RMSZ | $\#\|Z\| > 2$ |
| 3 | EDO | B | 411 | - | 3,3,3 | 0.13 | 0 | 2,2,2 | 0.14 | 0 |
| 2 | SO4 | B | 404 | - | 4,4,4 | 0.24 | 0 | 6,6,6 | 0.24 | 0 |
| 2 | SO4 | B | 403 | - | 4,4,4 | 0.33 | 0 | 6,6,6 | 0.12 | 0 |
| 3 | EDO | A | 406 | - | 3,3,3 | 0.44 | 0 | 2,2,2 | 0.48 | 0 |
| 2 | SO4 | B | 408 | - | 4,4,4 | 0.38 | 0 | 6,6,6 | 0.14 | 0 |
| 4 | A1H6H | A | 407 | - | 42,44,44 | 0.47 | 0 | 56,67,67 | 0.77 | 1 (1%) |
| 2 | SO4 | B | 409 | - | 4,4,4 | 0.31 | 0 | 6,6,6 | 0.17 | 0 |
| 2 | SO4 | B | 402 | - | 4,4,4 | 0.42 | 0 | 6,6,6 | 0.31 | 0 |
| 2 | SO4 | A | 409 | - | 4,4,4 | 0.25 | 0 | 6,6,6 | 0.33 | 0 |
| 3 | EDO | B | 412 | - | 3,3,3 | 0.13 | 0 | 2,2,2 | 0.45 | 0 |
| 3 | EDO | B | 406 | - | 3,3,3 | 0.22 | 0 | 2,2,2 | 0.62 | 0 |

WORLDWIDE PDB PROTEIN DATA BANK

For Manuscript Review

| Mol | Type | Chain | Res | Link | Bond lengths | | | Bond angles | | |
|---|---|---|---|---|---|---|---|---|---|---|
| | | | | | Counts | RMSZ | #\|Z\| > 2 | Counts | RMSZ | #\|Z\| > 2 |
| 2 | SO4 | A | 404 | - | 4,4,4 | 0.32 | 0 | 6,6,6 | 0.29 | 0 |
| 4 | A1H6H | B | 407 | - | 42,44,44 | 0.44 | 0 | 56,67,67 | 0.65 | 1 (1%) |
| 2 | SO4 | A | 402 | - | 4,4,4 | 0.24 | 0 | 6,6,6 | 0.09 | 0 |
| 2 | SO4 | A | 408 | - | 4,4,4 | 0.35 | 0 | 6,6,6 | 0.07 | 0 |
| 2 | SO4 | A | 401 | - | 4,4,4 | 0.27 | 0 | 6,6,6 | 0.06 | 0 |
| 2 | SO4 | B | 405 | - | 4,4,4 | 0.48 | 0 | 6,6,6 | 0.31 | 0 |
| 3 | EDO | B | 410 | - | 3,3,3 | 0.32 | 0 | 2,2,2 | 0.68 | 0 |
| 2 | SO4 | B | 401 | - | 4,4,4 | 0.28 | 0 | 6,6,6 | 0.41 | 0 |
| 3 | EDO | A | 403 | - | 3,3,3 | 0.39 | 0 | 2,2,2 | 0.55 | 0 |
| 2 | SO4 | A | 405 | - | 4,4,4 | 0.64 | 0 | 6,6,6 | 0.36 | 0 |

In the following table, the Chirals column lists the number of chiral outliers, the number of chiral centers analysed, the number of these observed in the model and the number defined in the Chemical Component Dictionary. Similar counts are reported in the Torsion and Rings columns. '-' means no outliers of that kind were identified.

| Mol | Type | Chain | Res | Link | Chirals | Torsions | Rings |
|---|---|---|---|---|---|---|---|
| 3 | EDO | A | 406 | - | - | 1/1/1/1 | - |
| 3 | EDO | B | 406 | - | - | 0/1/1/1 | - |
| 3 | EDO | B | 411 | - | - | 1/1/1/1 | - |
| 4 | A1H6H | A | 407 | - | - | 4/19/43/43 | 0/6/6/6 |
| 4 | A1H6H | B | 407 | - | - | 1/19/43/43 | 0/6/6/6 |
| 3 | EDO | B | 410 | - | - | 1/1/1/1 | - |
| 3 | EDO | B | 412 | - | - | 1/1/1/1 | - |
| 3 | EDO | A | 403 | - | - | 0/1/1/1 | - |

There are no bond length outliers.

All (2) bond angle outliers are listed below:

| Mol | Chain | Res | Type | Atoms | Z | Observed(°) | Ideal(°) |
|---|---|---|---|---|---|---|---|
| 4 | A | 407 | A1H6H | C5-N6-C7 | -2.96 | 109.91 | 114.14 |
| 4 | B | 407 | A1H6H | C5-N6-C7 | -2.00 | 111.28 | 114.14 |

There are no chirality outliers.

All (9) torsion outliers are listed below:

| Mol | Chain | Res | Type | Atoms |
|---|---|---|---|---|
| 4 | A | 407 | A1H6H | C2-C33-C34-C35 |
| 4 | A | 407 | A1H6H | C2-C33-C34-C39 |
| 4 | A | 407 | A1H6H | C32-C33-C34-C39 |
| 4 | A | 407 | A1H6H | C32-C33-C34-C35 |

*Continued on next page...*

For Manuscript Review

*Continued from previous page...*

| Mol | Chain | Res | Type | Atoms |
|---|---|---|---|---|
| 3 | A | 406 | EDO | O1-C1-C2-O2 |
| 3 | B | 412 | EDO | O1-C1-C2-O2 |
| 3 | B | 411 | EDO | O1-C1-C2-O2 |
| 4 | B | 407 | A1H6H | C4-C5-N6-C7 |
| 3 | B | 410 | EDO | O1-C1-C2-O2 |

There are no ring outliers.

6 monomers are involved in 10 short contacts:

| Mol | Chain | Res | Type | Clashes | Symm-Clashes |
|---|---|---|---|---|---|
| 3 | B | 411 | EDO | 2 | 0 |
| 4 | A | 407 | A1H6H | 1 | 0 |
| 2 | A | 409 | SO4 | 1 | 0 |
| 3 | B | 406 | EDO | 1 | 0 |
| 3 | B | 410 | EDO | 1 | 0 |
| 3 | A | 403 | EDO | 4 | 0 |

The following is a two-dimensional graphical depiction of Mogul quality analysis of bond lengths, bond angles, torsion angles, and ring geometry for all instances of the Ligand of Interest. In addition, ligands with molecular weight > 250 and outliers as shown on the validation Tables will also be included. For torsion angles, if less then 5% of the Mogul distribution of torsion angles is within 10 degrees of the torsion angle in question, then that torsion angle is considered an outlier. Any bond that is central to one or more torsion angles identified as an outlier by Mogul will be highlighted in the graph. For rings, the root-mean-square deviation (RMSD) between the ring in question and similar rings identified by Mogul is calculated over all ring torsion angles. If the average RMSD is greater than 60 degrees and the minimal RMSD between the ring in question and any Mogul-identified rings is also greater than 60 degrees, then that ring is considered an outlier. The outliers are highlighted in purple. The color gray indicates Mogul did not find sufficient equivalents in the CSD to analyse the geometry.

For Manuscript Review

Ligand A1H6H A 407

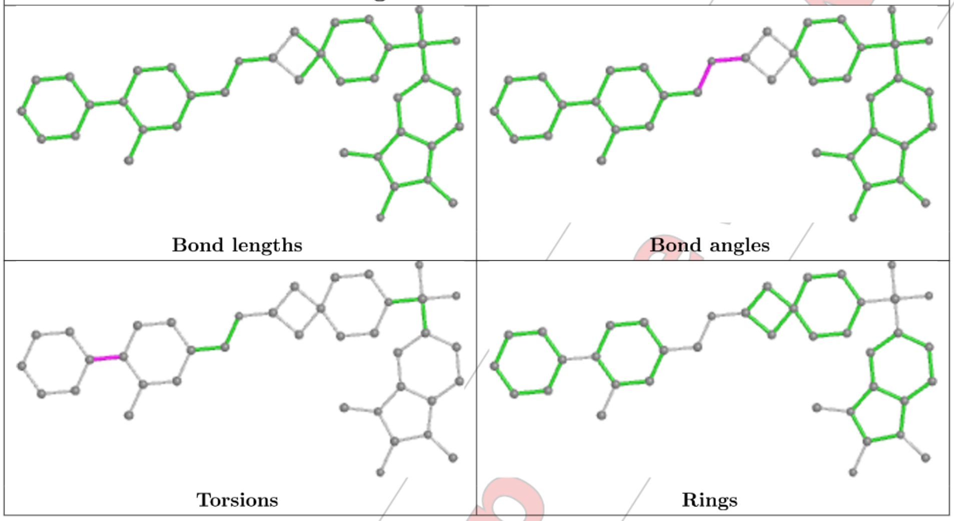

Ligand A1H6H B 407

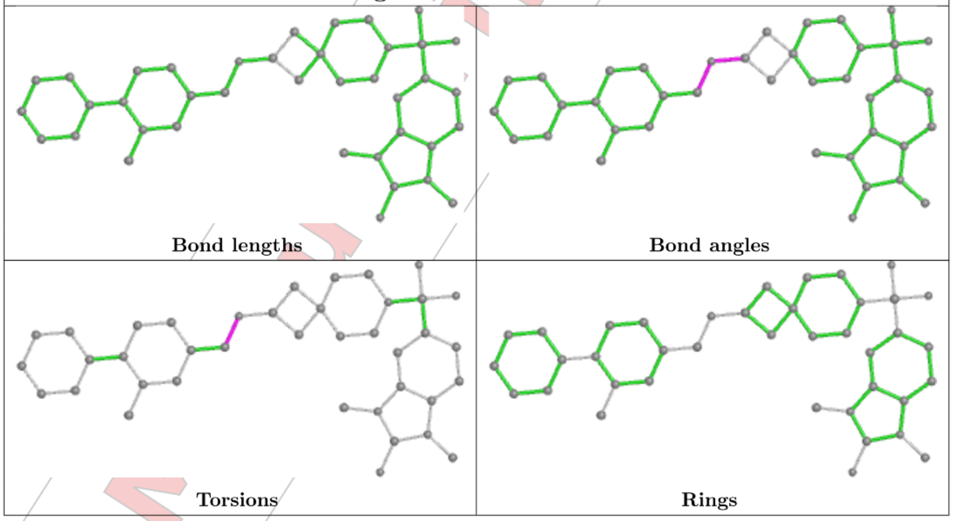

#### 5.7 Other polymers (i)

There are no such residues in this entry.

For Manuscript Review

#### 5.8 Polymer linkage issues (i)

There are no chain breaks in this entry.

For Manuscript Review

### 6 Fit of model and data (i)

#### 6.1 Protein, DNA and RNA chains (i)

In the following table, the column labelled '#RSRZ> 2' contains the number (and percentage) of RSRZ outliers, followed by percent RSRZ outliers for the chain as percentile scores relative to all X-ray entries and entries of similar resolution. The OWAB column contains the minimum, median, $95^{th}$ percentile and maximum values of the occupancy-weighted average B-factor per residue. The column labelled 'Q< 0.9' lists the number of (and percentage) of residues with an average occupancy less than 0.9.

| Mol | Chain | Analysed | <RSRZ> | #RSRZ>2 | | | OWAB(Å$^2$) | Q<0.9 |
|---|---|---|---|---|---|---|---|---|
| 1 | A | 329/338 (97%) | -0.21 | 4 (1%) | 79 | 83 | 23, 38, 74, 120 | 0 |
| 1 | B | 323/338 (95%) | -0.20 | 7 (2%) | 62 | 69 | 22, 37, 69, 88 | 0 |
| All | All | 652/676 (96%) | -0.20 | 11 (1%) | 70 | 76 | 22, 37, 70, 120 | 0 |

All (11) RSRZ outliers are listed below:

| Mol | Chain | Res | Type | RSRZ |
|---|---|---|---|---|
| 1 | B | 128 | LEU | 4.7 |
| 1 | A | 2 | SER | 3.6 |
| 1 | B | 50 | TYR | 3.5 |
| 1 | A | 127 | THR | 3.4 |
| 1 | B | 127 | THR | 3.1 |
| 1 | A | 271 | ASP | 2.8 |
| 1 | B | 273 | LEU | 2.7 |
| 1 | A | 124 | LEU | 2.6 |
| 1 | B | 271 | ASP | 2.5 |
| 1 | B | 126 | GLN | 2.3 |
| 1 | B | 166 | HIS | 2.3 |

#### 6.2 Non-standard residues in protein, DNA, RNA chains (i)

There are no non-standard protein/DNA/RNA residues in this entry.

#### 6.3 Carbohydrates (i)

There are no monosaccharides in this entry.

For Manuscript Review

#### 6.4 Ligands (i)

In the following table, the Atoms column lists the number of modelled atoms in the group and the number defined in the chemical component dictionary. The B-factors column lists the minimum, median, $95^{th}$ percentile and maximum values of B factors of atoms in the group. The column labelled 'Q< 0.9' lists the number of atoms with occupancy less than 0.9.

| Mol | Type | Chain | Res | Atoms | RSCC | RSR | B-factors($Å^2$) | Q<0.9 |
|---|---|---|---|---|---|---|---|---|
| 2 | SO4 | B | 403 | 5/5 | 0.74 | 0.55 | 101,111,117,119 | 0 |
| 3 | EDO | A | 406 | 4/4 | 0.80 | 0.14 | 45,52,61,61 | 0 |
| 2 | SO4 | A | 401 | 5/5 | 0.81 | 0.19 | 63,72,96,96 | 0 |
| 2 | SO4 | B | 409 | 5/5 | 0.84 | 0.26 | 73,78,93,102 | 0 |
| 4 | A1H6H | B | 407 | 39/39 | 0.84 | 0.22 | 40,59,83,91 | 0 |
| 4 | A1H6H | A | 407 | 39/39 | 0.88 | 0.18 | 41,53,74,132 | 0 |
| 2 | SO4 | A | 404 | 5/5 | 0.88 | 0.15 | 59,61,75,86 | 0 |
| 2 | SO4 | A | 408 | 5/5 | 0.90 | 0.25 | 76,92,114,117 | 0 |
| 2 | SO4 | B | 404 | 5/5 | 0.90 | 0.11 | 49,59,72,79 | 0 |
| 2 | SO4 | B | 401 | 5/5 | 0.90 | 0.17 | 57,59,74,79 | 0 |
| 2 | SO4 | B | 408 | 5/5 | 0.93 | 0.36 | 70,72,95,103 | 0 |
| 3 | EDO | B | 410 | 4/4 | 0.93 | 0.12 | 39,51,52,53 | 0 |
| 2 | SO4 | A | 402 | 5/5 | 0.94 | 0.11 | 47,55,61,64 | 0 |
| 2 | SO4 | A | 409 | 5/5 | 0.96 | 0.17 | 43,54,62,62 | 0 |
| 3 | EDO | B | 411 | 4/4 | 0.96 | 0.16 | 30,33,36,43 | 0 |
| 3 | EDO | A | 403 | 4/4 | 0.97 | 0.14 | 22,28,31,33 | 0 |
| 3 | EDO | B | 412 | 4/4 | 0.97 | 0.20 | 42,48,52,54 | 0 |
| 3 | EDO | B | 406 | 4/4 | 0.98 | 0.15 | 19,28,31,34 | 0 |
| 2 | SO4 | B | 402 | 5/5 | 0.98 | 0.10 | 45,46,52,56 | 0 |
| 2 | SO4 | A | 405 | 5/5 | 0.99 | 0.15 | 43,47,49,57 | 0 |
| 2 | SO4 | B | 405 | 5/5 | 0.99 | 0.09 | 44,44,60,66 | 0 |

The following is a graphical depiction of the model fit to experimental electron density of all instances of the Ligand of Interest. In addition, ligands with molecular weight > 250 and outliers as shown on the geometry validation Tables will also be included. Each fit is shown from different orientation to approximate a three-dimensional view.

For Manuscript Review

**Electron density around A1H6H B 407:**

$2mF_o$-$DF_c$ (at 0.7 rmsd) in gray
$mF_o$-$DF_c$ (at 3 rmsd) in purple (negative)
and green (positive)

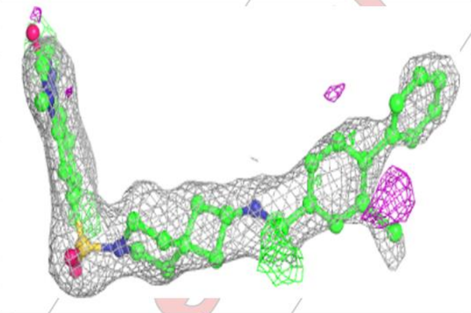

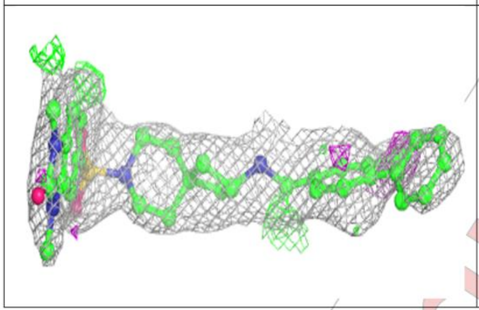

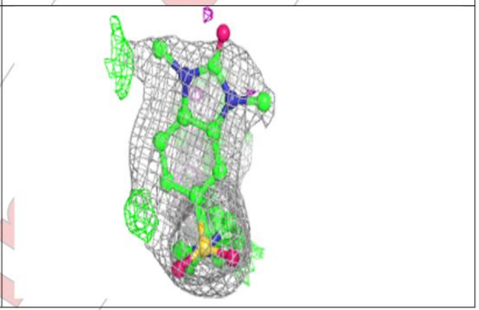

**Electron density around A1H6H A 407:**

$2mF_o$-$DF_c$ (at 0.7 rmsd) in gray
$mF_o$-$DF_c$ (at 3 rmsd) in purple (negative)
and green (positive)

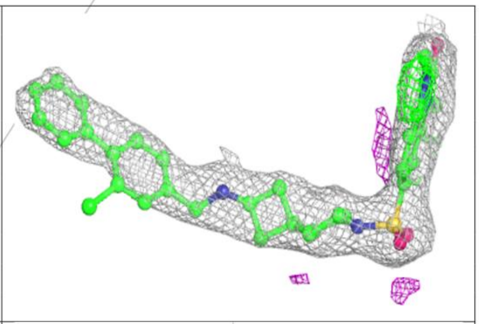

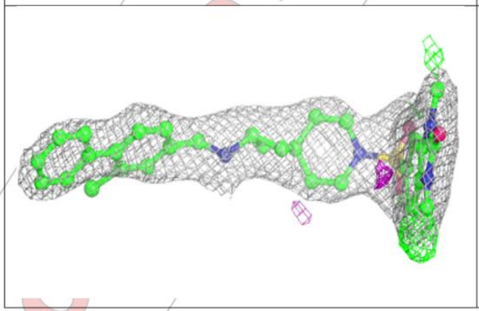

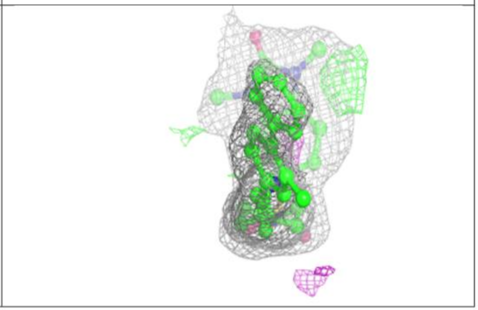

For Manuscript Review

#### 6.5 Other polymers ⓘ

There are no such residues in this entry.

For Manuscript Review

WORLDWIDE PDB PROTEIN DATA BANK

### Full wwPDB X-ray Structure Validation Report (i)

Mar 20, 2024 – 03:52 pm GMT

| | | |
|---|---|---|
| PDB ID | : | 9EPW |
| Title | : | Crystal structure of human Casein Kinase II subunit alpha (CK2a1) in complex with allosteric ligand FGC3336 |
| Deposited on | : | 2024-03-20 |
| Resolution | : | 2.30 Å(reported) |

**This wwPDB validation report is for manuscript review**

For Manuscript Review

This is a Full wwPDB X-ray Structure Validation Report.

This report is produced by the wwPDB biocuration pipeline after annotation of the structure.

We welcome your comments at **
A user guide is available at
https://www.wwpdb.org/validation/2017/XrayValidationReportHelp
with specific help available everywhere you see the (i) symbol.

The types of validation reports are described at
http://www.wwpdb.org/validation/2017/FAQs#types.

The following versions of software and data (see references (i)) were used in the production of this report:

| | | |
|---|---|---|
| MolProbity | : | 4.02b-467 |
| Mogul | : | 1.8.4, CSD as541be (2020) |
| Xtriage (Phenix) | : | 1.13 |
| EDS | : | 2.36 |
| buster-report | : | 1.1.7 (2018) |
| Percentile statistics | : | 20191225.v01 (using entries in the PDB archive December 25th 2019) |
| Refmac | : | 5.8.0158 |
| CCP4 | : | 7.0.044 (Gargrove) |
| Ideal geometry (proteins) | : | Engh & Huber (2001) |

### 1 Overall quality at a glance (i)

The following experimental techniques were used to determine the structure:
*X-RAY DIFFRACTION*

The reported resolution of this entry is 2.30 Å.

Percentile scores (ranging between 0-100) for global validation metrics of the entry are shown in the following graphic. The table shows the number of entries on which the scores are based.

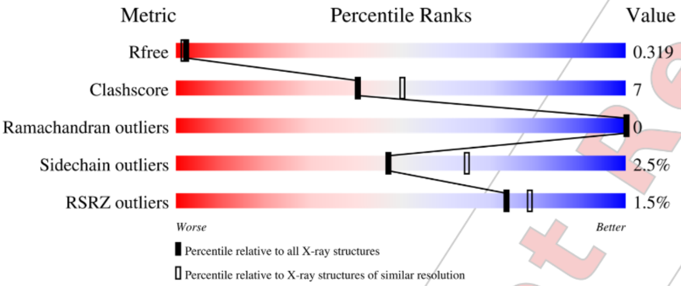

| Metric | Whole archive (#Entries) | Similar resolution (#Entries, resolution range(Å)) |
|---|---|---|
| $R_{free}$ | 130704 | 5042 (2.30-2.30) |
| Clashscore | 141614 | 5643 (2.30-2.30) |
| Ramachandran outliers | 138981 | 5575 (2.30-2.30) |
| Sidechain outliers | 138945 | 5575 (2.30-2.30) |
| RSRZ outliers | 127900 | 4938 (2.30-2.30) |

The table below summarises the geometric issues observed across the polymeric chains and their fit to the electron density. The red, orange, yellow and green segments of the lower bar indicate the fraction of residues that contain outliers for >=3, 2, 1 and 0 types of geometric quality criteria respectively. A grey segment represents the fraction of residues that are not modelled. The numeric value for each fraction is indicated below the corresponding segment, with a dot representing fractions <=5% The upper red bar (where present) indicates the fraction of residues that have poor fit to the electron density. The numeric value is given above the bar.

| Mol | Chain | Length | Quality of chain |
|---|---|---|---|
| 1 | B | 338 | 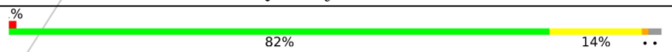 |

The following table lists non-polymeric compounds, carbohydrate monomers and non-standard residues in protein, DNA, RNA chains that are outliers for geometric or electron-density-fit criteria:

Ideal geometry (DNA, RNA) : Parkinson et al. (1996)
Validation Pipeline (wwPDB-VP) : 2.36

For Manuscript Review

| Mol | Type | Chain | Res | Chirality | Geometry | Clashes | Electron density |
|---|---|---|---|---|---|---|---|
| 2 | SO4 | B | 403 | - | - | - | X |

For Manuscript Review

#### 2 Entry composition (i)

There are 4 unique types of molecules in this entry. The entry contains 2779 atoms, of which 0 are hydrogens and 0 are deuteriums.

In the tables below, the ZeroOcc column contains the number of atoms modelled with zero occupancy, the AltConf column contains the number of residues with at least one atom in alternate conformation and the Trace column contains the number of residues modelled with at most 2 atoms.

- Molecule 1 is a protein called Casein kinase II subunit alpha.

| Mol | Chain | Residues | Atoms | | | | | ZeroOcc | AltConf | Trace |
|---|---|---|---|---|---|---|---|---|---|---|
| | | | Total | C | N | O | S | | | |
| 1 | B | 331 | 2684 | 1723 | 459 | 491 | 11 | 0 | 0 | 0 |

There is a discrepancy between the modelled and reference sequences:

| Chain | Residue | Modelled | Actual | Comment | Reference |
|---|---|---|---|---|---|
| B | 0 | SER | - | expression tag | UNP P68400 |

- Molecule 2 is SULFATE ION (three-letter code: SO4) (formula: $O_4S$).

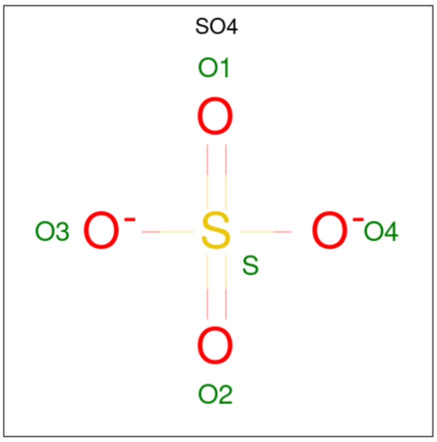

| Mol | Chain | Residues | Atoms | | | ZeroOcc | AltConf |
|---|---|---|---|---|---|---|---|
| | | | Total | O | S | | |
| 2 | B | 1 | 5 | 4 | 1 | 0 | 0 |
| 2 | B | 1 | 5 | 4 | 1 | 0 | 0 |
| 2 | B | 1 | 5 | 4 | 1 | 0 | 0 |

*Continued on next page...*

For Manuscript Review

*Continued from previous page...*

| Mol | Chain | Residues | Atoms | ZeroOcc | AltConf |
|---|---|---|---|---|---|
| 2 | B | 1 | Total O S<br>5 4 1 | 0 | 0 |
| 2 | B | 1 | Total O S<br>5 4 1 | 0 | 0 |

- Molecule 3 is {N}-[(3-chloranyl-4-phenyl-phenyl)methyl]-2-[1-(3-methylquinolin-8-yl)sulfonylpiperidin-4-yl]ethanamine (three-letter code: A1H6F) (formula: $C_{30}H_{32}ClN_3O_2S$) (labeled as "Ligand of Interest" by depositor).

A1H6F

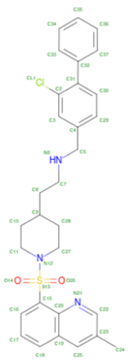

| Mol | Chain | Residues | Atoms | ZeroOcc | AltConf |
|---|---|---|---|---|---|
| 3 | B | 1 | Total C Cl N O S<br>37 30 1 3 2 1 | 0 | 0 |

- Molecule 4 is water.

| Mol | Chain | Residues | Atoms | ZeroOcc | AltConf |
|---|---|---|---|---|---|
| 4 | B | 33 | Total O<br>33 33 | 0 | 0 |

For Manuscript Review

#### 3 Residue-property plots (i)

These plots are drawn for all protein, RNA, DNA and oligosaccharide chains in the entry. The first graphic for a chain summarises the proportions of the various outlier classes displayed in the second graphic. The second graphic shows the sequence view annotated by issues in geometry and electron density. Residues are color-coded according to the number of geometric quality criteria for which they contain at least one outlier: green = 0, yellow = 1, orange = 2 and red = 3 or more. A red dot above a residue indicates a poor fit to the electron density (RSRZ > 2). Stretches of 2 or more consecutive residues without any outlier are shown as a green connector. Residues present in the sample, but not in the model, are shown in grey.

- Molecule 1: Casein kinase II subunit alpha

Chain B:

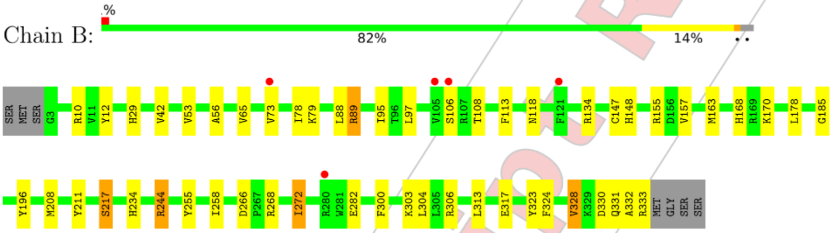

For Manuscript Review

### 4 Data and refinement statistics (i)

| Property | Value | Source |
|---|---|---|
| Space group | P 43 21 2 | Depositor |
| Cell constants<br>a, b, c, $\alpha$, $\beta$, $\gamma$ | 72.06Å 72.06Å 133.28Å<br>90.00° 90.00° 90.00° | Depositor |
| Resolution (Å) | 48.98 – 2.30<br>48.93 – 2.30 | Depositor<br>EDS |
| % Data completeness<br>(in resolution range) | 100.0 (48.98-2.30)<br>100.0 (48.93-2.30) | Depositor<br>EDS |
| $R_{merge}$ | (Not available) | Depositor |
| $R_{sym}$ | (Not available) | Depositor |
| $< I/\sigma(I) >$ [1] | 1.43 (at 2.29Å) | Xtriage |
| Refinement program | REFMAC 5.8.0419 | Depositor |
| R, $R_{free}$ | 0.259 , 0.308<br>0.265 , 0.319 | Depositor<br>DCC |
| $R_{free}$ test set | 809 reflections (4.97%) | wwPDB-VP |
| Wilson B-factor (Å$^2$) | 45.0 | Xtriage |
| Anisotropy | 0.036 | Xtriage |
| Bulk solvent $k_{sol}$(e/Å$^3$), $B_{sol}$(Å$^2$) | 0.35 , 35.0 | EDS |
| L-test for twinning[2] | $< \|L\| > = 0.45$, $< L^2 > = 0.27$ | Xtriage |
| Estimated twinning fraction | No twinning to report. | Xtriage |
| $F_o$,$F_c$ correlation | 0.92 | EDS |
| Total number of atoms | 2779 | wwPDB-VP |
| Average B, all atoms (Å$^2$) | 50.0 | wwPDB-VP |

Xtriage's analysis on translational NCS is as follows: *The largest off-origin peak in the Patterson function is 5.45% of the height of the origin peak. No significant pseudotranslation is detected.*

[1]Intensities estimated from amplitudes.
[2]Theoretical values of $< |L| >$, $< L^2 >$ for acentric reflections are 0.5, 0.333 respectively for untwinned datasets, and 0.375, 0.2 for perfectly twinned datasets.

For Manuscript Review

### 5 Model quality (i)

#### 5.1 Standard geometry (i)

Bond lengths and bond angles in the following residue types are not validated in this section: SO4, A1H6F

The Z score for a bond length (or angle) is the number of standard deviations the observed value is removed from the expected value. A bond length (or angle) with $|Z| > 5$ is considered an outlier worth inspection. RMSZ is the root-mean-square of all Z scores of the bond lengths (or angles).

| Mol | Chain | Bond lengths | | Bond angles | |
|---|---|---|---|---|---|
| | | RMSZ | #$\|Z\|$ >5 | RMSZ | #$\|Z\|$ >5 |
| 1 | B | 0.38 | 0/2758 | 0.75 | 0/3752 |

There are no bond length outliers.

There are no bond angle outliers.

There are no chirality outliers.

There are no planarity outliers.

#### 5.2 Too-close contacts (i)

In the following table, the Non-H and H(model) columns list the number of non-hydrogen atoms and hydrogen atoms in the chain respectively. The H(added) column lists the number of hydrogen atoms added and optimized by MolProbity. The Clashes column lists the number of clashes within the asymmetric unit, whereas Symm-Clashes lists symmetry-related clashes.

| Mol | Chain | Non-H | H(model) | H(added) | Clashes | Symm-Clashes |
|---|---|---|---|---|---|---|
| 1 | B | 2684 | 0 | 2515 | 38 | 0 |
| 2 | B | 25 | 0 | 0 | 0 | 0 |
| 3 | B | 37 | 0 | 0 | 2 | 0 |
| 4 | B | 33 | 0 | 0 | 0 | 0 |
| All | All | 2779 | 0 | 2515 | 38 | 0 |

The all-atom clashscore is defined as the number of clashes found per 1000 atoms (including hydrogen atoms). The all-atom clashscore for this structure is 7.

All (38) close contacts within the same asymmetric unit are listed below, sorted by their clash magnitude.

For Manuscript Review

| Atom-1 | Atom-2 | Interatomic distance (Å) | Clash overlap (Å) |
|---|---|---|---|
| 1:B:185:GLY:HA2 | 1:B:208:MET:HE2 | 1.41 | 0.99 |
| 1:B:185:GLY:HA2 | 1:B:208:MET:CE | 2.01 | 0.89 |
| 1:B:244:ARG:CG | 1:B:244:ARG:HH11 | 1.99 | 0.74 |
| 1:B:168:HIS:O | 1:B:170:LYS:HG2 | 1.95 | 0.67 |
| 1:B:29:HIS:O | 1:B:79:LYS:NZ | 2.29 | 0.66 |
| 1:B:282:GLU:N | 1:B:282:GLU:OE1 | 2.31 | 0.63 |
| 1:B:118:ASN:HB2 | 1:B:163:MET:CE | 2.29 | 0.63 |
| 1:B:155:ARG:HD3 | 1:B:178:LEU:O | 1.99 | 0.62 |
| 1:B:332:ALA:O | 1:B:333:ARG:C | 2.40 | 0.59 |
| 1:B:118:ASN:HB2 | 1:B:163:MET:HE2 | 1.85 | 0.58 |
| 1:B:244:ARG:HH11 | 1:B:244:ARG:HG2 | 1.68 | 0.58 |
| 1:B:266:ASP:OD2 | 1:B:268:ARG:NH2 | 2.37 | 0.57 |
| 1:B:134:ARG:HG2 | 1:B:323:TYR:CZ | 2.41 | 0.55 |
| 1:B:268:ARG:O | 1:B:272:ILE:HG23 | 2.07 | 0.55 |
| 1:B:73:VAL:HG13 | 1:B:78:ILE:HG13 | 1.89 | 0.54 |
| 1:B:255:TYR:O | 1:B:258:ILE:HG13 | 2.08 | 0.53 |
| 1:B:330:ASP:O | 1:B:333:ARG:HG3 | 2.09 | 0.52 |
| 1:B:196:TYR:CE1 | 1:B:234:HIS:HD2 | 2.29 | 0.51 |
| 1:B:106:SER:C | 1:B:108:THR:H | 2.14 | 0.49 |
| 1:B:244:ARG:HH11 | 1:B:244:ARG:HG3 | 1.73 | 0.49 |
| 1:B:95:ILE:HG13 | 1:B:113:PHE:CD1 | 2.47 | 0.49 |
| 1:B:157:VAL:HB | 1:B:217:SER:HB2 | 1.95 | 0.49 |
| 1:B:163:MET:HE1 | 3:B:406:A1H6F:C18 | 2.43 | 0.48 |
| 1:B:244:ARG:CG | 1:B:244:ARG:NH1 | 2.67 | 0.47 |
| 1:B:244:ARG:HG2 | 1:B:244:ARG:NH1 | 2.29 | 0.47 |
| 1:B:42:VAL:HG23 | 1:B:56:ALA:HA | 1.98 | 0.46 |
| 1:B:148:HIS:CD2 | 1:B:211:TYR:HB3 | 2.50 | 0.46 |
| 1:B:89:ARG:HD3 | 1:B:97:LEU:O | 2.17 | 0.44 |
| 1:B:303:LYS:HB3 | 1:B:313:LEU:HG | 2.00 | 0.43 |
| 1:B:324:PHE:O | 1:B:328:VAL:HG13 | 2.19 | 0.43 |
| 1:B:88:LEU:HD21 | 1:B:147:CYS:HA | 2.01 | 0.42 |
| 1:B:12:TYR:HB2 | 1:B:211:TYR:CE2 | 2.55 | 0.41 |
| 1:B:185:GLY:HA2 | 1:B:208:MET:HE3 | 1.95 | 0.41 |
| 1:B:118:ASN:HB2 | 1:B:163:MET:HE3 | 2.00 | 0.41 |
| 1:B:331:GLN:C | 1:B:333:ARG:H | 2.23 | 0.41 |
| 1:B:10:ARG:NH1 | 1:B:317:GLU:OE2 | 2.54 | 0.41 |
| 1:B:300:PHE:CZ | 1:B:304:LEU:HD11 | 2.55 | 0.41 |
| 1:B:53:VAL:HG11 | 3:B:406:A1H6F:C20 | 2.51 | 0.41 |

There are no symmetry-related clashes.

For Manuscript Review

#### 5.3 Torsion angles (i)

##### 5.3.1 Protein backbone (i)

In the following table, the Percentiles column shows the percent Ramachandran outliers of the chain as a percentile score with respect to all X-ray entries followed by that with respect to entries of similar resolution.

The Analysed column shows the number of residues for which the backbone conformation was analysed, and the total number of residues.

| Mol | Chain | Analysed | Favoured | Allowed | Outliers | Percentiles | |
|---|---|---|---|---|---|---|---|
| 1 | B | 329/338 (97%) | 318 (97%) | 11 (3%) | 0 | 100 | 100 |

There are no Ramachandran outliers to report.

##### 5.3.2 Protein sidechains (i)

In the following table, the Percentiles column shows the percent sidechain outliers of the chain as a percentile score with respect to all X-ray entries followed by that with respect to entries of similar resolution.

The Analysed column shows the number of residues for which the sidechain conformation was analysed, and the total number of residues.

| Mol | Chain | Analysed | Rotameric | Outliers | Percentiles | |
|---|---|---|---|---|---|---|
| 1 | B | 278/310 (90%) | 271 (98%) | 7 (2%) | 47 | 65 |

All (7) residues with a non-rotameric sidechain are listed below:

| Mol | Chain | Res | Type |
|---|---|---|---|
| 1 | B | 65 | VAL |
| 1 | B | 89 | ARG |
| 1 | B | 217 | SER |
| 1 | B | 244 | ARG |
| 1 | B | 272 | ILE |
| 1 | B | 306 | ARG |
| 1 | B | 328 | VAL |

Sometimes sidechains can be flipped to improve hydrogen bonding and reduce clashes. All (4) such sidechains are listed below:

| Mol | Chain | Res | Type |
|---|---|---|---|
| 1 | B | 148 | HIS |

*Continued on next page...*

For Manuscript Review

*Continued from previous page...*

| Mol | Chain | Res | Type |
|---|---|---|---|
| 1 | B | 161 | ASN |
| 1 | B | 186 | GLN |
| 1 | B | 234 | HIS |

###### 5.3.3 RNA (i)

There are no RNA molecules in this entry.

##### 5.4 Non-standard residues in protein, DNA, RNA chains (i)

There are no non-standard protein/DNA/RNA residues in this entry.

##### 5.5 Carbohydrates (i)

There are no monosaccharides in this entry.

##### 5.6 Ligand geometry (i)

6 ligands are modelled in this entry.

In the following table, the Counts columns list the number of bonds (or angles) for which Mogul statistics could be retrieved, the number of bonds (or angles) that are observed in the model and the number of bonds (or angles) that are defined in the Chemical Component Dictionary. The Link column lists molecule types, if any, to which the group is linked. The Z score for a bond length (or angle) is the number of standard deviations the observed value is removed from the expected value. A bond length (or angle) with $|Z| > 2$ is considered an outlier worth inspection. RMSZ is the root-mean-square of all Z scores of the bond lengths (or angles).

| Mol | Type | Chain | Res | Link | Bond lengths | | | Bond angles | | |
|---|---|---|---|---|---|---|---|---|---|---|
| | | | | | Counts | RMSZ | $\#\|Z\| > 2$ | Counts | RMSZ | $\#\|Z\| > 2$ |
| 2 | SO4 | B | 403 | - | 4,4,4 | 0.34 | 0 | 6,6,6 | 0.07 | 0 |
| 2 | SO4 | B | 405 | - | 4,4,4 | 0.42 | 0 | 6,6,6 | 0.12 | 0 |
| 2 | SO4 | B | 402 | - | 4,4,4 | 0.26 | 0 | 6,6,6 | 0.13 | 0 |
| 2 | SO4 | B | 401 | - | 4,4,4 | 0.33 | 0 | 6,6,6 | 0.13 | 0 |
| 2 | SO4 | B | 404 | - | 4,4,4 | 0.38 | 0 | 6,6,6 | 0.14 | 0 |
| 3 | A1H6F | B | 406 | - | 41,41,41 | 0.32 | 0 | 56,58,58 | 0.68 | 1 (1%) |

In the following table, the Chirals column lists the number of chiral outliers, the number of chiral centers analysed, the number of these observed in the model and the number defined in the Chemical Component Dictionary. Similar counts are reported in the Torsion and Rings columns.

For Manuscript Review

'-' means no outliers of that kind were identified.

| Mol | Type | Chain | Res | Link | Chirals | Torsions | Rings |
|---|---|---|---|---|---|---|---|
| 3 | A1H6F | B | 406 | - | - | 12/23/33/33 | 0/5/5/5 |

There are no bond length outliers.

All (1) bond angle outliers are listed below:

| Mol | Chain | Res | Type | Atoms | Z | Observed($^o$) | Ideal($^o$) |
|---|---|---|---|---|---|---|---|
| 3 | B | 406 | A1H6F | C7-C8-C9 | 2.52 | 120.73 | 114.32 |

There are no chirality outliers.

All (12) torsion outliers are listed below:

| Mol | Chain | Res | Type | Atoms |
|---|---|---|---|---|
| 3 | B | 406 | A1H6F | C20-C15-S13-O14 |
| 3 | B | 406 | A1H6F | C20-C15-S13-O26 |
| 3 | B | 406 | A1H6F | C27-N12-S13-O14 |
| 3 | B | 406 | A1H6F | C27-N12-S13-C15 |
| 3 | B | 406 | A1H6F | C27-N12-S13-O26 |
| 3 | B | 406 | A1H6F | C11-N12-S13-O14 |
| 3 | B | 406 | A1H6F | C11-N12-S13-O26 |
| 3 | B | 406 | A1H6F | C11-N12-S13-C15 |
| 3 | B | 406 | A1H6F | C16-C15-S13-O14 |
| 3 | B | 406 | A1H6F | C7-C8-C9-C28 |
| 3 | B | 406 | A1H6F | C30-C31-C32-C37 |
| 3 | B | 406 | A1H6F | C30-C31-C32-C33 |

There are no ring outliers.

1 monomer is involved in 2 short contacts:

| Mol | Chain | Res | Type | Clashes | Symm-Clashes |
|---|---|---|---|---|---|
| 3 | B | 406 | A1H6F | 2 | 0 |

The following is a two-dimensional graphical depiction of Mogul quality analysis of bond lengths, bond angles, torsion angles, and ring geometry for all instances of the Ligand of Interest. In addition, ligands with molecular weight > 250 and outliers as shown on the validation Tables will also be included. For torsion angles, if less then 5% of the Mogul distribution of torsion angles is within 10 degrees of the torsion angle in question, then that torsion angle is considered an outlier. Any bond that is central to one or more torsion angles identified as an outlier by Mogul will be highlighted in the graph. For rings, the root-mean-square deviation (RMSD) between the ring in question and similar rings identified by Mogul is calculated over all ring torsion angles. If the average RMSD is greater than 60 degrees and the minimal RMSD between the ring in question and

For Manuscript Review

any Mogul-identified rings is also greater than 60 degrees, then that ring is considered an outlier. The outliers are highlighted in purple. The color gray indicates Mogul did not find sufficient equivalents in the CSD to analyse the geometry.

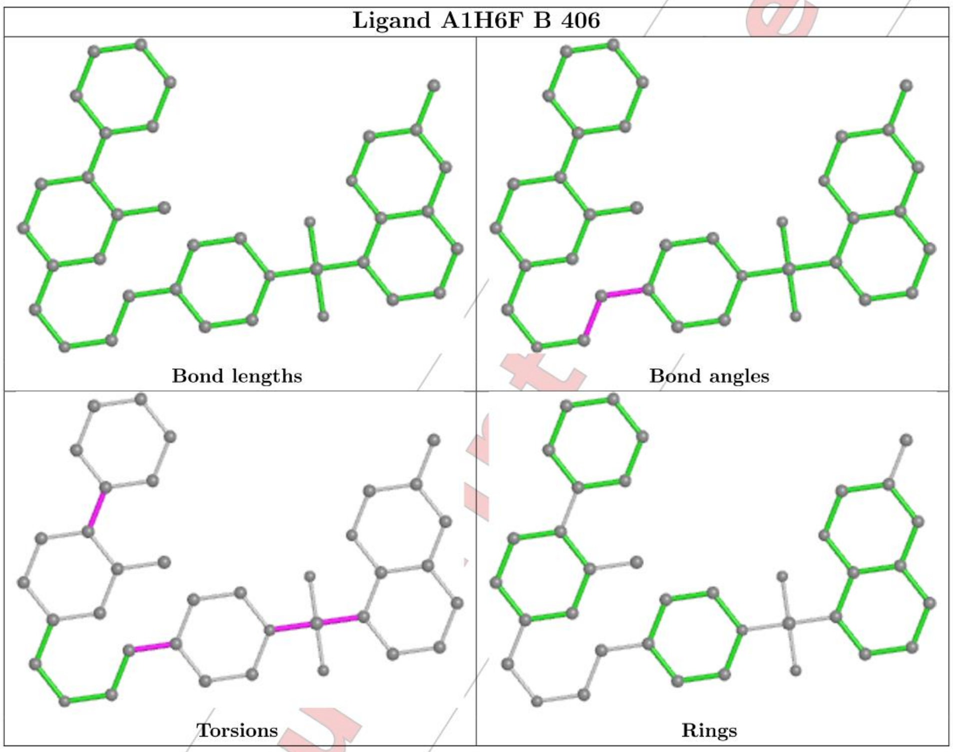

#### 5.7 Other polymers ⓘ

There are no such residues in this entry.

#### 5.8 Polymer linkage issues ⓘ

There are no chain breaks in this entry.

For Manuscript Review

### 6 Fit of model and data (i)

#### 6.1 Protein, DNA and RNA chains (i)

In the following table, the column labelled '#RSRZ> 2' contains the number (and percentage) of RSRZ outliers, followed by percent RSRZ outliers for the chain as percentile scores relative to all X-ray entries and entries of similar resolution. The OWAB column contains the minimum, median, 95$^{th}$ percentile and maximum values of the occupancy-weighted average B-factor per residue. The column labelled 'Q< 0.9' lists the number of (and percentage) of residues with an average occupancy less than 0.9.

| Mol | Chain | Analysed | <RSRZ> | #RSRZ>2 | | | OWAB(Å$^2$) | Q<0.9 |
|---|---|---|---|---|---|---|---|---|
| 1 | B | 331/338 (97%) | 0.22 | 5 (1%) | 73 | 79 | 26, 48, 80, 95 | 0 |

All (5) RSRZ outliers are listed below:

| Mol | Chain | Res | Type | RSRZ |
|---|---|---|---|---|
| 1 | B | 105 | VAL | 4.1 |
| 1 | B | 106 | SER | 2.3 |
| 1 | B | 280 | ARG | 2.3 |
| 1 | B | 121 | PHE | 2.2 |
| 1 | B | 73 | VAL | 2.2 |

#### 6.2 Non-standard residues in protein, DNA, RNA chains (i)

There are no non-standard protein/DNA/RNA residues in this entry.

#### 6.3 Carbohydrates (i)

There are no monosaccharides in this entry.

#### 6.4 Ligands (i)

In the following table, the Atoms column lists the number of modelled atoms in the group and the number defined in the chemical component dictionary. The B-factors column lists the minimum, median, 95$^{th}$ percentile and maximum values of B factors of atoms in the group. The column labelled 'Q< 0.9' lists the number of atoms with occupancy less than 0.9.

| Mol | Type | Chain | Res | Atoms | RSCC | RSR | B-factors(Å$^2$) | Q<0.9 |
|---|---|---|---|---|---|---|---|---|
| 2 | SO4 | B | 403 | 5/5 | 0.50 | 0.43 | 100,102,117,120 | 0 |
| 2 | SO4 | B | 402 | 5/5 | 0.64 | 0.25 | 63,67,87,93 | 0 |

*Continued on next page...*

For Manuscript Review

*Continued from previous page...*

| Mol | Type | Chain | Res | Atoms | RSCC | RSR | B-factors($Å^2$) | Q<0.9 |
|---|---|---|---|---|---|---|---|---|
| 2 | SO4 | B | 401 | 5/5 | 0.73 | 0.31 | 80,91,99,106 | 0 |
| 3 | A1H6F | B | 406 | 37/37 | 0.90 | 0.17 | 38,48,56,66 | 0 |
| 2 | SO4 | B | 404 | 5/5 | 0.91 | 0.20 | 74,76,78,83 | 0 |
| 2 | SO4 | B | 405 | 5/5 | 0.97 | 0.13 | 50,51,52,58 | 0 |

The following is a graphical depiction of the model fit to experimental electron density of all instances of the Ligand of Interest. In addition, ligands with molecular weight > 250 and outliers as shown on the geometry validation Tables will also be included. Each fit is shown from different orientation to approximate a three-dimensional view.

**Electron density around A1H6F B 406:**

$2mF_o$-$DF_c$ (at 0.7 rmsd) in gray
$mF_o$-$DF_c$ (at 3 rmsd) in purple (negative) and green (positive)

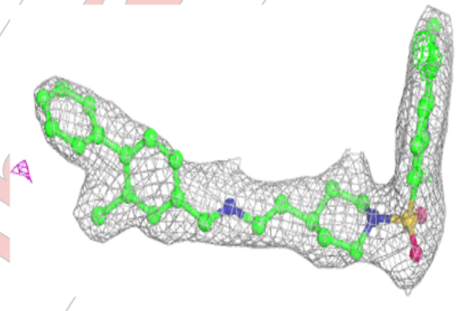

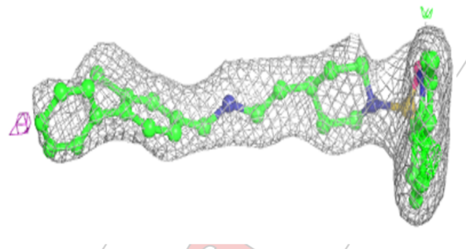

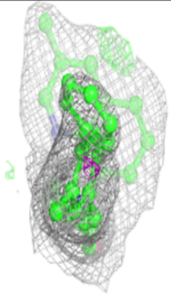

#### 6.5 Other polymers ⓘ

There are no such residues in this entry.

For Manuscript Review

### Full wwPDB X-ray Structure Validation Report (i)

Mar 20, 2024 – 02:51 pm GMT

| | | |
|---|---|---|
| PDB ID | : | 9EPX |
| Title | : | Crystal structure of human Casein Kinase II subunit alpha (CK2a1) in complex with allosteric ligand FGC3331 |
| Deposited on | : | 2024-03-20 |
| Resolution | : | 2.60 Å(reported) |

**This wwPDB validation report is for manuscript review**

For Manuscript Review

This is a Full wwPDB X-ray Structure Validation Report.

This report is produced by the wwPDB biocuration pipeline after annotation of the structure.

We welcome your comments at **
A user guide is available at
https://www.wwpdb.org/validation/2017/XrayValidationReportHelp
with specific help available everywhere you see the (i) symbol.

The types of validation reports are described at
http://www.wwpdb.org/validation/2017/FAQs#types.

The following versions of software and data (see references (i)) were used in the production of this report:

| | | |
|---|---|---|
| MolProbity | : | 4.02b-467 |
| Mogul | : | 1.8.4, CSD as541be (2020) |
| Xtriage (Phenix) | : | 1.13 |
| EDS | : | 2.36 |
| buster-report | : | 1.1.7 (2018) |
| Percentile statistics | : | 20191225.v01 (using entries in the PDB archive December 25th 2019) |
| Refmac | : | 5.8.0158 |
| CCP4 | : | 7.0.044 (Gargrove) |
| Ideal geometry (proteins) | : | Engh & Huber (2001) |

### 1 Overall quality at a glance (i)

The following experimental techniques were used to determine the structure:
*X-RAY DIFFRACTION*

The reported resolution of this entry is 2.60 Å.

Percentile scores (ranging between 0-100) for global validation metrics of the entry are shown in the following graphic. The table shows the number of entries on which the scores are based.

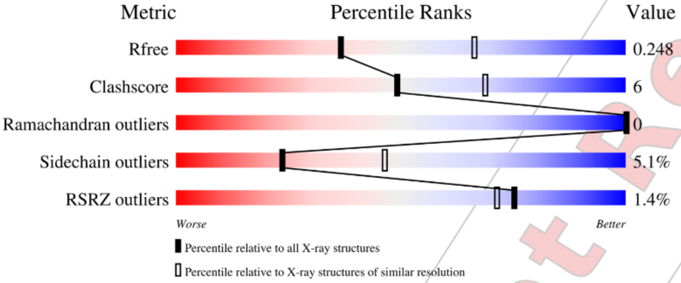

| Metric | Whole archive (#Entries) | Similar resolution (#Entries, resolution range(Å)) |
|---|---|---|
| $R_{free}$ | 130704 | 3163 (2.60-2.60) |
| Clashscore | 141614 | 3518 (2.60-2.60) |
| Ramachandran outliers | 138981 | 3455 (2.60-2.60) |
| Sidechain outliers | 138945 | 3455 (2.60-2.60) |
| RSRZ outliers | 127900 | 3104 (2.60-2.60) |

The table below summarises the geometric issues observed across the polymeric chains and their fit to the electron density. The red, orange, yellow and green segments of the lower bar indicate the fraction of residues that contain outliers for >=3, 2, 1 and 0 types of geometric quality criteria respectively. A grey segment represents the fraction of residues that are not modelled. The numeric value for each fraction is indicated below the corresponding segment, with a dot representing fractions <=5% The upper red bar (where present) indicates the fraction of residues that have poor fit to the electron density. The numeric value is given above the bar.

| Mol | Chain | Length | Quality of chain |
|---|---|---|---|
| 1 | A | 338 | % 86% 10% • • |
| 1 | B | 338 | 2% 81% 15% • • |

Ideal geometry (DNA, RNA) : Parkinson et al. (1996)
Validation Pipeline (wwPDB-VP) : 2.36

WORLDWIDE PDB PROTEIN DATA BANK

For Manuscript Review

#### 2 Entry composition (i)

There are 5 unique types of molecules in this entry. The entry contains 5643 atoms, of which 0 are hydrogens and 0 are deuteriums.

In the tables below, the ZeroOcc column contains the number of atoms modelled with zero occupancy, the AltConf column contains the number of residues with at least one atom in alternate conformation and the Trace column contains the number of residues modelled with at most 2 atoms.

- Molecule 1 is a protein called Casein kinase II subunit alpha.

| Mol | Chain | Residues | Atoms | | | | | ZeroOcc | AltConf | Trace |
|---|---|---|---|---|---|---|---|---|---|---|
| 1 | A | 329 | Total 2697 | C 1737 | N 469 | O 480 | S 11 | 0 | 0 | 0 |
| 1 | B | 329 | Total 2721 | C 1744 | N 475 | O 491 | S 11 | 0 | 0 | 0 |

There are 2 discrepancies between the modelled and reference sequences:

| Chain | Residue | Modelled | Actual | Comment | Reference |
|---|---|---|---|---|---|
| A | 0 | SER | - | expression tag | UNP P68400 |
| B | 0 | SER | - | expression tag | UNP P68400 |

- Molecule 2 is SULFATE ION (three-letter code: SO4) (formula: $O_4S$).

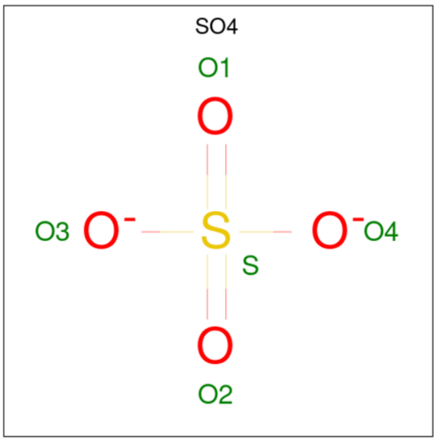

| Mol | Chain | Residues | Atoms | | | ZeroOcc | AltConf |
|---|---|---|---|---|---|---|---|
| 2 | A | 1 | Total 5 | O 4 | S 1 | 0 | 0 |

*Continued on next page...*

For Manuscript Review

*Continued from previous page...*

| Mol | Chain | Residues | Atoms | ZeroOcc | AltConf |
|---|---|---|---|---|---|
| 2 | A | 1 | Total O S<br>5 4 1 | 0 | 0 |
| 2 | A | 1 | Total O S<br>5 4 1 | 0 | 0 |
| 2 | A | 1 | Total O S<br>5 4 1 | 0 | 0 |
| 2 | A | 1 | Total O S<br>5 4 1 | 0 | 0 |
| 2 | A | 1 | Total O S<br>5 4 1 | 0 | 0 |
| 2 | B | 1 | Total O S<br>5 4 1 | 0 | 0 |
| 2 | B | 1 | Total O S<br>5 4 1 | 0 | 0 |
| 2 | B | 1 | Total O S<br>5 4 1 | 0 | 0 |
| 2 | B | 1 | Total O S<br>5 4 1 | 0 | 0 |
| 2 | B | 1 | Total O S<br>5 4 1 | 0 | 0 |
| 2 | B | 1 | Total O S<br>5 4 1 | 0 | 0 |
| 2 | B | 1 | Total O S<br>5 4 1 | 0 | 0 |
| 2 | B | 1 | Total O S<br>5 4 1 | 0 | 0 |
| 2 | B | 1 | Total O S<br>5 4 1 | 0 | 0 |
| 2 | B | 1 | Total O S<br>5 4 1 | 0 | 0 |

- Molecule 3 is 1,2-ETHANEDIOL (three-letter code: EDO) (formula: $C_2H_6O_2$).

For Manuscript Review

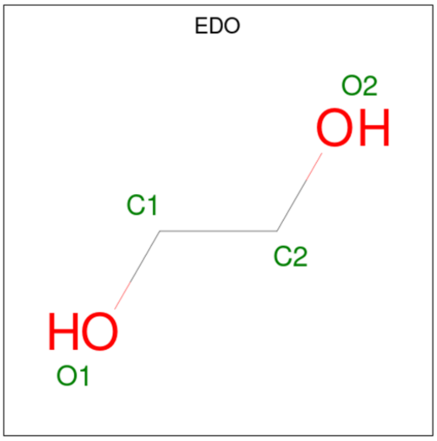

| Mol | Chain | Residues | Atoms | | | ZeroOcc | AltConf |
|---|---|---|---|---|---|---|---|
| | | | Total | C | O | | |
| 3 | A | 1 | 4 | 2 | 2 | 0 | 0 |

- Molecule 4 is 7-[(2-chloranyl-1,3-benzothiazol-6-yl)sulfonyl]- {N}-[(3-chloranyl-4-phenyl-phenyl)methyl]-7-azaspiro[3.5]nonan-2-amine (three-letter code: A1H6E) (formula: $C_{28}H_{27}Cl_2N_3O_2S_2$) (labeled as "Ligand of Interest" by depositor).

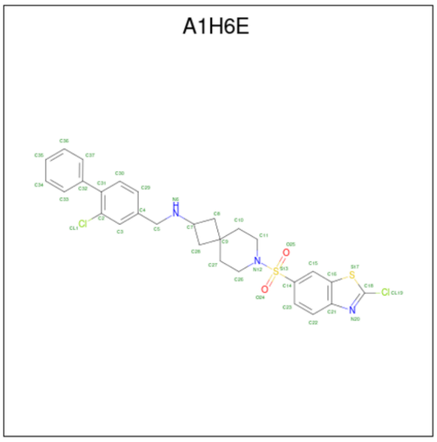

| Mol | Chain | Residues | Atoms | | | | | | ZeroOcc | AltConf |
|---|---|---|---|---|---|---|---|---|---|---|
| | | | Total | C | Cl | N | O | S | | |
| 4 | A | 1 | 37 | 28 | 2 | 3 | 2 | 2 | 0 | 0 |
| 4 | B | 1 | 37 | 28 | 2 | 3 | 2 | 2 | 0 | 0 |

For Manuscript Review

- Molecule 5 is water.

| Mol | Chain | Residues | Atoms | ZeroOcc | AltConf |
|---|---|---|---|---|---|
| 5 | A | 31 | Total O<br>31 31 | 0 | 0 |
| 5 | B | 36 | Total O<br>36 36 | 0 | 0 |

For Manuscript Review

### 3 Residue-property plots (i)

These plots are drawn for all protein, RNA, DNA and oligosaccharide chains in the entry. The first graphic for a chain summarises the proportions of the various outlier classes displayed in the second graphic. The second graphic shows the sequence view annotated by issues in geometry and electron density. Residues are color-coded according to the number of geometric quality criteria for which they contain at least one outlier: green = 0, yellow = 1, orange = 2 and red = 3 or more. A red dot above a residue indicates a poor fit to the electron density (RSRZ > 2). Stretches of 2 or more consecutive residues without any outlier are shown as a green connector. Residues present in the sample, but not in the model, are shown in grey.

• Molecule 1: Casein kinase II subunit alpha

Chain A:

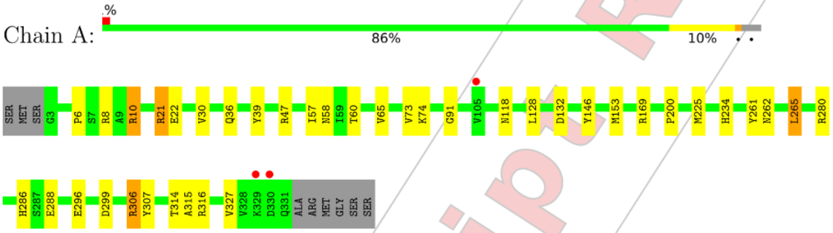

• Molecule 1: Casein kinase II subunit alpha

Chain B:

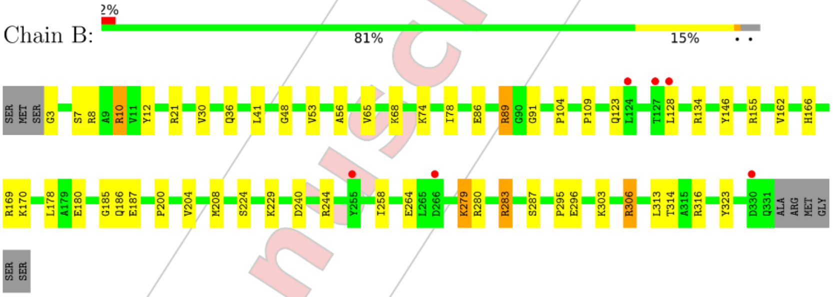

For Manuscript Review

#### 4 Data and refinement statistics (i)

| Property | Value | Source |
|---|---|---|
| Space group | P 43 21 2 | Depositor |
| Cell constants<br>a, b, c, $\alpha$, $\beta$, $\gamma$ | 128.16Å 128.16Å 124.01Å<br>90.00° 90.00° 90.00° | Depositor |
| Resolution (Å) | 45.31 – 2.60<br>45.31 – 2.60 | Depositor<br>EDS |
| % Data completeness<br>(in resolution range) | 100.0 (45.31-2.60)<br>100.0 (45.31-2.60) | Depositor<br>EDS |
| $R_{merge}$ | (Not available) | Depositor |
| $R_{sym}$ | (Not available) | Depositor |
| $< I/\sigma(I) >$ [1] | 2.09 (at 2.61Å) | Xtriage |
| Refinement program | REFMAC 5.8.0419 | Depositor |
| R, $R_{free}$ | 0.214 , 0.246<br>0.227 , 0.248 | Depositor<br>DCC |
| $R_{free}$ test set | 1664 reflections (5.14%) | wwPDB-VP |
| Wilson B-factor (Å$^2$) | 45.6 | Xtriage |
| Anisotropy | 0.484 | Xtriage |
| Bulk solvent $k_{sol}$(e/Å$^3$), $B_{sol}$(Å$^2$) | 0.34 , 37.4 | EDS |
| L-test for twinning[2] | $< \|L\| >$ = 0.50, $< L^2 >$ = 0.33 | Xtriage |
| Estimated twinning fraction | 0.000 for -h,-l,-k<br>0.000 for l,-k,h | Xtriage |
| $F_o$,$F_c$ correlation | 0.93 | EDS |
| Total number of atoms | 5643 | wwPDB-VP |
| Average B, all atoms (Å$^2$) | 52.0 | wwPDB-VP |

Xtriage's analysis on translational NCS is as follows: *The analyses of the Patterson function reveals a significant off-origin peak that is 30.33 % of the origin peak, indicating pseudo-translational symmetry. The chance of finding a peak of this or larger height randomly in a structure without pseudo-translational symmetry is equal to 1.3316e-03. The detected translational NCS is most likely also responsible for the elevated intensity ratio.*

[1]Intensities estimated from amplitudes.
[2]Theoretical values of $< |L| >$, $< L^2 >$ for acentric reflections are 0.5, 0.333 respectively for untwinned datasets, and 0.375, 0.2 for perfectly twinned datasets.

For Manuscript Review

### 5 Model quality (i)

#### 5.1 Standard geometry (i)

Bond lengths and bond angles in the following residue types are not validated in this section: A1H6E, SO4, EDO

The Z score for a bond length (or angle) is the number of standard deviations the observed value is removed from the expected value. A bond length (or angle) with $|Z| > 5$ is considered an outlier worth inspection. RMSZ is the root-mean-square of all Z scores of the bond lengths (or angles).

| Mol | Chain | Bond lengths | | Bond angles | |
|---|---|---|---|---|---|
| | | RMSZ | $\#|Z|>5$ | RMSZ | $\#|Z|>5$ |
| 1 | A | 0.42 | 0/2771 | 0.76 | 0/3760 |
| 1 | B | 0.42 | 0/2794 | 0.76 | 0/3787 |
| All | All | 0.42 | 0/5565 | 0.76 | 0/7547 |

There are no bond length outliers.

There are no bond angle outliers.

There are no chirality outliers.

There are no planarity outliers.

#### 5.2 Too-close contacts (i)

In the following table, the Non-H and H(model) columns list the number of non-hydrogen atoms and hydrogen atoms in the chain respectively. The H(added) column lists the number of hydrogen atoms added and optimized by MolProbity. The Clashes column lists the number of clashes within the asymmetric unit, whereas Symm-Clashes lists symmetry-related clashes.

| Mol | Chain | Non-H | H(model) | H(added) | Clashes | Symm-Clashes |
|---|---|---|---|---|---|---|
| 1 | A | 2697 | 0 | 2590 | 35 | 0 |
| 1 | B | 2721 | 0 | 2617 | 35 | 0 |
| 2 | A | 30 | 0 | 0 | 0 | 0 |
| 2 | B | 50 | 0 | 0 | 2 | 0 |
| 3 | A | 4 | 0 | 6 | 0 | 0 |
| 4 | A | 37 | 0 | 0 | 1 | 0 |
| 4 | B | 37 | 0 | 0 | 1 | 0 |
| 5 | A | 31 | 0 | 0 | 1 | 0 |
| 5 | B | 36 | 0 | 0 | 1 | 0 |
| All | All | 5643 | 0 | 5213 | 64 | 0 |

The all-atom clashscore is defined as the number of clashes found per 1000 atoms (including

For Manuscript Review

hydrogen atoms). The all-atom clashscore for this structure is 6.

All (64) close contacts within the same asymmetric unit are listed below, sorted by their clash magnitude.

| Atom-1 | Atom-2 | Interatomic distance (Å) | Clash overlap (Å) |
|---|---|---|---|
| 1:A:21:ARG:HH11 | 1:A:21:ARG:HG3 | 1.21 | 1.05 |
| 1:A:58:ASN:OD1 | 1:A:60:THR:HB | 1.76 | 0.86 |
| 1:A:280:ARG:HD3 | 1:B:306:ARG:NH2 | 1.94 | 0.83 |
| 1:A:21:ARG:HG3 | 1:A:21:ARG:NH1 | 1.85 | 0.82 |
| 1:A:314:THR:HG23 | 5:A:525:HOH:O | 1.79 | 0.81 |
| 1:A:280:ARG:CD | 1:B:306:ARG:NH2 | 2.45 | 0.79 |
| 1:A:128:LEU:HD11 | 1:A:225:MET:HE1 | 1.62 | 0.78 |
| 1:A:280:ARG:CD | 1:B:306:ARG:HH21 | 1.97 | 0.76 |
| 1:A:73:VAL:HG12 | 1:A:74:LYS:N | 2.04 | 0.72 |
| 1:A:128:LEU:CD1 | 1:A:225:MET:CE | 2.68 | 0.72 |
| 1:A:306:ARG:NH1 | 1:B:280:ARG:HD3 | 2.05 | 0.71 |
| 1:B:155:ARG:NH2 | 1:B:180:GLU:HB2 | 2.07 | 0.68 |
| 1:A:280:ARG:NE | 1:B:306:ARG:HH21 | 1.92 | 0.68 |
| 1:A:280:ARG:HD3 | 1:B:306:ARG:HH21 | 1.57 | 0.68 |
| 1:A:128:LEU:CD1 | 1:A:225:MET:HE1 | 2.24 | 0.67 |
| 1:B:155:ARG:HH21 | 1:B:180:GLU:HB2 | 1.61 | 0.64 |
| 1:A:128:LEU:HD11 | 1:A:225:MET:CE | 2.29 | 0.63 |
| 1:B:36:GLN:HG3 | 1:B:104:PRO:HD3 | 1.83 | 0.59 |
| 1:A:73:VAL:CG1 | 1:A:74:LYS:N | 2.66 | 0.58 |
| 1:B:240:ASP:O | 1:B:244:ARG:HG2 | 2.04 | 0.57 |
| 1:A:132:ASP:OD1 | 1:A:169:ARG:NH2 | 2.39 | 0.56 |
| 1:A:21:ARG:HH11 | 1:A:21:ARG:CG | 2.04 | 0.55 |
| 1:B:10:ARG:NH1 | 1:B:314:THR:HG23 | 2.23 | 0.53 |
| 1:B:41:LEU:HD23 | 1:B:56:ALA:HB2 | 1.90 | 0.53 |
| 1:B:3:GLY:N | 5:B:503:HOH:O | 2.43 | 0.51 |
| 1:B:185:GLY:HA2 | 1:B:208:MET:CE | 2.40 | 0.51 |
| 1:B:185:GLY:HA2 | 1:B:208:MET:HE3 | 1.94 | 0.50 |
| 1:A:73:VAL:HG12 | 1:A:74:LYS:H | 1.76 | 0.49 |
| 4:A:408:A1H6E:CL1 | 4:A:408:A1H6E:C37 | 2.99 | 0.48 |
| 1:A:286:HIS:ND1 | 1:A:288:GLU:OE1 | 2.47 | 0.47 |
| 1:B:279:LYS:HD3 | 1:B:283:ARG:CZ | 2.44 | 0.47 |
| 1:B:53:VAL:HG22 | 1:B:68:LYS:HG2 | 1.96 | 0.47 |
| 1:A:73:VAL:CG1 | 1:A:74:LYS:H | 2.26 | 0.47 |
| 1:B:12:TYR:OH | 1:B:316:ARG:NH1 | 2.48 | 0.46 |
| 1:B:166:HIS:CD2 | 1:B:169:ARG:HH11 | 2.34 | 0.46 |
| 1:A:261:TYR:O | 1:A:262:ASN:C | 2.53 | 0.46 |
| 1:B:303:LYS:HB3 | 1:B:313:LEU:HG | 1.97 | 0.46 |
| 1:B:155:ARG:HD3 | 1:B:178:LEU:O | 2.16 | 0.45 |

*Continued on next page...*

*Continued from previous page...*

| Atom-1 | Atom-2 | Interatomic distance (Å) | Clash overlap (Å) |
|---|---|---|---|
| 1:B:224:SER:HB2 | 1:B:229:LYS:O | 2.16 | 0.45 |
| 1:B:186:GLN:HG3 | 1:B:187:GLU:N | 2.33 | 0.44 |
| 1:A:91:GLY:HA3 | 1:A:146:TYR:CE2 | 2.53 | 0.44 |
| 1:B:128:LEU:HD23 | 1:B:128:LEU:HA | 1.81 | 0.44 |
| 1:A:299:ASP:HB2 | 1:B:295:PRO:HB3 | 2.00 | 0.43 |
| 1:B:86:GLU:HG2 | 1:B:89:ARG:NH2 | 2.33 | 0.43 |
| 1:A:128:LEU:HD23 | 1:A:128:LEU:HA | 1.82 | 0.43 |
| 1:B:7:SER:H | 1:B:208:MET:HE2 | 1.83 | 0.43 |
| 1:B:78:ILE:HD13 | 1:B:109:PRO:HG2 | 2.01 | 0.43 |
| 1:B:162:VAL:O | 4:B:411:A1H6E:N6 | 2.52 | 0.43 |
| 1:A:21:ARG:HG3 | 1:A:22:GLU:N | 2.34 | 0.42 |
| 1:B:200:PRO:O | 1:B:204:VAL:HG22 | 2.18 | 0.42 |
| 1:A:21:ARG:NH1 | 1:A:22:GLU:HA | 2.35 | 0.42 |
| 1:A:265:LEU:HD12 | 1:A:265:LEU:HA | 1.76 | 0.42 |
| 1:B:91:GLY:HA3 | 1:B:146:TYR:CE2 | 2.55 | 0.42 |
| 1:B:8:ARG:NH2 | 1:B:208:MET:CE | 2.82 | 0.42 |
| 1:B:229:LYS:HE3 | 2:B:407:SO4:O3 | 2.19 | 0.42 |
| 1:B:134:ARG:HG2 | 1:B:323:TYR:CZ | 2.55 | 0.42 |
| 1:A:200:PRO:HG2 | 1:A:307:TYR:HA | 2.01 | 0.42 |
| 1:A:314:THR:HG22 | 1:A:315:ALA:N | 2.35 | 0.42 |
| 1:A:10:ARG:H | 1:A:10:ARG:HG3 | 1.58 | 0.41 |
| 1:A:6:PRO:HB3 | 1:A:8:ARG:HH22 | 1.85 | 0.41 |
| 1:A:21:ARG:HH12 | 1:A:22:GLU:HA | 1.85 | 0.41 |
| 1:A:39:TYR:HA | 1:A:57:ILE:O | 2.21 | 0.40 |
| 1:A:21:ARG:HH11 | 1:A:22:GLU:N | 2.19 | 0.40 |
| 1:B:48:GLY:O | 2:B:408:SO4:O4 | 2.39 | 0.40 |

There are no symmetry-related clashes.

#### 5.3 Torsion angles (i)

##### 5.3.1 Protein backbone (i)

In the following table, the Percentiles column shows the percent Ramachandran outliers of the chain as a percentile score with respect to all X-ray entries followed by that with respect to entries of similar resolution.

The Analysed column shows the number of residues for which the backbone conformation was analysed, and the total number of residues.

For Manuscript Review

| Mol | Chain | Analysed | Favoured | Allowed | Outliers | Percentiles | |
|---|---|---|---|---|---|---|---|
| 1 | A | 327/338 (97%) | 304 (93%) | 23 (7%) | 0 | 100 | 100 |
| 1 | B | 327/338 (97%) | 312 (95%) | 15 (5%) | 0 | 100 | 100 |
| All | All | 654/676 (97%) | 616 (94%) | 38 (6%) | 0 | 100 | 100 |

There are no Ramachandran outliers to report.

##### 5.3.2 Protein sidechains (i)

In the following table, the Percentiles column shows the percent sidechain outliers of the chain as a percentile score with respect to all X-ray entries followed by that with respect to entries of similar resolution.

The Analysed column shows the number of residues for which the sidechain conformation was analysed, and the total number of residues.

| Mol | Chain | Analysed | Rotameric | Outliers | Percentiles | |
|---|---|---|---|---|---|---|
| 1 | A | 283/310 (91%) | 269 (95%) | 14 (5%) | 25 | 48 |
| 1 | B | 289/310 (93%) | 274 (95%) | 15 (5%) | 23 | 46 |
| All | All | 572/620 (92%) | 543 (95%) | 29 (5%) | 24 | 46 |

All (29) residues with a non-rotameric sidechain are listed below:

| Mol | Chain | Res | Type |
|---|---|---|---|
| 1 | A | 10 | ARG |
| 1 | A | 21 | ARG |
| 1 | A | 30 | VAL |
| 1 | A | 36 | GLN |
| 1 | A | 47 | ARG |
| 1 | A | 65 | VAL |
| 1 | A | 118 | ASN |
| 1 | A | 153 | MET |
| 1 | A | 234 | HIS |
| 1 | A | 265 | LEU |
| 1 | A | 296 | GLU |
| 1 | A | 306 | ARG |
| 1 | A | 316 | ARG |
| 1 | A | 327 | VAL |
| 1 | B | 10 | ARG |
| 1 | B | 21 | ARG |
| 1 | B | 30 | VAL |
| 1 | B | 65 | VAL |

*Continued on next page...*

For Manuscript Review

*Continued from previous page...*

| Mol | Chain | Res | Type |
|---|---|---|---|
| 1 | B | 74 | LYS |
| 1 | B | 89 | ARG |
| 1 | B | 123 | GLN |
| 1 | B | 170 | LYS |
| 1 | B | 258 | ILE |
| 1 | B | 264 | GLU |
| 1 | B | 279 | LYS |
| 1 | B | 283 | ARG |
| 1 | B | 287 | SER |
| 1 | B | 296 | GLU |
| 1 | B | 306 | ARG |

Sometimes sidechains can be flipped to improve hydrogen bonding and reduce clashes. All (1) such sidechains are listed below:

| Mol | Chain | Res | Type |
|---|---|---|---|
| 1 | B | 123 | GLN |

###### 5.3.3 RNA (i)

There are no RNA molecules in this entry.

##### 5.4 Non-standard residues in protein, DNA, RNA chains (i)

There are no non-standard protein/DNA/RNA residues in this entry.

##### 5.5 Carbohydrates (i)

There are no monosaccharides in this entry.

##### 5.6 Ligand geometry (i)

19 ligands are modelled in this entry.

In the following table, the Counts columns list the number of bonds (or angles) for which Mogul statistics could be retrieved, the number of bonds (or angles) that are observed in the model and the number of bonds (or angles) that are defined in the Chemical Component Dictionary. The Link column lists molecule types, if any, to which the group is linked. The Z score for a bond length (or angle) is the number of standard deviations the observed value is removed from the expected value. A bond length (or angle) with $|Z| > 2$ is considered an outlier worth inspection. RMSZ is the root-mean-square of all Z scores of the bond lengths (or angles).

For Manuscript Review

| Mol | Type | Chain | Res | Link | Bond lengths | | | Bond angles | | |
|---|---|---|---|---|---|---|---|---|---|---|
| | | | | | Counts | RMSZ | #\|Z\| > 2 | Counts | RMSZ | #\|Z\| > 2 |
| 2 | SO4 | B | 408 | - | 4,4,4 | 0.36 | 0 | 6,6,6 | 0.18 | 0 |
| 2 | SO4 | B | 401 | - | 4,4,4 | 0.33 | 0 | 6,6,6 | 0.30 | 0 |
| 2 | SO4 | A | 401 | - | 4,4,4 | 0.30 | 0 | 6,6,6 | 0.20 | 0 |
| 2 | SO4 | B | 403 | - | 4,4,4 | 0.31 | 0 | 6,6,6 | 0.10 | 0 |
| 2 | SO4 | B | 404 | - | 4,4,4 | 0.27 | 0 | 6,6,6 | 0.27 | 0 |
| 2 | SO4 | A | 406 | - | 4,4,4 | 0.34 | 0 | 6,6,6 | 0.25 | 0 |
| 2 | SO4 | A | 405 | - | 4,4,4 | 0.34 | 0 | 6,6,6 | 0.18 | 0 |
| 2 | SO4 | B | 406 | - | 4,4,4 | 0.32 | 0 | 6,6,6 | 0.11 | 0 |
| 2 | SO4 | B | 410 | - | 4,4,4 | 0.38 | 0 | 6,6,6 | 0.17 | 0 |
| 2 | SO4 | B | 409 | - | 4,4,4 | 0.29 | 0 | 6,6,6 | 0.31 | 0 |
| 3 | EDO | A | 407 | - | 3,3,3 | 0.17 | 0 | 2,2,2 | 0.09 | 0 |
| 4 | A1H6E | A | 408 | - | 36,42,42 | 0.47 | 0 | 46,63,63 | 0.75 | 1 (2%) |
| 4 | A1H6E | B | 411 | - | 36,42,42 | 0.48 | 0 | 46,63,63 | 0.74 | 1 (2%) |
| 2 | SO4 | A | 403 | - | 4,4,4 | 0.37 | 0 | 6,6,6 | 0.16 | 0 |
| 2 | SO4 | B | 402 | - | 4,4,4 | 0.28 | 0 | 6,6,6 | 0.11 | 0 |
| 2 | SO4 | A | 404 | - | 4,4,4 | 0.32 | 0 | 6,6,6 | 0.10 | 0 |
| 2 | SO4 | A | 402 | - | 4,4,4 | 0.32 | 0 | 6,6,6 | 0.13 | 0 |
| 2 | SO4 | B | 407 | - | 4,4,4 | 0.38 | 0 | 6,6,6 | 0.09 | 0 |
| 2 | SO4 | B | 405 | - | 4,4,4 | 0.33 | 0 | 6,6,6 | 0.18 | 0 |

In the following table, the Chirals column lists the number of chiral outliers, the number of chiral centers analysed, the number of these observed in the model and the number defined in the Chemical Component Dictionary. Similar counts are reported in the Torsion and Rings columns. '-' means no outliers of that kind were identified.

| Mol | Type | Chain | Res | Link | Chirals | Torsions | Rings |
|---|---|---|---|---|---|---|---|
| 4 | A1H6E | B | 411 | - | - | 0/19/43/43 | 0/6/6/6 |
| 3 | EDO | A | 407 | - | - | 1/1/1/1 | - |
| 4 | A1H6E | A | 408 | - | - | 0/19/43/43 | 0/6/6/6 |

There are no bond length outliers.

All (2) bond angle outliers are listed below:

| Mol | Chain | Res | Type | Atoms | Z | Observed(°) | Ideal(°) |
|---|---|---|---|---|---|---|---|
| 4 | B | 411 | A1H6E | C16-C21-N20 | 2.96 | 114.93 | 108.04 |
| 4 | A | 408 | A1H6E | C16-C21-N20 | 2.93 | 114.87 | 108.04 |

There are no chirality outliers.

All (1) torsion outliers are listed below:

For Manuscript Review

| Mol | Chain | Res | Type | Atoms |
|---|---|---|---|---|
| 3 | A | 407 | EDO | O1-C1-C2-O2 |

There are no ring outliers.

4 monomers are involved in 4 short contacts:

| Mol | Chain | Res | Type | Clashes | Symm-Clashes |
|---|---|---|---|---|---|
| 2 | B | 408 | SO4 | 1 | 0 |
| 4 | A | 408 | A1H6E | 1 | 0 |
| 4 | B | 411 | A1H6E | 1 | 0 |
| 2 | B | 407 | SO4 | 1 | 0 |

The following is a two-dimensional graphical depiction of Mogul quality analysis of bond lengths, bond angles, torsion angles, and ring geometry for all instances of the Ligand of Interest. In addition, ligands with molecular weight > 250 and outliers as shown on the validation Tables will also be included. For torsion angles, if less then 5% of the Mogul distribution of torsion angles is within 10 degrees of the torsion angle in question, then that torsion angle is considered an outlier. Any bond that is central to one or more torsion angles identified as an outlier by Mogul will be highlighted in the graph. For rings, the root-mean-square deviation (RMSD) between the ring in question and similar rings identified by Mogul is calculated over all ring torsion angles. If the average RMSD is greater than 60 degrees and the minimal RMSD between the ring in question and any Mogul-identified rings is also greater than 60 degrees, then that ring is considered an outlier. The outliers are highlighted in purple. The color gray indicates Mogul did not find sufficient equivalents in the CSD to analyse the geometry.

**Ligand A1H6E A 408**

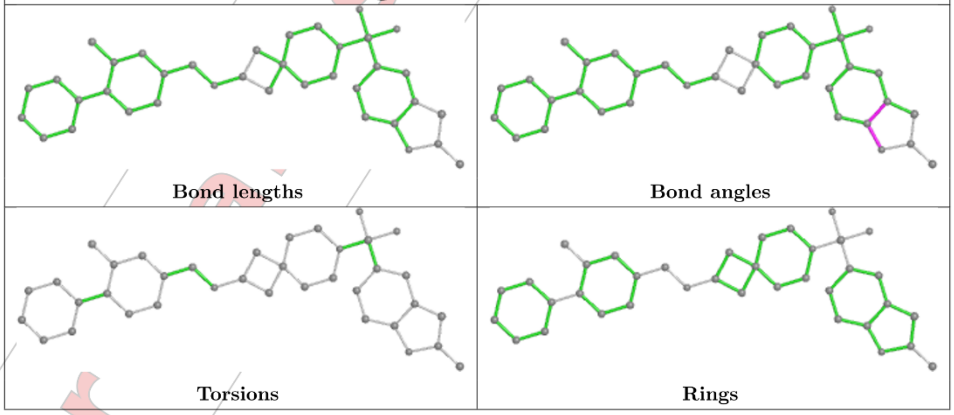

For Manuscript Review

**Ligand A1H6E B 411**

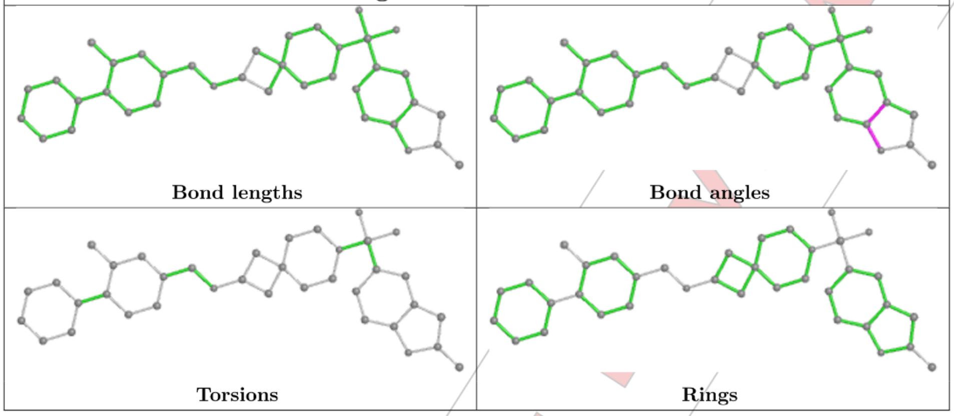

#### 5.7 Other polymers (i)

There are no such residues in this entry.

#### 5.8 Polymer linkage issues (i)

There are no chain breaks in this entry.

For Manuscript Review

### 6 Fit of model and data (i)

#### 6.1 Protein, DNA and RNA chains (i)

In the following table, the column labelled '#RSRZ> 2' contains the number (and percentage) of RSRZ outliers, followed by percent RSRZ outliers for the chain as percentile scores relative to all X-ray entries and entries of similar resolution. The OWAB column contains the minimum, median, $95^{th}$ percentile and maximum values of the occupancy-weighted average B-factor per residue. The column labelled 'Q< 0.9' lists the number of (and percentage) of residues with an average occupancy less than 0.9.

| Mol | Chain | Analysed | <RSRZ> | #RSRZ>2 | | | OWAB(Å$^2$) | Q<0.9 |
|---|---|---|---|---|---|---|---|---|
| 1 | A | 329/338 (97%) | -0.22 | 3 (0%) | 84 | 82 | 29, 48, 77, 108 | 0 |
| 1 | B | 329/338 (97%) | -0.10 | 6 (1%) | 68 | 64 | 30, 49, 86, 115 | 0 |
| All | All | 658/676 (97%) | -0.16 | 9 (1%) | 75 | 71 | 29, 49, 84, 115 | 0 |

All (9) RSRZ outliers are listed below:

| Mol | Chain | Res | Type | RSRZ |
|---|---|---|---|---|
| 1 | B | 330 | ASP | 4.2 |
| 1 | B | 127 | THR | 3.7 |
| 1 | A | 330 | ASP | 3.0 |
| 1 | B | 124 | LEU | 2.7 |
| 1 | B | 255 | TYR | 2.5 |
| 1 | A | 105 | VAL | 2.3 |
| 1 | A | 329 | LYS | 2.3 |
| 1 | B | 128 | LEU | 2.2 |
| 1 | B | 266 | ASP | 2.1 |

#### 6.2 Non-standard residues in protein, DNA, RNA chains (i)

There are no non-standard protein/DNA/RNA residues in this entry.

#### 6.3 Carbohydrates (i)

There are no monosaccharides in this entry.

#### 6.4 Ligands (i)

In the following table, the Atoms column lists the number of modelled atoms in the group and the number defined in the chemical component dictionary. The B-factors column lists the minimum,

For Manuscript Review

median, $95^{th}$ percentile and maximum values of B factors of atoms in the group. The column labelled 'Q< 0.9' lists the number of atoms with occupancy less than 0.9.

| Mol | Type | Chain | Res | Atoms | RSCC | RSR | B-factors($Å^2$) | Q<0.9 |
|---|---|---|---|---|---|---|---|---|
| 2 | SO4 | B | 405 | 5/5 | 0.68 | 0.23 | 113,122,123,124 | 0 |
| 2 | SO4 | B | 409 | 5/5 | 0.76 | 0.18 | 66,78,94,95 | 0 |
| 2 | SO4 | A | 405 | 5/5 | 0.82 | 0.18 | 74,76,86,105 | 0 |
| 2 | SO4 | B | 404 | 5/5 | 0.84 | 0.19 | 68,71,99,100 | 0 |
| 2 | SO4 | A | 404 | 5/5 | 0.84 | 0.17 | 89,97,102,108 | 0 |
| 2 | SO4 | B | 403 | 5/5 | 0.84 | 0.19 | 92,106,109,110 | 0 |
| 3 | EDO | A | 407 | 4/4 | 0.84 | 0.17 | 62,68,68,72 | 0 |
| 2 | SO4 | B | 410 | 5/5 | 0.88 | 0.19 | 60,79,83,83 | 0 |
| 2 | SO4 | B | 406 | 5/5 | 0.88 | 0.27 | 69,95,100,107 | 0 |
| 4 | A1H6E | B | 411 | 37/37 | 0.89 | 0.22 | 42,53,89,112 | 0 |
| 2 | SO4 | B | 407 | 5/5 | 0.91 | 0.30 | 93,103,110,126 | 0 |
| 4 | A1H6E | A | 408 | 37/37 | 0.92 | 0.20 | 39,50,82,91 | 0 |
| 2 | SO4 | B | 408 | 5/5 | 0.94 | 0.14 | 70,77,81,86 | 0 |
| 2 | SO4 | B | 402 | 5/5 | 0.96 | 0.08 | 63,65,79,83 | 0 |
| 2 | SO4 | A | 402 | 5/5 | 0.96 | 0.14 | 61,70,71,81 | 0 |
| 2 | SO4 | A | 401 | 5/5 | 0.97 | 0.11 | 44,48,57,59 | 0 |
| 2 | SO4 | A | 406 | 5/5 | 0.97 | 0.12 | 47,50,59,62 | 0 |
| 2 | SO4 | B | 401 | 5/5 | 0.97 | 0.12 | 49,58,61,63 | 0 |
| 2 | SO4 | A | 403 | 5/5 | 0.98 | 0.11 | 47,56,62,75 | 0 |

The following is a graphical depiction of the model fit to experimental electron density of all instances of the Ligand of Interest. In addition, ligands with molecular weight > 250 and outliers as shown on the geometry validation Tables will also be included. Each fit is shown from different orientation to approximate a three-dimensional view.

For Manuscript Review

**Electron density around A1H6E B 411:**

$2mF_o$-$DF_c$ (at 0.7 rmsd) in gray
$mF_o$-$DF_c$ (at 3 rmsd) in purple (negative)
and green (positive)

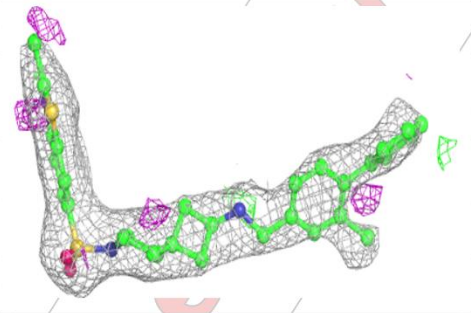

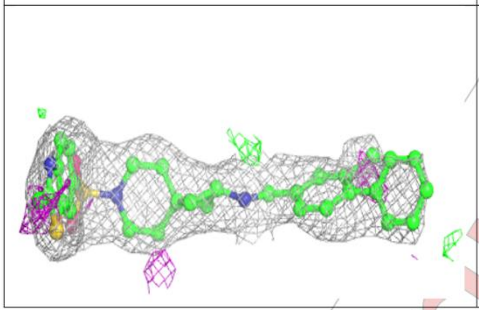

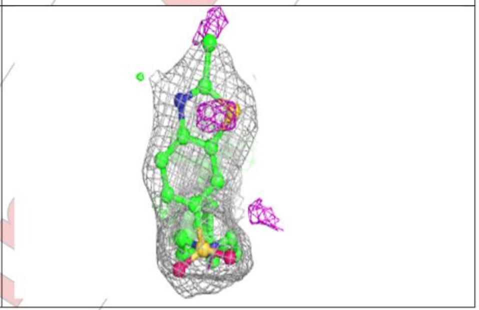

**Electron density around A1H6E A 408:**

$2mF_o$-$DF_c$ (at 0.7 rmsd) in gray
$mF_o$-$DF_c$ (at 3 rmsd) in purple (negative)
and green (positive)

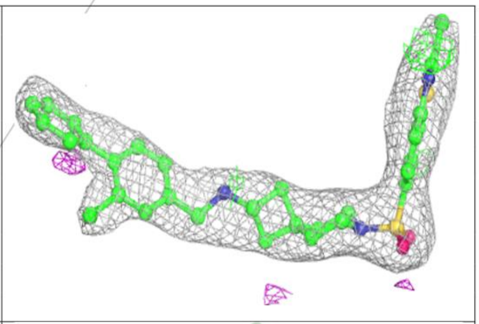

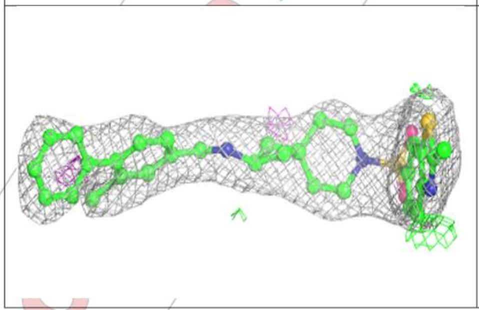

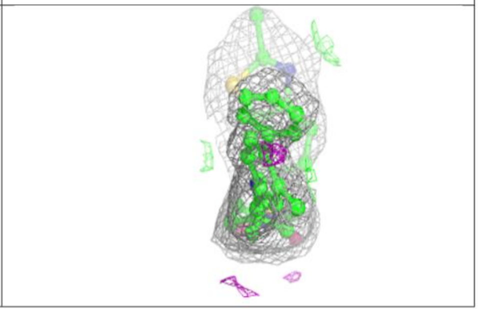

#### 6.5 Other polymers (i)

There are no such residues in this entry.

For Manuscript Review

### Full wwPDB X-ray Structure Validation Report ⓘ

Mar 20, 2024 – 02:50 pm GMT

| | | |
|---|---|---|
| PDB ID | : | 9EPY |
| Title | : | Crystal structure of human Casein Kinase II subunit alpha (CK2a1) in complex with allosteric ligand FGC3330 |
| Deposited on | : | 2024-03-20 |
| Resolution | : | 2.65 Å(reported) |

**This wwPDB validation report is for manuscript review**

For Manuscript Review

This is a Full wwPDB X-ray Structure Validation Report.

This report is produced by the wwPDB biocuration pipeline after annotation of the structure.

We welcome your comments at **
A user guide is available at
https://www.wwpdb.org/validation/2017/XrayValidationReportHelp
with specific help available everywhere you see the ⓘ symbol.

The types of validation reports are described at
http://www.wwpdb.org/validation/2017/FAQs#types.

The following versions of software and data (see references ⓘ) were used in the production of this report:

| | | |
|---|---|---|
| MolProbity | : | 4.02b-467 |
| Mogul | : | 1.8.4, CSD as541be (2020) |
| Xtriage (Phenix) | : | 1.13 |
| EDS | : | 2.36 |
| buster-report | : | 1.1.7 (2018) |
| Percentile statistics | : | 20191225.v01 (using entries in the PDB archive December 25th 2019) |
| Refmac | : | 5.8.0158 |
| CCP4 | : | 7.0.044 (Gargrove) |
| Ideal geometry (proteins) | : | Engh & Huber (2001) |

### 1 Overall quality at a glance (i)

The following experimental techniques were used to determine the structure:
*X-RAY DIFFRACTION*

The reported resolution of this entry is 2.65 Å.

Percentile scores (ranging between 0-100) for global validation metrics of the entry are shown in the following graphic. The table shows the number of entries on which the scores are based.

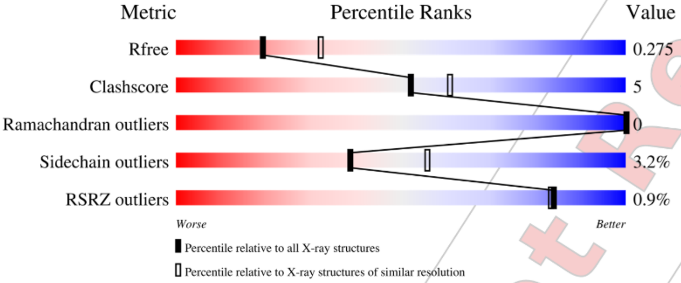

| Metric | Whole archive (#Entries) | Similar resolution (#Entries, resolution range(Å)) |
|---|---|---|
| $R_{free}$ | 130704 | 1332 (2.68-2.64) |
| Clashscore | 141614 | 1374 (2.68-2.64) |
| Ramachandran outliers | 138981 | 1349 (2.68-2.64) |
| Sidechain outliers | 138945 | 1349 (2.68-2.64) |
| RSRZ outliers | 127900 | 1318 (2.68-2.64) |

The table below summarises the geometric issues observed across the polymeric chains and their fit to the electron density. The red, orange, yellow and green segments of the lower bar indicate the fraction of residues that contain outliers for >=3, 2, 1 and 0 types of geometric quality criteria respectively. A grey segment represents the fraction of residues that are not modelled. The numeric value for each fraction is indicated below the corresponding segment, with a dot representing fractions <=5% The upper red bar (where present) indicates the fraction of residues that have poor fit to the electron density. The numeric value is given above the bar.

| Mol | Chain | Length | Quality of chain |
|---|---|---|---|
| 1 | A | 338 | % 86% 10% • • |
| 1 | B | 338 | % 83% 13% • • |

The following table lists non-polymeric compounds, carbohydrate monomers and non-standard

Ideal geometry (DNA, RNA) : Parkinson et al. (1996)
Validation Pipeline (wwPDB-VP) : 2.36

For Manuscript Review

residues in protein, DNA, RNA chains that are outliers for geometric or electron-density-fit criteria:

| Mol | Type | Chain | Res | Chirality | Geometry | Clashes | Electron density |
|---|---|---|---|---|---|---|---|
| 2 | SO4 | A | 405 | - | - | - | X |

For Manuscript Review

#### 2 Entry composition (i)

There are 5 unique types of molecules in this entry. The entry contains 5530 atoms, of which 0 are hydrogens and 0 are deuteriums.

In the tables below, the ZeroOcc column contains the number of atoms modelled with zero occupancy, the AltConf column contains the number of residues with at least one atom in alternate conformation and the Trace column contains the number of residues modelled with at most 2 atoms.

- Molecule 1 is a protein called Casein kinase II subunit alpha.

| Mol | Chain | Residues | Atoms | | | | | ZeroOcc | AltConf | Trace |
|---|---|---|---|---|---|---|---|---|---|---|
| | | | Total | C | N | O | S | | | |
| 1 | A | 329 | 2667 | 1718 | 460 | 478 | 11 | 0 | 0 | 0 |
| 1 | B | 329 | 2676 | 1715 | 466 | 484 | 11 | 0 | 0 | 0 |

There are 2 discrepancies between the modelled and reference sequences:

| Chain | Residue | Modelled | Actual | Comment | Reference |
|---|---|---|---|---|---|
| A | 0 | SER | - | expression tag | UNP P68400 |
| B | 0 | SER | - | expression tag | UNP P68400 |

- Molecule 2 is SULFATE ION (three-letter code: SO4) (formula: $O_4S$).

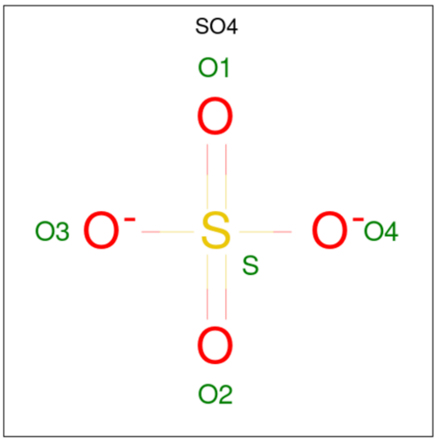

| Mol | Chain | Residues | Atoms | | | ZeroOcc | AltConf |
|---|---|---|---|---|---|---|---|
| | | | Total | O | S | | |
| 2 | A | 1 | 5 | 4 | 1 | 0 | 0 |

Continued on next page...

For Manuscript Review

*Continued from previous page...*

| Mol | Chain | Residues | Atoms | ZeroOcc | AltConf |
|---|---|---|---|---|---|
| 2 | A | 1 | Total O S<br>5 4 1 | 0 | 0 |
| 2 | A | 1 | Total O S<br>5 4 1 | 0 | 0 |
| 2 | A | 1 | Total O S<br>5 4 1 | 0 | 0 |
| 2 | A | 1 | Total O S<br>5 4 1 | 0 | 0 |
| 2 | A | 1 | Total O S<br>5 4 1 | 0 | 0 |
| 2 | A | 1 | Total O S<br>5 4 1 | 0 | 0 |
| 2 | A | 1 | Total O S<br>5 4 1 | 0 | 0 |
| 2 | A | 1 | Total O S<br>5 4 1 | 0 | 0 |
| 2 | B | 1 | Total O S<br>5 4 1 | 0 | 0 |
| 2 | B | 1 | Total O S<br>5 4 1 | 0 | 0 |
| 2 | B | 1 | Total O S<br>5 4 1 | 0 | 0 |
| 2 | B | 1 | Total O S<br>5 4 1 | 0 | 0 |
| 2 | B | 1 | Total O S<br>5 4 1 | 0 | 0 |
| 2 | B | 1 | Total O S<br>5 4 1 | 0 | 0 |
| 2 | B | 1 | Total O S<br>5 4 1 | 0 | 0 |
| 2 | B | 1 | Total O S<br>5 4 1 | 0 | 0 |

- Molecule 3 is 1,2-ETHANEDIOL (three-letter code: EDO) (formula: $C_2H_6O_2$).

For Manuscript Review

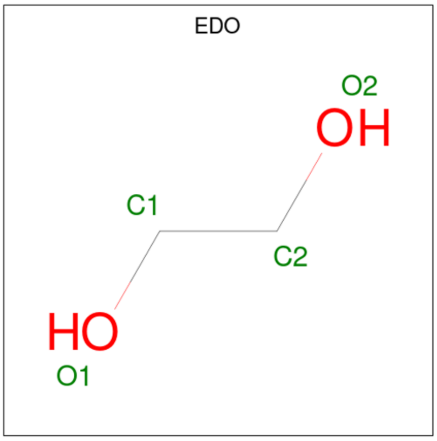

| Mol | Chain | Residues | Atoms | ZeroOcc | AltConf |
|---|---|---|---|---|---|
| 3 | A | 1 | Total C O<br>4 2 2 | 0 | 0 |
| 3 | B | 1 | Total C O<br>4 2 2 | 0 | 0 |

- Molecule 4 is {N}-[(3-chloranyl-4-phenyl-phenyl)methyl]-7-[(2-methyl-1,3-benzothiazol-6-yl)sulfonyl]-7-azaspiro[3.5]nonan-2-amine (three-letter code: A1H6G) (formula: $C_{29}H_{30}ClN_3O_2S_2$) (labeled as "Ligand of Interest" by depositor).

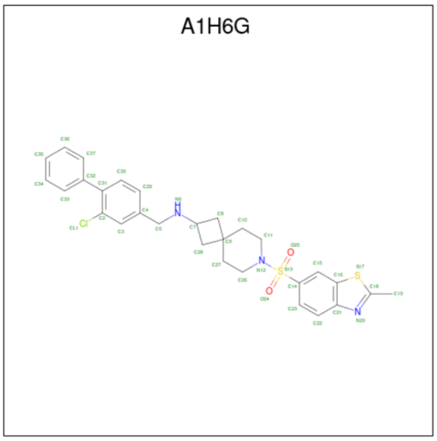

| Mol | Chain | Residues | Atoms | ZeroOcc | AltConf |
|---|---|---|---|---|---|
| 4 | A | 1 | Total C Cl N O S<br>37 29 1 3 2 2 | 0 | 0 |

Continued on next page...

For Manuscript Review

*Continued from previous page...*

| Mol | Chain | Residues | Atoms | | | | | | ZeroOcc | AltConf |
|---|---|---|---|---|---|---|---|---|---|---|
| | | | Total | C | Cl | N | O | S | | |
| 4 | B | 1 | 37 | 29 | 1 | 3 | 2 | 2 | 0 | 0 |

- Molecule 5 is water.

| Mol | Chain | Residues | Atoms | | ZeroOcc | AltConf |
|---|---|---|---|---|---|---|
| 5 | A | 8 | Total 8 | O 8 | 0 | 0 |
| 5 | B | 12 | Total 12 | O 12 | 0 | 0 |

For Manuscript Review

### 3 Residue-property plots ⓘ

These plots are drawn for all protein, RNA, DNA and oligosaccharide chains in the entry. The first graphic for a chain summarises the proportions of the various outlier classes displayed in the second graphic. The second graphic shows the sequence view annotated by issues in geometry and electron density. Residues are color-coded according to the number of geometric quality criteria for which they contain at least one outlier: green = 0, yellow = 1, orange = 2 and red = 3 or more. A red dot above a residue indicates a poor fit to the electron density (RSRZ > 2). Stretches of 2 or more consecutive residues without any outlier are shown as a green connector. Residues present in the sample, but not in the model, are shown in grey.

- Molecule 1: Casein kinase II subunit alpha

Chain A:

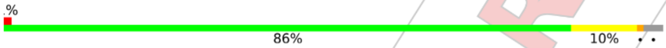

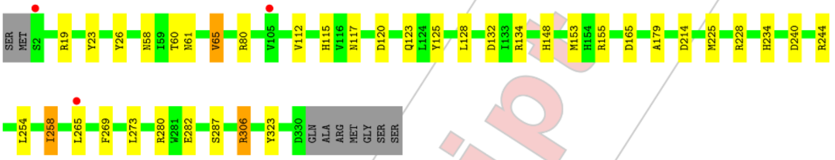

- Molecule 1: Casein kinase II subunit alpha

Chain B:

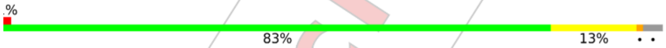

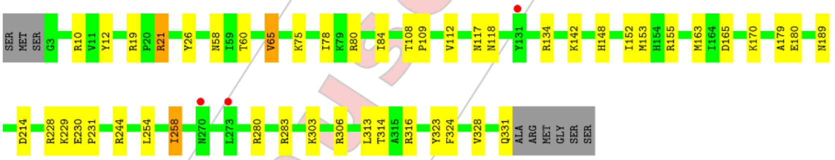

For Manuscript Review

#### 4 Data and refinement statistics (i)

| Property | Value | Source |
|---|---|---|
| Space group | P 43 21 2 | Depositor |
| Cell constants<br>a, b, c, $\alpha$, $\beta$, $\gamma$ | 126.64Å 126.64Å 124.25Å<br>90.00° 90.00° 90.00° | Depositor |
| Resolution (Å) | 44.82 – 2.65<br>44.78 – 2.65 | Depositor<br>EDS |
| % Data completeness<br>(in resolution range) | 99.9 (44.82-2.65)<br>100.0 (44.78-2.65) | Depositor<br>EDS |
| $R_{merge}$ | (Not available) | Depositor |
| $R_{sym}$ | (Not available) | Depositor |
| $< I/\sigma(I) >$ [1] | 1.81 (at 2.65Å) | Xtriage |
| Refinement program | REFMAC 5.8.0419 | Depositor |
| R, $R_{free}$ | 0.226 , 0.269<br>0.237 , 0.275 | Depositor<br>DCC |
| $R_{free}$ test set | 1480 reflections (4.94%) | wwPDB-VP |
| Wilson B-factor (Å$^2$) | 65.4 | Xtriage |
| Anisotropy | 0.503 | Xtriage |
| Bulk solvent $k_{sol}$(e/Å$^3$), $B_{sol}$(Å$^2$) | 0.33 , 46.7 | EDS |
| L-test for twinning[2] | $< \|L\| > = 0.48$, $< L^2 > = 0.31$ | Xtriage |
| Estimated twinning fraction | 0.008 for -h,-l,-k<br>0.000 for l,-k,h | Xtriage |
| $F_o$,$F_c$ correlation | 0.94 | EDS |
| Total number of atoms | 5530 | wwPDB-VP |
| Average B, all atoms (Å$^2$) | 77.0 | wwPDB-VP |

Xtriage's analysis on translational NCS is as follows: *The analyses of the Patterson function reveals a significant off-origin peak that is 25.16 % of the origin peak, indicating pseudo-translational symmetry. The chance of finding a peak of this or larger height randomly in a structure without pseudo-translational symmetry is equal to 3.3427e-03. The detected translational NCS is most likely also responsible for the elevated intensity ratio.*

[1] Intensities estimated from amplitudes.
[2] Theoretical values of $< |L| >$, $< L^2 >$ for acentric reflections are 0.5, 0.333 respectively for untwinned datasets, and 0.375, 0.2 for perfectly twinned datasets.

For Manuscript Review

### 5 Model quality (i)

#### 5.1 Standard geometry (i)

Bond lengths and bond angles in the following residue types are not validated in this section: A1H6G, SO4, EDO

The Z score for a bond length (or angle) is the number of standard deviations the observed value is removed from the expected value. A bond length (or angle) with $|Z| > 5$ is considered an outlier worth inspection. RMSZ is the root-mean-square of all Z scores of the bond lengths (or angles).

| Mol | Chain | Bond lengths | | Bond angles | |
|---|---|---|---|---|---|
| | | RMSZ | $\#\|Z\|>5$ | RMSZ | $\#\|Z\|>5$ |
| 1 | A | 0.39 | 0/2741 | 0.73 | 0/3726 |
| 1 | B | 0.39 | 0/2749 | 0.72 | 0/3736 |
| All | All | 0.39 | 0/5490 | 0.73 | 0/7462 |

Chiral center outliers are detected by calculating the chiral volume of a chiral center and verifying if the center is modelled as a planar moiety or with the opposite hand.A planarity outlier is detected by checking planarity of atoms in a peptide group, atoms in a mainchain group or atoms of a sidechain that are expected to be planar.

| Mol | Chain | #Chirality outliers | #Planarity outliers |
|---|---|---|---|
| 1 | A | 0 | 1 |

There are no bond length outliers.

There are no bond angle outliers.

There are no chirality outliers.

All (1) planarity outliers are listed below:

| Mol | Chain | Res | Type | Group |
|---|---|---|---|---|
| 1 | A | 155 | ARG | Sidechain |

#### 5.2 Too-close contacts (i)

In the following table, the Non-H and H(model) columns list the number of non-hydrogen atoms and hydrogen atoms in the chain respectively. The H(added) column lists the number of hydrogen atoms added and optimized by MolProbity. The Clashes column lists the number of clashes within the asymmetric unit, whereas Symm-Clashes lists symmetry-related clashes.

For Manuscript Review

| Mol | Chain | Non-H | H(model) | H(added) | Clashes | Symm-Clashes |
|---|---|---|---|---|---|---|
| 1 | A | 2667 | 0 | 2527 | 28 | 0 |
| 1 | B | 2676 | 0 | 2524 | 37 | 0 |
| 2 | A | 45 | 0 | 0 | 0 | 0 |
| 2 | B | 40 | 0 | 0 | 1 | 0 |
| 3 | A | 4 | 0 | 6 | 0 | 0 |
| 3 | B | 4 | 0 | 6 | 2 | 0 |
| 4 | A | 37 | 0 | 0 | 1 | 0 |
| 4 | B | 37 | 0 | 0 | 0 | 0 |
| 5 | A | 8 | 0 | 0 | 1 | 0 |
| 5 | B | 12 | 0 | 0 | 0 | 0 |
| All | All | 5530 | 0 | 5063 | 58 | 0 |

The all-atom clashscore is defined as the number of clashes found per 1000 atoms (including hydrogen atoms). The all-atom clashscore for this structure is 5.

All (58) close contacts within the same asymmetric unit are listed below, sorted by their clash magnitude.

| Atom-1 | Atom-2 | Interatomic distance (Å) | Clash overlap (Å) |
|---|---|---|---|
| 1:A:280:ARG:HD2 | 1:B:306:ARG:HH21 | 1.37 | 0.90 |
| 1:A:280:ARG:HD2 | 1:B:306:ARG:NH2 | 2.06 | 0.70 |
| 1:B:118:ASN:HD21 | 1:B:163:MET:HG2 | 1.57 | 0.69 |
| 1:B:10:ARG:NH1 | 1:B:314:THR:HG23 | 2.07 | 0.69 |
| 1:B:108:THR:HB | 1:B:109:PRO:HD2 | 1.75 | 0.68 |
| 1:B:230:GLU:HB3 | 1:B:231:PRO:HD3 | 1.77 | 0.67 |
| 1:A:306:ARG:NH2 | 1:B:280:ARG:HD3 | 2.12 | 0.64 |
| 1:A:306:ARG:NH2 | 1:B:280:ARG:CD | 2.61 | 0.64 |
| 1:B:142:LYS:NZ | 1:B:331:GLN:HE22 | 1.98 | 0.62 |
| 1:B:148:HIS:CD2 | 3:B:408:EDO:H12 | 2.34 | 0.61 |
| 1:A:58:ASN:OD1 | 1:A:60:THR:HB | 2.00 | 0.61 |
| 1:B:58:ASN:OD1 | 1:B:60:THR:HB | 2.01 | 0.60 |
| 1:A:80:ARG:HD3 | 1:A:179:ALA:O | 2.01 | 0.59 |
| 1:A:115:HIS:HE1 | 1:A:117:ASN:OD1 | 1.86 | 0.59 |
| 1:A:280:ARG:CD | 1:B:306:ARG:NH2 | 2.66 | 0.59 |
| 1:A:282:GLU:CD | 1:B:306:ARG:HH22 | 2.06 | 0.59 |
| 1:A:282:GLU:OE1 | 1:B:306:ARG:NH2 | 2.37 | 0.58 |
| 1:B:148:HIS:CD2 | 3:B:408:EDO:C1 | 2.87 | 0.58 |
| 1:B:65:VAL:CG1 | 1:B:112:VAL:HG13 | 2.34 | 0.57 |
| 1:B:118:ASN:ND2 | 1:B:163:MET:HG2 | 2.18 | 0.57 |
| 1:A:80:ARG:HD2 | 5:A:504:HOH:O | 2.05 | 0.56 |
| 1:B:80:ARG:HD3 | 1:B:179:ALA:O | 2.06 | 0.56 |
| 1:A:65:VAL:CG1 | 1:A:112:VAL:HG13 | 2.36 | 0.55 |
| 1:B:142:LYS:HZ2 | 1:B:331:GLN:HE22 | 1.55 | 0.54 |

*Continued on next page...*

For Manuscript Review

*Continued from previous page...*

| Atom-1 | Atom-2 | Interatomic distance (Å) | Clash overlap (Å) |
|---|---|---|---|
| 1:B:324:PHE:O | 1:B:328:VAL:HG13 | 2.07 | 0.53 |
| 1:B:12:TYR:OH | 1:B:316:ARG:NH1 | 2.41 | 0.53 |
| 1:A:254:LEU:O | 1:A:258:ILE:HG23 | 2.10 | 0.52 |
| 1:B:254:LEU:O | 1:B:258:ILE:HG23 | 2.10 | 0.52 |
| 1:A:265:LEU:HD22 | 1:A:269:PHE:CD2 | 2.45 | 0.51 |
| 1:B:108:THR:HB | 1:B:109:PRO:CD | 2.40 | 0.51 |
| 1:B:148:HIS:HE1 | 1:B:214:ASP:OD2 | 1.93 | 0.51 |
| 1:A:19:ARG:HD2 | 1:A:23:TYR:CG | 2.47 | 0.50 |
| 1:B:65:VAL:HG11 | 1:B:112:VAL:HG13 | 1.93 | 0.50 |
| 1:A:148:HIS:HE1 | 1:A:214:ASP:OD2 | 1.95 | 0.48 |
| 1:A:65:VAL:HG11 | 1:A:112:VAL:HG13 | 1.94 | 0.48 |
| 1:B:26:TYR:CE2 | 1:B:80:ARG:HG3 | 2.49 | 0.48 |
| 1:A:125:TYR:CE2 | 1:A:228:ARG:NH1 | 2.82 | 0.47 |
| 1:A:26:TYR:CE2 | 1:A:80:ARG:HG3 | 2.49 | 0.47 |
| 1:A:128:LEU:O | 1:A:132:ASP:HB2 | 2.14 | 0.47 |
| 1:A:19:ARG:HD2 | 1:A:23:TYR:CD2 | 2.51 | 0.46 |
| 1:B:303:LYS:HB3 | 1:B:313:LEU:HG | 1.99 | 0.45 |
| 1:B:155:ARG:NH2 | 1:B:180:GLU:HB2 | 2.32 | 0.43 |
| 1:A:265:LEU:HD23 | 1:A:265:LEU:HA | 1.86 | 0.43 |
| 1:B:117:ASN:O | 1:B:165:ASP:HB2 | 2.19 | 0.43 |
| 1:A:306:ARG:HH22 | 1:B:280:ARG:HD3 | 1.83 | 0.42 |
| 4:A:409:A1H6G:C37 | 4:A:409:A1H6G:CL1 | 3.04 | 0.42 |
| 1:B:134:ARG:HG2 | 1:B:323:TYR:CZ | 2.55 | 0.42 |
| 1:B:65:VAL:HG13 | 1:B:112:VAL:HG13 | 2.00 | 0.42 |
| 1:B:84:ILE:HG12 | 1:B:152:ILE:HD12 | 2.01 | 0.42 |
| 1:B:21:ARG:HE | 1:B:21:ARG:HB3 | 1.63 | 0.42 |
| 1:B:19:ARG:NH1 | 1:B:19:ARG:HG2 | 2.35 | 0.42 |
| 1:A:134:ARG:HG2 | 1:A:323:TYR:CZ | 2.55 | 0.41 |
| 1:B:75:LYS:HA | 1:B:78:ILE:HD12 | 2.01 | 0.41 |
| 1:A:120:ASP:OD1 | 1:A:123:GLN:CB | 2.69 | 0.41 |
| 1:A:128:LEU:O | 1:A:132:ASP:OD2 | 2.38 | 0.41 |
| 1:B:189:ASN:ND2 | 2:B:407:SO4:O4 | 2.52 | 0.41 |
| 1:A:117:ASN:O | 1:A:165:ASP:HB2 | 2.20 | 0.41 |
| 1:A:240:ASP:O | 1:A:244:ARG:HG2 | 2.21 | 0.40 |

There are no symmetry-related clashes.

For Manuscript Review

#### 5.3 Torsion angles (i)

##### 5.3.1 Protein backbone (i)

In the following table, the Percentiles column shows the percent Ramachandran outliers of the chain as a percentile score with respect to all X-ray entries followed by that with respect to entries of similar resolution.

The Analysed column shows the number of residues for which the backbone conformation was analysed, and the total number of residues.

| Mol | Chain | Analysed | Favoured | Allowed | Outliers | Percentiles | |
|---|---|---|---|---|---|---|---|
| 1 | A | 327/338 (97%) | 311 (95%) | 16 (5%) | 0 | 100 | 100 |
| 1 | B | 327/338 (97%) | 312 (95%) | 15 (5%) | 0 | 100 | 100 |
| All | All | 654/676 (97%) | 623 (95%) | 31 (5%) | 0 | 100 | 100 |

There are no Ramachandran outliers to report.

##### 5.3.2 Protein sidechains (i)

In the following table, the Percentiles column shows the percent sidechain outliers of the chain as a percentile score with respect to all X-ray entries followed by that with respect to entries of similar resolution.

The Analysed column shows the number of residues for which the sidechain conformation was analysed, and the total number of residues.

| Mol | Chain | Analysed | Rotameric | Outliers | Percentiles | |
|---|---|---|---|---|---|---|
| 1 | A | 277/310 (89%) | 268 (97%) | 9 (3%) | 39 | 56 |
| 1 | B | 278/310 (90%) | 269 (97%) | 9 (3%) | 39 | 56 |
| All | All | 555/620 (90%) | 537 (97%) | 18 (3%) | 39 | 56 |

All (18) residues with a non-rotameric sidechain are listed below:

| Mol | Chain | Res | Type |
|---|---|---|---|
| 1 | A | 61 | ASN |
| 1 | A | 65 | VAL |
| 1 | A | 153 | MET |
| 1 | A | 225 | MET |
| 1 | A | 234 | HIS |
| 1 | A | 258 | ILE |
| 1 | A | 273 | LEU |
| 1 | A | 287 | SER |

*Continued on next page...*

For Manuscript Review

*Continued from previous page...*

| Mol | Chain | Res | Type |
|---|---|---|---|
| 1 | A | 306 | ARG |
| 1 | B | 21 | ARG |
| 1 | B | 65 | VAL |
| 1 | B | 153 | MET |
| 1 | B | 170 | LYS |
| 1 | B | 228 | ARG |
| 1 | B | 229 | LYS |
| 1 | B | 244 | ARG |
| 1 | B | 258 | ILE |
| 1 | B | 283 | ARG |

Sometimes sidechains can be flipped to improve hydrogen bonding and reduce clashes. All (8) such sidechains are listed below:

| Mol | Chain | Res | Type |
|---|---|---|---|
| 1 | A | 61 | ASN |
| 1 | A | 115 | HIS |
| 1 | A | 148 | HIS |
| 1 | A | 310 | GLN |
| 1 | B | 118 | ASN |
| 1 | B | 148 | HIS |
| 1 | B | 291 | HIS |
| 1 | B | 331 | GLN |

##### 5.3.3 RNA ⓘ

There are no RNA molecules in this entry.

#### 5.4 Non-standard residues in protein, DNA, RNA chains ⓘ

There are no non-standard protein/DNA/RNA residues in this entry.

#### 5.5 Carbohydrates ⓘ

There are no monosaccharides in this entry.

#### 5.6 Ligand geometry ⓘ

21 ligands are modelled in this entry.

For Manuscript Review

In the following table, the Counts columns list the number of bonds (or angles) for which Mogul statistics could be retrieved, the number of bonds (or angles) that are observed in the model and the number of bonds (or angles) that are defined in the Chemical Component Dictionary. The Link column lists molecule types, if any, to which the group is linked. The Z score for a bond length (or angle) is the number of standard deviations the observed value is removed from the expected value. A bond length (or angle) with $|Z| > 2$ is considered an outlier worth inspection. RMSZ is the root-mean-square of all Z scores of the bond lengths (or angles).

| Mol | Type | Chain | Res | Link | Bond lengths | | | Bond angles | | |
|---|---|---|---|---|---|---|---|---|---|---|
| | | | | | Counts | RMSZ | #\|Z\| > 2 | Counts | RMSZ | #\|Z\| > 2 |
| 2 | SO4 | A | 401 | - | 4,4,4 | 0.23 | 0 | 6,6,6 | 0.13 | 0 |
| 2 | SO4 | B | 402 | - | 4,4,4 | 0.38 | 0 | 6,6,6 | 0.07 | 0 |
| 2 | SO4 | A | 407 | - | 4,4,4 | 0.43 | 0 | 6,6,6 | 0.17 | 0 |
| 2 | SO4 | A | 405 | - | 4,4,4 | 0.36 | 0 | 6,6,6 | 0.05 | 0 |
| 4 | A1H6G | A | 409 | - | 38,42,42 | 0.52 | 0 | 48,63,63 | 0.95 | 3 (6%) |
| 2 | SO4 | B | 404 | - | 4,4,4 | 0.36 | 0 | 6,6,6 | 0.07 | 0 |
| 2 | SO4 | A | 402 | - | 4,4,4 | 0.33 | 0 | 6,6,6 | 0.08 | 0 |
| 3 | EDO | B | 408 | - | 3,3,3 | 0.43 | 0 | 2,2,2 | 0.65 | 0 |
| 2 | SO4 | B | 407 | - | 4,4,4 | 0.37 | 0 | 6,6,6 | 0.14 | 0 |
| 2 | SO4 | A | 411 | - | 4,4,4 | 0.42 | 0 | 6,6,6 | 0.04 | 0 |
| 4 | A1H6G | B | 409 | - | 38,42,42 | 0.55 | 0 | 48,63,63 | 1.08 | 2 (4%) |
| 2 | SO4 | A | 403 | - | 4,4,4 | 0.34 | 0 | 6,6,6 | 0.08 | 0 |
| 2 | SO4 | B | 410 | - | 4,4,4 | 0.34 | 0 | 6,6,6 | 0.15 | 0 |
| 2 | SO4 | B | 401 | - | 4,4,4 | 0.37 | 0 | 6,6,6 | 0.08 | 0 |
| 2 | SO4 | B | 403 | - | 4,4,4 | 0.34 | 0 | 6,6,6 | 0.08 | 0 |
| 3 | EDO | A | 408 | - | 3,3,3 | 0.17 | 0 | 2,2,2 | 0.26 | 0 |
| 2 | SO4 | B | 406 | - | 4,4,4 | 0.39 | 0 | 6,6,6 | 0.18 | 0 |
| 2 | SO4 | A | 406 | - | 4,4,4 | 0.37 | 0 | 6,6,6 | 0.12 | 0 |
| 2 | SO4 | B | 405 | - | 4,4,4 | 0.33 | 0 | 6,6,6 | 0.05 | 0 |
| 2 | SO4 | A | 410 | - | 4,4,4 | 0.36 | 0 | 6,6,6 | 0.13 | 0 |
| 2 | SO4 | A | 404 | - | 4,4,4 | 0.37 | 0 | 6,6,6 | 0.05 | 0 |

In the following table, the Chirals column lists the number of chiral outliers, the number of chiral centers analysed, the number of these observed in the model and the number defined in the Chemical Component Dictionary. Similar counts are reported in the Torsion and Rings columns. '-' means no outliers of that kind were identified.

| Mol | Type | Chain | Res | Link | Chirals | Torsions | Rings |
|---|---|---|---|---|---|---|---|
| 3 | EDO | B | 408 | - | - | 1/1/1/1 | - |
| 4 | A1H6G | A | 409 | - | - | 0/19/43/43 | 0/6/6/6 |
| 4 | A1H6G | B | 409 | - | - | 0/19/43/43 | 0/6/6/6 |
| 3 | EDO | A | 408 | - | - | 0/1/1/1 | - |

There are no bond length outliers.

For Manuscript Review

All (5) bond angle outliers are listed below:

| Mol | Chain | Res | Type | Atoms | Z | Observed(°) | Ideal(°) |
|---|---|---|---|---|---|---|---|
| 4 | B | 409 | A1H6G | C5-N6-C7 | 5.36 | 121.80 | 114.14 |
| 4 | A | 409 | A1H6G | C16-C21-N20 | 2.98 | 114.97 | 108.04 |
| 4 | B | 409 | A1H6G | C16-C21-N20 | 2.94 | 114.89 | 108.04 |
| 4 | A | 409 | A1H6G | C5-N6-C7 | 2.74 | 118.06 | 114.14 |
| 4 | A | 409 | A1H6G | C26-N12-S13 | 2.23 | 121.10 | 117.05 |

There are no chirality outliers.

All (1) torsion outliers are listed below:

| Mol | Chain | Res | Type | Atoms |
|---|---|---|---|---|
| 3 | B | 408 | EDO | O1-C1-C2-O2 |

There are no ring outliers.

3 monomers are involved in 4 short contacts:

| Mol | Chain | Res | Type | Clashes | Symm-Clashes |
|---|---|---|---|---|---|
| 4 | A | 409 | A1H6G | 1 | 0 |
| 3 | B | 408 | EDO | 2 | 0 |
| 2 | B | 407 | SO4 | 1 | 0 |

The following is a two-dimensional graphical depiction of Mogul quality analysis of bond lengths, bond angles, torsion angles, and ring geometry for all instances of the Ligand of Interest. In addition, ligands with molecular weight > 250 and outliers as shown on the validation Tables will also be included. For torsion angles, if less then 5% of the Mogul distribution of torsion angles is within 10 degrees of the torsion angle in question, then that torsion angle is considered an outlier. Any bond that is central to one or more torsion angles identified as an outlier by Mogul will be highlighted in the graph. For rings, the root-mean-square deviation (RMSD) between the ring in question and similar rings identified by Mogul is calculated over all ring torsion angles. If the average RMSD is greater than 60 degrees and the minimal RMSD between the ring in question and any Mogul-identified rings is also greater than 60 degrees, then that ring is considered an outlier. The outliers are highlighted in purple. The color gray indicates Mogul did not find sufficient equivalents in the CSD to analyse the geometry.

For Manuscript Review

Ligand A1H6G A 409

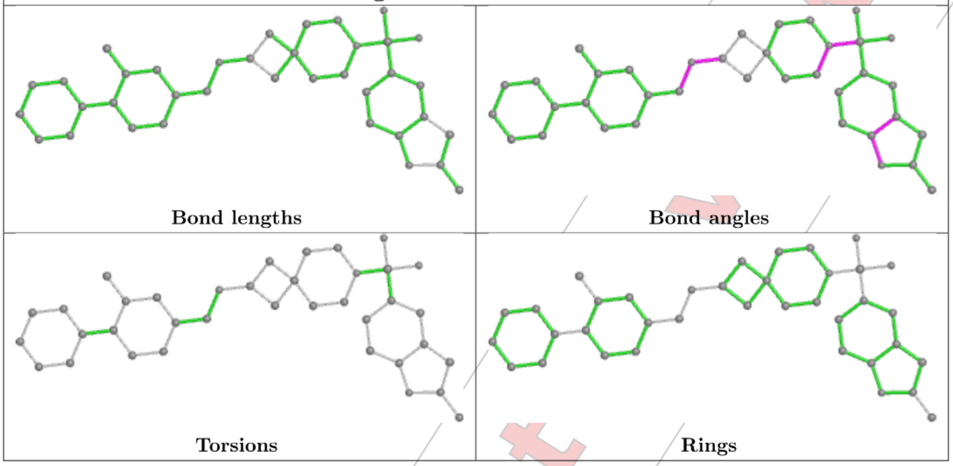

Ligand A1H6G B 409

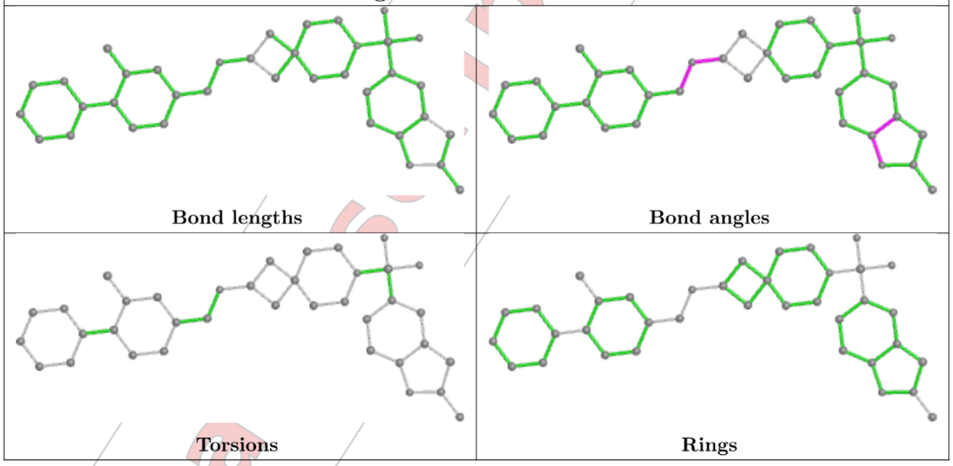

#### 5.7 Other polymers (i)

There are no such residues in this entry.

#### 5.8 Polymer linkage issues (i)

There are no chain breaks in this entry.

For Manuscript Review

### 6 Fit of model and data (i)

#### 6.1 Protein, DNA and RNA chains (i)

In the following table, the column labelled '#RSRZ> 2' contains the number (and percentage) of RSRZ outliers, followed by percent RSRZ outliers for the chain as percentile scores relative to all X-ray entries and entries of similar resolution. The OWAB column contains the minimum, median, $95^{th}$ percentile and maximum values of the occupancy-weighted average B-factor per residue. The column labelled 'Q< 0.9' lists the number of (and percentage) of residues with an average occupancy less than 0.9.

| Mol | Chain | Analysed | <RSRZ> | #RSRZ>2 | | | OWAB(Å$^2$) | Q<0.9 |
|---|---|---|---|---|---|---|---|---|
| 1 | A | 329/338 (97%) | -0.08 | 3 (0%) | 84 | 83 | 51, 73, 108, 130 | 0 |
| 1 | B | 329/338 (97%) | -0.02 | 3 (0%) | 84 | 83 | 50, 73, 110, 142 | 0 |
| All | All | 658/676 (97%) | -0.05 | 6 (0%) | 84 | 83 | 50, 73, 110, 142 | 0 |

All (6) RSRZ outliers are listed below:

| Mol | Chain | Res | Type | RSRZ |
|---|---|---|---|---|
| 1 | A | 105 | VAL | 3.5 |
| 1 | B | 273 | LEU | 3.1 |
| 1 | A | 265 | LEU | 2.5 |
| 1 | A | 2 | SER | 2.3 |
| 1 | B | 270 | ASN | 2.3 |
| 1 | B | 131 | TYR | 2.1 |

#### 6.2 Non-standard residues in protein, DNA, RNA chains (i)

There are no non-standard protein/DNA/RNA residues in this entry.

#### 6.3 Carbohydrates (i)

There are no monosaccharides in this entry.

#### 6.4 Ligands (i)

In the following table, the Atoms column lists the number of modelled atoms in the group and the number defined in the chemical component dictionary. The B-factors column lists the minimum, median, $95^{th}$ percentile and maximum values of B factors of atoms in the group. The column labelled 'Q< 0.9' lists the number of atoms with occupancy less than 0.9.

For Manuscript Review

| Mol | Type | Chain | Res | Atoms | RSCC | RSR | B-factors(Å$^2$) | Q<0.9 |
|---|---|---|---|---|---|---|---|---|
| 2 | SO4 | A | 405 | 5/5 | 0.71 | 0.40 | 142,155,162,167 | 0 |
| 3 | EDO | A | 408 | 4/4 | 0.77 | 0.18 | 91,95,99,100 | 0 |
| 2 | SO4 | B | 402 | 5/5 | 0.80 | 0.16 | 144,147,150,157 | 0 |
| 2 | SO4 | B | 406 | 5/5 | 0.81 | 0.14 | 84,107,112,113 | 0 |
| 2 | SO4 | B | 410 | 5/5 | 0.84 | 0.19 | 110,117,125,135 | 0 |
| 2 | SO4 | A | 410 | 5/5 | 0.84 | 0.23 | 97,100,113,128 | 0 |
| 2 | SO4 | B | 405 | 5/5 | 0.85 | 0.30 | 118,123,138,145 | 0 |
| 2 | SO4 | A | 411 | 5/5 | 0.86 | 0.74 | 30,30,30,30 | 0 |
| 2 | SO4 | A | 404 | 5/5 | 0.86 | 0.33 | 147,149,159,160 | 0 |
| 2 | SO4 | A | 406 | 5/5 | 0.87 | 0.14 | 92,105,111,131 | 0 |
| 2 | SO4 | B | 401 | 5/5 | 0.90 | 0.14 | 96,100,120,135 | 0 |
| 2 | SO4 | A | 401 | 5/5 | 0.91 | 0.13 | 78,88,117,125 | 0 |
| 2 | SO4 | B | 403 | 5/5 | 0.91 | 0.12 | 112,120,131,131 | 0 |
| 2 | SO4 | A | 403 | 5/5 | 0.91 | 0.17 | 96,99,102,103 | 0 |
| 3 | EDO | B | 408 | 4/4 | 0.92 | 0.20 | 57,61,61,63 | 0 |
| 4 | A1H6G | B | 409 | 37/37 | 0.92 | 0.24 | 62,74,119,136 | 0 |
| 2 | SO4 | B | 407 | 5/5 | 0.93 | 0.15 | 79,86,95,97 | 0 |
| 2 | SO4 | A | 402 | 5/5 | 0.94 | 0.11 | 103,113,117,119 | 0 |
| 2 | SO4 | A | 407 | 5/5 | 0.95 | 0.14 | 72,73,84,92 | 0 |
| 4 | A1H6G | A | 409 | 37/37 | 0.96 | 0.21 | 56,77,107,140 | 0 |
| 2 | SO4 | B | 404 | 5/5 | 0.96 | 0.09 | 91,94,101,102 | 0 |

The following is a graphical depiction of the model fit to experimental electron density of all instances of the Ligand of Interest. In addition, ligands with molecular weight > 250 and outliers as shown on the geometry validation Tables will also be included. Each fit is shown from different orientation to approximate a three-dimensional view.

For Manuscript Review

**Electron density around A1H6G B 409:**

2mF$_o$-DF$_c$ (at 0.7 rmsd) in gray
mF$_o$-DF$_c$ (at 3 rmsd) in purple (negative)
and green (positive)

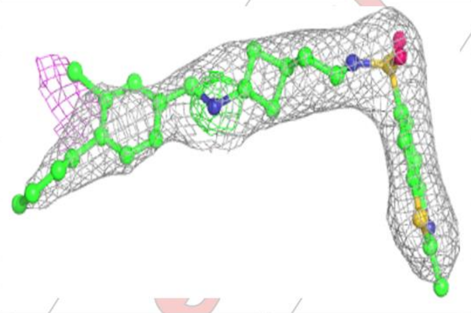

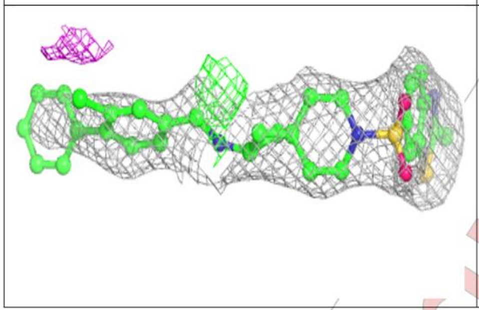

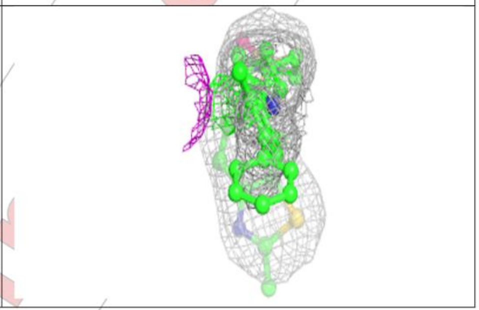

**Electron density around A1H6G A 409:**

2mF$_o$-DF$_c$ (at 0.7 rmsd) in gray
mF$_o$-DF$_c$ (at 3 rmsd) in purple (negative)
and green (positive)

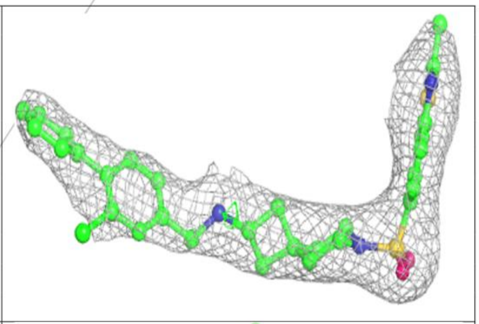

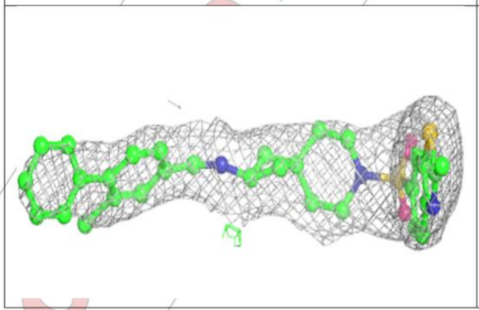

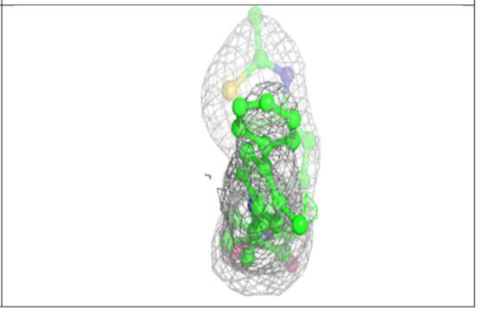

For Manuscript Review

#### 6.5 Other polymers ⓘ

There are no such residues in this entry.

For Manuscript Review

WORLDWIDE PDB PROTEIN DATA BANK

### Full wwPDB X-ray Structure Validation Report (i)

Mar 20, 2024 – 02:50 pm GMT

| | | |
|---|---|---|
| PDB ID | : | 9EPZ |
| Title | : | Crystal structure of human Casein Kinase II subunit alpha (CK2a1) in complex with allosteric ligand FGC3337 |
| Deposited on | : | 2024-03-20 |
| Resolution | : | 2.65 Å(reported) |

**This wwPDB validation report is for manuscript review**

For Manuscript Review

This is a Full wwPDB X-ray Structure Validation Report.

This report is produced by the wwPDB biocuration pipeline after annotation of the structure.

We welcome your comments at **
A user guide is available at
https://www.wwpdb.org/validation/2017/XrayValidationReportHelp
with specific help available everywhere you see the (i) symbol.

The types of validation reports are described at
http://www.wwpdb.org/validation/2017/FAQs#types.

The following versions of software and data (see references (i)) were used in the production of this report:

| | | |
|---|---|---|
| MolProbity | : | 4.02b-467 |
| Mogul | : | 1.8.4, CSD as541be (2020) |
| Xtriage (Phenix) | : | 1.13 |
| EDS | : | 2.36 |
| buster-report | : | 1.1.7 (2018) |
| Percentile statistics | : | 20191225.v01 (using entries in the PDB archive December 25th 2019) |
| Refmac | : | 5.8.0158 |
| CCP4 | : | 7.0.044 (Gargrove) |
| Ideal geometry (proteins) | : | Engh & Huber (2001) |

### 1 Overall quality at a glance (i)

The following experimental techniques were used to determine the structure:
*X-RAY DIFFRACTION*

The reported resolution of this entry is 2.65 Å.

Percentile scores (ranging between 0-100) for global validation metrics of the entry are shown in the following graphic. The table shows the number of entries on which the scores are based.

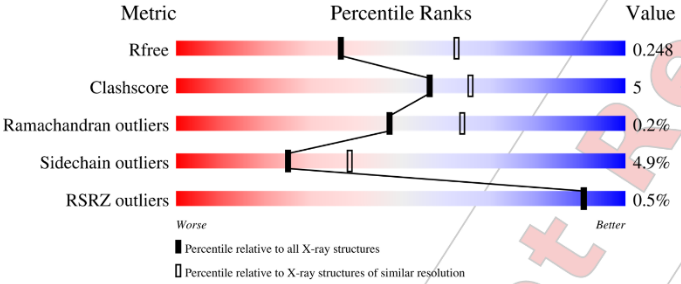

| Metric | Whole archive (#Entries) | Similar resolution (#Entries, resolution range(Å)) |
|---|---|---|
| $R_{free}$ | 130704 | 1332 (2.68-2.64) |
| Clashscore | 141614 | 1374 (2.68-2.64) |
| Ramachandran outliers | 138981 | 1349 (2.68-2.64) |
| Sidechain outliers | 138945 | 1349 (2.68-2.64) |
| RSRZ outliers | 127900 | 1318 (2.68-2.64) |

The table below summarises the geometric issues observed across the polymeric chains and their fit to the electron density. The red, orange, yellow and green segments of the lower bar indicate the fraction of residues that contain outliers for >=3, 2, 1 and 0 types of geometric quality criteria respectively. A grey segment represents the fraction of residues that are not modelled. The numeric value for each fraction is indicated below the corresponding segment, with a dot representing fractions <=5% The upper red bar (where present) indicates the fraction of residues that have poor fit to the electron density. The numeric value is given above the bar.

| Mol | Chain | Length | Quality of chain |
|---|---|---|---|
| 1 | A | 338 | 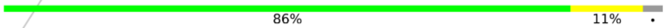 |
| 1 | B | 338 | 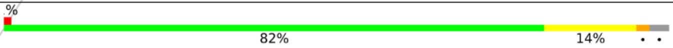 |

Ideal geometry (DNA, RNA) : Parkinson et al. (1996)
Validation Pipeline (wwPDB-VP) : 2.36

For Manuscript Review

#### 2 Entry composition (i)

There are 5 unique types of molecules in this entry. The entry contains 5677 atoms, of which 0 are hydrogens and 0 are deuteriums.

In the tables below, the ZeroOcc column contains the number of atoms modelled with zero occupancy, the AltConf column contains the number of residues with at least one atom in alternate conformation and the Trace column contains the number of residues modelled with at most 2 atoms.

- Molecule 1 is a protein called Casein kinase II subunit alpha.

| Mol | Chain | Residues | Atoms | | | | | ZeroOcc | AltConf | Trace |
|---|---|---|---|---|---|---|---|---|---|---|
| 1 | A | 329 | Total 2712 | C 1748 | N 470 | O 483 | S 11 | 0 | 0 | 0 |
| 1 | B | 329 | Total 2728 | C 1749 | N 476 | O 492 | S 11 | 0 | 0 | 0 |

There are 2 discrepancies between the modelled and reference sequences:

| Chain | Residue | Modelled | Actual | Comment | Reference |
|---|---|---|---|---|---|
| A | 0 | SER | - | expression tag | UNP P68400 |
| B | 0 | SER | - | expression tag | UNP P68400 |

- Molecule 2 is SULFATE ION (three-letter code: SO4) (formula: $O_4S$).

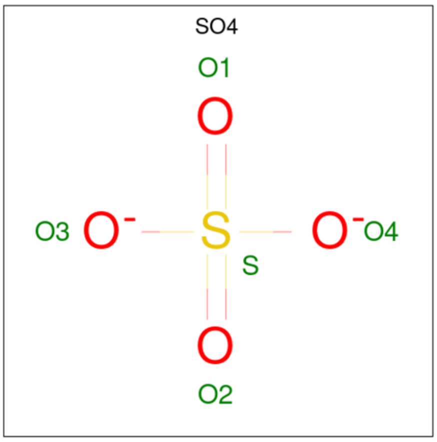

| Mol | Chain | Residues | Atoms | | | ZeroOcc | AltConf |
|---|---|---|---|---|---|---|---|
| 2 | A | 1 | Total 5 | O 4 | S 1 | 0 | 0 |

Continued on next page...

*Continued from previous page...*

| Mol | Chain | Residues | Atoms | ZeroOcc | AltConf |
|---|---|---|---|---|---|
| 2 | A | 1 | Total O S<br>5 4 1 | 0 | 0 |
| 2 | A | 1 | Total O S<br>5 4 1 | 0 | 0 |
| 2 | A | 1 | Total O S<br>5 4 1 | 0 | 0 |
| 2 | A | 1 | Total O S<br>5 4 1 | 0 | 0 |
| 2 | A | 1 | Total O S<br>5 4 1 | 0 | 0 |
| 2 | A | 1 | Total O S<br>5 4 1 | 0 | 0 |
| 2 | B | 1 | Total O S<br>5 4 1 | 0 | 0 |
| 2 | B | 1 | Total O S<br>5 4 1 | 0 | 0 |
| 2 | B | 1 | Total O S<br>5 4 1 | 0 | 0 |
| 2 | B | 1 | Total O S<br>5 4 1 | 0 | 0 |
| 2 | B | 1 | Total O S<br>5 4 1 | 0 | 0 |
| 2 | B | 1 | Total O S<br>5 4 1 | 0 | 0 |
| 2 | B | 1 | Total O S<br>5 4 1 | 0 | 0 |
| 2 | B | 1 | Total O S<br>5 4 1 | 0 | 0 |
| 2 | B | 1 | Total O S<br>5 4 1 | 0 | 0 |
| 2 | B | 1 | Total O S<br>5 4 1 | 0 | 0 |

- Molecule 3 is 1,2-ETHANEDIOL (three-letter code: EDO) (formula: $C_2H_6O_2$).

For Manuscript Review

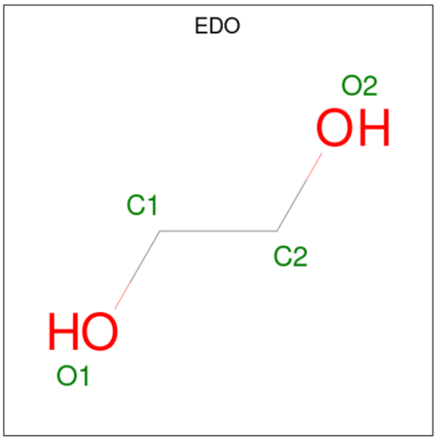

| Mol | Chain | Residues | Atoms | | | ZeroOcc | AltConf |
|---|---|---|---|---|---|---|---|
| | | | Total | C | O | | |
| 3 | A | 1 | 4 | 2 | 2 | 0 | 0 |

- Molecule 4 is {N}-[(3-chloranyl-4-phenyl-phenyl)methyl]-2-[1-[(2-methyl-1,3-benzothiazol-6-yl)sulfonyl]piperidin-4-yl]ethanamine (three-letter code: A1H6I) (formula: $C_{28}H_{30}ClN_3O_2S_2$) (labeled as "Ligand of Interest" by depositor).

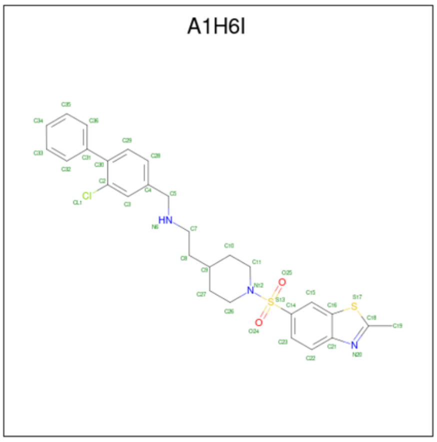

| Mol | Chain | Residues | Atoms | | | | | | ZeroOcc | AltConf |
|---|---|---|---|---|---|---|---|---|---|---|
| | | | Total | C | Cl | N | O | S | | |
| 4 | A | 1 | 36 | 28 | 1 | 3 | 2 | 2 | 0 | 0 |
| 4 | B | 1 | 36 | 28 | 1 | 3 | 2 | 2 | 0 | 0 |

For Manuscript Review

- Molecule 5 is water.

| Mol | Chain | Residues | Atoms | ZeroOcc | AltConf |
|---|---|---|---|---|---|
| 5 | A | 38 | Total O<br>38 38 | 0 | 0 |
| 5 | B | 38 | Total O<br>38 38 | 0 | 0 |

For Manuscript Review

WORLDWIDE PDB PROTEIN DATA BANK

#### 3 Residue-property plots (i)

These plots are drawn for all protein, RNA, DNA and oligosaccharide chains in the entry. The first graphic for a chain summarises the proportions of the various outlier classes displayed in the second graphic. The second graphic shows the sequence view annotated by issues in geometry and electron density. Residues are color-coded according to the number of geometric quality criteria for which they contain at least one outlier: green = 0, yellow = 1, orange = 2 and red = 3 or more. A red dot above a residue indicates a poor fit to the electron density (RSRZ > 2). Stretches of 2 or more consecutive residues without any outlier are shown as a green connector. Residues present in the sample, but not in the model, are shown in grey.

- Molecule 1: Casein kinase II subunit alpha

Chain A:

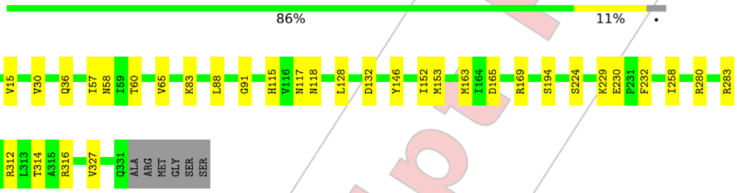

- Molecule 1: Casein kinase II subunit alpha

Chain B:

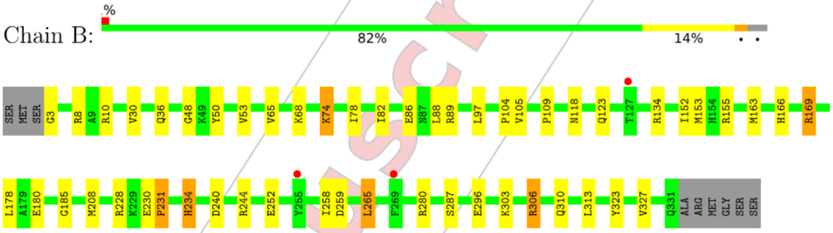

For Manuscript Review

#### 4 Data and refinement statistics ⓘ

| Property | Value | Source |
|---|---|---|
| Space group | P 43 21 2 | Depositor |
| Cell constants<br>a, b, c, $\alpha$, $\beta$, $\gamma$ | 128.16Å 128.16Å 124.01Å<br>90.00° 90.00° 90.00° | Depositor |
| Resolution (Å) | 45.31 – 2.65<br>45.31 – 2.65 | Depositor<br>EDS |
| % Data completeness<br>(in resolution range) | 100.0 (45.31-2.65)<br>100.0 (45.31-2.65) | Depositor<br>EDS |
| $R_{merge}$ | (Not available) | Depositor |
| $R_{sym}$ | (Not available) | Depositor |
| $< I/\sigma(I) >$ [1] | 2.22 (at 2.65Å) | Xtriage |
| Refinement program | REFMAC 5.8.0419 | Depositor |
| R, $R_{free}$ | 0.213 , 0.245<br>0.219 , 0.248 | Depositor<br>DCC |
| $R_{free}$ test set | 1573 reflections (5.14%) | wwPDB-VP |
| Wilson B-factor (Å$^2$) | 46.5 | Xtriage |
| Anisotropy | 0.498 | Xtriage |
| Bulk solvent $k_{sol}$(e/Å$^3$), $B_{sol}$(Å$^2$) | 0.34 , 44.2 | EDS |
| L-test for twinning[2] | $< \|L\| >$ = 0.50, $< L^2 >$ = 0.33 | Xtriage |
| Estimated twinning fraction | 0.000 for -h,-l,-k<br>0.000 for l,-k,h | Xtriage |
| $F_o$,$F_c$ correlation | 0.94 | EDS |
| Total number of atoms | 5677 | wwPDB-VP |
| Average B, all atoms (Å$^2$) | 52.0 | wwPDB-VP |

Xtriage's analysis on translational NCS is as follows: *The analyses of the Patterson function reveals a significant off-origin peak that is 30.36 % of the origin peak, indicating pseudo-translational symmetry. The chance of finding a peak of this or larger height randomly in a structure without pseudo-translational symmetry is equal to 1.3254e-03. The detected translational NCS is most likely also responsible for the elevated intensity ratio.*

[1] Intensities estimated from amplitudes.
[2] Theoretical values of $< |L| >$, $< L^2 >$ for acentric reflections are 0.5, 0.333 respectively for untwinned datasets, and 0.375, 0.2 for perfectly twinned datasets.

For Manuscript Review

### 5 Model quality (i)

#### 5.1 Standard geometry (i)

Bond lengths and bond angles in the following residue types are not validated in this section: EDO, A1H6I, SO4

The Z score for a bond length (or angle) is the number of standard deviations the observed value is removed from the expected value. A bond length (or angle) with $|Z| > 5$ is considered an outlier worth inspection. RMSZ is the root-mean-square of all Z scores of the bond lengths (or angles).

| Mol | Chain | Bond lengths | | Bond angles | |
|---|---|---|---|---|---|
| | | RMSZ | #\|Z\| >5 | RMSZ | #\|Z\| >5 |
| 1 | A | 0.41 | 0/2787 | 0.80 | 0/3780 |
| 1 | B | 0.39 | 0/2801 | 0.79 | 0/3796 |
| All | All | 0.40 | 0/5588 | 0.79 | 0/7576 |

There are no bond length outliers.

There are no bond angle outliers.

There are no chirality outliers.

There are no planarity outliers.

#### 5.2 Too-close contacts (i)

In the following table, the Non-H and H(model) columns list the number of non-hydrogen atoms and hydrogen atoms in the chain respectively. The H(added) column lists the number of hydrogen atoms added and optimized by MolProbity. The Clashes column lists the number of clashes within the asymmetric unit, whereas Symm-Clashes lists symmetry-related clashes.

| Mol | Chain | Non-H | H(model) | H(added) | Clashes | Symm-Clashes |
|---|---|---|---|---|---|---|
| 1 | A | 2712 | 0 | 2612 | 27 | 0 |
| 1 | B | 2728 | 0 | 2632 | 35 | 0 |
| 2 | A | 35 | 0 | 0 | 1 | 0 |
| 2 | B | 50 | 0 | 0 | 2 | 0 |
| 3 | A | 4 | 0 | 6 | 0 | 0 |
| 4 | A | 36 | 0 | 0 | 0 | 0 |
| 4 | B | 36 | 0 | 0 | 0 | 0 |
| 5 | A | 38 | 0 | 0 | 2 | 0 |
| 5 | B | 38 | 0 | 0 | 1 | 0 |
| All | All | 5677 | 0 | 5250 | 54 | 0 |

The all-atom clashscore is defined as the number of clashes found per 1000 atoms (including

For Manuscript Review

hydrogen atoms). The all-atom clashscore for this structure is 5.

All (54) close contacts within the same asymmetric unit are listed below, sorted by their clash magnitude.

| Atom-1 | Atom-2 | Interatomic distance (Å) | Clash overlap (Å) |
|---|---|---|---|
| 1:A:280:ARG:CD | 1:B:306:ARG:HH21 | 1.79 | 0.93 |
| 1:A:280:ARG:HD3 | 1:B:306:ARG:HH21 | 1.42 | 0.84 |
| 1:A:306:ARG:NH1 | 1:B:280:ARG:HD3 | 1.93 | 0.83 |
| 1:A:58:ASN:OD1 | 1:A:60:THR:HB | 1.84 | 0.77 |
| 1:A:224:SER:HB2 | 1:A:230:GLU:HG3 | 1.66 | 0.77 |
| 1:A:280:ARG:CD | 1:B:306:ARG:NH2 | 2.49 | 0.74 |
| 1:A:306:ARG:HH11 | 1:B:280:ARG:HD3 | 1.57 | 0.70 |
| 1:A:280:ARG:HD3 | 1:B:306:ARG:NH2 | 2.07 | 0.69 |
| 1:B:118:ASN:OD1 | 1:B:163:MET:HE3 | 2.01 | 0.61 |
| 1:A:118:ASN:HD21 | 1:A:163:MET:HG2 | 1.69 | 0.57 |
| 1:B:36:GLN:HG3 | 1:B:104:PRO:HD3 | 1.86 | 0.56 |
| 1:B:185:GLY:HA2 | 1:B:208:MET:CE | 2.36 | 0.55 |
| 1:A:280:ARG:NE | 1:B:306:ARG:HH21 | 2.06 | 0.54 |
| 1:A:10:ARG:HD2 | 1:A:310:GLN:O | 2.08 | 0.54 |
| 1:B:118:ASN:OD1 | 1:B:163:MET:CE | 2.56 | 0.54 |
| 1:A:128:LEU:O | 1:A:132:ASP:HB2 | 2.09 | 0.53 |
| 1:A:229:LYS:HD3 | 1:A:232:PHE:HA | 1.89 | 0.53 |
| 1:B:155:ARG:NH2 | 1:B:180:GLU:HB2 | 2.23 | 0.52 |
| 1:B:10:ARG:HD2 | 1:B:310:GLN:O | 2.09 | 0.52 |
| 1:B:74:LYS:HE2 | 2:B:402:SO4:O1 | 2.09 | 0.52 |
| 1:B:185:GLY:HA2 | 1:B:208:MET:HE1 | 1.91 | 0.51 |
| 1:A:115:HIS:NE2 | 1:A:117:ASN:OD1 | 2.41 | 0.51 |
| 1:B:78:ILE:HD13 | 1:B:109:PRO:HG2 | 1.95 | 0.48 |
| 1:A:91:GLY:HA3 | 1:A:146:TYR:CE2 | 2.49 | 0.48 |
| 1:A:283:ARG:NH1 | 2:A:401:SO4:O1 | 2.46 | 0.48 |
| 1:B:8:ARG:HH21 | 1:B:208:MET:HE2 | 1.79 | 0.47 |
| 1:B:89:ARG:NH1 | 1:B:97:LEU:O | 2.47 | 0.47 |
| 1:A:280:ARG:O | 1:A:283:ARG:HG2 | 2.15 | 0.47 |
| 1:B:230:GLU:CB | 1:B:231:PRO:HD3 | 2.45 | 0.46 |
| 1:A:314:THR:HG22 | 1:A:316:ARG:H | 1.79 | 0.46 |
| 1:B:82:ILE:O | 1:B:86:GLU:HG3 | 2.15 | 0.46 |
| 1:A:128:LEU:O | 1:A:132:ASP:OD2 | 2.33 | 0.45 |
| 1:B:155:ARG:HD3 | 1:B:178:LEU:O | 2.16 | 0.45 |
| 1:B:53:VAL:HG22 | 1:B:68:LYS:HG2 | 1.98 | 0.45 |
| 1:B:303:LYS:HB3 | 1:B:313:LEU:HG | 1.98 | 0.45 |
| 1:B:3:GLY:N | 5:B:501:HOH:O | 2.50 | 0.45 |
| 1:A:10:ARG:NH2 | 1:A:312:ARG:O | 2.50 | 0.45 |
| 1:B:48:GLY:O | 1:B:50:TYR:N | 2.49 | 0.44 |

*Continued on next page...*

For Manuscript Review

*Continued from previous page...*

| Atom-1 | Atom-2 | Interatomic distance (Å) | Clash overlap (Å) |
|---|---|---|---|
| 1:A:88:LEU:HD21 | 1:A:152:ILE:HD12 | 1.99 | 0.44 |
| 1:A:280:ARG:HD2 | 1:B:306:ARG:NH2 | 2.30 | 0.43 |
| 1:B:8:ARG:NH2 | 1:B:208:MET:CE | 2.81 | 0.43 |
| 1:A:132:ASP:OD1 | 1:A:169:ARG:NH2 | 2.51 | 0.43 |
| 1:B:134:ARG:HG2 | 1:B:323:TYR:CZ | 2.54 | 0.43 |
| 1:A:314:THR:HG22 | 5:A:506:HOH:O | 2.19 | 0.43 |
| 1:B:166:HIS:CD2 | 1:B:169:ARG:HH11 | 2.38 | 0.41 |
| 1:B:240:ASP:O | 1:B:244:ARG:HG2 | 2.20 | 0.41 |
| 1:B:234:HIS:O | 1:B:244:ARG:NH1 | 2.53 | 0.41 |
| 1:B:265:LEU:HD23 | 1:B:265:LEU:HA | 1.91 | 0.41 |
| 1:A:314:THR:CG2 | 5:A:506:HOH:O | 2.69 | 0.41 |
| 1:A:117:ASN:O | 1:A:165:ASP:HB2 | 2.21 | 0.40 |
| 1:B:48:GLY:O | 2:B:408:SO4:O4 | 2.39 | 0.40 |
| 1:B:88:LEU:HD21 | 1:B:152:ILE:HD12 | 2.04 | 0.40 |
| 1:B:185:GLY:HA2 | 1:B:208:MET:HE3 | 2.03 | 0.40 |
| 1:A:83:LYS:HA | 1:A:83:LYS:HD2 | 1.93 | 0.40 |

There are no symmetry-related clashes.

#### 5.3 Torsion angles ⓘ

##### 5.3.1 Protein backbone ⓘ

In the following table, the Percentiles column shows the percent Ramachandran outliers of the chain as a percentile score with respect to all X-ray entries followed by that with respect to entries of similar resolution.

The Analysed column shows the number of residues for which the backbone conformation was analysed, and the total number of residues.

| Mol | Chain | Analysed | Favoured | Allowed | Outliers | Percentiles | |
|---|---|---|---|---|---|---|---|
| 1 | A | 327/338 (97%) | 313 (96%) | 14 (4%) | 0 | 100 | 100 |
| 1 | B | 327/338 (97%) | 312 (95%) | 14 (4%) | 1 (0%) | 41 | 56 |
| All | All | 654/676 (97%) | 625 (96%) | 28 (4%) | 1 (0%) | 47 | 64 |

All (1) Ramachandran outliers are listed below:

| Mol | Chain | Res | Type |
|---|---|---|---|
| 1 | B | 231 | PRO |

For Manuscript Review

##### 5.3.2 Protein sidechains (i)

In the following table, the Percentiles column shows the percent sidechain outliers of the chain as a percentile score with respect to all X-ray entries followed by that with respect to entries of similar resolution.

The Analysed column shows the number of residues for which the sidechain conformation was analysed, and the total number of residues.

| Mol | Chain | Analysed | Rotameric | Outliers | Percentiles | |
|---|---|---|---|---|---|---|
| 1 | A | 286/310 (92%) | 275 (96%) | 11 (4%) | 33 | 49 |
| 1 | B | 291/310 (94%) | 274 (94%) | 17 (6%) | 20 | 31 |
| All | All | 577/620 (93%) | 549 (95%) | 28 (5%) | 25 | 38 |

All (28) residues with a non-rotameric sidechain are listed below:

| Mol | Chain | Res | Type |
|---|---|---|---|
| 1 | A | 15 | VAL |
| 1 | A | 30 | VAL |
| 1 | A | 36 | GLN |
| 1 | A | 57 | ILE |
| 1 | A | 65 | VAL |
| 1 | A | 153 | MET |
| 1 | A | 194 | SER |
| 1 | A | 258 | ILE |
| 1 | A | 296 | GLU |
| 1 | A | 306 | ARG |
| 1 | A | 327 | VAL |
| 1 | B | 30 | VAL |
| 1 | B | 65 | VAL |
| 1 | B | 74 | LYS |
| 1 | B | 105 | VAL |
| 1 | B | 123 | GLN |
| 1 | B | 153 | MET |
| 1 | B | 169 | ARG |
| 1 | B | 228 | ARG |
| 1 | B | 234 | HIS |
| 1 | B | 252 | GLU |
| 1 | B | 258 | ILE |
| 1 | B | 259 | ASP |
| 1 | B | 265 | LEU |
| 1 | B | 287 | SER |
| 1 | B | 296 | GLU |
| 1 | B | 306 | ARG |

*Continued on next page...*

For Manuscript Review

*Continued from previous page...*

| Mol | Chain | Res | Type |
|---|---|---|---|
| 1 | B | 327 | VAL |

Sometimes sidechains can be flipped to improve hydrogen bonding and reduce clashes. All (2) such sidechains are listed below:

| Mol | Chain | Res | Type |
|---|---|---|---|
| 1 | A | 62 | ASN |
| 1 | B | 186 | GLN |

##### 5.3.3 RNA (i)

There are no RNA molecules in this entry.

#### 5.4 Non-standard residues in protein, DNA, RNA chains (i)

There are no non-standard protein/DNA/RNA residues in this entry.

#### 5.5 Carbohydrates (i)

There are no monosaccharides in this entry.

#### 5.6 Ligand geometry (i)

20 ligands are modelled in this entry.

In the following table, the Counts columns list the number of bonds (or angles) for which Mogul statistics could be retrieved, the number of bonds (or angles) that are observed in the model and the number of bonds (or angles) that are defined in the Chemical Component Dictionary. The Link column lists molecule types, if any, to which the group is linked. The Z score for a bond length (or angle) is the number of standard deviations the observed value is removed from the expected value. A bond length (or angle) with $|Z| > 2$ is considered an outlier worth inspection. RMSZ is the root-mean-square of all Z scores of the bond lengths (or angles).

| Mol | Type | Chain | Res | Link | Bond lengths | | | Bond angles | | |
|---|---|---|---|---|---|---|---|---|---|---|
| | | | | | Counts | RMSZ | $\#\|Z\| > 2$ | Counts | RMSZ | $\#\|Z\| > 2$ |
| 4 | A1H6I | B | 411 | - | 38,40,40 | 0.52 | 0 | 52,57,57 | 1.00 | 4 (7%) |
| 2 | SO4 | B | 401 | - | 4,4,4 | 0.27 | 0 | 6,6,6 | 0.17 | 0 |
| 2 | SO4 | A | 404 | - | 4,4,4 | 0.32 | 0 | 6,6,6 | 0.09 | 0 |
| 2 | SO4 | A | 408 | - | 4,4,4 | 0.30 | 0 | 6,6,6 | 0.09 | 0 |
| 2 | SO4 | B | 409 | - | 4,4,4 | 0.28 | 0 | 6,6,6 | 0.29 | 0 |

For Manuscript Review

| Mol | Type | Chain | Res | Link | Bond lengths | | | Bond angles | | |
|---|---|---|---|---|---|---|---|---|---|---|
| | | | | | Counts | RMSZ | #\|Z\| > 2 | Counts | RMSZ | #\|Z\| > 2 |
| 2 | SO4 | A | 402 | - | 4,4,4 | 0.29 | 0 | 6,6,6 | 0.15 | 0 |
| 2 | SO4 | A | 401 | - | 4,4,4 | 0.27 | 0 | 6,6,6 | 0.17 | 0 |
| 2 | SO4 | B | 402 | - | 4,4,4 | 0.31 | 0 | 6,6,6 | 0.15 | 0 |
| 2 | SO4 | A | 406 | - | 4,4,4 | 0.34 | 0 | 6,6,6 | 0.16 | 0 |
| 2 | SO4 | A | 403 | - | 4,4,4 | 0.36 | 0 | 6,6,6 | 0.16 | 0 |
| 2 | SO4 | A | 405 | - | 4,4,4 | 0.37 | 0 | 6,6,6 | 0.16 | 0 |
| 2 | SO4 | B | 408 | - | 4,4,4 | 0.33 | 0 | 6,6,6 | 0.13 | 0 |
| 2 | SO4 | B | 407 | - | 4,4,4 | 0.36 | 0 | 6,6,6 | 0.10 | 0 |
| 2 | SO4 | B | 406 | - | 4,4,4 | 0.29 | 0 | 6,6,6 | 0.07 | 0 |
| 2 | SO4 | B | 403 | - | 4,4,4 | 0.30 | 0 | 6,6,6 | 0.10 | 0 |
| 3 | EDO | A | 407 | - | 3,3,3 | 0.14 | 0 | 2,2,2 | 0.24 | 0 |
| 2 | SO4 | B | 405 | - | 4,4,4 | 0.34 | 0 | 6,6,6 | 0.16 | 0 |
| 4 | A1H6I | A | 409 | - | 38,40,40 | 0.54 | 0 | 52,57,57 | 0.85 | 2 (3%) |
| 2 | SO4 | B | 404 | - | 4,4,4 | 0.28 | 0 | 6,6,6 | 0.19 | 0 |
| 2 | SO4 | B | 410 | - | 4,4,4 | 0.38 | 0 | 6,6,6 | 0.24 | 0 |

In the following table, the Chirals column lists the number of chiral outliers, the number of chiral centers analysed, the number of these observed in the model and the number defined in the Chemical Component Dictionary. Similar counts are reported in the Torsion and Rings columns. '-' means no outliers of that kind were identified.

| Mol | Type | Chain | Res | Link | Chirals | Torsions | Rings |
|---|---|---|---|---|---|---|---|
| 4 | A1H6I | A | 409 | - | - | 5/23/33/33 | 0/5/5/5 |
| 3 | EDO | A | 407 | - | - | 1/1/1/1 | - |
| 4 | A1H6I | B | 411 | - | - | 4/23/33/33 | 0/5/5/5 |

There are no bond length outliers.

All (6) bond angle outliers are listed below:

| Mol | Chain | Res | Type | Atoms | Z | Observed(°) | Ideal(°) |
|---|---|---|---|---|---|---|---|
| 4 | B | 411 | A1H6I | C7-C8-C9 | -3.21 | 106.16 | 114.32 |
| 4 | A | 409 | A1H6I | C16-C21-N20 | 3.03 | 115.09 | 108.04 |
| 4 | B | 411 | A1H6I | C16-C21-N20 | 2.96 | 114.94 | 108.04 |
| 4 | B | 411 | A1H6I | C26-C27-C9 | -2.48 | 105.72 | 111.99 |
| 4 | A | 409 | A1H6I | C7-C8-C9 | -2.22 | 108.69 | 114.32 |
| 4 | B | 411 | A1H6I | C11-C10-C9 | -2.04 | 106.82 | 111.99 |

There are no chirality outliers.

All (10) torsion outliers are listed below:

For Manuscript Review

| Mol | Chain | Res | Type | Atoms |
|---|---|---|---|---|
| 4 | A | 409 | A1H6I | N6-C7-C8-C9 |
| 4 | A | 409 | A1H6I | C7-C8-C9-C27 |
| 4 | B | 411 | A1H6I | N6-C7-C8-C9 |
| 4 | B | 411 | A1H6I | C4-C5-N6-C7 |
| 4 | A | 409 | A1H6I | C4-C5-N6-C7 |
| 3 | A | 407 | EDO | O1-C1-C2-O2 |
| 4 | B | 411 | A1H6I | C7-C8-C9-C27 |
| 4 | A | 409 | A1H6I | C7-C8-C9-C10 |
| 4 | B | 411 | A1H6I | C7-C8-C9-C10 |
| 4 | A | 409 | A1H6I | C8-C7-N6-C5 |

There are no ring outliers.

3 monomers are involved in 3 short contacts:

| Mol | Chain | Res | Type | Clashes | Symm-Clashes |
|---|---|---|---|---|---|
| 2 | A | 401 | SO4 | 1 | 0 |
| 2 | B | 402 | SO4 | 1 | 0 |
| 2 | B | 408 | SO4 | 1 | 0 |

The following is a two-dimensional graphical depiction of Mogul quality analysis of bond lengths, bond angles, torsion angles, and ring geometry for all instances of the Ligand of Interest. In addition, ligands with molecular weight > 250 and outliers as shown on the validation Tables will also be included. For torsion angles, if less then 5% of the Mogul distribution of torsion angles is within 10 degrees of the torsion angle in question, then that torsion angle is considered an outlier. Any bond that is central to one or more torsion angles identified as an outlier by Mogul will be highlighted in the graph. For rings, the root-mean-square deviation (RMSD) between the ring in question and similar rings identified by Mogul is calculated over all ring torsion angles. If the average RMSD is greater than 60 degrees and the minimal RMSD between the ring in question and any Mogul-identified rings is also greater than 60 degrees, then that ring is considered an outlier. The outliers are highlighted in purple. The color gray indicates Mogul did not find sufficient equivalents in the CSD to analyse the geometry.

For Manuscript Review

**Ligand A1H6I B 411**

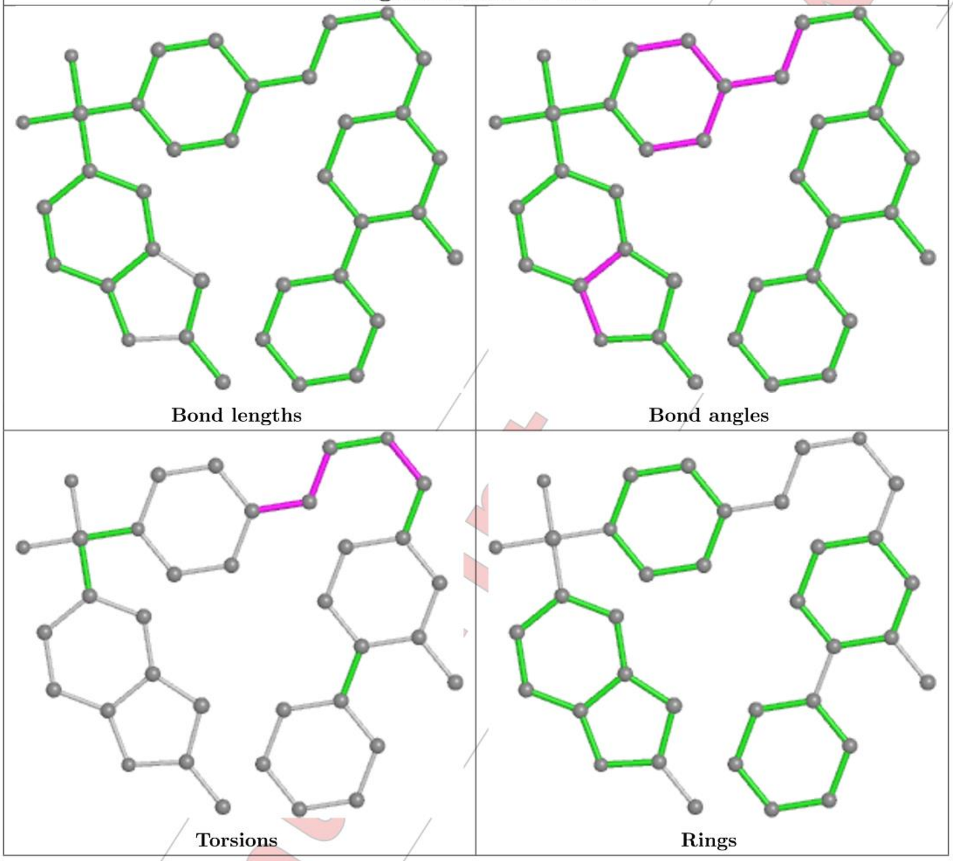

For Manuscript Review

**Ligand A1H6I A 409**

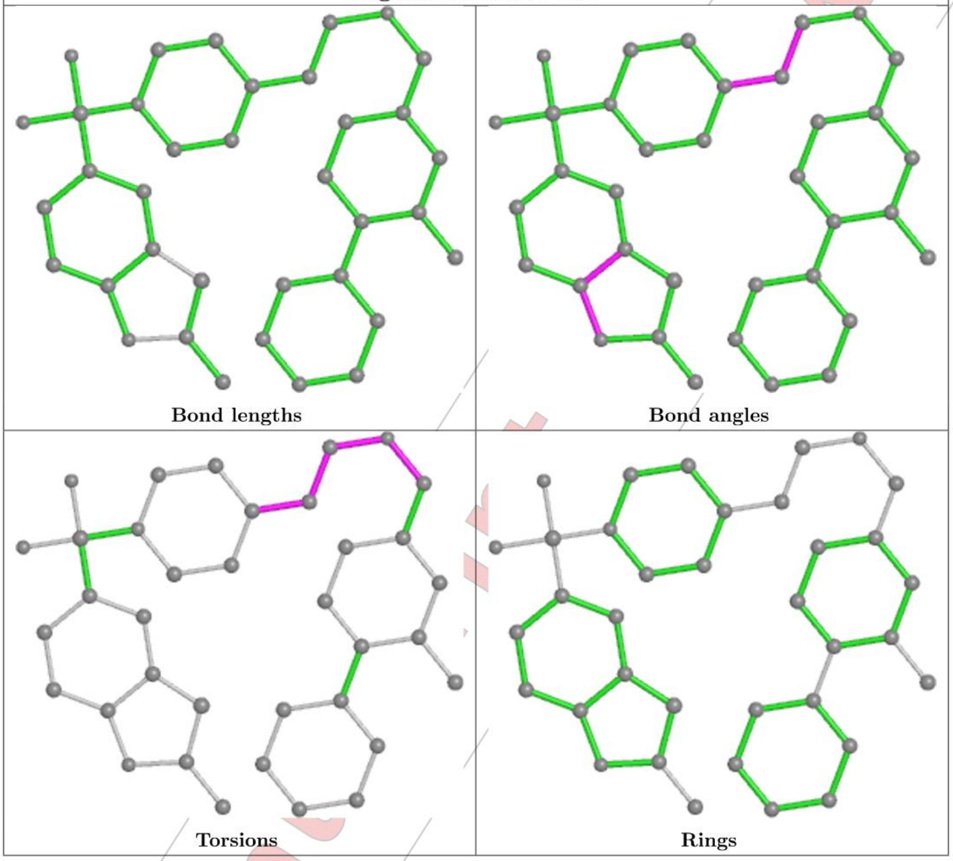

#### 5.7 Other polymers (i)

There are no such residues in this entry.

#### 5.8 Polymer linkage issues (i)

There are no chain breaks in this entry.

For Manuscript Review

### 6 Fit of model and data (i)

#### 6.1 Protein, DNA and RNA chains (i)

In the following table, the column labelled '#RSRZ> 2' contains the number (and percentage) of RSRZ outliers, followed by percent RSRZ outliers for the chain as percentile scores relative to all X-ray entries and entries of similar resolution. The OWAB column contains the minimum, median, $95^{th}$ percentile and maximum values of the occupancy-weighted average B-factor per residue. The column labelled 'Q< 0.9' lists the number of (and percentage) of residues with an average occupancy less than 0.9.

| Mol | Chain | Analysed | <RSRZ> | #RSRZ>2 | | | OWAB(Å$^2$) | Q<0.9 |
|---|---|---|---|---|---|---|---|---|
| 1 | A | 329/338 (97%) | -0.25 | 0 | 100 | 100 | 30, 48, 78, 103 | 0 |
| 1 | B | 329/338 (97%) | -0.14 | 3 (0%) | 84 | 83 | 30, 49, 86, 117 | 0 |
| All | All | 658/676 (97%) | -0.20 | 3 (0%) | 91 | 91 | 30, 48, 83, 117 | 0 |

All (3) RSRZ outliers are listed below:

| Mol | Chain | Res | Type | RSRZ |
|---|---|---|---|---|
| 1 | B | 255 | TYR | 2.9 |
| 1 | B | 269 | PHE | 2.3 |
| 1 | B | 127 | THR | 2.1 |

#### 6.2 Non-standard residues in protein, DNA, RNA chains (i)

There are no non-standard protein/DNA/RNA residues in this entry.

#### 6.3 Carbohydrates (i)

There are no monosaccharides in this entry.

#### 6.4 Ligands (i)

In the following table, the Atoms column lists the number of modelled atoms in the group and the number defined in the chemical component dictionary. The B-factors column lists the minimum, median, $95^{th}$ percentile and maximum values of B factors of atoms in the group. The column labelled 'Q< 0.9' lists the number of atoms with occupancy less than 0.9.

| Mol | Type | Chain | Res | Atoms | RSCC | RSR | B-factors(Å$^2$) | Q<0.9 |
|---|---|---|---|---|---|---|---|---|
| 2 | SO4 | B | 409 | 5/5 | 0.74 | 0.21 | 63,72,86,87 | 0 |
| 2 | SO4 | B | 406 | 5/5 | 0.79 | 0.27 | 74,101,109,109 | 0 |

*Continued on next page...*

For Manuscript Review

*Continued from previous page...*

| Mol | Type | Chain | Res | Atoms | RSCC | RSR | B-factors($Å^2$) | Q<0.9 |
|---|---|---|---|---|---|---|---|---|
| 2 | SO4 | A | 408 | 5/5 | 0.79 | 0.25 | 79,93,101,117 | 0 |
| 2 | SO4 | A | 405 | 5/5 | 0.83 | 0.14 | 78,79,91,103 | 0 |
| 2 | SO4 | B | 404 | 5/5 | 0.83 | 0.25 | 75,83,102,102 | 0 |
| 3 | EDO | A | 407 | 4/4 | 0.83 | 0.17 | 66,68,69,73 | 0 |
| 2 | SO4 | A | 404 | 5/5 | 0.86 | 0.18 | 88,103,105,108 | 0 |
| 2 | SO4 | B | 403 | 5/5 | 0.89 | 0.20 | 88,92,102,104 | 0 |
| 2 | SO4 | B | 405 | 5/5 | 0.90 | 0.15 | 105,113,118,120 | 0 |
| 2 | SO4 | B | 407 | 5/5 | 0.91 | 0.35 | 100,101,107,120 | 0 |
| 4 | A1H6I | A | 409 | 36/36 | 0.91 | 0.20 | 38,49,61,79 | 0 |
| 4 | A1H6I | B | 411 | 36/36 | 0.91 | 0.22 | 48,55,84,91 | 0 |
| 2 | SO4 | B | 408 | 5/5 | 0.94 | 0.17 | 73,76,79,79 | 0 |
| 2 | SO4 | A | 402 | 5/5 | 0.95 | 0.14 | 64,64,75,90 | 0 |
| 2 | SO4 | B | 410 | 5/5 | 0.96 | 0.14 | 65,66,79,83 | 0 |
| 2 | SO4 | B | 402 | 5/5 | 0.96 | 0.08 | 60,69,78,79 | 0 |
| 2 | SO4 | A | 403 | 5/5 | 0.97 | 0.12 | 51,56,62,74 | 0 |
| 2 | SO4 | A | 401 | 5/5 | 0.97 | 0.12 | 42,45,55,57 | 0 |
| 2 | SO4 | B | 401 | 5/5 | 0.97 | 0.11 | 47,52,66,66 | 0 |
| 2 | SO4 | A | 406 | 5/5 | 0.98 | 0.10 | 46,47,59,60 | 0 |

The following is a graphical depiction of the model fit to experimental electron density of all instances of the Ligand of Interest. In addition, ligands with molecular weight > 250 and outliers as shown on the geometry validation Tables will also be included. Each fit is shown from different orientation to approximate a three-dimensional view.

For Manuscript Review

**Electron density around A1H6I A 409:**

$2mF_o$-$DF_c$ (at 0.7 rmsd) in gray
$mF_o$-$DF_c$ (at 3 rmsd) in purple (negative) and green (positive)

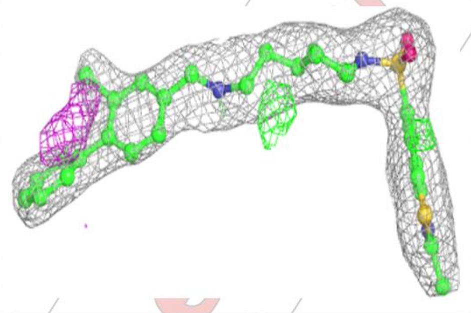

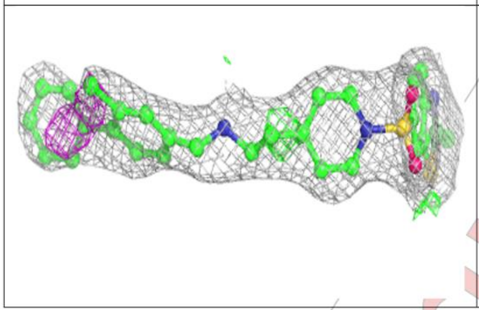

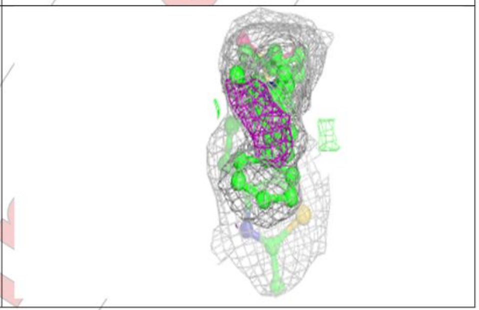

**Electron density around A1H6I B 411:**

$2mF_o$-$DF_c$ (at 0.7 rmsd) in gray
$mF_o$-$DF_c$ (at 3 rmsd) in purple (negative) and green (positive)

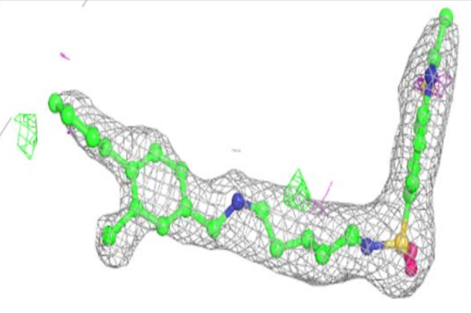

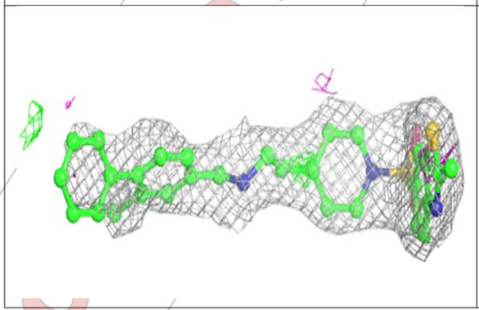

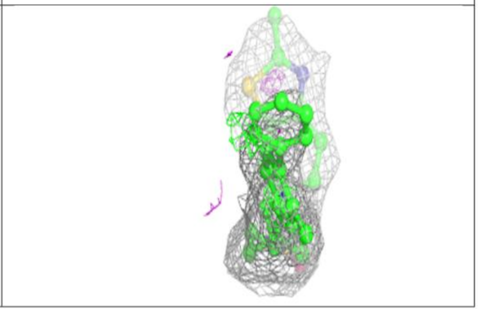

#### 6.5 Other polymers (i)

There are no such residues in this entry.

For Manuscript Review

### Full wwPDB X-ray Structure Validation Report (i)

Mar 20, 2024 – 02:51 pm GMT

| | | |
|---|---|---|
| PDB ID | : | 9EQ0 |
| Title | : | Crystal structure of human Casein Kinase II subunit alpha (CK2a1) in complex with allosteric ligand FGJG12 |
| Deposited on | : | 2024-03-20 |
| Resolution | : | 3.15 Å(reported) |

**This wwPDB validation report is for manuscript review**

This is a Full wwPDB X-ray Structure Validation Report.

This report is produced by the wwPDB biocuration pipeline after annotation of the structure.

We welcome your comments at **
A user guide is available at
https://www.wwpdb.org/validation/2017/XrayValidationReportHelp
with specific help available everywhere you see the (i) symbol.

The types of validation reports are described at
http://www.wwpdb.org/validation/2017/FAQs#types.

The following versions of software and data (see references (i)) were used in the production of this report:

| | | |
|---|---|---|
| MolProbity | : | 4.02b-467 |
| Mogul | : | 1.8.4, CSD as541be (2020) |
| Xtriage (Phenix) | : | 1.13 |
| EDS | : | 2.36 |
| buster-report | : | 1.1.7 (2018) |
| Percentile statistics | : | 20191225.v01 (using entries in the PDB archive December 25th 2019) |
| Refmac | : | 5.8.0158 |
| CCP4 | : | 7.0.044 (Gargrove) |
| Ideal geometry (proteins) | : | Engh & Huber (2001) |

For Manuscript Review

### 1 Overall quality at a glance (i)

The following experimental techniques were used to determine the structure:
*X-RAY DIFFRACTION*

The reported resolution of this entry is 3.15 Å.

Percentile scores (ranging between 0-100) for global validation metrics of the entry are shown in the following graphic. The table shows the number of entries on which the scores are based.

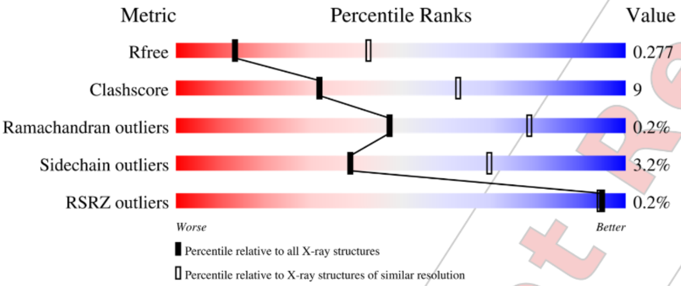

| Metric | Whole archive (#Entries) | Similar resolution (#Entries, resolution range(Å)) |
|---|---|---|
| $R_{free}$ | 130704 | 1665 (3.20-3.12) |
| Clashscore | 141614 | 1804 (3.20-3.12) |
| Ramachandran outliers | 138981 | 1770 (3.20-3.12) |
| Sidechain outliers | 138945 | 1769 (3.20-3.12) |
| RSRZ outliers | 127900 | 1616 (3.20-3.12) |

The table below summarises the geometric issues observed across the polymeric chains and their fit to the electron density. The red, orange, yellow and green segments of the lower bar indicate the fraction of residues that contain outliers for >=3, 2, 1 and 0 types of geometric quality criteria respectively. A grey segment represents the fraction of residues that are not modelled. The numeric value for each fraction is indicated below the corresponding segment, with a dot representing fractions <=5% The upper red bar (where present) indicates the fraction of residues that have poor fit to the electron density. The numeric value is given above the bar.

| Mol | Chain | Length | Quality of chain |
|---|---|---|---|
| 1 | A | 338 | 79% 15% • 5% |
| 1 | B | 338 | 81% 14% • • |

Ideal geometry (DNA, RNA) : Parkinson et al. (1996)
Validation Pipeline (wwPDB-VP) : 2.36

For Manuscript Review

#### 2 Entry composition (i)

There are 4 unique types of molecules in this entry. The entry contains 5285 atoms, of which 0 are hydrogens and 0 are deuteriums.

In the tables below, the ZeroOcc column contains the number of atoms modelled with zero occupancy, the AltConf column contains the number of residues with at least one atom in alternate conformation and the Trace column contains the number of residues modelled with at most 2 atoms.

- Molecule 1 is a protein called Casein kinase II subunit alpha.

| Mol | Chain | Residues | Atoms | | | | | ZeroOcc | AltConf | Trace |
|---|---|---|---|---|---|---|---|---|---|---|
| 1 | A | 321 | Total 2597 | C 1672 | N 442 | O 472 | S 11 | 0 | 0 | 0 |
| 1 | B | 324 | Total 2590 | C 1663 | N 445 | O 471 | S 11 | 0 | 0 | 0 |

There are 2 discrepancies between the modelled and reference sequences:

| Chain | Residue | Modelled | Actual | Comment | Reference |
|---|---|---|---|---|---|
| A | 0 | SER | - | expression tag | UNP P68400 |
| B | 0 | SER | - | expression tag | UNP P68400 |

- Molecule 2 is SULFATE ION (three-letter code: SO4) (formula: $O_4S$).

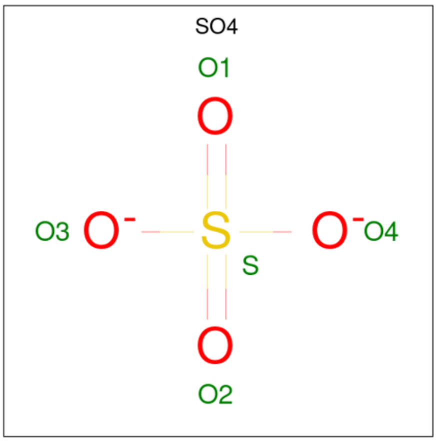

| Mol | Chain | Residues | Atoms | | | ZeroOcc | AltConf |
|---|---|---|---|---|---|---|---|
| 2 | A | 1 | Total 5 | O 4 | S 1 | 0 | 0 |

Continued on next page...

For Manuscript Review

Continued from previous page...

| Mol | Chain | Residues | Atoms | ZeroOcc | AltConf |
|---|---|---|---|---|---|
| 2 | A | 1 | Total O S<br>5 4 1 | 0 | 0 |
| 2 | A | 1 | Total O S<br>5 4 1 | 0 | 0 |
| 2 | B | 1 | Total O S<br>5 4 1 | 0 | 0 |
| 2 | B | 1 | Total O S<br>5 4 1 | 0 | 0 |
| 2 | B | 1 | Total O S<br>5 4 1 | 0 | 0 |

- Molecule 3 is {N}-[4-[(3-chloranyl-4-phenyl-phenyl)methylamino]butyl]isoquinoline-5-sulfonamide (three-letter code: A1H6D) (formula: $C_{26}H_{26}ClN_3O_2S$) (labeled as "Ligand of Interest" by depositor).

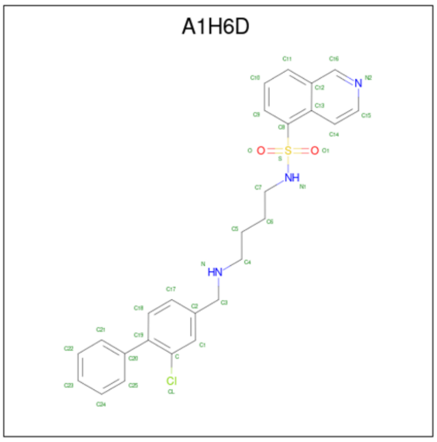

| Mol | Chain | Residues | Atoms | ZeroOcc | AltConf |
|---|---|---|---|---|---|
| 3 | A | 1 | Total C Cl N O S<br>33 26 1 3 2 1 | 0 | 0 |
| 3 | B | 1 | Total C Cl N O S<br>33 26 1 3 2 1 | 0 | 0 |

- Molecule 4 is water.

| Mol | Chain | Residues | Atoms | ZeroOcc | AltConf |
|---|---|---|---|---|---|
| 4 | A | 1 | Total O<br>1 1 | 0 | 0 |

Continued on next page...

For Manuscript Review

*Continued from previous page...*

| Mol | Chain | Residues | Atoms | ZeroOcc | AltConf |
|---|---|---|---|---|---|
| 4 | B | 1 | Total O<br>1 1 | 0 | 0 |

For Manuscript Review

### 3 Residue-property plots (i)

These plots are drawn for all protein, RNA, DNA and oligosaccharide chains in the entry. The first graphic for a chain summarises the proportions of the various outlier classes displayed in the second graphic. The second graphic shows the sequence view annotated by issues in geometry and electron density. Residues are color-coded according to the number of geometric quality criteria for which they contain at least one outlier: green = 0, yellow = 1, orange = 2 and red = 3 or more. A red dot above a residue indicates a poor fit to the electron density (RSRZ > 2). Stretches of 2 or more consecutive residues without any outlier are shown as a green connector. Residues present in the sample, but not in the model, are shown in grey.

- Molecule 1: Casein kinase II subunit alpha

Chain A:

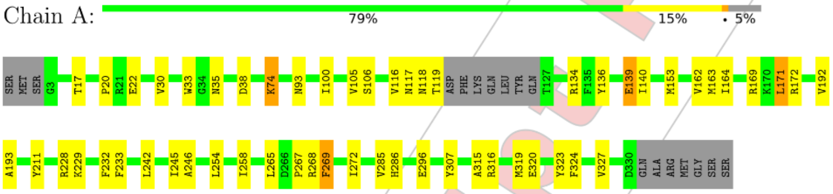

- Molecule 1: Casein kinase II subunit alpha

Chain B:

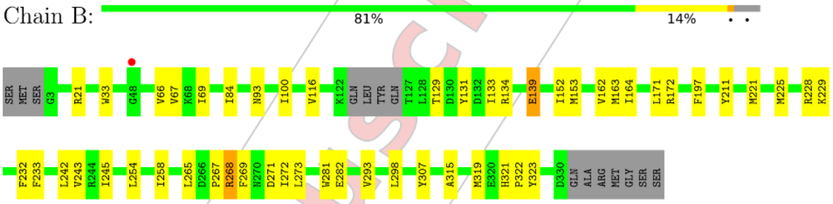

### 4 Data and refinement statistics (i)

| Property | Value | Source |
|---|---|---|
| Space group | P 43 21 2 | Depositor |
| Cell constants<br>a, b, c, $\alpha$, $\beta$, $\gamma$ | 126.50Å 126.50Å 126.44Å<br>90.00° 90.00° 90.00° | Depositor |
| Resolution (Å) | 44.76 – 3.15<br>44.72 – 3.15 | Depositor<br>EDS |
| % Data completeness<br>(in resolution range) | 99.9 (44.76-3.15)<br>100.0 (44.72-3.15) | Depositor<br>EDS |
| $R_{merge}$ | (Not available) | Depositor |
| $R_{sym}$ | (Not available) | Depositor |
| $< I/\sigma(I) >$ [1] | 1.98 (at 3.12Å) | Xtriage |
| Refinement program | REFMAC 5.8.0419 | Depositor |
| R, $R_{free}$ | 0.255 , 0.274<br>0.262 , 0.277 | Depositor<br>DCC |
| $R_{free}$ test set | 962 reflections (5.25%) | wwPDB-VP |
| Wilson B-factor (Å$^2$) | 55.3 | Xtriage |
| Anisotropy | 0.599 | Xtriage |
| Bulk solvent $k_{sol}$(e/Å$^3$), $B_{sol}$(Å$^2$) | 0.30 , 35.6 | EDS |
| L-test for twinning[2] | $< \|L\| > = 0.47$, $< L^2 > = 0.30$ | Xtriage |
| Estimated twinning fraction | 0.000 for -h,-l,-k<br>0.000 for l,-k,h | Xtriage |
| $F_o$,$F_c$ correlation | 0.89 | EDS |
| Total number of atoms | 5285 | wwPDB-VP |
| Average B, all atoms (Å$^2$) | 57.0 | wwPDB-VP |

Xtriage's analysis on translational NCS is as follows: *The analyses of the Patterson function reveals a significant off-origin peak that is 32.05 % of the origin peak, indicating pseudo-translational symmetry. The chance of finding a peak of this or larger height randomly in a structure without pseudo-translational symmetry is equal to 1.0005e-03. The detected translational NCS is most likely also responsible for the elevated intensity ratio.*

[1] Intensities estimated from amplitudes.
[2] Theoretical values of $< |L| >$, $< L^2 >$ for acentric reflections are 0.5, 0.333 respectively for untwinned datasets, and 0.375, 0.2 for perfectly twinned datasets.

For Manuscript Review

### 5 Model quality (i)

#### 5.1 Standard geometry (i)

Bond lengths and bond angles in the following residue types are not validated in this section: A1H6D, SO4

The Z score for a bond length (or angle) is the number of standard deviations the observed value is removed from the expected value. A bond length (or angle) with $|Z| > 5$ is considered an outlier worth inspection. RMSZ is the root-mean-square of all Z scores of the bond lengths (or angles).

| Mol | Chain | Bond lengths | | Bond angles | |
|---|---|---|---|---|---|
| | | RMSZ | $\#\|Z\|>5$ | RMSZ | $\#\|Z\|>5$ |
| 1 | A | 0.43 | 0/2668 | 0.76 | 0/3629 |
| 1 | B | 0.40 | 0/2659 | 0.76 | 0/3618 |
| All | All | 0.42 | 0/5327 | 0.76 | 0/7247 |

There are no bond length outliers.

There are no bond angle outliers.

There are no chirality outliers.

There are no planarity outliers.

#### 5.2 Too-close contacts (i)

In the following table, the Non-H and H(model) columns list the number of non-hydrogen atoms and hydrogen atoms in the chain respectively. The H(added) column lists the number of hydrogen atoms added and optimized by MolProbity. The Clashes column lists the number of clashes within the asymmetric unit, whereas Symm-Clashes lists symmetry-related clashes.

| Mol | Chain | Non-H | H(model) | H(added) | Clashes | Symm-Clashes |
|---|---|---|---|---|---|---|
| 1 | A | 2597 | 0 | 2445 | 44 | 0 |
| 1 | B | 2590 | 0 | 2419 | 45 | 0 |
| 2 | A | 15 | 0 | 0 | 0 | 0 |
| 2 | B | 15 | 0 | 0 | 0 | 0 |
| 3 | A | 33 | 0 | 0 | 5 | 0 |
| 3 | B | 33 | 0 | 0 | 3 | 0 |
| 4 | A | 1 | 0 | 0 | 0 | 0 |
| 4 | B | 1 | 0 | 0 | 0 | 0 |
| All | All | 5285 | 0 | 4864 | 90 | 0 |

The all-atom clashscore is defined as the number of clashes found per 1000 atoms (including hydrogen atoms). The all-atom clashscore for this structure is 9.

For Manuscript Review

All (90) close contacts within the same asymmetric unit are listed below, sorted by their clash magnitude.

| Atom-1 | Atom-2 | Interatomic distance (Å) | Clash overlap (Å) |
|---|---|---|---|
| 1:B:139:GLU:HB3 | 1:B:171:LEU:HD23 | 1.47 | 0.95 |
| 1:B:268:ARG:NH2 | 1:B:272:ILE:HG21 | 1.95 | 0.82 |
| 1:B:281:TRP:HB3 | 1:B:298:LEU:CD1 | 2.13 | 0.79 |
| 1:B:281:TRP:HB3 | 1:B:298:LEU:HD12 | 1.66 | 0.78 |
| 1:B:281:TRP:CB | 1:B:298:LEU:HD12 | 2.14 | 0.76 |
| 1:A:140:ILE:HG13 | 1:A:171:LEU:HD21 | 1.67 | 0.75 |
| 1:A:139:GLU:HB3 | 1:A:171:LEU:HD23 | 1.70 | 0.74 |
| 1:B:268:ARG:HH21 | 1:B:268:ARG:HG2 | 1.52 | 0.74 |
| 1:B:268:ARG:HH21 | 1:B:268:ARG:CG | 1.99 | 0.73 |
| 1:A:118:ASN:HA | 1:A:163:MET:SD | 2.30 | 0.71 |
| 1:B:293:VAL:HG13 | 1:B:298:LEU:CD2 | 2.22 | 0.70 |
| 1:B:164:ILE:HG12 | 1:B:171:LEU:HD13 | 1.76 | 0.68 |
| 1:B:162:VAL:O | 3:B:404:A1H6D:C4 | 2.43 | 0.67 |
| 1:B:293:VAL:HG13 | 1:B:298:LEU:HD21 | 1.77 | 0.67 |
| 1:B:172:ARG:HG2 | 1:B:172:ARG:HH11 | 1.63 | 0.63 |
| 1:A:163:MET:HE3 | 3:A:404:A1H6D:C11 | 2.29 | 0.63 |
| 1:A:20:PRO:HB2 | 1:A:22:GLU:HG2 | 1.82 | 0.62 |
| 1:A:315:ALA:O | 1:A:319:MET:HG3 | 2.00 | 0.61 |
| 1:B:268:ARG:HH22 | 1:B:272:ILE:HG21 | 1.66 | 0.61 |
| 1:B:268:ARG:NH2 | 1:B:268:ARG:CG | 2.63 | 0.61 |
| 1:A:267:PRO:O | 1:A:269:PHE:N | 2.33 | 0.60 |
| 1:A:119:THR:O | 1:A:119:THR:HG22 | 2.01 | 0.60 |
| 1:B:242:LEU:HD11 | 1:B:254:LEU:HD11 | 1.83 | 0.59 |
| 1:B:242:LEU:HD11 | 1:B:254:LEU:CD1 | 2.33 | 0.59 |
| 1:B:162:VAL:CG1 | 1:B:171:LEU:HD11 | 2.33 | 0.58 |
| 1:B:281:TRP:HB3 | 1:B:298:LEU:HD11 | 1.87 | 0.56 |
| 1:A:172:ARG:HH11 | 1:A:172:ARG:HG2 | 1.70 | 0.56 |
| 1:B:162:VAL:HG13 | 1:B:171:LEU:HD11 | 1.89 | 0.55 |
| 1:A:163:MET:HE1 | 3:A:404:A1H6D:C10 | 2.37 | 0.54 |
| 1:A:163:MET:O | 1:A:164:ILE:HD13 | 2.08 | 0.54 |
| 1:A:324:PHE:O | 1:A:327:VAL:HG22 | 2.08 | 0.54 |
| 3:B:404:A1H6D:CL | 3:B:404:A1H6D:C25 | 2.94 | 0.53 |
| 1:A:254:LEU:O | 1:A:258:ILE:HG12 | 2.09 | 0.53 |
| 1:A:242:LEU:HD11 | 1:A:254:LEU:HD11 | 1.90 | 0.52 |
| 1:A:163:MET:HE1 | 3:A:404:A1H6D:C9 | 2.40 | 0.52 |
| 1:A:33:TRP:CE3 | 1:A:100:ILE:HG22 | 2.43 | 0.52 |
| 1:A:153:MET:HG2 | 1:A:211:TYR:HA | 1.91 | 0.52 |
| 1:B:254:LEU:O | 1:B:258:ILE:HG12 | 2.10 | 0.51 |
| 1:B:269:PHE:HA | 1:B:272:ILE:HG12 | 1.91 | 0.51 |
| 1:A:269:PHE:HA | 1:A:272:ILE:HG12 | 1.93 | 0.51 |

*Continued on next page...*

For Manuscript Review

*Continued from previous page...*

| Atom-1 | Atom-2 | Interatomic distance (Å) | Clash overlap (Å) |
|---|---|---|---|
| 1:B:33:TRP:CE3 | 1:B:100:ILE:HG22 | 2.45 | 0.51 |
| 1:A:163:MET:CE | 3:A:404:A1H6D:C11 | 2.89 | 0.50 |
| 1:A:171:LEU:C | 1:A:171:LEU:HD12 | 2.32 | 0.50 |
| 1:A:242:LEU:HD11 | 1:A:254:LEU:CD1 | 2.42 | 0.50 |
| 1:B:163:MET:O | 1:B:171:LEU:HD12 | 2.11 | 0.50 |
| 1:B:315:ALA:O | 1:B:319:MET:HG3 | 2.12 | 0.50 |
| 1:B:254:LEU:HD12 | 1:B:307:TYR:CE1 | 2.47 | 0.49 |
| 1:B:229:LYS:HD2 | 1:B:232:PHE:HA | 1.96 | 0.48 |
| 1:B:221:MET:O | 1:B:225:MET:HG2 | 2.13 | 0.48 |
| 1:A:93:ASN:HD21 | 1:A:139:GLU:HG3 | 1.78 | 0.47 |
| 1:A:163:MET:CE | 3:A:404:A1H6D:C10 | 2.93 | 0.47 |
| 1:B:321:HIS:CG | 1:B:322:PRO:HD2 | 2.50 | 0.46 |
| 1:B:93:ASN:HD21 | 1:B:139:GLU:HG3 | 1.80 | 0.46 |
| 1:A:229:LYS:HD2 | 1:A:232:PHE:HA | 1.98 | 0.46 |
| 1:A:171:LEU:HD12 | 1:A:172:ARG:N | 2.30 | 0.46 |
| 1:A:74:LYS:HB3 | 1:A:74:LYS:HE3 | 1.66 | 0.45 |
| 1:A:119:THR:HB | 1:A:164:ILE:O | 2.17 | 0.45 |
| 1:A:254:LEU:HD12 | 1:A:307:TYR:CE1 | 2.51 | 0.45 |
| 1:A:162:VAL:HG13 | 1:A:171:LEU:HD13 | 1.99 | 0.44 |
| 1:A:172:ARG:HG2 | 1:A:172:ARG:NH1 | 2.30 | 0.44 |
| 1:B:129:THR:HG22 | 1:B:131:TYR:H | 1.82 | 0.44 |
| 1:B:172:ARG:HG2 | 1:B:172:ARG:NH1 | 2.31 | 0.44 |
| 1:B:129:THR:O | 1:B:133:ILE:HD13 | 2.16 | 0.44 |
| 1:A:162:VAL:HG12 | 1:A:164:ILE:HD11 | 2.00 | 0.43 |
| 1:B:243:VAL:HA | 1:B:273:LEU:HD23 | 1.99 | 0.43 |
| 1:A:316:ARG:O | 1:A:320:GLU:HG3 | 2.19 | 0.43 |
| 1:A:285:VAL:O | 1:A:286:HIS:HD2 | 2.01 | 0.43 |
| 1:B:197:PHE:CD1 | 1:B:197:PHE:N | 2.87 | 0.43 |
| 1:A:93:ASN:ND2 | 1:A:139:GLU:HG3 | 2.33 | 0.43 |
| 1:A:116:VAL:HG12 | 1:A:117:ASN:O | 2.19 | 0.43 |
| 1:A:162:VAL:HG13 | 1:A:171:LEU:CD1 | 2.50 | 0.42 |
| 1:B:134:ARG:HG2 | 1:B:323:TYR:CE1 | 2.55 | 0.42 |
| 1:A:192:VAL:O | 1:A:193:ALA:HB3 | 2.19 | 0.42 |
| 1:B:67:VAL:HG12 | 1:B:69:ILE:HD13 | 2.01 | 0.42 |
| 1:A:246:ALA:HA | 1:A:307:TYR:OH | 2.20 | 0.41 |
| 1:B:66:VAL:HG11 | 3:B:404:A1H6D:C16 | 2.49 | 0.41 |
| 1:A:134:ARG:HG2 | 1:A:323:TYR:CE1 | 2.54 | 0.41 |
| 1:A:265:LEU:HD23 | 1:A:265:LEU:HA | 1.88 | 0.41 |
| 1:B:84:ILE:HG23 | 1:B:152:ILE:HD13 | 2.01 | 0.41 |
| 1:A:140:ILE:CG1 | 1:A:171:LEU:HD21 | 2.45 | 0.41 |
| 1:B:93:ASN:ND2 | 1:B:139:GLU:HG3 | 2.35 | 0.41 |

*Continued on next page...*

For Manuscript Review

*Continued from previous page...*

| Atom-1 | Atom-2 | Interatomic distance (Å) | Clash overlap (Å) |
|---|---|---|---|
| 1:B:233:PHE:CD1 | 1:B:245:ILE:HA | 2.56 | 0.41 |
| 1:B:153:MET:HG2 | 1:B:211:TYR:HA | 2.02 | 0.41 |
| 1:B:267:PRO:O | 1:B:268:ARG:C | 2.58 | 0.41 |
| 1:B:281:TRP:CG | 1:B:298:LEU:HD12 | 2.54 | 0.40 |
| 1:A:233:PHE:CE1 | 1:A:245:ILE:HA | 2.56 | 0.40 |
| 1:B:268:ARG:HG2 | 1:B:272:ILE:HD13 | 2.03 | 0.40 |
| 1:B:265:LEU:HD12 | 1:B:265:LEU:HA | 1.78 | 0.40 |
| 1:A:105:VAL:HG13 | 1:A:106:SER:H | 1.87 | 0.40 |
| 1:A:136:TYR:OH | 1:A:169:ARG:HA | 2.21 | 0.40 |

There are no symmetry-related clashes.

#### 5.3 Torsion angles (i)

##### 5.3.1 Protein backbone (i)

In the following table, the Percentiles column shows the percent Ramachandran outliers of the chain as a percentile score with respect to all X-ray entries followed by that with respect to entries of similar resolution.

The Analysed column shows the number of residues for which the backbone conformation was analysed, and the total number of residues.

| Mol | Chain | Analysed | Favoured | Allowed | Outliers | Percentiles | |
|---|---|---|---|---|---|---|---|
| 1 | A | 317/338 (94%) | 303 (96%) | 13 (4%) | 1 (0%) | 41 | 73 |
| 1 | B | 320/338 (95%) | 305 (95%) | 15 (5%) | 0 | 100 | 100 |
| All | All | 637/676 (94%) | 608 (95%) | 28 (4%) | 1 (0%) | 47 | 78 |

All (1) Ramachandran outliers are listed below:

| Mol | Chain | Res | Type |
|---|---|---|---|
| 1 | A | 268 | ARG |

##### 5.3.2 Protein sidechains (i)

In the following table, the Percentiles column shows the percent sidechain outliers of the chain as a percentile score with respect to all X-ray entries followed by that with respect to entries of similar resolution.

The Analysed column shows the number of residues for which the sidechain conformation was

For Manuscript Review

analysed, and the total number of residues.

| Mol | Chain | Analysed | Rotameric | Outliers | Percentiles | |
|---|---|---|---|---|---|---|
| 1 | A | 270/310 (87%) | 260 (96%) | 10 (4%) | 34 | 66 |
| 1 | B | 265/310 (86%) | 258 (97%) | 7 (3%) | 46 | 74 |
| All | All | 535/620 (86%) | 518 (97%) | 17 (3%) | 39 | 70 |

All (17) residues with a non-rotameric sidechain are listed below:

| Mol | Chain | Res | Type |
|---|---|---|---|
| 1 | A | 17 | THR |
| 1 | A | 30 | VAL |
| 1 | A | 35 | ASN |
| 1 | A | 38 | ASP |
| 1 | A | 74 | LYS |
| 1 | A | 139 | GLU |
| 1 | A | 171 | LEU |
| 1 | A | 228 | ARG |
| 1 | A | 269 | PHE |
| 1 | A | 296 | GLU |
| 1 | B | 21 | ARG |
| 1 | B | 116 | VAL |
| 1 | B | 139 | GLU |
| 1 | B | 228 | ARG |
| 1 | B | 268 | ARG |
| 1 | B | 271 | ASP |
| 1 | B | 282 | GLU |

Sometimes sidechains can be flipped to improve hydrogen bonding and reduce clashes. All (6) such sidechains are listed below:

| Mol | Chain | Res | Type |
|---|---|---|---|
| 1 | A | 35 | ASN |
| 1 | A | 93 | ASN |
| 1 | A | 286 | HIS |
| 1 | B | 93 | ASN |
| 1 | B | 168 | HIS |
| 1 | B | 286 | HIS |

##### 5.3.3 RNA ⓘ

There are no RNA molecules in this entry.

For Manuscript Review

##### 5.4 Non-standard residues in protein, DNA, RNA chains (i)

There are no non-standard protein/DNA/RNA residues in this entry.

##### 5.5 Carbohydrates (i)

There are no monosaccharides in this entry.

##### 5.6 Ligand geometry (i)

8 ligands are modelled in this entry.

In the following table, the Counts columns list the number of bonds (or angles) for which Mogul statistics could be retrieved, the number of bonds (or angles) that are observed in the model and the number of bonds (or angles) that are defined in the Chemical Component Dictionary. The Link column lists molecule types, if any, to which the group is linked. The Z score for a bond length (or angle) is the number of standard deviations the observed value is removed from the expected value. A bond length (or angle) with $|Z| > 2$ is considered an outlier worth inspection. RMSZ is the root-mean-square of all Z scores of the bond lengths (or angles).

| Mol | Type | Chain | Res | Link | Bond lengths | | | Bond angles | | |
|---|---|---|---|---|---|---|---|---|---|---|
| | | | | | Counts | RMSZ | $\#\|Z\| > 2$ | Counts | RMSZ | $\#\|Z\| > 2$ |
| 2 | SO4 | B | 402 | - | 4,4,4 | 0.35 | 0 | 6,6,6 | 0.08 | 0 |
| 2 | SO4 | A | 402 | - | 4,4,4 | 0.32 | 0 | 6,6,6 | 0.12 | 0 |
| 3 | A1H6D | B | 404 | - | 36,36,36 | 0.19 | 0 | 49,49,49 | 0.38 | 0 |
| 2 | SO4 | B | 401 | - | 4,4,4 | 0.34 | 0 | 6,6,6 | 0.07 | 0 |
| 3 | A1H6D | A | 404 | - | 36,36,36 | 0.19 | 0 | 49,49,49 | 0.38 | 0 |
| 2 | SO4 | A | 401 | - | 4,4,4 | 0.36 | 0 | 6,6,6 | 0.17 | 0 |
| 2 | SO4 | A | 403 | - | 4,4,4 | 0.37 | 0 | 6,6,6 | 0.09 | 0 |
| 2 | SO4 | B | 403 | - | 4,4,4 | 0.35 | 0 | 6,6,6 | 0.18 | 0 |

In the following table, the Chirals column lists the number of chiral outliers, the number of chiral centers analysed, the number of these observed in the model and the number defined in the Chemical Component Dictionary. Similar counts are reported in the Torsion and Rings columns. '-' means no outliers of that kind were identified.

| Mol | Type | Chain | Res | Link | Chirals | Torsions | Rings |
|---|---|---|---|---|---|---|---|
| 3 | A1H6D | B | 404 | - | - | 11/21/21/21 | 0/4/4/4 |
| 3 | A1H6D | A | 404 | - | - | 8/21/21/21 | 0/4/4/4 |

There are no bond length outliers.

There are no bond angle outliers.

There are no chirality outliers.

For Manuscript Review

All (19) torsion outliers are listed below:

| Mol | Chain | Res | Type | Atoms |
|---|---|---|---|---|
| 3 | A | 404 | A1H6D | C5-C6-C7-N1 |
| 3 | B | 404 | A1H6D | C13-C8-S-O1 |
| 3 | A | 404 | A1H6D | C2-C3-N-C4 |
| 3 | B | 404 | A1H6D | N-C4-C5-C6 |
| 3 | B | 404 | A1H6D | C9-C8-S-O1 |
| 3 | B | 404 | A1H6D | C9-C8-S-O |
| 3 | B | 404 | A1H6D | C13-C8-S-O |
| 3 | B | 404 | A1H6D | C2-C3-N-C4 |
| 3 | B | 404 | A1H6D | C9-C8-S-N1 |
| 3 | B | 404 | A1H6D | C13-C8-S-N1 |
| 3 | B | 404 | A1H6D | C4-C5-C6-C7 |
| 3 | A | 404 | A1H6D | C-C19-C20-C25 |
| 3 | A | 404 | A1H6D | C5-C4-N-C3 |
| 3 | A | 404 | A1H6D | C13-C8-S-O1 |
| 3 | A | 404 | A1H6D | C7-N1-S-O1 |
| 3 | B | 404 | A1H6D | C5-C4-N-C3 |
| 3 | B | 404 | A1H6D | C7-N1-S-O |
| 3 | A | 404 | A1H6D | C13-C8-S-O |
| 3 | A | 404 | A1H6D | C7-N1-S-O |

There are no ring outliers.

2 monomers are involved in 8 short contacts:

| Mol | Chain | Res | Type | Clashes | Symm-Clashes |
|---|---|---|---|---|---|
| 3 | B | 404 | A1H6D | 3 | 0 |
| 3 | A | 404 | A1H6D | 5 | 0 |

The following is a two-dimensional graphical depiction of Mogul quality analysis of bond lengths, bond angles, torsion angles, and ring geometry for all instances of the Ligand of Interest. In addition, ligands with molecular weight > 250 and outliers as shown on the validation Tables will also be included. For torsion angles, if less then 5% of the Mogul distribution of torsion angles is within 10 degrees of the torsion angle in question, then that torsion angle is considered an outlier. Any bond that is central to one or more torsion angles identified as an outlier by Mogul will be highlighted in the graph. For rings, the root-mean-square deviation (RMSD) between the ring in question and similar rings identified by Mogul is calculated over all ring torsion angles. If the average RMSD is greater than 60 degrees and the minimal RMSD between the ring in question and any Mogul-identified rings is also greater than 60 degrees, then that ring is considered an outlier. The outliers are highlighted in purple. The color gray indicates Mogul did not find sufficient equivalents in the CSD to analyse the geometry.

For Manuscript Review

**Ligand A1H6D B 404**

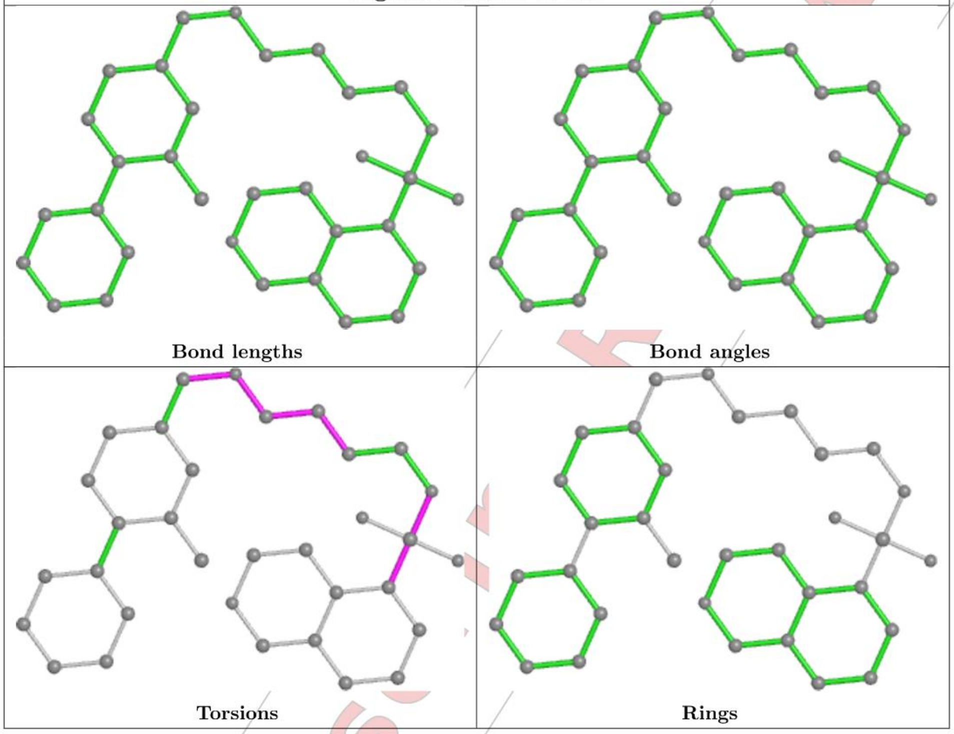

For Manuscript Review

**Ligand A1H6D A 404**

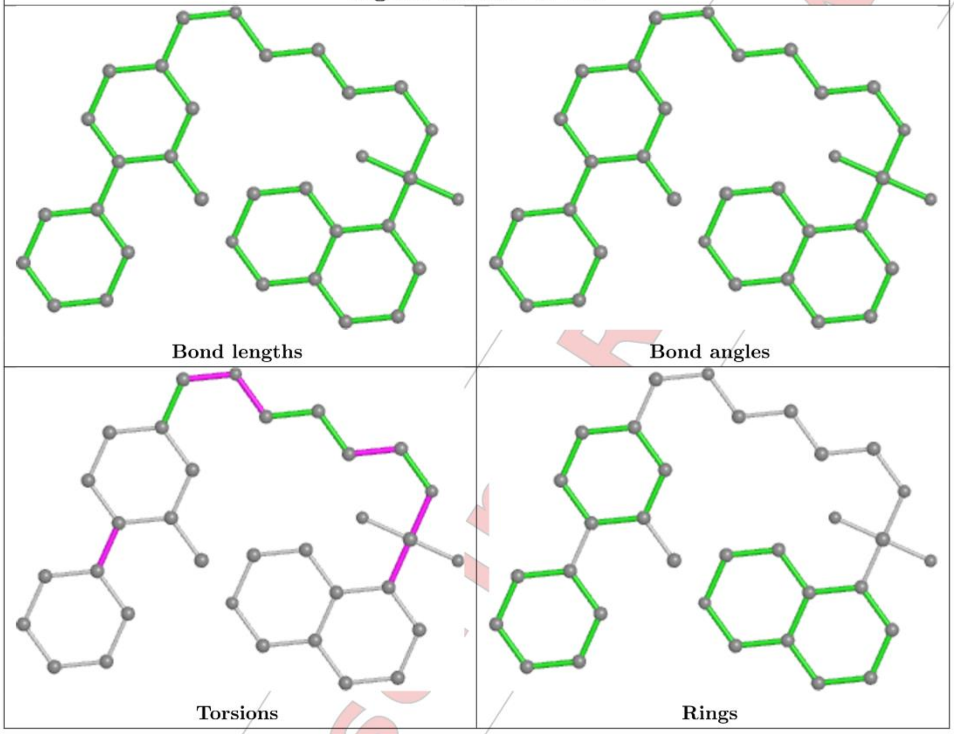

#### 5.7 Other polymers (i)

There are no such residues in this entry.

#### 5.8 Polymer linkage issues (i)

There are no chain breaks in this entry.

For Manuscript Review

### 6 Fit of model and data (i)

#### 6.1 Protein, DNA and RNA chains (i)

In the following table, the column labelled '#RSRZ> 2' contains the number (and percentage) of RSRZ outliers, followed by percent RSRZ outliers for the chain as percentile scores relative to all X-ray entries and entries of similar resolution. The OWAB column contains the minimum, median, 95$^{th}$ percentile and maximum values of the occupancy-weighted average B-factor per residue. The column labelled 'Q< 0.9' lists the number of (and percentage) of residues with an average occupancy less than 0.9.

| Mol | Chain | Analysed | <RSRZ> | #RSRZ>2 | | | OWAB(Å$^2$) | Q<0.9 |
|---|---|---|---|---|---|---|---|---|
| 1 | A | 321/338 (94%) | -0.27 | 0 | 100 | 100 | 30, 55, 82, 110 | 0 |
| 1 | B | 324/338 (95%) | -0.24 | 1 (0%) | 94 | 92 | 36, 54, 85, 114 | 0 |
| All | All | 645/676 (95%) | -0.25 | 1 (0%) | 95 | 94 | 30, 55, 84, 114 | 0 |

All (1) RSRZ outliers are listed below:

| Mol | Chain | Res | Type | RSRZ |
|---|---|---|---|---|
| 1 | B | 48 | GLY | 2.2 |

#### 6.2 Non-standard residues in protein, DNA, RNA chains (i)

There are no non-standard protein/DNA/RNA residues in this entry.

#### 6.3 Carbohydrates (i)

There are no monosaccharides in this entry.

#### 6.4 Ligands (i)

In the following table, the Atoms column lists the number of modelled atoms in the group and the number defined in the chemical component dictionary. The B-factors column lists the minimum, median, 95$^{th}$ percentile and maximum values of B factors of atoms in the group. The column labelled 'Q< 0.9' lists the number of atoms with occupancy less than 0.9.

| Mol | Type | Chain | Res | Atoms | RSCC | RSR | B-factors(Å$^2$) | Q<0.9 |
|---|---|---|---|---|---|---|---|---|
| 2 | SO4 | A | 402 | 5/5 | 0.86 | 0.21 | 80,88,94,97 | 0 |
| 2 | SO4 | A | 403 | 5/5 | 0.86 | 0.29 | 82,89,94,105 | 0 |
| 2 | SO4 | B | 403 | 5/5 | 0.88 | 0.23 | 80,80,86,89 | 0 |
| 2 | SO4 | B | 402 | 5/5 | 0.89 | 0.20 | 81,84,87,97 | 0 |

*Continued on next page...*

For Manuscript Review

*Continued from previous page...*

| Mol | Type | Chain | Res | Atoms | RSCC | RSR | B-factors($Å^2$) | Q<0.9 |
|---|---|---|---|---|---|---|---|---|
| 3 | A1H6D | A | 404 | 33/33 | 0.90 | 0.26 | 71,81,99,106 | 0 |
| 3 | A1H6D | B | 404 | 33/33 | 0.90 | 0.32 | 66,79,91,101 | 0 |
| 2 | SO4 | B | 401 | 5/5 | 0.91 | 0.17 | 80,83,85,86 | 0 |
| 2 | SO4 | A | 401 | 5/5 | 0.94 | 0.14 | 54,56,65,68 | 0 |

The following is a graphical depiction of the model fit to experimental electron density of all instances of the Ligand of Interest. In addition, ligands with molecular weight > 250 and outliers as shown on the geometry validation Tables will also be included. Each fit is shown from different orientation to approximate a three-dimensional view.

**Electron density around A1H6D A 404:**

$2mF_o$-$DF_c$ (at 0.7 rmsd) in gray
$mF_o$-$DF_c$ (at 3 rmsd) in purple (negative)
and green (positive)

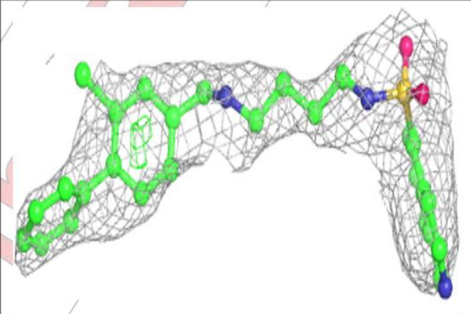

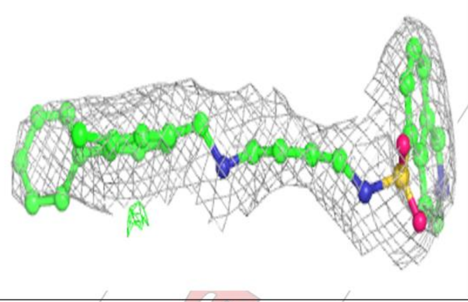

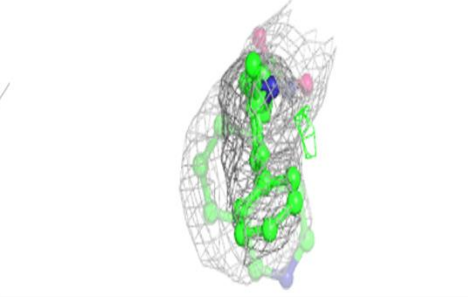

For Manuscript Review

**Electron density around A1H6D B 404:**

$2mF_o$-$DF_c$ (at 0.7 rmsd) in gray
$mF_o$-$DF_c$ (at 3 rmsd) in purple (negative)
and green (positive)

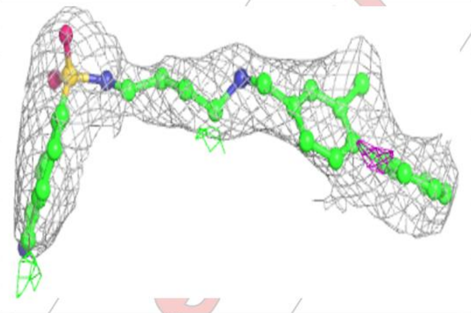

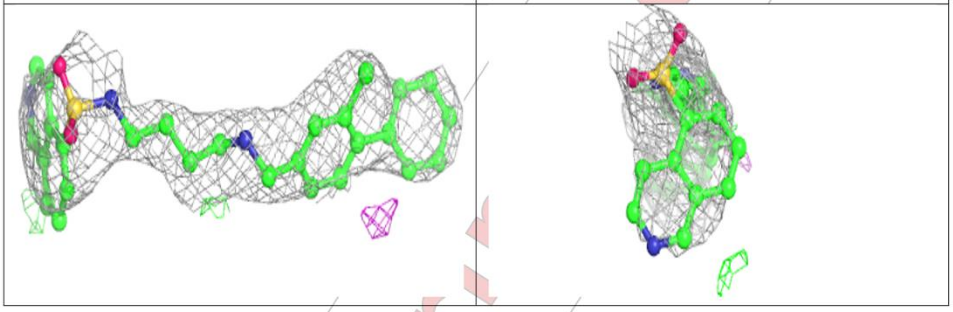

#### 6.5 Other polymers (i)

There are no such residues in this entry.

For Manuscript Review

### Full wwPDB X-ray Structure Validation Report (i)

Mar 20, 2024 – 03:57 pm GMT

| | | |
|---|---|---|
| PDB ID | : | 9EQ1 |
| Title | : | Crystal structure of human Casein Kinase II subunit alpha (CK2a1) in complex with allosteric ligand FGJM24 |
| Deposited on | : | 2024-03-20 |
| Resolution | : | 3.00 Å(reported) |

**This wwPDB validation report is for manuscript review**

For Manuscript Review

This is a Full wwPDB X-ray Structure Validation Report.

This report is produced by the wwPDB biocuration pipeline after annotation of the structure.

We welcome your comments at **
A user guide is available at
https://www.wwpdb.org/validation/2017/XrayValidationReportHelp
with specific help available everywhere you see the (i) symbol.

The types of validation reports are described at
http://www.wwpdb.org/validation/2017/FAQs#types.

The following versions of software and data (see references (i)) were used in the production of this report:

| | | |
|---|---|---|
| MolProbity | : | 4.02b-467 |
| Mogul | : | 1.8.4, CSD as541be (2020) |
| Xtriage (Phenix) | : | 1.13 |
| EDS | : | 2.36 |
| buster-report | : | 1.1.7 (2018) |
| Percentile statistics | : | 20191225.v01 (using entries in the PDB archive December 25th 2019) |
| Refmac | : | 5.8.0158 |
| CCP4 | : | 7.0.044 (Gargrove) |
| Ideal geometry (proteins) | : | Engh & Huber (2001) |

### 1 Overall quality at a glance (i)

The following experimental techniques were used to determine the structure:
*X-RAY DIFFRACTION*

The reported resolution of this entry is 3.00 Å.

Percentile scores (ranging between 0-100) for global validation metrics of the entry are shown in the following graphic. The table shows the number of entries on which the scores are based.

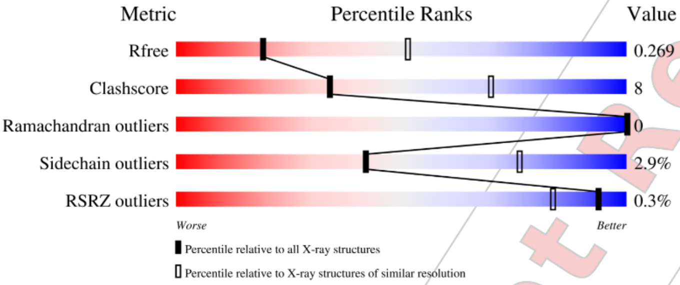

| Metric | Whole archive (#Entries) | Similar resolution (#Entries, resolution range(Å)) |
|---|---|---|
| $R_{free}$ | 130704 | 2092 (3.00-3.00) |
| Clashscore | 141614 | 2416 (3.00-3.00) |
| Ramachandran outliers | 138981 | 2333 (3.00-3.00) |
| Sidechain outliers | 138945 | 2336 (3.00-3.00) |
| RSRZ outliers | 127900 | 1990 (3.00-3.00) |

The table below summarises the geometric issues observed across the polymeric chains and their fit to the electron density. The red, orange, yellow and green segments of the lower bar indicate the fraction of residues that contain outliers for >=3, 2, 1 and 0 types of geometric quality criteria respectively. A grey segment represents the fraction of residues that are not modelled. The numeric value for each fraction is indicated below the corresponding segment, with a dot representing fractions <=5% The upper red bar (where present) indicates the fraction of residues that have poor fit to the electron density. The numeric value is given above the bar.

| Mol | Chain | Length | Quality of chain |
|---|---|---|---|
| 1 | A | 338 | 79% 19% • |
| 1 | B | 338 | 80% 17% • |

The following table lists non-polymeric compounds, carbohydrate monomers and non-standard

Ideal geometry (DNA, RNA) : Parkinson et al. (1996)
Validation Pipeline (wwPDB-VP) : 2.36

For Manuscript Review

residues in protein, DNA, RNA chains that are outliers for geometric or electron-density-fit criteria:

| Mol | Type | Chain | Res | Chirality | Geometry | Clashes | Electron density |
|---|---|---|---|---|---|---|---|
| 2 | SO4 | B | 405 | - | - | - | X |
| 2 | SO4 | B | 407 | - | - | - | X |
| 2 | SO4 | B | 409 | - | - | - | X |

For Manuscript Review

#### 2 Entry composition (i)

There are 5 unique types of molecules in this entry. The entry contains 5648 atoms, of which 0 are hydrogens and 0 are deuteriums.

In the tables below, the ZeroOcc column contains the number of atoms modelled with zero occupancy, the AltConf column contains the number of residues with at least one atom in alternate conformation and the Trace column contains the number of residues modelled with at most 2 atoms.

- Molecule 1 is a protein called Casein kinase II subunit alpha.

| Mol | Chain | Residues | Atoms | | | | | ZeroOcc | AltConf | Trace |
|---|---|---|---|---|---|---|---|---|---|---|
| 1 | A | 331 | Total 2721 | C 1748 | N 468 | O 494 | S 11 | 0 | 0 | 0 |
| 1 | B | 329 | Total 2738 | C 1755 | N 477 | O 495 | S 11 | 0 | 0 | 0 |

There are 2 discrepancies between the modelled and reference sequences:

| Chain | Residue | Modelled | Actual | Comment | Reference |
|---|---|---|---|---|---|
| A | 0 | SER | - | expression tag | UNP P68400 |
| B | 0 | SER | - | expression tag | UNP P68400 |

- Molecule 2 is SULFATE ION (three-letter code: SO4) (formula: $O_4S$).

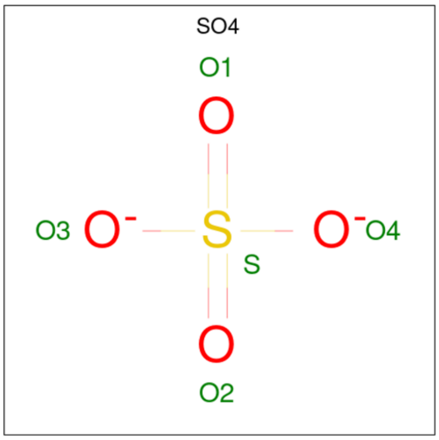

| Mol | Chain | Residues | Atoms | | | ZeroOcc | AltConf |
|---|---|---|---|---|---|---|---|
| 2 | A | 1 | Total 5 | O 4 | S 1 | 0 | 0 |

*Continued on next page...*

For Manuscript Review

*Continued from previous page...*

| Mol | Chain | Residues | Atoms | ZeroOcc | AltConf |
|---|---|---|---|---|---|
| 2 | A | 1 | Total O S<br>5 4 1 | 0 | 0 |
| 2 | A | 1 | Total O S<br>5 4 1 | 0 | 0 |
| 2 | A | 1 | Total O S<br>5 4 1 | 0 | 0 |
| 2 | A | 1 | Total O S<br>5 4 1 | 0 | 0 |
| 2 | A | 1 | Total O S<br>5 4 1 | 0 | 0 |
| 2 | A | 1 | Total O S<br>5 4 1 | 0 | 0 |
| 2 | B | 1 | Total O S<br>5 4 1 | 0 | 0 |
| 2 | B | 1 | Total O S<br>5 4 1 | 0 | 0 |
| 2 | B | 1 | Total O S<br>5 4 1 | 0 | 0 |
| 2 | B | 1 | Total O S<br>5 4 1 | 0 | 0 |
| 2 | B | 1 | Total O S<br>5 4 1 | 0 | 0 |
| 2 | B | 1 | Total O S<br>5 4 1 | 0 | 0 |
| 2 | B | 1 | Total O S<br>5 4 1 | 0 | 0 |
| 2 | B | 1 | Total O S<br>5 4 1 | 0 | 0 |
| 2 | B | 1 | Total O S<br>5 4 1 | 0 | 0 |
| 2 | B | 1 | Total O S<br>5 4 1 | 0 | 0 |
| 2 | B | 1 | Total O S<br>5 4 1 | 0 | 0 |
| 2 | B | 1 | Total O S<br>5 4 1 | 0 | 0 |

- Molecule 3 is 1,2-ETHANEDIOL (three-letter code: EDO) (formula: $C_2H_6O_2$).

For Manuscript Review

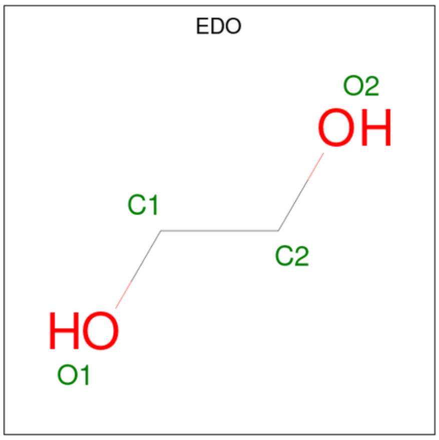

| Mol | Chain | Residues | Atoms | ZeroOcc | AltConf |
|---|---|---|---|---|---|
| 3 | A | 1 | Total C O<br>4 2 2 | 0 | 0 |
| 3 | B | 1 | Total C O<br>4 2 2 | 0 | 0 |

- Molecule 4 is methyl 2-[1,3-benzothiazol-6-ylsulfonyl-[5-[(3-chloranyl-4-phenyl-phenyl)methylamino]pentyl]amino]ethanoate (three-letter code: A1H6J) (formula: $C_{28}H_{30}ClN_3O_4S_2$) (labeled as "Ligand of Interest" by depositor).

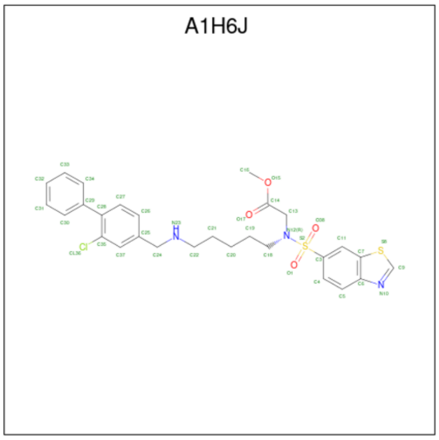

| Mol | Chain | Residues | Atoms | ZeroOcc | AltConf |
|---|---|---|---|---|---|
| 4 | A | 1 | Total C Cl N O S<br>38 28 1 3 4 2 | 0 | 0 |

*Continued on next page...*

*Continued from previous page...*

| Mol | Chain | Residues | Atoms | | | | | | ZeroOcc | AltConf |
|---|---|---|---|---|---|---|---|---|---|---|
| 4 | B | 1 | Total<br>38 | C<br>28 | Cl<br>1 | N<br>3 | O<br>4 | S<br>2 | 0 | 0 |

- Molecule 5 is water.

| Mol | Chain | Residues | Atoms | | ZeroOcc | AltConf |
|---|---|---|---|---|---|---|
| 5 | A | 6 | Total<br>6 | O<br>6 | 0 | 0 |
| 5 | B | 4 | Total<br>4 | O<br>4 | 0 | 0 |

For Manuscript Review

### 3 Residue-property plots ⓘ

These plots are drawn for all protein, RNA, DNA and oligosaccharide chains in the entry. The first graphic for a chain summarises the proportions of the various outlier classes displayed in the second graphic. The second graphic shows the sequence view annotated by issues in geometry and electron density. Residues are color-coded according to the number of geometric quality criteria for which they contain at least one outlier: green = 0, yellow = 1, orange = 2 and red = 3 or more. A red dot above a residue indicates a poor fit to the electron density (RSRZ > 2). Stretches of 2 or more consecutive residues without any outlier are shown as a green connector. Residues present in the sample, but not in the model, are shown in grey.

- Molecule 1: Casein kinase II subunit alpha

Chain A:

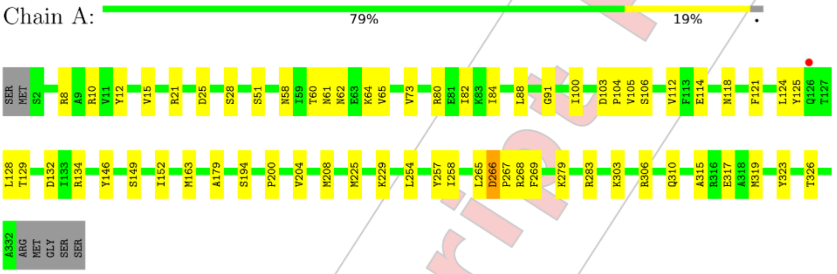

- Molecule 1: Casein kinase II subunit alpha

Chain B:

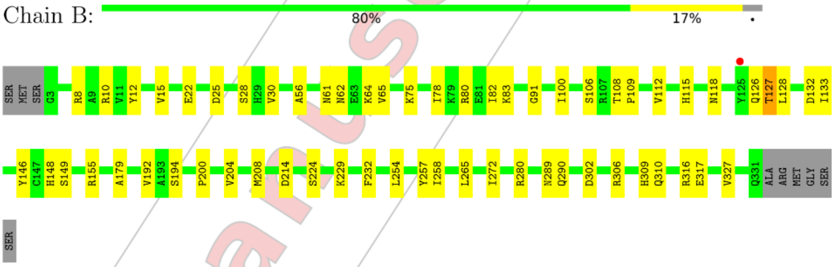

For Manuscript Review

### 4 Data and refinement statistics (i)

| Property | Value | Source |
|---|---|---|
| Space group | P 43 21 2 | Depositor |
| Cell constants<br>a, b, c, $\alpha$, $\beta$, $\gamma$ | 126.74Å 126.74Å 125.72Å<br>90.00° 90.00° 90.00° | Depositor |
| Resolution (Å) | 89.62 – 3.00<br>89.62 – 3.00 | Depositor<br>EDS |
| % Data completeness<br>(in resolution range) | 100.0 (89.62-3.00)<br>100.0 (89.62-3.00) | Depositor<br>EDS |
| $R_{merge}$ | (Not available) | Depositor |
| $R_{sym}$ | (Not available) | Depositor |
| $< I/\sigma(I) >$ [1] | 2.21 (at 3.01Å) | Xtriage |
| Refinement program | REFMAC 5.8.0419 | Depositor |
| R, $R_{free}$ | 0.210 , 0.251<br>0.226 , 0.269 | Depositor<br>DCC |
| $R_{free}$ test set | 976 reflections (4.63%) | wwPDB-VP |
| Wilson B-factor (Å$^2$) | 62.2 | Xtriage |
| Anisotropy | 0.477 | Xtriage |
| Bulk solvent $k_{sol}$(e/Å$^3$), $B_{sol}$(Å$^2$) | 0.35 , 53.5 | EDS |
| L-test for twinning[2] | $< \|L\| > = 0.51$, $< L^2 > = 0.35$ | Xtriage |
| Estimated twinning fraction | 0.000 for -h,-l,-k<br>0.000 for l,-k,h | Xtriage |
| $F_o$,$F_c$ correlation | 0.93 | EDS |
| Total number of atoms | 5648 | wwPDB-VP |
| Average B, all atoms (Å$^2$) | 65.0 | wwPDB-VP |

Xtriage's analysis on translational NCS is as follows: *The analyses of the Patterson function reveals a significant off-origin peak that is 32.27 % of the origin peak, indicating pseudo-translational symmetry. The chance of finding a peak of this or larger height randomly in a structure without pseudo-translational symmetry is equal to 9.6604e-04. The detected translational NCS is most likely also responsible for the elevated intensity ratio.*

[1] Intensities estimated from amplitudes.
[2] Theoretical values of $< |L| >$, $< L^2 >$ for acentric reflections are 0.5, 0.333 respectively for untwinned datasets, and 0.375, 0.2 for perfectly twinned datasets.

For Manuscript Review

### 5 Model quality (i)

#### 5.1 Standard geometry (i)

Bond lengths and bond angles in the following residue types are not validated in this section: EDO, SO4, A1H6J

The Z score for a bond length (or angle) is the number of standard deviations the observed value is removed from the expected value. A bond length (or angle) with $|Z| > 5$ is considered an outlier worth inspection. RMSZ is the root-mean-square of all Z scores of the bond lengths (or angles).

| Mol | Chain | Bond lengths | | Bond angles | |
|---|---|---|---|---|---|
| | | RMSZ | #\|Z\| >5 | RMSZ | #\|Z\| >5 |
| 1 | A | 0.46 | 0/2796 | 0.78 | 0/3796 |
| 1 | B | 0.45 | 0/2811 | 0.79 | 0/3809 |
| All | All | 0.45 | 0/5607 | 0.78 | 0/7605 |

There are no bond length outliers.

There are no bond angle outliers.

There are no chirality outliers.

There are no planarity outliers.

#### 5.2 Too-close contacts (i)

In the following table, the Non-H and H(model) columns list the number of non-hydrogen atoms and hydrogen atoms in the chain respectively. The H(added) column lists the number of hydrogen atoms added and optimized by MolProbity. The Clashes column lists the number of clashes within the asymmetric unit, whereas Symm-Clashes lists symmetry-related clashes.

| Mol | Chain | Non-H | H(model) | H(added) | Clashes | Symm-Clashes |
|---|---|---|---|---|---|---|
| 1 | A | 2721 | 0 | 2591 | 56 | 0 |
| 1 | B | 2738 | 0 | 2645 | 37 | 0 |
| 2 | A | 35 | 0 | 0 | 3 | 0 |
| 2 | B | 60 | 0 | 0 | 1 | 0 |
| 3 | A | 4 | 0 | 6 | 0 | 0 |
| 3 | B | 4 | 0 | 6 | 1 | 0 |
| 4 | A | 38 | 0 | 0 | 3 | 0 |
| 4 | B | 38 | 0 | 0 | 2 | 0 |
| 5 | A | 6 | 0 | 0 | 0 | 0 |
| 5 | B | 4 | 0 | 0 | 0 | 0 |
| All | All | 5648 | 0 | 5248 | 91 | 0 |

For Manuscript Review

The all-atom clashscore is defined as the number of clashes found per 1000 atoms (including hydrogen atoms). The all-atom clashscore for this structure is 8.

All (91) close contacts within the same asymmetric unit are listed below, sorted by their clash magnitude.

| Atom-1 | Atom-2 | Interatomic distance (Å) | Clash overlap (Å) |
|---|---|---|---|
| 1:A:303:LYS:HA | 1:A:306:ARG:HE | 1.09 | 1.15 |
| 1:A:225:MET:HG3 | 4:A:409:A1H6J:C33 | 2.04 | 0.87 |
| 1:A:265:LEU:HD11 | 1:A:269:PHE:CD2 | 2.12 | 0.85 |
| 1:B:56:ALA:O | 1:B:65:VAL:HG12 | 1.75 | 0.85 |
| 1:A:303:LYS:HA | 1:A:306:ARG:NE | 1.92 | 0.83 |
| 1:B:148:HIS:CD2 | 3:B:413:EDO:H12 | 2.18 | 0.78 |
| 1:A:80:ARG:NH2 | 2:A:407:SO4:O3 | 2.22 | 0.73 |
| 1:A:306:ARG:HH12 | 1:B:280:ARG:CD | 2.07 | 0.67 |
| 1:A:64:LYS:C | 1:A:65:VAL:HG23 | 2.14 | 0.67 |
| 1:A:80:ARG:HD3 | 1:A:179:ALA:O | 1.95 | 0.66 |
| 1:A:8:ARG:NH2 | 1:A:208:MET:CE | 2.59 | 0.66 |
| 1:B:78:ILE:HD13 | 1:B:109:PRO:HG2 | 1.79 | 0.65 |
| 1:B:80:ARG:HD3 | 1:B:179:ALA:O | 1.97 | 0.64 |
| 1:B:56:ALA:O | 1:B:65:VAL:CG1 | 2.46 | 0.63 |
| 1:A:64:LYS:O | 1:A:65:VAL:HG23 | 2.02 | 0.59 |
| 1:A:265:LEU:CD1 | 1:A:269:PHE:CD2 | 2.84 | 0.58 |
| 1:A:64:LYS:O | 1:A:65:VAL:CG2 | 2.51 | 0.58 |
| 1:A:105:VAL:HG23 | 1:A:106:SER:N | 2.19 | 0.58 |
| 1:A:118:ASN:HB2 | 1:A:163:MET:HE2 | 1.86 | 0.58 |
| 1:A:128:LEU:CD1 | 1:A:225:MET:HE1 | 2.34 | 0.57 |
| 1:A:265:LEU:HD11 | 1:A:269:PHE:CG | 2.40 | 0.57 |
| 1:A:303:LYS:CA | 1:A:306:ARG:HE | 2.01 | 0.57 |
| 1:A:84:ILE:HG23 | 1:A:152:ILE:HD13 | 1.88 | 0.55 |
| 1:B:25:ASP:O | 1:B:28:SER:HB2 | 2.07 | 0.55 |
| 1:A:128:LEU:O | 1:A:132:ASP:HB2 | 2.06 | 0.54 |
| 1:A:8:ARG:HH22 | 1:A:208:MET:CE | 2.19 | 0.54 |
| 1:B:10:ARG:NH1 | 1:B:317:GLU:OE2 | 2.41 | 0.54 |
| 1:A:25:ASP:O | 1:A:28:SER:HB2 | 2.08 | 0.54 |
| 1:A:105:VAL:HG23 | 1:A:106:SER:H | 1.73 | 0.52 |
| 1:A:121:PHE:CZ | 1:A:125:TYR:CD2 | 2.98 | 0.52 |
| 1:A:118:ASN:HB2 | 1:A:163:MET:CE | 2.38 | 0.52 |
| 1:A:128:LEU:O | 1:A:129:THR:OG1 | 2.26 | 0.52 |
| 1:A:8:ARG:NH2 | 1:A:208:MET:HE2 | 2.26 | 0.51 |
| 1:A:229:LYS:HE3 | 2:A:404:SO4:O2 | 2.11 | 0.51 |
| 1:B:148:HIS:HE1 | 1:B:214:ASP:OD2 | 1.93 | 0.50 |
| 1:B:232:PHE:O | 2:B:409:SO4:O2 | 2.28 | 0.50 |
| 1:A:61:ASN:O | 1:A:62:ASN:HB2 | 2.11 | 0.50 |

Continued on next page...

For Manuscript Review

*Continued from previous page...*

| Atom-1 | Atom-2 | Interatomic distance (Å) | Clash overlap (Å) |
|---|---|---|---|
| 1:A:64:LYS:C | 1:A:65:VAL:CG2 | 2.79 | 0.50 |
| 1:A:91:GLY:HA3 | 1:A:146:TYR:CE2 | 2.47 | 0.50 |
| 1:A:265:LEU:HD21 | 1:A:269:PHE:HB3 | 1.93 | 0.50 |
| 1:B:118:ASN:ND2 | 4:B:414:A1H6J:C21 | 2.75 | 0.50 |
| 1:B:200:PRO:O | 1:B:204:VAL:HG22 | 2.12 | 0.49 |
| 1:A:10:ARG:NH1 | 1:A:317:GLU:OE2 | 2.45 | 0.49 |
| 1:B:155:ARG:NH1 | 1:B:192:VAL:HG11 | 2.27 | 0.49 |
| 1:A:65:VAL:CG1 | 1:A:112:VAL:CG1 | 2.90 | 0.49 |
| 1:B:316:ARG:HG2 | 1:B:316:ARG:HH21 | 1.77 | 0.49 |
| 1:B:8:ARG:NH2 | 1:B:208:MET:CE | 2.76 | 0.49 |
| 1:B:65:VAL:CG2 | 1:B:112:VAL:CG1 | 2.91 | 0.49 |
| 1:A:200:PRO:O | 1:A:204:VAL:HG22 | 2.13 | 0.49 |
| 1:B:61:ASN:O | 1:B:62:ASN:HB2 | 2.12 | 0.49 |
| 1:B:302:ASP:O | 1:B:306:ARG:HD3 | 2.13 | 0.48 |
| 1:B:82:ILE:HD13 | 1:B:100:ILE:HD13 | 1.95 | 0.48 |
| 1:A:279:LYS:HB3 | 1:A:283:ARG:HD3 | 1.96 | 0.48 |
| 1:A:225:MET:CE | 4:A:409:A1H6J:CL36 | 2.99 | 0.47 |
| 1:A:254:LEU:O | 1:A:257:TYR:HB3 | 2.14 | 0.47 |
| 1:B:91:GLY:HA3 | 1:B:146:TYR:CE2 | 2.49 | 0.47 |
| 1:A:283:ARG:NH1 | 2:A:401:SO4:O1 | 2.48 | 0.47 |
| 1:B:118:ASN:HD21 | 4:B:414:A1H6J:C21 | 2.27 | 0.46 |
| 1:A:266:ASP:O | 1:A:267:PRO:C | 2.53 | 0.46 |
| 1:A:315:ALA:O | 1:A:319:MET:HG3 | 2.15 | 0.46 |
| 1:B:10:ARG:HD2 | 1:B:310:GLN:O | 2.16 | 0.46 |
| 1:B:265:LEU:HD23 | 1:B:265:LEU:HA | 1.75 | 0.46 |
| 1:A:124:LEU:O | 1:A:128:LEU:HG | 2.16 | 0.46 |
| 1:B:128:LEU:O | 1:B:132:ASP:HB2 | 2.16 | 0.45 |
| 1:A:82:ILE:HD13 | 1:A:100:ILE:HD13 | 1.98 | 0.45 |
| 1:B:254:LEU:O | 1:B:257:TYR:HB3 | 2.17 | 0.45 |
| 1:B:128:LEU:HD23 | 1:B:128:LEU:HA | 1.81 | 0.44 |
| 1:B:128:LEU:HD13 | 1:B:133:ILE:HG12 | 2.00 | 0.44 |
| 1:A:124:LEU:HD12 | 1:A:124:LEU:HA | 1.91 | 0.43 |
| 1:B:309:HIS:H | 1:B:309:HIS:CD2 | 2.35 | 0.43 |
| 1:A:58:ASN:OD1 | 1:A:60:THR:HB | 2.19 | 0.43 |
| 1:A:306:ARG:HH12 | 1:B:280:ARG:NE | 2.17 | 0.42 |
| 1:A:128:LEU:O | 1:A:129:THR:CB | 2.66 | 0.42 |
| 1:B:64:LYS:HG2 | 1:B:115:HIS:CD2 | 2.54 | 0.42 |
| 1:B:224:SER:HB3 | 1:B:229:LYS:O | 2.20 | 0.42 |
| 1:A:225:MET:CG | 4:A:409:A1H6J:C33 | 2.89 | 0.42 |
| 1:B:127:THR:O | 1:B:127:THR:CG2 | 2.67 | 0.42 |
| 1:B:106:SER:C | 1:B:108:THR:N | 2.72 | 0.42 |

*Continued on next page...*

For Manuscript Review

*Continued from previous page...*

| Atom-1 | Atom-2 | Interatomic distance (Å) | Clash overlap (Å) |
|---|---|---|---|
| 1:A:124:LEU:HG | 1:A:128:LEU:HD21 | 2.01 | 0.41 |
| 1:B:22:GLU:O | 1:B:83:LYS:HE2 | 2.20 | 0.41 |
| 1:A:8:ARG:HH22 | 1:A:208:MET:HE2 | 1.83 | 0.41 |
| 1:A:12:TYR:CD1 | 1:A:149:SER:HA | 2.55 | 0.41 |
| 1:A:266:ASP:O | 1:A:268:ARG:N | 2.54 | 0.41 |
| 1:A:103:ASP:OD1 | 1:A:104:PRO:HD2 | 2.21 | 0.41 |
| 1:A:114:GLU:O | 1:A:114:GLU:HG3 | 2.21 | 0.41 |
| 1:A:10:ARG:HD2 | 1:A:310:GLN:O | 2.20 | 0.41 |
| 1:A:134:ARG:HG2 | 1:A:323:TYR:CZ | 2.56 | 0.41 |
| 1:B:12:TYR:CD1 | 1:B:149:SER:HA | 2.55 | 0.41 |
| 1:B:127:THR:O | 1:B:127:THR:HG23 | 2.20 | 0.41 |
| 1:A:88:LEU:HD21 | 1:A:152:ILE:HD12 | 2.02 | 0.40 |
| 1:B:289:ASN:O | 1:B:290:GLN:C | 2.59 | 0.40 |

There are no symmetry-related clashes.

#### 5.3 Torsion angles (i)

##### 5.3.1 Protein backbone (i)

In the following table, the Percentiles column shows the percent Ramachandran outliers of the chain as a percentile score with respect to all X-ray entries followed by that with respect to entries of similar resolution.

The Analysed column shows the number of residues for which the backbone conformation was analysed, and the total number of residues.

| Mol | Chain | Analysed | Favoured | Allowed | Outliers | Percentiles | |
|---|---|---|---|---|---|---|---|
| 1 | A | 329/338 (97%) | 312 (95%) | 17 (5%) | 0 | 100 | 100 |
| 1 | B | 327/338 (97%) | 305 (93%) | 22 (7%) | 0 | 100 | 100 |
| All | All | 656/676 (97%) | 617 (94%) | 39 (6%) | 0 | 100 | 100 |

There are no Ramachandran outliers to report.

##### 5.3.2 Protein sidechains (i)

In the following table, the Percentiles column shows the percent sidechain outliers of the chain as a percentile score with respect to all X-ray entries followed by that with respect to entries of similar resolution.

For Manuscript Review

The Analysed column shows the number of residues for which the sidechain conformation was analysed, and the total number of residues.

| Mol | Chain | Analysed | Rotameric | Outliers | Percentiles | |
|---|---|---|---|---|---|---|
| 1 | A | 287/310 (93%) | 279 (97%) | 8 (3%) | 43 | 77 |
| 1 | B | 293/310 (94%) | 284 (97%) | 9 (3%) | 40 | 75 |
| All | All | 580/620 (94%) | 563 (97%) | 17 (3%) | 42 | 76 |

All (17) residues with a non-rotameric sidechain are listed below:

| Mol | Chain | Res | Type |
|---|---|---|---|
| 1 | A | 15 | VAL |
| 1 | A | 21 | ARG |
| 1 | A | 51 | SER |
| 1 | A | 73 | VAL |
| 1 | A | 194 | SER |
| 1 | A | 258 | ILE |
| 1 | A | 266 | ASP |
| 1 | A | 326 | THR |
| 1 | B | 15 | VAL |
| 1 | B | 30 | VAL |
| 1 | B | 75 | LYS |
| 1 | B | 126 | GLN |
| 1 | B | 127 | THR |
| 1 | B | 194 | SER |
| 1 | B | 258 | ILE |
| 1 | B | 272 | ILE |
| 1 | B | 327 | VAL |

Sometimes sidechains can be flipped to improve hydrogen bonding and reduce clashes. All (10) such sidechains are listed below:

| Mol | Chain | Res | Type |
|---|---|---|---|
| 1 | A | 126 | GLN |
| 1 | A | 186 | GLN |
| 1 | B | 115 | HIS |
| 1 | B | 118 | ASN |
| 1 | B | 148 | HIS |
| 1 | B | 186 | GLN |
| 1 | B | 290 | GLN |
| 1 | B | 309 | HIS |
| 1 | B | 310 | GLN |
| 1 | B | 331 | GLN |

For Manuscript Review

##### 5.3.3 RNA (i)

There are no RNA molecules in this entry.

#### 5.4 Non-standard residues in protein, DNA, RNA chains (i)

There are no non-standard protein/DNA/RNA residues in this entry.

#### 5.5 Carbohydrates (i)

There are no monosaccharides in this entry.

#### 5.6 Ligand geometry (i)

23 ligands are modelled in this entry.

In the following table, the Counts columns list the number of bonds (or angles) for which Mogul statistics could be retrieved, the number of bonds (or angles) that are observed in the model and the number of bonds (or angles) that are defined in the Chemical Component Dictionary. The Link column lists molecule types, if any, to which the group is linked. The Z score for a bond length (or angle) is the number of standard deviations the observed value is removed from the expected value. A bond length (or angle) with $|Z| > 2$ is considered an outlier worth inspection. RMSZ is the root-mean-square of all Z scores of the bond lengths (or angles).

| Mol | Type | Chain | Res | Link | Bond lengths | | | Bond angles | | |
|---|---|---|---|---|---|---|---|---|---|---|
| | | | | | Counts | RMSZ | $\#\|Z\| > 2$ | Counts | RMSZ | $\#\|Z\| > 2$ |
| 2 | SO4 | B | 403 | - | 4,4,4 | 0.39 | 0 | 6,6,6 | 0.11 | 0 |
| 2 | SO4 | A | 401 | - | 4,4,4 | 0.36 | 0 | 6,6,6 | 0.19 | 0 |
| 2 | SO4 | A | 405 | - | 4,4,4 | 0.32 | 0 | 6,6,6 | 0.20 | 0 |
| 2 | SO4 | A | 407 | - | 4,4,4 | 0.37 | 0 | 6,6,6 | 0.49 | 0 |
| 4 | A1H6J | A | 409 | - | 39,41,41 | 0.59 | 0 | 49,56,56 | 0.84 | 2 (4%) |
| 2 | SO4 | B | 406 | - | 4,4,4 | 0.28 | 0 | 6,6,6 | 0.18 | 0 |
| 2 | SO4 | B | 408 | - | 4,4,4 | 0.31 | 0 | 6,6,6 | 0.10 | 0 |
| 2 | SO4 | B | 407 | - | 4,4,4 | 0.34 | 0 | 6,6,6 | 0.14 | 0 |
| 2 | SO4 | B | 401 | - | 4,4,4 | 0.27 | 0 | 6,6,6 | 0.14 | 0 |
| 2 | SO4 | B | 410 | - | 4,4,4 | 0.28 | 0 | 6,6,6 | 0.10 | 0 |
| 2 | SO4 | B | 412 | - | 4,4,4 | 0.49 | 0 | 6,6,6 | 0.05 | 0 |
| 3 | EDO | A | 408 | - | 3,3,3 | 0.35 | 0 | 2,2,2 | 0.43 | 0 |
| 2 | SO4 | B | 402 | - | 4,4,4 | 0.23 | 0 | 6,6,6 | 0.17 | 0 |
| 2 | SO4 | B | 409 | - | 4,4,4 | 0.30 | 0 | 6,6,6 | 0.16 | 0 |
| 2 | SO4 | A | 402 | - | 4,4,4 | 0.37 | 0 | 6,6,6 | 0.24 | 0 |
| 2 | SO4 | B | 404 | - | 4,4,4 | 0.29 | 0 | 6,6,6 | 0.24 | 0 |
| 2 | SO4 | B | 411 | - | 4,4,4 | 0.31 | 0 | 6,6,6 | 0.28 | 0 |

For Manuscript Review

| Mol | Type | Chain | Res | Link | Bond lengths | | | Bond angles | | |
|---|---|---|---|---|---|---|---|---|---|---|
| | | | | | Counts | RMSZ | #\|Z\| > 2 | Counts | RMSZ | #\|Z\| > 2 |
| 4 | A1H6J | B | 414 | - | 39,41,41 | 0.57 | 0 | 49,56,56 | 0.99 | 2 (4%) |
| 2 | SO4 | A | 406 | - | 4,4,4 | 0.42 | 0 | 6,6,6 | 0.40 | 0 |
| 3 | EDO | B | 413 | - | 3,3,3 | 0.63 | 0 | 2,2,2 | 0.69 | 0 |
| 2 | SO4 | B | 405 | - | 4,4,4 | 0.35 | 0 | 6,6,6 | 0.15 | 0 |
| 2 | SO4 | A | 403 | - | 4,4,4 | 0.35 | 0 | 6,6,6 | 0.18 | 0 |
| 2 | SO4 | A | 404 | - | 4,4,4 | 0.37 | 0 | 6,6,6 | 0.08 | 0 |

In the following table, the Chirals column lists the number of chiral outliers, the number of chiral centers analysed, the number of these observed in the model and the number defined in the Chemical Component Dictionary. Similar counts are reported in the Torsion and Rings columns. '-' means no outliers of that kind were identified.

| Mol | Type | Chain | Res | Link | Chirals | Torsions | Rings |
|---|---|---|---|---|---|---|---|
| 3 | EDO | A | 408 | - | - | 0/1/1/1 | - |
| 3 | EDO | B | 413 | - | - | 0/1/1/1 | - |
| 4 | A1H6J | B | 414 | - | - | 16/32/32/32 | 0/4/4/4 |
| 4 | A1H6J | A | 409 | - | - | 15/32/32/32 | 0/4/4/4 |

There are no bond length outliers.

All (4) bond angle outliers are listed below:

| Mol | Chain | Res | Type | Atoms | Z | Observed($^o$) | Ideal($^o$) |
|---|---|---|---|---|---|---|---|
| 4 | B | 414 | A1H6J | C13-N12-C18 | -3.80 | 107.52 | 116.84 |
| 4 | A | 409 | A1H6J | C7-C6-N10 | 3.12 | 114.81 | 107.87 |
| 4 | B | 414 | A1H6J | C7-C6-N10 | 3.11 | 114.77 | 107.87 |
| 4 | A | 409 | A1H6J | C18-N12-S2 | -2.60 | 112.65 | 117.92 |

There are no chirality outliers.

All (31) torsion outliers are listed below:

| Mol | Chain | Res | Type | Atoms |
|---|---|---|---|---|
| 4 | A | 409 | A1H6J | C14-C13-N12-S2 |
| 4 | A | 409 | A1H6J | C14-C13-N12-C18 |
| 4 | A | 409 | A1H6J | C19-C18-N12-S2 |
| 4 | A | 409 | A1H6J | C19-C18-N12-C13 |
| 4 | A | 409 | A1H6J | N12-C13-C14-O15 |
| 4 | A | 409 | A1H6J | N12-C13-C14-O17 |
| 4 | B | 414 | A1H6J | C14-C13-N12-C18 |
| 4 | A | 409 | A1H6J | C13-N12-S2-O1 |
| 4 | A | 409 | A1H6J | C25-C24-N23-C22 |

*Continued on next page...*

WORLDWIDE PDB PROTEIN DATA BANK

For Manuscript Review

*Continued from previous page...*

| Mol | Chain | Res | Type | Atoms |
|---|---|---|---|---|
| 4 | A | 409 | A1H6J | C13-N12-S2-C3 |
| 4 | B | 414 | A1H6J | C20-C21-C22-N23 |
| 4 | B | 414 | A1H6J | C13-N12-S2-O1 |
| 4 | A | 409 | A1H6J | C19-C20-C21-C22 |
| 4 | B | 414 | A1H6J | C18-C19-C20-C21 |
| 4 | A | 409 | A1H6J | C21-C22-N23-C24 |
| 4 | B | 414 | A1H6J | C13-N12-S2-C3 |
| 4 | B | 414 | A1H6J | C14-C13-N12-S2 |
| 4 | A | 409 | A1H6J | N12-C18-C19-C20 |
| 4 | B | 414 | A1H6J | N12-C18-C19-C20 |
| 4 | B | 414 | A1H6J | C35-C28-C29-C30 |
| 4 | B | 414 | A1H6J | C4-C3-S2-O38 |
| 4 | B | 414 | A1H6J | N12-C13-C14-O17 |
| 4 | B | 414 | A1H6J | N12-C13-C14-O15 |
| 4 | A | 409 | A1H6J | C11-C3-S2-O38 |
| 4 | B | 414 | A1H6J | C27-C28-C29-C30 |
| 4 | B | 414 | A1H6J | C27-C28-C29-C34 |
| 4 | B | 414 | A1H6J | C4-C3-S2-N12 |
| 4 | A | 409 | A1H6J | C4-C3-S2-O38 |
| 4 | B | 414 | A1H6J | C11-C3-S2-O38 |
| 4 | B | 414 | A1H6J | C21-C22-N23-C24 |
| 4 | A | 409 | A1H6J | C27-C28-C29-C34 |

There are no ring outliers.

7 monomers are involved in 10 short contacts:

| Mol | Chain | Res | Type | Clashes | Symm-Clashes |
|---|---|---|---|---|---|
| 2 | A | 401 | SO4 | 1 | 0 |
| 2 | A | 407 | SO4 | 1 | 0 |
| 4 | A | 409 | A1H6J | 3 | 0 |
| 2 | B | 409 | SO4 | 1 | 0 |
| 4 | B | 414 | A1H6J | 2 | 0 |
| 3 | B | 413 | EDO | 1 | 0 |
| 2 | A | 404 | SO4 | 1 | 0 |

The following is a two-dimensional graphical depiction of Mogul quality analysis of bond lengths, bond angles, torsion angles, and ring geometry for all instances of the Ligand of Interest. In addition, ligands with molecular weight > 250 and outliers as shown on the validation Tables will also be included. For torsion angles, if less then 5% of the Mogul distribution of torsion angles is within 10 degrees of the torsion angle in question, then that torsion angle is considered an outlier. Any bond that is central to one or more torsion angles identified as an outlier by Mogul will be highlighted in the graph. For rings, the root-mean-square deviation (RMSD) between the ring

For Manuscript Review

in question and similar rings identified by Mogul is calculated over all ring torsion angles. If the average RMSD is greater than 60 degrees and the minimal RMSD between the ring in question and any Mogul-identified rings is also greater than 60 degrees, then that ring is considered an outlier. The outliers are highlighted in purple. The color gray indicates Mogul did not find sufficient equivalents in the CSD to analyse the geometry.

**Ligand A1H6J A 409**

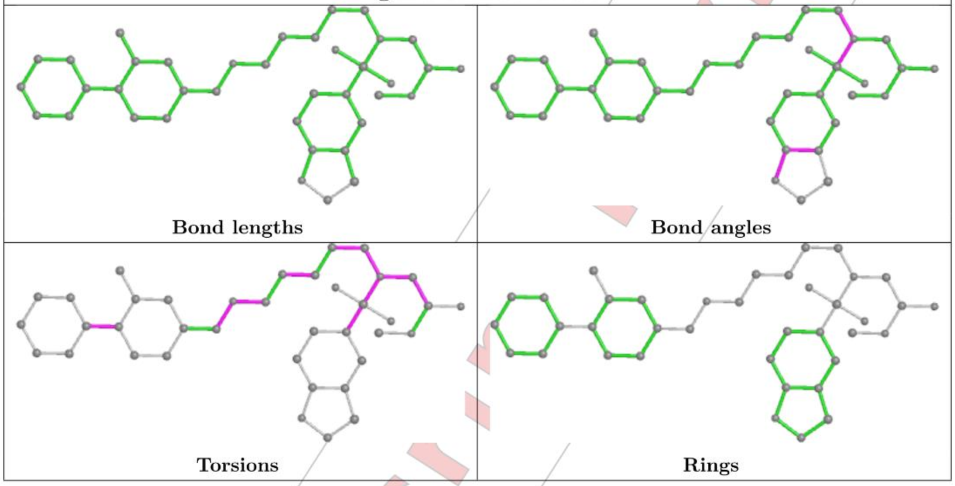

**Ligand A1H6J B 414**

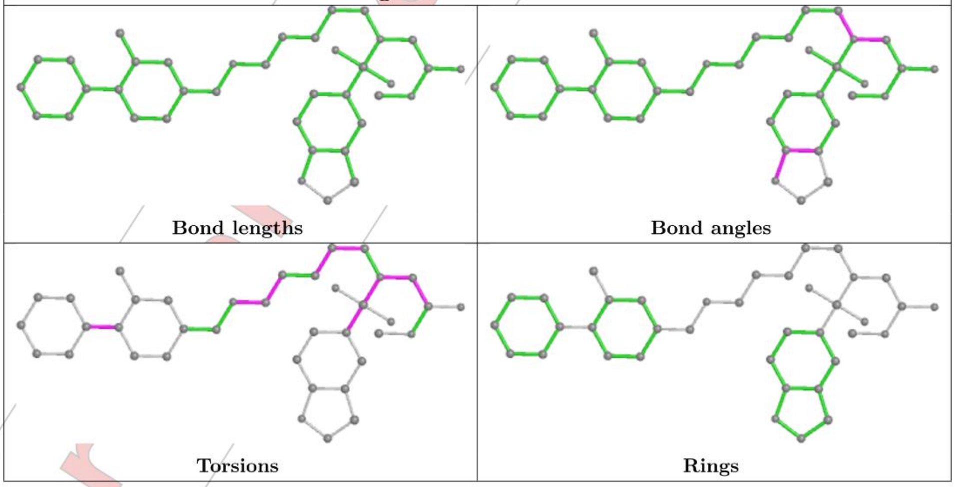

#### 5.7 Other polymers (i)

There are no such residues in this entry.

For Manuscript Review

#### 5.8 Polymer linkage issues (i)

There are no chain breaks in this entry.

For Manuscript Review

### 6 Fit of model and data (i)

#### 6.1 Protein, DNA and RNA chains (i)

In the following table, the column labelled '#RSRZ> 2' contains the number (and percentage) of RSRZ outliers, followed by percent RSRZ outliers for the chain as percentile scores relative to all X-ray entries and entries of similar resolution. The OWAB column contains the minimum, median, $95^{th}$ percentile and maximum values of the occupancy-weighted average B-factor per residue. The column labelled 'Q< 0.9' lists the number of (and percentage) of residues with an average occupancy less than 0.9.

| Mol | Chain | Analysed | <RSRZ> | #RSRZ>2 | | | OWAB(Å$^2$) | Q<0.9 |
|---|---|---|---|---|---|---|---|---|
| 1 | A | 331/338 (97%) | -0.05 | 1 (0%) | 94 | 84 | 30, 58, 102, 137 | 0 |
| 1 | B | 329/338 (97%) | 0.03 | 1 (0%) | 94 | 84 | 30, 60, 105, 136 | 0 |
| All | All | 660/676 (97%) | -0.01 | 2 (0%) | 94 | 84 | 30, 59, 104, 137 | 0 |

All (2) RSRZ outliers are listed below:

| Mol | Chain | Res | Type | RSRZ |
|---|---|---|---|---|
| 1 | A | 126 | GLN | 2.6 |
| 1 | B | 125 | TYR | 2.0 |

#### 6.2 Non-standard residues in protein, DNA, RNA chains (i)

There are no non-standard protein/DNA/RNA residues in this entry.

#### 6.3 Carbohydrates (i)

There are no monosaccharides in this entry.

#### 6.4 Ligands (i)

In the following table, the Atoms column lists the number of modelled atoms in the group and the number defined in the chemical component dictionary. The B-factors column lists the minimum, median, $95^{th}$ percentile and maximum values of B factors of atoms in the group. The column labelled 'Q< 0.9' lists the number of atoms with occupancy less than 0.9.

| Mol | Type | Chain | Res | Atoms | RSCC | RSR | B-factors(Å$^2$) | Q<0.9 |
|---|---|---|---|---|---|---|---|---|
| 2 | SO4 | B | 405 | 5/5 | 0.60 | 0.56 | 113,125,154,169 | 0 |
| 2 | SO4 | B | 407 | 5/5 | 0.66 | 0.67 | 126,153,172,180 | 0 |

Continued on next page...

For Manuscript Review

*Continued from previous page...*

| Mol | Type | Chain | Res | Atoms | RSCC | RSR | B-factors(Å$^2$) | Q<0.9 |
|---|---|---|---|---|---|---|---|---|
| 2 | SO4 | B | 408 | 5/5 | 0.70 | 0.28 | 119,132,142,160 | 0 |
| 2 | SO4 | A | 406 | 5/5 | 0.72 | 0.22 | 70,82,110,149 | 0 |
| 2 | SO4 | B | 404 | 5/5 | 0.76 | 0.27 | 84,109,130,145 | 0 |
| 2 | SO4 | B | 406 | 5/5 | 0.78 | 0.19 | 104,107,120,127 | 0 |
| 2 | SO4 | B | 409 | 5/5 | 0.80 | 0.55 | 100,131,142,152 | 0 |
| 2 | SO4 | A | 404 | 5/5 | 0.81 | 0.67 | 137,137,155,163 | 0 |
| 2 | SO4 | B | 403 | 5/5 | 0.81 | 0.20 | 115,131,142,148 | 0 |
| 2 | SO4 | B | 410 | 5/5 | 0.82 | 0.30 | 102,107,132,138 | 0 |
| 2 | SO4 | B | 401 | 5/5 | 0.83 | 0.36 | 96,102,125,154 | 0 |
| 4 | A1H6J | B | 414 | 38/38 | 0.85 | 0.35 | 62,84,120,135 | 0 |
| 4 | A1H6J | A | 409 | 38/38 | 0.87 | 0.31 | 53,84,117,136 | 0 |
| 3 | EDO | B | 413 | 4/4 | 0.88 | 0.32 | 38,47,50,54 | 0 |
| 2 | SO4 | A | 402 | 5/5 | 0.88 | 0.20 | 90,94,96,98 | 0 |
| 2 | SO4 | A | 405 | 5/5 | 0.88 | 0.26 | 112,127,138,148 | 0 |
| 2 | SO4 | B | 411 | 5/5 | 0.89 | 0.13 | 77,82,85,86 | 0 |
| 3 | EDO | A | 408 | 4/4 | 0.90 | 0.24 | 76,79,87,90 | 0 |
| 2 | SO4 | A | 403 | 5/5 | 0.92 | 0.15 | 72,87,92,98 | 0 |
| 2 | SO4 | B | 402 | 5/5 | 0.93 | 0.17 | 72,89,103,128 | 0 |
| 2 | SO4 | A | 401 | 5/5 | 0.94 | 0.32 | 74,80,101,103 | 0 |
| 2 | SO4 | B | 412 | 5/5 | 0.95 | 0.13 | 60,69,73,78 | 0 |
| 2 | SO4 | A | 407 | 5/5 | 0.97 | 0.17 | 55,57,58,59 | 0 |

The following is a graphical depiction of the model fit to experimental electron density of all instances of the Ligand of Interest. In addition, ligands with molecular weight > 250 and outliers as shown on the geometry validation Tables will also be included. Each fit is shown from different orientation to approximate a three-dimensional view.

For Manuscript Review

**Electron density around A1H6J B 414:**

2mF$_o$-DF$_c$ (at 0.7 rmsd) in gray
mF$_o$-DF$_c$ (at 3 rmsd) in purple (negative)
and green (positive)

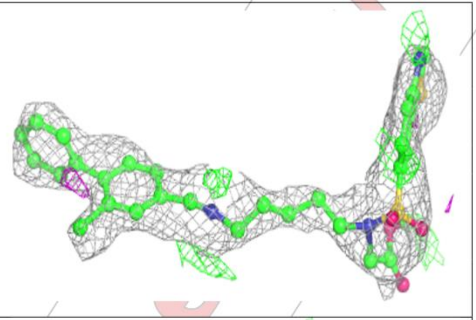

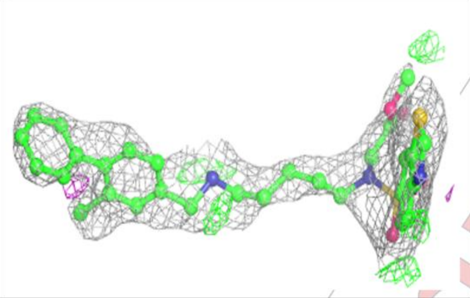

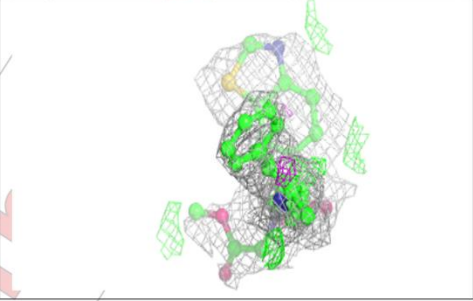

**Electron density around A1H6J A 409:**

2mF$_o$-DF$_c$ (at 0.7 rmsd) in gray
mF$_o$-DF$_c$ (at 3 rmsd) in purple (negative)
and green (positive)

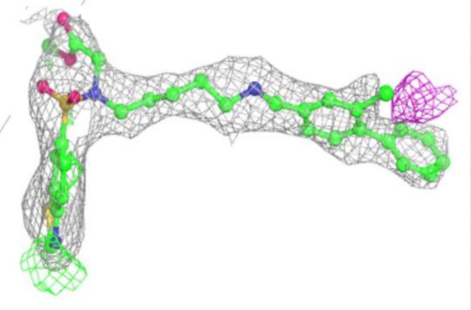

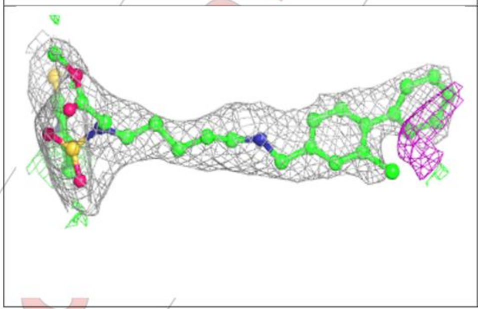

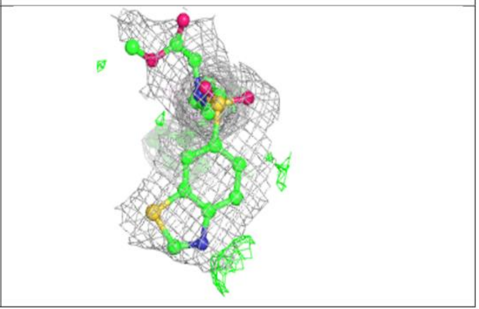

For Manuscript Review

#### 6.5 Other polymers (i)

There are no such residues in this entry.

For Manuscript Review

**References**

References

(1) Hirozane, Y.; Toyofuku, M.; Yogo, T.; Tanaka, Y.; Sameshima, T.; Miyahisa, I.; Yoshikawa, M. Structure-based rational design of staurosporine-based fluorescent probe with broad-ranging kinase affinity for kinase panel application. *Bioorganic & medicinal chemistry letters* **2019**, *29* (21), 126641. DOI: 10.1016/j.bmcl.2019.126641. Published Online: Sep. 7, 2019.

(2) Vasta, J. D.; Corona, C. R.; Wilkinson, J.; Zimprich, C. A.; Hartnett, J. R.; Ingold, M. R.; Zimmerman, K.; Machleidt, T.; Kirkland, T. A.; Huwiler, K. G.; Ohana, R. F.; Slater, M.; Otto, P.; Cong, M.; Wells, C. I.; Berger, B.-T.; Hanke, T.; Glas, C.; Ding, K.; Drewry, D. H.; Huber, K. V. M.; Willson, T. M.; Knapp, S.; Müller, S.; Meisenheimer, P. L.; Fan, F.; Wood, K. V.; Robers, M. B. Quantitative, Wide-Spectrum Kinase Profiling in Live Cells for Assessing the Effect of Cellular ATP on Target Engagement. *Cell chemical biology* **2018**, *25* (2), 206-214.e11. DOI: 10.1016/j.chembiol.2017.10.010. Published Online: Nov. 22, 2017.

(3) Schwalm, M. P.; Krämer, A.; Dölle, A.; Weckesser, J.; Yu, X.; Jin, J.; Saxena, K.; Knapp, S. Tracking the PROTAC degradation pathway in living cells highlights the importance of ternary complex measurement for PROTAC optimization. *Cell chemical biology* **2023**, *30* (7), 753-765.e8. DOI: 10.1016/j.chembiol.2023.06.002. Published Online: Jun. 23, 2023.

(4) Robers, M. B.; Friedman-Ohana, R.; Huber, K. V. M.; Kilpatrick, L.; Vasta, J. D.; Berger, B.-T.; Chaudhry, C.; Hill, S.; Müller, S.; Knapp, S.; Wood, K. V. Quantifying Target Occupancy of Small Molecules Within Living Cells. *Annual review of biochemistry* **2020**, *89*, 557–581. DOI: 10.1146/annurev-biochem-011420-092302. Published Online: Mar. 24, 2020.

(5) Wells, C. I.; Drewry, D. H.; Pickett, J. E.; Tjaden, A.; Krämer, A.; Müller, S.; Gyenis, L.; Menyhart, D.; Litchfield, D. W.; Knapp, S.; Axtman, A. D. Development of a potent and selective chemical probe for the pleiotropic kinase CK2. *Cell chemical biology* **2021**, *28* (4), 546-558.e10. DOI: 10.1016/j.chembiol.2020.12.013. Published Online: Jan. 22, 2021.

(6) Davis-Gilbert, Z. W.; Krämer, A.; Dunford, J. E.; Howell, S.; Senbabaoglu, F.; Wells, C. I.; Bashore, F. M.; Havener, T. M.; Smith, J. L.; Hossain, M. A.; Oppermann, U.; Drewry, D. H.; Axtman, A. D. Discovery of a Potent and Selective Naphthyridine-Based Chemical Probe for Casein Kinase 2. *ACS medicinal chemistry letters* **2023**, *14* (4), 432–441. DOI: 10.1021/acsmedchemlett.2c00530. Published Online: Mar. 14, 2023.

(7) Krämer, A.; Kurz, C. G.; Berger, B.-T.; Celik, I. E.; Tjaden, A.; Greco, F. A.; Knapp, S.; Hanke, T. Optimization of pyrazolo1,5-apyrimidines lead to the identification of a highly selective casein kinase 2 inhibitor. *European journal of medicinal chemistry* **2020**, *208*, 112770. DOI: 10.1016/j.ejmech.2020.112770. Published Online: Aug. 23, 2020.

(8) Kabsch, W. XDS. *Acta crystallographica. Section D, Biological crystallography* **2010**, *66* (Pt 2), 125–132. DOI: 10.1107/S0907444909047337. Published Online: Jan. 22, 2010.

(9) Evans, P. R.; Murshudov, G. N. How good are my data and what is the resolution? *Acta crystallographica. Section D, Biological crystallography* **2013**, *69* (Pt 7), 1204–1214. DOI: 10.1107/S0907444913000061. Published Online: Jun. 13, 2013.

(10) Lebedev, A. A.; Vagin, A. A.; Murshudov, G. N. Model preparation in MOLREP and examples of model improvement using X-ray data. *Acta crystallographica. Section D, Biological crystallography* **2008**, *64* (Pt 1), 33–39. DOI: 10.1107/S0907444907049839. Published Online: Dec. 5, 2007.

(11) Emsley, P.; Cowtan, K. Coot: model-building tools for molecular graphics. *Acta crystallographica. Section D, Biological crystallography* **2004**, *60* (Pt 12 Pt 1), 2126–2132. DOI: 10.1107/S0907444904019158. Published Online: Nov. 26, 2004.

(12) Vagin, A. A.; Steiner, R. A.; Lebedev, A. A.; Potterton, L.; McNicholas, S.; Long, F.; Murshudov, G. N. REFMAC5 dictionary: organization of prior chemical knowledge and guidelines for its use. *Acta crystallographica. Section D, Biological crystallography* **2004**, *60* (Pt 12 Pt 1), 2184–2195. DOI: 10.1107/S0907444904023510. Published Online: Nov. 26, 2004.

(13) Davies, A. T.; Curto, J. M.; Bagley, S. W.; Willis, M. C. One-pot palladium-catalyzed synthesis of sulfonyl fluorides from aryl bromides. *Chemical science* **2017**, *8* (2), 1233–1237. DOI: 10.1039/c6sc03924c. Published Online: Oct. 11, 2016.

(14) Iegre, J.; Brear, P.; Fusco, C. de; Yoshida, M.; Mitchell, S. L.; Rossmann, M.; Carro, L.; Sore, H. F.; Hyvönen, M.; Spring, D. R. Second-generation CK2α inhibitors targeting the αD pocket. *Chemical science* **2018**, *9* (11), 3041–3049. DOI: 10.1039/c7sc05122k. Published Online: Feb. 20, 2018.
